# Brainstem organoids from human pluripotent stem cells contain neural crest population

**DOI:** 10.1101/829275

**Authors:** Nobuyuki Eura, Takeshi K. Matsui, Joachim Luginbühl, Masaya Matsubayashi, Hitoki Nanaura, Tomo Shiota, Kaoru Kinugawa, Naohiko Iguchi, Takao Kiriyama, Canbin Zheng, Tsukasa Kouno, Yan Jun Lan, Pornparn Kongpracha, Pattama Wiriyasermkul, Yoshihiko M. Sakaguchi, Riko Nagata, Tomoya Komeda, Naritaka Morikawa, Fumika Kitayoshi, Miyong Jong, Shinko Kobashigawa, Mari Nakanishi, Masatoshi Hasegawa, Yasuhiko Saito, Takashi Shiromizu, Yuhei Nishimura, Takahiko Kasai, Maiko Takeda, Hiroshi Kobayashi, Yusuke Inagaki, Yasuhito Tanaka, Manabu Makinodan, Toshifumi Kishimoto, Hiroki Kuniyasu, Shushi Nagamori, Alysson R. Muotri, Jay W. Shin, Kazuma Sugie, Eiichiro Mori

**Affiliations:** Department of Neurology, Nara Medical University, Kashihara, Nara, Japan; Department of Future Basic Medicine, Nara Medical University, Kashihara, Nara, Japan; Division of Genomic Technologies, RIKEN Center for Life Science Technologies, Yokohama, Kanagawa, Japan; Departments of Pediatrics and Cellular & Molecular Medicine, University of California San Diego, San Diego, CA, USA; Laboratory of Biomolecular Dynamics, Department of Collaborative Research, Nara Medical University, Kashihara, Nara, Japan; Department of Radiation Oncology, Nara Medical University, Kashihara, Nara, Japan; Department of Neurophysiology, Nara Medical University, Kashihara, Nara, Japan; Department of Integrative Pharmacology, Mie University Graduate School of Medicine, Tsu, Mie, Japan; Department of Laboratory Medicine and Pathology, National Hospital Organization Kinki-chuo Chest Medical Center, Osaka, Japan; Department of Obstetrics and Gynecology, Nara Medical University, Kashihara, Nara, Japan; Department of Orthopaedic Surgery, Nara Medical University, Kashihara, Nara, Japan; Department of Psychiatry, Nara Medical University, Kashihara, Nara, Japan; Department of Molecular Pathology, Nara Medical University, Kashihara, Nara, Japan

**Keywords:** brain organoids, brainstem, neural crest, midbrain, dopaminergic neurons, human pluripotent stem cells, melanocyte

## Abstract

The brainstem controls heartbeat, blood pressure and respiration, which are life-sustaining functions, therefore, disorders of the brainstem can be lethal. Brain organoids derived from human pluripotent stem cells recapitulate the course of human brain development and are expected to be useful for medical research on central nervous system disorders. However, existing organoid models have limitations, hampering the elucidation of diseases affecting specific components of the brain. Here, we developed a method to generate human brainstem organoids (hBSOs), containing neural crest stem cells as well as midbrain/hindbrain progenitors, noradrenergic and cholinergic neurons, and dopaminergic neurons, demonstrated by specific electrophysiological signatures. Single-cell RNA sequence analysis, together with proteomics and electrophysiology, revealed that the cellular population in these organoids was similar to that of the human brainstem and neural crest, which raises the possibility of making use of hBSOs in grafting for transplantation, efficient drug screenings and modeling the neural crest diseases.

## Introduction

The brainstem is a posterior region of the brain between the deep structures of the cerebral hemispheres. It connects the cerebrum with the spinal cord and is divided into three parts: midbrain, pons and medulla oblongata. They contain multiple nuclei and small fiber tracts widely projecting to the cerebrum cortex, basal ganglia and other parts of the cerebrum. Brainstem functions such as alertness, heartbeat, blood pressure and respiration are considered to be more vital for life than that of the cortex. Therefore, damages to or disorders of brainstem such as infarction, hemorrhage, tumors, or any neurodegenerative diseases may lead to death.

Of those, Parkinson’s disease (PD) is the most well-known degenerative disease, showing progressive voluntary movement impairments, including rigidity, akinesia, tremor, and postural instability. These symptoms are caused by the loss of dopaminergic neurons at midbrain substantia nigra. In addition, non-motor symptoms, such as autonomic dysfunction, sleep disorder, and depression in PD patients, are thought to be derived from impairments of the serotonergic or noradrenergic system in the brainstem. However, the mechanisms driving these symptoms have yet to be determined. Hence, we need the development of models that can recapitulate the human midbrain and surrounding brainstem to elucidate the process of neural degeneration in this area.

Recent progress on protocols for inducing organs in-a-dish (organoids) provides potentials for the modeling of various diseases (Clevers, 2016). Organoids mimic the structure of organs composed of various cells such as the kidney (Takasato et al., 2015), brain (Dang et al., 2016), colon (Sato et al., 2009), and retina (Eiraku et al., 2011; Nakano et al., 2012). The use of brain organoids is a recognized method for the recapitulation of human fetal development during in vitro cultivation (Lancaster and Knoblich, 2014; Lancaster et al., 2013; Trujillo et al., 2019).

However, improvements to the protocols are still needed, particularly in the aspects such as the maturity, efficiency and the extent of recapitulation captured in the organoids. Recently, a protocol for generating human midbrain-like organoids from human pluripotent stem cells (hPSCs) has been reported (Jo et al., 2016). There are also reports on the effects of reagents or growth-factors on the differentiation of dopaminergic neurons (Ayton et al., 2016; Diaz et al., 2009; Lee et al., 2016). Based on these findings, we designed a new method for generating a human brainstem organoid (hBSO) model where the midbrain, surrounding brainstem parts, and neural crest region behind them are induced by the addition of basic fibroblast growth factor (bFGF) and epidermal growth factor (EGF) for neuronal stem/progenitor cells expansion. This is followed by treatment with brain derived neurotrophic factor (BDNF), glial cell line derived neurotrophic factor (GDNF), neurotrophin 3 (NT-3), cyclic adenosine monophosphate (cAMP) and ascorbic acid for the differentiation of neural crest cells and dopaminergic neurons under the condition in favor of the survival of dopaminergic neurons. In the present study, we established a novel method for inducing hBSOs, which contained neural crest cells surrounding the brainstem. We believe our methods will become a powerful tool in examining the pathology of neurodegenerative diseases affecting brainstem such as PD and neural crest disorders.

## Results

To generate hBSOs from hPSCs containing neural crest cells from hPSCs, we used a combination of several growth factors, including EGF/bFGF for the initial proliferation of neuronal stem/progenitor cells, and BDNF, GDNF, and NT-3 for the subsequent differentiation of dopaminergic neurons and neural crest cells; this procedure was different from the protocols in previously reported studies (Figures 1A and S1). A recent study by Muotri and colleagues reported the presence of neural networks in their cortical organoids with advanced maturity (Trujillo et al., 2018). We further modified their protocol to induce dopaminergic neurons by adding insulin, transferrin and progesterone, all of which have been shown to induce dopaminergic neuronal differentiation and survival in 2D culture (Ayton et al., 2016; Diaz et al., 2009; Lee et al., 2016). Our novel approach to generate hBSO yielded cells with dark granules between 22 – 28 days of cultivation (Figure 1A), an indication of neural crest origin, which is an observation absent at this stage in previous studies. (Thomas et al., 2017; Trujillo et al., 2018).

**Figure 1.**
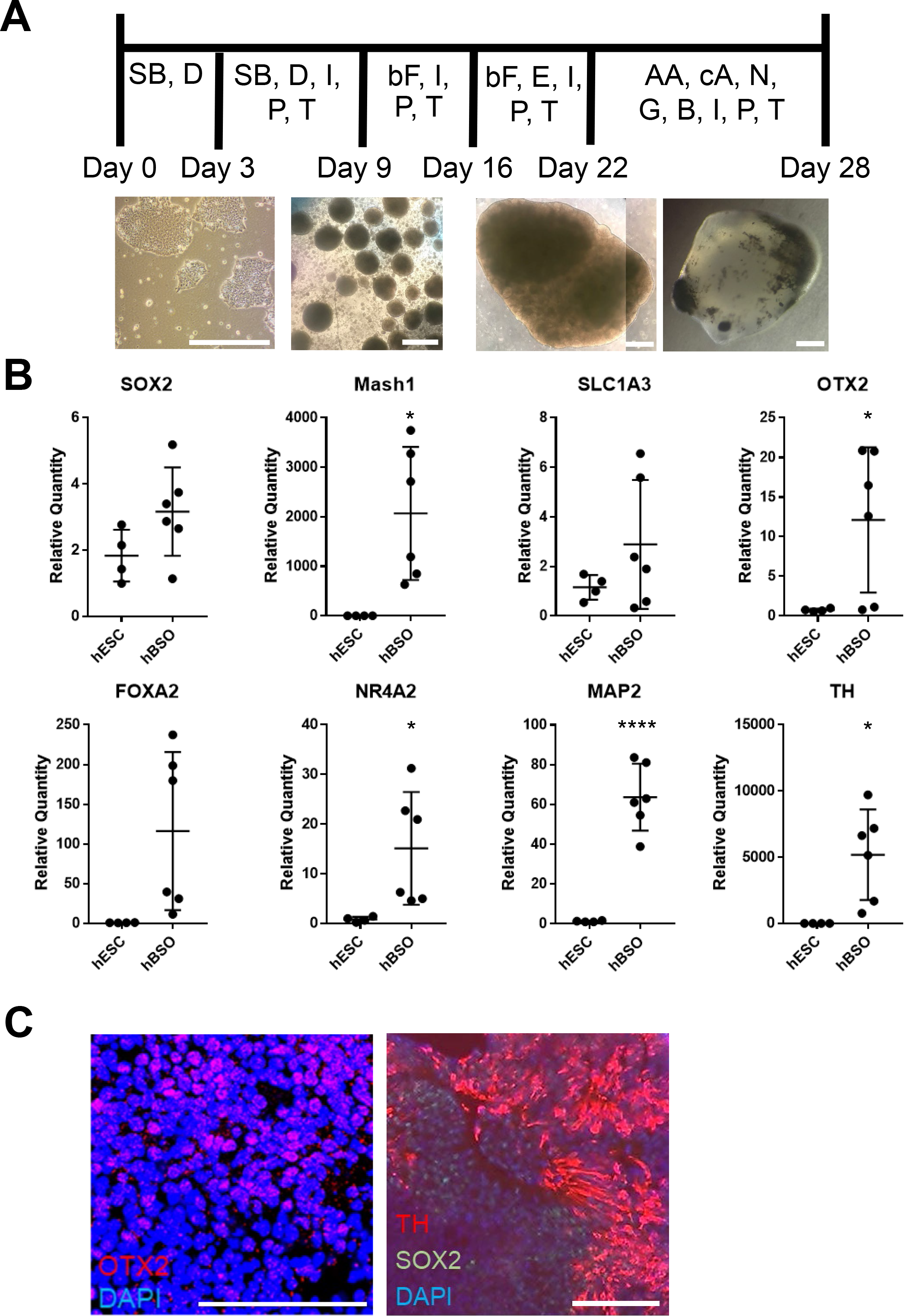
Schematic procedure of inducing hBSOs. (A) Schematic procedure of generating human brainstem organoids. SB, SB431542; D, dorsomorphin; I, insulin; P, progesterone; T, transferrin; bF, basic fibroblast growth factor; E, epidermal growth factor; AA, ascorbic acid; cA, cyclic adenosine monophosphate; N, neurotrophin 3; G, glial cell line derived neurotrophic factor; B, brain derived neurotrophic factor. Pictures of organoids were taken on day0, 9, 22 and 28. Bars = 500μm (B) qPCR analysis of 1-month old human brainstem organoids for the markers of neural stem/progenitor cell (SOX2, Mash1, SLC1A3), mature neuron (MAP2), midbrain (OTX2) and mDA (FOXA2, NR4A2, TH). Error bars indicate mean ± SEM; *p = 0.0167 (Mash1), *p = 0.0412 (OTX2), *p = 0.0387 (NR4A2), ****p < 0.0001 (MAP2), *p = 0.0178 (TH). (C) Immunohistochemical staining of midbrain marker (OTX2), mDA marker (TH) and neural stem cell marker (SOX2) at 1 month. Bars = 100μm.

Based on a quantitative PCR (qPCR) analysis of one-month old hBSOs, we confirmed distinct expression of various markers for neuronal cells. Additionally, we detected the neural stem/progenitor cell-markers *SOX2*, *Mash1*, *SLC1A3* and *OTX2*, which are necessary for the development of anterior brain structures including the midbrain (Wurst and Prakash, 2014) (Figure 1B). Our analyses also revealed expression of *FOXA2*, a potent inducer of midbrain dopaminergic (mDA) progenitors (Kittappa et al., 2007; Lin et al., 2009; Norton et al., 2005; Ribes et al., 2010; Sasaki and Hogan, 1993), and *NR4A2*, which is essential for both the survival and final differentiation of ventral mesencephalic late dopaminergic precursor neurons into dopaminergic neurons (Saucedo-Cardenas et al., 1998) (Figure 1B). Consistently, we detected the expression of mRNAs coding for the pan-neuronal marker *MAP2*, and the mature dopaminergic neuronal marker *TH* (Figure 1B). Using immunohistochemistry (IHC), we demonstrated that the organoids have immature/mature midbrain components via the detection of protein expressions of SOX2, OTX2, and TH, demonstrating that the organoids contain immature/mature midbrain components (Figure 1C). Other midbrain or mDA markers were also detected in one-month hBSOs by qPCR (Figure S2).

Furthermore, we observed expressions of *ChAT* on qPCR and immunohistochemistry (Figure 2A). GBX2 is a hindbrain marker that plays a role in the positioning of the midbrain/hindbrain boundary with OTX2. Its expression suggests that the hBSOs included midbrain and hindbrain population (Waters and Lewandoski, 2006) (Figure 2A). The expression of DBH, a marker for the central noradrenergic nervous system, may indicate that pons and medulla components are contained in the hBSOs (Swanson and Hartman, 1975). The detection of ChAT, a marker for cholinergic neurons, suggests the existence of medulla population (Stornetta et al., 2013). On IHC, melanin in the hBSOs were detected by hematoxylin-eosin, Fontana-Masson and HMB45 stainings (Figure 2B), showing that these dark cells are melanocytes derived from neural crest cells in the organoids. The expression of *SOX9*, which plays a role in the migratoin of neural crest cells (Spokony et al., 2002), also support the existence of neural crest population in the brainstem organoids (Fig. 2B). On the qPCR analysis for melanocyte markers such as *MLANA*, *PMEL*, *TYR*, we found the expression of these markers to be varied across samples (Fig.2B). Also, we detected *VGLUT1* and *GAD67*, markers for mature and functional excitatory and inhibitory neurons respectively (Fremeau et al., 2004; Soghomonian and Martin, 1998). The expression of *OLIG2* and *MBP* indicated that our organoids contained oligodendrocyte progenitors and mature oligodendrocytes (Wei et al., 2003), whereas the presence of S100β suggested the existence of astrocytes (Figure 2C). In addition, the qPCR analysis on three-month old hBSOs demonstrated the expressions of a variety of neuronal components (Figure S3).

**Figure 2.**
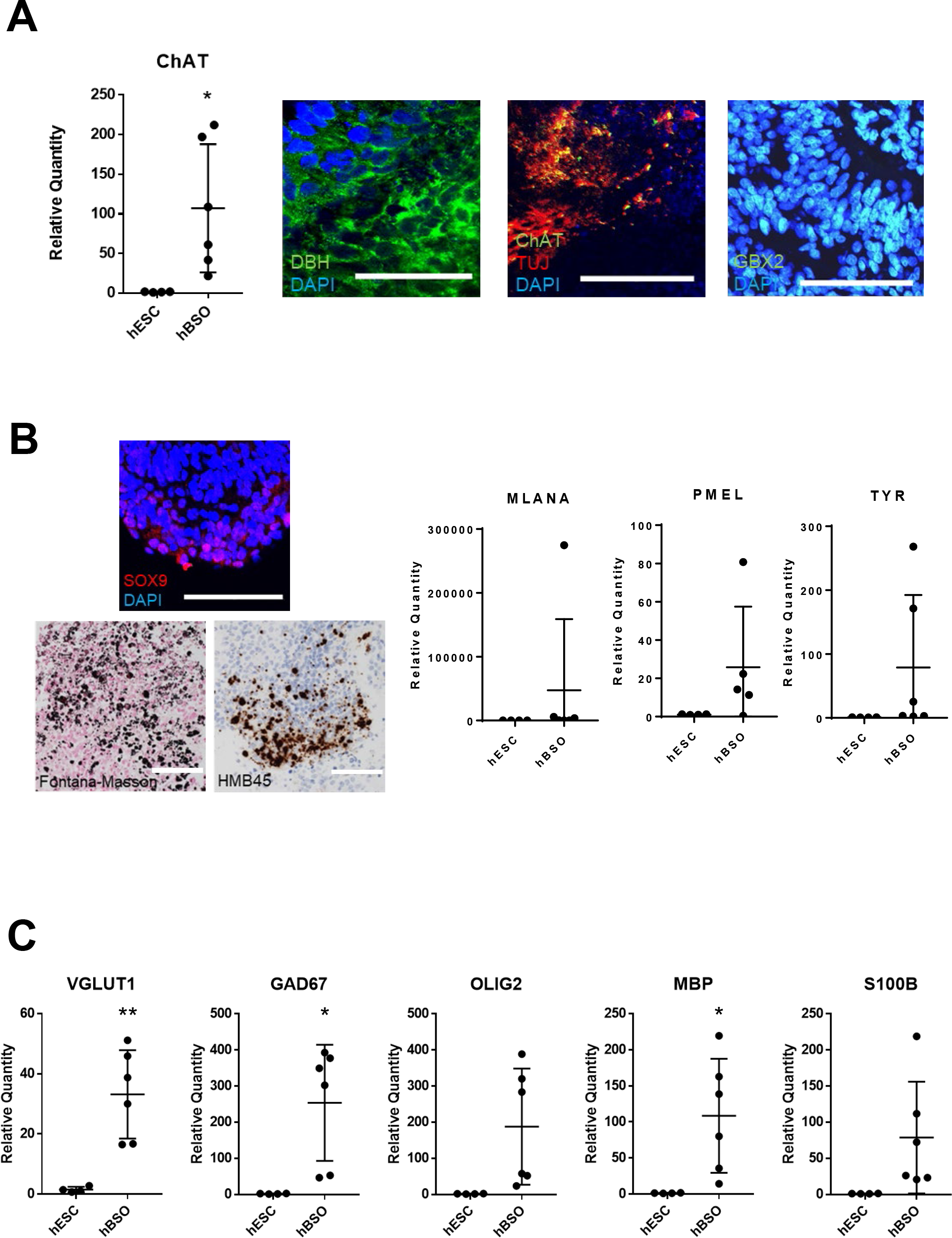
qPCR and immunohistochemical analysis of 1-month old hBSOs. (A) qPCR analysis for the marker of cholinergic neuron (ChAT). Error bars indicate mean ± SEM; *p = 0.0338 (ChAT). Immunohistochemical staining of DBH, the marker of noradrenergic neuron, ChAT and hindbrain marker (GBX2). Bars = 100μm. (B) qPCR analysis for the marker of melanocyte (MLANA, PMEL, TYR). Error bars indicate mean ± SEM. Immunohistochemistry of neural crest cell (SOX9) and melanocyte (Fontana-Masson, HMB45). Bars = 100μm. (C) qPCR analysis for the marker of excitatory neuron (VGLUT1), inhibitory neuron (GAD67), oligodendrocyte (OLIG2, MBP) and astrocyte (S100β). Error bars indicate mean ± SEM; **p = 0.0029 (VGLUT1), *p = 0.0155 (GAD67), *p = 0.0288 (MBP).

To further verify the translated products in the brainstem organoid, we performed protein mass spectrometric analysis of the organoids at one month. This analysis on the brain organoids demonstrated that the organoids expressed various neuroectoderm-specific proteins, such as NESTIN, GFAP, TBR1, TUJ1, MAP2, SYN1, S100β, and midbrain-specific protein, OTX2 (Figure 3).

**Figure 3.**
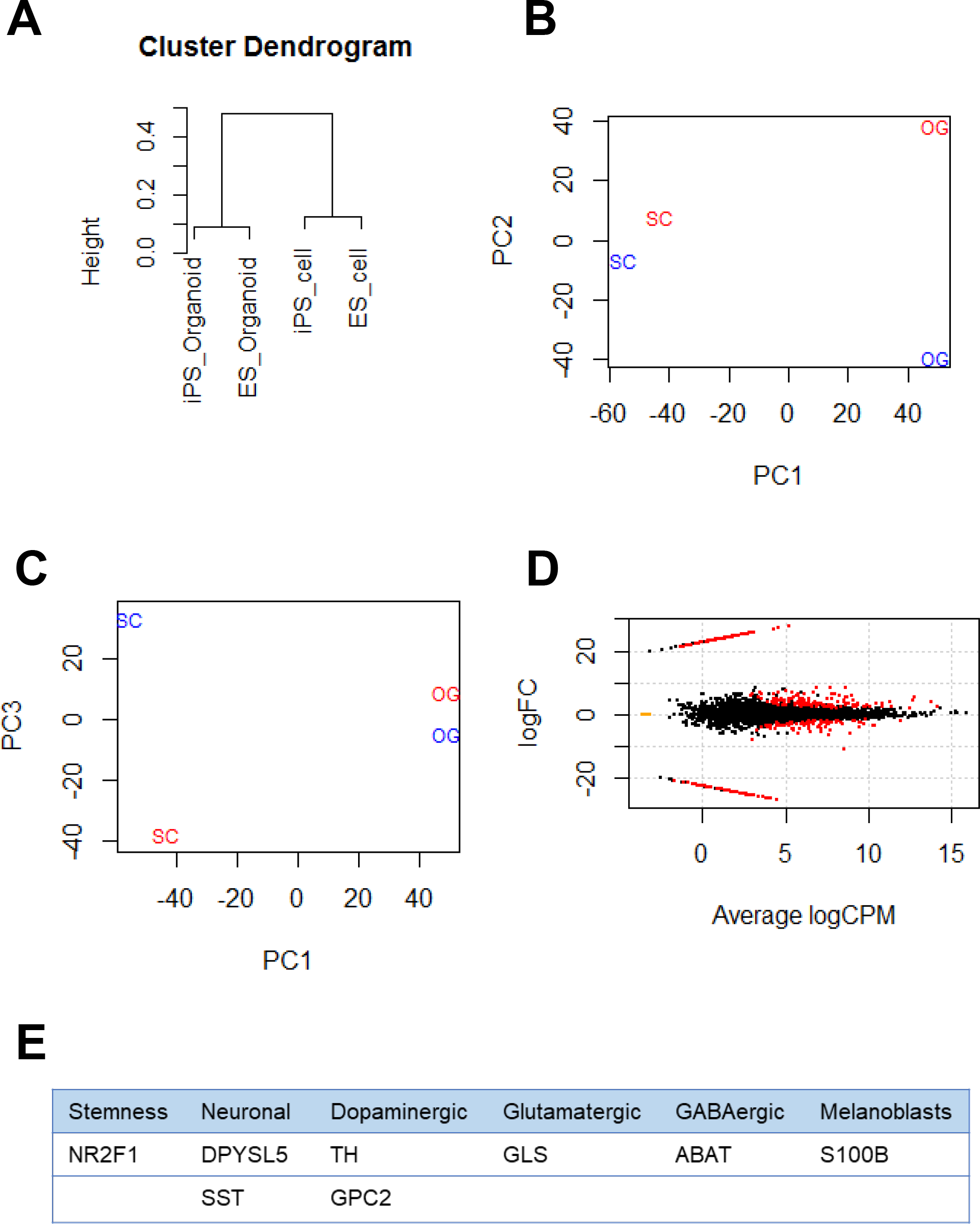
Mass spectrometric analysis of hBSOs. (A) Cluster dendrogram of hPSCs and human brain organoids. (B-C) PC analysis of hPSCs and human brain organoids. red: hiPS cell derived, blue: hES cell derived, SC: stem cell, OG: organoids. (D) MA plot for differential expression analysis (FDR<0.02) (E) Table shows list of differentially expressed protein by mass-spec. (FDR<0.05)

The detection of SYN1 protein, essential for synaptic transmission, prompted us to assess the electrical functionality of the neurons in the hBSOs. To this end, we performed electrophysiological characterization using the whole-cell patch clump method. Most cells displayed neither an action potential nor a membrane potential below −40 mV immediately after patch membrane rupture. One cell showed hyperpolarizing voltage responses with an obvious voltage sag, defined as a fast hyperpolarization, followed by a slow depolarization (Figure 4-left-1, arrow), whereas other cells (n = 8) showed no sag (Figure 4-middle-1, 4-right-1). In the neurons exhibiting repetitive firings, the spike overshot (52.9 ± 5.5 mV in amplitude) and its width was narrow (1.1 ± 0.3 ms of the half width). However, in the neuron exhibiting a few spikes, the spike amplitude was small (33.9 ± 7.9 mV) and the half width was wide (2.8 ± 1.6 ms), suggesting that these neurons were still in the course of development.

**Figure 4.**
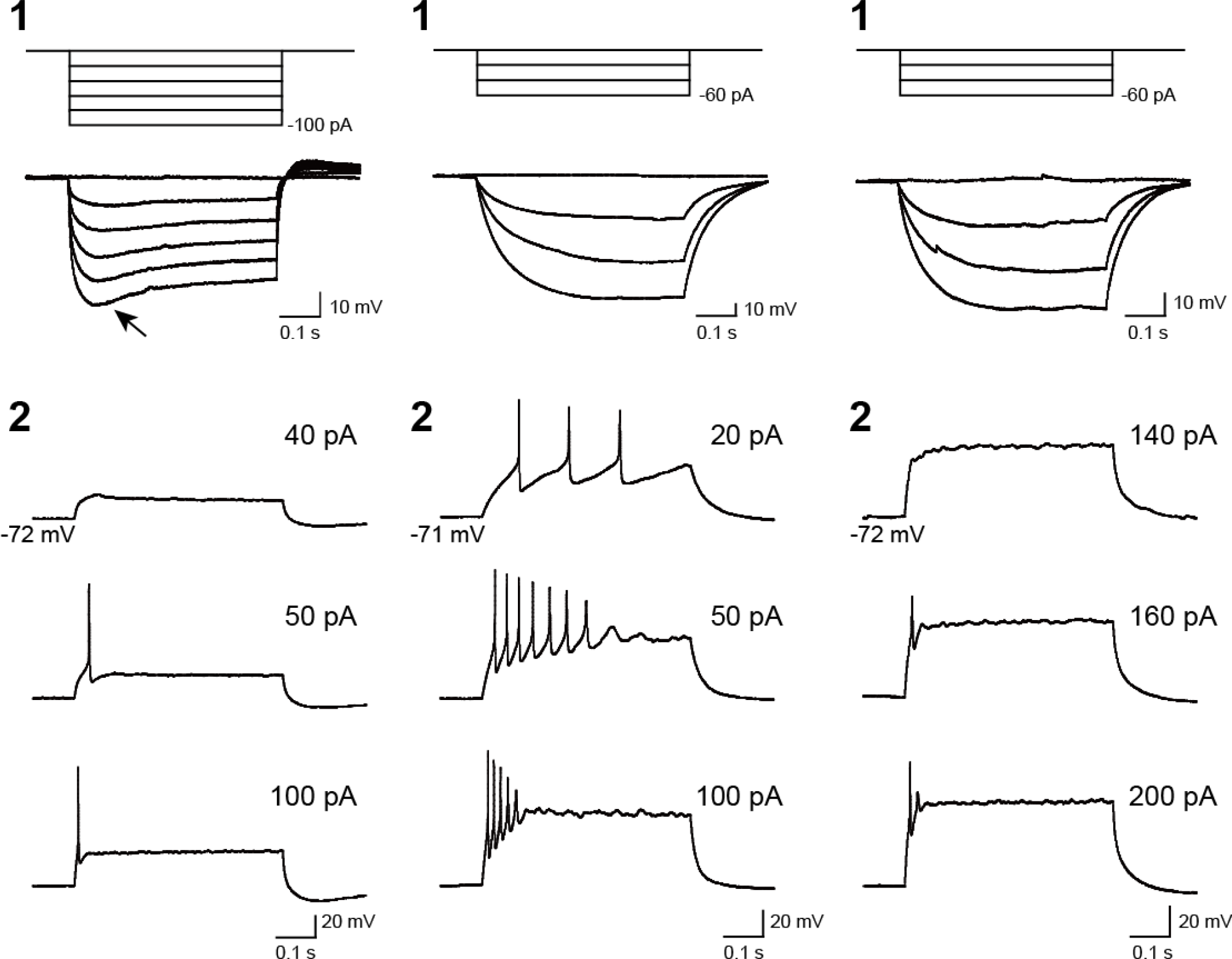
Voltage responses of recorded cells in hBSOs to current pulses. (left, middle, right) Three types of cells exhibiting different hyperpolarizing and firing responses. (1) Voltage responses to hyperpolarizing current pulses. Arrow: voltage responses characterized by a voltage sag. (2) Firing responses to depolarizing current pulses. Firing responses with multiple spikes (left), immature spikes (middle), and a few spikes (right). The values of depolarizing current pulses are given at *right*.

To further analyze the gene expression profile of hBSOs, we carried out total RNA sequencing (RNA-seq) analysis of human cerebral organoids (hCOs) induced by Lancaster et al.’s protocol (Lancaster and Knoblich, 2014; Lancaster et al., 2013) and the hBSOs at day 28. RNA-seq analysis revealed that the hBSOs contained cell populations similar to that of a human brainstem. At the age of one month, the hBSOs expressed genes that were characteristic of a fetal midbrain, such as *LMX1A* and *LMX1B*, and those indicating dopaminergic neuronal property, such as *EN1*, *EN2*, *TYR*, and *TH*, whose expression was stronger than in hCOs (Table S1). Additionally, *MLANA* and *MITF*, known as melanocyte-marker genes, and *MBP*, a marker for mature oligodendrocytes, also showed higher expressions in the hBSOs. We also observed significant expression of NGF and SOX9, specific for neural crest-stem cells. On the other hand, cortical neuron specific markers, such as Reelin and Lhx2, were lower in the hBSOs than the hCOs, indicating their distinct cellular populations.

To better understand the molecular mechanism regulating the differentiation of the hBSOs, we identified the differentially expressed genes (DEGs, Figure 5A) between the hBSOs and human embryonic stem cells (hESCs) (Table S2) and between the hCOs and hESCs (Table S3). We detected 91 DEGs that were selectively regulated in the hBSOs (Table S4) and 215 DEGs in the hCOs (Table S5).

**Figure 5.**
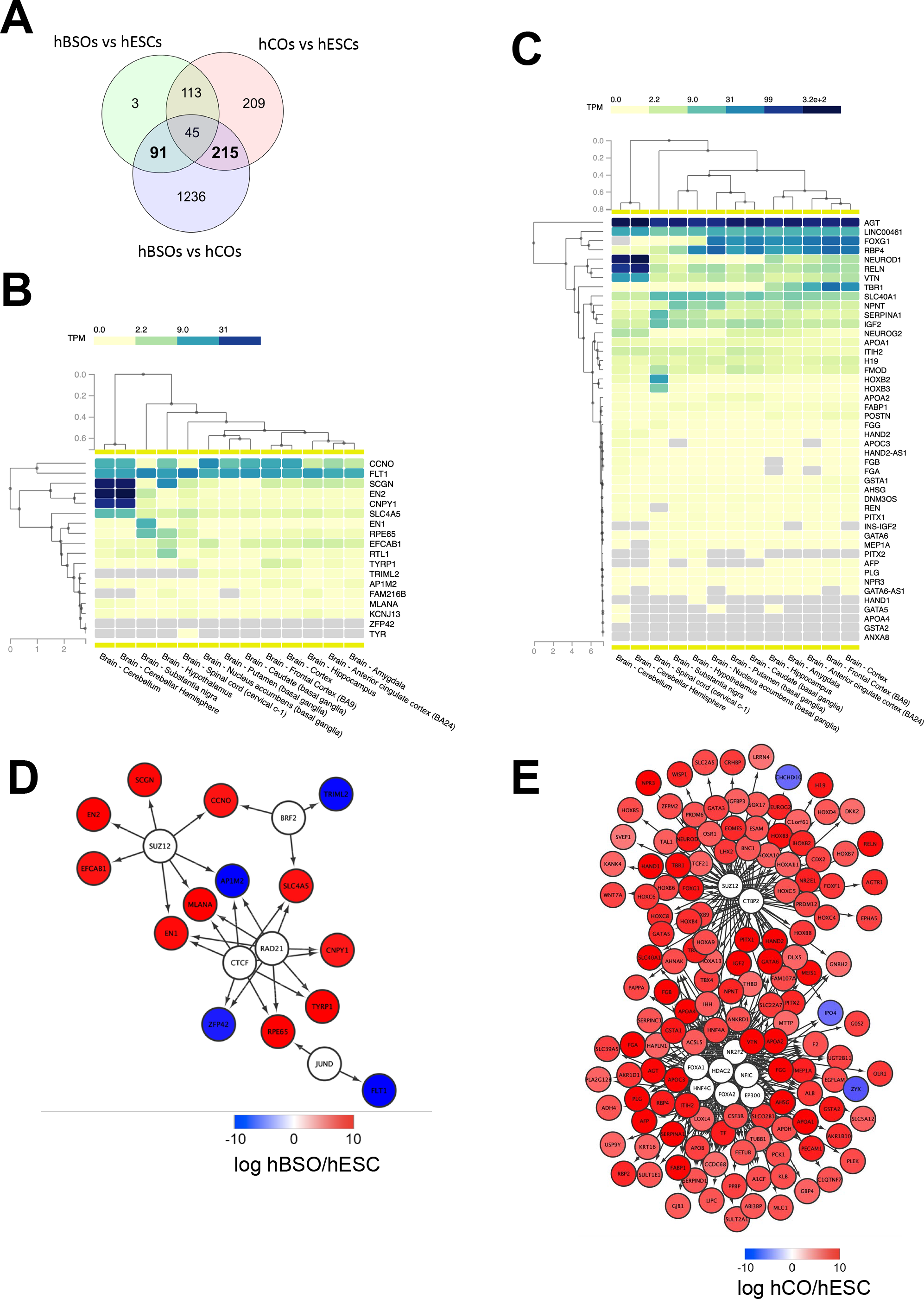
RNA-seq transcriptomic analysis of hBSOs. (A) Venn diagrams of the number of genes differentially expressed between human brainstem organoids (hBSOs), human cerebral organoids (hCOs) and human ES cells(hESCs). (B) Clustering brain regions based on the expression of differentially expressed genes selective in hBSOs (FDR<10%). (C) Clustering brain regions based on the expression of differentially expressed genes selective in hCOs (FDR<3%). (D) Transcription factors that potentially regulate the differentially expressed genes in hBSOs. (E) Transcription factors that potentially regulate the differentially expressed genes in hCOs.

To analyze the correlation between the genes selectively regulated in the hBSOs, and the hCOs and various parts of the brain, we used Genotype-Tissue Expression (GTEx), a comprehensive public resource to study tissue-specific gene expression and regulation (Consortium, 2013). High expressions of *EN2*, *CNPY1* reflected the link between the hBSOs, human cerebellum and substantia nigra, while the expression of *EN1* and *RPE65* demonstrated the relationship between the hBSOs, substantia nigra and hypothalamus (Figure 5B). Low expression of *TRIML2* in the hBSOs is characteristic of the cerebellum, substantia nigra, and hypothalamus (Figure 5B).

To identify transcription factors (TFs) potentially governing the genes selectively regulated in the hBSOs and the hCOs, we applied iRegulon (Janky et al., 2014), a computational method built upon the fact that genes co-regulated by the same TF contain common TF-binding sites, and uses the gene sets derived from ENCODE ChIP-seq data (Gerstein et al., 2012). *CTCF*, *RAD21*, *BRF2*, *JUND*, and *SUZ12* were identified as potential TFs for genes selectively regulated in the hBSOs (Figure 5D). *FOXA1*, *FOXA2*, *HDAC2*, *EP300*, *NFIC*, *HNF4G*, *NR2F2*, *CTBP2*, and *SUZ12* were detected in the hCOs (Figure 5E).

Finally, to investigate heterogeneity and gene expression dynamics in hBSOs, we performed single cell RNA sequencing (scRNA-seq) analysis on one month old hBSOs. After processing, quality control and filtering, we analyzed a total of 2,345 cells expressing 19,454 genes. To identify distinct cell populations based on shared and unique patterns of gene expression, we performed dimensionality reduction and unsupervised cell clustering using uniform manifold approximation and projection (UMAP) (Figure 6A). The UMAP plot revealed ten distinct cell populations composed of various cell types. Cell populations were identified based on cluster gene markers (Table S6) and the expression of known marker genes. We could not annotate cluster 1 and termed this cluster as “unknown” (U). Dot plot showed a selection of genes that can be used to identify cell population types (Figure 6B). Each cell population expressed canonical cell type markers. Violin plots showed the expression intensity distribution of the marker genes in each cluster (Figures 6C, S4). The neuronal progenitor cell (NP) clusters expressed genes of cell proliferation (e.g. *MKI67*) and neural stem cell markers (e.g. *PLAGL1*). The midbrain (MB) clusters expressed genes of midbrain progenitors (*OTX2*), dopaminergic (*FGFR2*), GABAergic (*ABAT*), and serotonergic (*HTR2C*) markers, indicating that the MB clusters contained various neuronal subtypes (Besse et al., 2015; Quilter et al., 2012; Ratzka et al., 2012). The cerebral cortex (CC) cluster expressed genes of mature neurons (*MAP2*, *NEFM*, *NEFL*), cholinergic (*ACHE*) and Glutamatergic (*SLC17A6*) markers. The cerebellum (CE) cluster expressed genes of cerebellar or medulla formation (*ZIC4*). The inflammation/melanocyte cluster expressed genes of microglia cells (*AIF1*), endothelial cells (*ICAM1*), and melanoblast (*PMEL*) markers. The scRNA-seq indicated that the organoids contain various cell types and neuronal subtypes.

**Figure 6.**
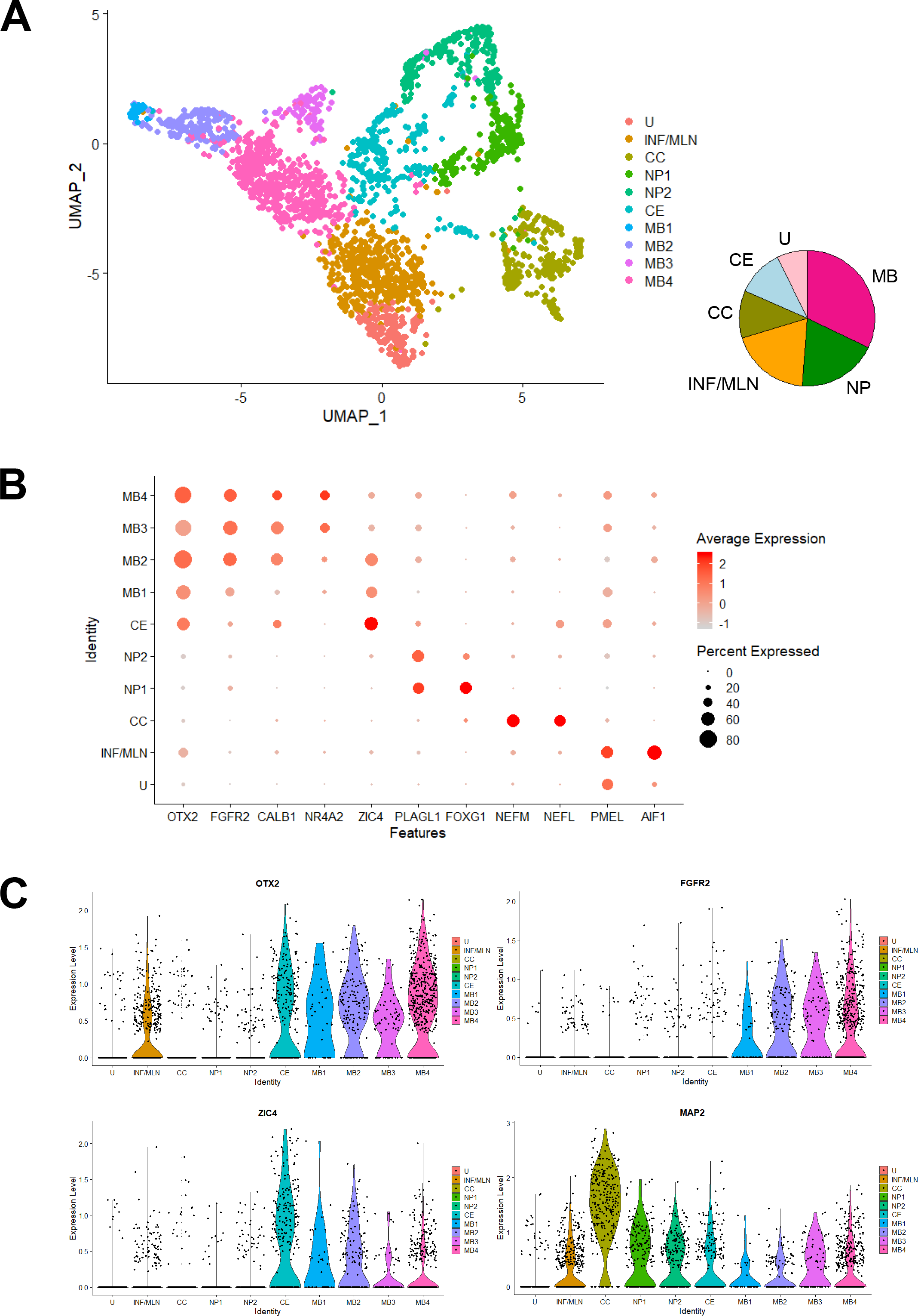
scRNA-seq analysis of the hBSOs. (A) Unsupervised clustering of all cells from human brainstem organoids. INF, inflammation; MLN; melaninocyte, CC; cerebral cortex; NP, neural progenitors; CE, cerebellum; MB, midbrain; U, unknown. (B) Dot plots showing a selection of genes that identify cell population types. (C) Cell distribution plot of OTX2, FGFR2, ZIC4 and MAP2.

## Discussion

To the best of our knowledge, our current study successfully induced hBSOs with dark cells like melanocytes for the first time. These cells are neural crest origin and derived from the fetal midbrain-hindbrain boundary. Using qPCR, IHC, RNA-seq, scRNA-seq, we observed a gene expression profile similar to that of a human fetal brainstem in hBSOs that were cultivated for 28 days. We built our current protocol upon existing protocols for human brain organoids. In particular, we were inspired by Lancaster and colleagues (Lancaster and Knoblich, 2014; Lancaster et al., 2013) who found that their organoids, acquired with the least use of growth factors, were composed of neuro-ectodermal tissues with multiple identities, such as cerebral cortex, hippocampus and retina. In another key study, Muotri and colleagues reported a protocol where cortical organoids were induced with electrophysiologically-active neurons with glutamatergic and GABAergic signaling (Trujillo et al., 2018). We designed our protocol based on these existing reports, and made use of selected hormones, such as human insulin, transferrin, and progesterone, that have been found to support the survival and differentiation of dopaminergic neurons and neural crest lineage cells (Ciarlo et al., 2017; Frith and Tsakiridis, 2019). Of note, our findings suggest that the use of an orbit shaker might have contributed to quick induction of hBSOs, through the achievement of an ideal concentration gradient of growth factors.

The hBSOs and previously reported midbrain organoids contained characteristic dark spots mimicking the existence of neuromelanin. Given that neuromelanin is derived over a few years, through an accumulation of catecholamine-derived wastes, the dark spots observed in previously-reported midbrain-organoid protocols may have melanocytes such as those found in the hBSOs. However, to our knowledge, no existing study on midbrain organoids has identified such neural crest-derived cells in their organoids.

Additionally, our RNA-seq data showed that the organoids in our study not only include monoaminergic neurons, but also melanoblasts at the dark region in the organoids, and that they highly express melanoblast-specific genes such as MELAN-A, DCT, MITF and PMEL17 (Baxter and Pavan, 2003). Our findings indicate that the hBSOs mock broad fetal brainstem region and surrounding neural crest, as cranial melanoblasts are known to originate from the neural crest around the midbrain and spread to the whole cranial region (Adameyko et al., 2012; Denecker et al., 2014). Our observations of the high Wnt1 expressions in the hBSOs, which characterize the neural crest (Adameyko et al., 2012), is consistent with this idea. Our current findings also provide the basis for future research on hereditary diseases caused by neural crest migration-disorder and PD; it has been suggested that the expression level of MITF, specific for melanocyte and melanoma, influences the risk of PD (Lubbe et al., 2016).

We would like to propose three potential applications based on our current findings: the application of hBSOs as grafts to neurodegenerative diseases, a tool for drug screening and an efficient tool for the modeling of neural crest disorders. Our observations suggest that hBSOs can be effectively used as grafts to treat patients suffering from diseases affecting the brainstem such as PD. The transplantation of human induced pluripotent stem cells (hiPSC)/hESC-derived dopaminergic neuronal progenitors to PD patients is currently under trial (Li et al., 2016). Despite the positive outcomes observed in the past studies, it is important to note that these existing trials use fetus-derived midbrain tissue, composed of multiple cells, such as neurons, astrocytes, oligodendrocytes, and neural stem cells as grafts, not the purified cell populations under trial in the current study. Given that functional defects in non-neuronal cells, such as astrocytes and microglia, contribute to the onset of PD (Joe et al., 2018), hBSOs composed of multiple brainstem-like cells similar to a fetal brainstem instead of purified dopaminergic neurons, may become better grafts for transplantation therapy against PD.

Secondly, the quick induction of hBSOs will enable more efficient drug screenings and accelerated research on the molecular mechanisms driving brainstem neurodegenerative diseases. In a recent study by Quadrato and colleagues where Lancaster et al.’s protocol (Lancaster and Knoblich, 2014; Lancaster et al., 2013) was applied, they reported the presence of SYN1+ neurons in cerebral organoids at the age of three months, but not in one month (Quadrato et al., 2017). Using our current protocol, we observed SYN1 protein expression even at one month; this may be a consequence of the regional identity of each organoid. Electrophysiological analysis of the hBSOs at the age of 1.5 months revealed neurons that exhibited action potentials and hyperpolarizing responses with voltage sags attributed to the activation of hyperpolarization-activated cyclic nucleotide-gated (HCN) cation channels (He et al., 2014; Robinson and Siegelbaum, 2003). This finding suggests that HCN channels, as well as spike generating Na^+^ and K^+^ channels, are expressed at an early stage of the human brainstem organoid. It also suggests the presence of a heterogeneous neuronal population that is capable of exhibiting distinct electrophysiological properties in the organoid.

A third potential application of the hBSOs lies in their utility in investigations into the interaction between the brainstem and neural crest cells. For example, brainstem functions are reported to be affected in representative neural crest disorders, DiGeorge syndrome and Waardenburg-Shah syndrome (Nusrat et al., 2018; Wang et al., 2017). Nonetheless, the disease models of such diseases have yet to be established and their pathologies remain to be known. We see the potential in applying the hBSOs developed in our current study as they contain neural crest cells and can be powerful tools for elucidating the mechanisms driving such neural crest diseases.

## Experimental Procedures

### Cell culture

Human iPSCs and ESCs are maintained in feeder-free condition with mTESR-TM1 media. Human iPSC line (XY) was obtained from Takara, and Human H9 ESC line (WA09, Wicell Research Institute, Madison, WI, USA) was purchased from WiCell. ES and iPS cells were cultivated in mTeSRTM1 medium (catalog # 05851, Stemcell Technologies, Vancouver, British Columbia, Canada), based upon feeder-free culture protocols on six-well plates (catalog # 3506, Corning, Corning, New York, USA) coated with growth factors reduced Matrigel (catalog # 356230, BD Biosciences, San Jose, CA, USA). At the time of passage, we added ROCK inhibitor (final concentration 10 μM, catalog # S-1049, Selleck Chemicals, Houston, Texas, USA).These cells were maintained with daily medium change without ROCK inhibitor until they reached about 70% confluency. Then, they were detached by versene solution (catalog # 15040-066, Thermo Fisher Scientific, Waltham, MA, USA) and seeded by 1:20 dilution ratio.

### Human cerebral organoid generation

The hCOs were generated as previously reported (Lancaster and Knoblich, 2014; Lancaster et al., 2013). Human iPSCs/ESCs were detached and subjected to EB induction using the protocol. After four days, half of the media was replaced by human EB medium without ROCK inhibitor and basic FGF. After two days, the EBs were transferred into a neural induction media, and embedded in matrigel after five days. The organoids were subsequently induced by the use of an orbital shaker, following the original protocol.

### Human brainstem organoid generation

The hBSOs were generated with some modifications on the cortical organoid protocol (*9, 14*). Human iPSCs/ESCs were gently dissociated by ten minutes of treatment with 50% Accutase in PBS. Detached cells were transferred to six-well plates at the density of four million cells in 5ml mTESRTM1 medium with 5μM ROCK inhibitor, 1mM dorsomorphin and 10mM SB431542 per one well in six-well plates on the orbit shaker to keep the cells under suspension. For neural induction from day 3, media was switched to one composed of Neurobasal media and 2× Gem21NeuroPlex, 1× NEAA, 1× Glutamax, 1mM dorsomorphin, 10mM SB431542, 10 μM Transferrin, 5mg/l human Insulin, and 0.063mg/l Progesterone. After nine days of exposure to dorsomorphin and SB431542, we treated the cells with 20ng/ml bFGF to induce NPC proliferation in the presence of Neurobasal-A media, supplemented with 2× Gem21NeuroPlex, 1× NEAA, 1× Glutamax, 10 μM Transferrin, 5mg/l human Insulin, and 0.063mg/l Progesterone till day 16. Cells were then kept in the same media containing not only 20ng/ml bFGF, but also 20ng/ml EGF until day 22. After day 22, EGF and bFGF were replaced by Ascorbic acid, cAMP, BDNF, GDNF, and NT-3. After day 28, cells were cultivated without any growth factors for neuronal maturation. Organoid results were combined from at least three separate batches of inductions.

### Immunohistochemical analysis

Each human cerebral or brainstem organoid was fixed in 4% paraformaldehyde in Phosphate-Buffered Saline (PBS) overnight at 4°C, dehydrated with 30% sucrose in PBS and embedded in OCT Compound (23-730-571, Thermo Fisher Scientific). Cryostat sections (14 μm) were cut and mounted onto slides (Thermo Fisher Scientific). Mounted sections were incubated for 1 hour at room temperature with blocking solution [3% normal donkey serum+0.3% Triton X-100 in Tris-Buffered Saline (TBS)] and incubated with primary antibodies diluted in blocking solution overnight at 4 °C. After three washes with TBS, corresponding fluorophore-conjugated secondary antibodies diluted in the blocking solution were added and incubated for 2 hours at room temperature and followed by DAPI staining. Finally, stained slides were rinsed with TBS three times, mounted and analyzed using a microscope. Antibodies specific for TUJ1 (1:400, T8660) and S100β (1:200, S2532) purchased from Sigma were used for immunostaining.

### RNA isolation, RT-PCR and quantitative PCR

RNA from hCOs/hBSOs and ES cells was extracted according to the protocol supplied with TRIzol reagent (15596018, Thermo Fisher Scientific). The concentration and purity of the RNA samples were measured using Spectrophotometer (Beckman Coulter). Extracted RNA samples were either shipped to Bioengineering lab for RNA sequencing analysis or subjected to RT-PCR. For RT-PCR, the extracted RNAs were reverse transcribed according to the protocol supplied with ReverTra Ace qPCR RT Master Mix (FSQ-201, TOYOBO). StepOne Plus Real-time PCR System (Thermo Fisher Scientific) was used to amplify and quantify levels of target gene cDNA. Quantitative real-time PCR (qRT-PCR) was performed with SsoAdvanced Universal SYBR Green Supermix (172-5271, Bio-Rad Laboratories) and specific primers for qRT-PCR. Reactions were run in triplicate. The expression of each gene was normalized to the geometric mean of β-actin as a housekeeping gene and analyzed using the ΔΔCT method. Statistical significance was calculated by the Mann–Whitney test.

### RNA sequencing

Total RNA was isolated from the cells using the PureLink RNA Mini Kit (12183018A) according to the manufacturer’s instructions. RNA concentration was analyzed by Qubit RNA HS Assay Kit (Thermo Fisher Scientific) and the purity was assessed using the Qsep100 DNA Fragment Analyzer and RNA R1 Cartridge (BiOptic). Afterwards, total RNA was converted to cDNA, and used for Illumina sequencing library preparation based on the KAPA Stranded mRNA-Seq Kit protocols (KAPA BIOSYSTEMS). DNA fragments were then subjected to adapter ligation, where dsDNA adapters with 3’-dTMP overhangs were ligated to A-tailed library insert fragments by FastGene Adapter Kit (FastGene). The purified cDNA library products were evaluated using Qubit and dsDNA HS Assay Kit (Thermo Fisher Scientific), followed by quality assessment using the Fragment Analyzer and dsDNA 915 Reagent Kit (Advanced Analytical Technologies), and finally, by sequencing (2×75bp) on NextSeq 500 (Illumina).

### Transcriptome analysis

A count-based differential expression analysis “TCC” was used to identify DEGs in the RNA-seq data with a thresholded false discovery rate of 20% (Sun et al., 2013). iRegulon (*25*) was used to identify TFs potentially regulating the DEG with normalized enrichment scores>4 as the threshold. GTEx (*27*) was used to analyze the similarity of expression pattern between organoids and various tissues in brain.

### Electrophysiology

Electrophysiological recordings of the cells in hBSOs at three months were performed. An organoid was transferred to a glass-bottom recording chamber on an upright microscope (Leica DM LFS; Leica, Wetzlar, Germany) and continuously perfused with an extracellular solution containing (in mM) 125 NaCl, 2.5 KCl, 2 CaCl2, 1 MgCl2, 1.25 NaH2PO4, 26 NaHCO3, and 25 glucose and aeration with 95% O2 and 5% CO2 (pH 7.4), at a rate of 2 ml/min. The organoid was held down by a weighted net to prevent it from moving. The bath temperature was maintained at 30—32°C using an in-line heater (TC-324B; Warner Instruments, Hamden, CT). Whole cell current-clamp recordings were performed using an EPC-8 patch-clamp amplifier (HEKA, Darmstadt, Germany). Patch pipettes were prepared from borosilicate glass capillaries and filled with an internal solution containing (in mM) 120 K-methylsulfate, 10 KCl, 0.2 EGTA, 2 MgATP, 0.3 NaGTP, 10 HEPES, 10 Na2-phosphocreatine, and 0.1 spermine, adjusted to pH 7.3 with KOH. The osmolarity of the internal solution was 280 – 290 mOsm/L and the resistance of the patch electrodes was 4 – 8 MΩ in the bath solution. The voltage signals were low-pass filtered at 3 kHz and digitized at 10 kHz. The calculated liquid junction potential of −5 mV was corrected. The data were acquired using a pClamp 9 system (Molecular Devices, Sunnyvale, CA). Voltage responses of the cells were investigated by the application of depolarizing and hyperpolarizing current pulses (400 ms in duration). Off-line analysis was performed using AxoGraph X software (AxoGraph Scientific). The input capacitance was estimated based on the current induced by a 10 mV-voltage step from a holding potential of −70 mV. The input resistance was estimated based on the voltage change induced by an applied hyperpolarizing current pulse of −40 pA. The spike amplitude was determined as the spike height from its threshold, defined as the membrane potential at which the derivative of the voltage trace reached 10 V/s. The maximum firing frequency was obtained from cells that exhibited more than 1 spike and calculated as the reciprocal of the shortest inter-spike interval between successive pairs of spikes.

### Mass spectrometric analysis

Human ES cells, iPS cells, ESC-derived organoids and iPSC-derived organoids were washed with ice-cold PBS, harvested by scraping and centrifugation, and then frozen in liquid nitrogen. The frozen cells and organoids were crushed by using Multi-beads shocker (Yasui Kikai, Japan) and subsequently lysed by sonication in 9.8 M urea with protease inhibitor cocktail (cOmplete™; Roche) and phosphatase inhibitor cocktail (PhosSTOP™; Roche). The clear lysate was collected by centrifugation and protein concentration was measured by BCA protein assay. Twenty μg of proteins were mixed with an internal standard protein mixture (10 fmol/ml) (MassPREPTM; Water) and incubated with 2 mM Tri(2-carboxyethyl)phosphine hydro-chloride (TCEP-HCl) for 30 min at 37 °C for reduction, followed by alkylation with 55 mM iodoacetamide (IAA) or 30 min at room temperature. The mixture was then diluted four-fold with 0.1 M Triethylamonium bicarbonate (TEAB) and subjected to trypsin digestion (1:40 trypsin: sample ratio) for three hrs at 37 °C. The digestion was terminated by Trifluoroacetic acid (TFA), following by desalting with a SDB-StageTip (styrene divinylbenzene) (2007). The samples were fractionated into eight fractions by using SDB-Stage Tip. Each fraction was dried by vacuum and dissolved in the measurement buffer (3% Acetonitrile (ACN) and 0.1% Formic acid (FA)). Mass spectrometry was performed as described previously (Uetsuka et al., 2015). To identify the proteins, raw data of peptides were analyzed using Proteome Discoverer 2.2 (Thermo Fisher Scientific) and Mascot 2.6 (Matrix Science, London, UK). The peptide results from all eight fractions were combined and subjected to search for the matching proteins in UniProt human database (released in August 2018). Maximum numbers of missed cleavages, precursor mass tolerance, and fragment mass tolerance were set to 3, 10 ppm and 0.01 Da, respectively. The carbamidomethylation on Cys was set as a fixed modification, whereas oxidation of Met and deamidation of Asn and Gln were set as variable modifications. A filter of false discovery rate < 1% was applied to the data.

The Minora Feature Detector node was used for label-free quantification and the consensus workflow included the Feature Mapper and the Precursor Ion Quantifier nodes using intensity for the precursor quantification. The protein intensities were normalized by the total peptides intensity. In addition, annotations from the Ingenuity Knowledge Base (IKB; released in autumn, 2018; Qiagen, Redwood City, CA, USA) and the database of Ingenuity Pathway Analysis (IPA) were used to determine the localization and functional categories of the identified proteins.

### Single cell RNA sequencing and data analysis

To dissociate hBSOs into single cells, they were incubated for ~30min in Accutase (STEMCELL TECHNOLOGIES) at 37°C. Droplet-based scRNA-seq libraries were generated using the Chromium™ Single Cell 3’ Reagent kits V2 (10× Genomics). Briefly, cell number and cell viability were assessed using the Countess II Automated Cell Counter (ThermoFisher). Thereafter, cells were mixed with the Single Cell Master Mix and loaded together with Single Cell 3’ Gel beads and Partitioning Oil into a Single Cell 3’ Chip. RNA transcripts were uniquely barcoded and reverse-transcribed in droplets. cDNAs were pooled and amplified according to the manufacturer’s protocol. Libraries were quantified by High Sensitivity DNA Reagents (Agilent Technologies) and the KAPA Library Quantification kit (KAPA BIOSYSTEMS). Libraries then were sequenced by Illumina Hiseq 2500 in rapid mode.

Raw sequencing data from the organoid was preprocessed using the Cell Ranger (v 2.2.0, 10× Genomics) software (Zheng et al., 2017). Reads were aligned to the GRCh38 human reference genome using STAR. After processing by Cell Ranger, the scRNA-seq data was analyzed using the Seurat v.3.0.0 R package (Satija et al., 2015). Cells with more than 8000 or less than 750 detected genes, as well as cells expressing more than 5% mitochondrial genes were excluded. Genes expressed in less than 3 cells were excluded. We collected a total of 2345 cells expressing a total of 19454 genes. The datasets were log normalized and scaled to 10000 transcripts per cell. The top 2000 highly variable genes were determined using the vst method. The datasets then were scaled and principal component analysis (PCA) was performed. Clustering was performed based on the top 45 principal components (PCs) using the shared nearest neighbor (SNN) modularity optimization with a resolution of 0.8. Cluster identities were assigned based on cluster gene markers determined by the ‘FindAllMarkers’ function in Seurat (Table S6).

## Author contributions

N.E., T.K.M, J.L., J.W.S., K.S., and E.M. designed the study; N.E., T.K.M, J.L., M.Matsubayashi, H.N., T.Shiota., N.I., T.Kiriyama., C.Z., T.Kouno, Y.J.L., P.K., P.W., Y.M.S., R.N., T.Komeda, N.M., F.K., M.J., S.K., M.N., M.H., Y.S., T.Shiromizu., Y.N., T.Kasai, M.T., H.K., Y.I., Y.T., M.Makinodan, T.Kishimoto, H.K., S.N., J.W.S., K.S. and E.M. conducted the research; N.E., T.K.M, J.L., Y.S., T.Shiromizu., Y.N., T.Kasai, M.T., S.N., A.R.M, J.W.S., K.S. and E.M. analyzed the data; N.E., T.K.M, J.L., K.K., Y.S., T.Shiromizu., Y.N., T.Kasai, M.T., S.N., J.W.S., K.S. and E.M. wrote the paper.

## Acknowledgements

The authors thank Jenny Hsieh (University of Texas at San Antonio), Keren-Happuch E Fan Fen, Noriko Horii (Department of Anatomy and Cell Biology, Nara Medical University) and Jens Christian Schwamborn (Luxembourg Centre for Systems Biomedicine, University of Luxembourg) for their critical reading of the manuscript. This work was supported by grants from JSPS KAKENHI [JP17H07031 to E.M., JP18H06202 to H.N., JP19K16925 to T.M., JP19K23952 to K.K., JP19K17044 to N.I., JP19K07978 to T.K., JP19K08150 to S.K., JP19K23976 to M.N.], AMED The Program for Technological Innovation of Regenerative Medicine [JP19bm0704039h to T.K.M.], AMED Osaka University Seeds (A) to T.K.M, Takeda Science Foundation to E.M. and T.K.M., Kanzawa Medical Research Foundation to E.M., Uehara Memorial Foundation to E.M., Nakatomi Foundation to E.M., Konica Minolta Science and Technology Foundation to E.M., Naito Foundation to E.M., MSD Life Science Foundation to E.M., Mochida Memorial Foundation for Medical and Pharmaceutical Research to E.M., SENSHIN Medical Research Foundation to E.M., Terumo Foundation for Life Sciences and Arts to E.M., Nara Kidney Disease Research Foundation to E.M., Novartis Research Grants to E.M., K.S. and N.E., Sumitomo Dainippon Pharma Research Grant to T.K.M., Nara Medical University Grant-in-Aid for Collaborative Research Projects to K.S. and H.K., Nara Medical University Grant-in-Aid for Young Scientists to N.E. and T.K.M., and by unrestricted funds provided to E.M. from Dr. Taichi Noda (KTX Corp., Aichi, Japan) and Dr. Yasuhiro Horii (Koseikai, Nara, Japan).

---

**Table S1: Differentially expressed genes between hBSOs and hCOs in RNA-seq analysis. Related to Figure 5.**

**Table S2: Differentially expressed genes between hBSOs and hESCs in RNA-seq analysis. Related to Figure 5.**

**Table S3: Differentially expressed genes between hCOs and hESCs in RNA-seq analysis. Related to Figure 5.**

**Table S4: Differentially expressed genes selective in hBSOs in RNA-seq analysis. Related to Figure 5.**

**Table S5: Differentially expressed genes selective in hCOs in RNA-seq analysis. Related to Figure 5.**

**Table S6: Cell populations identified based on each cluster gene markers in scRNA-seq analysis. Related to Figure 6.**

