## Supplemental Figures for "Brainstem organoids from human pluripotent stem cells contain neural crest population"

### Supplemental Information

#### Supplemental Figures and Legends

**Figure S1**

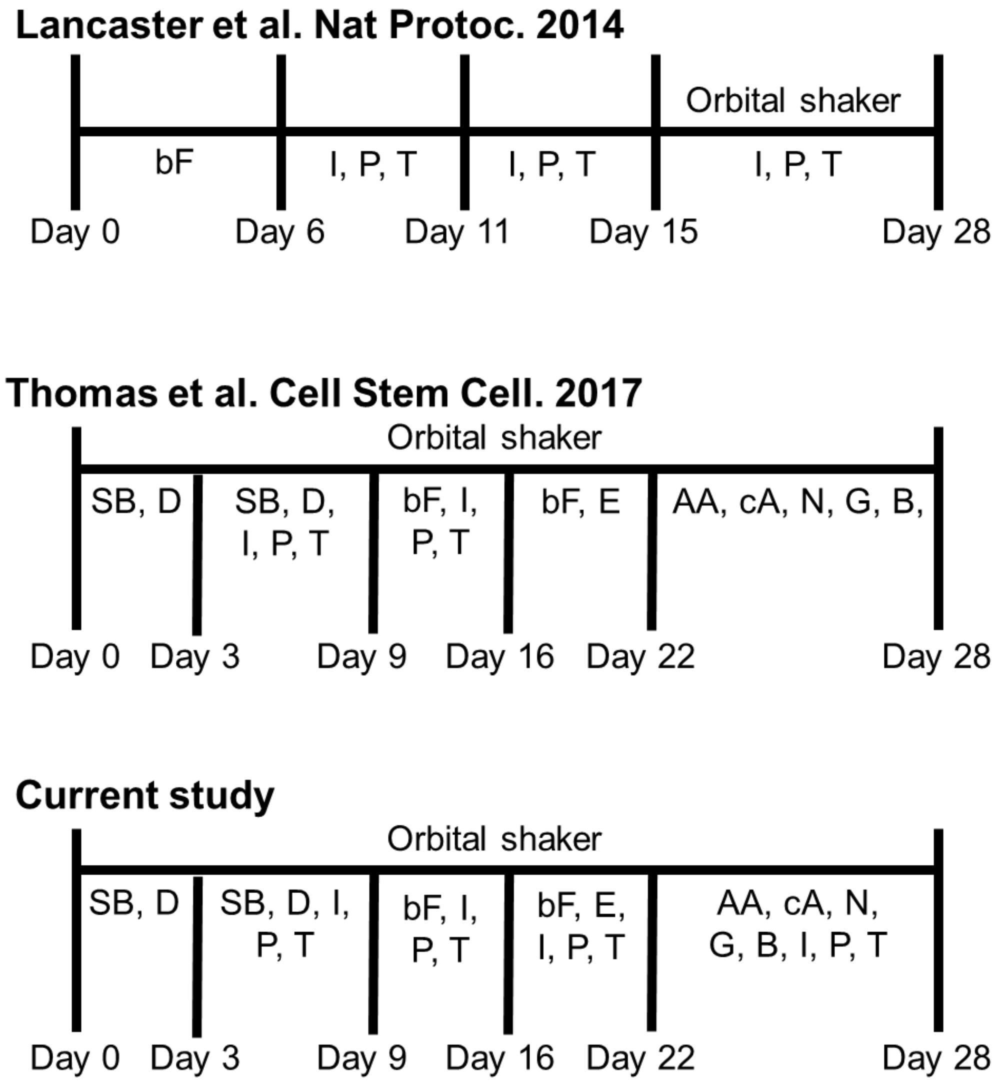


**Figure S1: Schematic procedures of brain organoid by different groups. Related to Figure 1.**

Procedures of organoid generation by three different groups, Lancaster et al., Thomas et al., and current study. SB, SB431542; D, dorsomorphin; I, insulin; P, progesterone; T, transferrin; bF, basic fibroblast growth factor; E, epidermal growth factor; AA, ascorbic acid; cA, cyclic adenosine monophosphate; N, neurotrophin 3; G, glial cell line derived neurotrophic factor; B, brain derived neurotrophic factor.

### Figure S2

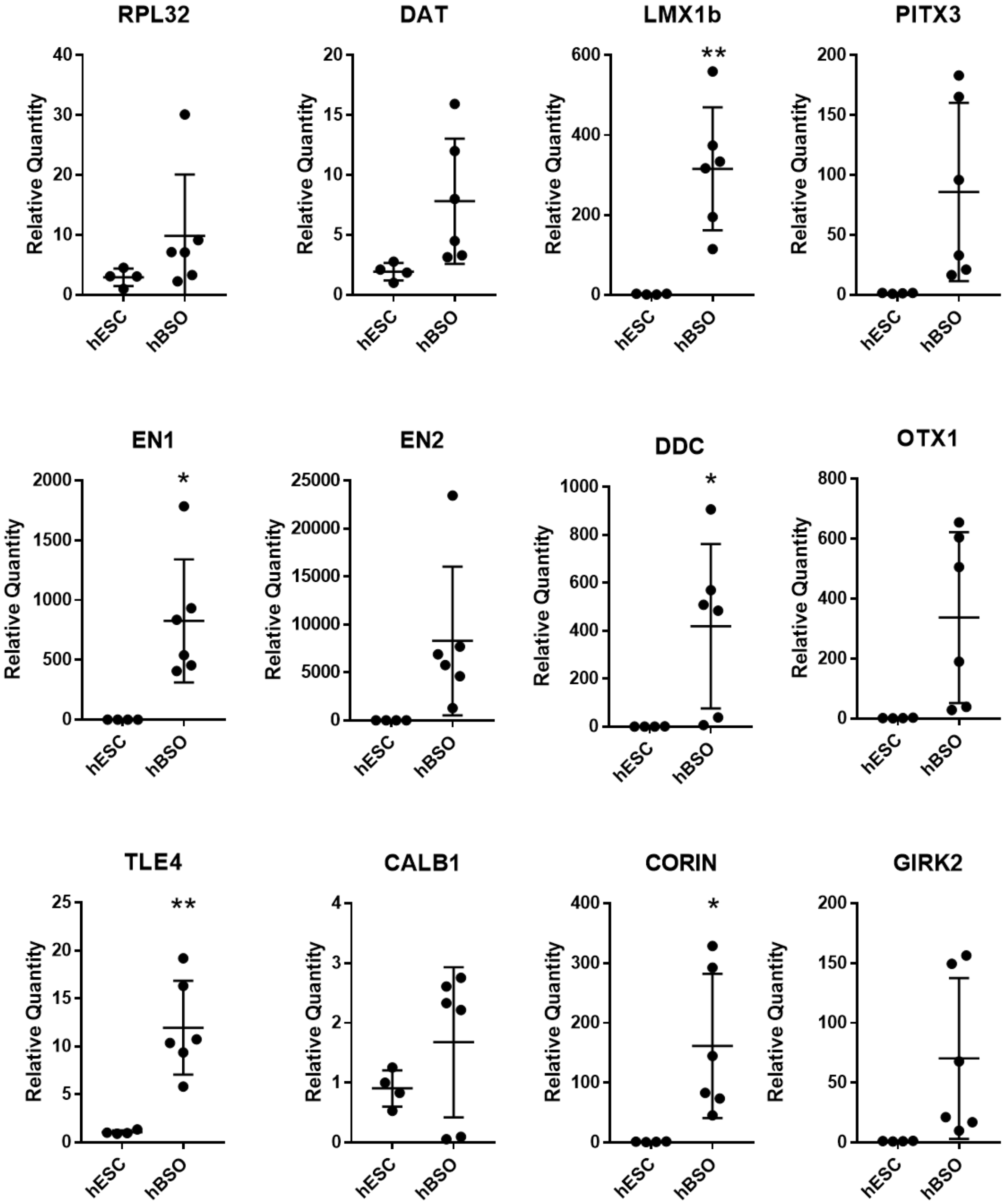


**Figure S2: qPCR analysis for midbrain or mDA markers of hBSOs at 1 month from hESCs. Related to Figures 1 and 2.**

qPCR analysis for the marker of midbrain or dopaminergic (RPL32, DAT, LMX1b, PITX3, EN1, EN2, DDC, OTX1, TLE4, CALB1, CORIN, GIRK2). Error bars indicate mean ± SEM; **p = 0.004 (LMX1b), *p = 0.0138 (EN1), *p = 0.0438 (DDC), **p = 0.0024 (TLE4), *p = 0.0316 (CORIN).

### Figure S3

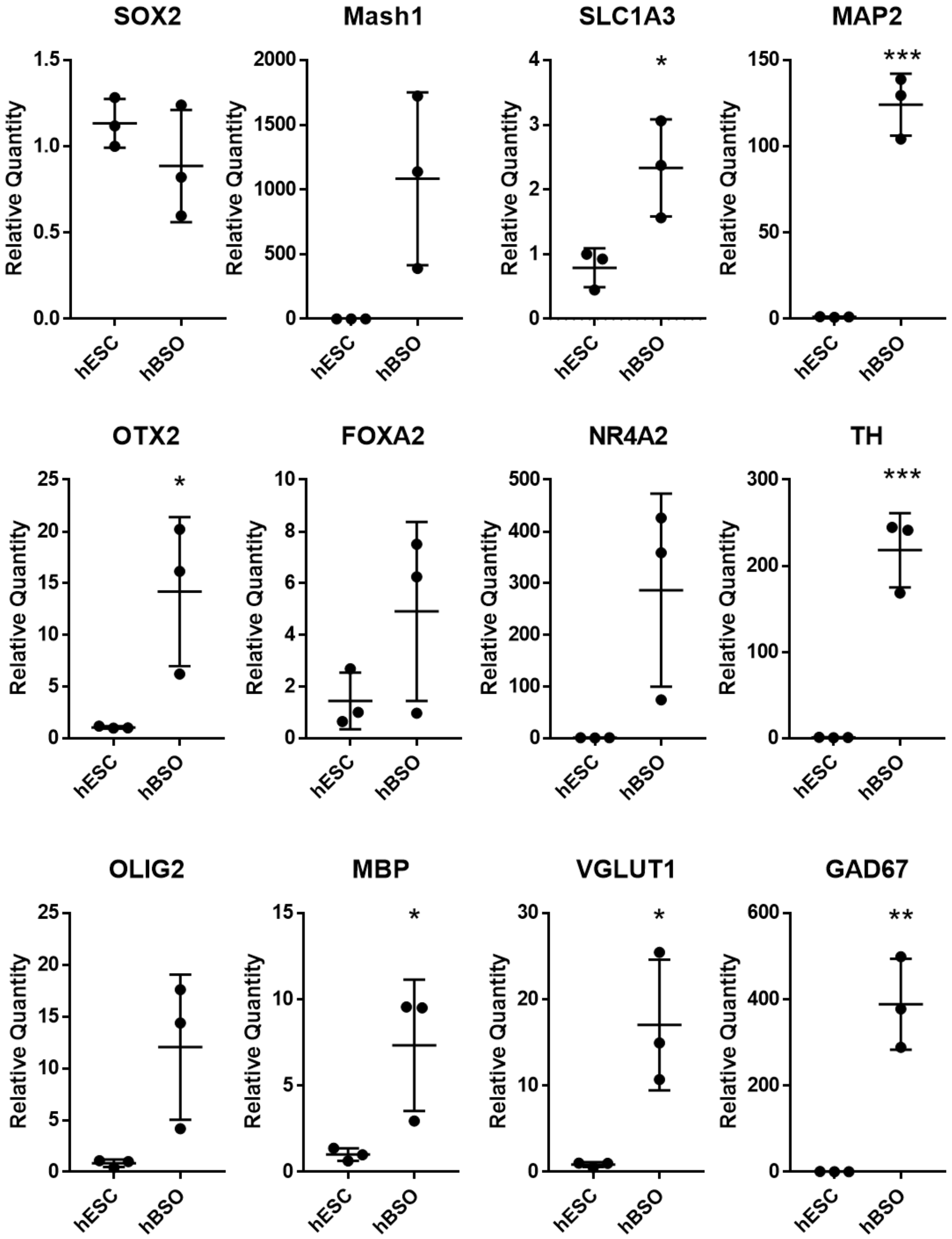


**Figure S3: qPCR analysis of hBSOs at 3 months old from hESCs. Related to Figures 1 and 2.**

qPCR analysis of 3-month old human brainstem organoids for the markers of neural stem/progenitor cell (SOX2, Mash1, SLC1A3), mature neuron (MAP2), midbrain (OTX2) and mDA (FOXA2, NR4A2, TH), excitatory neuron (VGLUT1), inhibitory neuron (GAD67), oligodendrocyte (OLIG2, MBP). Error bars indicate mean ± SEM; *p = 0.0298 (SLC1A3), ***p = 0.0003 (MAP2), *p = 0.0343 (OTX2), ***p = 0.0009 (TH), *p = 0.0453 (MBP), *p = 0.0209 (VGLUT1), **p = 0.0031 (GAD67).

### Figure S4

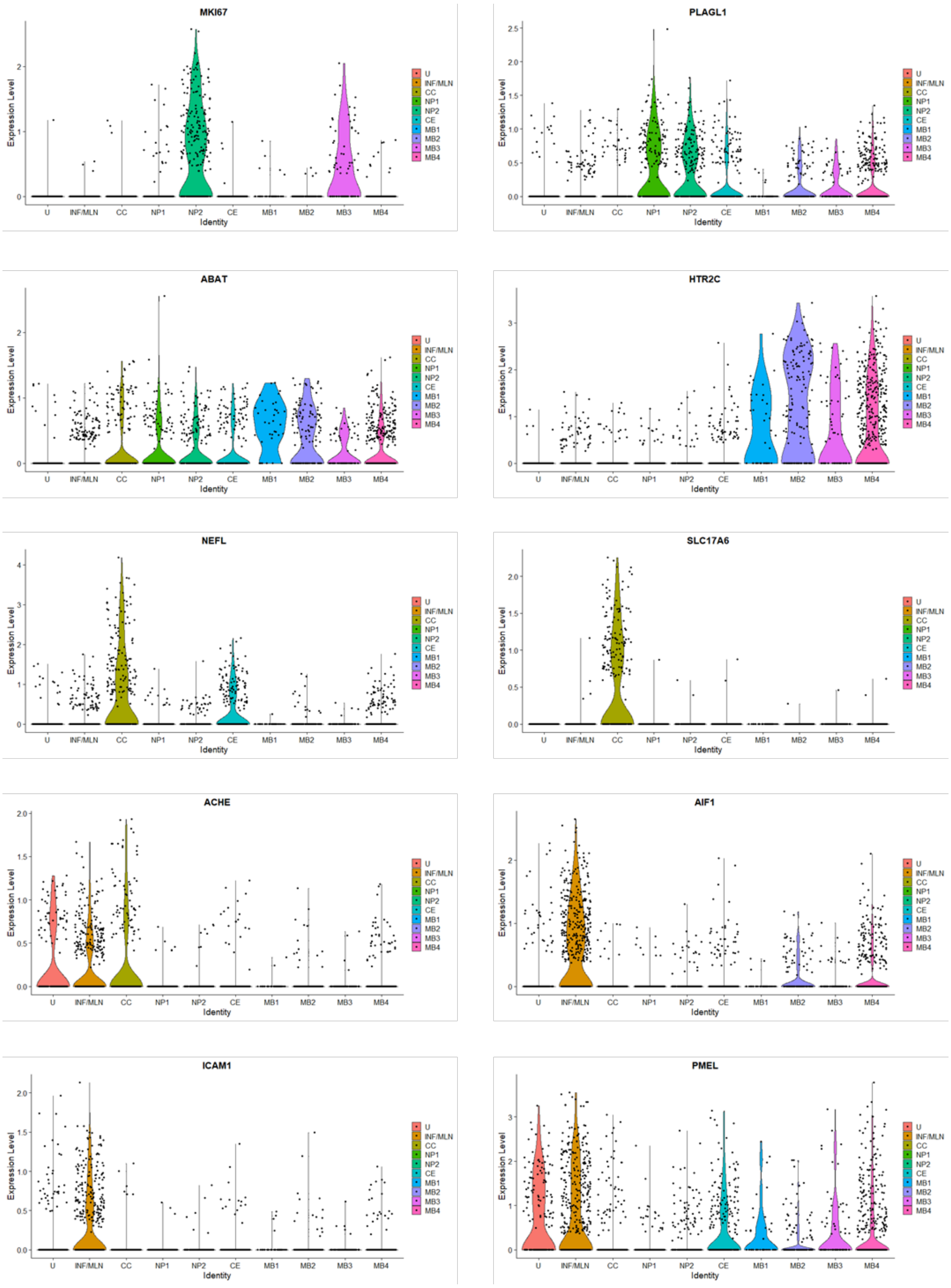


**Figure S4: scRNA-seq analysis of hBSOs. Related to Figure 6.**

Cell distribution plot of MKI67, PLAGL1, ABAT, HTR2C, NEFL, SLC17A6, ACHE, AIF1.
