## Supplemental Table 1 for "Brainstem organoids from human pluripotent stem cells contain neural crest population"

**Supplemental Table 1 Differentially expressed genes between hBSOs and hCOs**

| Gene_name | log (hBSO/hCOs) | FDR (hBSOs/hCOs) | A (hBSOs/hCOs) |
|---|---|---|---|
| A1CF | -6.98 | 5.01E-05 | -0.97 |
| A2M | -3.56 | 5.95E-03 | 9.17 |
| A4GALT | 3.39 | 2.75E-02 | 5.37 |
| AACSP1 | -4.22 | 1.29E-01 | -0.97 |
| ABCA13 | 2.68 | 1.69E-01 | 4.70 |
| ABCB5 | 8.13 | 6.27E-08 | -0.97 |
| ABCC9 | 3.63 | 9.85E-03 | 6.26 |
| ABCG2 | -5.98 | 8.50E-05 | 4.29 |
| ABHD12B | 4.22 | 2.19E-02 | -0.97 |
| ABI3BP | -2.65 | 1.18E-01 | 5.95 |
| ABLIM2 | 2.49 | 1.78E-01 | 5.60 |
| ABR | -3.25 | 3.75E-02 | 5.25 |
| ACACA | -3.34 | 5.58E-02 | 3.97 |
| ACAN | -4.43 | 8.00E-04 | 5.93 |
| ACKR3 | -2.32 | 1.78E-01 | 8.38 |
| ACP5 | 5.12 | 6.80E-05 | 6.92 |
| ACPP | -5.87 | 2.97E-04 | 3.65 |
| ACSL5 | -6.45 | 2.63E-04 | -0.97 |
| ACSM1 | 3.89 | 4.99E-02 | -0.97 |
| ACSM3 | -3.64 | 1.29E-02 | 5.12 |
| ACTC1 | -2.75 | 9.32E-02 | 5.49 |
| ADAMTS1 | -2.30 | 1.78E-01 | 9.89 |
| ADAMTS19 | -2.49 | 1.38E-01 | 6.52 |
| ADCY10 | -3.37 | 1.72E-01 | 2.99 |
| ADGB | 5.36 | 9.52E-04 | -0.97 |
| ADGRF5 | -5.73 | 4.84E-04 | 3.58 |
| ADGRG2 | 2.46 | 1.70E-01 | 6.29 |
| ADGRG6 | 2.53 | 1.15E-01 | 8.03 |
| ADH4 | -6.77 | 8.75E-05 | -0.97 |
| ADM5 | -3.30 | 1.95E-01 | 2.95 |
| ADTRP | 4.57 | 8.47E-02 | 2.64 |
| AFP | -15.69 | 3.72E-21 | 8.56 |
| AGPAT4-IT1 | -3.50 | 8.75E-02 | 3.46 |
| AGPAT5 | 3.14 | 1.14E-01 | 3.93 |
| AGT | -3.13 | 2.19E-02 | 8.11 |
| AGTR1 | -8.48 | 1.71E-07 | -0.97 |
| AGXT2 | -4.73 | 3.57E-02 | -0.97 |
| AHNAK | -3.14 | 1.84E-02 | 12.52 |
| AHSG | -11.51 | 2.68E-13 | 6.47 |
| AK7 | 2.53 | 1.13E-01 | 8.01 |
| AK8 | 3.02 | 5.78E-02 | 5.45 |
| AKAP14 | 4.78 | 1.17E-03 | 4.74 |
| AKAP4 | -5.16 | 1.26E-02 | -0.97 |
| AKAP5 | -3.92 | 4.22E-03 | 5.76 |
| AKR1B10 | -8.27 | 3.47E-07 | 3.85 |
| AKR1B15 | -6.06 | 8.75E-04 | -0.97 |
| AKR1D1 | -8.00 | 1.08E-06 | 3.71 |
| ALB | -6.97 | 6.94E-06 | 4.20 |
| ALDH1A1 | -5.70 | 1.12E-05 | 6.89 |
| ALDH1A3 | 2.38 | 1.58E-01 | 8.22 |
| ALDOB | -6.00 | 1.13E-03 | -0.97 |
| ALKAL2 | 4.95 | 3.31E-02 | 2.83 |
| ALOX15 | -2.66 | 1.42E-01 | 5.37 |
| ALOX15B | -5.28 | 8.53E-04 | 3.94 |
| ALPK2 | -5.32 | 5.83E-05 | 5.96 |
| ALPPL2 | -3.32 | 1.16E-01 | 3.70 |

| | | | |
|---|---|---|---|
| AMBN | -6.92 | 5.83E-05 | -0.97 |
| AMBP | -8.51 | 1.52E-07 | 3.97 |
| AMDHD1 | -3.54 | 1.88E-01 | 2.49 |
| AMN | -2.89 | 9.56E-02 | 4.86 |
| AMPD3 | 3.26 | 2.28E-02 | 6.44 |
| ANGPT1 | -3.05 | 5.23E-02 | 5.49 |
| ANGPTL3 | -5.92 | 1.34E-03 | -0.97 |
| ANKRD1 | -2.98 | 7.70E-02 | 5.11 |
| ANKRD18B | -2.41 | 1.78E-01 | 6.17 |
| ANKRD37 | 3.10 | 3.35E-02 | 6.60 |
| ANKRD66 | 5.47 | 8.75E-04 | 4.09 |
| ANO4 | 4.75 | 6.10E-04 | 5.54 |
| ANO9 | -3.73 | 1.19E-01 | 2.58 |
| ANPEP | -3.25 | 2.95E-02 | 5.43 |
| ANXA3 | -3.13 | 3.03E-02 | 6.32 |
| ANXA8 | -8.10 | 9.63E-08 | 4.77 |
| ANXA8L1 | -7.88 | 1.64E-06 | -0.97 |
| AP1M2 | -7.05 | 4.19E-05 | 3.24 |
| APELA | -6.39 | 3.05E-04 | 2.91 |
| APLNR | -2.77 | 7.49E-02 | 6.39 |
| APOA1 | -8.36 | 2.30E-10 | 8.90 |
| APOA1-AS | -5.39 | 2.46E-02 | 2.41 |
| APOA2 | -11.80 | 6.11E-13 | -0.97 |
| APOA4 | -11.47 | 2.16E-12 | 5.45 |
| APOB | -9.36 | 8.97E-12 | 8.92 |
| APOC2 | -5.73 | 2.35E-03 | -0.97 |
| APOC3 | -12.03 | 2.81E-13 | -0.97 |
| APOC4-APOC2 | -5.77 | 2.11E-03 | -0.97 |
| APOD | 2.99 | 8.30E-02 | 4.85 |
| APOH | -7.05 | 4.19E-05 | 3.24 |
| APOL6 | -2.89 | 1.46E-01 | 4.16 |
| AQP1 | -2.79 | 4.99E-02 | 11.03 |
| ARG1 | -4.89 | 2.55E-02 | -0.97 |
| ARHGAP22 | 4.02 | 2.20E-02 | 3.95 |
| ARHGEF39 | -2.67 | 1.60E-01 | 4.51 |
| ARID3C | -3.10 | 9.70E-02 | 3.85 |
| ARMC3 | 4.80 | 4.44E-04 | 5.92 |
| ARNTL2 | 2.43 | 1.61E-01 | 6.85 |
| ARSE | -2.46 | 1.56E-01 | 6.34 |
| ASCL1 | 3.09 | 2.25E-02 | 9.52 |
| ASCL2 | 4.79 | 5.44E-03 | -0.97 |
| ASGR2 | -6.52 | 2.01E-04 | -0.97 |
| ASIC4 | 2.87 | 7.20E-02 | 5.96 |
| ASMTL | -5.34 | 7.07E-04 | 3.97 |
| ASPA | 6.58 | 2.06E-05 | -0.97 |
| ASPG | -5.16 | 1.26E-02 | -0.97 |
| ATAD5 | -2.88 | 4.66E-02 | 7.33 |
| ATAT1 | 6.22 | 8.10E-04 | 3.46 |
| ATP10A | 5.69 | 4.24E-04 | 4.20 |
| ATP6V1C2 | 4.43 | 3.48E-03 | 4.89 |
| ATP7B | -2.48 | 1.14E-01 | 10.23 |
| ATP8A2 | 2.92 | 3.92E-02 | 8.74 |
| ATP8B1 | -3.85 | 1.28E-02 | 4.45 |
| AURKB | -2.45 | 1.47E-01 | 7.07 |
| AVPR1A | 3.29 | 2.52E-02 | 5.90 |
| B3GNT4 | 3.82 | 8.75E-02 | 3.26 |
| BAAT | 5.67 | 4.63E-03 | 3.19 |
| BAG6 | -3.56 | 1.08E-02 | 6.02 |
| BCAN | 5.02 | 5.01E-05 | 10.03 |

| | | | |
|---|---|---|---|
| BCAS1 | 3.61 | 8.72E-02 | -0.97 |
| BCO1 | 3.92 | 5.18E-03 | 5.78 |
| BDNF | 2.50 | 1.53E-01 | 6.19 |
| BEX5 | 2.32 | 1.95E-01 | 7.44 |
| BHLHE40-AS1 | 3.68 | 5.14E-02 | 3.78 |
| BHLHE41 | 4.89 | 1.05E-04 | 7.61 |
| BNC1 | -6.89 | 1.58E-07 | 6.62 |
| BNIPL | -3.30 | 1.95E-01 | 2.95 |
| BRINP3 | -3.21 | 3.26E-02 | 5.32 |
| BSX | 5.41 | 1.01E-02 | 3.06 |
| C10orf105 | 2.72 | 1.34E-01 | 5.41 |
| C10orf90 | 6.16 | 7.72E-05 | -0.97 |
| C11orf88 | 5.54 | 9.15E-05 | 5.45 |
| C11orf97 | 4.19 | 3.64E-02 | 3.45 |
| C1GALT1C1L | -3.96 | 1.95E-01 | -0.97 |
| C1orf158 | 6.03 | 1.50E-03 | 3.37 |
| C1orf189 | 5.49 | 7.75E-03 | 3.10 |
| C1orf194 | 3.17 | 5.98E-02 | 5.11 |
| C1orf210 | -3.58 | 1.07E-01 | 3.09 |
| C1orf61 | -9.43 | 4.96E-09 | 4.43 |
| C1orf87 | 4.22 | 2.19E-02 | -0.97 |
| C1QL4 | 3.33 | 2.45E-02 | 5.83 |
| C1QTNF7 | -4.22 | 4.81E-03 | 4.41 |
| C20orf85 | 5.85 | 2.56E-04 | 4.28 |
| C22orf15 | 4.12 | 4.27E-02 | 3.41 |
| C22orf23 | 3.23 | 9.68E-02 | 4.29 |
| C2orf48 | -3.64 | 9.50E-02 | 3.12 |
| C2orf50 | 3.78 | 9.29E-03 | 5.57 |
| C2orf71 | 4.48 | 6.33E-03 | 4.18 |
| C2orf72 | -3.18 | 2.41E-02 | 6.80 |
| C2orf73 | 3.63 | 6.01E-02 | 3.76 |
| C2orf81 | 2.36 | 1.95E-01 | 6.74 |
| C2orf88 | 3.46 | 1.55E-02 | 5.99 |
| C3 | -6.53 | 1.28E-06 | 5.98 |
| C3orf35 | -2.92 | 9.07E-02 | 4.98 |
| C3orf80 | 4.41 | 1.77E-03 | 5.37 |
| C3orf85 | -4.64 | 4.99E-02 | -0.97 |
| C4orf47 | 2.52 | 1.56E-01 | 6.26 |
| C4orf54 | 3.61 | 8.72E-02 | -0.97 |
| C5AR1 | 3.11 | 4.51E-02 | 5.49 |
| C5orf49 | 3.83 | 5.29E-03 | 6.18 |
| C5orf63 | -4.64 | 4.99E-02 | -0.97 |
| C5orf66-AS1 | -6.66 | 1.31E-04 | -0.97 |
| C6orf141 | -3.01 | 1.26E-01 | 3.80 |
| C7 | 3.29 | 3.91E-02 | 5.00 |
| C7orf57 | 2.91 | 7.51E-02 | 5.62 |
| C8B | -4.22 | 1.29E-01 | -0.97 |
| C8orf34 | 4.57 | 2.23E-03 | 4.64 |
| C9orf135 | 3.46 | 2.43E-02 | 5.08 |
| C9orf24 | 5.02 | 3.57E-04 | 5.45 |
| CA10 | 4.13 | 1.23E-03 | 7.59 |
| CA12 | -2.63 | 9.14E-02 | 7.10 |
| CA2 | 3.49 | 7.33E-03 | 9.83 |
| CABLES1 | 3.30 | 1.49E-02 | 7.47 |
| CACFD1 | 4.89 | 4.03E-02 | 2.80 |
| CACNA1S | 3.68 | 8.72E-02 | -0.97 |
| CALB1 | 3.40 | 1.25E-02 | 7.18 |
| CALB2 | 6.14 | 8.02E-05 | -0.97 |
| CALHM5 | 3.45 | 9.70E-02 | 3.66 |

| | | | |
|---|---|---|---|
| CAPN3 | 3.57 | 7.22E-03 | 7.69 |
| CAPSL | 5.90 | 6.51E-05 | 4.89 |
| CARD11 | -2.70 | 1.23E-01 | 5.15 |
| CARD16 | -4.45 | 7.16E-02 | -0.97 |
| CARNS1 | -3.20 | 2.85E-02 | 5.78 |
| CASC1 | 3.29 | 2.23E-02 | 6.39 |
| CASC2 | 2.86 | 8.12E-02 | 5.87 |
| CASP10 | -4.34 | 8.72E-02 | -0.97 |
| CAT | -4.96 | 2.19E-02 | -0.97 |
| CATIP | 2.98 | 1.65E-01 | 3.84 |
| CAVIN1 | -3.04 | 4.09E-02 | 5.93 |
| CAVIN2 | -3.74 | 1.75E-02 | 4.39 |
| CBLN1 | 5.49 | 9.05E-06 | 10.37 |
| CCDC103 | 2.91 | 6.85E-02 | 5.72 |
| CCDC114 | 2.95 | 8.72E-02 | 5.15 |
| CCDC140 | 3.95 | 1.43E-02 | 4.65 |
| CCDC157 | 2.65 | 1.76E-01 | 5.00 |
| CCDC169 | -3.64 | 9.50E-02 | 3.12 |
| CCDC169-SOHLI | -3.78 | 3.59E-02 | 3.93 |
| CCDC170 | 4.51 | 7.74E-04 | 6.07 |
| CCDC194 | -3.45 | 1.34E-01 | 3.02 |
| CCDC39 | 4.46 | 2.07E-03 | 5.17 |
| CCDC60 | 4.75 | 6.00E-04 | 5.54 |
| CCDC65 | 3.21 | 3.47E-02 | 5.77 |
| CCDC68 | -3.06 | 4.95E-02 | 5.50 |
| CCDC78 | 2.75 | 1.07E-01 | 5.82 |
| CCDC81 | 2.47 | 1.99E-01 | 5.49 |
| CCDC96 | 3.67 | 9.75E-03 | 5.77 |
| CCNO | 5.34 | 5.95E-05 | 6.35 |
| CD244 | -4.73 | 1.19E-01 | 2.08 |
| CD300E | -6.59 | 1.57E-04 | 3.01 |
| CD34 | -5.19 | 1.46E-04 | 5.63 |
| CD44 | -3.40 | 1.14E-02 | 7.25 |
| CD84 | -5.34 | 7.75E-03 | -0.97 |
| CD93 | -7.06 | 1.64E-06 | 4.83 |
| CDA | -4.34 | 8.72E-02 | -0.97 |
| CDC20B | 4.91 | 7.49E-05 | 9.04 |
| CDC42EP5 | 6.33 | 5.88E-04 | 3.52 |
| CDC45 | -3.54 | 3.12E-02 | 4.07 |
| CDCA7 | -2.61 | 9.25E-02 | 7.63 |
| CDCP1 | -6.04 | 6.98E-05 | 4.32 |
| CDH13 | -2.86 | 8.44E-02 | 5.32 |
| CDH18 | 4.77 | 2.07E-04 | 7.13 |
| CDH5 | -8.21 | 4.58E-07 | 3.82 |
| CDHR3 | 2.78 | 8.72E-02 | 5.99 |
| CDK15 | 4.15 | 4.20E-03 | 5.23 |
| CDK7 | 3.89 | 7.61E-03 | 5.30 |
| CDKN2B | 3.30 | 3.02E-02 | 5.47 |
| CDKN2C | -3.00 | 1.11E-01 | 4.21 |
| CDX2 | -7.34 | 1.37E-05 | -0.97 |
| CEACAM1 | 3.27 | 1.50E-01 | 3.57 |
| CEACAM19 | -2.66 | 1.60E-01 | 4.86 |
| CERKL | 2.96 | 5.85E-02 | 6.00 |
| CES1 | -5.22 | 1.11E-02 | -0.97 |
| CFAP126 | 3.97 | 2.25E-03 | 7.25 |
| CFAP157 | 3.73 | 1.38E-02 | 5.03 |
| CFAP161 | 3.89 | 3.12E-02 | 3.88 |
| CFAP206 | 2.90 | 5.21E-02 | 6.61 |
| CFAP299 | 4.85 | 4.19E-03 | -0.97 |

| | | | |
|---|---|---|---|
| CFAP43 | 3.14 | 2.41E-02 | 7.48 |
| CFAP45 | 3.79 | 4.73E-03 | 6.57 |
| CFAP47 | 3.03 | 5.10E-02 | 5.96 |
| CFAP52 | 4.27 | 1.44E-03 | 6.30 |
| CFAP54 | 3.03 | 4.80E-02 | 5.87 |
| CFAP57 | 3.35 | 2.67E-02 | 5.49 |
| CFAP61 | 4.59 | 7.14E-04 | 5.97 |
| CFAP65 | 5.67 | 3.85E-04 | -0.97 |
| CFAP70 | 2.69 | 8.18E-02 | 7.02 |
| CFAP73 | 3.61 | 3.78E-02 | 4.16 |
| CFAP74 | 3.74 | 7.87E-03 | 5.81 |
| CFAP77 | 2.49 | 1.92E-01 | 5.69 |
| CFAP97D2 | 3.53 | 1.07E-01 | -0.97 |
| CFI | 2.48 | 1.60E-01 | 6.45 |
| CFL1P1 | 3.36 | 1.60E-01 | -0.97 |
| CHCHD10 | 5.66 | 4.93E-04 | 4.18 |
| CHL1 | 2.99 | 6.35E-02 | 5.43 |
| CHMP4A | 5.61 | 5.83E-03 | 3.16 |
| CHRNB1 | 3.53 | 1.07E-01 | -0.97 |
| CHRND | 2.84 | 1.78E-01 | 4.36 |
| CHST9 | 2.37 | 1.59E-01 | 8.14 |
| CKM | 3.13 | 1.95E-01 | 3.51 |
| CLCF1 | -3.19 | 7.22E-02 | 4.12 |
| CLDN1 | 3.48 | 8.52E-03 | 8.50 |
| CLDN11 | 2.34 | 1.95E-01 | 6.92 |
| CLDN16 | 3.27 | 1.95E-01 | -0.97 |
| CLDN7 | -3.57 | 1.04E-02 | 6.02 |
| CLEC12A | -4.45 | 7.16E-02 | -0.97 |
| CLEC14A | -3.89 | 8.86E-02 | 2.66 |
| CLEC1B | -4.22 | 1.29E-01 | -0.97 |
| CLEC4A | -3.41 | 4.45E-02 | 4.01 |
| CLIC3 | -5.45 | 2.19E-02 | 2.44 |
| CLIC6 | 5.53 | 2.23E-05 | 6.93 |
| CLRN3 | -5.54 | 4.19E-03 | -0.97 |
| CLUL1 | 3.47 | 1.01E-02 | 7.09 |
| CNKSR1 | -2.79 | 1.20E-01 | 4.57 |
| CNPY1 | 4.24 | 1.10E-03 | 6.80 |
| CNTNAP2 | 2.51 | 1.32E-01 | 7.00 |
| COBLL1 | -3.42 | 9.16E-03 | 9.06 |
| COL11A2 | 5.57 | 6.27E-03 | 3.14 |
| COL12A1 | -2.51 | 1.22E-01 | 7.28 |
| COL13A1 | -3.50 | 9.82E-03 | 6.96 |
| COL14A1 | -5.45 | 2.92E-05 | 6.53 |
| COL16A1 | -2.36 | 1.85E-01 | 6.59 |
| COL1A1 | -3.38 | 9.57E-03 | 13.18 |
| COL21A1 | -5.32 | 4.59E-05 | 6.62 |
| COL3A1 | -3.75 | 2.84E-03 | 15.25 |
| COL5A1 | -2.87 | 4.03E-02 | 12.57 |
| COL6A3 | -2.90 | 3.77E-02 | 10.74 |
| COL8A2 | -2.34 | 1.70E-01 | 8.47 |
| CORIN | 3.10 | 5.03E-02 | 5.61 |
| CORO7 | 4.82 | 4.45E-02 | 2.76 |
| CORO7-PAM16 | 4.79 | 4.95E-02 | 2.75 |
| COX6B2 | -3.84 | 4.46E-03 | 6.03 |
| CPEB1 | 3.57 | 1.60E-01 | 3.14 |
| CPLX3 | 4.48 | 1.45E-03 | 5.40 |
| CPNE9 | 3.32 | 7.80E-02 | 4.01 |
| CPSF1 | 3.42 | 2.96E-02 | 5.23 |
| CPVL | 2.33 | 1.74E-01 | 8.27 |

| | | | |
|---|---|---|---|
| CPZ | -2.85 | 4.84E-02 | 7.92 |
| CRB3 | -3.89 | 3.34E-02 | 3.66 |
| CREB3L3 | -3.50 | 8.75E-02 | 3.46 |
| CREG2 | -2.54 | 1.28E-01 | 6.16 |
| CRHBP | -4.62 | 1.02E-03 | 5.19 |
| CRIP1 | 4.21 | 1.39E-03 | 6.46 |
| CRISPLD2 | -4.16 | 8.97E-04 | 8.75 |
| CROCC2 | 5.03 | 1.08E-03 | 4.46 |
| CRTAC1 | -2.61 | 1.05E-01 | 6.57 |
| CRX | 5.19 | 3.05E-04 | 5.27 |
| CRYAA2 | 3.27 | 1.95E-01 | -0.97 |
| CRYAB | 3.57 | 6.48E-03 | 8.29 |
| CRYBG1 | -3.73 | 1.19E-01 | 2.58 |
| CRYZ | -4.45 | 7.16E-02 | -0.97 |
| CSF2RB | -4.54 | 1.88E-01 | 1.99 |
| CSF3R | -4.51 | 1.88E-03 | 4.56 |
| CST1 | -6.73 | 1.05E-04 | -0.97 |
| CTSF | -5.34 | 2.78E-02 | 2.39 |
| CTSV | -2.49 | 1.23E-01 | 7.82 |
| CXCL1 | -3.96 | 1.95E-01 | -0.97 |
| CXCL13 | 6.86 | 6.97E-06 | -0.97 |
| CXCR4 | 2.29 | 1.85E-01 | 9.23 |
| CXorf36 | -6.57 | 1.79E-04 | -0.97 |
| CYP1A1 | -3.65 | 1.26E-02 | 5.00 |
| CYP26A1 | -2.43 | 1.77E-01 | 5.88 |
| CYP27B1 | 3.34 | 4.16E-02 | 4.83 |
| CYP2A6 | -4.39 | 2.46E-02 | 2.91 |
| CYP2B7P | -5.34 | 7.75E-03 | -0.97 |
| CYP2W1 | -5.16 | 4.54E-02 | 2.30 |
| DAPP1 | -5.10 | 1.43E-02 | -0.97 |
| DAW1 | 4.61 | 1.22E-02 | 3.66 |
| DCN | -3.26 | 1.32E-02 | 11.16 |
| DCT | 8.56 | 7.27E-11 | 11.63 |
| DDAH2 | 7.44 | 1.45E-05 | 4.08 |
| DDN | 3.60 | 1.26E-02 | 5.86 |
| DDR1 | -2.58 | 1.95E-01 | 4.59 |
| DDX3Y | -5.81 | 1.94E-03 | -0.97 |
| DGAT1 | 5.43 | 8.10E-04 | -0.97 |
| DGKG | 3.80 | 2.19E-02 | 4.26 |
| DHRS2 | 3.62 | 1.92E-02 | 4.97 |
| DHRSX | 3.89 | 1.75E-02 | 4.30 |
| DIAPH3 | -2.71 | 9.68E-02 | 5.77 |
| DIO1 | -5.34 | 7.75E-03 | -0.97 |
| DIO2 | 2.56 | 1.41E-01 | 6.28 |
| DIO3 | 2.41 | 1.60E-01 | 7.23 |
| DKK1 | -3.22 | 6.01E-02 | 4.65 |
| DKK2 | -4.51 | 4.51E-04 | 6.67 |
| DLEC1 | 3.26 | 2.53E-02 | 5.98 |
| DLK1 | 2.44 | 1.25E-01 | 13.61 |
| DLX5 | -6.06 | 8.75E-04 | 2.75 |
| DLX6 | -3.58 | 1.07E-01 | 3.09 |
| DLX6-AS1 | -5.96 | 1.23E-03 | 2.70 |
| DNAAF1 | 3.73 | 1.07E-02 | 5.54 |
| DNAAF3 | 2.96 | 4.66E-02 | 6.59 |
| DNAH11 | 4.39 | 1.89E-03 | 5.36 |
| DNAH12 | 5.43 | 9.57E-03 | 3.07 |
| DNAH17 | -4.10 | 1.60E-01 | -0.97 |
| DNAH3 | 4.65 | 7.75E-03 | -0.97 |
| DNAH5 | 2.92 | 8.13E-02 | 5.28 |

| | | | |
|---|---|---|---|
| DNAH7 | 3.00 | 4.00E-02 | 7.06 |
| DNAH9 | 4.02 | 2.55E-03 | 6.54 |
| DNAI2 | 5.08 | 2.92E-03 | 3.89 |
| DNM3OS | -5.16 | 9.83E-05 | 5.76 |
| DOCK8 | 2.49 | 1.78E-01 | 5.99 |
| DPPA2 | -4.73 | 1.19E-01 | 2.08 |
| DPYS | -4.34 | 8.72E-02 | -0.97 |
| DRC1 | 3.04 | 2.96E-02 | 8.02 |
| DRC3 | 3.30 | 3.57E-02 | 5.32 |
| DRC7 | 4.16 | 8.16E-03 | 4.43 |
| DRGX | 2.49 | 1.68E-01 | 6.06 |
| DSC3 | -6.11 | 1.30E-06 | 7.77 |
| DSG4 | -4.34 | 8.72E-02 | -0.97 |
| DTHD1 | 3.49 | 1.95E-01 | 3.10 |
| DUSP13 | 3.17 | 1.95E-01 | -0.97 |
| DYNC1I1 | 2.39 | 1.67E-01 | 7.22 |
| DYNLRB2 | 3.42 | 3.35E-02 | 4.87 |
| E2F8 | -2.60 | 1.89E-01 | 4.48 |
| EBLN2 | -2.90 | 1.33E-01 | 4.49 |
| ECEL1 | 3.39 | 9.58E-03 | 10.24 |
| ECEL1P1 | -3.07 | 8.69E-02 | 4.57 |
| ECSCR | -6.22 | 5.52E-04 | -0.97 |
| EDAR | 4.88 | 5.57E-04 | 5.38 |
| EFCAB1 | 3.72 | 4.94E-03 | 7.30 |
| EFCAB12 | 4.16 | 8.17E-03 | 4.76 |
| EFHC2 | 3.15 | 2.47E-02 | 7.06 |
| EGFLAM | -4.50 | 8.75E-04 | 5.67 |
| EGLN3 | 2.77 | 6.82E-02 | 7.35 |
| ELF3 | -3.17 | 7.17E-02 | 4.47 |
| ELMOD1 | -2.65 | 1.14E-01 | 5.80 |
| EMCN | -5.89 | 1.48E-03 | 2.66 |
| EMP1 | 5.82 | 2.80E-06 | 8.96 |
| EMX1 | -6.94 | 5.35E-05 | -0.97 |
| EN1 | 5.86 | 9.62E-06 | 6.60 |
| EN2 | 5.08 | 5.35E-05 | 8.25 |
| ENKUR | 2.75 | 7.21E-02 | 6.90 |
| ENO3 | 2.33 | 1.74E-01 | 8.29 |
| ENO4 | 5.01 | 5.52E-04 | 5.18 |
| ENPP1 | -3.22 | 2.10E-02 | 6.75 |
| EOMES | -2.90 | 4.88E-02 | 6.69 |
| EPCAM | -3.43 | 9.44E-03 | 8.24 |
| EPHA5 | -2.88 | 4.99E-02 | 7.30 |
| EPHA6 | 2.67 | 1.16E-01 | 5.94 |
| EPSTI1 | -3.17 | 4.75E-02 | 5.00 |
| ERFE | 2.83 | 8.52E-02 | 5.68 |
| ERN2 | 4.17 | 1.95E-01 | 2.44 |
| ERVH48-1 | -8.38 | 3.14E-08 | 4.91 |
| ESAM | -3.85 | 1.19E-02 | 4.64 |
| ESM1 | 2.31 | 1.85E-01 | 8.09 |
| ESPL1 | -3.38 | 1.73E-02 | 5.86 |
| ESRG | -6.73 | 1.76E-05 | 4.08 |
| ESRP1 | -3.38 | 1.26E-02 | 6.93 |
| ETV1 | 2.30 | 1.79E-01 | 9.73 |
| ETV3L | 4.61 | 8.71E-03 | -0.97 |
| EVA1A | 2.58 | 1.22E-01 | 6.69 |
| EXTL1 | 3.08 | 3.75E-02 | 6.22 |
| F11R | -2.24 | 2.00E-01 | 9.55 |
| F2 | -6.94 | 5.35E-05 | -0.97 |
| FABP1 | -10.08 | 3.76E-10 | -0.97 |

| | | | |
|---|---|---|---|
| FABP6 | 2.63 | 1.91E-01 | 4.99 |
| FADS6 | -3.64 | 1.60E-01 | 2.53 |
| FAM107A | -2.93 | 4.56E-02 | 6.99 |
| FAM110C | 3.34 | 2.15E-02 | 5.93 |
| FAM124B | -3.70 | 2.10E-02 | 4.15 |
| FAM157B | -3.16 | 1.95E-01 | 3.30 |
| FAM160A1-DT | 3.49 | 1.95E-01 | 3.10 |
| FAM163B | -3.12 | 9.04E-02 | 4.08 |
| FAM166B | 4.38 | 2.73E-03 | 5.13 |
| FAM181B | 3.01 | 3.13E-02 | 8.40 |
| FAM183A | 4.14 | 3.32E-03 | 5.42 |
| FAM184B | 4.74 | 8.48E-03 | 3.72 |
| FAM187A | 3.10 | 9.64E-02 | 4.49 |
| FAM213B | 3.34 | 7.22E-02 | 4.02 |
| FAM216B | 5.64 | 2.86E-05 | 6.34 |
| FAM221B | 4.68 | 6.80E-02 | 2.70 |
| FAM238C | -3.36 | 9.70E-02 | 3.72 |
| FAM72A | -3.10 | 9.70E-02 | 3.85 |
| FAM72B | -2.85 | 1.08E-01 | 4.60 |
| FAM81B | 4.34 | 1.56E-03 | 5.85 |
| FAM83B | -6.70 | 1.11E-04 | -0.97 |
| FAM92B | 3.74 | 1.77E-02 | 4.81 |
| FANK1 | 2.85 | 6.13E-02 | 6.69 |
| FBL | -5.52 | 6.16E-05 | 5.29 |
| FBXO17 | -6.39 | 3.05E-04 | -0.97 |
| FBXO4 | -2.81 | 1.60E-01 | 4.44 |
| FCRLA | 5.99 | 1.31E-04 | -0.97 |
| FENDRR | -5.77 | 9.18E-03 | 2.60 |
| FERMT1 | -3.58 | 2.43E-02 | 4.67 |
| FERMT3 | -3.05 | 6.54E-02 | 4.83 |
| FETUB | -6.79 | 8.39E-05 | -0.97 |
| FFAR4 | 6.29 | 6.73E-04 | 3.50 |
| FGA | -11.91 | 4.41E-13 | -0.97 |
| FGB | -12.51 | 6.85E-14 | -0.97 |
| FGF10 | -3.20 | 6.36E-02 | 4.64 |
| FGF12 | 2.40 | 1.58E-01 | 7.53 |
| FGF17 | 3.01 | 7.47E-02 | 5.18 |
| FGF19 | -4.03 | 5.93E-02 | 2.73 |
| FGFBP3 | -3.76 | 3.00E-03 | 9.40 |
| FGFR3 | -3.11 | 2.10E-02 | 10.73 |
| FGFR4 | -2.50 | 1.47E-01 | 6.25 |
| FGG | -11.53 | 1.93E-12 | -0.97 |
| FIBIN | -3.61 | 8.40E-03 | 6.38 |
| FKBP11 | -2.99 | 4.13E-02 | 6.42 |
| FLI1 | -2.38 | 1.80E-01 | 6.49 |
| FLJ12825 | -5.34 | 2.78E-02 | 2.39 |
| FLOT1 | -2.96 | 1.16E-01 | 4.37 |
| FLT1 | -8.81 | 6.66E-09 | 5.12 |
| FMN1 | 4.68 | 1.56E-03 | 5.02 |
| FMO1 | -4.30 | 3.09E-03 | 4.86 |
| FMO5 | -3.22 | 7.56E-02 | 3.91 |
| FMOD | -3.16 | 2.09E-02 | 7.85 |
| FN1 | -2.25 | 1.95E-01 | 12.99 |
| FNDC10 | 2.36 | 1.88E-01 | 7.03 |
| FOLR1 | 4.06 | 4.89E-03 | 5.55 |
| FOXA1 | -5.92 | 1.34E-03 | 2.68 |
| FOXA2 | -5.89 | 1.48E-03 | -0.97 |
| FOXD1 | -3.71 | 1.85E-02 | 4.38 |
| FOXD2 | -4.03 | 5.93E-02 | 2.73 |

| | | | |
|---|---|---|---|
| FOXF1 | -3.06 | 4.01E-02 | 6.10 |
| FOXG1 | -10.54 | 7.27E-11 | 4.99 |
| FOXH1 | -2.74 | 1.40E-01 | 4.89 |
| FOXJ1 | 3.85 | 2.27E-03 | 9.32 |
| FOXL2NB | -4.45 | 7.16E-02 | -0.97 |
| FREM1 | -3.84 | 2.80E-03 | 7.59 |
| FRG2DP | 3.45 | 1.29E-01 | -0.97 |
| FRMD3 | 3.10 | 3.72E-02 | 6.15 |
| FRMD7 | 3.68 | 5.14E-02 | 3.78 |
| FRZB | -3.85 | 2.13E-03 | 10.37 |
| FSHR | -4.45 | 7.16E-02 | -0.97 |
| FST | -3.93 | 2.97E-03 | 6.49 |
| FTCD | -4.37 | 1.36E-02 | 3.49 |
| FUT9 | -3.09 | 2.53E-02 | 7.53 |
| FZD10 | 2.48 | 1.18E-01 | 9.29 |
| G0S2 | -5.50 | 1.96E-05 | 7.11 |
| GABRG1 | -4.73 | 3.57E-02 | -0.97 |
| GABRG3 | -2.67 | 1.60E-01 | 4.75 |
| GABRP | -5.65 | 4.43E-05 | 5.35 |
| GAL | 3.33 | 2.85E-02 | 5.48 |
| GAPLINC | 5.09 | 2.11E-03 | -0.97 |
| GAS6-AS2 | 4.50 | 5.71E-03 | 4.19 |
| GATA3 | -2.46 | 1.58E-01 | 6.11 |
| GATA4 | -4.10 | 1.60E-01 | -0.97 |
| GATA5 | -4.67 | 4.97E-04 | 5.96 |
| GATA6 | -8.61 | 4.96E-09 | 5.61 |
| GATA6-AS1 | -8.68 | 8.94E-08 | -0.97 |
| GBA3 | -5.85 | 7.75E-03 | 2.64 |
| GBP2 | -2.82 | 1.14E-01 | 4.59 |
| GBP3 | -4.81 | 3.02E-02 | -0.97 |
| GBP4 | -4.58 | 7.75E-03 | 3.59 |
| GCK | 2.66 | 1.57E-01 | 5.14 |
| GCSAML | -4.10 | 1.60E-01 | -0.97 |
| GCSAML-AS1 | 5.03 | 2.66E-03 | -0.97 |
| GDA | 3.57 | 1.60E-01 | 3.14 |
| GDF5 | 4.67 | 1.65E-03 | 5.01 |
| GFI1B | -4.81 | 3.02E-02 | -0.97 |
| GIMAP1 | -3.96 | 1.95E-01 | -0.97 |
| GIMAP4 | -5.16 | 1.26E-02 | -0.97 |
| GIMAP6 | -4.54 | 6.01E-02 | -0.97 |
| GJA4 | -5.92 | 2.41E-04 | 3.68 |
| GJA5 | -3.08 | 6.22E-02 | 4.84 |
| GJB1 | -4.54 | 5.64E-03 | 3.99 |
| GLI1 | -6.25 | 8.39E-05 | 3.84 |
| GLIS3 | 3.05 | 3.94E-02 | 6.40 |
| GLYAT | -4.10 | 1.60E-01 | -0.97 |
| GMNC | 5.57 | 4.36E-05 | 5.95 |
| GNG13 | 3.95 | 4.20E-02 | -0.97 |
| GNG8 | 2.66 | 1.57E-01 | 5.14 |
| GNGT1 | 5.29 | 1.54E-03 | 4.00 |
| GNL1 | 3.82 | 3.89E-02 | 3.85 |
| GNRH2 | -7.39 | 1.12E-05 | 3.41 |
| GOLGA2P10 | -3.40 | 4.03E-02 | 4.59 |
| GOLGA6A | 4.14 | 3.98E-02 | 3.43 |
| GOLGA6D | 3.89 | 4.99E-02 | -0.97 |
| GOLGA6L10 | -4.89 | 2.55E-02 | -0.97 |
| GP1BA | -5.16 | 4.54E-02 | 2.30 |
| GP9 | -4.34 | 8.72E-02 | -0.97 |
| GPC5-AS1 | 5.36 | 9.52E-04 | -0.97 |

| | | | |
|---|---|---|---|
| GPD1 | -3.28 | 3.58E-02 | 5.06 |
| GPNMB | 6.61 | 1.90E-07 | 8.36 |
| GPR101 | -5.39 | 2.46E-02 | 2.41 |
| GPR135 | 2.48 | 1.83E-01 | 5.59 |
| GPR158 | 5.12 | 7.80E-05 | 6.82 |
| GPR85 | 2.55 | 1.12E-01 | 7.74 |
| GPRC5C | -2.26 | 1.95E-01 | 9.11 |
| GRB7 | -4.69 | 9.57E-04 | 4.87 |
| GRHL3 | -3.05 | 1.16E-01 | 3.83 |
| GRIA2 | -2.31 | 1.76E-01 | 8.58 |
| GRIK1 | 4.48 | 5.23E-04 | 6.91 |
| GRPR | -5.08 | 2.64E-04 | 5.06 |
| GSDMB | -2.86 | 4.82E-02 | 7.50 |
| GSG1L | 3.04 | 5.48E-02 | 5.68 |
| GSTA1 | -6.65 | 1.08E-06 | 5.62 |
| GSTA2 | -9.29 | 9.31E-09 | -0.97 |
| GSTA3 | 5.34 | 1.03E-03 | -0.97 |
| GSX2 | -4.64 | 4.99E-02 | -0.97 |
| GUCY1A1 | -3.39 | 1.29E-02 | 6.90 |
| GUCY2C | -5.39 | 1.32E-03 | 3.41 |
| GUCY2F | 5.67 | 3.85E-04 | -0.97 |
| GZMK | 3.36 | 1.60E-01 | -0.97 |
| H19 | -7.33 | 1.44E-08 | 8.29 |
| HAL | -4.22 | 1.29E-01 | -0.97 |
| HAND1 | -9.10 | 7.67E-11 | 6.85 |
| HAND2 | -9.35 | 7.19E-10 | 5.39 |
| HAND2-AS1 | -9.71 | 1.63E-09 | 4.57 |
| HAPLN1 | -4.71 | 1.79E-04 | 7.62 |
| HAVCR1 | -6.42 | 2.84E-04 | 2.93 |
| HBE1 | -7.90 | 1.59E-06 | 3.67 |
| HBG1 | -5.22 | 1.11E-02 | -0.97 |
| HBG2 | -6.83 | 7.72E-05 | -0.97 |
| HDC | 3.90 | 8.92E-03 | 5.11 |
| HEBP1 | 2.74 | 5.88E-02 | 10.00 |
| HEY2 | -3.05 | 3.38E-02 | 6.67 |
| HGF | -3.00 | 3.64E-02 | 6.95 |
| HHAT | 4.01 | 4.20E-02 | -0.97 |
| HHIP | -6.68 | 1.17E-04 | -0.97 |
| HHIP-AS1 | -4.18 | 2.26E-02 | 3.39 |
| HJURP | -3.02 | 3.28E-02 | 7.31 |
| HKDC1 | 3.41 | 3.51E-02 | 4.87 |
| HLA-A | 2.27 | 1.88E-01 | 9.85 |
| HLA-E | 3.43 | 2.85E-02 | 5.24 |
| HLX | -4.14 | 6.33E-03 | 4.37 |
| HMX1 | -5.68 | 2.98E-03 | -0.97 |
| HNF1A | -4.96 | 7.68E-02 | 2.20 |
| HNF4A | -6.50 | 4.19E-05 | 3.96 |
| HNRNPA1P8 | -3.58 | 1.07E-01 | 3.09 |
| HOPX | -4.58 | 1.54E-03 | 4.59 |
| HORMAD2 | 4.72 | 6.07E-02 | 2.71 |
| HOXA10 | -6.45 | 2.63E-04 | -0.97 |
| HOXA10-HOXA9 | -6.96 | 5.23E-05 | -0.97 |
| HOXA11 | -6.31 | 4.10E-04 | -0.97 |
| HOXA13 | -6.19 | 5.90E-04 | -0.97 |
| HOXA2 | -5.59 | 3.72E-03 | -0.97 |
| HOXA3 | -5.85 | 1.77E-03 | -0.97 |
| HOXA4 | -4.10 | 1.60E-01 | -0.97 |
| HOXA5 | -4.81 | 3.02E-02 | -0.97 |
| HOXA7 | -4.54 | 1.88E-01 | 1.99 |

| | | | |
|---|---|---|---|
| HOXA9 | -7.05 | 4.19E-05 | -0.97 |
| HOXB-AS1 | -4.26 | 1.75E-02 | 3.43 |
| HOXB-AS3 | -4.73 | 3.57E-02 | -0.97 |
| HOXB1 | -4.45 | 7.16E-02 | -0.97 |
| HOXB2 | -5.89 | 1.83E-05 | 5.47 |
| HOXB3 | -8.43 | 2.57E-08 | 4.93 |
| HOXB4 | -8.33 | 2.94E-07 | -0.97 |
| HOXB5 | -4.48 | 1.99E-03 | 4.54 |
| HOXB6 | -7.52 | 6.97E-06 | -0.97 |
| HOXB7 | -5.10 | 1.50E-03 | 3.85 |
| HOXB8 | -3.18 | 2.71E-02 | 6.00 |
| HOXB9 | -7.67 | 3.55E-06 | 3.55 |
| HOXC10 | -5.50 | 5.44E-03 | -0.97 |
| HOXC4 | -6.14 | 5.35E-05 | 4.37 |
| HOXC5 | -6.77 | 8.75E-05 | -0.97 |
| HOXC6 | -8.01 | 1.03E-06 | 3.72 |
| HOXC8 | -8.39 | 2.28E-07 | -0.97 |
| HOXC9 | -5.34 | 7.75E-03 | -0.97 |
| HOXD3 | -4.10 | 1.60E-01 | -0.97 |
| HOXD4 | -6.59 | 1.57E-04 | -0.97 |
| HP | -4.10 | 1.60E-01 | -0.97 |
| HPN | -3.03 | 5.56E-02 | 5.55 |
| HPSE2 | -6.47 | 4.58E-05 | 3.95 |
| HPX | -3.75 | 1.43E-02 | 4.76 |
| HSPB6 | -4.54 | 1.88E-01 | 1.99 |
| HSPB7 | -2.73 | 1.41E-01 | 4.54 |
| HTR1E | 2.54 | 1.57E-01 | 5.87 |
| HTR2A | 4.24 | 3.09E-02 | 3.48 |
| HTR2C | 6.65 | 1.40E-07 | 9.04 |
| HTR7 | -3.89 | 3.34E-02 | 3.66 |
| HTR7P1 | 5.30 | 6.55E-05 | 6.33 |
| HYDIN | 3.41 | 1.79E-02 | 5.97 |
| ICAM1 | 2.50 | 1.23E-01 | 7.81 |
| ICAM2 | -5.73 | 1.01E-02 | 2.58 |
| IFI27L1 | 4.61 | 7.49E-02 | 2.66 |
| IFIH1 | -3.64 | 1.28E-02 | 5.24 |
| IGF2 | -4.71 | 1.18E-04 | 13.45 |
| IGF2-AS | -2.69 | 1.12E-01 | 5.52 |
| IGFBP3 | -4.10 | 9.73E-04 | 10.13 |
| IGFBP5 | 3.44 | 7.91E-03 | 13.28 |
| IGFBP7 | 3.39 | 1.11E-02 | 7.93 |
| IGFBP7-AS1 | 4.41 | 1.20E-01 | 2.56 |
| IGSF11 | 2.76 | 1.04E-01 | 5.82 |
| IHH | -4.17 | 1.17E-02 | 4.12 |
| IL11 | -2.93 | 5.92E-02 | 5.57 |
| IL12A-AS1 | 3.27 | 1.95E-01 | -0.97 |
| IL13RA2 | 2.89 | 1.60E-01 | 4.38 |
| IL16 | 3.09 | 5.34E-02 | 5.60 |
| IL17RD | 2.88 | 3.96E-02 | 10.97 |
| IL2RB | -2.89 | 1.38E-01 | 4.33 |
| IL33 | -4.96 | 2.19E-02 | -0.97 |
| IL5RA | 5.57 | 5.10E-04 | -0.97 |
| IL6R | -4.68 | 2.09E-04 | 7.22 |
| INS-IGF2 | -4.69 | 1.28E-04 | 13.40 |
| INSM2 | 3.62 | 1.10E-02 | 5.75 |
| INTS3 | 2.68 | 1.95E-01 | 4.50 |
| IPO4 | 4.72 | 7.00E-03 | -0.97 |
| IQCD | 2.80 | 1.00E-01 | 5.34 |
| IQCG | 2.66 | 8.93E-02 | 7.41 |

| | | | |
|---|---|---|---|
| IQCH | 3.57 | 9.57E-03 | 6.46 |
| IQGAP3 | -2.60 | 1.03E-01 | 6.85 |
| IRF4 | 3.64 | 9.48E-03 | 6.08 |
| IRF5 | 3.27 | 1.95E-01 | -0.97 |
| IRF6 | -2.84 | 7.83E-02 | 5.78 |
| IRS4 | 2.99 | 3.02E-02 | 9.49 |
| IRX3 | -2.95 | 4.13E-02 | 6.86 |
| ISPD | 2.88 | 5.78E-02 | 6.55 |
| ITGA1 | -3.34 | 1.32E-02 | 7.22 |
| ITGA11 | -2.86 | 7.03E-02 | 5.54 |
| ITGB2 | -3.58 | 2.92E-02 | 4.09 |
| ITGB3 | -3.01 | 3.84E-02 | 6.68 |
| ITGB6 | 2.61 | 1.46E-01 | 5.57 |
| ITIH1 | -5.81 | 8.42E-03 | 2.62 |
| ITIH2 | -7.66 | 1.67E-07 | 5.13 |
| ITIH5 | 3.22 | 1.65E-02 | 8.55 |
| ITLN2 | -4.45 | 7.16E-02 | -0.97 |
| ITPK1 | 6.13 | 1.09E-03 | 3.42 |
| IYD | -4.64 | 4.99E-02 | -0.97 |
| JDP2 | -2.71 | 1.98E-01 | 4.39 |
| JHY | 2.32 | 1.97E-01 | 6.97 |
| JMJD1C-AS1 | -3.37 | 1.72E-01 | 2.99 |
| KANK4 | -2.90 | 4.40E-02 | 7.66 |
| KCNA5 | 3.03 | 9.10E-02 | 4.68 |
| KCNAB1 | 2.57 | 1.35E-01 | 5.96 |
| KCNJ10 | -3.45 | 8.25E-02 | 3.76 |
| KCNJ13 | 4.62 | 3.26E-04 | 7.13 |
| KCNJ16 | 3.75 | 1.11E-02 | 5.23 |
| KCNJ2 | 3.34 | 1.26E-02 | 8.11 |
| KCNK13 | 4.07 | 7.55E-03 | 4.97 |
| KCNMB1 | 3.53 | 1.07E-01 | -0.97 |
| KCNQ3 | 2.30 | 1.80E-01 | 9.00 |
| KCTD8 | 3.72 | 1.43E-02 | 5.02 |
| KDF1 | -5.10 | 5.14E-02 | 2.26 |
| KDM5D | -5.28 | 8.71E-03 | -0.97 |
| KDR | -5.60 | 6.97E-06 | 8.63 |
| KIAA2012 | 2.77 | 1.37E-01 | 5.06 |
| KIF18B | -2.62 | 1.05E-01 | 6.61 |
| KIF20A | -2.51 | 1.14E-01 | 8.26 |
| KIF2C | -2.41 | 1.43E-01 | 8.35 |
| KIF5C | 6.11 | 8.32E-06 | 5.99 |
| KIF6 | 4.36 | 1.34E-01 | 2.53 |
| KIF9 | 3.47 | 1.38E-02 | 6.26 |
| KLB | -8.02 | 1.01E-06 | -0.97 |
| KLHL40 | -4.39 | 2.46E-02 | 2.91 |
| KLHL6 | 2.96 | 1.07E-01 | 4.64 |
| KLK6 | -2.60 | 1.89E-01 | 4.48 |
| KLRA1P | -2.51 | 1.40E-01 | 6.43 |
| KNG1 | -5.85 | 1.77E-03 | -0.97 |
| KREMEN1 | 2.56 | 1.07E-01 | 7.86 |
| KRT16 | -6.16 | 6.37E-04 | 2.80 |
| KRT16P3 | -4.81 | 3.02E-02 | -0.97 |
| KRT19 | -3.57 | 5.84E-03 | 9.40 |
| KRT24 | -6.90 | 6.12E-05 | -0.97 |
| KRT7 | -3.77 | 4.66E-02 | 3.60 |
| KYNU | -4.10 | 1.60E-01 | -0.97 |
| L1TD1 | -4.23 | 8.75E-04 | 7.38 |
| LAD1 | -5.59 | 3.72E-03 | -0.97 |
| LAMA2 | 2.88 | 4.17E-02 | 9.36 |

| | | | |
|---|---|---|---|
| LAMB3 | -4.75 | 6.27E-04 | 5.41 |
| LAMC2 | -3.41 | 2.40E-02 | 5.01 |
| LANCL3 | 3.70 | 2.95E-02 | 4.20 |
| LAPTM5 | -4.63 | 1.17E-03 | 4.84 |
| LCA5L | 3.49 | 4.88E-02 | 4.10 |
| LCAT | -2.92 | 6.04E-02 | 5.57 |
| LCP1 | -4.60 | 6.37E-04 | 5.72 |
| LEAP2 | -3.51 | 1.20E-01 | 3.06 |
| LENG8 | -5.85 | 1.77E-03 | -0.97 |
| LGALS2 | -5.39 | 7.00E-03 | -0.97 |
| LGI1 | 2.98 | 4.09E-02 | 7.09 |
| LGI3 | 5.12 | 4.10E-04 | 4.91 |
| LGR6 | -3.68 | 5.57E-02 | 3.56 |
| LGSN | 3.57 | 1.60E-01 | 3.14 |
| LHFPL6 | 2.40 | 1.47E-01 | 8.78 |
| LHX2 | -2.73 | 6.82E-02 | 7.88 |
| LIN28A | -3.37 | 1.09E-02 | 8.73 |
| LINC00261 | -6.98 | 5.01E-05 | -0.97 |
| LINC00271 | 4.89 | 4.03E-02 | 2.80 |
| LINC00305 | -4.10 | 5.14E-02 | 2.76 |
| LINC00312 | -3.00 | 1.22E-01 | 4.02 |
| LINC00379 | -4.10 | 1.60E-01 | -0.97 |
| LINC00461 | -4.67 | 1.69E-04 | 8.75 |
| LINC00473 | 5.65 | 3.44E-05 | 5.99 |
| LINC00488 | 6.14 | 1.03E-03 | 3.43 |
| LINC00518 | 3.89 | 4.99E-02 | -0.97 |
| LINC00616 | 3.17 | 1.95E-01 | -0.97 |
| LINC00643 | 3.18 | 2.17E-02 | 7.59 |
| LINC00678 | -3.96 | 1.95E-01 | -0.97 |
| LINC00906 | 4.17 | 2.55E-02 | -0.97 |
| LINC01011 | 2.70 | 1.84E-01 | 4.87 |
| LINC01091 | 5.77 | 2.63E-04 | -0.97 |
| LINC01198 | -2.81 | 1.78E-01 | 4.12 |
| LINC01266 | -6.19 | 5.90E-04 | 2.81 |
| LINC01356 | -3.16 | 7.44E-02 | 4.30 |
| LINC01436 | 3.68 | 8.72E-02 | -0.97 |
| LINC01497 | 4.06 | 3.57E-02 | -0.97 |
| LINC01551 | -9.69 | 1.74E-09 | 4.56 |
| LINC01621 | 4.32 | 1.53E-01 | 2.51 |
| LINC01679 | 6.50 | 3.26E-05 | 4.60 |
| LINC01697 | -4.81 | 1.03E-01 | 2.12 |
| LINC01765 | 3.79 | 9.68E-02 | 3.25 |
| LINC01844 | 4.27 | 2.85E-02 | 3.49 |
| LINC01933 | 3.17 | 1.14E-01 | 3.94 |
| LINC02006 | -4.54 | 1.88E-01 | 1.99 |
| LINC02154 | 6.04 | 1.31E-04 | 4.37 |
| LINC02268 | -5.34 | 7.75E-03 | -0.97 |
| LINC02380 | 5.09 | 2.11E-03 | -0.97 |
| LINC02381 | -2.76 | 6.15E-02 | 7.93 |
| LINC02488 | -4.10 | 1.60E-01 | -0.97 |
| LINC02520 | 3.89 | 4.99E-02 | -0.97 |
| LIPC | -6.34 | 3.85E-04 | -0.97 |
| LMCD1 | -2.67 | 8.59E-02 | 7.18 |
| LMNTD1 | 4.85 | 4.19E-03 | -0.97 |
| LMO7 | -2.83 | 7.16E-02 | 5.88 |
| LMX1A | 3.67 | 4.19E-03 | 9.41 |
| LMX1B | 3.24 | 1.63E-02 | 8.31 |
| LNCPRESS1 | -4.81 | 3.02E-02 | -0.97 |
| LNCPRESS2 | -3.96 | 1.95E-01 | -0.97 |

| | | | |
|---|---|---|---|
| LOC100130331 | -4.50 | 1.77E-02 | 2.96 |
| LOC100240735 | -4.10 | 1.60E-01 | -0.97 |
| LOC100286906 | 4.75 | 6.14E-03 | -0.97 |
| LOC100287425 | 4.41 | 1.20E-01 | 2.56 |
| LOC100505841 | 3.02 | 5.29E-02 | 5.77 |
| LOC100506114 | 3.27 | 1.95E-01 | -0.97 |
| LOC100507477 | 3.61 | 8.72E-02 | -0.97 |
| LOC101927013 | 3.82 | 8.75E-02 | 3.26 |
| LOC101927057 | 5.70 | 3.57E-04 | -0.97 |
| LOC101927120 | 3.58 | 9.82E-03 | 6.54 |
| LOC101927263 | -4.54 | 1.88E-01 | 1.99 |
| LOC101927267 | 2.82 | 1.35E-01 | 4.93 |
| LOC101927888 | 3.68 | 8.72E-02 | -0.97 |
| LOC101928202 | -3.96 | 1.95E-01 | -0.97 |
| LOC101928217 | 5.77 | 2.63E-04 | -0.97 |
| LOC101928499 | 5.77 | 3.27E-03 | 3.24 |
| LOC101928635 | 4.36 | 1.34E-01 | 2.53 |
| LOC101928817 | 4.89 | 8.05E-04 | 5.12 |
| LOC101928842 | 3.86 | 3.34E-02 | 3.87 |
| LOC101929007 | -5.10 | 5.14E-02 | 2.26 |
| LOC101929322 | 4.08 | 7.27E-03 | 4.98 |
| LOC102723694 | -2.65 | 1.16E-01 | 5.90 |
| LOC102723989 | 3.61 | 6.54E-02 | 3.74 |
| LOC102724165 | -4.54 | 6.01E-02 | -0.97 |
| LOC102724680 | 3.17 | 1.14E-01 | 3.94 |
| LOC102724731 | -3.08 | 8.63E-02 | 4.43 |
| LOC102724785 | 4.06 | 3.57E-02 | -0.97 |
| LOC102724858 | -5.10 | 5.14E-02 | 2.26 |
| LOC102724859 | 4.27 | 9.71E-04 | 7.01 |
| LOC102724945 | 4.49 | 1.26E-02 | -0.97 |
| LOC102725258 | -4.22 | 1.29E-01 | -0.97 |
| LOC105369151 | -2.59 | 9.57E-02 | 7.99 |
| LOC105369212 | -2.60 | 1.57E-01 | 5.18 |
| LOC105369315 | -3.54 | 1.88E-01 | 2.49 |
| LOC105369486 | 3.53 | 1.07E-01 | -0.97 |
| LOC105369795 | 5.51 | 7.75E-03 | 3.11 |
| LOC105369869 | 4.68 | 7.00E-03 | -0.97 |
| LOC105370169 | 4.17 | 2.55E-02 | -0.97 |
| LOC105370202 | 5.80 | 3.05E-03 | 3.26 |
| LOC105370259 | 3.89 | 7.99E-02 | 3.30 |
| LOC105370421 | -4.22 | 1.29E-01 | -0.97 |
| LOC105370463 | -3.96 | 1.95E-01 | -0.97 |
| LOC105370733 | -4.54 | 6.01E-02 | -0.97 |
| LOC105370767 | 4.88 | 5.66E-04 | 5.38 |
| LOC105370809 | 3.68 | 1.29E-01 | 3.20 |
| LOC105371114 | 3.45 | 1.29E-01 | -0.97 |
| LOC105371240 | 3.00 | 6.85E-02 | 5.31 |
| LOC105371241 | 3.53 | 1.07E-01 | -0.97 |
| LOC105371267 | 4.02 | 2.24E-03 | 6.82 |
| LOC105371455 | -7.24 | 2.06E-05 | 3.33 |
| LOC105371492 | -3.37 | 1.72E-01 | 2.99 |
| LOC105371509 | 3.45 | 1.95E-01 | 3.08 |
| LOC105371561 | 3.17 | 1.95E-01 | -0.97 |
| LOC105371611 | -4.73 | 1.19E-01 | 2.08 |
| LOC105371629 | -5.73 | 1.01E-02 | 2.58 |
| LOC105371908 | 3.36 | 1.60E-01 | -0.97 |
| LOC105371932 | 3.29 | 8.44E-02 | 4.00 |
| LOC105371941 | 3.17 | 1.95E-01 | -0.97 |
| LOC105372097 | 3.78 | 1.22E-02 | 5.05 |

| | | | |
|---|---|---|---|
| LOC105372135 | 3.27 | 1.95E-01 | -0.97 |
| LOC105372310 | -2.67 | 1.60E-01 | 4.51 |
| LOC105373042 | 3.82 | 6.01E-02 | -0.97 |
| LOC105373530 | 5.41 | 8.75E-04 | -0.97 |
| LOC105373605 | -4.81 | 3.02E-02 | -0.97 |
| LOC105373654 | 4.68 | 7.00E-03 | -0.97 |
| LOC105373730 | 6.18 | 9.37E-04 | 3.45 |
| LOC105373957 | 3.36 | 1.60E-01 | -0.97 |
| LOC105374013 | -2.35 | 1.74E-01 | 7.46 |
| LOC105374042 | 3.12 | 1.22E-01 | 3.91 |
| LOC105374336 | 3.51 | 4.88E-02 | 4.11 |
| LOC105374353 | 3.45 | 1.29E-01 | -0.97 |
| LOC105374396 | 3.17 | 1.95E-01 | -0.97 |
| LOC105374474 | -4.81 | 1.03E-01 | 2.12 |
| LOC105374557 | -3.40 | 8.93E-02 | 3.74 |
| LOC105374591 | 3.82 | 6.01E-02 | -0.97 |
| LOC105374794 | 11.62 | 1.98E-12 | 6.16 |
| LOC105374815 | -4.22 | 1.29E-01 | -0.97 |
| LOC105374972 | 8.58 | 1.17E-08 | -0.97 |
| LOC105375213 | -5.84 | 1.94E-05 | 5.64 |
| LOC105375704 | 4.45 | 1.07E-01 | 2.58 |
| LOC105375906 | -5.03 | 5.93E-02 | 2.23 |
| LOC105375943 | 4.95 | 3.30E-03 | -0.97 |
| LOC105375988 | 3.36 | 1.60E-01 | -0.97 |
| LOC105376218 | -4.34 | 8.72E-02 | -0.97 |
| LOC105376267 | -5.39 | 2.46E-02 | 2.41 |
| LOC105376277 | 4.01 | 5.57E-02 | 3.36 |
| LOC105376654 | 4.75 | 5.48E-02 | 2.73 |
| LOC105377008 | 3.45 | 1.29E-01 | -0.97 |
| LOC105377375 | 3.36 | 1.60E-01 | -0.97 |
| LOC105377412 | -4.64 | 4.99E-02 | -0.97 |
| LOC105377632 | -4.81 | 1.03E-01 | 2.12 |
| LOC105377876 | 3.04 | 7.00E-02 | 5.20 |
| LOC105377901 | 5.45 | 8.85E-03 | 3.08 |
| LOC105378008 | -3.05 | 1.16E-01 | 3.83 |
| LOC105378029 | 6.61 | 1.89E-05 | -0.97 |
| LOC105378114 | 4.32 | 9.98E-03 | 4.10 |
| LOC105378119 | -3.45 | 1.07E-01 | 3.44 |
| LOC105378308 | -6.62 | 1.47E-04 | -0.97 |
| LOC105378309 | -5.22 | 4.03E-02 | 2.33 |
| LOC105378589 | 3.17 | 1.95E-01 | -0.97 |
| LOC105378597 | 4.41 | 1.43E-02 | -0.97 |
| LOC105378696 | 3.89 | 4.99E-02 | -0.97 |
| LOC105378936 | -3.79 | 1.32E-02 | 4.78 |
| LOC105379041 | 7.23 | 3.55E-05 | 3.97 |
| LOC105379335 | -4.10 | 1.60E-01 | -0.97 |
| LOC105379506 | 7.76 | 2.14E-07 | -0.97 |
| LOC107984328 | 4.02 | 8.45E-03 | 4.95 |
| LOC107984414 | 4.32 | 1.90E-02 | -0.97 |
| LOC107984498 | 6.70 | 1.78E-04 | 3.70 |
| LOC107984900 | -5.03 | 5.93E-02 | 2.23 |
| LOC107984919 | -4.16 | 4.54E-02 | 2.80 |
| LOC107985143 | 3.68 | 8.72E-02 | -0.97 |
| LOC107985177 | 3.01 | 9.59E-02 | 4.67 |
| LOC107985221 | -3.07 | 1.95E-01 | 3.57 |
| LOC107985634 | 3.45 | 1.29E-01 | -0.97 |
| LOC107985653 | 3.68 | 8.72E-02 | -0.97 |
| LOC107985678 | -2.70 | 1.53E-01 | 4.65 |
| LOC107985681 | 4.32 | 1.53E-01 | 2.51 |

| | | | |
|---|---|---|---|
| LOC107985780 | -3.55 | 1.63E-02 | 4.95 |
| LOC107985939 | -2.52 | 1.35E-01 | 6.15 |
| LOC107985944 | 5.77 | 5.01E-05 | 5.24 |
| LOC107985992 | 6.23 | 6.31E-05 | -0.97 |
| LOC107986058 | -5.26 | 1.61E-04 | 4.93 |
| LOC107986298 | 5.15 | 7.40E-04 | 4.51 |
| LOC107986523 | 7.06 | 3.12E-06 | -0.97 |
| LOC107986655 | 4.01 | 4.20E-02 | -0.97 |
| LOC107986669 | 5.95 | 1.79E-04 | 4.33 |
| LOC107986673 | -3.08 | 9.70E-02 | 4.06 |
| LOC107986725 | 2.88 | 1.21E-01 | 4.96 |
| LOC107986862 | -4.22 | 1.29E-01 | -0.97 |
| LOC107987116 | 3.82 | 8.75E-02 | 3.26 |
| LOC107987208 | 4.12 | 9.15E-03 | 4.73 |
| LOC109864269 | -5.66 | 5.98E-04 | 3.55 |
| LOC112267873 | 4.41 | 7.75E-03 | 4.14 |
| LOC112267931 | 6.28 | 5.35E-05 | -0.97 |
| LOC112268037 | 5.01 | 2.72E-02 | 2.86 |
| LOC112268114 | -3.67 | 1.19E-02 | 5.25 |
| LOC112268177 | -5.39 | 7.00E-03 | -0.97 |
| LOC112268254 | -2.66 | 1.61E-01 | 4.95 |
| LOC112268256 | -2.62 | 1.29E-01 | 5.42 |
| LOC112268449 | 3.65 | 1.02E-02 | 5.76 |
| LOC112268460 | -3.67 | 4.45E-02 | 3.87 |
| LOC145694 | 4.06 | 3.57E-02 | -0.97 |
| LOC145845 | -4.34 | 2.78E-02 | 2.89 |
| LOC400499 | -3.96 | 7.68E-02 | 2.70 |
| LOC400706 | -4.54 | 6.01E-02 | -0.97 |
| LOC401261 | -3.11 | 4.34E-02 | 5.44 |
| LOC440982 | 3.34 | 2.35E-02 | 5.83 |
| LOC441204 | 2.51 | 1.30E-01 | 6.93 |
| LOC645529 | 3.75 | 7.16E-02 | -0.97 |
| LOC91548 | 3.17 | 1.95E-01 | -0.97 |
| LOXHD1 | 3.45 | 1.29E-01 | -0.97 |
| LOXL4 | -3.48 | 3.89E-02 | 4.04 |
| LPA | -4.22 | 1.29E-01 | -0.97 |
| LPCAT1 | -3.60 | 2.11E-02 | 4.84 |
| LRAT | 2.42 | 1.63E-01 | 6.85 |
| LRP2BP | 3.55 | 1.01E-02 | 6.59 |
| LRRC10B | 2.56 | 1.15E-01 | 7.36 |
| LRRC18 | 4.22 | 2.19E-02 | -0.97 |
| LRRC19 | -6.06 | 8.75E-04 | -0.97 |
| LRRC2 | -3.34 | 1.29E-01 | 3.39 |
| LRRC32 | -5.12 | 8.02E-05 | 6.36 |
| LRRC37A6P | 2.78 | 1.98E-01 | 4.33 |
| LRRC43 | 2.88 | 7.10E-02 | 5.79 |
| LRRC46 | 5.29 | 4.85E-04 | 4.59 |
| LRRC53 | -3.50 | 8.75E-02 | 3.46 |
| LRRC71 | 4.41 | 2.11E-02 | 3.56 |
| LRRC73 | 3.48 | 1.47E-02 | 6.00 |
| LRRC74B | 4.34 | 2.43E-02 | 3.52 |
| LRRC77P | 3.87 | 1.85E-02 | 4.29 |
| LRRC9 | 2.39 | 1.75E-01 | 7.07 |
| LRRN4 | -9.07 | 1.89E-08 | 4.25 |
| LSS | 3.27 | 1.95E-01 | -0.97 |
| LUM | -3.09 | 2.17E-02 | 11.52 |
| LY6H | -5.28 | 3.13E-02 | 2.36 |
| LYVE1 | -5.39 | 2.46E-02 | 2.41 |
| LYZ | -4.41 | 1.27E-02 | 3.51 |

| | | | |
|---|---|---|---|
| MACC1 | -3.45 | 4.03E-02 | 4.25 |
| MAEL | 3.27 | 1.95E-01 | -0.97 |
| MAFB | 3.68 | 4.57E-03 | 8.40 |
| MAK | 4.38 | 8.54E-03 | 4.13 |
| MAML1 | -5.03 | 4.12E-03 | 3.23 |
| MAN1A1 | -2.24 | 2.00E-01 | 9.62 |
| MAP1LC3C | 5.58 | 8.51E-06 | 8.43 |
| MAP3K19 | 6.83 | 2.09E-06 | 5.35 |
| MARCO | 5.06 | 2.35E-03 | -0.97 |
| MARVELD3 | -4.13 | 1.80E-02 | 3.78 |
| MASCRNA | -2.77 | 8.55E-02 | 5.96 |
| MASP1 | -2.75 | 7.08E-02 | 7.24 |
| MAT1A | -4.40 | 7.33E-04 | 6.38 |
| MB | 4.01 | 5.57E-02 | 3.36 |
| MBP | 3.73 | 5.66E-03 | 6.74 |
| MCHR1 | 2.68 | 1.61E-01 | 5.02 |
| MCIDAS | 3.93 | 1.49E-02 | 4.32 |
| MCOLN2 | -5.16 | 4.54E-02 | 2.30 |
| MCOLN3 | -7.02 | 5.63E-06 | 4.23 |
| MCPH1-AS1 | 3.36 | 1.60E-01 | -0.97 |
| MCTP2 | -5.82 | 1.47E-04 | 4.21 |
| MDGA2 | 2.43 | 1.56E-01 | 7.09 |
| MDH1B | 2.76 | 9.11E-02 | 5.98 |
| MEIOB | -4.45 | 2.19E-02 | 2.94 |
| MEIS1 | -2.84 | 4.95E-02 | 8.08 |
| MEOX1 | 4.65 | 2.49E-04 | 7.44 |
| MEP1A | -8.84 | 4.70E-08 | -0.97 |
| MEP1B | -3.96 | 1.95E-01 | -0.97 |
| MET | 3.38 | 1.32E-02 | 7.26 |
| METTL7A | -2.80 | 5.57E-02 | 7.98 |
| METTL7B | -4.50 | 1.77E-02 | 2.96 |
| MFAP3L | 2.29 | 1.95E-01 | 7.94 |
| MFAP4 | -4.15 | 9.19E-04 | 8.47 |
| MGAT4B | -3.73 | 1.97E-02 | 4.16 |
| MGC16275 | 2.84 | 1.78E-01 | 4.36 |
| MIA2 | -4.64 | 4.99E-02 | -0.97 |
| MINOS1-NBL1 | 2.35 | 1.60E-01 | 9.31 |
| MIR1-1HG-AS1 | -3.57 | 1.01E-02 | 5.89 |
| MIR1282 | 2.99 | 1.27E-01 | 4.43 |
| MIR17HG | -3.61 | 1.10E-02 | 5.61 |
| MIR205HG | -5.10 | 1.43E-02 | -0.97 |
| MIR214 | -4.64 | 4.99E-02 | -0.97 |
| MIR217HG | 3.20 | 7.72E-02 | 4.54 |
| MIR302D | -4.22 | 1.29E-01 | -0.97 |
| MIR34AHG | -2.80 | 6.04E-02 | 7.09 |
| MIR3606 | -3.93 | 1.71E-03 | 9.72 |
| MIR3615 | -3.06 | 2.72E-02 | 8.16 |
| MIR503HG | -4.32 | 3.51E-03 | 4.46 |
| MIR6513 | -3.12 | 1.78E-01 | 3.60 |
| MIR675 | -6.70 | 1.11E-04 | -0.97 |
| MIR9-2 | -3.22 | 1.36E-01 | 3.65 |
| MIR9-3HG | -4.82 | 2.15E-04 | 6.12 |
| MITF | 3.83 | 4.20E-03 | 6.59 |
| MIXL1 | 3.12 | 1.22E-01 | 3.91 |
| MKI67 | -2.77 | 5.55E-02 | 9.65 |
| MLANA | 9.07 | 7.27E-11 | 7.89 |
| MLC1 | -4.28 | 3.90E-03 | 4.44 |
| MLPH | 2.81 | 6.53E-02 | 6.76 |
| MME | -5.18 | 4.19E-05 | 7.66 |

| | | | |
|---|---|---|---|
| MMP1 | -5.73 | 4.84E-04 | 3.58 |
| MND1 | -2.71 | 1.25E-01 | 5.39 |
| MORN5 | 3.67 | 1.64E-02 | 5.00 |
| MPO | -8.32 | 2.99E-07 | -0.97 |
| MPPED1 | -3.00 | 3.94E-02 | 6.64 |
| MPZ | -3.91 | 4.38E-03 | 5.84 |
| MRC1 | -5.34 | 2.78E-02 | 2.39 |
| MRNIP | -5.03 | 5.93E-02 | 2.23 |
| MS4A10 | -5.34 | 7.75E-03 | -0.97 |
| MS4A6A | 5.82 | 4.36E-05 | 5.26 |
| MS4A8 | 3.53 | 7.56E-02 | 3.70 |
| MSC-AS1 | 3.17 | 1.09E-01 | 4.26 |
| MSR1 | -3.18 | 8.25E-02 | 3.89 |
| MT1E | -4.92 | 5.95E-03 | 3.18 |
| MT1G | -6.00 | 1.13E-03 | -0.97 |
| MT1H | -5.85 | 1.77E-03 | -0.97 |
| MTTP | -3.94 | 2.11E-03 | 7.41 |
| MUC12 | 3.03 | 1.54E-01 | 3.87 |
| MUSK | 3.26 | 2.77E-02 | 5.89 |
| MYCT1 | -5.68 | 5.40E-04 | 3.56 |
| MYH7 | 2.54 | 1.38E-01 | 6.48 |
| MYL4 | -6.45 | 4.93E-05 | 3.94 |
| MYL7 | -3.30 | 1.95E-01 | 2.95 |
| MYO3B | 3.10 | 6.41E-02 | 4.91 |
| MYRF | -4.29 | 5.41E-04 | 9.76 |
| MYRIP | 2.58 | 1.69E-01 | 5.45 |
| N4BP2L2-IT2 | -3.03 | 1.04E-01 | 4.23 |
| NALCN | 2.96 | 5.43E-02 | 6.08 |
| NANOG | -4.22 | 1.29E-01 | -0.97 |
| NANOGP8 | -3.96 | 1.95E-01 | -0.97 |
| NANOS3 | 3.36 | 1.60E-01 | -0.97 |
| NAP1L4 | 2.65 | 9.66E-02 | 6.81 |
| NAPRT | -3.75 | 6.80E-02 | 3.18 |
| NBL1 | 3.26 | 1.50E-02 | 8.44 |
| NDP | -3.40 | 8.93E-02 | 3.74 |
| NDUFA3 | 3.61 | 8.72E-02 | -0.97 |
| NDUFS1 | 4.90 | 7.85E-04 | 5.12 |
| NDUFS3 | -7.06 | 4.07E-05 | -0.97 |
| NEB | 5.32 | 4.25E-04 | 4.60 |
| NECAB1 | 3.68 | 7.51E-03 | 6.44 |
| NECTIN4 | -4.89 | 6.53E-03 | 3.16 |
| NEFL | 2.82 | 4.56E-02 | 12.89 |
| NEK10 | 4.82 | 4.75E-03 | -0.97 |
| NEK11 | 2.38 | 1.87E-01 | 6.67 |
| NEUROD1 | -3.42 | 1.15E-02 | 7.21 |
| NEUROD2 | -5.73 | 2.35E-03 | -0.97 |
| NEUROD6 | -6.79 | 8.39E-05 | 3.11 |
| NEUROG2 | -5.46 | 6.06E-05 | 5.77 |
| NFE2L3 | 2.76 | 6.42E-02 | 8.00 |
| NFYC-AS1 | -2.57 | 1.61E-01 | 5.09 |
| NID2 | -2.41 | 1.34E-01 | 10.59 |
| NKX1-2 | -3.22 | 1.77E-01 | 3.33 |
| NKX2-3 | -5.39 | 7.00E-03 | -0.97 |
| NLRC5 | -4.73 | 3.57E-02 | -0.97 |
| NME5 | 2.70 | 9.57E-02 | 6.29 |
| NMRAL1 | 3.17 | 1.95E-01 | -0.97 |
| NOX4 | -2.40 | 1.97E-01 | 5.61 |
| NPAS2 | 2.68 | 1.19E-01 | 5.86 |
| NPC1L1 | -4.42 | 7.98E-03 | 3.93 |

| | | | |
|---|---|---|---|
| NPNT | -8.40 | 3.31E-10 | 8.08 |
| NPR1 | -3.06 | 9.68E-02 | 4.25 |
| NPR3 | -3.26 | 1.43E-02 | 8.91 |
| NPY5R | 3.05 | 7.21E-02 | 4.88 |
| NQO1 | 2.68 | 7.16E-02 | 9.24 |
| NR2E1 | -3.03 | 3.27E-02 | 7.06 |
| NR2F2-AS1 | -3.22 | 6.82E-02 | 4.13 |
| NRAP | 4.27 | 1.72E-01 | 2.49 |
| NRG1 | 2.27 | 2.00E-01 | 8.15 |
| NRGN | -3.67 | 4.45E-02 | 3.87 |
| NTNG1 | 3.06 | 3.58E-02 | 6.64 |
| NTNG2 | 3.51 | 8.71E-03 | 7.72 |
| NUSAP1 | -2.31 | 1.78E-01 | 9.14 |
| NXPE4 | -4.34 | 8.72E-02 | -0.97 |
| NXPH2 | -3.96 | 7.91E-03 | 5.02 |
| OC90 | 7.24 | 1.63E-06 | -0.97 |
| OCA2 | 5.69 | 2.95E-05 | 6.00 |
| ODAM | -4.45 | 7.16E-02 | -0.97 |
| ODF3B | 2.62 | 1.09E-01 | 6.87 |
| OIT3 | -5.03 | 1.63E-02 | -0.97 |
| OLIG1 | -4.73 | 3.57E-02 | -0.97 |
| OLIG2 | -3.89 | 3.34E-02 | 3.66 |
| OLR1 | -7.99 | 1.13E-06 | -0.97 |
| ONECUT2 | 2.86 | 4.28E-02 | 10.06 |
| OPRK1 | -4.99 | 9.72E-04 | 4.53 |
| OPRM1 | -4.34 | 1.01E-02 | 3.89 |
| OR2C3 | 6.89 | 6.48E-06 | -0.97 |
| OR51E1 | -4.64 | 4.99E-02 | -0.97 |
| ORM1 | -8.02 | 1.01E-06 | -0.97 |
| OSR1 | -5.05 | 2.70E-04 | 5.24 |
| OSTM1-AS1 | -3.96 | 1.95E-01 | -0.97 |
| OTOL1 | 4.53 | 1.46E-02 | 3.62 |
| OTX2-AS1 | 3.68 | 1.04E-02 | 5.66 |
| OVOS2 | -3.81 | 4.27E-02 | 3.62 |
| P2RX1 | -4.73 | 1.19E-01 | 2.08 |
| PACRG | 3.93 | 1.49E-02 | 4.64 |
| PAEP | 5.14 | 1.77E-03 | -0.97 |
| PAH | -3.42 | 3.93E-02 | 4.43 |
| PALMD | 3.12 | 3.11E-02 | 6.50 |
| PANCR | -5.03 | 1.63E-02 | -0.97 |
| PANK4 | 4.01 | 4.20E-02 | -0.97 |
| PAPLN | -4.81 | 1.03E-01 | 2.12 |
| PAPPA | -3.04 | 2.62E-02 | 9.04 |
| PAPSS2 | 3.22 | 1.52E-02 | 9.71 |
| PARPBP | -2.57 | 1.49E-01 | 5.46 |
| PAX2 | 4.35 | 4.70E-03 | 4.53 |
| PAX3 | 3.55 | 7.56E-03 | 7.71 |
| PAX5 | 2.69 | 9.57E-02 | 6.28 |
| PAX8 | 4.45 | 3.27E-03 | 4.90 |
| PCAT14 | -4.32 | 3.01E-03 | 4.68 |
| PCAT19 | -4.34 | 8.72E-02 | -0.97 |
| PCDHAC2 | -2.92 | 5.27E-02 | 6.22 |
| PCDHB1 | 3.72 | 1.18E-01 | 3.21 |
| PCK1 | -4.01 | 7.69E-03 | 4.72 |
| PDE4B | 2.79 | 5.32E-02 | 9.30 |
| PDE4D | 2.28 | 1.87E-01 | 9.37 |
| PDE7B | 2.65 | 9.70E-02 | 6.68 |
| PDPN | -2.59 | 8.72E-02 | 9.66 |
| PDYN | -4.39 | 2.46E-02 | 2.91 |

| | | | |
|---|---|---|---|
| PDZRN4 | 2.88 | 4.49E-02 | 7.97 |
| PEAR1 | -3.16 | 2.31E-02 | 6.99 |
| PECAM1 | -6.98 | 1.27E-06 | 5.20 |
| PF4 | -7.89 | 1.64E-06 | -0.97 |
| PHF1 | -4.45 | 2.19E-02 | 2.94 |
| PIEZO1 | -2.37 | 1.59E-01 | 8.13 |
| PIF1 | -2.99 | 6.04E-02 | 5.53 |
| PIFO | 3.21 | 1.66E-02 | 8.91 |
| PIGAP1 | -4.22 | 1.29E-01 | -0.97 |
| PIH1D3 | 4.22 | 1.72E-01 | 2.46 |
| PINX1 | 3.53 | 1.07E-01 | -0.97 |
| PIP5K1B | 2.88 | 6.05E-02 | 6.49 |
| PITX1 | -7.74 | 1.70E-08 | 6.17 |
| PITX2 | -7.91 | 2.39E-08 | 5.99 |
| PIWIL2 | -2.47 | 1.68E-01 | 6.08 |
| PKHD1 | 3.61 | 8.72E-02 | -0.97 |
| PKHD1L1 | -6.98 | 2.94E-07 | 5.79 |
| PKP2 | -3.25 | 1.47E-02 | 8.98 |
| PKP3 | -5.34 | 1.64E-03 | 3.39 |
| PLA1A | 4.17 | 1.95E-01 | 2.44 |
| PLA2G12B | -5.10 | 3.43E-03 | 3.26 |
| PLA2G2A | -9.48 | 4.12E-09 | -0.97 |
| PLA2G3 | -3.05 | 6.43E-02 | 5.15 |
| PLAT | -2.80 | 5.08E-02 | 9.24 |
| PLAU | -2.50 | 1.74E-01 | 5.43 |
| PLCXD1 | 4.01 | 5.57E-02 | 3.36 |
| PLEK | -3.72 | 1.07E-02 | 5.03 |
| PLEKHB1 | 2.32 | 1.86E-01 | 7.68 |
| PLEKHD1 | 4.85 | 5.96E-03 | 3.78 |
| PLG | -10.78 | 4.13E-11 | -0.97 |
| PLLP | -4.20 | 5.10E-03 | 4.40 |
| PLSCR4 | -3.70 | 8.40E-03 | 5.82 |
| PLVAP | -5.53 | 1.61E-04 | 4.80 |
| PMEL | 4.83 | 8.02E-05 | 12.71 |
| PODXL | -6.71 | 8.60E-08 | 10.47 |
| POLR3E | 3.65 | 3.32E-02 | 4.18 |
| PON1 | -4.45 | 5.42E-03 | 4.26 |
| POSTN | -3.74 | 2.98E-03 | 11.87 |
| POU5F1 | -4.22 | 1.29E-01 | -0.97 |
| POU5F1P3 | -4.22 | 1.29E-01 | -0.97 |
| PPARGC1A | 4.03 | 1.95E-03 | 6.95 |
| PPBP | -8.54 | 1.40E-07 | -0.97 |
| PPP1R42 | 3.59 | 3.78E-02 | 4.15 |
| PRDM12 | -3.39 | 2.55E-02 | 5.32 |
| PRDM13 | -5.16 | 1.26E-02 | -0.97 |
| PRDM6 | -4.85 | 7.75E-03 | 3.14 |
| PRDM8 | -3.41 | 4.45E-02 | 4.01 |
| PRELP | 2.58 | 1.07E-01 | 7.57 |
| PRH1 | 3.53 | 1.07E-01 | -0.97 |
| PRH1-TAS2R14 | 3.45 | 1.29E-01 | -0.97 |
| PROCR | -3.21 | 3.33E-02 | 5.41 |
| PRODH2 | -5.16 | 4.54E-02 | 2.30 |
| PROZ | -4.45 | 7.16E-02 | -0.97 |
| PRPF31 | -4.92 | 5.95E-03 | 3.18 |
| PRPH | 2.33 | 1.61E-01 | 10.84 |
| PRR15L | -4.73 | 3.57E-02 | -0.97 |
| PRR18 | 3.65 | 1.41E-02 | 5.18 |
| PRR29 | 3.49 | 1.32E-02 | 6.10 |
| PRRT4 | -2.40 | 1.72E-01 | 6.67 |

| | | | |
|---|---|---|---|
| PRSS16 | -2.48 | 1.96E-01 | 5.04 |
| PRSS33 | 3.71 | 1.22E-02 | 5.21 |
| PRSS56 | 3.66 | 7.87E-03 | 6.58 |
| PSAPL1 | -5.03 | 5.93E-02 | 2.23 |
| PSMD6-AS2 | -3.17 | 1.50E-01 | 3.62 |
| PSME1 | 4.76 | 1.22E-03 | 5.06 |
| PTCH1 | -3.08 | 2.39E-02 | 8.77 |
| PTEN | -6.54 | 1.91E-04 | 2.99 |
| PTGDS | 4.89 | 8.26E-05 | 8.86 |
| PTGER4 | -3.39 | 1.18E-01 | 3.41 |
| PTGFR | -4.42 | 5.99E-03 | 4.25 |
| PTPRC | 6.74 | 1.25E-05 | 4.72 |
| PTPRN2 | 2.48 | 1.30E-01 | 7.48 |
| PUS7L | -6.28 | 4.40E-04 | -0.97 |
| QPCT | 2.34 | 1.98E-01 | 6.53 |
| RAB11FIP3 | -4.48 | 1.01E-02 | 3.54 |
| RAB25 | -5.68 | 2.28E-04 | 4.14 |
| RAB37 | -2.87 | 5.62E-02 | 6.64 |
| RAB38 | 2.48 | 1.23E-01 | 8.13 |
| RAC2 | -6.59 | 1.09E-05 | 4.60 |
| RAD51 | -2.71 | 1.25E-01 | 5.39 |
| RAD51AP1 | -2.36 | 1.70E-01 | 7.36 |
| RAG2 | 3.75 | 7.16E-02 | -0.97 |
| RAMP3 | 3.36 | 1.60E-01 | -0.97 |
| RASEF | 4.02 | 3.57E-03 | 6.06 |
| RASL11B | 2.43 | 1.45E-01 | 7.87 |
| RASSF6 | -4.34 | 1.47E-02 | 3.47 |
| RBFOX1 | 3.17 | 2.01E-02 | 8.43 |
| RBM24 | 2.55 | 1.13E-01 | 7.58 |
| RBP2 | -8.60 | 1.12E-07 | -0.97 |
| RBP4 | -3.93 | 1.75E-03 | 9.15 |
| RCSD1 | -3.16 | 1.95E-01 | 3.30 |
| RD3 | 3.99 | 6.04E-03 | 5.52 |
| RDH10 | 3.21 | 1.64E-02 | 9.06 |
| RELN | -3.11 | 2.10E-02 | 11.10 |
| REN | -5.29 | 6.31E-05 | 6.27 |
| RESP18 | 3.17 | 5.37E-02 | 4.94 |
| RET | 3.13 | 2.19E-02 | 8.36 |
| RFX6 | -4.10 | 1.60E-01 | -0.97 |
| RGS13 | -6.50 | 2.14E-04 | -0.97 |
| RGS18 | -5.54 | 4.19E-03 | -0.97 |
| RGS22 | 4.21 | 7.26E-03 | 4.46 |
| RGS6 | -3.30 | 1.95E-01 | 2.95 |
| RGS9 | 2.56 | 1.33E-01 | 6.34 |
| RHAG | -4.81 | 3.02E-02 | -0.97 |
| RHBG | -4.73 | 3.57E-02 | -0.97 |
| RHOH | 4.45 | 1.32E-03 | 5.75 |
| RIBC1 | 3.86 | 3.34E-02 | 3.87 |
| RIBC2 | 4.18 | 7.72E-03 | 4.77 |
| RIPOR3 | -4.88 | 5.34E-04 | 4.96 |
| RN7SL608P | -4.54 | 1.88E-01 | 1.99 |
| RNASE1 | -4.61 | 7.28E-03 | 3.60 |
| RNF128 | -3.70 | 2.10E-02 | 4.15 |
| RNF187 | -3.69 | 1.14E-02 | 5.02 |
| RNF219-AS1 | 4.36 | 1.34E-01 | 2.53 |
| RNLS | 2.78 | 9.84E-02 | 5.83 |
| ROBO4 | -3.54 | 1.88E-01 | 2.49 |
| ROPN1 | 4.45 | 1.07E-01 | 2.58 |
| ROPN1L | 5.33 | 1.98E-04 | 5.02 |

| | | | |
|---|---|---|---|
| RPE65 | 7.29 | 1.42E-07 | 6.32 |
| RPH3A | 3.10 | 3.44E-02 | 6.43 |
| RPL29P19 | -4.16 | 4.54E-02 | 2.80 |
| RPRM | 3.00 | 3.28E-02 | 8.08 |
| RPS17 | -5.39 | 1.45E-05 | 8.95 |
| RPS3AP38 | -3.39 | 1.18E-01 | 3.41 |
| RPS4Y1 | -7.05 | 4.19E-05 | -0.97 |
| RPS9 | -3.77 | 5.72E-03 | 6.06 |
| RSPH1 | 4.14 | 1.65E-03 | 6.67 |
| RSPH10B | 3.95 | 4.20E-02 | -0.97 |
| RSPH4A | 3.89 | 3.65E-03 | 6.55 |
| RTL1 | 4.99 | 7.97E-05 | 7.61 |
| RUFY1 | -3.66 | 2.19E-02 | 4.36 |
| S100A10 | -3.16 | 1.90E-02 | 8.92 |
| S100A14 | -8.51 | 1.93E-08 | 4.97 |
| S100A16 | -2.79 | 6.91E-02 | 6.28 |
| SAMD15 | 2.49 | 1.63E-01 | 5.92 |
| SAMD3 | -3.11 | 8.11E-02 | 4.44 |
| SAMSN1 | -3.64 | 1.60E-01 | 2.53 |
| SAXO2 | 2.55 | 1.95E-01 | 5.09 |
| SCARA5 | 5.21 | 7.51E-05 | 6.42 |
| SCARNA17 | -2.68 | 9.57E-02 | 6.30 |
| SCARNA9 | -2.82 | 9.36E-02 | 5.37 |
| SCGB3A2 | -4.34 | 8.72E-02 | -0.97 |
| SCGN | 5.84 | 8.38E-06 | 6.73 |
| SCN9A | 2.46 | 1.35E-01 | 7.87 |
| SCRG1 | -3.38 | 2.61E-02 | 5.11 |
| SCRIB | 5.95 | 1.47E-04 | -0.97 |
| SELP | -5.22 | 1.11E-02 | -0.97 |
| SEMA3C | 2.42 | 1.29E-01 | 10.62 |
| SEMA3E | -2.93 | 7.16E-02 | 5.43 |
| SEMA3G | -3.12 | 5.31E-02 | 5.18 |
| SERPINA1 | -10.50 | 7.27E-11 | -0.97 |
| SERPINA11 | -4.34 | 8.72E-02 | -0.97 |
| SERPINA3 | 3.15 | 1.16E-01 | 4.25 |
| SERPINA4 | -4.22 | 1.29E-01 | -0.97 |
| SERPINA7 | -4.73 | 3.57E-02 | -0.97 |
| SERPINC1 | -6.73 | 1.05E-04 | 3.08 |
| SERPIND1 | -6.75 | 9.90E-05 | 3.09 |
| SERPINE3 | -6.42 | 2.10E-05 | 4.51 |
| SERPINF1 | 2.55 | 9.57E-02 | 11.15 |
| SERTM1 | 2.68 | 1.43E-01 | 5.40 |
| SFN | -4.54 | 1.88E-01 | 1.99 |
| SFRP5 | -5.64 | 3.30E-03 | -0.97 |
| SGCD | 4.24 | 6.66E-03 | 4.80 |
| SGCG | -2.88 | 1.02E-01 | 4.61 |
| SGPP2 | 4.90 | 1.85E-04 | 6.39 |
| SH2D3A | -4.73 | 3.57E-02 | -0.97 |
| SH3RF2 | -4.60 | 1.00E-03 | 5.34 |
| SH3RF3 | 4.01 | 3.92E-03 | 5.82 |
| SH3RF3-AS1 | 4.92 | 3.65E-02 | 2.81 |
| SHISA3 | -7.38 | 1.17E-05 | 3.41 |
| SHISA9 | 4.14 | 1.21E-03 | 7.67 |
| SI | -6.31 | 4.10E-04 | -0.97 |
| SIDT1 | 3.95 | 6.68E-02 | 3.33 |
| SIK1 | 2.97 | 9.47E-02 | 5.01 |
| SIK1B | 2.68 | 8.19E-02 | 7.39 |
| SINHCAF | -4.48 | 6.73E-04 | 5.96 |
| SIX2 | -3.49 | 2.96E-02 | 4.78 |

| | | | |
|---|---|---|---|
| SLC13A4 | 6.28 | 2.26E-06 | 6.66 |
| SLC13A5 | -4.24 | 1.97E-03 | 5.54 |
| SLC16A6 | 3.00 | 3.89E-02 | 6.86 |
| SLC17A1 | -4.10 | 1.60E-01 | -0.97 |
| SLC17A8 | 4.00 | 2.78E-03 | 6.52 |
| SLC22A3 | -6.13 | 7.50E-04 | 2.78 |
| SLC22A7 | -6.39 | 5.44E-05 | 3.91 |
| SLC23A1 | -5.50 | 5.44E-03 | -0.97 |
| SLC24A4 | 3.25 | 6.68E-02 | 4.57 |
| SLC24A5 | 5.79 | 3.17E-04 | 4.25 |
| SLC26A3 | -4.81 | 1.03E-01 | 2.12 |
| SLC2A2 | -4.22 | 1.29E-01 | -0.97 |
| SLC2A5 | -3.83 | 1.19E-02 | 4.80 |
| SLC34A2 | -8.78 | 1.32E-09 | 6.10 |
| SLC35F2 | -3.17 | 2.61E-02 | 6.35 |
| SLC39A12 | 5.59 | 4.79E-04 | -0.97 |
| SLC39A2 | -3.58 | 2.92E-02 | 4.09 |
| SLC39A5 | -6.11 | 5.83E-05 | 4.36 |
| SLC40A1 | -5.08 | 5.01E-05 | 8.36 |
| SLC45A2 | 8.06 | 8.12E-08 | -0.97 |
| SLC47A1 | -4.62 | 1.02E-03 | 5.19 |
| SLC4A5 | 4.82 | 9.66E-05 | 9.45 |
| SLC51A | -6.03 | 9.52E-04 | 2.73 |
| SLC5A10 | 5.38 | 1.12E-03 | 4.05 |
| SLC5A12 | -6.17 | 1.68E-06 | 7.05 |
| SLC6A15 | 2.27 | 1.89E-01 | 9.59 |
| SLC6A17 | 2.84 | 6.01E-02 | 6.90 |
| SLC7A2 | -2.67 | 7.16E-02 | 9.04 |
| SLC7A5 | 2.68 | 6.85E-02 | 10.45 |
| SLC7A7 | -5.05 | 6.28E-05 | 7.49 |
| SLC7A8 | 2.42 | 1.34E-01 | 9.66 |
| SLC9A3R1 | -2.94 | 3.36E-02 | 10.72 |
| SLCO1B1 | -3.45 | 1.34E-01 | 3.02 |
| SLCO1B3 | 4.12 | 3.02E-02 | -0.97 |
| SLCO1C1 | 4.83 | 1.99E-03 | 4.36 |
| SLCO2B1 | -5.86 | 5.62E-05 | 4.97 |
| SLN | -4.45 | 7.16E-02 | -0.97 |
| SLPI | -4.65 | 1.22E-03 | 4.63 |
| SMG1P7 | -2.57 | 1.27E-01 | 5.95 |
| SMIM24 | -2.83 | 1.78E-01 | 3.94 |
| SMKR1 | 3.12 | 1.22E-01 | 3.91 |
| SMLR1 | -6.06 | 8.75E-04 | -0.97 |
| SMN1 | -4.73 | 1.19E-01 | 2.08 |
| SMTN | -2.43 | 1.37E-01 | 7.82 |
| SNCA | 2.79 | 5.26E-02 | 9.34 |
| SNCB | 3.41 | 1.55E-02 | 6.31 |
| SNHG22 | -3.12 | 9.04E-02 | 4.08 |
| SNORA24 | -3.64 | 1.60E-01 | 2.53 |
| SNORA4 | -2.64 | 1.48E-01 | 5.36 |
| SNORA48 | -3.70 | 7.49E-02 | 3.15 |
| SNORA63B | -3.34 | 4.49E-02 | 4.71 |
| SNORA73A | -3.54 | 1.88E-01 | 2.49 |
| SNORD10 | -3.22 | 6.01E-02 | 4.65 |
| SNORD12B | -4.54 | 1.88E-01 | 1.99 |
| SNORD144 | -3.04 | 1.06E-01 | 4.04 |
| SNORD14B | -3.64 | 1.60E-01 | 2.53 |
| SNORD19 | -3.05 | 1.16E-01 | 3.83 |
| SNORD2 | -4.48 | 1.01E-02 | 3.54 |
| SNORD22 | -2.32 | 1.96E-01 | 6.90 |

| | | | |
|---|---|---|---|
| SNORD26 | -3.78 | 3.59E-02 | 3.93 |
| SNORD29 | -2.73 | 1.16E-01 | 5.08 |
| SNORD30 | -5.92 | 1.34E-03 | 2.68 |
| SNORD31 | -2.41 | 1.95E-01 | 5.62 |
| SNORD5 | -2.71 | 1.48E-01 | 4.88 |
| SNORD59B | -2.83 | 1.78E-01 | 3.94 |
| SNORD69 | -4.64 | 1.60E-01 | 2.03 |
| SNORD76 | -2.58 | 1.95E-01 | 4.71 |
| SNORD87 | -3.07 | 1.95E-01 | 3.57 |
| SNORD97 | -3.75 | 6.80E-02 | 3.18 |
| SNTN | 4.92 | 3.72E-03 | -0.97 |
| SOAT2 | -6.06 | 8.75E-04 | -0.97 |
| SOCS1 | -2.69 | 1.56E-01 | 4.97 |
| SOHLH2 | -3.58 | 1.07E-01 | 3.09 |
| SOST | 4.03 | 3.44E-03 | 6.07 |
| SOX14 | -4.54 | 6.01E-02 | -0.97 |
| SOX17 | -6.64 | 1.40E-04 | -0.97 |
| SOX18 | -3.64 | 1.60E-01 | 2.53 |
| SOX2-OT | 2.32 | 1.77E-01 | 8.90 |
| SOX21 | -2.93 | 4.15E-02 | 7.25 |
| SPACA9 | 2.53 | 1.29E-01 | 6.62 |
| SPAG17 | 3.15 | 2.40E-02 | 7.42 |
| SPAG6 | 4.46 | 4.22E-04 | 7.94 |
| SPAG8 | 2.60 | 1.20E-01 | 6.24 |
| SPARCL1 | 2.55 | 1.03E-01 | 8.51 |
| SPATA17 | 3.54 | 1.16E-02 | 6.12 |
| SPATA31C1 | 4.82 | 4.45E-02 | 2.76 |
| SPATA31C2 | 5.30 | 1.46E-03 | 4.01 |
| SPATS1 | 5.84 | 2.68E-03 | 3.27 |
| SPEF1 | 4.51 | 2.75E-03 | 4.93 |
| SPINK1 | -5.89 | 1.48E-03 | -0.97 |
| SPINK5 | -3.69 | 5.53E-03 | 6.73 |
| SPON1 | -4.30 | 5.66E-04 | 8.82 |
| SPR | -3.00 | 7.29E-02 | 5.02 |
| SPRN | 3.21 | 3.13E-02 | 6.05 |
| SRGAP3-AS2 | 3.82 | 6.01E-02 | -0.97 |
| SRGN | -5.85 | 7.75E-03 | 2.64 |
| SSBP3-AS1 | -4.03 | 5.93E-02 | 2.73 |
| SST | 3.11 | 2.10E-02 | 10.23 |
| ST14 | -3.61 | 2.44E-02 | 4.33 |
| ST6GALNAC2 | 2.87 | 1.16E-01 | 4.79 |
| STAB1 | -5.41 | 5.63E-04 | 4.01 |
| STAB2 | -5.81 | 1.94E-03 | -0.97 |
| STAM-AS1 | -2.83 | 1.78E-01 | 3.94 |
| STK32A | 3.05 | 4.28E-02 | 6.27 |
| STK32A-AS1 | 4.32 | 1.90E-02 | -0.97 |
| STK32B | 4.47 | 1.48E-03 | 5.40 |
| STMND1 | 3.73 | 4.80E-02 | 3.80 |
| STOML3 | 5.33 | 4.25E-04 | 4.60 |
| STPG2-AS1 | 3.17 | 1.95E-01 | -0.97 |
| STXBP5-AS1 | 2.86 | 9.68E-02 | 5.24 |
| SUCNR1 | -5.22 | 1.11E-02 | -0.97 |
| SULT1E1 | -6.79 | 8.39E-05 | -0.97 |
| SULT2A1 | -6.68 | 1.17E-04 | -0.97 |
| SV2B | 6.85 | 2.94E-07 | 6.59 |
| SV2C | 4.79 | 5.41E-04 | 5.56 |
| SVEP1 | -7.18 | 1.12E-07 | 6.31 |
| SYN2 | 2.39 | 1.89E-01 | 6.36 |
| SYTL3 | 3.68 | 2.96E-02 | 4.52 |

| | | | |
|---|---|---|---|
| TAC3 | -4.89 | 6.53E-03 | 3.16 |
| TAL1 | -6.34 | 3.85E-04 | 2.89 |
| TAL2 | 3.83 | 2.05E-02 | 4.59 |
| TBC1D7 | 2.48 | 1.31E-01 | 7.48 |
| TBCD | 2.78 | 8.72E-02 | 5.99 |
| TBCE | 5.75 | 3.48E-03 | 3.23 |
| TBR1 | -6.87 | 3.19E-07 | 6.15 |
| TBX1 | -4.16 | 4.54E-02 | 2.80 |
| TBX18 | -4.61 | 7.28E-03 | 3.60 |
| TBX20 | -9.18 | 1.26E-08 | -0.97 |
| TBX4 | -5.16 | 1.28E-03 | 3.88 |
| TBX5 | -5.14 | 1.34E-03 | 3.87 |
| TBX5-AS1 | -4.34 | 8.72E-02 | -0.97 |
| TCAF1 | -3.51 | 1.84E-02 | 5.17 |
| TCF21 | -7.77 | 2.52E-06 | -0.97 |
| TCN1 | 6.57 | 2.16E-05 | -0.97 |
| TCTEX1D1 | 3.61 | 8.44E-03 | 6.48 |
| TCTEX1D4 | 4.65 | 7.75E-03 | -0.97 |
| TDGF1 | -6.22 | 8.97E-05 | 3.83 |
| TDGF1P3 | -4.45 | 7.16E-02 | -0.97 |
| TEAD4 | -3.22 | 4.10E-02 | 4.79 |
| TECRL | -6.59 | 1.57E-04 | -0.97 |
| TEK | -5.95 | 1.25E-05 | 5.69 |
| TEKT1 | 4.30 | 1.29E-03 | 6.31 |
| TEKT4 | 5.65 | 5.01E-03 | 3.18 |
| TEX26 | 3.17 | 6.54E-02 | 4.75 |
| TEX29 | -3.54 | 1.88E-01 | 2.49 |
| TF | -3.94 | 2.66E-03 | 6.77 |
| TFEC | -3.06 | 9.68E-02 | 4.25 |
| TFPI2 | 2.47 | 1.32E-01 | 7.64 |
| TGFBR3 | -2.88 | 4.15E-02 | 9.23 |
| TH | 3.83 | 2.98E-03 | 8.10 |
| THBD | -5.33 | 4.50E-05 | 6.70 |
| THCAT155 | -2.53 | 1.57E-01 | 5.89 |
| THRA1/BTR | 4.27 | 2.19E-02 | -0.97 |
| THSD1 | -4.16 | 5.99E-03 | 4.38 |
| THSD7B | -3.06 | 3.09E-02 | 6.77 |
| TICRR | -3.19 | 2.33E-02 | 6.52 |
| TINAGL1 | -4.39 | 2.46E-02 | 2.91 |
| TJP2 | -2.27 | 1.95E-01 | 8.58 |
| TK1 | -2.72 | 9.59E-02 | 5.94 |
| TLE2 | -3.31 | 1.99E-02 | 6.28 |
| TLR4 | -4.39 | 2.46E-02 | 2.91 |
| TLX1 | -4.64 | 1.31E-02 | 3.03 |
| TM4SF18 | -3.14 | 7.59E-02 | 4.46 |
| TMC8 | -4.50 | 1.77E-02 | 2.96 |
| TMEM119 | -2.42 | 1.67E-01 | 6.54 |
| TMEM14C | 6.27 | 7.60E-06 | 5.49 |
| TMEM161B-AS1 | -2.38 | 1.81E-01 | 6.69 |
| TMEM232 | 3.22 | 3.04E-02 | 5.87 |
| TMEM235 | -2.83 | 1.58E-01 | 4.30 |
| TMEM30B | -3.53 | 3.12E-02 | 4.29 |
| TMEM37 | -3.18 | 3.01E-02 | 5.70 |
| TMEM40 | -4.96 | 2.19E-02 | -0.97 |
| TMEM72 | 3.66 | 2.26E-02 | 4.77 |
| TMEM88 | -3.41 | 1.15E-02 | 7.33 |
| TMEM92 | -4.86 | 3.27E-03 | 3.73 |
| TMIGD3 | 3.61 | 8.72E-02 | -0.97 |
| TMPRSS13 | -4.54 | 1.88E-01 | 1.99 |

| | | | |
|---|---|---|---|
| TMPRSS6 | -6.31 | 4.10E-04 | -0.97 |
| TNFRSF11A | 5.11 | 4.14E-04 | 4.91 |
| TNFRSF11B | 3.09 | 7.83E-02 | 4.71 |
| TNFRSF14 | 3.17 | 1.95E-01 | -0.97 |
| TNFRSF1B | -2.60 | 1.28E-01 | 5.71 |
| TNFSF15 | -6.42 | 2.84E-04 | -0.97 |
| TNK2-AS1 | 2.95 | 1.41E-01 | 4.41 |
| TNNI1 | -2.60 | 1.09E-01 | 6.80 |
| TNNT2 | -5.85 | 1.77E-03 | -0.97 |
| TNR | 4.49 | 1.07E-01 | 2.60 |
| TPBG | 4.06 | 1.97E-02 | 3.97 |
| TPH1 | 3.75 | 1.07E-01 | 3.23 |
| TPPP3 | 3.39 | 9.95E-03 | 9.55 |
| TRHDE-AS1 | -2.89 | 9.56E-02 | 4.86 |
| TRIM14 | -3.03 | 1.04E-01 | 4.23 |
| TRIM63 | 5.62 | 5.52E-04 | 4.16 |
| TRIML2 | -5.73 | 2.35E-03 | -0.97 |
| tRNA-Arg | -2.68 | 7.72E-02 | 7.77 |
| tRNA-Leu | -2.89 | 1.46E-01 | 4.16 |
| TROAP | -2.42 | 1.73E-01 | 6.42 |
| TRPC5 | 4.65 | 6.80E-02 | 2.68 |
| TRPM1 | 5.57 | 5.10E-04 | -0.97 |
| TRPM3 | 3.89 | 2.09E-03 | 8.88 |
| TRPM8 | 3.10 | 9.64E-02 | 4.49 |
| TSGA10IP | 3.61 | 8.72E-02 | -0.97 |
| TSHR | 4.72 | 9.19E-04 | 5.30 |
| TSNAXIP1 | 2.82 | 7.27E-02 | 6.09 |
| TSPAN10 | 6.38 | 4.94E-05 | 4.54 |
| TSPYL5 | -5.96 | 1.23E-03 | 2.70 |
| TSTA3 | -3.45 | 1.34E-01 | 3.02 |
| TTC16 | 3.75 | 7.16E-02 | -0.97 |
| TTC29 | 2.71 | 1.08E-01 | 5.71 |
| TTLL10 | 5.09 | 2.11E-03 | -0.97 |
| TTR | 2.90 | 3.71E-02 | 14.25 |
| TTTY15 | -5.39 | 7.00E-03 | -0.97 |
| TUBB1 | -6.13 | 7.50E-04 | 2.78 |
| TUBB2BP1 | 2.48 | 1.22E-01 | 8.34 |
| TUBBP5 | -3.37 | 1.72E-01 | 2.99 |
| TXLNB | 3.87 | 1.77E-02 | 4.61 |
| TYR | 10.35 | 1.81E-11 | 6.53 |
| TYRP1 | 7.22 | 1.29E-08 | 9.97 |
| TYW3 | -4.22 | 1.29E-01 | -0.97 |
| U2AF1L5 | -3.91 | 5.95E-03 | 5.48 |
| UBA7 | -3.78 | 1.49E-02 | 4.42 |
| UBAC2-AS1 | 3.45 | 9.70E-02 | 3.66 |
| UBXN10 | 2.79 | 6.80E-02 | 7.00 |
| UGT2B11 | -7.45 | 9.20E-06 | -0.97 |
| UGT2B4 | -4.54 | 6.01E-02 | -0.97 |
| UMODL1-AS1 | 4.57 | 8.47E-02 | 2.64 |
| UNC93A | -6.13 | 7.50E-04 | -0.97 |
| UPK3B | -4.89 | 2.55E-02 | -0.97 |
| URAD | -5.34 | 7.75E-03 | -0.97 |
| USP32P1 | -3.54 | 1.88E-01 | 2.49 |
| USP9Y | -6.10 | 8.10E-04 | 2.76 |
| UTS2B | 3.49 | 1.95E-01 | 3.10 |
| UTS2R | 3.31 | 8.11E-02 | 4.33 |
| UTY | -4.22 | 1.29E-01 | -0.97 |
| VGF | 3.52 | 8.36E-03 | 7.73 |
| VIL1 | -7.89 | 1.64E-06 | 3.66 |

| | | | |
|---|---|---|---|
| VIPR1 | -3.45 | 4.15E-02 | 4.02 |
| VIT | 4.19 | 3.64E-02 | 3.45 |
| VRTN | -5.54 | 4.19E-03 | -0.97 |
| VSIG1 | 5.55 | 6.63E-04 | 4.13 |
| VSIR | -2.85 | 8.43E-02 | 5.23 |
| VSNL1 | -3.45 | 8.25E-02 | 3.76 |
| VSTM2L | 2.30 | 1.79E-01 | 9.29 |
| VTCN1 | -3.50 | 3.32E-02 | 4.27 |
| VTN | -4.78 | 1.24E-04 | 8.19 |
| VWA3B | 6.05 | 4.36E-05 | 4.96 |
| VWCE | -3.00 | 4.36E-02 | 5.92 |
| VWDE | -2.58 | 1.95E-01 | 4.71 |
| WDR38 | 4.78 | 2.38E-03 | 4.33 |
| WDR46 | -5.34 | 7.75E-03 | -0.97 |
| WDR49 | 2.94 | 5.81E-02 | 6.07 |
| WDR63 | 2.98 | 4.03E-02 | 7.20 |
| WDR64 | 3.17 | 1.95E-01 | -0.97 |
| WDR66 | 2.55 | 1.32E-01 | 6.53 |
| WDR78 | 2.58 | 1.18E-01 | 6.73 |
| WDR93 | 3.61 | 6.54E-02 | 3.74 |
| WFIKKN2 | 5.88 | 3.46E-05 | 5.61 |
| WISP1 | -2.43 | 1.54E-01 | 7.16 |
| WNT1 | 3.47 | 8.61E-03 | 8.50 |
| WNT10B | 3.17 | 2.19E-02 | 7.52 |
| WNT2 | -4.54 | 6.01E-02 | -0.97 |
| WNT2B | 3.85 | 2.52E-03 | 8.56 |
| WNT3A | 2.84 | 6.12E-02 | 6.86 |
| WNT4 | 2.70 | 1.14E-01 | 5.87 |
| WNT6 | -4.45 | 7.16E-02 | -0.97 |
| WNT7A | -2.78 | 9.21E-02 | 5.75 |
| WSCD2 | 2.93 | 4.36E-02 | 7.37 |
| XK | 2.59 | 1.16E-01 | 6.86 |
| XRCC2 | -2.59 | 1.32E-01 | 5.82 |
| YJEFN3 | -2.99 | 6.12E-02 | 5.46 |
| ZBED1 | 4.12 | 3.02E-02 | -0.97 |
| ZC3HAV1L | -2.57 | 1.49E-01 | 5.46 |
| ZFP42 | -4.59 | 6.68E-04 | 5.60 |
| ZFPM2 | -3.23 | 1.85E-02 | 7.16 |
| ZG16 | -4.22 | 1.29E-01 | -0.97 |
| ZGRF1 | -3.36 | 1.65E-02 | 6.20 |
| ZIC1 | 2.32 | 1.67E-01 | 11.95 |
| ZIC4 | 3.52 | 8.45E-03 | 7.64 |
| ZMYND10 | 4.83 | 2.01E-04 | 6.85 |
| ZNF295-AS1 | 4.27 | 1.72E-01 | 2.49 |
| ZNF385B | -4.77 | 9.18E-03 | 3.10 |
| ZNF396 | 3.24 | 9.04E-02 | 3.98 |
| ZNF436-AS1 | 3.95 | 4.20E-02 | -0.97 |
| ZNF474 | 3.02 | 4.24E-02 | 6.39 |
| ZNF528-AS1 | -5.16 | 1.26E-02 | -0.97 |
| ZNF560 | -4.64 | 4.99E-02 | -0.97 |
| ZNF578 | -3.96 | 1.04E-02 | 4.28 |
| ZNF648 | 2.72 | 1.88E-01 | 4.52 |
| ZNF732 | -3.01 | 5.69E-02 | 5.31 |
| ZNF90 | -3.05 | 3.35E-02 | 6.63 |
| ZSCAN10 | -3.73 | 1.97E-02 | 4.16 |
| ZYX | 4.63 | 1.14E-02 | 3.67 |
