## Supplemental Table 2 for "Brainstem organoids from human pluripotent stem cells contain neural crest population"

**Supplemental Table 2  Differentially expressed genes between hBSOs and hESCs**

| Gene_name | log (hBSO/hESC) | FDR (hBSO/hESC) | A (hBSO/hESC) |
|---|---|---|---|
| A2M | 7.51 | 1.89E-01 | -1.00 |
| ABCB5 | 8.59 | 1.18E-01 | -1.00 |
| ADCYAP1 | 7.77 | 1.65E-01 | -1.00 |
| AKAP6 | 7.33 | 1.92E-01 | 7.66 |
| ALX4 | 8.89 | 1.41E-01 | 4.53 |
| AMER2 | 7.40 | 1.89E-01 | 6.60 |
| AP1M2 | -10.88 | 6.73E-02 | 5.36 |
| APELA | -10.04 | 9.87E-02 | 4.93 |
| AQP1 | 8.76 | 1.25E-01 | 5.47 |
| ARMC3 | 8.43 | 1.25E-01 | -1.00 |
| ASCL1 | 11.18 | 6.73E-02 | 5.68 |
| ASIC4 | 7.51 | 1.89E-01 | -1.00 |
| ASPA | 7.73 | 1.65E-01 | -1.00 |
| BARHL1 | 8.32 | 1.28E-01 | -1.00 |
| BARHL2 | 8.53 | 1.19E-01 | -1.00 |
| BMP5 | 8.83 | 1.08E-01 | -1.00 |
| C11orf88 | 8.33 | 1.28E-01 | -1.00 |
| CABP7 | 9.41 | 1.18E-01 | 4.79 |
| CAPN3 | 8.02 | 1.77E-01 | 5.10 |
| CAPSL | 7.94 | 1.50E-01 | -1.00 |
| CCNO | 9.22 | 8.61E-02 | -1.00 |
| CDC20B | 11.61 | 6.73E-02 | 5.89 |
| CDCP1 | -7.78 | 1.95E-01 | 5.39 |
| CDH18 | 7.65 | 1.82E-01 | 5.91 |
| CDK15 | 7.42 | 1.95E-01 | -1.00 |
| CELF6 | 7.76 | 1.65E-01 | -1.00 |
| CFAP126 | 9.34 | 8.17E-02 | -1.00 |
| CFAP206 | 8.17 | 1.32E-01 | -1.00 |
| CFAP47 | 8.05 | 1.40E-01 | -1.00 |
| CFAP52 | 8.54 | 1.19E-01 | -1.00 |
| CFAP54 | 7.42 | 1.95E-01 | -1.00 |
| CFAP61 | 8.13 | 1.35E-01 | -1.00 |
| CFI | 7.84 | 1.60E-01 | -1.00 |
| CHL1 | 10.40 | 6.73E-02 | 6.29 |
| CHST4 | -7.92 | 1.65E-01 | -1.00 |
| CLUL1 | 7.99 | 1.79E-01 | 5.09 |
| CNPY1 | 9.03 | 9.75E-02 | -1.00 |
| CNTNAP2 | 8.37 | 1.25E-01 | -1.00 |
| COL3A1 | 11.90 | 5.99E-02 | 7.62 |
| CPED1 | 8.15 | 1.35E-01 | -1.00 |
| CRYAB | 9.19 | 1.08E-01 | 5.68 |
| CRYBG1 | -8.68 | 1.35E-01 | 5.25 |

| | | | |
|---|---|---|---|
| D21S2088E | -9.79 | 1.11E-01 | 4.81 |
| DACH1 | 8.23 | 1.29E-01 | 6.79 |
| DCN | 9.87 | 6.73E-02 | -1.00 |
| DCT | 16.02 | 6.49E-03 | 8.10 |
| DDC | 8.66 | 1.16E-01 | -1.00 |
| DIO3 | 8.54 | 1.71E-01 | 4.36 |
| DLK1 | 10.94 | 6.36E-02 | 9.56 |
| DMRTA2 | 8.02 | 1.42E-01 | -1.00 |
| DNMT3B | -7.25 | 1.89E-01 | 11.73 |
| DRC1 | 9.66 | 1.08E-01 | 4.92 |
| DRGX | 8.98 | 1.01E-01 | -1.00 |
| EBF1 | 9.18 | 1.08E-01 | 5.68 |
| EBF2 | 9.85 | 6.73E-02 | -1.00 |
| EBF3 | 8.48 | 1.29E-01 | 5.91 |
| EDAR | 7.93 | 1.51E-01 | -1.00 |
| EFCAB1 | 9.22 | 8.61E-02 | -1.00 |
| ELAVL4 | 10.71 | 6.73E-02 | 5.45 |
| EMP1 | 8.99 | 1.02E-01 | 7.58 |
| EMX2 | 9.61 | 7.82E-02 | -1.00 |
| EMX2OS | 9.79 | 6.79E-02 | -1.00 |
| EN1 | 9.64 | 7.57E-02 | -1.00 |
| EN2 | 10.91 | 6.73E-02 | 5.54 |
| ENKUR | 8.47 | 1.79E-01 | 4.33 |
| ERBB4 | 8.58 | 1.32E-01 | 5.38 |
| ESM1 | 9.36 | 8.17E-02 | -1.00 |
| ESRG | -13.27 | 3.19E-02 | 7.55 |
| FAM198B | 8.37 | 1.25E-01 | -1.00 |
| FAM216B | 9.28 | 8.30E-02 | -1.00 |
| FAM81B | 8.20 | 1.32E-01 | -1.00 |
| FEZF2 | 8.14 | 1.35E-01 | -1.00 |
| FLT1 | -10.91 | 6.73E-02 | 6.37 |
| FOXC1 | 9.51 | 7.98E-02 | -1.00 |
| FZD10-AS1 | 7.92 | 1.51E-01 | -1.00 |
| GDF10 | 7.99 | 1.79E-01 | 5.09 |
| GDF3 | -7.64 | 1.91E-01 | -1.00 |
| GDF5 | 7.46 | 1.92E-01 | -1.00 |
| GDF7 | 12.21 | 5.99E-02 | 6.20 |
| GLIS3 | 9.63 | 1.11E-01 | 4.90 |
| GPNMB | 8.08 | 1.32E-01 | 7.83 |
| GREB1L | 8.03 | 1.58E-01 | 5.69 |
| GRIA2 | 7.53 | 1.89E-01 | -1.00 |
| HES3 | -7.90 | 1.67E-01 | -1.00 |
| HES4 | 7.49 | 1.89E-01 | 6.16 |
| HMGCLL1 | 8.36 | 1.25E-01 | -1.00 |
| HSF4 | 8.12 | 1.35E-01 | -1.00 |

| | | | |
|---|---|---|---|
| IDO1 | -9.61 | 8.17E-02 | -1.00 |
| IGFBP5 | 10.78 | 6.59E-02 | 9.80 |
| INSM1 | 9.95 | 9.17E-02 | 5.06 |
| ISL1 | 9.96 | 6.73E-02 | -1.00 |
| ISL2 | 7.74 | 1.65E-01 | -1.00 |
| ISLR2 | 9.59 | 7.98E-02 | 6.88 |
| ITGA8 | 9.01 | 1.34E-01 | 4.59 |
| KCNJ13 | 9.57 | 7.91E-02 | -1.00 |
| KLKB1 | -8.12 | 1.50E-01 | -1.00 |
| KLRG2 | -8.80 | 1.65E-01 | 4.32 |
| L1TD1 | -9.49 | 7.15E-02 | 10.21 |
| LAD1 | -9.14 | 1.02E-01 | -1.00 |
| LCK | -10.32 | 8.30E-02 | 5.08 |
| LGI1 | 9.41 | 1.18E-01 | 4.80 |
| LHX9 | 8.81 | 1.08E-01 | -1.00 |
| LIN28A | -7.71 | 1.50E-01 | 11.10 |
| LINC00428 | -11.20 | 6.73E-02 | 5.52 |
| LINC00473 | 8.92 | 1.05E-01 | -1.00 |
| LINC00643 | 9.29 | 8.30E-02 | -1.00 |
| LINC00678 | -11.60 | 5.99E-02 | -1.00 |
| LINC00982 | 8.77 | 1.51E-01 | 4.48 |
| LINC02154 | 7.51 | 1.89E-01 | -1.00 |
| LIX1 | 10.61 | 6.73E-02 | -1.00 |
| LMX1A | 11.51 | 5.99E-02 | -1.00 |
| LMX1B | 9.05 | 1.16E-01 | 5.61 |
| LNCPRESS1 | -9.26 | 9.21E-02 | -1.00 |
| LNCPRESS2 | -10.88 | 6.73E-02 | -1.00 |
| LOC100126447 | -9.34 | 8.99E-02 | -1.00 |
| LOC100130370 | 7.48 | 1.89E-01 | -1.00 |
| LOC101927120 | 8.44 | 1.25E-01 | -1.00 |
| LOC101928817 | 7.68 | 1.71E-01 | -1.00 |
| LOC105369436 | -10.79 | 6.73E-02 | -1.00 |
| LOC105370482 | -9.35 | 8.93E-02 | -1.00 |
| LOC105370767 | 7.92 | 1.51E-01 | -1.00 |
| LOC105373409 | -8.63 | 1.25E-01 | -1.00 |
| LOC105374794 | 12.08 | 5.79E-02 | -1.00 |
| LOC105374972 | 9.04 | 9.67E-02 | -1.00 |
| LOC105378853 | 8.74 | 1.13E-01 | -1.00 |
| LOC105379041 | 7.69 | 1.71E-01 | -1.00 |
| LOC105379506 | 8.23 | 1.32E-01 | -1.00 |
| LOC107986523 | 7.52 | 1.89E-01 | -1.00 |
| LOC107987188 | 9.98 | 9.07E-02 | 5.08 |
| LOC109864269 | -11.67 | 6.36E-02 | 6.75 |
| LOC112268449 | 7.70 | 1.70E-01 | -1.00 |
| LOC440982 | 7.61 | 1.79E-01 | -1.00 |

| | | | |
|---|---|---|---|
| LOC441204 | 8.19 | 1.32E-01 | -1.00 |
| LOC642366 | 8.89 | 1.40E-01 | 4.54 |
| LOC653513 | 7.75 | 1.65E-01 | -1.00 |
| LRP1B | 8.70 | 1.16E-01 | -1.00 |
| LRRTM2 | 8.80 | 1.08E-01 | -1.00 |
| LUM | 10.08 | 6.73E-02 | -1.00 |
| MAB21L1 | 9.29 | 8.30E-02 | -1.00 |
| MAB21L2 | 9.41 | 8.17E-02 | -1.00 |
| MAP1LC3C | 7.25 | 1.96E-01 | 7.80 |
| MAP3K19 | 8.88 | 1.07E-01 | -1.00 |
| MEIS2 | 10.20 | 6.73E-02 | -1.00 |
| MEOX1 | 8.98 | 1.01E-01 | -1.00 |
| MGP | 8.69 | 1.16E-01 | -1.00 |
| MIR100HG | 8.42 | 1.25E-01 | 6.62 |
| MIR3606 | 7.86 | 1.57E-01 | -1.00 |
| MIR99AHG | 8.22 | 1.32E-01 | -1.00 |
| MLANA | 12.53 | 3.69E-02 | -1.00 |
| MMRN1 | 11.35 | 5.99E-02 | -1.00 |
| MS4A6A | 8.28 | 1.29E-01 | -1.00 |
| MSX1 | 8.47 | 1.29E-01 | 5.91 |
| MSX2 | 8.32 | 1.32E-01 | 6.25 |
| MUSK | 7.63 | 1.78E-01 | -1.00 |
| MYH7 | 7.86 | 1.57E-01 | -1.00 |
| MYO16 | 9.23 | 1.25E-01 | 4.71 |
| MYT1L | 7.76 | 1.65E-01 | -1.00 |
| NANOG | -10.38 | 6.73E-02 | -1.00 |
| NANOGP8 | -9.53 | 8.17E-02 | -1.00 |
| NDST4 | 9.35 | 1.22E-01 | 4.77 |
| NEUROD4 | 8.38 | 1.25E-01 | -1.00 |
| NKX1-2 | -7.95 | 1.68E-01 | 5.89 |
| NR2F1 | 10.93 | 6.73E-02 | -1.00 |
| NR2F2 | 10.51 | 6.73E-02 | -1.00 |
| NRN1 | 10.87 | 6.73E-02 | -1.00 |
| NSG2 | 8.49 | 1.18E-01 | 7.14 |
| NTRK2 | 10.22 | 8.17E-02 | 5.20 |
| OC90 | 7.71 | 1.68E-01 | -1.00 |
| OTX2-AS1 | 7.61 | 1.80E-01 | -1.00 |
| P2RX3 | 11.14 | 6.73E-02 | 5.66 |
| PALMD | 8.17 | 1.33E-01 | -1.00 |
| PAPPA2 | 8.31 | 1.25E-01 | 7.83 |
| PAX3 | 9.60 | 7.86E-02 | -1.00 |
| PAX6 | 11.33 | 5.99E-02 | -1.00 |
| PDE7B | 8.11 | 1.35E-01 | -1.00 |
| PDZRN4 | 9.52 | 7.98E-02 | -1.00 |
| PGM5 | 8.25 | 1.96E-01 | 4.21 |

| | | | |
|---|---|---|---|
| PIRT | 8.00 | 1.44E-01 | -1.00 |
| PODXL | -7.82 | 1.40E-01 | 11.22 |
| POU3F3 | 7.90 | 1.60E-01 | 6.04 |
| POU4F1 | 8.43 | 1.25E-01 | 6.30 |
| POU5F1 | -10.25 | 6.73E-02 | -1.00 |
| POU5F1P3 | -10.61 | 6.73E-02 | -1.00 |
| POU5F1P4 | -8.29 | 1.35E-01 | -1.00 |
| PRDM14 | -8.69 | 1.18E-01 | 6.84 |
| PRDM16 | 7.48 | 1.89E-01 | 6.15 |
| PRPH | 8.94 | 1.02E-01 | 7.73 |
| PRRX1 | 8.21 | 1.32E-01 | -1.00 |
| PRSS56 | 8.52 | 1.20E-01 | -1.00 |
| PTGDS | 8.25 | 1.32E-01 | 7.38 |
| PTPRC | 8.21 | 1.32E-01 | -1.00 |
| RARB | 9.44 | 1.18E-01 | 4.81 |
| RFX4 | 9.13 | 1.29E-01 | 4.66 |
| RPE65 | 10.07 | 6.73E-02 | -1.00 |
| RPH3A | 8.11 | 1.35E-01 | -1.00 |
| RSPO1 | 10.72 | 6.73E-02 | 5.45 |
| RSPO3 | 8.71 | 1.28E-01 | 5.44 |
| RTL1 | 10.21 | 6.73E-02 | -1.00 |
| RTN4RL1 | 9.29 | 8.30E-02 | -1.00 |
| RXRG | 9.75 | 6.96E-02 | -1.00 |
| SAMD11 | 9.85 | 6.73E-02 | -1.00 |
| SAMD5 | 8.04 | 1.57E-01 | 5.69 |
| SCARA5 | 8.13 | 1.65E-01 | 5.16 |
| SCG2 | 7.75 | 1.65E-01 | 7.13 |
| SCGN | 9.76 | 6.96E-02 | -1.00 |
| SEMA3C | 8.94 | 1.05E-01 | 7.56 |
| SEZ6L | 7.62 | 1.68E-01 | 6.71 |
| SLC16A6 | 8.47 | 1.79E-01 | 4.32 |
| SLC45A2 | 8.53 | 1.19E-01 | -1.00 |
| SLC4A5 | 9.34 | 8.17E-02 | 8.08 |
| SLITRK6 | 7.73 | 1.86E-01 | 5.54 |
| SOX6 | 9.05 | 1.16E-01 | 5.62 |
| SPAG6 | 9.29 | 1.05E-01 | 5.73 |
| SST | 9.89 | 6.73E-02 | 7.04 |
| STMN2 | 9.25 | 8.17E-02 | 8.72 |
| STMN4 | 9.25 | 1.07E-01 | 5.71 |
| SV2C | 7.42 | 1.89E-01 | 6.61 |
| TBX2 | 8.10 | 1.36E-01 | -1.00 |
| TBX3 | 7.70 | 1.89E-01 | 5.52 |
| TCIM | 8.61 | 1.65E-01 | 4.40 |
| TDGF1 | -12.54 | 5.99E-02 | 7.19 |
| TDGF1P3 | -10.17 | 6.73E-02 | -1.00 |

| | | | |
|---|---|---|---|
| TEKT1 | 8.57 | 1.18E-01 | -1.00 |
| TERT | -8.94 | 1.53E-01 | 4.38 |
| TFAP2A | 9.69 | 7.27E-02 | -1.00 |
| TFAP2B | 9.57 | 7.91E-02 | -1.00 |
| TH | 10.13 | 6.73E-02 | -1.00 |
| TNFRSF8 | -8.54 | 1.25E-01 | -1.00 |
| TPPP3 | 8.76 | 1.08E-01 | 7.06 |
| TRH | 11.52 | 5.99E-02 | -1.00 |
| TRIM67 | 9.15 | 1.29E-01 | 4.67 |
| TRIML2 | -9.73 | 7.93E-02 | -1.00 |
| TSHR | 7.77 | 1.65E-01 | -1.00 |
| TSHZ1 | 8.06 | 1.56E-01 | 5.71 |
| TTR | 15.81 | 6.49E-03 | 8.00 |
| TYR | 11.82 | 5.99E-02 | -1.00 |
| TYRP1 | 13.71 | 2.97E-02 | 6.95 |
| VRTN | -12.06 | 5.99E-02 | -1.00 |
| VSNL1 | -8.28 | 1.36E-01 | 6.38 |
| VWA3B | 8.10 | 1.36E-01 | -1.00 |
| WLS | 7.66 | 1.57E-01 | 8.92 |
| WNT1 | 10.34 | 6.73E-02 | -1.00 |
| WNT10B | 8.84 | 1.08E-01 | -1.00 |
| WNT3A | 8.39 | 1.25E-01 | -1.00 |
| WNT4 | 7.76 | 1.82E-01 | 5.55 |
| ZEB2 | 7.98 | 1.40E-01 | 6.67 |
| ZFHX4-AS1 | 9.02 | 9.78E-02 | -1.00 |
| ZFP42 | -8.91 | 9.75E-02 | 7.54 |
| ZIC1 | 10.05 | 6.73E-02 | 8.28 |
| ZIC4 | 11.24 | 6.11E-02 | -1.00 |
| ZNF503 | 10.01 | 6.73E-02 | -1.00 |
| ZSCAN10 | -9.39 | 8.30E-02 | 7.20 |
