## Supplemental Table 3 for "Brainstem organoids from human pluripotent stem cells contain neural crest population"

**Supplemental Table 3  Differentially expressed genes between hCOs and hESCs**

| Gene_name | log (hCO/hESC) | FDR (hCO/hESC) | A (hCO/hESC) |
|---|---|---|---|
| A1CF | 6.83 | 1.67E-01 | 3.19 |
| A2M | 11.09 | 3.47E-03 | -0.98 |
| AATF | -7.44 | 3.98E-02 | -0.98 |
| ABCA8 | 6.62 | 1.07E-01 | -0.98 |
| ABI3BP | 6.45 | 1.51E-01 | 4.00 |
| ACKR3 | 5.22 | 1.84E-01 | 6.84 |
| ACOXL | -6.88 | 1.09E-01 | 3.71 |
| ACSL5 | 6.30 | 1.42E-01 | -0.98 |
| ACVRL1 | 6.54 | 1.15E-01 | -0.98 |
| ADAM12 | 5.64 | 1.38E-01 | 6.68 |
| ADAMTS18 | 9.58 | 1.79E-02 | 4.56 |
| ADAMTS9 | 7.21 | 4.10E-02 | 7.63 |
| ADAMTS9-AS2 | 6.47 | 1.21E-01 | -0.98 |
| ADCYAP1 | 7.65 | 4.89E-02 | -0.98 |
| ADGRD1 | 6.40 | 9.91E-02 | 4.56 |
| ADGRF5 | 6.64 | 1.96E-01 | 3.10 |
| ADGRG1 | 6.18 | 8.63E-02 | 7.91 |
| ADGRG3 | 5.51 | 1.51E-01 | 6.53 |
| ADH4 | 6.62 | 1.07E-01 | -0.98 |
| AFP | 10.74 | 2.93E-03 | 10.95 |
| AGT | 9.82 | 1.54E-02 | 4.68 |
| AGTR1 | 8.33 | 5.26E-02 | 3.94 |
| AHNAK | 6.89 | 4.64E-02 | 10.56 |
| AHSG | 10.78 | 3.47E-03 | 6.75 |
| AKAP6 | 6.19 | 9.10E-02 | 6.04 |
| AKR1B10 | 8.13 | 3.42E-02 | -0.98 |
| AKR1C2 | 7.03 | 1.46E-01 | 3.29 |
| AKR1D1 | 7.85 | 4.26E-02 | -0.98 |
| ALB | 7.82 | 7.87E-02 | 3.69 |
| ALDH1A1 | 6.29 | 8.61E-02 | 6.50 |
| ALPK2 | 6.18 | 1.08E-01 | 5.45 |
| ALX4 | 9.33 | 2.42E-02 | 4.44 |
| AMBN | 6.78 | 9.54E-02 | -0.98 |
| AMER2 | 6.63 | 7.75E-02 | 5.89 |
| ANGPT2 | 6.85 | 8.99E-02 | -0.98 |
| ANKFN1 | 6.22 | 1.49E-01 | -0.98 |
| ANKRD1 | 6.74 | 1.81E-01 | 3.14 |
| ANXA1 | 6.60 | 7.89E-02 | 5.66 |
| ANXA8 | 8.96 | 1.61E-02 | -0.98 |
| ANXA8L1 | 7.73 | 4.52E-02 | -0.98 |
| APOA1 | 11.63 | 2.60E-03 | 7.18 |
| APOA2 | 11.66 | 4.25E-03 | 5.60 |
| APOA4 | 11.32 | 3.00E-03 | -0.98 |

| | | | |
|---|---|---|---|
| APOB | 7.36 | 3.35E-02 | 9.83 |
| APOC3 | 11.88 | 2.60E-03 | -0.98 |
| APOH | 6.90 | 8.63E-02 | -0.98 |
| APOLD1 | 6.18 | 1.17E-01 | 4.45 |
| AQP1 | 11.57 | 3.00E-03 | 6.56 |
| ARID5B | 5.21 | 1.93E-01 | 6.55 |
| ARX | 8.38 | 1.99E-02 | 5.55 |
| ASCL1 | 8.12 | 6.10E-02 | 3.83 |
| ASXL3 | 5.22 | 1.87E-01 | 6.39 |
| ATP10A | -5.74 | 1.80E-01 | 4.14 |
| ATP6V0D2 | 6.56 | 1.15E-01 | -0.98 |
| B3GALT2 | 6.64 | 1.06E-01 | -0.98 |
| BAHCC1 | 7.24 | 4.13E-02 | 6.56 |
| BARHL1 | 7.70 | 4.66E-02 | -0.98 |
| BARHL2 | 7.42 | 6.02E-02 | -0.98 |
| BCO2 | 6.63 | 1.33E-01 | 4.09 |
| BMP5 | 9.60 | 1.19E-02 | -0.98 |
| BNC1 | 6.51 | 7.44E-02 | 6.73 |
| BRINP3 | 7.30 | 6.24E-02 | -0.98 |
| C11orf96 | 5.47 | 1.96E-01 | 4.83 |
| C13orf42 | -6.68 | 8.98E-02 | 5.19 |
| C1orf61 | 5.88 | 1.17E-01 | 5.88 |
| C1orf94 | -7.32 | 7.87E-02 | 3.93 |
| C1QTNF7 | 6.66 | 1.04E-01 | -0.98 |
| C20orf203 | 6.45 | 1.27E-01 | -0.98 |
| C5orf66-AS1 | 6.52 | 1.17E-01 | -0.98 |
| CABP7 | 7.38 | 1.09E-01 | 3.46 |
| CAMK2A | 6.11 | 1.20E-01 | 4.83 |
| CBLN2 | 6.72 | 1.84E-01 | 3.14 |
| CCDC33 | 5.95 | 1.84E-01 | -0.98 |
| CCDC68 | 5.58 | 1.90E-01 | 4.15 |
| CD300E | 6.45 | 1.27E-01 | -0.98 |
| CD93 | 8.50 | 4.52E-02 | 4.03 |
| CDC20B | 6.72 | 1.84E-01 | 3.14 |
| CDH10 | 5.96 | 1.42E-01 | 4.34 |
| CDH12 | 7.12 | 7.51E-02 | -0.98 |
| CDH19 | 7.60 | 5.06E-02 | -0.98 |
| CDH20 | 7.27 | 1.18E-01 | 3.41 |
| CDX2 | 7.19 | 7.03E-02 | -0.98 |
| CELF3 | 6.44 | 7.86E-02 | 6.70 |
| CELF6 | 7.19 | 7.03E-02 | -0.98 |
| CHCHD10 | -5.79 | 1.70E-01 | 4.16 |
| CHL1 | 8.08 | 4.32E-02 | 4.82 |
| CHORDC1 | -5.79 | 1.83E-01 | 4.75 |
| CHRM2 | 6.08 | 1.64E-01 | -0.98 |
| CLIP4 | 5.95 | 1.45E-01 | 4.34 |

| | | | |
|---|---|---|---|
| CNGB1 | 6.89 | 8.80E-02 | -0.98 |
| CNMD | -6.37 | 7.81E-02 | 7.45 |
| CNTFR | 6.20 | 8.65E-02 | 6.87 |
| CNTN5 | 8.55 | 2.43E-02 | -0.98 |
| CNTN6 | 7.36 | 6.10E-02 | -0.98 |
| CNTNAP2 | 5.89 | 1.92E-01 | -0.98 |
| CNTNAP4 | 7.30 | 7.87E-02 | 4.42 |
| COL15A1 | 8.89 | 1.32E-02 | 6.80 |
| COL22A1 | 5.85 | 1.42E-01 | 5.50 |
| COL3A1 | 15.68 | 1.13E-04 | 9.20 |
| COL6A3 | 8.88 | 1.23E-02 | 7.67 |
| COL6A6 | 6.68 | 1.90E-01 | 3.12 |
| COPG2IT1 | 6.99 | 4.97E-02 | 6.44 |
| CPED1 | 9.59 | 1.19E-02 | -0.98 |
| CR2 | -6.79 | 1.16E-01 | 3.66 |
| CRB1 | 7.54 | 9.54E-02 | 3.54 |
| CREB5 | 7.30 | 6.24E-02 | -0.98 |
| CRHBP | 7.64 | 4.93E-02 | -0.98 |
| CSF3R | 6.95 | 8.36E-02 | -0.98 |
| CSNK2B | -5.36 | 1.97E-01 | -0.98 |
| CUBN | 7.02 | 9.54E-02 | 4.29 |
| CUZD1 | -5.74 | 1.89E-01 | 4.72 |
| CXCL5 | -5.67 | 1.74E-01 | 5.10 |
| CXCL6 | -5.70 | 1.87E-01 | 4.12 |
| CXorf36 | 6.42 | 1.29E-01 | -0.98 |
| CYP26B1 | 6.74 | 1.81E-01 | 3.14 |
| CYSLTR2 | 7.00 | 8.08E-02 | -0.98 |
| D21S2088E | -5.54 | 1.49E-01 | 6.62 |
| DACH1 | 7.31 | 4.46E-02 | 6.01 |
| DCC | 6.05 | 1.03E-01 | 6.38 |
| DCLK2 | 6.70 | 7.75E-02 | 5.12 |
| DCN | 13.06 | 1.48E-03 | -0.98 |
| DCT | 7.49 | 9.91E-02 | 3.52 |
| DCX | 5.35 | 1.55E-01 | 10.02 |
| DDC | 6.95 | 8.36E-02 | -0.98 |
| DDR2 | 8.07 | 4.32E-02 | 4.81 |
| DKK2 | 5.60 | 1.49E-01 | 6.04 |
| DLK1 | 8.53 | 1.41E-02 | 8.04 |
| DLX1 | 7.50 | 5.63E-02 | -0.98 |
| DLX5 | 5.92 | 1.87E-01 | -0.98 |
| DMRT3 | 7.26 | 8.08E-02 | 4.41 |
| DMRTA2 | 9.59 | 1.19E-02 | -0.98 |
| DNAH2 | 7.08 | 9.10E-02 | 4.31 |
| DNM3OS | 9.35 | 1.31E-02 | -0.98 |
| DNMT3B | -5.50 | 1.42E-01 | 11.57 |
| DPF3 | 6.38 | 1.33E-01 | -0.98 |

| | | | |
|---|---|---|---|
| DRC1 | 6.66 | 1.93E-01 | 3.11 |
| DRGX | 6.89 | 8.80E-02 | -0.98 |
| DSCAML1 | 6.58 | 1.13E-01 | -0.98 |
| EBF1 | 8.31 | 3.56E-02 | 4.93 |
| EBF2 | 9.37 | 1.29E-02 | -0.98 |
| EBF3 | 8.49 | 1.79E-02 | 5.61 |
| EBF4 | 5.23 | 1.98E-01 | 5.71 |
| EGFLAM | 7.06 | 9.31E-02 | 4.30 |
| ELAVL3 | 5.37 | 1.57E-01 | 7.85 |
| ELAVL4 | 10.45 | 1.07E-02 | 5.00 |
| EMX2 | 10.48 | 5.41E-03 | -0.98 |
| EMX2OS | 9.74 | 1.07E-02 | -0.98 |
| ENPP2 | 5.33 | 1.67E-01 | 6.96 |
| EOMES | 8.39 | 4.99E-02 | 3.97 |
| EPHA3 | 7.46 | 3.51E-02 | 6.67 |
| EPHA5 | 7.40 | 4.52E-02 | 5.06 |
| ERBB4 | 8.56 | 3.10E-02 | 5.05 |
| ERVMER34-1 | -5.35 | 1.71E-01 | 6.85 |
| ESAM | 6.70 | 1.87E-01 | 3.13 |
| ESM1 | 7.11 | 7.63E-02 | -0.98 |
| ESRG | -6.51 | 6.15E-02 | 10.61 |
| EXD3 | 6.14 | 1.96E-01 | 3.84 |
| F2 | 6.80 | 1.70E-01 | 3.17 |
| F2RL2 | 7.39 | 1.07E-01 | 3.47 |
| FABP1 | 9.93 | 9.93E-03 | -0.98 |
| FAM107A | 7.01 | 6.10E-02 | 4.87 |
| FAM198B | 9.11 | 1.52E-02 | -0.98 |
| FAM24B-CUZD1 | -5.74 | 1.89E-01 | 4.72 |
| FAP | 7.08 | 7.75E-02 | -0.98 |
| FETUB | 6.64 | 1.06E-01 | -0.98 |
| FEZF2 | 9.57 | 1.19E-02 | -0.98 |
| FGA | 11.77 | 2.60E-03 | -0.98 |
| FGB | 12.36 | 2.32E-03 | -0.98 |
| FGF9 | 6.90 | 8.63E-02 | -0.98 |
| FGG | 11.37 | 3.00E-03 | -0.98 |
| FIBIN | 8.33 | 3.10E-02 | -0.98 |
| FILIP1L | 8.04 | 6.41E-02 | 3.79 |
| FLRT2 | 6.85 | 8.99E-02 | -0.98 |
| FLRT3 | 6.25 | 8.45E-02 | 7.22 |
| FMO1 | 7.55 | 9.44E-02 | 3.55 |
| FMOD | 8.35 | 2.06E-02 | 5.53 |
| FNDC7 | -5.85 | 1.38E-01 | -0.98 |
| FOXC1 | 8.57 | 2.42E-02 | -0.98 |
| FOXF1 | 7.77 | 4.41E-02 | -0.98 |
| FOXG1 | 9.40 | 1.48E-02 | 5.47 |
| FOXP2 | 6.90 | 6.70E-02 | 4.81 |

| | | | |
|---|---|---|---|
| FRZB | 5.75 | 1.15E-01 | 9.33 |
| FUT5 | 6.54 | 1.15E-01 | -0.98 |
| FYB2 | 7.57 | 9.29E-02 | 3.56 |
| FZD10-AS1 | 6.80 | 9.29E-02 | -0.98 |
| G0S2 | 7.41 | 4.21E-02 | 6.07 |
| GABRB1 | 6.34 | 1.65E-01 | 3.94 |
| GAL | -5.52 | 1.53E-01 | 6.49 |
| GALNT5 | 6.74 | 1.81E-01 | 3.14 |
| GATA2 | 7.31 | 1.15E-01 | 3.43 |
| GATA3 | 7.48 | 9.97E-02 | 3.52 |
| GATA5 | 8.44 | 2.74E-02 | -0.98 |
| GATA6 | 10.05 | 8.53E-03 | -0.98 |
| GATA6-AS1 | 8.53 | 2.46E-02 | -0.98 |
| GBP2 | 6.14 | 1.58E-01 | -0.98 |
| GBP4 | 6.02 | 1.75E-01 | -0.98 |
| GDF10 | 6.33 | 1.67E-01 | 3.94 |
| GDF7 | 10.73 | 9.47E-03 | 5.14 |
| GGT5 | 6.56 | 1.15E-01 | -0.98 |
| GJB1 | 6.40 | 1.31E-01 | -0.98 |
| GLB1L3 | -8.27 | 3.71E-02 | 4.40 |
| GLIS3 | 7.08 | 1.41E-01 | 3.31 |
| GNRH2 | 5.66 | 1.78E-01 | 4.19 |
| GPM6A | 6.10 | 8.87E-02 | 7.95 |
| GPR176 | -6.17 | 8.36E-02 | 8.94 |
| GREB1L | 7.72 | 3.56E-02 | 5.22 |
| GREM2 | 7.55 | 6.24E-02 | 4.55 |
| GRIA2 | 9.88 | 1.05E-02 | -0.98 |
| GRIN3A | 6.99 | 1.50E-01 | 3.27 |
| GRM7 | 5.92 | 1.87E-01 | -0.98 |
| GSTA1 | 9.09 | 1.54E-02 | -0.98 |
| GSTA2 | 9.14 | 1.48E-02 | -0.98 |
| GUCY1A1 | 9.02 | 1.55E-02 | -0.98 |
| H19 | 9.09 | 1.19E-02 | 7.32 |
| HAND1 | 11.54 | 2.93E-03 | -0.98 |
| HAND2 | 10.20 | 6.98E-03 | -0.98 |
| HAND2-AS1 | 9.57 | 1.19E-02 | -0.98 |
| HAPLN1 | 5.99 | 1.02E-01 | 6.86 |
| HBE1 | 7.76 | 4.42E-02 | -0.98 |
| HBG2 | 6.68 | 1.03E-01 | -0.98 |
| HES4 | 7.59 | 3.69E-02 | 5.89 |
| HGF | 7.62 | 6.10E-02 | 4.58 |
| HHLA1 | -5.39 | 1.92E-01 | -0.98 |
| HLA-A | -5.88 | 1.33E-01 | -0.98 |
| HMGCLL1 | 7.90 | 4.06E-02 | -0.98 |
| HMGCS2 | 5.89 | 1.92E-01 | -0.98 |
| HNF4A | 7.35 | 6.10E-02 | -0.98 |

| | | | |
|---|---|---|---|
| HOPX | 7.02 | 8.06E-02 | -0.98 |
| HOXA10 | 6.30 | 1.42E-01 | -0.98 |
| HOXA10-HOXA9 | 6.82 | 9.14E-02 | -0.98 |
| HOXA11 | 6.17 | 1.55E-01 | -0.98 |
| HOXA13 | 6.05 | 1.67E-01 | -0.98 |
| HOXA9 | 6.90 | 8.63E-02 | -0.98 |
| HOXB2 | 8.55 | 2.44E-02 | -0.98 |
| HOXB3 | 9.35 | 1.30E-02 | -0.98 |
| HOXB4 | 8.19 | 3.25E-02 | -0.98 |
| HOXB5 | 6.92 | 8.56E-02 | -0.98 |
| HOXB6 | 7.38 | 6.10E-02 | -0.98 |
| HOXB7 | 6.54 | 1.15E-01 | -0.98 |
| HOXB8 | 7.73 | 4.52E-02 | -0.98 |
| HOXB9 | 7.53 | 5.51E-02 | -0.98 |
| HOXC4 | 7.58 | 9.19E-02 | 3.56 |
| HOXC5 | 6.62 | 1.07E-01 | -0.98 |
| HOXC6 | 7.87 | 4.19E-02 | -0.98 |
| HOXC8 | 8.25 | 3.14E-02 | -0.98 |
| HOXD4 | 6.45 | 1.27E-01 | -0.98 |
| HPD | 6.11 | 1.61E-01 | -0.98 |
| HSD17B2 | 6.25 | 1.47E-01 | -0.98 |
| HSF4 | 6.25 | 1.47E-01 | -0.98 |
| HSPB7 | 6.17 | 1.55E-01 | -0.98 |
| HYMAI | 6.33 | 1.41E-01 | -0.98 |
| ID4 | 5.88 | 1.03E-01 | 9.28 |
| IDO1 | -7.36 | 5.47E-02 | 5.53 |
| IFI44L | 7.02 | 8.06E-02 | -0.98 |
| IGF1 | 6.80 | 9.29E-02 | -0.98 |
| IGF2 | 11.02 | 2.60E-03 | 10.14 |
| IGF2-AS | 7.00 | 8.08E-02 | -0.98 |
| IGFBP3 | 6.07 | 8.95E-02 | 9.06 |
| IGFBP5 | 7.37 | 3.56E-02 | 7.78 |
| IHH | 6.35 | 1.35E-01 | -0.98 |
| INPP4B | 5.75 | 1.55E-01 | 4.97 |
| INS-IGF2 | 11.02 | 2.60E-03 | 10.14 |
| INSM1 | 8.94 | 3.28E-02 | 4.24 |
| IPO4 | -5.59 | 1.64E-01 | -0.98 |
| ISL1 | 8.47 | 2.63E-02 | -0.98 |
| ISL2 | 7.29 | 6.29E-02 | -0.98 |
| ISLR2 | 7.48 | 4.20E-02 | 5.51 |
| ITGA8 | 10.46 | 1.07E-02 | 5.00 |
| ITIH2 | 9.10 | 1.52E-02 | -0.98 |
| JAKMIP2-AS1 | -8.52 | 3.14E-02 | 4.53 |
| KANK4 | 5.94 | 1.13E-01 | 6.33 |
| KCNJ9 | 7.12 | 7.51E-02 | -0.98 |
| KCTD16 | 7.20 | 8.36E-02 | 4.37 |

| | | | |
|---|---|---|---|
| KIAA0895 | 5.91 | 1.35E-01 | 5.53 |
| KIAA1755 | 6.25 | 1.47E-01 | -0.98 |
| KLB | 6.88 | 1.07E-01 | 4.21 |
| KLHDC1 | 6.22 | 1.49E-01 | -0.98 |
| KLHDC7A | -7.41 | 5.15E-02 | 5.56 |
| KLKB1 | -7.45 | 7.12E-02 | 3.99 |
| KLRG2 | -5.81 | 1.45E-01 | 5.50 |
| KRT16 | 6.02 | 1.75E-01 | -0.98 |
| KRT24 | 6.76 | 9.61E-02 | -0.98 |
| L1TD1 | -5.23 | 1.69E-01 | 12.03 |
| LCK | -8.66 | 1.68E-02 | 5.60 |
| LEF1 | 7.13 | 4.41E-02 | 6.51 |
| LHX2 | 7.79 | 3.43E-02 | 5.26 |
| LHX5 | 7.06 | 6.10E-02 | 4.89 |
| LHX5-AS1 | 7.78 | 4.41E-02 | -0.98 |
| LHX9 | 8.04 | 3.59E-02 | -0.98 |
| LINC00261 | 6.83 | 1.67E-01 | 3.19 |
| LINC00428 | -7.73 | 3.00E-02 | 6.94 |
| LINC00461 | 11.23 | 5.41E-03 | 5.39 |
| LINC00643 | 6.14 | 1.58E-01 | -0.98 |
| LINC00678 | -7.61 | 3.14E-02 | 7.39 |
| LINC00982 | 7.71 | 8.36E-02 | 3.63 |
| LINC01158 | 7.45 | 5.91E-02 | -0.98 |
| LINC01266 | 6.05 | 1.67E-01 | -0.98 |
| LINC01551 | 9.55 | 1.21E-02 | -0.98 |
| LINC01833 | 5.92 | 1.87E-01 | -0.98 |
| LINC02381 | 9.45 | 1.24E-02 | -0.98 |
| LIPC | 6.62 | 1.07E-01 | -0.98 |
| LIX1 | 10.87 | 4.30E-03 | -0.98 |
| LMX1A | 7.89 | 4.12E-02 | -0.98 |
| LNCPRESS2 | -6.89 | 5.57E-02 | 7.03 |
| LOC100126447 | -6.67 | 8.00E-02 | 5.60 |
| LOC100130370 | 6.30 | 1.42E-01 | -0.98 |
| LOC101928499 | -6.79 | 1.16E-01 | 3.66 |
| LOC102723512 | 6.68 | 1.03E-01 | -0.98 |
| LOC102725048 | -6.14 | 1.06E-01 | -0.98 |
| LOC105369212 | 6.62 | 1.07E-01 | -0.98 |
| LOC105369436 | -7.31 | 4.01E-02 | 6.73 |
| LOC105370482 | -8.69 | 1.47E-02 | -0.98 |
| LOC105371455 | 7.09 | 7.71E-02 | -0.98 |
| LOC105372426 | 5.60 | 1.70E-01 | 5.16 |
| LOC105373409 | -5.38 | 1.87E-01 | 5.54 |
| LOC105376569 | -6.31 | 9.14E-02 | -0.98 |
| LOC105377647 | 6.40 | 1.31E-01 | -0.98 |
| LOC105378853 | 6.68 | 1.03E-01 | -0.98 |
| LOC107986058 | 7.70 | 4.66E-02 | -0.98 |

| | | | |
|---|---|---|---|
| LOC107987188 | 9.13 | 2.95E-02 | 4.34 |
| LOC109864269 | -6.06 | 9.10E-02 | 9.13 |
| LOC112268113 | 6.40 | 1.57E-01 | 3.97 |
| LOC642366 | 7.56 | 9.38E-02 | 3.55 |
| LOC644285 | 6.70 | 9.98E-02 | -0.98 |
| LOC653513 | 6.30 | 1.42E-01 | -0.98 |
| LOXL4 | 5.92 | 1.87E-01 | -0.98 |
| LRP1B | 7.45 | 5.91E-02 | -0.98 |
| LRP2 | 6.15 | 9.10E-02 | 7.17 |
| LRRC19 | 5.92 | 1.87E-01 | -0.98 |
| LRRC7 | 6.62 | 1.07E-01 | -0.98 |
| LRRN4 | 5.30 | 1.89E-01 | 6.01 |
| LRRTM2 | 7.73 | 4.52E-02 | -0.98 |
| LUM | 13.20 | 1.48E-03 | -0.98 |
| LURAP1L | 5.66 | 1.78E-01 | 4.19 |
| LYST | 6.46 | 9.38E-02 | 4.59 |
| MAB21L1 | 9.04 | 1.55E-02 | -0.98 |
| MAB21L2 | 9.62 | 1.17E-02 | -0.98 |
| MAPT | 5.75 | 1.51E-01 | 5.46 |
| MBNL2 | 5.18 | 1.99E-01 | 6.61 |
| MDGA1 | 6.73 | 1.22E-01 | 4.14 |
| MECOM | 5.82 | 1.49E-01 | 5.00 |
| MEIS1 | 9.63 | 1.73E-02 | 4.59 |
| MEIS2 | 10.89 | 4.25E-03 | -0.98 |
| MEP1A | 8.69 | 2.04E-02 | -0.98 |
| METTL24 | 5.54 | 1.99E-01 | 4.13 |
| MFAP4 | 5.64 | 1.35E-01 | 7.64 |
| MGP | 7.15 | 7.24E-02 | -0.98 |
| MIR100HG | 7.67 | 3.54E-02 | 5.93 |
| MIR124-2HG | 7.19 | 4.80E-02 | 6.37 |
| MIR302C | -5.75 | 1.47E-01 | -0.98 |
| MIR3606 | 11.83 | 2.60E-03 | -0.98 |
| MIR5004 | -5.77 | 1.45E-01 | -0.98 |
| MIR9-3HG | 6.72 | 7.64E-02 | 5.13 |
| MIR99AHG | 7.05 | 7.89E-02 | -0.98 |
| MLC1 | 6.74 | 9.79E-02 | -0.98 |
| MMP1 | 6.58 | 1.13E-01 | -0.98 |
| MMP25 | -7.25 | 4.50E-02 | -0.98 |
| MMRN1 | 9.49 | 1.23E-02 | -0.98 |
| MPO | 8.17 | 6.10E-02 | 3.86 |
| MPPED1 | 5.47 | 1.87E-01 | 5.32 |
| MSX1 | 6.94 | 6.42E-02 | 4.83 |
| MSX2 | 7.20 | 5.14E-02 | 5.38 |
| MTTP | 5.72 | 1.39E-01 | 6.44 |
| MVP | 6.96 | 6.06E-02 | 6.06 |
| MYL4 | 7.30 | 6.24E-02 | -0.98 |

| | | | |
|---|---|---|---|
| MYO16 | 7.12 | 1.36E-01 | 3.34 |
| MYT1L | 7.47 | 5.87E-02 | -0.98 |
| NANOG | -6.13 | 9.14E-02 | 6.92 |
| NANOGP1 | -6.60 | 7.63E-02 | -0.98 |
| NANOGP8 | -5.54 | 1.55E-01 | 6.36 |
| NDRG1 | 5.35 | 1.64E-01 | 7.36 |
| NDST3 | 6.29 | 1.70E-01 | 3.92 |
| NDST4 | 8.14 | 6.10E-02 | 3.84 |
| NETO1 | -6.09 | 1.46E-01 | 4.90 |
| NEUROD1 | 9.06 | 1.54E-02 | -0.98 |
| NEUROD4 | 7.49 | 5.70E-02 | -0.98 |
| NEUROD6 | 6.64 | 1.06E-01 | -0.98 |
| NEUROG1 | 7.12 | 7.51E-02 | -0.98 |
| NEUROG2 | 8.63 | 2.21E-02 | -0.98 |
| NFE2L3 | -5.22 | 1.74E-01 | 9.14 |
| NHLH1 | 7.21 | 4.97E-02 | 5.70 |
| NPNT | 8.53 | 1.48E-02 | 7.95 |
| NPR3 | 10.67 | 1.01E-02 | 5.11 |
| NR2E1 | 8.72 | 1.98E-02 | -0.98 |
| NR2F1 | 12.30 | 2.36E-03 | -0.98 |
| NR2F1-AS1 | 6.24 | 1.13E-01 | 4.48 |
| NR2F2 | 12.21 | 2.49E-03 | -0.98 |
| NR2F2-AS1 | 6.30 | 1.42E-01 | -0.98 |
| NRN1 | 9.20 | 1.44E-02 | -0.98 |
| NSG2 | 7.89 | 2.99E-02 | 6.53 |
| NTRK2 | 10.10 | 1.05E-02 | 5.82 |
| OLR1 | 7.84 | 4.28E-02 | -0.98 |
| OSR1 | 6.32 | 1.05E-01 | 4.52 |
| P2RX3 | 9.95 | 1.47E-02 | 4.75 |
| PAPPA | 7.17 | 4.37E-02 | 6.68 |
| PAPPA2 | 6.35 | 8.31E-02 | 6.53 |
| PAX3 | 6.23 | 1.14E-01 | 4.47 |
| PAX6 | 11.31 | 3.00E-03 | -0.98 |
| PCK1 | 6.87 | 8.89E-02 | -0.98 |
| PDE1A | 6.43 | 9.60E-02 | 4.58 |
| PDZRN4 | 6.78 | 9.54E-02 | -0.98 |
| PEAR1 | 5.25 | 1.96E-01 | 5.86 |
| PECAM1 | 8.83 | 3.56E-02 | 4.19 |
| PF4 | 6.15 | 1.22E-01 | 4.43 |
| PGAM1P5 | -5.55 | 1.67E-01 | -0.98 |
| PGM5 | 9.19 | 2.78E-02 | 4.37 |
| PHACTR3 | 5.83 | 1.48E-01 | 5.01 |
| PIRT | 5.92 | 1.87E-01 | -0.98 |
| PITX1 | 10.18 | 7.19E-03 | -0.98 |
| PITX2 | 8.50 | 1.79E-02 | 5.61 |
| PKHD1L1 | 7.83 | 3.31E-02 | 5.28 |

| | | | |
|---|---|---|---|
| PLA2G12B | 5.95 | 1.84E-01 | -0.98 |
| PLAGL1 | 5.74 | 1.28E-01 | 6.89 |
| PLCL1 | 6.55 | 1.42E-01 | 4.05 |
| PLEK | 7.03 | 7.99E-02 | -0.98 |
| PLG | 10.64 | 4.89E-03 | -0.98 |
| PLVAP | 6.70 | 1.27E-01 | 4.13 |
| PLXNA2 | 5.44 | 1.51E-01 | 7.85 |
| PNPLA5 | -6.52 | 1.47E-01 | 3.53 |
| POLR3G | -6.49 | 6.34E-02 | 9.16 |
| POSTN | 8.65 | 1.27E-02 | 9.56 |
| POU3F2 | 5.22 | 1.86E-01 | 6.97 |
| POU3F3 | 8.63 | 1.55E-02 | 6.09 |
| POU4F1 | 7.41 | 4.37E-02 | 5.48 |
| POU5F1 | -6.00 | 1.02E-01 | 6.85 |
| POU5F1B | -6.14 | 9.32E-02 | 7.15 |
| POU5F1P3 | -6.36 | 7.87E-02 | 7.03 |
| POU5F1P4 | -6.62 | 8.89E-02 | 4.58 |
| PPBP | 7.39 | 7.37E-02 | 4.47 |
| PRDM12 | 7.15 | 7.24E-02 | -0.98 |
| PRDM14 | -6.16 | 8.73E-02 | 7.52 |
| PRDM16 | 7.40 | 4.32E-02 | 5.80 |
| PRDM6 | 6.72 | 1.23E-01 | 4.14 |
| PRPH | 6.64 | 6.35E-02 | 6.26 |
| PRRX1 | 9.45 | 1.24E-02 | -0.98 |
| PSD2 | 5.97 | 1.33E-01 | 5.08 |
| PTH1R | 7.74 | 5.80E-02 | 4.64 |
| PTPRH | 5.89 | 1.51E-01 | 4.30 |
| PUS7L | 6.14 | 1.58E-01 | -0.98 |
| RAB27B | 7.18 | 1.30E-01 | 3.36 |
| RAI1 | 5.18 | 1.87E-01 | 7.82 |
| RALYL | 6.40 | 1.31E-01 | -0.98 |
| RARB | 8.12 | 6.10E-02 | 3.83 |
| RBP2 | 8.45 | 4.72E-02 | 4.00 |
| RBP4 | 8.93 | 1.30E-02 | 6.56 |
| RELN | 9.47 | 7.85E-03 | 7.83 |
| REN | 9.05 | 1.54E-02 | -0.98 |
| RFX4 | 8.70 | 4.01E-02 | 4.12 |
| RGS13 | 6.35 | 1.35E-01 | -0.98 |
| RGS4 | 7.91 | 2.86E-02 | 6.73 |
| RHOJ | 6.41 | 1.56E-01 | 3.98 |
| RIMBP2 | 5.96 | 1.32E-01 | 5.34 |
| RNASE1 | 6.05 | 1.67E-01 | -0.98 |
| RPS4Y1 | 6.90 | 8.63E-02 | -0.98 |
| RSPO1 | 9.37 | 2.27E-02 | 4.46 |
| RSPO2 | 6.57 | 6.44E-02 | 7.15 |
| RSPO3 | 8.44 | 3.23E-02 | 5.00 |

| | | | |
|---|---|---|---|
| RTN4RL1 | 7.88 | 4.17E-02 | -0.98 |
| RXRG | 7.81 | 4.37E-02 | -0.98 |
| S100A14 | 5.27 | 1.82E-01 | 6.50 |
| SAMD11 | 7.81 | 4.37E-02 | -0.98 |
| SAMD5 | 6.89 | 6.81E-02 | 4.80 |
| SCG2 | 5.65 | 1.42E-01 | 5.77 |
| SCN3A | 6.50 | 8.36E-02 | 5.61 |
| SCRG1 | 6.94 | 8.44E-02 | -0.98 |
| SEMA3C | 6.55 | 8.06E-02 | 6.05 |
| SEMG1 | -5.46 | 1.84E-01 | -0.98 |
| SERPINA1 | 10.36 | 5.63E-03 | -0.98 |
| SERPINC1 | 6.58 | 1.13E-01 | -0.98 |
| SERPIND1 | 6.60 | 1.09E-01 | -0.98 |
| SERPINE3 | 6.27 | 1.09E-01 | 4.50 |
| SEZ6L | 6.22 | 1.02E-01 | 5.69 |
| SGCG | 6.19 | 1.51E-01 | -0.98 |
| SHOX2 | 6.14 | 1.23E-01 | 4.43 |
| SI | 6.17 | 1.55E-01 | -0.98 |
| SIX1 | 5.87 | 1.42E-01 | 5.29 |
| SIX3 | 6.65 | 1.32E-01 | 4.10 |
| SKOR2 | 6.11 | 1.61E-01 | -0.98 |
| SLC16A3 | 5.51 | 1.46E-01 | 7.70 |
| SLC17A6 | 6.13 | 9.58E-02 | 6.30 |
| SLC22A7 | 7.82 | 4.32E-02 | -0.98 |
| SLC2A5 | 6.97 | 8.36E-02 | -0.98 |
| SLC30A4 | 7.69 | 8.52E-02 | 3.62 |
| SLC38A4 | -5.94 | 1.17E-01 | 6.41 |
| SLC39A5 | 7.55 | 9.44E-02 | 3.55 |
| SLC40A1 | 10.04 | 1.07E-02 | 5.79 |
| SLC5A12 | 6.10 | 9.41E-02 | 6.99 |
| SLC7A4 | 5.99 | 1.80E-01 | -0.98 |
| SLCO2B1 | 8.03 | 3.64E-02 | -0.98 |
| SLIT1 | 5.81 | 1.15E-01 | 7.27 |
| SLIT3 | 6.42 | 7.14E-02 | 8.54 |
| SLITRK6 | 7.15 | 5.80E-02 | 4.94 |
| SLPI | 7.09 | 7.71E-02 | -0.98 |
| SMLR1 | 5.92 | 1.87E-01 | -0.98 |
| SMPDL3B | -8.28 | 2.47E-02 | 5.41 |
| SNAI2 | 6.51 | 7.49E-02 | 6.83 |
| SOX1 | 7.25 | 4.77E-02 | 5.72 |
| SOX17 | 6.49 | 1.18E-01 | -0.98 |
| SOX6 | 9.73 | 1.23E-02 | 5.64 |
| SST | 6.81 | 7.01E-02 | 5.18 |
| ST18 | 9.02 | 3.14E-02 | 4.28 |
| STMN2 | 8.56 | 1.41E-02 | 8.05 |
| STMN4 | 8.63 | 1.55E-02 | 6.09 |

| | | | |
|---|---|---|---|
| SULT1E1 | 6.64 | 1.06E-01 | -0.98 |
| SULT2A1 | 6.54 | 1.15E-01 | -0.98 |
| SVEP1 | 5.13 | 1.97E-01 | 7.25 |
| TAL1 | 6.19 | 1.51E-01 | -0.98 |
| TBR1 | 9.73 | 1.60E-02 | 4.64 |
| TBX2 | 8.61 | 2.29E-02 | -0.98 |
| TBX20 | 9.04 | 3.14E-02 | 4.29 |
| TBX3 | 7.91 | 3.14E-02 | 5.31 |
| TBX4 | 7.69 | 4.71E-02 | -0.98 |
| TCF21 | 6.04 | 1.35E-01 | 4.38 |
| TCIM | 8.75 | 3.77E-02 | 4.15 |
| TDGF1 | -6.29 | 7.73E-02 | 10.00 |
| TDGF1P3 | -5.69 | 1.37E-01 | 6.92 |
| TECRL | 6.45 | 1.27E-01 | -0.98 |
| TERT | -5.68 | 1.49E-01 | 5.69 |
| TF | 8.91 | 3.40E-02 | 4.23 |
| TFAP2A | 9.39 | 1.28E-02 | -0.98 |
| TFAP2B | 9.24 | 1.41E-02 | -0.98 |
| TFEC | 5.92 | 1.87E-01 | -0.98 |
| TGFB2 | 5.28 | 1.93E-01 | 6.32 |
| TH | 6.35 | 1.35E-01 | -0.98 |
| THBD | 5.70 | 1.41E-01 | 6.43 |
| THCAT155 | 7.29 | 1.16E-01 | 3.42 |
| TKTL1 | 6.62 | 1.07E-01 | -0.98 |
| TLX3 | 7.21 | 1.27E-01 | 3.38 |
| TMEM132C | 5.67 | 1.33E-01 | 6.61 |
| TMEM132E | 5.44 | 2.00E-01 | 4.82 |
| TMEM88 | 5.97 | 1.09E-01 | 6.08 |
| TNFRSF11A | -5.62 | 1.64E-01 | 5.40 |
| TNFRSF19 | 6.11 | 1.16E-01 | 5.15 |
| TNFRSF8 | -5.54 | 1.75E-01 | 5.36 |
| TNFSF11 | -6.52 | 1.47E-01 | 3.53 |
| TNNI1 | 6.65 | 8.21E-02 | 4.68 |
| TNRC18 | 5.81 | 1.10E-01 | 8.44 |
| TPPP3 | 5.45 | 1.93E-01 | 5.08 |
| TRH | 9.99 | 9.30E-03 | -0.98 |
| TRIM67 | 8.42 | 4.85E-02 | 3.99 |
| TSHZ1 | 8.05 | 2.86E-02 | 5.39 |
| TSHZ2 | 7.46 | 4.37E-02 | 5.09 |
| TTR | 12.94 | 2.49E-03 | 6.25 |
| TUBB1 | 5.99 | 1.80E-01 | -0.98 |
| TWIST1 | 6.78 | 6.95E-02 | 5.49 |
| TXNIP | 6.42 | 6.80E-02 | 10.26 |
| UBA7 | 7.00 | 8.08E-02 | -0.98 |
| UGT2B11 | 7.30 | 6.24E-02 | -0.98 |
| UNC13C | 6.11 | 1.61E-01 | -0.98 |

| | | | |
|---|---|---|---|
| UNCX | 7.89 | 7.51E-02 | 3.72 |
| USP44 | -5.56 | 1.37E-01 | 9.05 |
| USP9Y | 5.95 | 1.84E-01 | -0.98 |
| VCAM1 | 8.36 | 2.98E-02 | -0.98 |
| VRTN | -6.56 | 6.13E-02 | 8.36 |
| VTN | 10.72 | 4.89E-03 | -0.98 |
| WISP1 | 8.53 | 2.46E-02 | -0.98 |
| WLS | 6.60 | 6.10E-02 | 7.93 |
| WNT1 | 6.90 | 8.63E-02 | -0.98 |
| WNT11 | 7.48 | 9.97E-02 | 3.52 |
| WNT4 | 5.89 | 1.51E-01 | 4.30 |
| WNT7A | 7.27 | 1.18E-01 | 3.41 |
| WNT7B | 9.34 | 1.54E-02 | 5.44 |
| WRBP1 | -6.16 | 1.04E-01 | -0.98 |
| YPEL4 | 5.64 | 1.82E-01 | 4.18 |
| ZBTB16 | 7.48 | 4.08E-02 | 5.84 |
| ZBTB20 | 6.43 | 8.85E-02 | 5.58 |
| ZEB1 | 6.70 | 6.02E-02 | 7.71 |
| ZEB2 | 8.22 | 2.09E-02 | 6.47 |
| ZFHX3 | 6.53 | 7.27E-02 | 6.74 |
| ZFHX4 | 5.68 | 1.27E-01 | 8.04 |
| ZFHX4-AS1 | 7.18 | 7.12E-02 | -0.98 |
| ZFPM2 | 6.98 | 6.10E-02 | 5.26 |
| ZIC1 | 7.76 | 2.81E-02 | 6.83 |
| ZIC4 | 8.23 | 3.14E-02 | -0.98 |
| ZNF467 | 6.67 | 7.87E-02 | 5.11 |
| ZNF503 | 10.54 | 5.39E-03 | -0.98 |
| ZNF503-AS2 | 6.19 | 1.51E-01 | -0.98 |
| ZNF536 | 8.06 | 3.56E-02 | -0.98 |
| ZNF540 | 5.65 | 1.80E-01 | 4.18 |
| ZNF804A | 6.70 | 1.87E-01 | 3.13 |
| ZSCAN10 | -5.63 | 1.31E-01 | 8.76 |
| ZYX | -6.58 | 9.10E-02 | 4.56 |
