## Supplemental Table 4 for "Brainstem organoids from human pluripotent stem cells contain neural crest population"

**Supplemental Table 4  Differentially expressed genes selective in hBSOs**

| Gene symbol | log (hBSO/hESC) | FDR (hBSO/hESC) | A (hBSO/hESC) | log (hBSO/hCO) | FDR (hBSO/hCO) | A (hBSO/hCO) |
|---|---|---|---|---|---|---|
| FLT1 | -10.91 | 6.73E-02 | 6.37 | -8.81 | 6.66E-09 | 5.12 |
| AP1M2 | -10.88 | 6.73E-02 | 5.36 | -7.05 | 4.19E-05 | 3.24 |
| APELA | -10.04 | 9.87E-02 | 4.93 | -6.39 | 3.05E-04 | 2.91 |
| TRIML2 | -9.73 | 7.93E-02 | -1.00 | -5.73 | 2.35E-03 | -0.97 |
| LNCPRESS1 | -9.26 | 9.21E-02 | -1.00 | -4.81 | 3.02E-02 | -0.97 |
| LAD1 | -9.14 | 1.02E-01 | -1.00 | -5.59 | 3.72E-03 | -0.97 |
| ZFP42 | -8.91 | 9.75E-02 | 7.54 | -4.59 | 6.68E-04 | 5.60 |
| CRYBG1 | -8.68 | 1.35E-01 | 5.25 | -3.73 | 1.19E-01 | 2.58 |
| VSNL1 | -8.28 | 1.36E-01 | 6.38 | -3.45 | 8.25E-02 | 3.76 |
| NKX1-2 | -7.95 | 1.68E-01 | 5.89 | -3.22 | 1.77E-01 | 3.33 |
| PODXL | -7.82 | 1.40E-01 | 11.22 | -6.71 | 8.60E-08 | 10.47 |
| CDCP1 | -7.78 | 1.95E-01 | 5.39 | -6.04 | 6.98E-05 | 4.32 |
| LIN28A | -7.71 | 1.50E-01 | 11.10 | -3.37 | 1.09E-02 | 8.73 |
| MAP1LC3C | 7.25 | 1.96E-01 | 7.80 | 5.58 | 8.51E-06 | 8.43 |
| CDK15 | 7.42 | 1.95E-01 | -1.00 | 4.15 | 4.20E-03 | 5.23 |
| CFAP54 | 7.42 | 1.95E-01 | -1.00 | 3.03 | 4.80E-02 | 5.87 |
| SV2C | 7.42 | 1.89E-01 | 6.61 | 4.79 | 5.41E-04 | 5.56 |
| GDF5 | 7.46 | 1.92E-01 | -1.00 | 4.67 | 1.65E-03 | 5.01 |
| ASIC4 | 7.51 | 1.89E-01 | -1.00 | 2.87 | 7.20E-02 | 5.96 |
| LINC02154 | 7.51 | 1.89E-01 | -1.00 | 6.04 | 1.31E-04 | 4.37 |
| LOC107986523 | 7.52 | 1.89E-01 | -1.00 | 7.06 | 3.12E-06 | -0.97 |
| OTX2-AS1 | 7.61 | 1.80E-01 | -1.00 | 3.68 | 1.04E-02 | 5.66 |
| LOC440982 | 7.61 | 1.79E-01 | -1.00 | 3.34 | 2.35E-02 | 5.83 |
| MUSK | 7.63 | 1.78E-01 | -1.00 | 3.26 | 2.77E-02 | 5.89 |
| CDH18 | 7.65 | 1.82E-01 | 5.91 | 4.77 | 2.07E-04 | 7.13 |
| LOC101928817 | 7.68 | 1.71E-01 | -1.00 | 4.89 | 8.05E-04 | 5.12 |
| LOC105379041 | 7.69 | 1.71E-01 | -1.00 | 7.23 | 3.55E-05 | 3.97 |
| LOC112268449 | 7.70 | 1.70E-01 | -1.00 | 3.65 | 1.02E-02 | 5.76 |
| OC90 | 7.71 | 1.68E-01 | -1.00 | 7.24 | 1.63E-06 | -0.97 |
| ASPA | 7.73 | 1.65E-01 | -1.00 | 6.58 | 2.06E-05 | -0.97 |
| TSHR | 7.77 | 1.65E-01 | -1.00 | 4.72 | 9.19E-04 | 5.30 |
| CFI | 7.84 | 1.60E-01 | -1.00 | 2.48 | 1.60E-01 | 6.45 |
| MYH7 | 7.86 | 1.57E-01 | -1.00 | 2.54 | 1.38E-01 | 6.48 |
| LOC105370767 | 7.92 | 1.51E-01 | -1.00 | 4.88 | 5.66E-04 | 5.38 |
| EDAR | 7.93 | 1.51E-01 | -1.00 | 4.88 | 5.57E-04 | 5.38 |
| CAPSL | 7.94 | 1.50E-01 | -1.00 | 5.90 | 6.51E-05 | 4.89 |
| CLUL1 | 7.99 | 1.79E-01 | 5.09 | 3.47 | 1.01E-02 | 7.09 |
| CAPN3 | 8.02 | 1.77E-01 | 5.10 | 3.57 | 7.22E-03 | 7.69 |
| CFAP47 | 8.05 | 1.40E-01 | -1.00 | 3.03 | 5.10E-02 | 5.96 |
| GPNMB | 8.08 | 1.32E-01 | 7.83 | 6.61 | 1.90E-07 | 8.36 |
| VWA3B | 8.10 | 1.36E-01 | -1.00 | 6.05 | 4.36E-05 | 4.96 |
| PDE7B | 8.11 | 1.35E-01 | -1.00 | 2.65 | 9.70E-02 | 6.68 |
| RPH3A | 8.11 | 1.35E-01 | -1.00 | 3.10 | 3.44E-02 | 6.43 |
| CFAP61 | 8.13 | 1.35E-01 | -1.00 | 4.59 | 7.14E-04 | 5.97 |
| SCARA5 | 8.13 | 1.65E-01 | 5.16 | 5.21 | 7.51E-05 | 6.42 |
| PALMD | 8.17 | 1.33E-01 | -1.00 | 3.12 | 3.11E-02 | 6.50 |
| CFAP206 | 8.17 | 1.32E-01 | -1.00 | 2.90 | 5.21E-02 | 6.61 |
| LOC441204 | 8.19 | 1.32E-01 | -1.00 | 2.51 | 1.30E-01 | 6.93 |
| FAM81B | 8.20 | 1.32E-01 | -1.00 | 4.34 | 1.56E-03 | 5.85 |
| PTPRC | 8.21 | 1.32E-01 | -1.00 | 6.74 | 1.25E-05 | 4.72 |
| LOC105379506 | 8.23 | 1.32E-01 | -1.00 | 7.76 | 2.14E-07 | -0.97 |
| PTGDS | 8.25 | 1.32E-01 | 7.38 | 4.89 | 8.26E-05 | 8.86 |
| MS4A6A | 8.28 | 1.29E-01 | -1.00 | 5.82 | 4.36E-05 | 5.26 |
| C11orf88 | 8.33 | 1.28E-01 | -1.00 | 5.54 | 9.15E-05 | 5.45 |
| WNT3A | 8.39 | 1.25E-01 | -1.00 | 2.84 | 6.12E-02 | 6.86 |
| ARMC3 | 8.43 | 1.25E-01 | -1.00 | 4.80 | 4.44E-04 | 5.92 |
| LOC101927120 | 8.44 | 1.25E-01 | -1.00 | 3.58 | 9.82E-03 | 6.54 |
| SLC16A6 | 8.47 | 1.79E-01 | 4.32 | 3.00 | 3.89E-02 | 6.86 |
| ENKUR | 8.47 | 1.79E-01 | 4.33 | 2.75 | 7.21E-02 | 6.90 |
| PRSS56 | 8.52 | 1.20E-01 | -1.00 | 3.66 | 7.87E-03 | 6.58 |
| SLC45A2 | 8.53 | 1.19E-01 | -1.00 | 8.06 | 8.12E-08 | -0.97 |
| CFAP52 | 8.54 | 1.19E-01 | -1.00 | 4.27 | 1.44E-03 | 6.30 |
| DIO3 | 8.54 | 1.71E-01 | 4.36 | 2.41 | 1.60E-01 | 7.23 |
| TEKT1 | 8.57 | 1.18E-01 | -1.00 | 4.30 | 1.29E-03 | 6.31 |
| ABCB5 | 8.59 | 1.18E-01 | -1.00 | 8.13 | 6.27E-08 | -0.97 |
| WNT10B | 8.84 | 1.08E-01 | -1.00 | 3.17 | 2.19E-02 | 7.52 |
| MAP3K19 | 8.88 | 1.07E-01 | -1.00 | 6.83 | 2.09E-06 | 5.35 |
| LINC00473 | 8.92 | 1.05E-01 | -1.00 | 5.65 | 3.44E-05 | 5.99 |
| MEOX1 | 8.98 | 1.01E-01 | -1.00 | 4.65 | 2.49E-04 | 7.44 |
| EMP1 | 8.99 | 1.02E-01 | 7.58 | 5.82 | 2.80E-06 | 8.96 |
| CNPY1 | 9.03 | 9.75E-02 | -1.00 | 4.24 | 1.10E-03 | 6.80 |
| LOC105374972 | 9.04 | 9.67E-02 | -1.00 | 8.58 | 1.17E-08 | -0.97 |
| LMX1B | 9.05 | 1.16E-01 | 5.61 | 3.24 | 1.63E-02 | 8.31 |
| CRYAB | 9.19 | 1.08E-01 | 5.68 | 3.57 | 6.48E-03 | 8.29 |
| CCNO | 9.22 | 8.61E-02 | -1.00 | 5.34 | 5.95E-05 | 6.35 |
| EFCAB1 | 9.22 | 8.61E-02 | -1.00 | 3.72 | 4.94E-03 | 7.30 |
| FAM216B | 9.28 | 8.30E-02 | -1.00 | 5.64 | 2.86E-05 | 6.34 |
| SPAG6 | 9.29 | 1.05E-01 | 5.73 | 4.46 | 4.22E-04 | 7.94 |
| SLC4A5 | 9.34 | 8.17E-02 | 8.08 | 4.82 | 9.66E-05 | 9.45 |
| CFAP126 | 9.34 | 8.17E-02 | -1.00 | 3.97 | 2.25E-03 | 7.25 |
| LGI1 | 9.41 | 1.18E-01 | 4.80 | 2.98 | 4.09E-02 | 7.09 |
| KCNJ13 | 9.57 | 7.91E-02 | -1.00 | 4.62 | 3.26E-04 | 7.13 |
| EN1 | 9.64 | 7.57E-02 | -1.00 | 5.86 | 9.62E-06 | 6.60 |
| SCGN | 9.76 | 6.96E-02 | -1.00 | 5.84 | 8.38E-06 | 6.73 |
| RPE65 | 10.07 | 6.73E-02 | -1.00 | 7.29 | 1.42E-07 | 6.32 |
| RTL1 | 10.21 | 6.73E-02 | -1.00 | 4.99 | 7.97E-05 | 7.61 |
| EN2 | 10.91 | 6.73E-02 | 5.54 | 5.08 | 5.35E-05 | 8.25 |
| TYR | 11.82 | 5.99E-02 | -1.00 | 10.35 | 1.81E-11 | 6.53 |
| LOC105374794 | 12.08 | 5.79E-02 | -1.00 | 11.62 | 1.98E-12 | 6.16 |
| MLANA | 12.53 | 3.69E-02 | -1.00 | 9.07 | 7.27E-11 | 7.89 |
| TYRP1 | 13.71 | 2.97E-02 | 6.95 | 7.22 | 1.29E-08 | 9.97 |
