## Supplemental Table 5 for "Brainstem organoids from human pluripotent stem cells contain neural crest population"

**Supplemental Table 5  Differentially expressed genes selective in hCOs**

| Gene symbol | log (hCO/hESC) | FDR (hCO/hESC) | A (hCO/hESC) | log (hCO/hBSO) | FDR (hCO/hBSO) | A (hCO/hBSO) |
|---|---|---|---|---|---|---|
| LOC101928499 | -6.79 | 1.16E-01 | 3.66 | -5.77 | 3.27E-03 | 3.24 |
| ZYX | -6.58 | 9.10E-02 | 4.56 | -4.63 | 1.14E-02 | 3.67 |
| HLA-A | -5.88 | 1.33E-01 | -0.98 | -2.27 | 1.88E-01 | 9.85 |
| CHCHD10 | -5.79 | 1.70E-01 | 4.16 | -5.66 | 4.93E-04 | 4.18 |
| ATP10A | -5.74 | 1.80E-01 | 4.14 | -5.69 | 4.24E-04 | 4.20 |
| TNFRSF11A | -5.62 | 1.64E-01 | 5.40 | -5.11 | 4.14E-04 | 4.91 |
| IPO4 | -5.59 | 1.64E-01 | -0.98 | -4.72 | 7.00E-03 | -0.97 |
| GAL | -5.52 | 1.53E-01 | 6.49 | -3.33 | 2.85E-02 | 5.48 |
| NFE2L3 | -5.22 | 1.74E-01 | 9.14 | -2.76 | 6.42E-02 | 8.00 |
| SVEP1 | 5.13 | 1.97E-01 | 7.25 | 7.18 | 1.12E-07 | 6.31 |
| ACKR3 | 5.22 | 1.84E-01 | 6.84 | 2.32 | 1.78E-01 | 8.38 |
| PEAR1 | 5.25 | 1.96E-01 | 5.86 | 3.16 | 2.31E-02 | 6.99 |
| S100A14 | 5.27 | 1.82E-01 | 6.50 | 8.51 | 1.93E-08 | 4.97 |
| LRRN4 | 5.30 | 1.89E-01 | 6.01 | 9.07 | 1.89E-08 | 4.25 |
| MPPED1 | 5.47 | 1.87E-01 | 5.32 | 3.00 | 3.94E-02 | 6.64 |
| CCDC68 | 5.58 | 1.90E-01 | 4.15 | 3.06 | 4.95E-02 | 5.50 |
| DKK2 | 5.60 | 1.49E-01 | 6.04 | 4.51 | 4.51E-04 | 6.67 |
| MFAP4 | 5.64 | 1.35E-01 | 7.64 | 4.15 | 9.19E-04 | 8.47 |
| GNRH2 | 5.66 | 1.78E-01 | 4.19 | 7.39 | 1.12E-05 | 3.41 |
| THBD | 5.70 | 1.41E-01 | 6.43 | 5.33 | 4.50E-05 | 6.70 |
| MTTP | 5.72 | 1.39E-01 | 6.44 | 3.94 | 2.11E-03 | 7.41 |
| FRZB | 5.75 | 1.15E-01 | 9.33 | 3.85 | 2.13E-03 | 10.37 |
| C1orf61 | 5.88 | 1.17E-01 | 5.88 | 9.43 | 4.96E-09 | 4.43 |
| TFEC | 5.92 | 1.87E-01 | -0.98 | 3.06 | 9.68E-02 | 4.25 |
| LOXL4 | 5.92 | 1.87E-01 | -0.98 | 3.48 | 3.89E-02 | 4.04 |
| DLX5 | 5.92 | 1.87E-01 | -0.98 | 6.06 | 8.75E-04 | 2.75 |
| LRRC19 | 5.92 | 1.87E-01 | -0.98 | 6.06 | 8.75E-04 | -0.97 |
| SMLR1 | 5.92 | 1.87E-01 | -0.98 | 6.06 | 8.75E-04 | -0.97 |
| KANK4 | 5.94 | 1.13E-01 | 6.33 | 2.90 | 4.40E-02 | 7.66 |
| PLA2G12B | 5.95 | 1.84E-01 | -0.98 | 5.10 | 3.43E-03 | 3.26 |
| USP9Y | 5.95 | 1.84E-01 | -0.98 | 6.10 | 8.10E-04 | 2.76 |
| TMEM88 | 5.97 | 1.09E-01 | 6.08 | 3.41 | 1.15E-02 | 7.33 |
| TUBB1 | 5.99 | 1.80E-01 | -0.98 | 6.13 | 7.50E-04 | 2.78 |
| HAPLN1 | 5.99 | 1.02E-01 | 6.86 | 4.71 | 1.79E-04 | 7.62 |
| GBP4 | 6.02 | 1.75E-01 | -0.98 | 4.58 | 7.75E-03 | 3.59 |
| KRT16 | 6.02 | 1.75E-01 | -0.98 | 6.16 | 6.37E-04 | 2.80 |
| TCF21 | 6.04 | 1.35E-01 | 4.38 | 7.77 | 2.52E-06 | -0.97 |
| RNASE1 | 6.05 | 1.67E-01 | -0.98 | 4.61 | 7.28E-03 | 3.60 |
| HOXA13 | 6.05 | 1.67E-01 | -0.98 | 6.19 | 5.90E-04 | -0.97 |
| LINC01266 | 6.05 | 1.67E-01 | -0.98 | 6.19 | 5.90E-04 | 2.81 |
| IGFBP3 | 6.07 | 8.95E-02 | 9.06 | 4.10 | 9.73E-04 | 10.13 |
| SLC5A12 | 6.10 | 9.41E-02 | 6.99 | 6.17 | 1.68E-06 | 7.05 |
| GBP2 | 6.14 | 1.58E-01 | -0.98 | 2.82 | 1.14E-01 | 4.59 |
| PUS7L | 6.14 | 1.58E-01 | -0.98 | 6.28 | 4.40E-04 | -0.97 |
| PF4 | 6.15 | 1.22E-01 | 4.43 | 7.89 | 1.64E-06 | -0.97 |
| HSPB7 | 6.17 | 1.55E-01 | -0.98 | 2.73 | 1.41E-01 | 4.54 |
| HOXA11 | 6.17 | 1.55E-01 | -0.98 | 6.31 | 4.10E-04 | -0.97 |
| SI | 6.17 | 1.55E-01 | -0.98 | 6.31 | 4.10E-04 | -0.97 |
| ALPK2 | 6.18 | 1.08E-01 | 5.45 | 5.32 | 5.83E-05 | 5.96 |
| SGCG | 6.19 | 1.51E-01 | -0.98 | 2.88 | 1.02E-01 | 4.61 |
| TAL1 | 6.19 | 1.51E-01 | -0.98 | 6.34 | 3.85E-04 | 2.89 |
| SERPINE3 | 6.27 | 1.09E-01 | 4.50 | 6.42 | 2.10E-05 | 4.51 |
| ALDH1A1 | 6.29 | 8.61E-02 | 6.50 | 5.70 | 1.12E-05 | 6.89 |
| NR2F2-AS1 | 6.30 | 1.42E-01 | -0.98 | 3.22 | 6.82E-02 | 4.13 |
| ACSL5 | 6.30 | 1.42E-01 | -0.98 | 6.45 | 2.63E-04 | -0.97 |
| HOXA10 | 6.30 | 1.42E-01 | -0.98 | 6.45 | 2.63E-04 | -0.97 |
| OSR1 | 6.32 | 1.05E-01 | 4.52 | 5.05 | 2.70E-04 | 5.24 |
| IHH | 6.35 | 1.35E-01 | -0.98 | 4.17 | 1.17E-02 | 4.12 |
| RGS13 | 6.35 | 1.35E-01 | -0.98 | 6.50 | 2.14E-04 | -0.97 |
| GJB1 | 6.40 | 1.31E-01 | -0.98 | 4.54 | 5.64E-03 | 3.99 |
| CXorf36 | 6.42 | 1.29E-01 | -0.98 | 6.57 | 1.79E-04 | -0.97 |
| ABI3BP | 6.45 | 1.51E-01 | 4.00 | 2.65 | 1.18E-01 | 5.95 |
| CD300E | 6.45 | 1.27E-01 | -0.98 | 6.59 | 1.57E-04 | 3.01 |
| HOXD4 | 6.45 | 1.27E-01 | -0.98 | 6.59 | 1.57E-04 | -0.97 |
| TECRL | 6.45 | 1.27E-01 | -0.98 | 6.59 | 1.57E-04 | -0.97 |
| SOX17 | 6.49 | 1.18E-01 | -0.98 | 6.64 | 1.40E-04 | -0.97 |
| BNC1 | 6.51 | 7.44E-02 | 6.73 | 6.89 | 1.58E-07 | 6.62 |
| C5orf66-AS1 | 6.52 | 1.17E-01 | -0.98 | 6.66 | 1.31E-04 | -0.97 |
| HOXB7 | 6.54 | 1.15E-01 | -0.98 | 5.10 | 1.50E-03 | 3.85 |
| SULT2A1 | 6.54 | 1.15E-01 | -0.98 | 6.68 | 1.17E-04 | -0.97 |
| MMP1 | 6.58 | 1.13E-01 | -0.98 | 5.73 | 4.84E-04 | 3.58 |
| SERPINC1 | 6.58 | 1.13E-01 | -0.98 | 6.73 | 1.05E-04 | 3.08 |
| SERPIND1 | 6.60 | 1.09E-01 | -0.98 | 6.75 | 9.90E-05 | 3.09 |
| LOC105369212 | 6.62 | 1.07E-01 | -0.98 | 2.60 | 1.57E-01 | 5.18 |
| LIPC | 6.62 | 1.07E-01 | -0.98 | 6.34 | 3.85E-04 | -0.97 |
| ADH4 | 6.62 | 1.07E-01 | -0.98 | 6.77 | 8.75E-05 | -0.97 |
| HOXC5 | 6.62 | 1.07E-01 | -0.98 | 6.77 | 8.75E-05 | -0.97 |
| ADGRF5 | 6.64 | 1.96E-01 | 3.10 | 5.73 | 4.84E-04 | 3.58 |
| FETUB | 6.64 | 1.06E-01 | -0.98 | 6.79 | 8.39E-05 | -0.97 |
| NEUROD6 | 6.64 | 1.06E-01 | -0.98 | 6.79 | 8.39E-05 | 3.11 |
| SULT1E1 | 6.64 | 1.06E-01 | -0.98 | 6.79 | 8.39E-05 | -0.97 |
| TNNI1 | 6.65 | 8.21E-02 | 4.68 | 2.60 | 1.09E-01 | 6.80 |
| C1QTNF7 | 6.66 | 1.04E-01 | -0.98 | 4.22 | 4.81E-03 | 4.41 |
| HBG2 | 6.68 | 1.03E-01 | -0.98 | 6.83 | 7.72E-05 | -0.97 |
| ESAM | 6.70 | 1.87E-01 | 3.13 | 3.85 | 1.19E-02 | 4.64 |
| PLVAP | 6.70 | 1.27E-01 | 4.13 | 5.53 | 1.61E-04 | 4.80 |
| MIR9-3HG | 6.72 | 7.64E-02 | 5.13 | 4.82 | 2.15E-04 | 6.12 |
| PRDM6 | 6.72 | 1.23E-01 | 4.14 | 4.85 | 7.75E-03 | 3.14 |
| ANKRD1 | 6.74 | 1.81E-01 | 3.14 | 2.98 | 7.70E-02 | 5.11 |
| MLC1 | 6.74 | 9.79E-02 | -0.98 | 4.28 | 3.90E-03 | 4.44 |
| KRT24 | 6.76 | 9.61E-02 | -0.98 | 6.90 | 6.12E-05 | -0.97 |
| AMBN | 6.78 | 9.54E-02 | -0.98 | 6.92 | 5.83E-05 | -0.97 |
| F2 | 6.80 | 1.70E-01 | 3.17 | 6.94 | 5.35E-05 | -0.97 |
| HOXA10-HOXA9 | 6.82 | 9.14E-02 | -0.98 | 6.96 | 5.23E-05 | -0.97 |
| A1CF | 6.83 | 1.67E-01 | 3.19 | 6.98 | 5.01E-05 | -0.97 |

| | | | | | | |
|---|---|---|---|---|---|---|
| LINC00261 | 6.83 | 1.67E-01 | 3.19 | 6.98 | 5.01E-05 | -0.97 |
| PCK1 | 6.87 | 8.89E-02 | -0.98 | 4.01 | 7.69E-03 | 4.72 |
| KLB | 6.88 | 1.07E-01 | 4.21 | 8.02 | 1.01E-06 | -0.97 |
| AHNAK | 6.89 | 4.64E-02 | 10.56 | 3.14 | 1.84E-02 | 12.52 |
| APOH | 6.90 | 8.63E-02 | -0.98 | 7.05 | 4.19E-05 | 3.24 |
| HOXA9 | 6.90 | 8.63E-02 | -0.98 | 7.05 | 4.19E-05 | -0.97 |
| RPS4Y1 | 6.90 | 8.63E-02 | -0.98 | 7.05 | 4.19E-05 | -0.97 |
| HOXB5 | 6.92 | 8.56E-02 | -0.98 | 4.48 | 1.99E-03 | 4.54 |
| SCRG1 | 6.94 | 8.44E-02 | -0.98 | 3.38 | 2.61E-02 | 5.11 |
| CSF3R | 6.95 | 8.36E-02 | -0.98 | 4.51 | 1.88E-03 | 4.56 |
| SLC2A5 | 6.97 | 8.36E-02 | -0.98 | 3.83 | 1.19E-02 | 4.80 |
| ZFPM2 | 6.98 | 6.10E-02 | 5.26 | 3.23 | 1.85E-02 | 7.16 |
| IGF2-AS | 7.00 | 8.08E-02 | -0.98 | 2.69 | 1.12E-01 | 5.52 |
| UBA7 | 7.00 | 8.08E-02 | -0.98 | 3.78 | 1.49E-02 | 4.42 |
| FAM107A | 7.01 | 6.10E-02 | 4.87 | 2.93 | 4.56E-02 | 6.99 |
| HOPX | 7.02 | 8.06E-02 | -0.98 | 4.58 | 1.54E-03 | 4.59 |
| PLEK | 7.03 | 7.99E-02 | -0.98 | 3.72 | 1.07E-02 | 5.03 |
| EGFLAM | 7.06 | 9.31E-02 | 4.30 | 4.50 | 8.75E-04 | 5.67 |
| SLPI | 7.09 | 7.71E-02 | -0.98 | 4.65 | 1.22E-03 | 4.63 |
| LOC105371455 | 7.09 | 7.71E-02 | -0.98 | 7.24 | 2.06E-05 | 3.33 |
| PRDM12 | 7.15 | 7.24E-02 | -0.98 | 3.39 | 2.55E-02 | 5.32 |
| PAPPA | 7.17 | 4.37E-02 | 6.68 | 3.04 | 2.62E-02 | 9.04 |
| CDX2 | 7.19 | 7.03E-02 | -0.98 | 7.34 | 1.37E-05 | -0.97 |
| WNT7A | 7.27 | 1.18E-01 | 3.41 | 2.78 | 9.21E-02 | 5.75 |
| THCAT155 | 7.29 | 1.16E-01 | 3.42 | 2.53 | 1.57E-01 | 5.89 |
| BRINP3 | 7.30 | 6.24E-02 | -0.98 | 3.21 | 3.26E-02 | 5.32 |
| MYL4 | 7.30 | 6.24E-02 | -0.98 | 6.45 | 4.93E-05 | 3.94 |
| UGT2B11 | 7.30 | 6.24E-02 | -0.98 | 7.45 | 9.20E-06 | -0.97 |
| HNF4A | 7.35 | 6.10E-02 | -0.98 | 6.50 | 4.19E-05 | 3.96 |
| APOB | 7.36 | 3.35E-02 | 9.83 | 9.36 | 8.97E-12 | 8.92 |
| HOXB6 | 7.38 | 6.10E-02 | -0.98 | 7.52 | 6.97E-06 | -0.97 |
| PPBP | 7.39 | 7.37E-02 | 4.47 | 8.54 | 1.40E-07 | -0.97 |
| EPHA5 | 7.40 | 4.52E-02 | 5.06 | 2.88 | 4.99E-02 | 7.30 |
| G0S2 | 7.41 | 4.21E-02 | 6.07 | 5.50 | 1.96E-05 | 7.11 |
| GATA3 | 7.48 | 9.97E-02 | 3.52 | 2.46 | 1.58E-01 | 6.11 |
| HOXB9 | 7.53 | 5.51E-02 | -0.98 | 7.67 | 3.55E-06 | 3.55 |
| FMO1 | 7.55 | 9.44E-02 | 3.55 | 4.30 | 3.09E-03 | 4.86 |
| SLC39A5 | 7.55 | 9.44E-02 | 3.55 | 6.11 | 5.83E-05 | 4.36 |
| HOXC4 | 7.58 | 9.19E-02 | 3.56 | 6.14 | 5.35E-05 | 4.37 |
| HGF | 7.62 | 6.10E-02 | 4.58 | 3.00 | 3.64E-02 | 6.95 |
| CRHBP | 7.64 | 4.93E-02 | -0.98 | 4.62 | 1.02E-03 | 5.19 |
| TBX4 | 7.69 | 4.71E-02 | -0.98 | 5.16 | 1.28E-03 | 3.88 |
| LOC107986058 | 7.70 | 4.66E-02 | -0.98 | 5.26 | 1.61E-04 | 4.93 |
| HOXB8 | 7.73 | 4.52E-02 | -0.98 | 3.18 | 2.71E-02 | 6.00 |
| ANXA8L1 | 7.73 | 4.52E-02 | -0.98 | 7.88 | 1.64E-06 | -0.97 |
| HBE1 | 7.76 | 4.42E-02 | -0.98 | 7.90 | 1.59E-06 | 3.67 |
| FOXF1 | 7.77 | 4.41E-02 | -0.98 | 3.06 | 4.01E-02 | 6.10 |
| LHX2 | 7.79 | 3.43E-02 | 5.26 | 2.73 | 6.82E-02 | 7.88 |
| SLC22A7 | 7.82 | 4.32E-02 | -0.98 | 6.39 | 5.44E-05 | 3.91 |
| ALB | 7.82 | 7.87E-02 | 3.69 | 6.97 | 6.94E-06 | 4.20 |
| PKHD1L1 | 7.83 | 3.31E-02 | 5.28 | 6.98 | 2.94E-07 | 5.79 |
| OLR1 | 7.84 | 4.28E-02 | -0.98 | 7.99 | 1.13E-06 | -0.97 |
| AKR1D1 | 7.85 | 4.26E-02 | -0.98 | 8.00 | 1.08E-06 | 3.71 |
| HOXC6 | 7.87 | 4.19E-02 | -0.98 | 8.01 | 1.03E-06 | 3.72 |
| SLCO2B1 | 8.03 | 3.64E-02 | -0.98 | 5.86 | 5.62E-05 | 4.97 |
| AKR1B10 | 8.13 | 3.42E-02 | -0.98 | 8.27 | 3.47E-07 | 3.85 |
| MPO | 8.17 | 6.10E-02 | 3.86 | 8.32 | 2.99E-07 | -0.97 |
| HOXB4 | 8.19 | 3.25E-02 | -0.98 | 8.33 | 2.94E-07 | -0.97 |
| HOXC8 | 8.25 | 3.14E-02 | -0.98 | 8.39 | 2.28E-07 | -0.97 |
| FIBIN | 8.33 | 3.10E-02 | -0.98 | 3.61 | 8.40E-03 | 6.38 |
| AGTR1 | 8.33 | 5.26E-02 | 3.94 | 8.48 | 1.71E-07 | -0.97 |
| FMOD | 8.35 | 2.06E-02 | 5.53 | 3.16 | 2.09E-02 | 7.85 |
| EOMES | 8.39 | 4.99E-02 | 3.97 | 2.90 | 4.88E-02 | 6.69 |
| GATA5 | 8.44 | 2.74E-02 | -0.98 | 4.67 | 4.97E-04 | 5.96 |
| RBP2 | 8.45 | 4.72E-02 | 4.00 | 8.60 | 1.12E-07 | -0.97 |
| PITX2 | 8.50 | 1.79E-02 | 5.61 | 7.91 | 2.39E-08 | 5.99 |
| CD93 | 8.50 | 4.52E-02 | 4.03 | 7.06 | 1.64E-06 | 4.83 |
| WISP1 | 8.53 | 2.46E-02 | -0.98 | 2.43 | 1.54E-01 | 7.16 |
| NPNT | 8.53 | 1.48E-02 | 7.95 | 8.40 | 3.31E-10 | 8.08 |
| GATA6-AS1 | 8.53 | 2.46E-02 | -0.98 | 8.68 | 8.94E-08 | -0.97 |
| HOXB2 | 8.55 | 2.44E-02 | -0.98 | 5.89 | 1.83E-05 | 5.47 |
| NEUROG2 | 8.63 | 2.21E-02 | -0.98 | 5.46 | 6.06E-05 | 5.77 |
| POSTN | 8.65 | 1.27E-02 | 9.56 | 3.74 | 2.98E-03 | 11.87 |
| MEP1A | 8.69 | 2.04E-02 | -0.98 | 8.84 | 4.70E-08 | -0.97 |
| NR2E1 | 8.72 | 1.98E-02 | -0.98 | 3.03 | 3.27E-02 | 7.06 |
| PECAM1 | 8.83 | 3.56E-02 | 4.19 | 6.98 | 1.27E-06 | 5.20 |
| COL6A3 | 8.88 | 1.23E-02 | 7.67 | 2.90 | 3.77E-02 | 10.74 |
| TF | 8.91 | 3.40E-02 | 4.23 | 3.94 | 2.66E-03 | 6.77 |
| RBP4 | 8.93 | 1.30E-02 | 6.56 | 3.93 | 1.75E-03 | 9.15 |
| ANXA8 | 8.96 | 1.61E-02 | -0.98 | 8.10 | 9.63E-08 | 4.77 |
| GUCY1A1 | 9.02 | 1.55E-02 | -0.98 | 3.39 | 1.29E-02 | 6.90 |
| TBX20 | 9.04 | 3.14E-02 | 4.29 | 9.18 | 1.26E-08 | -0.97 |
| REN | 9.05 | 1.54E-02 | -0.98 | 5.29 | 6.31E-05 | 6.27 |
| NEUROD1 | 9.06 | 1.54E-02 | -0.98 | 3.42 | 1.15E-02 | 7.21 |
| GSTA1 | 9.09 | 1.54E-02 | -0.98 | 6.65 | 1.08E-06 | 5.62 |
| H19 | 9.09 | 1.19E-02 | 7.32 | 7.33 | 1.44E-08 | 8.29 |
| ITIH2 | 9.10 | 1.52E-02 | -0.98 | 7.66 | 1.67E-07 | 5.13 |
| GSTA2 | 9.14 | 1.48E-02 | -0.98 | 9.29 | 9.31E-09 | -0.97 |
| DNM3OS | 9.35 | 1.31E-02 | -0.98 | 5.16 | 9.83E-05 | 5.76 |
| HOXB3 | 9.35 | 1.30E-02 | -0.98 | 8.43 | 2.57E-08 | 4.93 |
| FOXG1 | 9.40 | 1.48E-02 | 5.47 | 10.54 | 7.27E-11 | 4.99 |
| LINC02381 | 9.45 | 1.24E-02 | -0.98 | 2.76 | 6.15E-02 | 7.93 |
| RELN | 9.47 | 7.85E-03 | 7.83 | 3.11 | 2.10E-02 | 11.10 |
| LINC01551 | 9.55 | 1.21E-02 | -0.98 | 9.69 | 1.74E-09 | 4.56 |
| HAND2-AS1 | 9.57 | 1.19E-02 | -0.98 | 9.71 | 1.63E-09 | 4.57 |
| MEIS1 | 9.63 | 1.73E-02 | 4.59 | 2.84 | 4.95E-02 | 8.08 |
| TBR1 | 9.73 | 1.60E-02 | 4.64 | 6.87 | 3.19E-07 | 6.15 |
| AGT | 9.82 | 1.54E-02 | 4.68 | 3.13 | 2.19E-02 | 8.11 |

| | | | | | | |
|---|---|---|---|---|---|---|
| FABP1 | 9.93 | 9.93E-03 | -0.98 | 10.08 | 3.76E-10 | -0.97 |
| SLC40A1 | 10.04 | 1.07E-02 | 5.79 | 5.08 | 5.01E-05 | 8.36 |
| GATA6 | 10.05 | 8.53E-03 | -0.98 | 8.61 | 4.96E-09 | 5.61 |
| PITX1 | 10.18 | 7.19E-03 | -0.98 | 7.74 | 1.70E-08 | 6.17 |
| HAND2 | 10.20 | 6.98E-03 | -0.98 | 9.35 | 7.19E-10 | 5.39 |
| SERPINA1 | 10.36 | 5.63E-03 | -0.98 | 10.50 | 7.27E-11 | -0.97 |
| PLG | 10.64 | 4.89E-03 | -0.98 | 10.78 | 4.13E-11 | -0.97 |
| NPR3 | 10.67 | 1.01E-02 | 5.11 | 3.26 | 1.43E-02 | 8.91 |
| VTN | 10.72 | 4.89E-03 | -0.98 | 4.78 | 1.24E-04 | 8.19 |
| AFP | 10.74 | 2.93E-03 | 10.95 | 15.69 | 3.72E-21 | 8.56 |
| AHSG | 10.78 | 3.47E-03 | 6.75 | 11.51 | 2.68E-13 | 6.47 |
| INS-IGF2 | 11.02 | 2.60E-03 | 10.14 | 4.69 | 1.28E-04 | 13.40 |
| IGF2 | 11.02 | 2.60E-03 | 10.14 | 4.71 | 1.18E-04 | 13.45 |
| LINC00461 | 11.23 | 5.41E-03 | 5.39 | 4.67 | 1.69E-04 | 8.75 |
| APOA4 | 11.32 | 3.00E-03 | -0.98 | 11.47 | 2.16E-12 | 5.45 |
| FGG | 11.37 | 3.00E-03 | -0.98 | 11.53 | 1.93E-12 | -0.97 |
| HAND1 | 11.54 | 2.93E-03 | -0.98 | 9.10 | 7.67E-11 | 6.85 |
| APOA1 | 11.63 | 2.60E-03 | 7.18 | 8.36 | 2.30E-10 | 8.90 |
| APOA2 | 11.66 | 4.25E-03 | 5.60 | 11.80 | 6.11E-13 | -0.97 |
| FGA | 11.77 | 2.60E-03 | -0.98 | 11.91 | 4.41E-13 | -0.97 |
| APOC3 | 11.88 | 2.60E-03 | -0.98 | 12.03 | 2.81E-13 | -0.97 |
| FGB | 12.36 | 2.32E-03 | -0.98 | 12.51 | 6.85E-14 | -0.97 |
