## Supplemental Table 6 for "Brainstem organoids from human pluripotent stem cells contain neural crest population"

| | p_val | avg_logFC | pct.1 | pct.2 | p_val_adj | cluster | gene |
|---|---|---|---|---|---|---|---|
| EIF1 | 2.31E-73 | 0.831921 | 1 | 1 | 4.50E-69 | U | EIF1 |
| RPS27 | 1.34E-69 | 0.773046 | 1 | 0.999 | 2.60E-65 | U | RPS27 |
| RPL21 | 6.05E-67 | 0.800269 | 1 | 1 | 1.18E-62 | U | RPL21 |
| FAU | 2.22E-66 | 0.571711 | 1 | 1 | 4.32E-62 | U | FAU |
| TRMT112 | 1.15E-65 | 0.873833 | 0.994 | 0.963 | 2.23E-61 | U | TRMT112 |
| RPS16 | 4.07E-64 | 0.641277 | 1 | 0.999 | 7.92E-60 | U | RPS16 |
| RPS25 | 1.05E-59 | 0.65538 | 1 | 1 | 2.04E-55 | U | RPS25 |
| RPS13 | 1.48E-59 | 0.664245 | 1 | 1 | 2.88E-55 | U | RPS13 |
| RPL27A | 5.38E-57 | 0.518285 | 1 | 1 | 1.05E-52 | U | RPL27A |
| RPL18 | 1.47E-56 | 0.548587 | 1 | 1 | 2.86E-52 | U | RPL18 |
| RPS15A | 1.08E-54 | 0.561632 | 1 | 1 | 2.10E-50 | U | RPS15A |
| FTL | 1.24E-54 | 0.969292 | 1 | 1 | 2.41E-50 | U | FTL |
| RPL13A | 3.38E-54 | 0.439923 | 1 | 1 | 6.58E-50 | U | RPL13A |
| RPS19 | 4.92E-54 | 0.519729 | 1 | 1 | 9.58E-50 | U | RPS19 |
| HSP90AB1 | 2.60E-53 | 0.783016 | 1 | 1 | 5.06E-49 | U | HSP90AB1 |
| RPL24 | 1.30E-52 | 0.562987 | 1 | 0.998 | 2.53E-48 | U | RPL24 |
| RPL35A | 2.94E-51 | 0.576451 | 1 | 1 | 5.71E-47 | U | RPL35A |
| RPS27A | 1.14E-50 | 0.515569 | 1 | 1 | 2.22E-46 | U | RPS27A |
| RPLP2 | 5.69E-50 | 0.530019 | 1 | 1 | 1.11E-45 | U | RPLP2 |
| UBA52 | 1.85E-49 | 0.511546 | 1 | 0.999 | 3.60E-45 | U | UBA52 |
| RPL9 | 9.84E-48 | 0.47564 | 1 | 1 | 1.91E-43 | U | RPL9 |
| NEAT1 | 9.00E-47 | 1.234255 | 0.97 | 0.938 | 1.75E-42 | U | NEAT1 |
| GOLT1B | 1.08E-46 | 0.978221 | 0.84 | 0.511 | 2.11E-42 | U | GOLT1B |
| RPS18 | 1.06E-45 | 0.478238 | 1 | 1 | 2.06E-41 | U | RPS18 |
| RPL10 | 1.15E-45 | 0.447187 | 1 | 1 | 2.24E-41 | U | RPL10 |
| LAPTM4A | 2.28E-45 | 0.740927 | 0.982 | 0.928 | 4.44E-41 | U | LAPTM4A |
| MALAT1 | 3.52E-45 | 0.68865 | 1 | 1 | 6.85E-41 | U | MALAT1 |
| RPL36 | 5.97E-45 | 0.51038 | 1 | 1 | 1.16E-40 | U | RPL36 |
| RPL18A | 6.39E-45 | 0.429746 | 1 | 1 | 1.24E-40 | U | RPL18A |
| RPS17 | 1.45E-44 | 0.472121 | 1 | 1 | 2.81E-40 | U | RPS17 |
| RPL31 | 1.83E-44 | 0.578716 | 1 | 0.999 | 3.57E-40 | U | RPL31 |
| RPS3A | 2.02E-44 | 0.513028 | 1 | 1 | 3.93E-40 | U | RPS3A |
| RPL22 | 2.91E-44 | 0.615682 | 1 | 0.997 | 5.65E-40 | U | RPL22 |
| ZFAS1 | 4.84E-44 | 0.696954 | 0.994 | 0.977 | 9.41E-40 | U | ZFAS1 |
| RPL26 | 5.58E-43 | 0.746805 | 1 | 0.999 | 1.08E-38 | U | RPL26 |
| COX7A2L | 3.03E-42 | 0.552134 | 0.982 | 0.987 | 5.89E-38 | U | COX7A2L |
| RPL34 | 3.93E-42 | 0.492678 | 1 | 1 | 7.65E-38 | U | RPL34 |
| CAMLG | 5.99E-41 | 0.635164 | 0.964 | 0.886 | 1.16E-36 | U | CAMLG |
| NEU1 | 1.70E-40 | 0.889254 | 0.882 | 0.639 | 3.31E-36 | U | NEU1 |

| | | | | | | | |
|---|---|---|---|---|---|---|---|
| RPS11 | 3.53E-40 | 0.429378 | 1 | 0.999 | 6.86E-36 | U | RPS11 |
| RPS5 | 2.27E-39 | 0.458448 | 1 | 1 | 4.41E-35 | U | RPS5 |
| RPS21 | 9.45E-39 | 0.571206 | 0.982 | 0.986 | 1.84E-34 | U | RPS21 |
| RPS8 | 1.35E-38 | 0.421575 | 1 | 1 | 2.63E-34 | U | RPS8 |
| RPL6 | 5.95E-37 | 0.393958 | 0.994 | 1 | 1.16E-32 | U | RPL6 |
| RPS29 | 2.06E-36 | 0.456566 | 0.994 | 0.999 | 4.02E-32 | U | RPS29 |
| RPS23 | 3.97E-36 | 0.415894 | 1 | 1 | 7.72E-32 | U | RPS23 |
| RPL27 | 7.43E-36 | 0.451022 | 1 | 0.999 | 1.44E-31 | U | RPL27 |
| RPL32 | 1.03E-35 | 0.383427 | 1 | 1 | 2.00E-31 | U | RPL32 |
| RPL8 | 3.25E-35 | 0.333993 | 1 | 1 | 6.32E-31 | U | RPL8 |
| RPS7 | 4.49E-35 | 0.398194 | 1 | 1 | 8.74E-31 | U | RPS7 |
| ST13 | 3.76E-34 | 0.544349 | 0.982 | 0.953 | 7.31E-30 | U | ST13 |
| RPL39 | 8.56E-34 | 0.456413 | 0.994 | 0.997 | 1.67E-29 | U | RPL39 |
| NACA | 1.47E-33 | 0.383624 | 1 | 1 | 2.86E-29 | U | NACA |
| RPL35 | 1.95E-33 | 0.35884 | 1 | 1 | 3.79E-29 | U | RPL35 |
| RPS24 | 5.58E-33 | 0.705135 | 0.994 | 0.991 | 1.09E-28 | U | RPS24 |
| RSL24D1 | 4.46E-32 | 0.545924 | 0.959 | 0.941 | 8.68E-28 | U | RSL24D1 |
| SYF2 | 4.97E-32 | 0.577223 | 0.876 | 0.75 | 9.67E-28 | U | SYF2 |
| RPL30 | 5.59E-32 | 0.426014 | 1 | 1 | 1.09E-27 | U | RPL30 |
| RPL7 | 7.17E-32 | 0.422352 | 1 | 1 | 1.39E-27 | U | RPL7 |
| PTP4A1 | 1.66E-31 | 0.806383 | 0.923 | 0.893 | 3.22E-27 | U | PTP4A1 |
| RPL41 | 2.65E-31 | 0.329992 | 1 | 1 | 5.16E-27 | U | RPL41 |
| CKS2 | 3.24E-31 | 0.706219 | 0.917 | 0.774 | 6.31E-27 | U | CKS2 |
| RPL37A | 5.45E-31 | 0.379181 | 1 | 1 | 1.06E-26 | U | RPL37A |
| TOMM20 | 1.90E-30 | 0.555597 | 0.911 | 0.87 | 3.70E-26 | U | TOMM20 |
| RPS20 | 2.16E-30 | 0.389936 | 1 | 0.999 | 4.20E-26 | U | RPS20 |
| HNRNPC | 4.46E-30 | 0.51783 | 0.988 | 0.989 | 8.68E-26 | U | HNRNPC |
| UBE2B | 2.22E-29 | 0.64901 | 0.87 | 0.72 | 4.32E-25 | U | UBE2B |
| PIM3 | 6.69E-29 | 0.687922 | 0.716 | 0.436 | 1.30E-24 | U | PIM3 |
| SCD | 6.78E-29 | 0.810905 | 0.899 | 0.696 | 1.32E-24 | U | SCD |
| RPL14 | 9.51E-29 | 0.356993 | 1 | 0.999 | 1.85E-24 | U | RPL14 |
| RPL5 | 1.65E-28 | 0.394726 | 1 | 0.999 | 3.22E-24 | U | RPL5 |
| SELK | 5.52E-28 | 0.529273 | 0.976 | 0.914 | 1.07E-23 | U | SELK |
| NPC2 | 3.89E-27 | 0.56116 | 0.97 | 0.899 | 7.56E-23 | U | NPC2 |
| RPL36A | 5.92E-27 | 0.541112 | 0.976 | 0.946 | 1.15E-22 | U | RPL36A |
| TPT1 | 7.08E-27 | 0.44543 | 1 | 0.998 | 1.38E-22 | U | TPT1 |
| TUBA4A | 1.93E-26 | 0.4957 | 0.373 | 0.111 | 3.76E-22 | U | TUBA4A |
| RPL15 | 3.03E-26 | 0.281882 | 1 | 1 | 5.89E-22 | U | RPL15 |
| RPS15 | 4.45E-26 | 0.323729 | 1 | 1 | 8.65E-22 | U | RPS15 |
| MAP1LC3I | 5.54E-26 | 0.657958 | 0.935 | 0.859 | 1.08E-21 | U | MAP1LC3B |

| | | | | | | | |
|---|---|---|---|---|---|---|---|
| RPS28 | 6.24E-26 | 0.323101 | 1 | 0.999 | 1.21E-21 | U | RPS28 |
| RPLP0 | 1.30E-25 | 0.337008 | 1 | 1 | 2.54E-21 | U | RPLP0 |
| PHLDA2 | 1.42E-25 | 0.465929 | 0.391 | 0.12 | 2.75E-21 | U | PHLDA2 |
| SSR3 | 1.81E-25 | 0.549875 | 0.852 | 0.749 | 3.51E-21 | U | SSR3 |
| COMMD6 | 1.85E-25 | 0.366468 | 0.988 | 0.994 | 3.61E-21 | U | COMMD6 |
| RPL13 | 4.24E-25 | 0.285784 | 1 | 1 | 8.26E-21 | U | RPL13 |
| HIST1H2B | 8.62E-25 | 0.430893 | 0.361 | 0.109 | 1.68E-20 | U | HIST1H2BN |
| CCNB1IP1 | 1.03E-24 | 0.54309 | 0.811 | 0.673 | 2.00E-20 | U | CCNB1IP1 |
| CLK1 | 1.16E-24 | 0.603715 | 0.722 | 0.463 | 2.25E-20 | U | CLK1 |
| DNAJB9 | 2.05E-24 | 0.683902 | 0.822 | 0.716 | 3.98E-20 | U | DNAJB9 |
| RPL23 | 2.43E-24 | 0.325795 | 1 | 0.998 | 4.72E-20 | U | RPL23 |
| RPL17 | 3.22E-24 | 0.531295 | 0.935 | 0.887 | 6.27E-20 | U | RPL17 |
| EEF1A1 | 1.00E-23 | 0.300587 | 1 | 1 | 1.95E-19 | U | EEF1A1 |
| KRT10 | 1.28E-23 | 0.555224 | 0.935 | 0.921 | 2.49E-19 | U | KRT10 |
| IMPDH2 | 2.65E-23 | 0.486877 | 0.929 | 0.877 | 5.16E-19 | U | IMPDH2 |
| RPL37 | 2.53E-22 | 0.396578 | 1 | 0.998 | 4.92E-18 | U | RPL37 |
| COX7C | 3.60E-22 | 0.289251 | 1 | 1 | 7.01E-18 | U | COX7C |
| RPL38 | 5.96E-22 | 0.34758 | 1 | 0.997 | 1.16E-17 | U | RPL38 |
| C6orf48 | 6.49E-22 | 0.450698 | 0.964 | 0.925 | 1.26E-17 | U | C6orf48 |
| RPS14 | 1.17E-21 | 0.267748 | 1 | 1 | 2.27E-17 | U | RPS14 |
| HLA-B | 1.38E-21 | 0.61494 | 0.621 | 0.366 | 2.68E-17 | U | HLA-B |
| RAB3A | 1.65E-21 | 0.608372 | 0.746 | 0.501 | 3.21E-17 | U | RAB3A |
| EIF3E | 2.89E-21 | 0.452226 | 1 | 0.993 | 5.62E-17 | U | EIF3E |
| MED10 | 4.49E-21 | 0.515317 | 0.834 | 0.726 | 8.74E-17 | U | MED10 |
| ALDOA | 6.41E-21 | 0.385327 | 0.994 | 0.989 | 1.25E-16 | U | ALDOA |
| RALA | 2.24E-20 | 0.481916 | 0.763 | 0.61 | 4.35E-16 | U | RALA |
| SBDS | 3.73E-20 | 0.575066 | 0.746 | 0.629 | 7.25E-16 | U | SBDS |
| TOMM7 | 6.67E-20 | 0.421096 | 0.959 | 0.962 | 1.30E-15 | U | TOMM7 |
| GUK1 | 1.34E-19 | 0.409691 | 0.953 | 0.958 | 2.61E-15 | U | GUK1 |
| ATP5L | 1.50E-19 | 0.296916 | 0.994 | 0.998 | 2.92E-15 | U | ATP5L |
| SNRPD2 | 2.41E-19 | 0.356794 | 0.994 | 0.975 | 4.68E-15 | U | SNRPD2 |
| EEF1D | 2.51E-19 | 0.349074 | 0.994 | 0.981 | 4.87E-15 | U | EEF1D |
| P4HB | 2.76E-19 | 0.4977 | 0.941 | 0.906 | 5.36E-15 | U | P4HB |
| POLE3 | 1.62E-18 | 0.512753 | 0.763 | 0.63 | 3.15E-14 | U | POLE3 |
| RPL12 | 1.67E-18 | 0.323348 | 0.994 | 1 | 3.25E-14 | U | RPL12 |
| SLC25A3 | 2.60E-18 | 0.324523 | 0.988 | 0.994 | 5.07E-14 | U | SLC25A3 |
| SERP1 | 5.73E-18 | 0.426604 | 0.893 | 0.909 | 1.11E-13 | U | SERP1 |
| USMG5 | 7.78E-18 | 0.467046 | 0.893 | 0.859 | 1.51E-13 | U | USMG5 |
| CRYAB | 8.52E-18 | 1.645584 | 0.562 | 0.325 | 1.66E-13 | U | CRYAB |
| KLHL28 | 9.03E-18 | 0.476203 | 0.592 | 0.362 | 1.76E-13 | U | KLHL28 |

| | | | | | | | |
|---|---|---|---|---|---|---|---|
| ADM | 9.89E-18 | 0.488681 | 0.402 | 0.165 | 1.92E-13 | U | ADM |
| SLC2A3 | 1.81E-17 | 0.481986 | 0.497 | 0.259 | 3.51E-13 | U | SLC2A3 |
| SNU13 | 5.05E-17 | 0.401552 | 0.941 | 0.958 | 9.83E-13 | U | SNU13 |
| TCP1 | 1.55E-16 | 0.470298 | 0.876 | 0.857 | 3.01E-12 | U | TCP1 |
| CALU | 2.14E-16 | 0.440617 | 0.899 | 0.862 | 4.16E-12 | U | CALU |
| SNHG8 | 2.78E-16 | 0.642905 | 0.698 | 0.552 | 5.40E-12 | U | SNHG8 |
| GNAI3 | 2.82E-16 | 0.43445 | 0.811 | 0.818 | 5.48E-12 | U | GNAI3 |
| MORF4L2 | 5.48E-16 | 0.328302 | 0.982 | 0.997 | 1.07E-11 | U | MORF4L2 |
| GDF15 | 7.32E-16 | 1.132033 | 0.426 | 0.207 | 1.42E-11 | U | GDF15 |
| VDAC2 | 7.50E-16 | 0.308552 | 0.97 | 0.968 | 1.46E-11 | U | VDAC2 |
| EIF3F | 7.92E-16 | 0.277567 | 0.982 | 0.975 | 1.54E-11 | U | EIF3F |
| GFPT1 | 1.14E-15 | 0.50406 | 0.651 | 0.449 | 2.21E-11 | U | GFPT1 |
| H3F3B | 2.99E-15 | 0.387526 | 1 | 1 | 5.82E-11 | U | H3F3B |
| YIPF5 | 5.81E-15 | 0.460715 | 0.74 | 0.635 | 1.13E-10 | U | YIPF5 |
| ALKBH5 | 6.69E-15 | 0.407746 | 0.811 | 0.706 | 1.30E-10 | U | ALKBH5 |
| EIF3H | 7.18E-15 | 0.325897 | 0.941 | 0.95 | 1.40E-10 | U | EIF3H |
| RPS10 | 7.19E-15 | 0.287062 | 1 | 0.997 | 1.40E-10 | U | RPS10 |
| CDKN1A | 8.29E-15 | 0.409682 | 0.71 | 0.479 | 1.61E-10 | U | CDKN1A |
| RPS12 | 8.42E-15 | 0.269059 | 1 | 1 | 1.64E-10 | U | RPS12 |
| ZMAT3 | 1.10E-14 | 0.469862 | 0.598 | 0.412 | 2.14E-10 | U | ZMAT3 |
| MXD1 | 1.75E-14 | 0.43905 | 0.402 | 0.197 | 3.40E-10 | U | MXD1 |
| LAMTOR3 | 3.06E-14 | 0.491646 | 0.716 | 0.621 | 5.95E-10 | U | LAMTOR3 |
| IFRD1 | 3.13E-14 | 0.459942 | 0.74 | 0.647 | 6.10E-10 | U | IFRD1 |
| FABP3 | 5.26E-14 | 0.393047 | 0.308 | 0.123 | 1.02E-09 | U | FABP3 |
| CD63 | 7.44E-14 | 0.323934 | 0.994 | 0.984 | 1.45E-09 | U | CD63 |
| CTA-29F1 | 1.03E-13 | 0.408183 | 0.491 | 0.298 | 2.01E-09 | U | CTA-29F11.1 |
| B2M | 1.05E-13 | 0.519638 | 0.87 | 0.859 | 2.04E-09 | U | B2M |
| EPB41L4A | 1.75E-13 | 0.43684 | 0.74 | 0.676 | 3.41E-09 | U | EPB41L4A-AS1 |
| EIF4A2 | 1.75E-13 | 0.322008 | 0.976 | 0.993 | 3.41E-09 | U | EIF4A2 |
| DDIT4 | 2.63E-13 | 0.538991 | 0.876 | 0.813 | 5.11E-09 | U | DDIT4 |
| SRP54 | 3.03E-13 | 0.451258 | 0.799 | 0.684 | 5.89E-09 | U | SRP54 |
| SNRPB2 | 4.27E-13 | 0.35949 | 0.893 | 0.933 | 8.30E-09 | U | SNRPB2 |
| STX3 | 4.41E-13 | 0.45939 | 0.456 | 0.273 | 8.57E-09 | U | STX3 |
| PLCG2 | 5.55E-13 | 0.457806 | 0.225 | 0.075 | 1.08E-08 | U | PLCG2 |
| SAT1 | 5.74E-13 | 0.302856 | 0.959 | 0.899 | 1.12E-08 | U | SAT1 |
| DDX24 | 1.52E-12 | 0.454953 | 0.805 | 0.802 | 2.96E-08 | U | DDX24 |
| DCAF13 | 1.57E-12 | 0.436596 | 0.763 | 0.683 | 3.06E-08 | U | DCAF13 |
| SLC3A2 | 1.76E-12 | 0.403308 | 0.935 | 0.911 | 3.42E-08 | U | SLC3A2 |
| OST4 | 2.51E-12 | 0.278338 | 0.988 | 0.975 | 4.89E-08 | U | OST4 |
| CSTB | 4.12E-12 | 0.377654 | 0.888 | 0.918 | 8.02E-08 | U | CSTB |

| | | | | | | | |
|---|---|---|---|---|---|---|---|
| MLEC | 5.38E-12 | 0.392137 | 0.728 | 0.674 | 1.05E-07 | U | MLEC |
| TMEM38B | 5.66E-12 | 0.356111 | 0.728 | 0.613 | 1.10E-07 | U | TMEM38B |
| M6PR | 5.68E-12 | 0.459262 | 0.669 | 0.602 | 1.10E-07 | U | M6PR |
| MAFF | 9.42E-12 | 0.362471 | 0.408 | 0.223 | 1.83E-07 | U | MAFF |
| FNIP1 | 9.70E-12 | 0.40978 | 0.574 | 0.433 | 1.89E-07 | U | FNIP1 |
| ANKRD37 | 9.77E-12 | 0.439075 | 0.402 | 0.223 | 1.90E-07 | U | ANKRD37 |
| RABGGTB | 1.31E-11 | 0.420613 | 0.74 | 0.682 | 2.54E-07 | U | RABGGTB |
| TMED3 | 1.93E-11 | 0.380329 | 0.751 | 0.715 | 3.76E-07 | U | TMED3 |
| HSPA6 | 2.02E-11 | 0.475153 | 0.497 | 0.278 | 3.93E-07 | U | HSPA6 |
| TSPYL2 | 2.41E-11 | 0.382909 | 0.456 | 0.283 | 4.68E-07 | U | TSPYL2 |
| SAR1A | 2.45E-11 | 0.374092 | 0.71 | 0.636 | 4.76E-07 | U | SAR1A |
| ATP6V1E1 | 2.91E-11 | 0.416041 | 0.663 | 0.612 | 5.67E-07 | U | ATP6V1E1 |
| CNN1 | 4.40E-11 | 0.253923 | 0.213 | 0.075 | 8.56E-07 | U | CNN1 |
| JMJD6 | 5.68E-11 | 0.377058 | 0.515 | 0.347 | 1.10E-06 | U | JMJD6 |
| DDIT3 | 7.33E-11 | 0.406024 | 0.822 | 0.708 | 1.43E-06 | U | DDIT3 |
| HSP90AA1 | 8.09E-11 | 0.344193 | 0.982 | 0.995 | 1.57E-06 | U | HSP90AA1 |
| UQCRH | 9.01E-11 | 0.253549 | 0.982 | 0.981 | 1.75E-06 | U | UQCRH |
| SEC61B | 9.54E-11 | 0.297454 | 0.947 | 0.968 | 1.86E-06 | U | SEC61B |
| SCML1 | 1.05E-10 | 0.373979 | 0.497 | 0.338 | 2.04E-06 | U | SCML1 |
| ATP6V0B | 1.10E-10 | 0.312375 | 0.917 | 0.925 | 2.14E-06 | U | ATP6V0B |
| C1orf35 | 1.36E-10 | 0.397612 | 0.68 | 0.622 | 2.65E-06 | U | C1orf35 |
| CALCB | 1.41E-10 | 0.250557 | 0.26 | 0.107 | 2.75E-06 | U | CALCB |
| CHMP5 | 1.73E-10 | 0.324132 | 0.87 | 0.896 | 3.36E-06 | U | CHMP5 |
| MZT1 | 2.47E-10 | 0.345509 | 0.763 | 0.664 | 4.80E-06 | U | MZT1 |
| UBL5 | 2.53E-10 | 0.297154 | 0.994 | 0.983 | 4.91E-06 | U | UBL5 |
| GID8 | 2.96E-10 | 0.318381 | 0.633 | 0.53 | 5.77E-06 | U | GID8 |
| BHLHE40 | 3.44E-10 | 0.353128 | 0.367 | 0.201 | 6.70E-06 | U | BHLHE40 |
| LSM3 | 3.52E-10 | 0.374845 | 0.686 | 0.643 | 6.85E-06 | U | LSM3 |
| WBP5 | 3.60E-10 | 0.351278 | 0.864 | 0.857 | 7.01E-06 | U | WBP5 |
| PFDN2 | 4.58E-10 | 0.297957 | 0.87 | 0.879 | 8.91E-06 | U | PFDN2 |
| SIVA1 | 4.83E-10 | 0.358425 | 0.805 | 0.774 | 9.39E-06 | U | SIVA1 |
| ATP6V1G1 | 5.89E-10 | 0.284238 | 0.976 | 0.99 | 1.15E-05 | U | ATP6V1G1 |
| POLR2C | 6.18E-10 | 0.315426 | 0.787 | 0.758 | 1.20E-05 | U | POLR2C |
| TAF1D | 7.37E-10 | 0.399076 | 0.657 | 0.59 | 1.43E-05 | U | TAF1D |
| TIMM17A | 8.55E-10 | 0.345098 | 0.71 | 0.639 | 1.66E-05 | U | TIMM17A |
| RPL36AL | 8.93E-10 | 0.341663 | 0.864 | 0.899 | 1.74E-05 | U | RPL36AL |
| SIKE1 | 1.03E-09 | 0.355162 | 0.592 | 0.48 | 2.00E-05 | U | SIKE1 |
| LAMTOR5 | 1.12E-09 | 0.36133 | 0.935 | 0.949 | 2.19E-05 | U | LAMTOR5 |
| P4HA1 | 1.15E-09 | 0.392253 | 0.734 | 0.626 | 2.24E-05 | U | P4HA1 |
| HIST2H2B | 1.38E-09 | 0.469397 | 0.651 | 0.5 | 2.68E-05 | U | HIST2H2BE |

| | | | | | | | |
|---|---|---|---|---|---|---|---|
| A1BG | 1.38E-09 | 0.403028 | 0.657 | 0.591 | 2.69E-05 | U | A1BG |
| ZFAND2A | 1.40E-09 | 0.314285 | 0.562 | 0.404 | 2.73E-05 | U | ZFAND2A |
| PLEKHB2 | 1.49E-09 | 0.324768 | 0.692 | 0.607 | 2.89E-05 | U | PLEKHB2 |
| RP11-727I | 1.74E-09 | 0.263365 | 0.231 | 0.095 | 3.38E-05 | U | RP11-727F15.9 |
| COX14 | 1.97E-09 | 0.372209 | 0.728 | 0.676 | 3.82E-05 | U | COX14 |
| TIPRL | 2.33E-09 | 0.368605 | 0.675 | 0.636 | 4.53E-05 | U | TIPRL |
| LUM | 2.52E-09 | 0.306419 | 0.207 | 0.08 | 4.90E-05 | U | LUM |
| OAZ1 | 2.55E-09 | 0.268213 | 0.982 | 0.993 | 4.97E-05 | U | OAZ1 |
| MAP2K1 | 2.89E-09 | 0.313435 | 0.604 | 0.511 | 5.63E-05 | U | MAP2K1 |
| RIOK3 | 2.90E-09 | 0.359674 | 0.521 | 0.38 | 5.64E-05 | U | RIOK3 |
| ZNF622 | 3.57E-09 | 0.361537 | 0.574 | 0.483 | 6.94E-05 | U | ZNF622 |
| HSPA1B | 3.64E-09 | 0.312255 | 0.627 | 0.441 | 7.08E-05 | U | HSPA1B |
| CIB1 | 3.76E-09 | 0.32375 | 0.799 | 0.746 | 7.31E-05 | U | CIB1 |
| ZNF330 | 4.24E-09 | 0.399846 | 0.562 | 0.468 | 8.25E-05 | U | ZNF330 |
| SF3B6 | 4.65E-09 | 0.295143 | 0.888 | 0.888 | 9.05E-05 | U | SF3B6 |
| BZW1 | 4.85E-09 | 0.302464 | 0.87 | 0.907 | 9.43E-05 | U | BZW1 |
| SEC61G | 4.86E-09 | 0.292685 | 0.982 | 0.99 | 9.45E-05 | U | SEC61G |
| RAB11A | 7.00E-09 | 0.284914 | 0.84 | 0.869 | 0.000136 | U | RAB11A |
| CLK3 | 8.08E-09 | 0.297801 | 0.55 | 0.412 | 0.000157 | U | CLK3 |
| ALDOC | 8.17E-09 | 0.407221 | 0.58 | 0.443 | 0.000159 | U | ALDOC |
| ATP6V0D1 | 8.23E-09 | 0.374973 | 0.757 | 0.77 | 0.00016 | U | ATP6V0D1 |
| TAF13 | 9.37E-09 | 0.276444 | 0.29 | 0.149 | 0.000182 | U | TAF13 |
| DNAJC3 | 1.02E-08 | 0.364518 | 0.538 | 0.417 | 0.000198 | U | DNAJC3 |
| CTTN | 1.05E-08 | 0.409736 | 0.497 | 0.393 | 0.000204 | U | CTTN |
| VIMP | 1.08E-08 | 0.333404 | 0.781 | 0.763 | 0.000209 | U | VIMP |
| KLHL24 | 1.12E-08 | 0.259859 | 0.485 | 0.329 | 0.000218 | U | KLHL24 |
| CUTA | 1.27E-08 | 0.257469 | 0.982 | 0.985 | 0.000247 | U | CUTA |
| ENO2 | 1.34E-08 | 0.31341 | 0.858 | 0.833 | 0.000261 | U | ENO2 |
| ARL1 | 1.48E-08 | 0.289831 | 0.769 | 0.67 | 0.000289 | U | ARL1 |
| SEC24A | 1.53E-08 | 0.286802 | 0.355 | 0.211 | 0.000297 | U | SEC24A |
| OSER1 | 1.96E-08 | 0.346075 | 0.538 | 0.435 | 0.000381 | U | OSER1 |
| EIF3M | 2.09E-08 | 0.277652 | 0.87 | 0.889 | 0.000407 | U | EIF3M |
| ATP6V1D | 2.59E-08 | 0.415591 | 0.763 | 0.745 | 0.000504 | U | ATP6V1D |
| ARPP19 | 2.60E-08 | 0.305443 | 0.822 | 0.823 | 0.000506 | U | ARPP19 |
| PJA2 | 2.97E-08 | 0.381823 | 0.627 | 0.585 | 0.000578 | U | PJA2 |
| TMBIM6 | 3.20E-08 | 0.3158 | 0.941 | 0.966 | 0.000623 | U | TMBIM6 |
| CTD-2521 | 3.42E-08 | 1.701392 | 0.142 | 0.047 | 0.000664 | U | CTD-2521M24.8 |
| FAM96B | 4.10E-08 | 0.294829 | 0.893 | 0.91 | 0.000798 | U | FAM96B |
| MTCH1 | 4.56E-08 | 0.279625 | 0.917 | 0.917 | 0.000887 | U | MTCH1 |
| ASAH1 | 4.60E-08 | 0.409365 | 0.74 | 0.734 | 0.000894 | U | ASAH1 |

| | | | | | | | |
|---|---|---|---|---|---|---|---|
| METTL12 | 4.65E-08 | 0.709371 | 0.29 | 0.157 | 0.000905 | U | METTL12 |
| ICAM5 | 4.91E-08 | 0.282208 | 0.325 | 0.177 | 0.000954 | U | ICAM5 |
| TMED7 | 5.06E-08 | 0.322306 | 0.627 | 0.561 | 0.000985 | U | TMED7 |
| FAM177A1 | 5.24E-08 | 0.339785 | 0.734 | 0.705 | 0.00102 | U | FAM177A1 |
| DGUOK | 5.62E-08 | 0.275662 | 0.888 | 0.861 | 0.001093 | U | DGUOK |
| RNMT | 5.86E-08 | 0.401863 | 0.586 | 0.534 | 0.00114 | U | RNMT |
| PRRC1 | 6.59E-08 | 0.326423 | 0.497 | 0.382 | 0.001282 | U | PRRC1 |
| INSIG2 | 7.37E-08 | 0.424825 | 0.598 | 0.542 | 0.001434 | U | INSIG2 |
| OSTC | 7.60E-08 | 0.258905 | 0.888 | 0.893 | 0.001478 | U | OSTC |
| CHPF | 8.33E-08 | 0.290429 | 0.657 | 0.553 | 0.00162 | U | CHPF |
| ARHGDIA | 9.25E-08 | 0.324543 | 0.675 | 0.663 | 0.0018 | U | ARHGDIA |
| PSMC4 | 9.30E-08 | 0.299486 | 0.769 | 0.727 | 0.001809 | U | PSMC4 |
| PMEL | 1.26E-07 | 0.3283 | 0.503 | 0.335 | 0.00246 | U | PMEL |
| UHMK1 | 1.35E-07 | 0.348342 | 0.391 | 0.27 | 0.002622 | U | UHMK1 |
| NAPA | 1.38E-07 | 0.323076 | 0.722 | 0.685 | 0.002689 | U | NAPA |
| HSPA1A | 2.09E-07 | 0.30088 | 0.479 | 0.316 | 0.004058 | U | HSPA1A |
| LAMP2 | 2.38E-07 | 0.328225 | 0.562 | 0.484 | 0.004623 | U | LAMP2 |
| OAT | 2.87E-07 | 0.316777 | 0.586 | 0.535 | 0.005579 | U | OAT |
| SRPRA | 3.07E-07 | 0.313689 | 0.58 | 0.469 | 0.00598 | U | SRPRA |
| NXT1 | 4.45E-07 | 0.31634 | 0.592 | 0.538 | 0.008664 | U | NXT1 |
| NUDT21 | 4.48E-07 | 0.300855 | 0.604 | 0.542 | 0.008708 | U | NUDT21 |
| XAB2 | 4.99E-07 | 0.319773 | 0.509 | 0.418 | 0.009704 | U | XAB2 |
| LAGE3 | 6.05E-07 | 0.315023 | 0.769 | 0.717 | 0.011777 | U | LAGE3 |
| SPTY2D1 | 6.39E-07 | 0.331326 | 0.438 | 0.325 | 0.012426 | U | SPTY2D1 |
| SCNM1 | 7.52E-07 | 0.271072 | 0.556 | 0.47 | 0.014626 | U | SCNM1 |
| RECQL | 7.90E-07 | 0.442581 | 0.367 | 0.242 | 0.015377 | U | RECQL |
| DAD1 | 8.68E-07 | 0.304736 | 0.911 | 0.947 | 0.016894 | U | DAD1 |
| NSA2 | 8.75E-07 | 0.272806 | 0.663 | 0.648 | 0.017014 | U | NSA2 |
| HSPA9 | 1.09E-06 | 0.289559 | 0.805 | 0.826 | 0.021282 | U | HSPA9 |
| NUP58 | 1.15E-06 | 0.275464 | 0.467 | 0.354 | 0.022308 | U | NUP58 |
| SCAMP3 | 1.70E-06 | 0.281073 | 0.604 | 0.521 | 0.033015 | U | SCAMP3 |
| SYAP1 | 1.72E-06 | 0.250164 | 0.751 | 0.784 | 0.033544 | U | SYAP1 |
| HSPB1 | 1.85E-06 | 0.298269 | 0.947 | 0.93 | 0.035911 | U | HSPB1 |
| IDI1 | 1.89E-06 | 0.44826 | 0.799 | 0.823 | 0.036733 | U | IDI1 |
| TMEM50B | 1.92E-06 | 0.295444 | 0.615 | 0.577 | 0.037345 | U | TMEM50B |
| SUCO | 2.69E-06 | 0.297177 | 0.479 | 0.385 | 0.05236 | U | SUCO |
| COG3 | 2.77E-06 | 0.288132 | 0.302 | 0.19 | 0.053979 | U | COG3 |
| HIST1H1C | 3.35E-06 | 0.484688 | 0.503 | 0.418 | 0.065163 | U | HIST1H1C |
| DDX18 | 3.44E-06 | 0.319269 | 0.663 | 0.628 | 0.066897 | U | DDX18 |
| GADD45B | 3.63E-06 | 0.349705 | 0.503 | 0.389 | 0.070522 | U | GADD45B |

| | | | | | | | |
|---|---|---|---|---|---|---|---|
| THOC3 | 3.67E-06 | 0.265117 | 0.663 | 0.625 | 0.07148 | U | THOC3 |
| HIST3H2A | 4.19E-06 | 0.34746 | 0.533 | 0.446 | 0.081543 | U | HIST3H2A |
| CAP1 | 5.06E-06 | 0.263247 | 0.746 | 0.783 | 0.098528 | U | CAP1 |
| PCMT1 | 6.14E-06 | 0.28684 | 0.74 | 0.825 | 0.11935 | U | PCMT1 |
| NAMPT | 6.82E-06 | 0.337596 | 0.521 | 0.449 | 0.132596 | U | NAMPT |
| RAB6A | 7.44E-06 | 0.254685 | 0.728 | 0.731 | 0.144821 | U | RAB6A |
| PPP1R15A | 7.72E-06 | 0.298301 | 0.686 | 0.687 | 0.150193 | U | PPP1R15A |
| UPP1 | 7.91E-06 | 0.345802 | 0.32 | 0.207 | 0.1539 | U | UPP1 |
| HSPH1 | 9.00E-06 | 0.272427 | 0.645 | 0.55 | 0.175101 | U | HSPH1 |
| G3BP2 | 1.05E-05 | 0.297302 | 0.556 | 0.52 | 0.204503 | U | G3BP2 |
| SRRM1 | 1.09E-05 | 0.276468 | 0.751 | 0.775 | 0.211952 | U | SRRM1 |
| SNHG21 | 1.17E-05 | 0.299266 | 0.325 | 0.224 | 0.227408 | U | SNHG21 |
| DDX21 | 1.28E-05 | 0.26883 | 0.621 | 0.576 | 0.248733 | U | DDX21 |
| SAR1B | 1.44E-05 | 0.271602 | 0.609 | 0.54 | 0.280085 | U | SAR1B |
| ANXA5 | 1.52E-05 | 0.260031 | 0.663 | 0.681 | 0.296099 | U | ANXA5 |
| VAT1 | 1.56E-05 | 0.312777 | 0.58 | 0.546 | 0.302925 | U | VAT1 |
| VGF | 1.78E-05 | 0.309287 | 0.207 | 0.106 | 0.347084 | U | VGF |
| NRBP1 | 2.02E-05 | 0.306331 | 0.615 | 0.656 | 0.393472 | U | NRBP1 |
| C1orf52 | 2.24E-05 | 0.273504 | 0.592 | 0.564 | 0.434865 | U | C1orf52 |
| YKT6 | 2.25E-05 | 0.276029 | 0.456 | 0.371 | 0.437013 | U | YKT6 |
| PGM3 | 2.32E-05 | 0.255807 | 0.473 | 0.386 | 0.450946 | U | PGM3 |
| LCMT1 | 2.47E-05 | 0.297758 | 0.669 | 0.71 | 0.479889 | U | LCMT1 |
| C12orf49 | 2.48E-05 | 0.294874 | 0.367 | 0.274 | 0.482781 | U | C12orf49 |
| BCAS2 | 2.48E-05 | 0.269468 | 0.609 | 0.612 | 0.483343 | U | BCAS2 |
| STARD4 | 2.52E-05 | 0.266733 | 0.467 | 0.392 | 0.48998 | U | STARD4 |
| RAB18 | 2.83E-05 | 0.27823 | 0.728 | 0.779 | 0.550703 | U | RAB18 |
| MRPL10 | 2.88E-05 | 0.25964 | 0.598 | 0.61 | 0.55984 | U | MRPL10 |
| SEC31A | 3.10E-05 | 0.263085 | 0.621 | 0.601 | 0.603596 | U | SEC31A |
| SMNDC1 | 3.80E-05 | 0.284675 | 0.497 | 0.448 | 0.739442 | U | SMNDC1 |
| ARRDC3 | 4.13E-05 | 0.324102 | 0.615 | 0.565 | 0.804394 | U | ARRDC3 |
| TMEM87A | 4.47E-05 | 0.290772 | 0.562 | 0.532 | 0.869948 | U | TMEM87A |
| SELM | 4.80E-05 | 0.29557 | 0.71 | 0.683 | 0.933622 | U | SELM |
| SPSB3 | 6.77E-05 | 0.274758 | 0.574 | 0.553 | 1 | U | SPSB3 |
| SQSTM1 | 7.25E-05 | 0.270999 | 0.669 | 0.69 | 1 | U | SQSTM1 |
| NR4A1 | 7.65E-05 | 0.257847 | 0.266 | 0.171 | 1 | U | NR4A1 |
| SDCBP | 8.24E-05 | 0.290977 | 0.864 | 0.923 | 1 | U | SDCBP |
| TBC1D15 | 0.000102 | 0.253875 | 0.396 | 0.317 | 1 | U | TBC1D15 |
| TMEM183 | 0.000104 | 0.257735 | 0.592 | 0.601 | 1 | U | TMEM183A |
| SNAPC1 | 0.000114 | 0.445943 | 0.467 | 0.404 | 1 | U | SNAPC1 |
| JUNB | 0.000117 | 0.366376 | 0.58 | 0.529 | 1 | U | JUNB |

| | | | | | | | |
|---|---|---|---|---|---|---|---|
| DPM1 | 0.000147 | 0.270958 | 0.598 | 0.617 | 1 | U | DPM1 |
| NOLC1 | 0.000149 | 0.278989 | 0.497 | 0.442 | 1 | U | NOLC1 |
| BUD31 | 0.000194 | 0.264126 | 0.787 | 0.857 | 1 | U | BUD31 |
| LGALS1 | 0.000203 | 0.310762 | 0.331 | 0.239 | 1 | U | LGALS1 |
| INSIG1 | 0.000233 | 0.298243 | 0.639 | 0.598 | 1 | U | INSIG1 |
| EIF5B | 0.000246 | 0.284722 | 0.686 | 0.766 | 1 | U | EIF5B |
| PNPLA8 | 0.000256 | 0.284929 | 0.503 | 0.469 | 1 | U | PNPLA8 |
| STARD3N | 0.000277 | 0.286257 | 0.74 | 0.768 | 1 | U | STARD3NL |
| HMGCS1 | 0.00032 | 0.327261 | 0.805 | 0.817 | 1 | U | HMGCS1 |
| NMRK2 | 0.00032 | 0.278756 | 0.308 | 0.219 | 1 | U | NMRK2 |
| RRAGD | 0.000329 | 0.252031 | 0.355 | 0.283 | 1 | U | RRAGD |
| ATG101 | 0.00034 | 0.255489 | 0.574 | 0.555 | 1 | U | ATG101 |
| ERO1A | 0.000581 | 0.255115 | 0.373 | 0.298 | 1 | U | ERO1A |
| PHF1 | 0.000602 | 0.255961 | 0.432 | 0.374 | 1 | U | PHF1 |
| GNL3 | 0.000658 | 0.281677 | 0.55 | 0.551 | 1 | U | GNL3 |
| GPI | 0.000686 | 0.270215 | 0.639 | 0.665 | 1 | U | GPI |
| SPTSSA | 0.000811 | 0.256533 | 0.692 | 0.739 | 1 | U | SPTSSA |
| TIMM44 | 0.000895 | 0.250558 | 0.527 | 0.502 | 1 | U | TIMM44 |
| RPL22L1 | 0.000979 | 0.277228 | 0.58 | 0.564 | 1 | U | RPL22L1 |
| CIR1 | 0.002641 | 0.255676 | 0.45 | 0.435 | 1 | U | CIR1 |
| HIST1H2B | 0.003103 | 0.290687 | 0.254 | 0.187 | 1 | U | HIST1H2BD |
| MDH1 | 0.003303 | 0.297857 | 0.817 | 0.894 | 1 | U | MDH1 |
| SYP | 0.005492 | 1.071435 | 0.083 | 0.173 | 1 | U | SYP |
| P4HB1 | 3.87E-190 | 1.619696 | 0.998 | 0.887 | 7.52E-186 | INF/MLN | P4HB |
| ARF4 | 2.43E-183 | 1.357702 | 1 | 0.953 | 4.72E-179 | INF/MLN | ARF4 |
| MPC1 | 6.31E-155 | 1.066277 | 0.971 | 0.668 | 1.23E-150 | INF/MLN | MPC1 |
| FTL1 | 1.49E-152 | 1.078796 | 1 | 1 | 2.91E-148 | INF/MLN | FTL |
| KDELR2 | 1.52E-152 | 0.865136 | 0.991 | 0.88 | 2.96E-148 | INF/MLN | KDELR2 |
| SERP11 | 7.34E-150 | 0.852628 | 0.989 | 0.889 | 1.43E-145 | INF/MLN | SERP1 |
| YIPF2 | 3.17E-148 | 0.821875 | 0.938 | 0.462 | 6.16E-144 | INF/MLN | YIPF2 |
| SRP541 | 4.96E-148 | 0.915295 | 0.975 | 0.626 | 9.64E-144 | INF/MLN | SRP54 |
| CALU1 | 8.15E-139 | 0.792892 | 0.996 | 0.833 | 1.58E-134 | INF/MLN | CALU |
| ARF1 | 1.44E-125 | 0.612859 | 0.991 | 0.954 | 2.80E-121 | INF/MLN | ARF1 |
| ICAM51 | 2.12E-125 | 0.500957 | 0.578 | 0.096 | 4.12E-121 | INF/MLN | ICAM5 |
| ALDOA1 | 1.52E-124 | 0.771309 | 1 | 0.986 | 2.96E-120 | INF/MLN | ALDOA |
| ARL11 | 4.20E-123 | 0.694141 | 0.962 | 0.61 | 8.17E-119 | INF/MLN | ARL1 |
| NPRL2 | 7.82E-122 | 0.735533 | 0.85 | 0.372 | 1.52E-117 | INF/MLN | NPRL2 |
| RCN1 | 1.67E-120 | 0.633428 | 1 | 0.899 | 3.26E-116 | INF/MLN | RCN1 |
| TRMT112 | 2.75E-114 | 0.57288 | 1 | 0.957 | 5.34E-110 | INF/MLN | TRMT112 |
| ALKBH51 | 3.88E-114 | 0.886918 | 0.935 | 0.662 | 7.56E-110 | INF/MLN | ALKBH5 |

| | | | | | | | |
|---|---|---|---|---|---|---|---|
| MFSD2A | 6.99E-113 | 0.539996 | 0.67 | 0.167 | 1.36E-108 | INF/MLN | MFSD2A |
| SEC61G1 | 1.11E-112 | 0.582696 | 0.996 | 0.988 | 2.16E-108 | INF/MLN | SEC61G |
| LAPTM4A | 2.96E-111 | 0.658129 | 0.998 | 0.916 | 5.76E-107 | INF/MLN | LAPTM4A |
| AIF1 | 1.07E-110 | 0.728669 | 0.627 | 0.159 | 2.09E-106 | INF/MLN | AIF1 |
| CIB11 | 2.05E-110 | 0.663132 | 0.971 | 0.697 | 3.99E-106 | INF/MLN | CIB1 |
| MAGEH1 | 2.83E-110 | 0.719677 | 0.971 | 0.716 | 5.51E-106 | INF/MLN | MAGEH1 |
| DAD11 | 1.18E-108 | 0.577933 | 0.991 | 0.933 | 2.29E-104 | INF/MLN | DAD1 |
| CD631 | 7.05E-108 | 0.571732 | 1 | 0.981 | 1.37E-103 | INF/MLN | CD63 |
| HSPA91 | 9.32E-108 | 0.800922 | 0.971 | 0.79 | 1.81E-103 | INF/MLN | HSPA9 |
| FAU1 | 2.73E-105 | 0.392478 | 1 | 1 | 5.32E-101 | INF/MLN | FAU |
| CREB3 | 1.20E-101 | 0.600299 | 0.908 | 0.53 | 2.34E-97 | INF/MLN | CREB3 |
| RPL181 | 3.03E-101 | 0.421594 | 1 | 0.999 | 5.90E-97 | INF/MLN | RPL18 |
| MAP1LC3I | 1.19E-100 | 0.806297 | 0.978 | 0.838 | 2.32E-96 | INF/MLN | MAP1LC3B |
| HERPUD1 | 4.27E-100 | 0.874696 | 0.978 | 0.778 | 8.31E-96 | INF/MLN | HERPUD1 |
| SEC13 | 3.53E-99 | 0.632216 | 0.935 | 0.704 | 6.87E-95 | INF/MLN | SEC13 |
| TRIB3 | 6.62E-99 | 0.740112 | 0.612 | 0.166 | 1.29E-94 | INF/MLN | TRIB3 |
| MED101 | 1.05E-98 | 0.619501 | 0.951 | 0.683 | 2.04E-94 | INF/MLN | MED10 |
| DCAF131 | 2.68E-98 | 0.686253 | 0.933 | 0.631 | 5.21E-94 | INF/MLN | DCAF13 |
| SRPRA1 | 7.16E-97 | 0.564718 | 0.844 | 0.39 | 1.39E-92 | INF/MLN | SRPRA |
| DNAJB2 | 7.11E-96 | 0.637018 | 0.904 | 0.555 | 1.38E-91 | INF/MLN | DNAJB2 |
| CHPF1 | 1.01E-94 | 0.595531 | 0.886 | 0.484 | 1.97E-90 | INF/MLN | CHPF |
| DDIT41 | 5.21E-94 | 0.822943 | 0.982 | 0.779 | 1.01E-89 | INF/MLN | DDIT4 |
| EIF11 | 1.14E-93 | 0.452457 | 1 | 1 | 2.21E-89 | INF/MLN | EIF1 |
| GOLT1B1 | 3.14E-93 | 0.575135 | 0.875 | 0.454 | 6.12E-89 | INF/MLN | GOLT1B |
| RPS161 | 4.90E-93 | 0.397849 | 1 | 0.999 | 9.53E-89 | INF/MLN | RPS16 |
| TPT11 | 1.90E-92 | 0.550656 | 1 | 0.998 | 3.69E-88 | INF/MLN | TPT1 |
| AURKAIP1 | 5.96E-92 | 0.570924 | 0.975 | 0.881 | 1.16E-87 | INF/MLN | AURKAIP1 |
| GFPT11 | 1.24E-91 | 0.561896 | 0.817 | 0.38 | 2.41E-87 | INF/MLN | GFPT1 |
| SND1 | 1.44E-91 | 0.598027 | 0.906 | 0.611 | 2.81E-87 | INF/MLN | SND1 |
| TCP11 | 9.49E-91 | 0.607077 | 0.975 | 0.831 | 1.85E-86 | INF/MLN | TCP1 |
| SSR4 | 2.15E-90 | 0.483324 | 0.989 | 0.944 | 4.18E-86 | INF/MLN | SSR4 |
| ICAM3 | 4.85E-89 | 0.382586 | 0.484 | 0.096 | 9.43E-85 | INF/MLN | ICAM3 |
| JMJD61 | 1.65E-88 | 0.525653 | 0.73 | 0.271 | 3.21E-84 | INF/MLN | JMJD6 |
| LAMTOR5 | 2.10E-88 | 0.537606 | 0.989 | 0.938 | 4.09E-84 | INF/MLN | LAMTOR5 |
| C11orf24 | 6.30E-88 | 0.454994 | 0.679 | 0.222 | 1.23E-83 | INF/MLN | C11orf24 |
| HSPA5 | 2.81E-86 | 0.940077 | 0.993 | 0.906 | 5.47E-82 | INF/MLN | HSPA5 |
| ICAM1 | 3.63E-86 | 0.354348 | 0.402 | 0.061 | 7.06E-82 | INF/MLN | ICAM1 |
| RPS51 | 3.77E-86 | 0.394565 | 1 | 0.999 | 7.34E-82 | INF/MLN | RPS5 |
| POLR2C1 | 1.15E-85 | 0.572238 | 0.942 | 0.717 | 2.23E-81 | INF/MLN | POLR2C |
| MORF4L2 | 1.32E-85 | 0.446382 | 1 | 0.995 | 2.57E-81 | INF/MLN | MORF4L2 |

| | | | | | | | |
|---|---|---|---|---|---|---|---|
| BNIP3 | 5.63E-85 | 0.673627 | 0.989 | 0.817 | 1.10E-80 | INF/MLN | BNIP3 |
| FAM177A1 | 1.20E-84 | 0.571744 | 0.922 | 0.657 | 2.33E-80 | INF/MLN | FAM177A1 |
| YIF1A | 2.33E-84 | 0.671043 | 0.926 | 0.678 | 4.54E-80 | INF/MLN | YIF1A |
| RER1 | 3.20E-84 | 0.588761 | 0.913 | 0.629 | 6.23E-80 | INF/MLN | RER1 |
| SLC30A5 | 3.67E-84 | 0.497302 | 0.779 | 0.343 | 7.13E-80 | INF/MLN | SLC30A5 |
| RPL27A1 | 5.31E-84 | 0.334179 | 1 | 0.999 | 1.03E-79 | INF/MLN | RPL27A |
| PGK1 | 8.34E-84 | 0.546866 | 0.996 | 0.929 | 1.62E-79 | INF/MLN | PGK1 |
| CDKN1A1 | 9.86E-84 | 0.841728 | 0.848 | 0.413 | 1.92E-79 | INF/MLN | CDKN1A |
| HLA-B1 | 1.12E-83 | 0.521002 | 0.752 | 0.298 | 2.18E-79 | INF/MLN | HLA-B |
| DDIT31 | 2.78E-81 | 0.774429 | 0.94 | 0.664 | 5.41E-77 | INF/MLN | DDIT3 |
| PSMD8 | 2.90E-80 | 0.475692 | 0.984 | 0.927 | 5.64E-76 | INF/MLN | PSMD8 |
| VIMP1 | 3.33E-80 | 0.544066 | 0.955 | 0.72 | 6.47E-76 | INF/MLN | VIMP |
| ATP5B | 5.47E-79 | 0.403599 | 1 | 0.995 | 1.06E-74 | INF/MLN | ATP5B |
| KRT101 | 9.71E-79 | 0.536501 | 0.975 | 0.909 | 1.89E-74 | INF/MLN | KRT10 |
| EPRS | 1.88E-78 | 0.534033 | 0.855 | 0.463 | 3.66E-74 | INF/MLN | EPRS |
| TIMP1 | 2.23E-78 | 0.686269 | 0.922 | 0.694 | 4.33E-74 | INF/MLN | TIMP1 |
| SLC31A1 | 4.88E-78 | 0.48101 | 0.737 | 0.317 | 9.50E-74 | INF/MLN | SLC31A1 |
| SYAP11 | 1.17E-77 | 0.560821 | 0.94 | 0.745 | 2.28E-73 | INF/MLN | SYAP1 |
| CYCS | 2.27E-77 | 0.57215 | 0.984 | 0.926 | 4.42E-73 | INF/MLN | CYCS |
| TM9SF2 | 3.22E-77 | 0.569007 | 0.9 | 0.624 | 6.26E-73 | INF/MLN | TM9SF2 |
| SERPINH1 | 1.73E-76 | 0.67577 | 0.938 | 0.675 | 3.36E-72 | INF/MLN | SERPINH1 |
| PLPP5 | 2.00E-76 | 0.515184 | 0.746 | 0.341 | 3.89E-72 | INF/MLN | PLPP5 |
| PSMC41 | 4.25E-76 | 0.517097 | 0.935 | 0.682 | 8.27E-72 | INF/MLN | PSMC4 |
| THOC31 | 4.65E-76 | 0.514258 | 0.886 | 0.567 | 9.05E-72 | INF/MLN | THOC3 |
| SNU131 | 9.94E-75 | 0.459821 | 0.989 | 0.949 | 1.93E-70 | INF/MLN | SNU13 |
| NEAT11 | 1.23E-73 | 0.788647 | 0.996 | 0.927 | 2.39E-69 | INF/MLN | NEAT1 |
| EAF2 | 2.64E-73 | 0.272315 | 0.424 | 0.085 | 5.14E-69 | INF/MLN | EAF2 |
| NPW | 3.34E-73 | 0.601532 | 0.48 | 0.126 | 6.50E-69 | INF/MLN | NPW |
| NEU11 | 4.29E-73 | 0.57598 | 0.893 | 0.601 | 8.35E-69 | INF/MLN | NEU1 |
| PDXDC1 | 6.33E-73 | 0.479294 | 0.75 | 0.351 | 1.23E-68 | INF/MLN | PDXDC1 |
| DNTTIP1 | 1.53E-72 | 0.503419 | 0.868 | 0.578 | 2.97E-68 | INF/MLN | DNTTIP1 |
| PSMD13 | 2.51E-72 | 0.513834 | 0.935 | 0.755 | 4.88E-68 | INF/MLN | PSMD13 |
| SCD1 | 4.91E-72 | 1.032346 | 0.92 | 0.661 | 9.55E-68 | INF/MLN | SCD |
| OCIAD1 | 8.68E-72 | 0.526138 | 0.96 | 0.819 | 1.69E-67 | INF/MLN | OCIAD1 |
| PHLDA21 | 6.70E-71 | 0.410796 | 0.402 | 0.078 | 1.30E-66 | INF/MLN | PHLDA2 |
| PSMA5 | 1.52E-70 | 0.53851 | 0.938 | 0.727 | 2.96E-66 | INF/MLN | PSMA5 |
| RNH1 | 4.06E-70 | 0.478962 | 0.933 | 0.726 | 7.89E-66 | INF/MLN | RNH1 |
| ECE2 | 6.00E-70 | 0.313299 | 0.504 | 0.132 | 1.17E-65 | INF/MLN | ECE2 |
| SLC16A3 | 1.47E-69 | 0.503899 | 0.766 | 0.343 | 2.85E-65 | INF/MLN | SLC16A3 |
| SPTY2D11 | 2.92E-69 | 0.410177 | 0.674 | 0.253 | 5.68E-65 | INF/MLN | SPTY2D1 |

| | | | | | | | |
|---|---|---|---|---|---|---|---|
| YIPF51 | 3.66E-69 | 0.499029 | 0.886 | 0.585 | 7.11E-65 | INF/MLN | YIPF5 |
| MRPL3 | 4.01E-69 | 0.495332 | 0.917 | 0.681 | 7.79E-65 | INF/MLN | MRPL3 |
| SURF4 | 4.24E-69 | 0.492747 | 0.875 | 0.564 | 8.26E-65 | INF/MLN | SURF4 |
| DEDD2 | 4.33E-69 | 0.48443 | 0.763 | 0.368 | 8.42E-65 | INF/MLN | DEDD2 |
| GUK11 | 1.91E-68 | 0.420636 | 0.989 | 0.95 | 3.71E-64 | INF/MLN | GUK1 |
| COX7A2L1 | 2.36E-68 | 0.385608 | 1 | 0.984 | 4.58E-64 | INF/MLN | COX7A2L |
| COPB2 | 2.40E-68 | 0.513846 | 0.868 | 0.563 | 4.67E-64 | INF/MLN | COPB2 |
| MDH11 | 1.44E-67 | 0.576353 | 0.962 | 0.871 | 2.79E-63 | INF/MLN | MDH1 |
| ARPC3 | 1.84E-67 | 0.430373 | 0.978 | 0.905 | 3.59E-63 | INF/MLN | ARPC3 |
| GPX4 | 2.54E-67 | 0.346137 | 1 | 0.991 | 4.95E-63 | INF/MLN | GPX4 |
| SELM1 | 6.14E-67 | 0.529624 | 0.913 | 0.632 | 1.19E-62 | INF/MLN | SELM |
| IMPDH21 | 6.63E-67 | 0.515738 | 0.971 | 0.86 | 1.29E-62 | INF/MLN | IMPDH2 |
| COQ10B | 7.92E-67 | 0.466018 | 0.772 | 0.393 | 1.54E-62 | INF/MLN | COQ10B |
| SEC61A1 | 1.92E-66 | 0.472144 | 0.855 | 0.522 | 3.74E-62 | INF/MLN | SEC61A1 |
| SRPRB | 2.05E-66 | 0.437245 | 0.801 | 0.441 | 3.99E-62 | INF/MLN | SRPRB |
| TMBIM61 | 1.41E-65 | 0.405499 | 0.998 | 0.957 | 2.75E-61 | INF/MLN | TMBIM6 |
| GORASP2 | 1.73E-65 | 0.473293 | 0.904 | 0.636 | 3.36E-61 | INF/MLN | GORASP2 |
| RALA1 | 2.05E-65 | 0.604618 | 0.857 | 0.566 | 3.99E-61 | INF/MLN | RALA |
| STX4 | 1.96E-64 | 0.418898 | 0.647 | 0.264 | 3.81E-60 | INF/MLN | STX4 |
| SSSCA1 | 2.04E-64 | 0.505276 | 0.799 | 0.453 | 3.98E-60 | INF/MLN | SSSCA1 |
| STC2 | 2.44E-64 | 0.456029 | 0.554 | 0.181 | 4.75E-60 | INF/MLN | STC2 |
| VDAC21 | 3.94E-64 | 0.393344 | 0.993 | 0.962 | 7.67E-60 | INF/MLN | VDAC2 |
| SYT13 | 4.26E-64 | 0.276078 | 0.426 | 0.099 | 8.29E-60 | INF/MLN | SYT13 |
| MZT11 | 4.45E-64 | 0.453186 | 0.92 | 0.613 | 8.66E-60 | INF/MLN | MZT1 |
| INSIG21 | 3.30E-63 | 0.520849 | 0.833 | 0.478 | 6.42E-59 | INF/MLN | INSIG2 |
| LGALS3 | 4.90E-63 | 0.420775 | 0.426 | 0.104 | 9.53E-59 | INF/MLN | LGALS3 |
| SEC61B1 | 6.69E-63 | 0.38854 | 0.989 | 0.962 | 1.30E-58 | INF/MLN | SEC61B |
| SELK1 | 9.14E-63 | 0.473216 | 0.98 | 0.904 | 1.78E-58 | INF/MLN | SELK |
| NANS | 1.33E-62 | 0.431566 | 0.783 | 0.423 | 2.60E-58 | INF/MLN | NANS |
| MYDGF | 1.45E-62 | 0.464649 | 0.942 | 0.762 | 2.82E-58 | INF/MLN | MYDGF |
| NARS | 2.01E-62 | 0.470511 | 0.942 | 0.731 | 3.91E-58 | INF/MLN | NARS |
| PIM31 | 2.39E-62 | 0.514707 | 0.761 | 0.384 | 4.65E-58 | INF/MLN | PIM3 |
| SEC31A1 | 2.57E-62 | 0.431484 | 0.864 | 0.54 | 5.00E-58 | INF/MLN | SEC31A |
| SCAMP31 | 2.66E-62 | 0.451455 | 0.81 | 0.46 | 5.18E-58 | INF/MLN | SCAMP3 |
| FAM162A | 4.84E-62 | 0.531686 | 0.951 | 0.724 | 9.42E-58 | INF/MLN | FAM162A |
| NAPA1 | 6.70E-62 | 0.473191 | 0.9 | 0.637 | 1.30E-57 | INF/MLN | NAPA |
| YKT61 | 7.09E-62 | 0.384849 | 0.699 | 0.302 | 1.38E-57 | INF/MLN | YKT6 |
| RPS131 | 1.25E-61 | 0.301504 | 1 | 0.999 | 2.43E-57 | INF/MLN | RPS13 |
| SLC1A5 | 1.72E-61 | 0.418562 | 0.65 | 0.267 | 3.35E-57 | INF/MLN | SLC1A5 |
| POLE31 | 2.10E-61 | 0.467815 | 0.897 | 0.579 | 4.08E-57 | INF/MLN | POLE3 |

| | | | | | | | |
|---|---|---|---|---|---|---|---|
| GADD45B | 5.01E-61 | 0.566881 | 0.71 | 0.323 | 9.75E-57 | INF/MLN | GADD45B |
| CAMLG1 | 1.78E-60 | 0.427474 | 0.973 | 0.873 | 3.45E-56 | INF/MLN | CAMLG |
| PTGDS | 2.40E-60 | 0.871738 | 0.569 | 0.213 | 4.67E-56 | INF/MLN | PTGDS |
| POLR3D | 2.62E-60 | 0.382373 | 0.712 | 0.317 | 5.09E-56 | INF/MLN | POLR3D |
| TMEM208 | 3.30E-60 | 0.459403 | 0.911 | 0.671 | 6.42E-56 | INF/MLN | TMEM208 |
| MLF2 | 1.09E-59 | 0.433407 | 0.98 | 0.91 | 2.13E-55 | INF/MLN | MLF2 |
| CCT6A | 1.48E-59 | 0.418183 | 0.967 | 0.835 | 2.88E-55 | INF/MLN | CCT6A |
| PGM31 | 2.26E-59 | 0.401588 | 0.703 | 0.319 | 4.39E-55 | INF/MLN | PGM3 |
| CEBPB | 2.41E-59 | 0.388736 | 0.547 | 0.191 | 4.69E-55 | INF/MLN | CEBPB |
| GOLGA2 | 3.34E-59 | 0.397236 | 0.739 | 0.352 | 6.50E-55 | INF/MLN | GOLGA2 |
| HM13 | 8.50E-59 | 0.50006 | 0.866 | 0.611 | 1.65E-54 | INF/MLN | HM13 |
| EIF3M1 | 1.96E-58 | 0.416316 | 0.973 | 0.868 | 3.81E-54 | INF/MLN | EIF3M |
| DDX181 | 8.85E-58 | 0.462048 | 0.871 | 0.574 | 1.72E-53 | INF/MLN | DDX18 |
| FAM96B1 | 9.78E-58 | 0.402212 | 0.973 | 0.894 | 1.90E-53 | INF/MLN | FAM96B |
| FKBP10 | 1.11E-57 | 0.48259 | 0.893 | 0.635 | 2.15E-53 | INF/MLN | FKBP10 |
| TMED31 | 1.21E-57 | 0.433121 | 0.906 | 0.673 | 2.36E-53 | INF/MLN | TMED3 |
| COPE | 1.61E-57 | 0.372519 | 0.991 | 0.948 | 3.14E-53 | INF/MLN | COPE |
| SAR1B1 | 1.85E-57 | 0.411106 | 0.821 | 0.48 | 3.59E-53 | INF/MLN | SAR1B |
| TMEM14C | 3.79E-57 | 0.443004 | 0.969 | 0.851 | 7.37E-53 | INF/MLN | TMEM14C |
| BLVRB | 5.04E-57 | 0.455313 | 0.922 | 0.666 | 9.81E-53 | INF/MLN | BLVRB |
| MRPL55 | 1.19E-56 | 0.450983 | 0.944 | 0.783 | 2.32E-52 | INF/MLN | MRPL55 |
| EHD4 | 1.82E-56 | 0.292309 | 0.487 | 0.152 | 3.53E-52 | INF/MLN | EHD4 |
| RPS111 | 2.05E-56 | 0.304395 | 1 | 0.999 | 3.98E-52 | INF/MLN | RPS11 |
| PUSL1 | 2.42E-56 | 0.380043 | 0.634 | 0.279 | 4.70E-52 | INF/MLN | PUSL1 |
| MGST3 | 3.82E-56 | 0.40529 | 0.982 | 0.916 | 7.42E-52 | INF/MLN | MGST3 |
| SPSB31 | 7.33E-56 | 0.415061 | 0.821 | 0.492 | 1.43E-51 | INF/MLN | SPSB3 |
| IFRD11 | 1.21E-55 | 0.436417 | 0.891 | 0.598 | 2.36E-51 | INF/MLN | IFRD1 |
| TMED2 | 1.97E-55 | 0.387288 | 0.987 | 0.919 | 3.83E-51 | INF/MLN | TMED2 |
| ZNF598 | 2.83E-55 | 0.329854 | 0.529 | 0.182 | 5.51E-51 | INF/MLN | ZNF598 |
| NXT11 | 2.88E-55 | 0.444753 | 0.815 | 0.477 | 5.59E-51 | INF/MLN | NXT1 |
| RPL271 | 1.00E-54 | 0.297866 | 1 | 0.999 | 1.95E-50 | INF/MLN | RPL27 |
| SLC39A7 | 3.57E-54 | 0.427606 | 0.873 | 0.608 | 6.95E-50 | INF/MLN | SLC39A7 |
| DNAJC31 | 3.77E-54 | 0.376902 | 0.732 | 0.354 | 7.33E-50 | INF/MLN | DNAJC3 |
| TIPRL1 | 4.91E-54 | 0.410127 | 0.868 | 0.585 | 9.55E-50 | INF/MLN | TIPRL |
| P3H4 | 5.61E-54 | 0.428801 | 0.85 | 0.531 | 1.09E-49 | INF/MLN | P3H4 |
| RPS211 | 2.47E-53 | 0.348903 | 1 | 0.982 | 4.80E-49 | INF/MLN | RPS21 |
| PRELID1 | 3.79E-53 | 0.41185 | 0.962 | 0.891 | 7.37E-49 | INF/MLN | PRELID1 |
| YIF1B | 9.28E-53 | 0.400009 | 0.833 | 0.558 | 1.81E-48 | INF/MLN | YIF1B |
| MED8 | 1.12E-52 | 0.402166 | 0.75 | 0.413 | 2.17E-48 | INF/MLN | MED8 |
| SIKE11 | 1.46E-52 | 0.367054 | 0.781 | 0.419 | 2.85E-48 | INF/MLN | SIKE1 |

| | | | | | | | |
|---|---|---|---|---|---|---|---|
| TXNL4B | 4.39E-52 | 0.315941 | 0.478 | 0.157 | 8.53E-48 | INF/MLN | TXNL4B |
| DERL1 | 4.51E-52 | 0.395501 | 0.808 | 0.497 | 8.77E-48 | INF/MLN | DERL1 |
| TBC1D10/ | 4.65E-52 | 0.264997 | 0.473 | 0.15 | 9.05E-48 | INF/MLN | TBC1D10A |
| TMEM263 | 6.99E-52 | 0.375287 | 0.79 | 0.44 | 1.36E-47 | INF/MLN | TMEM263 |
| FABP31 | 1.19E-51 | 0.272483 | 0.359 | 0.083 | 2.32E-47 | INF/MLN | FABP3 |
| DNAJC10 | 1.26E-51 | 0.406892 | 0.737 | 0.392 | 2.45E-47 | INF/MLN | DNAJC10 |
| RRBP1 | 1.50E-51 | 0.34552 | 0.681 | 0.319 | 2.92E-47 | INF/MLN | RRBP1 |
| LRRC59 | 1.65E-51 | 0.4241 | 0.853 | 0.568 | 3.20E-47 | INF/MLN | LRRC59 |
| UFM1 | 2.15E-51 | 0.416786 | 0.893 | 0.617 | 4.19E-47 | INF/MLN | UFM1 |
| TRAP1 | 4.24E-51 | 0.429559 | 0.877 | 0.651 | 8.24E-47 | INF/MLN | TRAP1 |
| RPLP21 | 4.80E-51 | 0.264986 | 1 | 1 | 9.33E-47 | INF/MLN | RPLP2 |
| MLEC1 | 7.85E-51 | 0.415333 | 0.9 | 0.625 | 1.53E-46 | INF/MLN | MLEC |
| MALAT11 | 1.42E-50 | 0.351592 | 1 | 1 | 2.76E-46 | INF/MLN | MALAT1 |
| MAP2K11 | 1.93E-50 | 0.384584 | 0.797 | 0.451 | 3.76E-46 | INF/MLN | MAP2K1 |
| SEC23IP | 2.19E-50 | 0.363632 | 0.694 | 0.342 | 4.26E-46 | INF/MLN | SEC23IP |
| SUCO1 | 2.64E-50 | 0.332336 | 0.685 | 0.323 | 5.15E-46 | INF/MLN | SUCO |
| LAGE31 | 3.49E-50 | 0.469808 | 0.893 | 0.68 | 6.78E-46 | INF/MLN | LAGE3 |
| HSPA13 | 3.93E-50 | 0.364777 | 0.743 | 0.373 | 7.64E-46 | INF/MLN | HSPA13 |
| TXNIP | 5.62E-50 | 0.611056 | 0.795 | 0.483 | 1.09E-45 | INF/MLN | TXNIP |
| EBLN3 | 8.71E-50 | 0.387949 | 0.746 | 0.418 | 1.70E-45 | INF/MLN | EBLN3 |
| CUTA1 | 9.13E-50 | 0.394951 | 0.996 | 0.982 | 1.78E-45 | INF/MLN | CUTA |
| GNL31 | 1.05E-49 | 0.398334 | 0.81 | 0.489 | 2.04E-45 | INF/MLN | GNL3 |
| COG6 | 1.32E-49 | 0.287824 | 0.567 | 0.221 | 2.57E-45 | INF/MLN | COG6 |
| TGDS | 2.04E-49 | 0.314283 | 0.605 | 0.249 | 3.97E-45 | INF/MLN | TGDS |
| P4HA11 | 2.15E-49 | 0.517471 | 0.859 | 0.58 | 4.18E-45 | INF/MLN | P4HA1 |
| SLC35E1 | 2.50E-49 | 0.312592 | 0.547 | 0.212 | 4.87E-45 | INF/MLN | SLC35E1 |
| RAB181 | 4.90E-49 | 0.391907 | 0.931 | 0.739 | 9.53E-45 | INF/MLN | RAB18 |
| ALG3 | 8.24E-49 | 0.38442 | 0.757 | 0.426 | 1.60E-44 | INF/MLN | ALG3 |
| HSPA1B1 | 1.20E-48 | 0.461263 | 0.748 | 0.385 | 2.34E-44 | INF/MLN | HSPA1B |
| SAT11 | 1.56E-48 | 0.482663 | 0.993 | 0.882 | 3.03E-44 | INF/MLN | SAT1 |
| TMEM55B | 1.77E-48 | 0.400491 | 0.754 | 0.428 | 3.44E-44 | INF/MLN | TMEM55B |
| TMCO1 | 1.94E-48 | 0.403914 | 0.94 | 0.809 | 3.78E-44 | INF/MLN | TMCO1 |
| COX141 | 1.96E-48 | 0.389874 | 0.888 | 0.63 | 3.81E-44 | INF/MLN | COX14 |
| ATP6V0B1 | 2.60E-48 | 0.361094 | 0.991 | 0.909 | 5.05E-44 | INF/MLN | ATP6V0B |
| ERLEC1 | 3.82E-48 | 0.405076 | 0.862 | 0.577 | 7.42E-44 | INF/MLN | ERLEC1 |
| DNAJB91 | 4.93E-48 | 0.454221 | 0.924 | 0.676 | 9.60E-44 | INF/MLN | DNAJB9 |
| C1orf43 | 6.97E-48 | 0.369121 | 0.953 | 0.844 | 1.36E-43 | INF/MLN | C1orf43 |
| RPS181 | 7.21E-48 | 0.2883 | 1 | 1 | 1.40E-43 | INF/MLN | RPS18 |
| TMEM205 | 7.81E-48 | 0.425086 | 0.855 | 0.625 | 1.52E-43 | INF/MLN | TMEM205 |
| PNPLA81 | 8.56E-48 | 0.360686 | 0.761 | 0.403 | 1.67E-43 | INF/MLN | PNPLA8 |

| | | | | | | | |
|---|---|---|---|---|---|---|---|
| EIF5A | 1.07E-47 | 0.378681 | 0.984 | 0.94 | 2.09E-43 | INF/MLN | EIF5A |
| SEC24D | 1.52E-47 | 0.295431 | 0.518 | 0.197 | 2.96E-43 | INF/MLN | SEC24D |
| LCMT11 | 2.14E-47 | 0.386626 | 0.886 | 0.665 | 4.17E-43 | INF/MLN | LCMT1 |
| SAR1A1 | 2.27E-47 | 0.386497 | 0.862 | 0.589 | 4.41E-43 | INF/MLN | SAR1A |
| CHMP2B | 2.99E-47 | 0.4114 | 0.81 | 0.547 | 5.82E-43 | INF/MLN | CHMP2B |
| ATP5G1 | 4.56E-47 | 0.433825 | 0.94 | 0.83 | 8.86E-43 | INF/MLN | ATP5G1 |
| AKR1A1 | 5.93E-47 | 0.388579 | 0.913 | 0.748 | 1.15E-42 | INF/MLN | AKR1A1 |
| ST131 | 6.58E-47 | 0.333532 | 0.989 | 0.947 | 1.28E-42 | INF/MLN | ST13 |
| ENO21 | 1.09E-46 | 0.44889 | 0.973 | 0.802 | 2.13E-42 | INF/MLN | ENO2 |
| C12orf57 | 1.14E-46 | 0.386509 | 0.978 | 0.949 | 2.21E-42 | INF/MLN | C12orf57 |
| TMED9 | 1.59E-46 | 0.338746 | 0.984 | 0.89 | 3.10E-42 | INF/MLN | TMED9 |
| PDHX | 1.64E-46 | 0.331948 | 0.737 | 0.4 | 3.19E-42 | INF/MLN | PDHX |
| ATP6V1D1 | 2.58E-46 | 0.417969 | 0.897 | 0.711 | 5.01E-42 | INF/MLN | ATP6V1D |
| GNL2 | 4.08E-46 | 0.373467 | 0.746 | 0.418 | 7.93E-42 | INF/MLN | GNL2 |
| EBNA1BP2 | 5.09E-46 | 0.43538 | 0.801 | 0.533 | 9.91E-42 | INF/MLN | EBNA1BP2 |
| ARFGAP3 | 5.96E-46 | 0.376623 | 0.795 | 0.51 | 1.16E-41 | INF/MLN | ARFGAP3 |
| RAB6A1 | 6.27E-46 | 0.360612 | 0.915 | 0.687 | 1.22E-41 | INF/MLN | RAB6A |
| ATG1011 | 2.77E-45 | 0.391332 | 0.781 | 0.503 | 5.40E-41 | INF/MLN | ATG101 |
| KDELR3 | 2.84E-45 | 0.301086 | 0.513 | 0.2 | 5.53E-41 | INF/MLN | KDELR3 |
| ALDOC1 | 3.76E-45 | 0.428161 | 0.721 | 0.39 | 7.31E-41 | INF/MLN | ALDOC |
| ZCCHC7 | 4.35E-45 | 0.362249 | 0.705 | 0.365 | 8.46E-41 | INF/MLN | ZCCHC7 |
| GTF2F1 | 5.46E-45 | 0.349343 | 0.833 | 0.558 | 1.06E-40 | INF/MLN | GTF2F1 |
| USE1 | 6.88E-45 | 0.419986 | 0.717 | 0.41 | 1.34E-40 | INF/MLN | USE1 |
| ENO3 | 9.15E-45 | 0.414824 | 0.679 | 0.357 | 1.78E-40 | INF/MLN | ENO3 |
| NOL7 | 1.48E-44 | 0.348983 | 0.917 | 0.725 | 2.89E-40 | INF/MLN | NOL7 |
| CMPK1 | 1.53E-44 | 0.367698 | 0.906 | 0.706 | 2.98E-40 | INF/MLN | CMPK1 |
| NDUFV3 | 4.79E-44 | 0.384165 | 0.924 | 0.779 | 9.32E-40 | INF/MLN | NDUFV3 |
| ZNF331 | 9.14E-44 | 0.333538 | 0.482 | 0.183 | 1.78E-39 | INF/MLN | ZNF331 |
| IMP4 | 9.85E-44 | 0.339101 | 0.779 | 0.467 | 1.92E-39 | INF/MLN | IMP4 |
| PLA2G16 | 1.62E-43 | 0.396301 | 0.775 | 0.479 | 3.15E-39 | INF/MLN | PLA2G16 |
| USMG51 | 2.06E-43 | 0.347987 | 0.955 | 0.84 | 4.02E-39 | INF/MLN | USMG5 |
| STRAP | 2.27E-43 | 0.307087 | 0.991 | 0.945 | 4.41E-39 | INF/MLN | STRAP |
| GDF151 | 3.28E-43 | 0.783885 | 0.455 | 0.168 | 6.38E-39 | INF/MLN | GDF15 |
| DNAJB1 | 3.54E-43 | 0.582244 | 0.844 | 0.609 | 6.90E-39 | INF/MLN | DNAJB1 |
| UTP6 | 7.02E-43 | 0.320853 | 0.614 | 0.288 | 1.37E-38 | INF/MLN | UTP6 |
| SWI5 | 8.64E-43 | 0.349186 | 0.799 | 0.506 | 1.68E-38 | INF/MLN | SWI5 |
| ATF3 | 1.04E-42 | 0.530672 | 0.701 | 0.382 | 2.03E-38 | INF/MLN | ATF3 |
| WIPI1 | 1.21E-42 | 0.302822 | 0.576 | 0.253 | 2.35E-38 | INF/MLN | WIPI1 |
| MDH2 | 1.31E-42 | 0.317878 | 0.991 | 0.923 | 2.54E-38 | INF/MLN | MDH2 |
| CD68 | 1.49E-42 | 0.256892 | 0.422 | 0.142 | 2.90E-38 | INF/MLN | CD68 |

| | | | | | | | |
|---|---|---|---|---|---|---|---|
| ENO1 | 1.67E-42 | 0.363875 | 0.998 | 0.977 | 3.25E-38 | INF/MLN | ENO1 |
| SESN2 | 2.47E-42 | 0.374332 | 0.498 | 0.2 | 4.81E-38 | INF/MLN | SESN2 |
| MRPL2 | 2.70E-42 | 0.354452 | 0.783 | 0.5 | 5.26E-38 | INF/MLN | MRPL2 |
| RPL101 | 4.20E-42 | 0.274234 | 1 | 1 | 8.17E-38 | INF/MLN | RPL10 |
| EIF2S3 | 4.29E-42 | 0.409567 | 0.946 | 0.778 | 8.35E-38 | INF/MLN | EIF2S3 |
| POMP | 4.52E-42 | 0.319356 | 0.993 | 0.941 | 8.79E-38 | INF/MLN | POMP |
| CRYAB1 | 5.97E-42 | 0.677748 | 0.58 | 0.286 | 1.16E-37 | INF/MLN | CRYAB |
| EIF4EBP1 | 8.11E-42 | 0.499372 | 0.79 | 0.499 | 1.58E-37 | INF/MLN | EIF4EBP1 |
| SMIM14 | 1.62E-41 | 0.350607 | 0.833 | 0.565 | 3.15E-37 | INF/MLN | SMIM14 |
| ADK | 3.04E-41 | 0.345563 | 0.676 | 0.374 | 5.92E-37 | INF/MLN | ADK |
| TMF1 | 3.61E-41 | 0.333235 | 0.708 | 0.385 | 7.02E-37 | INF/MLN | TMF1 |
| A1BG1 | 4.39E-41 | 0.335565 | 0.842 | 0.538 | 8.54E-37 | INF/MLN | A1BG |
| COASY | 4.65E-41 | 0.323326 | 0.754 | 0.426 | 9.04E-37 | INF/MLN | COASY |
| HSPA61 | 5.18E-41 | 0.946951 | 0.529 | 0.238 | 1.01E-36 | INF/MLN | HSPA6 |
| UBE2B1 | 6.12E-41 | 0.358902 | 0.886 | 0.694 | 1.19E-36 | INF/MLN | UBE2B |
| HSPB11 | 7.00E-41 | 0.474265 | 0.993 | 0.917 | 1.36E-36 | INF/MLN | HSPB1 |
| GPX8 | 7.12E-41 | 0.362546 | 0.696 | 0.379 | 1.39E-36 | INF/MLN | GPX8 |
| UBC | 9.20E-41 | 0.370453 | 0.998 | 0.99 | 1.79E-36 | INF/MLN | UBC |
| ATP6AP2 | 1.05E-40 | 0.33454 | 0.955 | 0.798 | 2.04E-36 | INF/MLN | ATP6AP2 |
| SQSTM11 | 1.06E-40 | 0.471374 | 0.871 | 0.646 | 2.06E-36 | INF/MLN | SQSTM1 |
| SEC23A | 1.15E-40 | 0.361269 | 0.842 | 0.584 | 2.24E-36 | INF/MLN | SEC23A |
| PMM1 | 1.48E-40 | 0.335195 | 0.725 | 0.423 | 2.87E-36 | INF/MLN | PMM1 |
| RABAC1 | 2.41E-40 | 0.339199 | 0.98 | 0.929 | 4.69E-36 | INF/MLN | RABAC1 |
| BRF2 | 3.11E-40 | 0.295251 | 0.634 | 0.321 | 6.06E-36 | INF/MLN | BRF2 |
| MMADHC | 3.14E-40 | 0.321319 | 0.915 | 0.771 | 6.10E-36 | INF/MLN | MMADHC |
| RAB1A | 3.83E-40 | 0.309257 | 0.975 | 0.871 | 7.44E-36 | INF/MLN | RAB1A |
| UAP1 | 8.12E-40 | 0.353273 | 0.676 | 0.37 | 1.58E-35 | INF/MLN | UAP1 |
| SAFB2 | 1.08E-39 | 0.367416 | 0.714 | 0.423 | 2.11E-35 | INF/MLN | SAFB2 |
| C19orf25 | 1.25E-39 | 0.345473 | 0.737 | 0.446 | 2.43E-35 | INF/MLN | C19orf25 |
| SEC63 | 2.12E-39 | 0.376676 | 0.839 | 0.622 | 4.13E-35 | INF/MLN | SEC63 |
| ZFPL1 | 2.45E-39 | 0.356344 | 0.728 | 0.441 | 4.77E-35 | INF/MLN | ZFPL1 |
| COPG1 | 2.56E-39 | 0.321421 | 0.696 | 0.381 | 4.98E-35 | INF/MLN | COPG1 |
| CDC37 | 2.57E-39 | 0.352443 | 0.897 | 0.745 | 5.00E-35 | INF/MLN | CDC37 |
| ADM1 | 3.06E-39 | 0.311215 | 0.4 | 0.131 | 5.96E-35 | INF/MLN | ADM |
| ODF2 | 3.48E-39 | 0.293036 | 0.719 | 0.389 | 6.76E-35 | INF/MLN | ODF2 |
| SNHG81 | 3.70E-39 | 0.399782 | 0.781 | 0.511 | 7.19E-35 | INF/MLN | SNHG8 |
| MRPL18 | 4.00E-39 | 0.404957 | 0.853 | 0.635 | 7.78E-35 | INF/MLN | MRPL18 |
| COPA | 4.89E-39 | 0.334719 | 0.746 | 0.432 | 9.51E-35 | INF/MLN | COPA |
| USO1 | 4.90E-39 | 0.329699 | 0.732 | 0.433 | 9.54E-35 | INF/MLN | USO1 |
| DIS3 | 4.92E-39 | 0.295978 | 0.654 | 0.323 | 9.58E-35 | INF/MLN | DIS3 |

| | | | | | | | |
|---|---|---|---|---|---|---|---|
| C19orf81 | 5.28E-39 | 0.310832 | 0.676 | 0.36 | 1.03E-34 | INF/MLN | C19orf81 |
| LAMTOR3 | 5.41E-39 | 0.319878 | 0.835 | 0.579 | 1.05E-34 | INF/MLN | LAMTOR3 |
| CALCB1 | 7.06E-39 | 0.261052 | 0.299 | 0.075 | 1.37E-34 | INF/MLN | CALCB |
| SDHAF1 | 7.52E-39 | 0.301769 | 0.612 | 0.308 | 1.46E-34 | INF/MLN | SDHAF1 |
| NUBP2 | 8.66E-39 | 0.352689 | 0.837 | 0.603 | 1.68E-34 | INF/MLN | NUBP2 |
| TRAPPC2F | 8.99E-39 | 0.292099 | 0.623 | 0.31 | 1.75E-34 | INF/MLN | TRAPPC2P1 |
| SERF2 | 2.36E-38 | 0.298521 | 1 | 0.997 | 4.59E-34 | INF/MLN | SERF2 |
| BUD311 | 2.63E-38 | 0.352177 | 0.949 | 0.829 | 5.11E-34 | INF/MLN | BUD31 |
| ACBD3 | 2.93E-38 | 0.337429 | 0.701 | 0.405 | 5.71E-34 | INF/MLN | ACBD3 |
| PGP | 3.50E-38 | 0.347015 | 0.804 | 0.529 | 6.82E-34 | INF/MLN | PGP |
| DNAJC25 | 1.98E-37 | 0.268351 | 0.498 | 0.209 | 3.86E-33 | INF/MLN | DNAJC25 |
| SLC35A2 | 2.21E-37 | 0.319514 | 0.71 | 0.424 | 4.30E-33 | INF/MLN | SLC35A2 |
| NABP2 | 3.06E-37 | 0.332637 | 0.812 | 0.579 | 5.95E-33 | INF/MLN | NABP2 |
| DHDDS | 3.48E-37 | 0.25083 | 0.509 | 0.224 | 6.78E-33 | INF/MLN | DHDDS |
| NDUFS8 | 3.75E-37 | 0.288671 | 0.975 | 0.929 | 7.29E-33 | INF/MLN | NDUFS8 |
| RNF167 | 4.91E-37 | 0.340096 | 0.799 | 0.546 | 9.54E-33 | INF/MLN | RNF167 |
| NPC21 | 4.95E-37 | 0.323713 | 0.982 | 0.886 | 9.63E-33 | INF/MLN | NPC2 |
| WHAMM | 5.34E-37 | 0.276403 | 0.542 | 0.245 | 1.04E-32 | INF/MLN | WHAMM |
| VMP1 | 5.49E-37 | 0.306378 | 0.81 | 0.535 | 1.07E-32 | INF/MLN | VMP1 |
| ASAH11 | 8.43E-37 | 0.466712 | 0.895 | 0.696 | 1.64E-32 | INF/MLN | ASAH1 |
| MCFD2 | 8.86E-37 | 0.355701 | 0.77 | 0.502 | 1.72E-32 | INF/MLN | MCFD2 |
| DPAGT1 | 1.17E-36 | 0.276801 | 0.589 | 0.29 | 2.27E-32 | INF/MLN | DPAGT1 |
| MTHFD2 | 1.31E-36 | 0.380334 | 0.634 | 0.325 | 2.55E-32 | INF/MLN | MTHFD2 |
| WBP2 | 2.06E-36 | 0.352221 | 0.835 | 0.582 | 4.01E-32 | INF/MLN | WBP2 |
| TMX2 | 2.08E-36 | 0.344383 | 0.77 | 0.502 | 4.04E-32 | INF/MLN | TMX2 |
| CKS21 | 2.17E-36 | 0.369478 | 0.942 | 0.747 | 4.22E-32 | INF/MLN | CKS2 |
| STT3A | 2.76E-36 | 0.266145 | 0.592 | 0.283 | 5.36E-32 | INF/MLN | STT3A |
| TNFRSF1( | 3.11E-36 | 0.285689 | 0.578 | 0.273 | 6.04E-32 | INF/MLN | TNFRSF10B |
| GADD45G | 3.27E-36 | 0.274852 | 0.569 | 0.251 | 6.36E-32 | INF/MLN | GADD45G |
| PLP2 | 4.71E-36 | 0.280069 | 0.52 | 0.238 | 9.17E-32 | INF/MLN | PLP2 |
| BAG3 | 4.81E-36 | 0.50882 | 0.585 | 0.302 | 9.35E-32 | INF/MLN | BAG3 |
| SBDS1 | 7.75E-36 | 0.357797 | 0.839 | 0.59 | 1.51E-31 | INF/MLN | SBDS |
| PHKG2 | 9.12E-36 | 0.280854 | 0.605 | 0.308 | 1.77E-31 | INF/MLN | PHKG2 |
| RNMT1 | 1.40E-35 | 0.328567 | 0.763 | 0.485 | 2.72E-31 | INF/MLN | RNMT |
| TMEM41B | 1.48E-35 | 0.358435 | 0.717 | 0.43 | 2.89E-31 | INF/MLN | TMEM41B |
| COPB1 | 1.81E-35 | 0.333598 | 0.79 | 0.523 | 3.53E-31 | INF/MLN | COPB1 |
| ALDH18A1 | 2.62E-35 | 0.266527 | 0.625 | 0.322 | 5.09E-31 | INF/MLN | ALDH18A1 |
| PCMT11 | 3.25E-35 | 0.308334 | 0.949 | 0.789 | 6.33E-31 | INF/MLN | PCMT1 |
| IDI11 | 3.38E-35 | 0.634105 | 0.924 | 0.797 | 6.57E-31 | INF/MLN | IDI1 |
| CLK31 | 4.76E-35 | 0.27632 | 0.667 | 0.364 | 9.25E-31 | INF/MLN | CLK3 |

| | | | | | | | |
|---|---|---|---|---|---|---|---|
| STX5 | 4.84E-35 | 0.296801 | 0.721 | 0.44 | 9.42E-31 | INF/MLN | STX5 |
| TOMM40 | 5.26E-35 | 0.368518 | 0.873 | 0.694 | 1.02E-30 | INF/MLN | TOMM40 |
| FADS1 | 5.45E-35 | 0.427187 | 0.911 | 0.772 | 1.06E-30 | INF/MLN | FADS1 |
| PPP1CB | 5.83E-35 | 0.357284 | 0.906 | 0.713 | 1.13E-30 | INF/MLN | PPP1CB |
| TMEM14B | 7.36E-35 | 0.323903 | 0.953 | 0.803 | 1.43E-30 | INF/MLN | TMEM14B |
| GOSR2 | 7.42E-35 | 0.286141 | 0.596 | 0.307 | 1.44E-30 | INF/MLN | GOSR2 |
| BEX5 | 7.67E-35 | 0.317893 | 0.67 | 0.373 | 1.49E-30 | INF/MLN | BEX5 |
| OST41 | 1.30E-34 | 0.262053 | 0.996 | 0.971 | 2.52E-30 | INF/MLN | OST4 |
| NKAP | 1.44E-34 | 0.305889 | 0.781 | 0.515 | 2.80E-30 | INF/MLN | NKAP |
| MICA | 1.56E-34 | 0.276029 | 0.562 | 0.279 | 3.04E-30 | INF/MLN | MICA |
| ZFP36 | 1.97E-34 | 0.260337 | 0.542 | 0.245 | 3.83E-30 | INF/MLN | ZFP36 |
| EMD | 2.79E-34 | 0.297793 | 0.795 | 0.529 | 5.42E-30 | INF/MLN | EMD |
| YIPF3 | 3.11E-34 | 0.312788 | 0.897 | 0.785 | 6.05E-30 | INF/MLN | YIPF3 |
| RNF181 | 3.47E-34 | 0.316055 | 0.913 | 0.778 | 6.76E-30 | INF/MLN | RNF181 |
| EIF3H1 | 3.79E-34 | 0.268775 | 0.984 | 0.941 | 7.37E-30 | INF/MLN | EIF3H |
| PPP2R5B | 4.31E-34 | 0.250486 | 0.529 | 0.234 | 8.39E-30 | INF/MLN | PPP2R5B |
| ANKHD1 | 5.31E-34 | 0.266686 | 0.518 | 0.239 | 1.03E-29 | INF/MLN | ANKHD1 |
| M6PR1 | 6.79E-34 | 0.339423 | 0.795 | 0.562 | 1.32E-29 | INF/MLN | M6PR |
| PJA21 | 9.47E-34 | 0.286868 | 0.821 | 0.533 | 1.84E-29 | INF/MLN | PJA2 |
| BST2 | 1.36E-33 | 0.288754 | 0.538 | 0.247 | 2.64E-29 | INF/MLN | BST2 |
| GAR1 | 2.11E-33 | 0.288515 | 0.656 | 0.363 | 4.10E-29 | INF/MLN | GAR1 |
| PREB | 2.20E-33 | 0.28211 | 0.672 | 0.373 | 4.27E-29 | INF/MLN | PREB |
| TFG | 2.63E-33 | 0.325911 | 0.868 | 0.695 | 5.11E-29 | INF/MLN | TFG |
| UQCRQ | 4.08E-33 | 0.279404 | 0.991 | 0.978 | 7.94E-29 | INF/MLN | UQCRQ |
| GHITM | 4.66E-33 | 0.283213 | 0.98 | 0.91 | 9.06E-29 | INF/MLN | GHITM |
| DPH7 | 9.61E-33 | 0.261314 | 0.616 | 0.322 | 1.87E-28 | INF/MLN | DPH7 |
| FADS2 | 1.72E-32 | 0.400318 | 0.761 | 0.522 | 3.34E-28 | INF/MLN | FADS2 |
| EIF4H | 2.17E-32 | 0.317083 | 0.933 | 0.831 | 4.23E-28 | INF/MLN | EIF4H |
| SPCS1 | 3.00E-32 | 0.25532 | 0.993 | 0.936 | 5.84E-28 | INF/MLN | SPCS1 |
| LONP1 | 4.17E-32 | 0.275365 | 0.554 | 0.29 | 8.12E-28 | INF/MLN | LONP1 |
| SLC25A39 | 4.19E-32 | 0.324207 | 0.902 | 0.733 | 8.15E-28 | INF/MLN | SLC25A39 |
| ZNF6221 | 5.42E-32 | 0.320138 | 0.699 | 0.44 | 1.05E-27 | INF/MLN | ZNF622 |
| EIF3F1 | 5.58E-32 | 0.258765 | 0.991 | 0.972 | 1.09E-27 | INF/MLN | EIF3F |
| LTN1 | 6.01E-32 | 0.2518 | 0.58 | 0.293 | 1.17E-27 | INF/MLN | LTN1 |
| FUCA1 | 6.55E-32 | 0.256914 | 0.612 | 0.323 | 1.27E-27 | INF/MLN | FUCA1 |
| SDHA | 1.09E-31 | 0.310358 | 0.797 | 0.556 | 2.13E-27 | INF/MLN | SDHA |
| CYB561D2 | 1.14E-31 | 0.255722 | 0.498 | 0.234 | 2.22E-27 | INF/MLN | CYB561D2 |
| SRSF5 | 1.43E-31 | 0.295711 | 0.96 | 0.917 | 2.79E-27 | INF/MLN | SRSF5 |
| WDR45B | 2.01E-31 | 0.313159 | 0.79 | 0.579 | 3.91E-27 | INF/MLN | WDR45B |
| C14orf166 | 2.16E-31 | 0.251463 | 0.969 | 0.921 | 4.21E-27 | INF/MLN | C14orf166 |

| | | | | | | | |
|---|---|---|---|---|---|---|---|
| GADD45A | 2.34E-31 | 0.343032 | 0.714 | 0.435 | 4.55E-27 | INF/MLN | GADD45A |
| ILVBL | 2.67E-31 | 0.275961 | 0.67 | 0.402 | 5.20E-27 | INF/MLN | ILVBL |
| PYCR1 | 3.65E-31 | 0.312397 | 0.759 | 0.494 | 7.11E-27 | INF/MLN | PYCR1 |
| ADIPOR1 | 3.80E-31 | 0.304489 | 0.815 | 0.579 | 7.39E-27 | INF/MLN | ADIPOR1 |
| SSR31 | 4.01E-31 | 0.299858 | 0.893 | 0.724 | 7.80E-27 | INF/MLN | SSR3 |
| CRIPT | 4.33E-31 | 0.286638 | 0.748 | 0.485 | 8.42E-27 | INF/MLN | CRIPT |
| SCNM11 | 1.01E-30 | 0.282899 | 0.696 | 0.424 | 1.96E-26 | INF/MLN | SCNM1 |
| EIF2B2 | 1.02E-30 | 0.276532 | 0.661 | 0.399 | 1.99E-26 | INF/MLN | EIF2B2 |
| ZNF503 | 1.71E-30 | 0.365419 | 0.833 | 0.58 | 3.33E-26 | INF/MLN | ZNF503 |
| DAPK3 | 1.96E-30 | 0.280516 | 0.681 | 0.416 | 3.80E-26 | INF/MLN | DAPK3 |
| PPAT | 1.98E-30 | 0.266461 | 0.643 | 0.368 | 3.85E-26 | INF/MLN | PPAT |
| APMAP | 2.00E-30 | 0.311231 | 0.844 | 0.664 | 3.90E-26 | INF/MLN | APMAP |
| C6orf481 | 2.13E-30 | 0.282052 | 0.975 | 0.917 | 4.15E-26 | INF/MLN | C6orf48 |
| ATP6V1E1 | 3.62E-30 | 0.301691 | 0.799 | 0.572 | 7.05E-26 | INF/MLN | ATP6V1E1 |
| EIF2AK1 | 4.19E-30 | 0.301592 | 0.828 | 0.619 | 8.15E-26 | INF/MLN | EIF2AK1 |
| BNIP3L | 4.74E-30 | 0.314677 | 0.96 | 0.875 | 9.22E-26 | INF/MLN | BNIP3L |
| CHMP2A | 4.83E-30 | 0.274999 | 0.955 | 0.894 | 9.40E-26 | INF/MLN | CHMP2A |
| DDX50 | 5.02E-30 | 0.275279 | 0.759 | 0.49 | 9.76E-26 | INF/MLN | DDX50 |
| EIF3I | 5.71E-30 | 0.285895 | 0.967 | 0.909 | 1.11E-25 | INF/MLN | EIF3I |
| GNAI31 | 5.75E-30 | 0.278822 | 0.92 | 0.793 | 1.12E-25 | INF/MLN | GNAI3 |
| TMED71 | 6.25E-30 | 0.261722 | 0.777 | 0.516 | 1.22E-25 | INF/MLN | TMED7 |
| IFT20 | 7.50E-30 | 0.29769 | 0.748 | 0.488 | 1.46E-25 | INF/MLN | IFT20 |
| BCAS21 | 7.65E-30 | 0.290011 | 0.817 | 0.564 | 1.49E-25 | INF/MLN | BCAS2 |
| GPI1 | 9.17E-30 | 0.368487 | 0.848 | 0.62 | 1.78E-25 | INF/MLN | GPI |
| NELFE | 1.14E-29 | 0.314579 | 0.864 | 0.703 | 2.21E-25 | INF/MLN | NELFE |
| NAGK | 1.27E-29 | 0.285492 | 0.6 | 0.347 | 2.47E-25 | INF/MLN | NAGK |
| SHMT2 | 1.51E-29 | 0.324406 | 0.703 | 0.443 | 2.94E-25 | INF/MLN | SHMT2 |
| DYRK4 | 1.64E-29 | 0.280366 | 0.719 | 0.477 | 3.18E-25 | INF/MLN | DYRK4 |
| TOMM201 | 1.67E-29 | 0.275547 | 0.958 | 0.853 | 3.25E-25 | INF/MLN | TOMM20 |
| SRM | 1.91E-29 | 0.305229 | 0.786 | 0.53 | 3.71E-25 | INF/MLN | SRM |
| SPNS1 | 4.85E-29 | 0.312028 | 0.759 | 0.52 | 9.43E-25 | INF/MLN | SPNS1 |
| ATF4 | 5.19E-29 | 0.395955 | 0.893 | 0.783 | 1.01E-24 | INF/MLN | ATF4 |
| PRKAB1 | 6.37E-29 | 0.27848 | 0.58 | 0.323 | 1.24E-24 | INF/MLN | PRKAB1 |
| HSPH11 | 7.27E-29 | 0.373634 | 0.757 | 0.509 | 1.42E-24 | INF/MLN | HSPH1 |
| RARS | 1.01E-28 | 0.287319 | 0.703 | 0.442 | 1.97E-24 | INF/MLN | RARS |
| NUCB2 | 1.40E-28 | 0.310419 | 0.873 | 0.67 | 2.71E-24 | INF/MLN | NUCB2 |
| CCPG1 | 1.49E-28 | 0.268784 | 0.634 | 0.37 | 2.89E-24 | INF/MLN | CCPG1 |
| GPT2 | 1.51E-28 | 0.293347 | 0.511 | 0.258 | 2.93E-24 | INF/MLN | GPT2 |
| TIMM441 | 1.94E-28 | 0.293528 | 0.712 | 0.455 | 3.76E-24 | INF/MLN | TIMM44 |
| SSR1 | 2.09E-28 | 0.276637 | 0.83 | 0.603 | 4.06E-24 | INF/MLN | SSR1 |

| | | | | | | | |
|---|---|---|---|---|---|---|---|
| EMC7 | 2.51E-28 | 0.292462 | 0.855 | 0.685 | 4.87E-24 | INF/MLN | EMC7 |
| GPX1 | 4.47E-28 | 0.25979 | 0.98 | 0.936 | 8.69E-24 | INF/MLN | GPX1 |
| GOLIM4 | 5.31E-28 | 0.276417 | 0.636 | 0.367 | 1.03E-23 | INF/MLN | GOLIM4 |
| TMEM87A | 6.21E-28 | 0.277028 | 0.754 | 0.482 | 1.21E-23 | INF/MLN | TMEM87A |
| APH1A | 6.62E-28 | 0.279024 | 0.902 | 0.702 | 1.29E-23 | INF/MLN | APH1A |
| PPP1R11 | 7.35E-28 | 0.299959 | 0.779 | 0.549 | 1.43E-23 | INF/MLN | PPP1R11 |
| AP3D1 | 9.45E-28 | 0.29743 | 0.781 | 0.588 | 1.84E-23 | INF/MLN | AP3D1 |
| KLHDC2 | 1.29E-27 | 0.285421 | 0.826 | 0.647 | 2.52E-23 | INF/MLN | KLHDC2 |
| PPM1K | 1.38E-27 | 0.277117 | 0.656 | 0.387 | 2.69E-23 | INF/MLN | PPM1K |
| GOLGA4 | 1.43E-27 | 0.259521 | 0.688 | 0.426 | 2.79E-23 | INF/MLN | GOLGA4 |
| PMEL1 | 1.90E-27 | 0.648209 | 0.542 | 0.301 | 3.69E-23 | INF/MLN | PMEL |
| ASNS | 1.93E-27 | 0.285556 | 0.612 | 0.326 | 3.75E-23 | INF/MLN | ASNS |
| ZFAND2A1 | 2.12E-27 | 0.389848 | 0.621 | 0.367 | 4.12E-23 | INF/MLN | ZFAND2A |
| CNPY2 | 2.26E-27 | 0.29769 | 0.897 | 0.794 | 4.39E-23 | INF/MLN | CNPY2 |
| SGTA | 2.36E-27 | 0.288656 | 0.766 | 0.582 | 4.58E-23 | INF/MLN | SGTA |
| NOC4L | 6.33E-27 | 0.270665 | 0.629 | 0.375 | 1.23E-22 | INF/MLN | NOC4L |
| EIF3D | 6.83E-27 | 0.254237 | 0.929 | 0.799 | 1.33E-22 | INF/MLN | EIF3D |
| RAB5A | 9.43E-27 | 0.283584 | 0.819 | 0.61 | 1.83E-22 | INF/MLN | RAB5A |
| PDHA1 | 1.13E-26 | 0.25834 | 0.855 | 0.68 | 2.21E-22 | INF/MLN | PDHA1 |
| EIF2S2 | 1.48E-26 | 0.283929 | 0.917 | 0.812 | 2.89E-22 | INF/MLN | EIF2S2 |
| SF3B2 | 2.38E-26 | 0.25005 | 0.975 | 0.903 | 4.63E-22 | INF/MLN | SF3B2 |
| STARD3NI | 2.82E-26 | 0.271882 | 0.886 | 0.737 | 5.49E-22 | INF/MLN | STARD3NL |
| SUCLG1 | 4.52E-26 | 0.258162 | 0.891 | 0.746 | 8.80E-22 | INF/MLN | SUCLG1 |
| SEC23B | 5.25E-26 | 0.253406 | 0.696 | 0.438 | 1.02E-21 | INF/MLN | SEC23B |
| AAMP | 7.78E-26 | 0.279653 | 0.833 | 0.653 | 1.51E-21 | INF/MLN | AAMP |
| LSM31 | 1.13E-25 | 0.259233 | 0.808 | 0.608 | 2.20E-21 | INF/MLN | LSM3 |
| ARHGDIA1 | 1.22E-25 | 0.3212 | 0.808 | 0.629 | 2.37E-21 | INF/MLN | ARHGDIA |
| SARS | 1.93E-25 | 0.299099 | 0.859 | 0.688 | 3.76E-21 | INF/MLN | SARS |
| GOSR1 | 2.41E-25 | 0.270954 | 0.67 | 0.413 | 4.68E-21 | INF/MLN | GOSR1 |
| NAMPT1 | 2.48E-25 | 0.281057 | 0.652 | 0.407 | 4.82E-21 | INF/MLN | NAMPT |
| BLOC1S6 | 2.54E-25 | 0.280068 | 0.759 | 0.511 | 4.94E-21 | INF/MLN | BLOC1S6 |
| PPP1R15A | 3.33E-25 | 0.355132 | 0.85 | 0.648 | 6.48E-21 | INF/MLN | PPP1R15A |
| DUSP1 | 3.76E-25 | 0.418273 | 0.833 | 0.636 | 7.31E-21 | INF/MLN | DUSP1 |
| LAMP1 | 3.98E-25 | 0.281437 | 0.875 | 0.715 | 7.75E-21 | INF/MLN | LAMP1 |
| CCDC47 | 7.41E-25 | 0.251558 | 0.694 | 0.458 | 1.44E-20 | INF/MLN | CCDC47 |
| C6orf62 | 1.01E-24 | 0.251448 | 0.67 | 0.41 | 1.97E-20 | INF/MLN | C6orf62 |
| XBP1 | 1.08E-24 | 0.292353 | 0.879 | 0.707 | 2.09E-20 | INF/MLN | XBP1 |
| LMAN1 | 1.54E-24 | 0.287699 | 0.801 | 0.569 | 3.00E-20 | INF/MLN | LMAN1 |
| TMEM101 | 1.68E-24 | 0.252248 | 0.734 | 0.493 | 3.27E-20 | INF/MLN | TMEM101 |
| ACAT1 | 1.87E-24 | 0.321475 | 0.676 | 0.451 | 3.63E-20 | INF/MLN | ACAT1 |

| | | | | | | | |
|---|---|---|---|---|---|---|---|
| STK19 | 3.24E-24 | 0.251109 | 0.638 | 0.392 | 6.31E-20 | INF/MLN | STK19 |
| MAF1 | 3.45E-24 | 0.256472 | 0.844 | 0.65 | 6.71E-20 | INF/MLN | MAF1 |
| ANKRD37 | 3.52E-24 | 0.273678 | 0.417 | 0.193 | 6.86E-20 | INF/MLN | ANKRD37 |
| MRPL17 | 4.88E-24 | 0.275793 | 0.768 | 0.559 | 9.49E-20 | INF/MLN | MRPL17 |
| ZRANB2 | 5.38E-24 | 0.256656 | 0.799 | 0.577 | 1.05E-19 | INF/MLN | ZRANB2 |
| CHCHD10 | 8.58E-24 | 0.270977 | 0.879 | 0.69 | 1.67E-19 | INF/MLN | CHCHD10 |
| BET1 | 1.03E-23 | 0.257902 | 0.775 | 0.539 | 2.00E-19 | INF/MLN | BET1 |
| DDX211 | 1.04E-23 | 0.269359 | 0.757 | 0.538 | 2.03E-19 | INF/MLN | DDX21 |
| MVD | 1.15E-23 | 0.255138 | 0.674 | 0.418 | 2.24E-19 | INF/MLN | MVD |
| YARS | 1.50E-23 | 0.272715 | 0.685 | 0.462 | 2.92E-19 | INF/MLN | YARS |
| GTPBP4 | 1.52E-23 | 0.259658 | 0.721 | 0.5 | 2.96E-19 | INF/MLN | GTPBP4 |
| AARS | 3.75E-23 | 0.253079 | 0.574 | 0.337 | 7.30E-19 | INF/MLN | AARS |
| JUNB1 | 6.72E-23 | 0.312822 | 0.725 | 0.487 | 1.31E-18 | INF/MLN | JUNB |
| SMIM15 | 1.37E-22 | 0.268832 | 0.748 | 0.526 | 2.66E-18 | INF/MLN | SMIM15 |
| VCP | 1.91E-22 | 0.276856 | 0.94 | 0.818 | 3.71E-18 | INF/MLN | VCP |
| HNRNPUL | 3.27E-22 | 0.250773 | 0.844 | 0.705 | 6.37E-18 | INF/MLN | HNRNPUL1 |
| EIF4A3 | 6.04E-22 | 0.256539 | 0.857 | 0.665 | 1.17E-17 | INF/MLN | EIF4A3 |
| JUN | 6.43E-22 | 0.326015 | 0.877 | 0.705 | 1.25E-17 | INF/MLN | JUN |
| C1orf351 | 1.24E-21 | 0.277387 | 0.772 | 0.592 | 2.41E-17 | INF/MLN | C1orf35 |
| COX17 | 1.58E-21 | 0.261829 | 0.906 | 0.729 | 3.07E-17 | INF/MLN | COX17 |
| CLPP | 1.75E-21 | 0.251248 | 0.844 | 0.666 | 3.41E-17 | INF/MLN | CLPP |
| WFDC2 | 3.50E-21 | 0.258752 | 0.533 | 0.317 | 6.81E-17 | INF/MLN | WFDC2 |
| SLC3A21 | 4.13E-21 | 0.252111 | 0.971 | 0.899 | 8.04E-17 | INF/MLN | SLC3A2 |
| TSN | 6.61E-21 | 0.26653 | 0.821 | 0.635 | 1.29E-16 | INF/MLN | TSN |
| LGALS3BI | 3.34E-19 | 0.291187 | 0.79 | 0.603 | 6.51E-15 | INF/MLN | LGALS3BP |
| HILPDA | 1.35E-18 | 0.40491 | 0.531 | 0.33 | 2.62E-14 | INF/MLN | HILPDA |
| CSTB1 | 2.09E-18 | 0.256371 | 0.975 | 0.901 | 4.06E-14 | INF/MLN | CSTB |
| GARS | 3.44E-17 | 0.328524 | 0.69 | 0.521 | 6.69E-13 | INF/MLN | GARS |
| NMRK21 | 5.78E-17 | 0.250612 | 0.368 | 0.191 | 1.12E-12 | INF/MLN | NMRK2 |
| ERO1A1 | 1.06E-16 | 0.260931 | 0.46 | 0.267 | 2.07E-12 | INF/MLN | ERO1A |
| CDK2AP2 | 1.11E-15 | 0.299648 | 0.69 | 0.483 | 2.16E-11 | INF/MLN | CDK2AP2 |
| HSPA1A1 | 6.14E-15 | 0.263095 | 0.475 | 0.293 | 1.19E-10 | INF/MLN | HSPA1A |
| INSIG11 | 1.83E-13 | 0.321365 | 0.721 | 0.573 | 3.56E-09 | INF/MLN | INSIG1 |
| HSP90B1 | 2.76E-13 | 0.272915 | 0.962 | 0.899 | 5.36E-09 | INF/MLN | HSP90B1 |
| LDHA | 9.48E-06 | 0.282182 | 0.683 | 0.583 | 0.184336 | INF/MLN | LDHA |
| SLC2A1 | 1.79E-05 | 0.300342 | 0.797 | 0.723 | 0.348298 | INF/MLN | SLC2A1 |
| SCG3 | 1.70E-250 | 1.198221 | 0.799 | 0.062 | 3.31E-246 | CC | SCG3 |
| HMP19 | 6.65E-248 | 1.259672 | 0.689 | 0.031 | 1.29E-243 | CC | HMP19 |
| ELAVL4 | 1.70E-215 | 1.549002 | 0.841 | 0.108 | 3.31E-211 | CC | ELAVL4 |
| SLC17A6 | 1.19E-202 | 0.783212 | 0.47 | 0.006 | 2.31E-198 | CC | SLC17A6 |

| | | | | | | | |
|---|---|---|---|---|---|---|---|
| STMN2 | 7.85E-199 | 3.047955 | 0.966 | 0.245 | 1.53E-194 | CC | STMN2 |
| INA | 6.27E-190 | 1.478519 | 0.811 | 0.124 | 1.22E-185 | CC | INA |
| FAM57B | 5.34E-184 | 1.16574 | 0.826 | 0.13 | 1.04E-179 | CC | FAM57B |
| TAGLN3 | 8.90E-174 | 1.86471 | 0.936 | 0.267 | 1.73E-169 | CC | TAGLN3 |
| CELF3 | 8.44E-168 | 0.741132 | 0.595 | 0.049 | 1.64E-163 | CC | CELF3 |
| TTC9B | 8.17E-167 | 0.689555 | 0.466 | 0.018 | 1.59E-162 | CC | TTC9B |
| ELAVL3 | 1.99E-165 | 1.293835 | 0.913 | 0.232 | 3.87E-161 | CC | ELAVL3 |
| NSG1 | 9.31E-156 | 1.476985 | 0.837 | 0.197 | 1.81E-151 | CC | NSG1 |
| LHX9 | 6.52E-151 | 0.941112 | 0.458 | 0.024 | 1.27E-146 | CC | LHX9 |
| RTN1 | 1.36E-146 | 2.08865 | 0.864 | 0.261 | 2.64E-142 | CC | RTN1 |
| CRMP1 | 1.40E-144 | 1.513454 | 0.966 | 0.457 | 2.73E-140 | CC | CRMP1 |
| TBR1 | 5.36E-144 | 0.889293 | 0.409 | 0.016 | 1.04E-139 | CC | TBR1 |
| MLLT11 | 1.36E-143 | 1.995841 | 0.996 | 0.754 | 2.65E-139 | CC | MLLT11 |
| ELAVL2 | 6.60E-140 | 1.048545 | 0.652 | 0.096 | 1.28E-135 | CC | ELAVL2 |
| GDAP1L1 | 2.22E-137 | 1.070187 | 0.765 | 0.169 | 4.31E-133 | CC | GDAP1L1 |
| ATP1A3 | 1.64E-136 | 0.9943 | 0.723 | 0.141 | 3.20E-132 | CC | ATP1A3 |
| CHGA | 3.02E-136 | 0.583234 | 0.413 | 0.02 | 5.88E-132 | CC | CHGA |
| CHGB | 7.28E-135 | 0.597566 | 0.481 | 0.037 | 1.42E-130 | CC | CHGB |
| CD24 | 1.81E-133 | 1.634469 | 0.992 | 0.641 | 3.51E-129 | CC | CD24 |
| KIF5C | 8.40E-133 | 1.469161 | 0.92 | 0.397 | 1.63E-128 | CC | KIF5C |
| ACTL6B | 4.48E-131 | 0.531218 | 0.383 | 0.016 | 8.72E-127 | CC | ACTL6B |
| STMN4 | 5.30E-131 | 1.913102 | 0.928 | 0.385 | 1.03E-126 | CC | STMN4 |
| GAP43 | 6.40E-131 | 2.062235 | 0.89 | 0.337 | 1.25E-126 | CC | GAP43 |
| TMSB10 | 1.31E-129 | 1.532994 | 1 | 0.992 | 2.54E-125 | CC | TMSB10 |
| BASP1 | 4.67E-128 | 1.408996 | 0.996 | 0.821 | 9.08E-124 | CC | BASP1 |
| TUBB2B | 1.14E-127 | 1.534828 | 1 | 0.98 | 2.21E-123 | CC | TUBB2B |
| NRXN1 | 1.46E-124 | 0.652477 | 0.436 | 0.032 | 2.85E-120 | CC | NRXN1 |
| HN1 | 2.46E-123 | 1.274437 | 1 | 0.859 | 4.79E-119 | CC | HN1 |
| PLPPR1 | 4.35E-120 | 0.773645 | 0.424 | 0.032 | 8.46E-116 | CC | PLPPR1 |
| DCX | 6.74E-120 | 1.44035 | 0.962 | 0.527 | 1.31E-115 | CC | DCX |
| GNG3 | 7.55E-118 | 1.612284 | 0.886 | 0.414 | 1.47E-113 | CC | GNG3 |
| NCAM1 | 1.49E-117 | 0.89399 | 0.663 | 0.131 | 2.90E-113 | CC | NCAM1 |
| TUBB2A | 4.54E-117 | 1.557213 | 0.989 | 0.759 | 8.83E-113 | CC | TUBB2A |
| GPC2 | 1.11E-116 | 1.219718 | 0.83 | 0.301 | 2.17E-112 | CC | GPC2 |
| SOX4 | 1.60E-114 | 1.516226 | 0.985 | 0.872 | 3.12E-110 | CC | SOX4 |
| TUBA1A | 6.16E-113 | 1.499576 | 1 | 0.994 | 1.20E-108 | CC | TUBA1A |
| CCNI | 1.75E-111 | 0.940661 | 1 | 0.986 | 3.40E-107 | CC | CCNI |
| PCSK2 | 2.16E-111 | 0.656462 | 0.39 | 0.027 | 4.20E-107 | CC | PCSK2 |
| RUNX1T1 | 6.06E-111 | 0.6379 | 0.383 | 0.026 | 1.18E-106 | CC | RUNX1T1 |
| SRRM4 | 3.13E-110 | 0.499494 | 0.364 | 0.022 | 6.09E-106 | CC | SRRM4 |

| | | | | | | | |
|---|---|---|---|---|---|---|---|
| SMIM18 | 5.60E-110 | 0.370595 | 0.277 | 0.006 | 1.09E-105 | CC | SMIM18 |
| MAP2 | 1.30E-109 | 1.191088 | 0.898 | 0.456 | 2.54E-105 | CC | MAP2 |
| RUNDC3A | 8.70E-108 | 0.572123 | 0.47 | 0.054 | 1.69E-103 | CC | RUNDC3A |
| STMN1 | 1.54E-107 | 1.344846 | 0.996 | 0.977 | 3.00E-103 | CC | STMN1 |
| CCDC184 | 3.20E-106 | 0.684496 | 0.511 | 0.072 | 6.23E-102 | CC | CCDC184 |
| MAP1B | 1.82E-105 | 1.149066 | 0.996 | 0.965 | 3.54E-101 | CC | MAP1B |
| AKR1C1 | 3.71E-105 | 0.472889 | 0.383 | 0.03 | 7.22E-101 | CC | AKR1C1 |
| MIAT | 4.06E-104 | 1.204391 | 0.833 | 0.321 | 7.90E-100 | CC | MIAT |
| CHST8 | 1.43E-99 | 0.652622 | 0.379 | 0.032 | 2.77E-95 | CC | CHST8 |
| KLC1 | 4.68E-99 | 1.103457 | 0.898 | 0.561 | 9.10E-95 | CC | KLC1 |
| CDK5R1 | 4.41E-98 | 0.619415 | 0.473 | 0.065 | 8.57E-94 | CC | CDK5R1 |
| GRIA2 | 1.88E-97 | 0.419379 | 0.273 | 0.009 | 3.66E-93 | CC | GRIA2 |
| TERF2IP | 4.64E-97 | 0.876559 | 0.989 | 0.883 | 9.03E-93 | CC | TERF2IP |
| CD200 | 3.02E-94 | 0.909715 | 0.822 | 0.352 | 5.87E-90 | CC | CD200 |
| RBFOX2 | 1.15E-93 | 0.894885 | 0.742 | 0.263 | 2.24E-89 | CC | RBFOX2 |
| RAB3A1 | 4.30E-91 | 0.999898 | 0.902 | 0.47 | 8.36E-87 | CC | RAB3A |
| MIR7-3HG | 5.24E-91 | 0.619276 | 0.348 | 0.029 | 1.02E-86 | CC | MIR7-3HG |
| SYP1 | 2.92E-90 | 0.481869 | 0.564 | 0.116 | 5.67E-86 | CC | SYP |
| GREM2 | 1.24E-89 | 0.411288 | 0.227 | 0.005 | 2.41E-85 | CC | GREM2 |
| FXYD7 | 2.52E-89 | 0.613031 | 0.383 | 0.041 | 4.90E-85 | CC | FXYD7 |
| SOX11 | 4.46E-89 | 1.243657 | 0.958 | 0.726 | 8.69E-85 | CC | SOX11 |
| MAPT | 1.68E-88 | 0.901217 | 0.606 | 0.149 | 3.26E-84 | CC | MAPT |
| SVOP | 4.78E-88 | 0.283882 | 0.227 | 0.005 | 9.31E-84 | CC | SVOP |
| SYT5 | 1.72E-83 | 0.677785 | 0.523 | 0.103 | 3.35E-79 | CC | SYT5 |
| TUBB | 4.20E-83 | 0.962665 | 1 | 0.99 | 8.16E-79 | CC | TUBB |
| TMEM163 | 9.17E-83 | 0.441381 | 0.295 | 0.021 | 1.78E-78 | CC | TMEM163 |
| ONECUT2 | 4.99E-82 | 0.578706 | 0.258 | 0.013 | 9.71E-78 | CC | ONECUT2 |
| POU2F2 | 1.86E-81 | 0.748569 | 0.417 | 0.06 | 3.63E-77 | CC | POU2F2 |
| SYT1 | 2.00E-81 | 1.013771 | 0.826 | 0.408 | 3.90E-77 | CC | SYT1 |
| RUFY3 | 2.73E-81 | 0.792017 | 0.871 | 0.54 | 5.32E-77 | CC | RUFY3 |
| DPYSL3 | 5.39E-80 | 0.852429 | 0.663 | 0.213 | 1.05E-75 | CC | DPYSL3 |
| NEUROD1 | 3.29E-78 | 0.55618 | 0.242 | 0.012 | 6.40E-74 | CC | NEUROD1 |
| ST18 | 5.63E-78 | 0.380268 | 0.231 | 0.01 | 1.09E-73 | CC | ST18 |
| UCHL1 | 1.04E-77 | 1.078443 | 0.97 | 0.903 | 2.03E-73 | CC | UCHL1 |
| GDI1 | 1.18E-77 | 0.863322 | 0.894 | 0.628 | 2.30E-73 | CC | GDI1 |
| RTN4 | 2.12E-76 | 0.737549 | 0.992 | 0.929 | 4.12E-72 | CC | RTN4 |
| MARCKSL | 2.52E-76 | 0.814838 | 1 | 0.977 | 4.90E-72 | CC | MARCKSL1 |
| MYT1L | 3.95E-76 | 0.26365 | 0.205 | 0.006 | 7.68E-72 | CC | MYT1L |
| MYT1 | 1.42E-75 | 0.360668 | 0.288 | 0.023 | 2.76E-71 | CC | MYT1 |
| GPM6A | 1.30E-74 | 1.018832 | 0.867 | 0.531 | 2.53E-70 | CC | GPM6A |

| | | | | | | | |
|---|---|---|---|---|---|---|---|
| EBF1 | 1.37E-74 | 0.481897 | 0.231 | 0.011 | 2.67E-70 | CC | EBF1 |
| YWHAQ | 2.18E-73 | 0.702307 | 0.989 | 0.972 | 4.24E-69 | CC | YWHAQ |
| CTD-3064 | 4.95E-73 | 0.267489 | 0.178 | 0.003 | 9.63E-69 | CC | CTD-3064C13.1 |
| NNAT | 8.47E-73 | 2.067138 | 0.871 | 0.642 | 1.65E-68 | CC | NNAT |
| DCC | 1.17E-71 | 0.686093 | 0.485 | 0.106 | 2.27E-67 | CC | DCC |
| RTN3 | 1.57E-71 | 0.688558 | 0.985 | 0.924 | 3.05E-67 | CC | RTN3 |
| ACTG1 | 5.02E-71 | 0.652593 | 1 | 1 | 9.76E-67 | CC | ACTG1 |
| PAFAH1B3 | 1.49E-70 | 0.950747 | 0.928 | 0.764 | 2.91E-66 | CC | PAFAH1B3 |
| OLFM1 | 2.13E-70 | 0.742657 | 0.515 | 0.126 | 4.14E-66 | CC | OLFM1 |
| SCN3B | 4.15E-70 | 0.399121 | 0.269 | 0.022 | 8.08E-66 | CC | SCN3B |
| MARCKS | 4.42E-70 | 0.733818 | 0.973 | 0.9 | 8.61E-66 | CC | MARCKS |
| ATCAY | 1.18E-69 | 0.566245 | 0.508 | 0.124 | 2.30E-65 | CC | ATCAY |
| MAB21L1 | 5.76E-69 | 0.723784 | 0.295 | 0.03 | 1.12E-64 | CC | MAB21L1 |
| TMEM35 | 6.05E-69 | 0.703712 | 0.617 | 0.205 | 1.18E-64 | CC | TMEM35 |
| NHLH1 | 7.93E-69 | 0.831199 | 0.284 | 0.027 | 1.54E-64 | CC | NHLH1 |
| VAMP2 | 8.24E-69 | 0.73025 | 0.989 | 0.93 | 1.60E-64 | CC | VAMP2 |
| APLP1 | 8.97E-69 | 0.904226 | 0.951 | 0.857 | 1.75E-64 | CC | APLP1 |
| EBF2 | 2.58E-68 | 0.300947 | 0.182 | 0.005 | 5.03E-64 | CC | EBF2 |
| SLCO5A1 | 2.82E-68 | 0.390165 | 0.265 | 0.022 | 5.48E-64 | CC | SLCO5A1 |
| CELF4 | 2.25E-67 | 0.811395 | 0.519 | 0.137 | 4.38E-63 | CC | CELF4 |
| EBF3 | 3.12E-67 | 0.255449 | 0.186 | 0.006 | 6.07E-63 | CC | EBF3 |
| GNG2 | 4.01E-67 | 0.714308 | 0.686 | 0.285 | 7.79E-63 | CC | GNG2 |
| PCBP4 | 1.59E-66 | 1.082106 | 0.867 | 0.616 | 3.10E-62 | CC | PCBP4 |
| DCLK1 | 3.83E-66 | 0.481218 | 0.348 | 0.049 | 7.44E-62 | CC | DCLK1 |
| ACTB | 5.42E-66 | 0.687543 | 1 | 1 | 1.05E-61 | CC | ACTB |
| DMTN | 1.37E-65 | 0.480656 | 0.375 | 0.062 | 2.66E-61 | CC | DMTN |
| MAP4K4 | 4.45E-65 | 0.788566 | 0.659 | 0.256 | 8.65E-61 | CC | MAP4K4 |
| NRN1 | 4.49E-65 | 0.925007 | 0.686 | 0.268 | 8.73E-61 | CC | NRN1 |
| UBE2E3 | 8.80E-65 | 0.76017 | 0.943 | 0.813 | 1.71E-60 | CC | UBE2E3 |
| H3F3A | 2.77E-64 | 0.52301 | 1 | 0.998 | 5.40E-60 | CC | H3F3A |
| 6-Sep | 4.06E-64 | 0.739394 | 0.739 | 0.358 | 7.89E-60 | CC | 6-Sep |
| NFASC | 6.18E-64 | 0.476899 | 0.379 | 0.064 | 1.20E-59 | CC | NFASC |
| GJD2 | 8.77E-64 | 0.253056 | 0.174 | 0.005 | 1.71E-59 | CC | GJD2 |
| AKR1C2 | 2.95E-63 | 0.326813 | 0.231 | 0.017 | 5.74E-59 | CC | AKR1C2 |
| LRRN3 | 4.93E-63 | 0.367784 | 0.258 | 0.023 | 9.60E-59 | CC | LRRN3 |
| SYT4 | 5.22E-63 | 0.581963 | 0.409 | 0.074 | 1.02E-58 | CC | SYT4 |
| SNAP25 | 2.04E-62 | 0.604464 | 0.527 | 0.15 | 3.96E-58 | CC | SNAP25 |
| L1CAM | 2.42E-61 | 0.377607 | 0.314 | 0.042 | 4.70E-57 | CC | L1CAM |
| INSM1 | 1.95E-60 | 0.440514 | 0.288 | 0.034 | 3.80E-56 | CC | INSM1 |
| NEFM | 2.29E-60 | 1.480336 | 0.576 | 0.194 | 4.45E-56 | CC | NEFM |

| | | | | | | | |
|---|---|---|---|---|---|---|---|
| ENC1 | 8.12E-59 | 0.89505 | 0.439 | 0.105 | 1.58E-54 | CC | ENC1 |
| SEZ6L2 | 1.29E-58 | 0.718674 | 0.837 | 0.529 | 2.51E-54 | CC | SEZ6L2 |
| LHX1 | 2.20E-57 | 0.508007 | 0.205 | 0.014 | 4.28E-53 | CC | LHX1 |
| TTC3 | 2.70E-57 | 0.65421 | 0.973 | 0.914 | 5.26E-53 | CC | TTC3 |
| PGM2L1 | 2.71E-57 | 0.732862 | 0.591 | 0.222 | 5.27E-53 | CC | PGM2L1 |
| ADGRG1 | 7.28E-57 | 0.687393 | 0.655 | 0.284 | 1.42E-52 | CC | ADGRG1 |
| TMSB15A | 1.09E-56 | 0.74445 | 0.924 | 0.622 | 2.12E-52 | CC | TMSB15A |
| PBX3 | 1.62E-56 | 0.80952 | 0.511 | 0.158 | 3.15E-52 | CC | PBX3 |
| ATAT1 | 5.02E-56 | 0.69341 | 0.807 | 0.538 | 9.77E-52 | CC | ATAT1 |
| TUBB3 | 5.54E-56 | 0.478861 | 0.409 | 0.093 | 1.08E-51 | CC | TUBB3 |
| DBN1 | 1.25E-55 | 0.704853 | 0.875 | 0.651 | 2.42E-51 | CC | DBN1 |
| RP11-445I | 1.84E-55 | 0.381807 | 0.182 | 0.01 | 3.59E-51 | CC | RP11-445F12.1 |
| BEX1 | 7.04E-55 | 0.762764 | 0.92 | 0.8 | 1.37E-50 | CC | BEX1 |
| PHYHIPL | 1.37E-54 | 0.577999 | 0.473 | 0.133 | 2.66E-50 | CC | PHYHIPL |
| SCG2 | 2.13E-54 | 0.608368 | 0.348 | 0.063 | 4.15E-50 | CC | SCG2 |
| EOMES | 3.68E-54 | 0.483819 | 0.186 | 0.012 | 7.17E-50 | CC | EOMES |
| ATL1 | 9.14E-54 | 0.530189 | 0.481 | 0.14 | 1.78E-49 | CC | ATL1 |
| NEFL | 9.26E-53 | 1.32854 | 0.492 | 0.151 | 1.80E-48 | CC | NEFL |
| THSD7A | 2.33E-52 | 0.480446 | 0.333 | 0.061 | 4.53E-48 | CC | THSD7A |
| CFL1 | 2.49E-52 | 0.525558 | 1 | 0.997 | 4.85E-48 | CC | CFL1 |
| NREP | 6.06E-52 | 0.714439 | 0.939 | 0.819 | 1.18E-47 | CC | NREP |
| SYN1 | 1.15E-51 | 0.261217 | 0.205 | 0.017 | 2.23E-47 | CC | SYN1 |
| RCAN2 | 2.19E-51 | 0.409043 | 0.288 | 0.044 | 4.25E-47 | CC | RCAN2 |
| CELF5 | 2.22E-51 | 0.547495 | 0.466 | 0.137 | 4.31E-47 | CC | CELF5 |
| EIF4G2 | 2.39E-51 | 0.558143 | 0.992 | 0.94 | 4.65E-47 | CC | EIF4G2 |
| CDKN2D | 2.57E-51 | 0.681294 | 0.621 | 0.271 | 5.00E-47 | CC | CDKN2D |
| YWHAH | 3.41E-51 | 0.726728 | 0.879 | 0.744 | 6.64E-47 | CC | YWHAH |
| AKAP6 | 6.91E-51 | 0.338605 | 0.299 | 0.049 | 1.34E-46 | CC | AKAP6 |
| DLL3 | 1.28E-50 | 0.921517 | 0.542 | 0.197 | 2.48E-46 | CC | DLL3 |
| CXADR | 1.65E-50 | 0.635931 | 0.951 | 0.854 | 3.21E-46 | CC | CXADR |
| DAAM1 | 1.84E-50 | 0.69249 | 0.864 | 0.602 | 3.59E-46 | CC | DAAM1 |
| MPP3 | 1.77E-49 | 0.301777 | 0.242 | 0.029 | 3.44E-45 | CC | MPP3 |
| FAM110A | 2.61E-49 | 0.340137 | 0.303 | 0.051 | 5.08E-45 | CC | FAM110A |
| SLIT1 | 3.03E-49 | 0.296474 | 0.197 | 0.017 | 5.89E-45 | CC | SLIT1 |
| FXYD6 | 1.06E-48 | 0.848614 | 0.89 | 0.769 | 2.06E-44 | CC | FXYD6 |
| ASPHD1 | 1.73E-48 | 0.357605 | 0.299 | 0.052 | 3.37E-44 | CC | ASPHD1 |
| GDAP1 | 4.36E-48 | 0.535067 | 0.553 | 0.21 | 8.48E-44 | CC | GDAP1 |
| YWHAG | 8.30E-48 | 0.654415 | 0.788 | 0.559 | 1.61E-43 | CC | YWHAG |
| CEP170 | 1.19E-47 | 0.629329 | 0.576 | 0.249 | 2.32E-43 | CC | CEP170 |
| NKAIN1 | 1.34E-47 | 0.336438 | 0.314 | 0.058 | 2.62E-43 | CC | NKAIN1 |

| | | | | | | | |
|---|---|---|---|---|---|---|---|
| CPE | 1.86E-47 | 0.643832 | 0.886 | 0.692 | 3.62E-43 | CC | CPE |
| NOVA1 | 5.80E-47 | 0.746361 | 0.758 | 0.501 | 1.13E-42 | CC | NOVA1 |
| DTD1 | 7.60E-47 | 0.61945 | 0.837 | 0.679 | 1.48E-42 | CC | DTD1 |
| ATP6V1G2 | 1.36E-46 | 0.653841 | 0.629 | 0.307 | 2.65E-42 | CC | ATP6V1G2 |
| GABRB3 | 2.44E-46 | 0.30109 | 0.288 | 0.049 | 4.74E-42 | CC | GABRB3 |
| LBH | 3.11E-46 | 0.606412 | 0.621 | 0.295 | 6.05E-42 | CC | LBH |
| KLF7 | 1.69E-45 | 0.565122 | 0.523 | 0.197 | 3.28E-41 | CC | KLF7 |
| CREG2 | 2.44E-45 | 0.286303 | 0.17 | 0.013 | 4.75E-41 | CC | CREG2 |
| GNG4 | 2.70E-45 | 0.471914 | 0.477 | 0.153 | 5.25E-41 | CC | GNG4 |
| SMARCA4 | 3.65E-45 | 0.609827 | 0.913 | 0.814 | 7.11E-41 | CC | SMARCA4 |
| TUBB4A | 4.22E-45 | 0.509361 | 0.432 | 0.131 | 8.21E-41 | CC | TUBB4A |
| NHLH2 | 4.67E-45 | 0.326069 | 0.14 | 0.007 | 9.09E-41 | CC | NHLH2 |
| CNTN2 | 5.14E-45 | 0.28834 | 0.148 | 0.008 | 1.00E-40 | CC | CNTN2 |
| PHF21B | 5.41E-45 | 0.423082 | 0.402 | 0.11 | 1.05E-40 | CC | PHF21B |
| C10orf35 | 1.52E-44 | 0.640082 | 0.667 | 0.379 | 2.96E-40 | CC | C10orf35 |
| STMN3 | 2.27E-44 | 0.587262 | 0.648 | 0.322 | 4.41E-40 | CC | STMN3 |
| SUMO2 | 2.81E-44 | 0.398984 | 1 | 0.994 | 5.46E-40 | CC | SUMO2 |
| SRRM3 | 2.91E-44 | 0.345048 | 0.273 | 0.046 | 5.66E-40 | CC | SRRM3 |
| STXBP1 | 3.43E-44 | 0.575067 | 0.583 | 0.263 | 6.68E-40 | CC | STXBP1 |
| MAB21L2 | 3.65E-43 | 0.327563 | 0.155 | 0.011 | 7.10E-39 | CC | MAB21L2 |
| CACNA2D | 4.85E-43 | 0.297082 | 0.227 | 0.03 | 9.43E-39 | CC | CACNA2D1 |
| GNAO1 | 1.18E-42 | 0.44169 | 0.379 | 0.102 | 2.31E-38 | CC | GNAO1 |
| HDAC2 | 1.42E-42 | 0.533273 | 0.958 | 0.906 | 2.75E-38 | CC | HDAC2 |
| PCSK1N | 1.48E-42 | 0.781221 | 0.939 | 0.805 | 2.87E-38 | CC | PCSK1N |
| PODXL2 | 1.49E-42 | 0.661667 | 0.761 | 0.526 | 2.89E-38 | CC | PODXL2 |
| TRIM36 | 2.46E-42 | 0.479519 | 0.447 | 0.147 | 4.78E-38 | CC | TRIM36 |
| TFAP2A | 7.30E-42 | 0.284734 | 0.125 | 0.005 | 1.42E-37 | CC | TFAP2A |
| EIF1B | 1.60E-41 | 0.485372 | 0.981 | 0.944 | 3.12E-37 | CC | EIF1B |
| FAM65B | 1.66E-41 | 0.310639 | 0.216 | 0.029 | 3.23E-37 | CC | FAM65B |
| PPP1R1A | 2.14E-41 | 0.533181 | 0.542 | 0.233 | 4.17E-37 | CC | PPP1R1A |
| C4orf48 | 3.14E-41 | 0.537823 | 0.955 | 0.918 | 6.11E-37 | CC | C4orf48 |
| NCAN | 3.48E-41 | 0.35176 | 0.295 | 0.06 | 6.76E-37 | CC | NCAN |
| BLCAP | 3.75E-41 | 0.744958 | 0.731 | 0.513 | 7.29E-37 | CC | BLCAP |
| CALM3 | 4.14E-41 | 0.517449 | 0.97 | 0.94 | 8.06E-37 | CC | CALM3 |
| DCTN2 | 6.10E-41 | 0.51938 | 0.89 | 0.799 | 1.19E-36 | CC | DCTN2 |
| TSPAN7 | 5.05E-40 | 0.696486 | 0.538 | 0.237 | 9.83E-36 | CC | TSPAN7 |
| GABARAP | 6.99E-40 | 0.454595 | 0.955 | 0.944 | 1.36E-35 | CC | GABARAPL2 |
| CCDC112 | 9.06E-40 | 0.58483 | 0.629 | 0.343 | 1.76E-35 | CC | CCDC112 |
| GSTA4 | 1.03E-39 | 0.616441 | 0.83 | 0.695 | 2.01E-35 | CC | GSTA4 |
| LZTS1 | 8.08E-39 | 0.257688 | 0.189 | 0.023 | 1.57E-34 | CC | LZTS1 |

| | | | | | | | |
|---|---|---|---|---|---|---|---|
| FAM155A | 8.94E-39 | 0.366861 | 0.269 | 0.052 | 1.74E-34 | CC | FAM155A |
| CADM3 | 1.09E-38 | 0.478559 | 0.489 | 0.194 | 2.13E-34 | CC | CADM3 |
| NOL4 | 1.16E-38 | 0.272524 | 0.193 | 0.024 | 2.26E-34 | CC | NOL4 |
| REEP1 | 1.44E-38 | 0.290636 | 0.261 | 0.049 | 2.80E-34 | CC | REEP1 |
| NRCAM | 2.58E-38 | 0.35572 | 0.292 | 0.063 | 5.01E-34 | CC | NRCAM |
| KIF3C | 5.40E-38 | 0.353675 | 0.295 | 0.066 | 1.05E-33 | CC | KIF3C |
| AMER2 | 2.47E-37 | 0.350345 | 0.299 | 0.068 | 4.80E-33 | CC | AMER2 |
| GLRX | 8.03E-37 | 0.455949 | 0.375 | 0.114 | 1.56E-32 | CC | GLRX |
| MAP4 | 1.68E-36 | 0.657447 | 0.648 | 0.403 | 3.27E-32 | CC | MAP4 |
| KLHL35 | 2.36E-36 | 0.574761 | 0.367 | 0.112 | 4.59E-32 | CC | KLHL35 |
| DPYSL2 | 3.07E-36 | 0.598087 | 0.617 | 0.346 | 5.98E-32 | CC | DPYSL2 |
| RNF165 | 3.62E-36 | 0.366633 | 0.371 | 0.111 | 7.04E-32 | CC | RNF165 |
| ANK3 | 5.31E-36 | 0.454682 | 0.409 | 0.144 | 1.03E-31 | CC | ANK3 |
| RELN | 6.47E-36 | 0.250609 | 0.121 | 0.007 | 1.26E-31 | CC | RELN |
| KIDINS22( | 7.79E-36 | 0.586656 | 0.572 | 0.299 | 1.52E-31 | CC | KIDINS220 |
| PAK3 | 1.17E-35 | 0.610115 | 0.693 | 0.432 | 2.28E-31 | CC | PAK3 |
| MEIS2 | 1.37E-35 | 0.913663 | 0.398 | 0.133 | 2.66E-31 | CC | MEIS2 |
| PTMS | 1.54E-35 | 0.457288 | 0.989 | 0.952 | 3.00E-31 | CC | PTMS |
| CORO1A | 1.59E-35 | 0.370982 | 0.333 | 0.091 | 3.09E-31 | CC | CORO1A |
| RTN2 | 1.81E-35 | 0.276816 | 0.231 | 0.041 | 3.53E-31 | CC | RTN2 |
| YWHAZ | 3.20E-35 | 0.545864 | 0.913 | 0.883 | 6.22E-31 | CC | YWHAZ |
| FNBP1L | 5.57E-35 | 0.637822 | 0.678 | 0.463 | 1.08E-30 | CC | FNBP1L |
| SBK1 | 5.61E-35 | 0.407763 | 0.402 | 0.138 | 1.09E-30 | CC | SBK1 |
| RPS6KL1 | 8.35E-35 | 0.374258 | 0.345 | 0.099 | 1.62E-30 | CC | RPS6KL1 |
| DPYSL4 | 8.95E-35 | 0.540775 | 0.686 | 0.434 | 1.74E-30 | CC | DPYSL4 |
| CLASP2 | 9.55E-35 | 0.502097 | 0.489 | 0.209 | 1.86E-30 | CC | CLASP2 |
| KLHDC8A | 1.52E-34 | 0.458252 | 0.314 | 0.081 | 2.95E-30 | CC | KLHDC8A |
| CDC42 | 1.58E-34 | 0.595816 | 0.875 | 0.819 | 3.08E-30 | CC | CDC42 |
| MIR124-2I | 1.94E-34 | 0.342852 | 0.36 | 0.108 | 3.77E-30 | CC | MIR124-2HG |
| GAD2 | 2.37E-34 | 0.445348 | 0.136 | 0.012 | 4.60E-30 | CC | GAD2 |
| 3-Sep | 3.14E-34 | 0.361721 | 0.322 | 0.088 | 6.10E-30 | CC | 3-Sep |
| PRKX | 3.19E-34 | 0.517976 | 0.576 | 0.296 | 6.21E-30 | CC | PRKX |
| TCEAL7 | 3.33E-34 | 0.694853 | 0.807 | 0.634 | 6.48E-30 | CC | TCEAL7 |
| FAM89B | 3.38E-34 | 0.558363 | 0.591 | 0.335 | 6.57E-30 | CC | FAM89B |
| CAMK2N1 | 4.88E-34 | 0.530175 | 0.386 | 0.128 | 9.50E-30 | CC | CAMK2N1 |
| CIRBP | 6.05E-34 | 0.401822 | 0.989 | 0.988 | 1.18E-29 | CC | CIRBP |
| GTF2I | 1.09E-33 | 0.47267 | 0.867 | 0.716 | 2.11E-29 | CC | GTF2I |
| DYNC1I2 | 1.17E-33 | 0.55322 | 0.856 | 0.752 | 2.27E-29 | CC | DYNC1I2 |
| ACTR1A | 1.26E-33 | 0.529793 | 0.716 | 0.536 | 2.45E-29 | CC | ACTR1A |
| DDX5 | 2.29E-33 | 0.395153 | 0.992 | 0.983 | 4.45E-29 | CC | DDX5 |

| | | | | | | | |
|---|---|---|---|---|---|---|---|
| TMEFF2 | 2.79E-33 | 0.355013 | 0.201 | 0.032 | 5.42E-29 | CC | TMEFF2 |
| YWHAB | 2.89E-33 | 0.54751 | 0.894 | 0.8 | 5.63E-29 | CC | YWHAB |
| DLX5 | 3.29E-33 | 0.337008 | 0.102 | 0.005 | 6.41E-29 | CC | DLX5 |
| SH3BP5 | 4.27E-33 | 0.506303 | 0.504 | 0.221 | 8.31E-29 | CC | SH3BP5 |
| SCG5 | 2.20E-32 | 0.642218 | 0.617 | 0.363 | 4.27E-28 | CC | SCG5 |
| C16orf45 | 3.56E-32 | 0.514106 | 0.591 | 0.346 | 6.93E-28 | CC | C16orf45 |
| PBX1 | 7.93E-32 | 0.664383 | 0.705 | 0.492 | 1.54E-27 | CC | PBX1 |
| DDAH2 | 1.80E-31 | 0.49531 | 0.951 | 0.928 | 3.51E-27 | CC | DDAH2 |
| CACNB3 | 2.27E-31 | 0.331923 | 0.326 | 0.096 | 4.42E-27 | CC | CACNB3 |
| ZNF428 | 3.54E-31 | 0.542753 | 0.92 | 0.881 | 6.88E-27 | CC | ZNF428 |
| RBP1 | 4.92E-31 | 0.825508 | 0.826 | 0.647 | 9.57E-27 | CC | RBP1 |
| NAV1 | 5.40E-31 | 0.482667 | 0.398 | 0.145 | 1.05E-26 | CC | NAV1 |
| HIP1R | 6.16E-31 | 0.287467 | 0.28 | 0.071 | 1.20E-26 | CC | HIP1R |
| PTMA | 7.02E-31 | 0.278689 | 1 | 1 | 1.37E-26 | CC | PTMA |
| NDRG4 | 8.06E-31 | 0.497465 | 0.523 | 0.264 | 1.57E-26 | CC | NDRG4 |
| RGMB | 1.28E-30 | 0.514619 | 0.356 | 0.122 | 2.50E-26 | CC | RGMB |
| C3orf14 | 1.47E-30 | 0.480587 | 0.538 | 0.277 | 2.86E-26 | CC | C3orf14 |
| DCTN3 | 2.00E-30 | 0.426095 | 0.886 | 0.837 | 3.89E-26 | CC | DCTN3 |
| SSTR2 | 3.61E-30 | 0.624599 | 0.269 | 0.067 | 7.03E-26 | CC | SSTR2 |
| TMEM57 | 6.21E-30 | 0.427968 | 0.455 | 0.198 | 1.21E-25 | CC | TMEM57 |
| TBCB | 6.30E-30 | 0.465474 | 0.898 | 0.875 | 1.23E-25 | CC | TBCB |
| JPH4 | 6.62E-30 | 0.411159 | 0.348 | 0.117 | 1.29E-25 | CC | JPH4 |
| IGFBPL1 | 9.85E-30 | 0.358928 | 0.311 | 0.09 | 1.92E-25 | CC | IGFBPL1 |
| TMEM59L | 1.54E-29 | 0.64764 | 0.625 | 0.4 | 3.00E-25 | CC | TMEM59L |
| ACAP3 | 3.39E-29 | 0.400762 | 0.432 | 0.181 | 6.60E-25 | CC | ACAP3 |
| MAPK8 | 3.66E-29 | 0.497661 | 0.553 | 0.321 | 7.12E-25 | CC | MAPK8 |
| NACAD | 6.47E-29 | 0.332584 | 0.303 | 0.089 | 1.26E-24 | CC | NACAD |
| STX12 | 1.12E-28 | 0.479984 | 0.625 | 0.409 | 2.17E-24 | CC | STX12 |
| SULT4A1 | 1.15E-28 | 0.281528 | 0.246 | 0.059 | 2.23E-24 | CC | SULT4A1 |
| KIF5A | 1.63E-28 | 0.262055 | 0.227 | 0.05 | 3.17E-24 | CC | KIF5A |
| RNF24 | 1.89E-28 | 0.459323 | 0.553 | 0.309 | 3.68E-24 | CC | RNF24 |
| STARD3NI | 2.24E-28 | 0.416197 | 0.848 | 0.755 | 4.36E-24 | CC | STARD3NL |
| REM2 | 2.41E-28 | 0.324913 | 0.322 | 0.101 | 4.68E-24 | CC | REM2 |
| DUSP8 | 5.12E-28 | 0.283201 | 0.25 | 0.062 | 9.95E-24 | CC | DUSP8 |
| MAP6 | 6.27E-28 | 0.488916 | 0.598 | 0.373 | 1.22E-23 | CC | MAP6 |
| RGS17 | 1.71E-27 | 0.293507 | 0.258 | 0.068 | 3.33E-23 | CC | RGS17 |
| DLX2 | 2.88E-27 | 0.356125 | 0.14 | 0.018 | 5.60E-23 | CC | DLX2 |
| SERTAD4 | 3.07E-27 | 0.26943 | 0.174 | 0.03 | 5.97E-23 | CC | SERTAD4 |
| MEX3B | 3.96E-27 | 0.446434 | 0.447 | 0.202 | 7.71E-23 | CC | MEX3B |
| SLC38A1 | 4.79E-27 | 0.467785 | 0.61 | 0.367 | 9.32E-23 | CC | SLC38A1 |

| | | | | | | | |
|---|---|---|---|---|---|---|---|
| AP1S2 | 6.20E-27 | 0.755934 | 0.754 | 0.677 | 1.21E-22 | CC | AP1S2 |
| SVBP | 6.55E-27 | 0.481685 | 0.746 | 0.611 | 1.27E-22 | CC | SVBP |
| CRABP1 | 8.69E-27 | 1.980953 | 0.686 | 0.563 | 1.69E-22 | CC | CRABP1 |
| NOVA2 | 1.53E-26 | 0.416447 | 0.428 | 0.192 | 2.97E-22 | CC | NOVA2 |
| CNTFR | 1.70E-26 | 0.482308 | 0.648 | 0.431 | 3.30E-22 | CC | CNTFR |
| LHX5 | 1.81E-26 | 0.693428 | 0.405 | 0.173 | 3.52E-22 | CC | LHX5 |
| OPTN | 2.48E-26 | 0.439686 | 0.489 | 0.243 | 4.83E-22 | CC | OPTN |
| DYNLRB1 | 2.66E-26 | 0.367539 | 0.932 | 0.884 | 5.17E-22 | CC | DYNLRB1 |
| FAM127A | 3.40E-26 | 0.451053 | 0.803 | 0.684 | 6.61E-22 | CC | FAM127A |
| ZC2HC1A | 3.40E-26 | 0.481831 | 0.561 | 0.339 | 6.62E-22 | CC | ZC2HC1A |
| RNF11 | 8.37E-26 | 0.432185 | 0.701 | 0.555 | 1.63E-21 | CC | RNF11 |
| MYL6 | 8.57E-26 | 0.4181 | 0.989 | 0.989 | 1.67E-21 | CC | MYL6 |
| CLTB | 8.91E-26 | 0.439449 | 0.761 | 0.607 | 1.73E-21 | CC | CLTB |
| CHN1 | 1.21E-25 | 0.480209 | 0.545 | 0.314 | 2.35E-21 | CC | CHN1 |
| DYNLT1 | 1.56E-25 | 0.386958 | 0.936 | 0.874 | 3.04E-21 | CC | DYNLT1 |
| MAPRE3 | 1.59E-25 | 0.426443 | 0.489 | 0.259 | 3.09E-21 | CC | MAPRE3 |
| DYNC1H1 | 1.69E-25 | 0.551557 | 0.652 | 0.489 | 3.29E-21 | CC | DYNC1H1 |
| CALM1 | 1.77E-25 | 0.47683 | 0.985 | 0.963 | 3.44E-21 | CC | CALM1 |
| CRIP2 | 1.94E-25 | 0.569181 | 0.67 | 0.511 | 3.78E-21 | CC | CRIP2 |
| IP6K2 | 2.43E-25 | 0.478291 | 0.75 | 0.658 | 4.73E-21 | CC | IP6K2 |
| ARID4A | 2.56E-25 | 0.464943 | 0.568 | 0.352 | 4.98E-21 | CC | ARID4A |
| PLPPR2 | 3.79E-25 | 0.388003 | 0.367 | 0.149 | 7.38E-21 | CC | PLPPR2 |
| CNRIP1 | 4.50E-25 | 0.337776 | 0.352 | 0.133 | 8.76E-21 | CC | CNRIP1 |
| PLPPR3 | 1.02E-24 | 0.461675 | 0.61 | 0.41 | 1.98E-20 | CC | PLPPR3 |
| PPP1R17 | 1.37E-24 | 0.312859 | 0.133 | 0.018 | 2.66E-20 | CC | PPP1R17 |
| BEX2 | 1.55E-24 | 0.468865 | 0.814 | 0.693 | 3.02E-20 | CC | BEX2 |
| ETFB | 1.82E-24 | 0.518725 | 0.867 | 0.817 | 3.54E-20 | CC | ETFB |
| HNRNPA0 | 2.31E-24 | 0.349126 | 0.962 | 0.958 | 4.49E-20 | CC | HNRNPA0 |
| MALAT12 | 2.41E-24 | 0.325016 | 1 | 1 | 4.70E-20 | CC | MALAT1 |
| PPP1R18 | 2.76E-24 | 0.425223 | 0.432 | 0.207 | 5.37E-20 | CC | PPP1R18 |
| MEAF6 | 4.00E-24 | 0.397729 | 0.883 | 0.855 | 7.78E-20 | CC | MEAF6 |
| SV2A | 4.54E-24 | 0.429948 | 0.458 | 0.237 | 8.84E-20 | CC | SV2A |
| PPP2R5B1 | 5.69E-24 | 0.419064 | 0.5 | 0.263 | 1.11E-19 | CC | PPP2R5B |
| MEX3A | 5.92E-24 | 0.527197 | 0.542 | 0.335 | 1.15E-19 | CC | MEX3A |
| H3F3B1 | 6.21E-24 | 0.250275 | 1 | 1 | 1.21E-19 | CC | H3F3B |
| NMNAT2 | 6.50E-24 | 0.270481 | 0.242 | 0.067 | 1.26E-19 | CC | NMNAT2 |
| PFDN4 | 6.97E-24 | 0.40045 | 0.826 | 0.751 | 1.36E-19 | CC | PFDN4 |
| KIFAP3 | 7.10E-24 | 0.479747 | 0.538 | 0.337 | 1.38E-19 | CC | KIFAP3 |
| HES6 | 8.71E-24 | 1.225125 | 0.633 | 0.417 | 1.70E-19 | CC | HES6 |
| DNAJB6 | 1.10E-23 | 0.401798 | 0.905 | 0.885 | 2.15E-19 | CC | DNAJB6 |

| | | | | | | | |
|---|---|---|---|---|---|---|---|
| VPS28 | 1.44E-23 | 0.424859 | 0.826 | 0.769 | 2.80E-19 | CC | VPS28 |
| RFTN1 | 1.53E-23 | 0.287487 | 0.167 | 0.032 | 2.97E-19 | CC | RFTN1 |
| KIF1A | 1.94E-23 | 0.416143 | 0.731 | 0.574 | 3.77E-19 | CC | KIF1A |
| AUTS2 | 2.33E-23 | 0.359751 | 0.364 | 0.152 | 4.53E-19 | CC | AUTS2 |
| ATP6V0E2 | 2.95E-23 | 0.462021 | 0.731 | 0.628 | 5.75E-19 | CC | ATP6V0E2 |
| SNN | 3.35E-23 | 0.383245 | 0.443 | 0.225 | 6.51E-19 | CC | SNN |
| NEDD4L | 6.15E-23 | 0.342127 | 0.326 | 0.124 | 1.20E-18 | CC | NEDD4L |
| RUFY2 | 7.51E-23 | 0.367692 | 0.405 | 0.19 | 1.46E-18 | CC | RUFY2 |
| ACOT7 | 7.52E-23 | 0.475166 | 0.614 | 0.432 | 1.46E-18 | CC | ACOT7 |
| RAB6A2 | 8.25E-23 | 0.41583 | 0.807 | 0.721 | 1.61E-18 | CC | RAB6A |
| SYT11 | 8.57E-23 | 0.44642 | 0.598 | 0.422 | 1.67E-18 | CC | SYT11 |
| SARAF | 1.21E-22 | 0.398341 | 0.958 | 0.96 | 2.36E-18 | CC | SARAF |
| SNRPN | 1.54E-22 | 0.331844 | 0.977 | 0.934 | 2.99E-18 | CC | SNRPN |
| BCL11A | 3.12E-22 | 0.440783 | 0.322 | 0.125 | 6.08E-18 | CC | BCL11A |
| FYN | 3.39E-22 | 0.438048 | 0.576 | 0.388 | 6.59E-18 | CC | FYN |
| APC2 | 4.10E-22 | 0.254264 | 0.223 | 0.061 | 7.98E-18 | CC | APC2 |
| C14orf132 | 6.96E-22 | 0.397768 | 0.432 | 0.219 | 1.35E-17 | CC | C14orf132 |
| ABRACL | 8.62E-22 | 0.473947 | 0.678 | 0.532 | 1.68E-17 | CC | ABRACL |
| CSDE1 | 9.72E-22 | 0.327433 | 0.913 | 0.907 | 1.89E-17 | CC | CSDE1 |
| CTNNA2 | 1.07E-21 | 0.255531 | 0.25 | 0.075 | 2.08E-17 | CC | CTNNA2 |
| BARHL2 | 1.12E-21 | 0.270956 | 0.258 | 0.074 | 2.18E-17 | CC | BARHL2 |
| FSD1 | 1.21E-21 | 0.386578 | 0.489 | 0.28 | 2.35E-17 | CC | FSD1 |
| DHX36 | 1.63E-21 | 0.41145 | 0.784 | 0.697 | 3.18E-17 | CC | DHX36 |
| DPF1 | 1.65E-21 | 0.396989 | 0.371 | 0.165 | 3.20E-17 | CC | DPF1 |
| CYTH2 | 1.69E-21 | 0.401519 | 0.667 | 0.535 | 3.30E-17 | CC | CYTH2 |
| USP11 | 2.05E-21 | 0.36748 | 0.943 | 0.923 | 3.99E-17 | CC | USP11 |
| MTSS1 | 2.24E-21 | 0.250253 | 0.235 | 0.069 | 4.36E-17 | CC | MTSS1 |
| RP11-247( | 3.22E-21 | 0.254411 | 0.212 | 0.058 | 6.26E-17 | CC | RP11-247C2.2 |
| PTH2 | 4.22E-21 | 0.504719 | 0.159 | 0.033 | 8.21E-17 | CC | PTH2 |
| DNAJB5 | 4.91E-21 | 0.321784 | 0.428 | 0.214 | 9.55E-17 | CC | DNAJB5 |
| SERINC1 | 4.99E-21 | 0.424505 | 0.742 | 0.63 | 9.71E-17 | CC | SERINC1 |
| BTF3L4 | 6.38E-21 | 0.410737 | 0.867 | 0.85 | 1.24E-16 | CC | BTF3L4 |
| THY1 | 6.60E-21 | 0.282113 | 0.284 | 0.099 | 1.28E-16 | CC | THY1 |
| ACTR10 | 6.92E-21 | 0.409283 | 0.754 | 0.692 | 1.35E-16 | CC | ACTR10 |
| RP4-665J2 | 8.91E-21 | 0.408598 | 0.409 | 0.182 | 1.73E-16 | CC | RP4-665J23.1 |
| UBE2D1 | 1.04E-20 | 0.48029 | 0.655 | 0.544 | 2.02E-16 | CC | UBE2D1 |
| SEMA6A | 1.21E-20 | 0.408757 | 0.436 | 0.233 | 2.36E-16 | CC | SEMA6A |
| ST7 | 1.71E-20 | 0.326278 | 0.303 | 0.118 | 3.33E-16 | CC | ST7 |
| ZNF738 | 2.05E-20 | 0.415461 | 0.674 | 0.536 | 4.00E-16 | CC | ZNF738 |
| EIF4A21 | 2.64E-20 | 0.303483 | 0.996 | 0.991 | 5.14E-16 | CC | EIF4A2 |

| | | | | | | | |
|---|---|---|---|---|---|---|---|
| ARG2 | 2.79E-20 | 0.37235 | 0.424 | 0.218 | 5.42E-16 | CC | ARG2 |
| UBA1 | 3.40E-20 | 0.359586 | 0.852 | 0.793 | 6.62E-16 | CC | UBA1 |
| CORO7 | 3.79E-20 | 0.268751 | 0.299 | 0.114 | 7.37E-16 | CC | CORO7 |
| TMEM176 | 9.68E-20 | 0.367815 | 0.231 | 0.072 | 1.88E-15 | CC | TMEM176A |
| SORBS2 | 9.91E-20 | 0.449333 | 0.477 | 0.288 | 1.93E-15 | CC | SORBS2 |
| TMSB4X | 1.24E-19 | 0.364074 | 1 | 1 | 2.42E-15 | CC | TMSB4X |
| 7-Sep | 1.45E-19 | 0.362837 | 0.955 | 0.904 | 2.82E-15 | CC | 7-Sep |
| MAP1A | 1.50E-19 | 0.445607 | 0.625 | 0.494 | 2.91E-15 | CC | MAP1A |
| APBB3 | 1.52E-19 | 0.382624 | 0.367 | 0.178 | 2.95E-15 | CC | APBB3 |
| FAIM2 | 1.58E-19 | 0.274119 | 0.235 | 0.075 | 3.07E-15 | CC | FAIM2 |
| CAMK2B | 1.99E-19 | 0.260078 | 0.239 | 0.079 | 3.88E-15 | CC | CAMK2B |
| PPP2R1A | 2.32E-19 | 0.294525 | 0.939 | 0.933 | 4.52E-15 | CC | PPP2R1A |
| AKIRIN2 | 2.70E-19 | 0.412118 | 0.568 | 0.419 | 5.24E-15 | CC | AKIRIN2 |
| ARPC5 | 3.73E-19 | 0.410409 | 0.792 | 0.771 | 7.26E-15 | CC | ARPC5 |
| TMEM169 | 3.80E-19 | 0.262372 | 0.239 | 0.079 | 7.39E-15 | CC | TMEM169 |
| PKIA | 4.36E-19 | 0.354699 | 0.413 | 0.218 | 8.48E-15 | CC | PKIA |
| WHSC1L1 | 4.58E-19 | 0.413848 | 0.742 | 0.679 | 8.91E-15 | CC | WHSC1L1 |
| FEZ1 | 4.89E-19 | 0.430103 | 0.78 | 0.727 | 9.51E-15 | CC | FEZ1 |
| LHX2 | 5.16E-19 | 0.516003 | 0.348 | 0.148 | 1.00E-14 | CC | LHX2 |
| SMARCB1 | 5.64E-19 | 0.335264 | 0.909 | 0.867 | 1.10E-14 | CC | SMARCB1 |
| POU3F2 | 6.68E-19 | 0.41269 | 0.402 | 0.198 | 1.30E-14 | CC | POU3F2 |
| FSCN1 | 7.13E-19 | 0.342439 | 0.92 | 0.912 | 1.39E-14 | CC | FSCN1 |
| BTG1 | 7.47E-19 | 0.390319 | 0.905 | 0.878 | 1.45E-14 | CC | BTG1 |
| TH | 1.26E-18 | 0.375625 | 0.163 | 0.038 | 2.44E-14 | CC | TH |
| GNB1 | 1.36E-18 | 0.350392 | 0.716 | 0.615 | 2.65E-14 | CC | GNB1 |
| PRKACB | 1.39E-18 | 0.369391 | 0.348 | 0.164 | 2.70E-14 | CC | PRKACB |
| ENO22 | 1.49E-18 | 0.392564 | 0.875 | 0.83 | 2.89E-14 | CC | ENO2 |
| KIAA0895l | 1.51E-18 | 0.319743 | 0.33 | 0.147 | 2.94E-14 | CC | KIAA0895L |
| ADD2 | 1.53E-18 | 0.252822 | 0.265 | 0.097 | 2.97E-14 | CC | ADD2 |
| KIF1B | 1.81E-18 | 0.38241 | 0.545 | 0.37 | 3.52E-14 | CC | KIF1B |
| ZNF821 | 2.72E-18 | 0.364013 | 0.458 | 0.271 | 5.29E-14 | CC | ZNF821 |
| TSPAN13 | 3.21E-18 | 0.42809 | 0.773 | 0.669 | 6.25E-14 | CC | TSPAN13 |
| MKRN1 | 3.41E-18 | 0.387161 | 0.758 | 0.707 | 6.63E-14 | CC | MKRN1 |
| LDOC1 | 5.68E-18 | 0.405515 | 0.614 | 0.491 | 1.10E-13 | CC | LDOC1 |
| DPYSL5 | 6.48E-18 | 0.359472 | 0.402 | 0.214 | 1.26E-13 | CC | DPYSL5 |
| FAXC | 7.03E-18 | 0.254159 | 0.261 | 0.097 | 1.37E-13 | CC | FAXC |
| KMT2E | 7.16E-18 | 0.367037 | 0.845 | 0.806 | 1.39E-13 | CC | KMT2E |
| BAG6 | 1.17E-17 | 0.3396 | 0.754 | 0.717 | 2.28E-13 | CC | BAG6 |
| APBB1 | 1.30E-17 | 0.410934 | 0.58 | 0.446 | 2.53E-13 | CC | APBB1 |
| PTPRN2 | 1.39E-17 | 0.251881 | 0.269 | 0.104 | 2.70E-13 | CC | PTPRN2 |

| | | | | | | | |
|---|---|---|---|---|---|---|---|
| MYL6B | 1.53E-17 | 0.306816 | 0.898 | 0.869 | 2.97E-13 | CC | MYL6B |
| DYNLL2 | 1.64E-17 | 0.388747 | 0.515 | 0.346 | 3.19E-13 | CC | DYNLL2 |
| CKB | 1.81E-17 | 0.379629 | 1 | 0.981 | 3.51E-13 | CC | CKB |
| PFDN21 | 2.23E-17 | 0.29463 | 0.867 | 0.879 | 4.35E-13 | CC | PFDN2 |
| CAMKV | 2.39E-17 | 0.314529 | 0.235 | 0.083 | 4.65E-13 | CC | CAMKV |
| DUSP4 | 3.23E-17 | 0.409172 | 0.511 | 0.328 | 6.28E-13 | CC | DUSP4 |
| SHD | 3.30E-17 | 0.349959 | 0.261 | 0.1 | 6.42E-13 | CC | SHD |
| SLC2A6 | 3.35E-17 | 0.257371 | 0.295 | 0.125 | 6.52E-13 | CC | SLC2A6 |
| DRAXIN | 5.05E-17 | 0.406227 | 0.527 | 0.347 | 9.83E-13 | CC | DRAXIN |
| CLCN4 | 5.26E-17 | 0.250297 | 0.227 | 0.079 | 1.02E-12 | CC | CLCN4 |
| GADD45G | 6.34E-17 | 0.579511 | 0.489 | 0.29 | 1.23E-12 | CC | GADD45G |
| RAB2A | 8.11E-17 | 0.310124 | 0.902 | 0.88 | 1.58E-12 | CC | RAB2A |
| SNCG | 8.51E-17 | 0.897273 | 0.386 | 0.217 | 1.65E-12 | CC | SNCG |
| KHDRBS1 | 9.66E-17 | 0.319703 | 0.883 | 0.9 | 1.88E-12 | CC | KHDRBS1 |
| SIAH1 | 1.00E-16 | 0.399619 | 0.553 | 0.414 | 1.95E-12 | CC | SIAH1 |
| SRGAP3 | 1.09E-16 | 0.412839 | 0.568 | 0.416 | 2.12E-12 | CC | SRGAP3 |
| LMBR1L | 1.28E-16 | 0.32968 | 0.398 | 0.219 | 2.49E-12 | CC | LMBR1L |
| IGDCC3 | 1.43E-16 | 0.580751 | 0.561 | 0.405 | 2.78E-12 | CC | IGDCC3 |
| PBRM1 | 1.46E-16 | 0.365 | 0.515 | 0.35 | 2.84E-12 | CC | PBRM1 |
| SCAMP5 | 1.95E-16 | 0.296457 | 0.33 | 0.159 | 3.78E-12 | CC | SCAMP5 |
| VGF1 | 2.19E-16 | 0.377925 | 0.254 | 0.096 | 4.26E-12 | CC | VGF |
| LCOR | 2.82E-16 | 0.344373 | 0.458 | 0.292 | 5.48E-12 | CC | LCOR |
| SMARCE1 | 3.46E-16 | 0.303183 | 0.773 | 0.697 | 6.74E-12 | CC | SMARCE1 |
| MIDN | 3.63E-16 | 0.420768 | 0.602 | 0.484 | 7.05E-12 | CC | MIDN |
| ATP1B1 | 4.18E-16 | 0.404119 | 0.72 | 0.612 | 8.12E-12 | CC | ATP1B1 |
| CORO1C | 4.92E-16 | 0.349081 | 0.455 | 0.286 | 9.58E-12 | CC | CORO1C |
| VBP1 | 5.05E-16 | 0.282515 | 0.886 | 0.847 | 9.83E-12 | CC | VBP1 |
| CSNK1E | 6.18E-16 | 0.420823 | 0.708 | 0.634 | 1.20E-11 | CC | CSNK1E |
| KIF3A | 6.31E-16 | 0.400357 | 0.519 | 0.378 | 1.23E-11 | CC | KIF3A |
| CLIP3 | 7.31E-16 | 0.317591 | 0.458 | 0.28 | 1.42E-11 | CC | CLIP3 |
| PNMA1 | 7.37E-16 | 0.359398 | 0.697 | 0.626 | 1.43E-11 | CC | PNMA1 |
| ATRX | 7.64E-16 | 0.402269 | 0.652 | 0.527 | 1.49E-11 | CC | ATRX |
| ZCCHC12 | 8.33E-16 | 0.299286 | 0.314 | 0.146 | 1.62E-11 | CC | ZCCHC12 |
| DLGAP4 | 9.76E-16 | 0.439598 | 0.572 | 0.462 | 1.90E-11 | CC | DLGAP4 |
| VEZT | 1.05E-15 | 0.352742 | 0.568 | 0.439 | 2.04E-11 | CC | VEZT |
| AP1S1 | 1.19E-15 | 0.381783 | 0.667 | 0.586 | 2.32E-11 | CC | AP1S1 |
| ZC4H2 | 1.56E-15 | 0.305695 | 0.371 | 0.198 | 3.03E-11 | CC | ZC4H2 |
| NDUFB8 | 1.74E-15 | 0.319596 | 0.924 | 0.92 | 3.39E-11 | CC | NDUFB8 |
| DCTN1 | 1.89E-15 | 0.350662 | 0.553 | 0.41 | 3.68E-11 | CC | DCTN1 |
| RAB33A | 2.18E-15 | 0.326776 | 0.348 | 0.181 | 4.24E-11 | CC | RAB33A |

| | | | | | | | |
|---|---|---|---|---|---|---|---|
| RP11-382/ | 2.19E-15 | 0.378712 | 0.583 | 0.474 | 4.26E-11 | CC | RP11-382A20.3 |
| FABP5 | 2.32E-15 | 0.350035 | 0.655 | 0.49 | 4.51E-11 | CC | FABP5 |
| RAB6B | 2.50E-15 | 0.283116 | 0.348 | 0.182 | 4.87E-11 | CC | RAB6B |
| BRK1 | 2.64E-15 | 0.265177 | 0.951 | 0.964 | 5.13E-11 | CC | BRK1 |
| PLEKHO1 | 2.66E-15 | 0.352762 | 0.534 | 0.379 | 5.18E-11 | CC | PLEKHO1 |
| STX7 | 2.87E-15 | 0.323805 | 0.477 | 0.317 | 5.59E-11 | CC | STX7 |
| RBM3 | 3.09E-15 | 0.315601 | 0.818 | 0.779 | 6.01E-11 | CC | RBM3 |
| NARF | 3.64E-15 | 0.335292 | 0.773 | 0.741 | 7.07E-11 | CC | NARF |
| NPDC1 | 3.69E-15 | 0.327326 | 0.727 | 0.637 | 7.17E-11 | CC | NPDC1 |
| LEMD1 | 4.01E-15 | 0.326929 | 0.398 | 0.22 | 7.81E-11 | CC | LEMD1 |
| KDM6B | 4.12E-15 | 0.326983 | 0.413 | 0.244 | 8.02E-11 | CC | KDM6B |
| FHL1 | 4.44E-15 | 0.315745 | 0.667 | 0.552 | 8.64E-11 | CC | FHL1 |
| PHLDA1 | 4.47E-15 | 0.565883 | 0.424 | 0.263 | 8.70E-11 | CC | PHLDA1 |
| FAM49B | 4.59E-15 | 0.339995 | 0.659 | 0.572 | 8.93E-11 | CC | FAM49B |
| FAM219A | 5.43E-15 | 0.298845 | 0.345 | 0.178 | 1.06E-10 | CC | FAM219A |
| NAPB | 7.11E-15 | 0.312278 | 0.307 | 0.148 | 1.38E-10 | CC | NAPB |
| PTPRN | 8.26E-15 | 0.303758 | 0.269 | 0.115 | 1.61E-10 | CC | PTPRN |
| DYNC1LI2 | 1.03E-14 | 0.381448 | 0.625 | 0.523 | 2.01E-10 | CC | DYNC1LI2 |
| TDG | 1.11E-14 | 0.421284 | 0.644 | 0.57 | 2.16E-10 | CC | TDG |
| TMEM178 | 1.77E-14 | 0.298761 | 0.28 | 0.128 | 3.45E-10 | CC | TMEM178A |
| BRD3 | 1.86E-14 | 0.362212 | 0.629 | 0.547 | 3.62E-10 | CC | BRD3 |
| BRWD1 | 1.90E-14 | 0.359568 | 0.568 | 0.447 | 3.70E-10 | CC | BRWD1 |
| APBA2 | 2.20E-14 | 0.337205 | 0.383 | 0.221 | 4.28E-10 | CC | APBA2 |
| ARPC1A | 2.25E-14 | 0.330585 | 0.754 | 0.713 | 4.37E-10 | CC | ARPC1A |
| KDM5B | 2.53E-14 | 0.334412 | 0.655 | 0.552 | 4.91E-10 | CC | KDM5B |
| DST | 2.81E-14 | 0.381222 | 0.394 | 0.237 | 5.47E-10 | CC | DST |
| CDC42EP3 | 3.05E-14 | 0.33949 | 0.341 | 0.184 | 5.93E-10 | CC | CDC42EP3 |
| SMARCC2 | 3.21E-14 | 0.357014 | 0.508 | 0.367 | 6.25E-10 | CC | SMARCC2 |
| CADM2 | 3.37E-14 | 0.33247 | 0.386 | 0.218 | 6.56E-10 | CC | CADM2 |
| MMD | 3.43E-14 | 0.359711 | 0.508 | 0.366 | 6.67E-10 | CC | MMD |
| XPR1 | 3.72E-14 | 0.377622 | 0.428 | 0.27 | 7.24E-10 | CC | XPR1 |
| KCTD13 | 3.93E-14 | 0.250732 | 0.311 | 0.154 | 7.65E-10 | CC | KCTD13 |
| RPAIN | 4.24E-14 | 0.325303 | 0.803 | 0.801 | 8.25E-10 | CC | RPAIN |
| ATP5O | 4.30E-14 | 0.265082 | 0.951 | 0.962 | 8.37E-10 | CC | ATP5O |
| ITFG1 | 4.91E-14 | 0.314043 | 0.583 | 0.479 | 9.56E-10 | CC | ITFG1 |
| PRKAR2B | 1.02E-13 | 0.329914 | 0.345 | 0.191 | 1.98E-09 | CC | PRKAR2B |
| CAPRIN1 | 1.04E-13 | 0.303494 | 0.701 | 0.637 | 2.02E-09 | CC | CAPRIN1 |
| OCIAD2 | 1.06E-13 | 0.492625 | 0.621 | 0.55 | 2.07E-09 | CC | OCIAD2 |
| C19orf43 | 1.10E-13 | 0.277134 | 0.902 | 0.891 | 2.14E-09 | CC | C19orf43 |
| MAP1LC3/ | 1.18E-13 | 0.451774 | 0.576 | 0.442 | 2.30E-09 | CC | MAP1LC3A |

| | | | | | | | |
|---|---|---|---|---|---|---|---|
| GRIPAP1 | 1.30E-13 | 0.346255 | 0.485 | 0.348 | 2.52E-09 | CC | GRIPAP1 |
| ARMCX3 | 1.31E-13 | 0.352378 | 0.761 | 0.743 | 2.55E-09 | CC | ARMCX3 |
| TAOK3 | 1.53E-13 | 0.345453 | 0.47 | 0.325 | 2.98E-09 | CC | TAOK3 |
| CELF1 | 1.58E-13 | 0.316811 | 0.648 | 0.579 | 3.07E-09 | CC | CELF1 |
| NUDT3 | 1.64E-13 | 0.364619 | 0.568 | 0.462 | 3.19E-09 | CC | NUDT3 |
| APP | 1.76E-13 | 0.318658 | 0.894 | 0.877 | 3.42E-09 | CC | APP |
| EPC1 | 1.86E-13 | 0.396093 | 0.617 | 0.543 | 3.63E-09 | CC | EPC1 |
| MAPK8IP1 | 1.94E-13 | 0.333837 | 0.371 | 0.225 | 3.77E-09 | CC | MAPK8IP1 |
| RSF1 | 2.47E-13 | 0.310592 | 0.689 | 0.63 | 4.81E-09 | CC | RSF1 |
| TMTC4 | 2.95E-13 | 0.287048 | 0.341 | 0.187 | 5.74E-09 | CC | TMTC4 |
| GABARAP | 3.42E-13 | 0.338849 | 0.598 | 0.506 | 6.66E-09 | CC | GABARAPL1 |
| BZW2 | 3.59E-13 | 0.362556 | 0.617 | 0.539 | 6.99E-09 | CC | BZW2 |
| ARL8A | 4.89E-13 | 0.341944 | 0.489 | 0.372 | 9.51E-09 | CC | ARL8A |
| CHRNA3 | 5.03E-13 | 0.270275 | 0.155 | 0.049 | 9.79E-09 | CC | CHRNA3 |
| MAPK10 | 5.44E-13 | 0.34879 | 0.496 | 0.357 | 1.06E-08 | CC | MAPK10 |
| REEP2 | 5.80E-13 | 0.353989 | 0.523 | 0.417 | 1.13E-08 | CC | REEP2 |
| EPS8L1 | 6.64E-13 | 0.272548 | 0.28 | 0.134 | 1.29E-08 | CC | EPS8L1 |
| TSC22D1 | 8.06E-13 | 0.326701 | 0.905 | 0.898 | 1.57E-08 | CC | TSC22D1 |
| CADM1 | 9.13E-13 | 0.432821 | 0.636 | 0.559 | 1.78E-08 | CC | CADM1 |
| DNER | 9.21E-13 | 0.395053 | 0.803 | 0.695 | 1.79E-08 | CC | DNER |
| SH3BGRL: | 9.85E-13 | 0.348251 | 0.913 | 0.927 | 1.92E-08 | CC | SH3BGRL3 |
| GRB2 | 9.91E-13 | 0.352811 | 0.572 | 0.479 | 1.93E-08 | CC | GRB2 |
| OS9 | 1.24E-12 | 0.310999 | 0.705 | 0.633 | 2.41E-08 | CC | OS9 |
| TPGS2 | 1.31E-12 | 0.357662 | 0.735 | 0.72 | 2.55E-08 | CC | TPGS2 |
| TTLL7 | 1.45E-12 | 0.291856 | 0.326 | 0.18 | 2.82E-08 | CC | TTLL7 |
| VAT11 | 1.60E-12 | 0.321076 | 0.633 | 0.538 | 3.12E-08 | CC | VAT1 |
| BEX4 | 1.76E-12 | 0.296052 | 0.856 | 0.847 | 3.42E-08 | CC | BEX4 |
| PJA1 | 2.05E-12 | 0.351843 | 0.534 | 0.434 | 3.99E-08 | CC | PJA1 |
| SLC32A1 | 2.63E-12 | 0.333281 | 0.129 | 0.036 | 5.11E-08 | CC | SLC32A1 |
| MLLT4 | 2.64E-12 | 0.368527 | 0.489 | 0.356 | 5.14E-08 | CC | MLLT4 |
| AMN1 | 2.71E-12 | 0.301469 | 0.428 | 0.289 | 5.28E-08 | CC | AMN1 |
| NR2F2 | 2.74E-12 | 0.501692 | 0.523 | 0.424 | 5.33E-08 | CC | NR2F2 |
| ARID4B | 2.80E-12 | 0.359578 | 0.614 | 0.539 | 5.45E-08 | CC | ARID4B |
| CARHSP1 | 3.38E-12 | 0.276483 | 0.807 | 0.728 | 6.58E-08 | CC | CARHSP1 |
| PAPSS1 | 3.82E-12 | 0.287363 | 0.663 | 0.59 | 7.42E-08 | CC | PAPSS1 |
| OAZ2 | 3.98E-12 | 0.276314 | 0.83 | 0.808 | 7.73E-08 | CC | OAZ2 |
| SSBP3 | 4.01E-12 | 0.34245 | 0.587 | 0.503 | 7.81E-08 | CC | SSBP3 |
| CASP3 | 5.58E-12 | 0.639522 | 0.553 | 0.491 | 1.09E-07 | CC | CASP3 |
| DYNC1LI1 | 6.08E-12 | 0.329884 | 0.583 | 0.516 | 1.18E-07 | CC | DYNC1LI1 |
| NDFIP1 | 6.27E-12 | 0.253384 | 0.917 | 0.897 | 1.22E-07 | CC | NDFIP1 |

| | | | | | | | |
|---|---|---|---|---|---|---|---|
| TRAF4 | 6.39E-12 | 0.311618 | 0.652 | 0.584 | 1.24E-07 | CC | TRAF4 |
| MTF2 | 6.82E-12 | 0.327953 | 0.652 | 0.6 | 1.33E-07 | CC | MTF2 |
| WSB1 | 6.98E-12 | 0.259901 | 0.966 | 0.941 | 1.36E-07 | CC | WSB1 |
| TP53I11 | 7.12E-12 | 0.321775 | 0.311 | 0.173 | 1.38E-07 | CC | TP53I11 |
| IGLON5 | 7.84E-12 | 0.290855 | 0.318 | 0.177 | 1.53E-07 | CC | IGLON5 |
| H2AFY2 | 8.02E-12 | 0.296211 | 0.705 | 0.632 | 1.56E-07 | CC | H2AFY2 |
| ZFHX3 | 8.24E-12 | 0.37697 | 0.394 | 0.255 | 1.60E-07 | CC | ZFHX3 |
| MUM1 | 8.30E-12 | 0.316527 | 0.492 | 0.357 | 1.62E-07 | CC | MUM1 |
| CMIP | 8.49E-12 | 0.318785 | 0.402 | 0.262 | 1.65E-07 | CC | CMIP |
| DHPS | 8.62E-12 | 0.290012 | 0.689 | 0.661 | 1.68E-07 | CC | DHPS |
| ARPC2 | 8.97E-12 | 0.254584 | 0.894 | 0.915 | 1.74E-07 | CC | ARPC2 |
| JAKMIP2 | 9.28E-12 | 0.280831 | 0.348 | 0.206 | 1.81E-07 | CC | JAKMIP2 |
| GSK3B | 1.01E-11 | 0.330365 | 0.538 | 0.44 | 1.97E-07 | CC | GSK3B |
| SARS1 | 1.21E-11 | 0.36043 | 0.723 | 0.72 | 2.36E-07 | CC | SARS |
| PHF14 | 1.27E-11 | 0.316379 | 0.629 | 0.559 | 2.47E-07 | CC | PHF14 |
| MGEA5 | 1.33E-11 | 0.34617 | 0.557 | 0.478 | 2.59E-07 | CC | MGEA5 |
| MFNG | 1.38E-11 | 0.285255 | 0.17 | 0.062 | 2.69E-07 | CC | MFNG |
| NAP1L3 | 1.47E-11 | 0.343264 | 0.587 | 0.51 | 2.85E-07 | CC | NAP1L3 |
| PSD3 | 1.80E-11 | 0.264331 | 0.311 | 0.173 | 3.50E-07 | CC | PSD3 |
| SPATS2 | 1.80E-11 | 0.303133 | 0.496 | 0.381 | 3.50E-07 | CC | SPATS2 |
| AES | 1.85E-11 | 0.310544 | 0.591 | 0.519 | 3.59E-07 | CC | AES |
| OLFM2 | 2.27E-11 | 0.372215 | 0.428 | 0.305 | 4.41E-07 | CC | OLFM2 |
| TIA1 | 2.57E-11 | 0.286659 | 0.511 | 0.391 | 4.99E-07 | CC | TIA1 |
| NTM | 3.13E-11 | 0.278552 | 0.197 | 0.08 | 6.09E-07 | CC | NTM |
| WSB2 | 3.38E-11 | 0.296247 | 0.428 | 0.297 | 6.58E-07 | CC | WSB2 |
| ATP6V1B2 | 4.44E-11 | 0.269707 | 0.659 | 0.584 | 8.63E-07 | CC | ATP6V1B2 |
| DCTN6 | 5.46E-11 | 0.287506 | 0.621 | 0.551 | 1.06E-06 | CC | DCTN6 |
| NEUROG1 | 5.59E-11 | 0.375101 | 0.148 | 0.049 | 1.09E-06 | CC | NEUROG1 |
| FKBP1A | 6.19E-11 | 0.268243 | 0.886 | 0.873 | 1.20E-06 | CC | FKBP1A |
| TMEM55A | 6.22E-11 | 0.328647 | 0.549 | 0.453 | 1.21E-06 | CC | TMEM55A |
| ANKRD46 | 6.66E-11 | 0.33208 | 0.564 | 0.505 | 1.30E-06 | CC | ANKRD46 |
| RAB11B | 6.73E-11 | 0.27253 | 0.814 | 0.822 | 1.31E-06 | CC | RAB11B |
| TBPL1 | 8.53E-11 | 0.310993 | 0.538 | 0.457 | 1.66E-06 | CC | TBPL1 |
| NAP1L5 | 9.56E-11 | 0.309291 | 0.333 | 0.206 | 1.86E-06 | CC | NAP1L5 |
| CAMSAP1 | 9.60E-11 | 0.257417 | 0.284 | 0.154 | 1.87E-06 | CC | CAMSAP1 |
| TMEM14A | 9.66E-11 | 0.361719 | 0.614 | 0.564 | 1.88E-06 | CC | TMEM14A |
| TPD52 | 1.00E-10 | 0.276356 | 0.242 | 0.118 | 1.95E-06 | CC | TPD52 |
| TSG101 | 1.04E-10 | 0.305791 | 0.693 | 0.681 | 2.02E-06 | CC | TSG101 |
| CSRNP3 | 1.15E-10 | 0.312004 | 0.405 | 0.277 | 2.24E-06 | CC | CSRNP3 |
| TRIB2 | 1.23E-10 | 0.304352 | 0.383 | 0.249 | 2.40E-06 | CC | TRIB2 |

| | | | | | | | |
|---|---|---|---|---|---|---|---|
| BAZ2B | 1.26E-10 | 0.3068 | 0.481 | 0.357 | 2.45E-06 | CC | BAZ2B |
| MTMR9 | 1.30E-10 | 0.275936 | 0.417 | 0.288 | 2.54E-06 | CC | MTMR9 |
| MAP7D1 | 1.44E-10 | 0.287544 | 0.572 | 0.522 | 2.80E-06 | CC | MAP7D1 |
| CCDC167 | 1.47E-10 | 0.277084 | 0.739 | 0.708 | 2.86E-06 | CC | CCDC167 |
| TRIM2 | 1.79E-10 | 0.357569 | 0.527 | 0.43 | 3.47E-06 | CC | TRIM2 |
| PARP6 | 1.81E-10 | 0.299509 | 0.402 | 0.278 | 3.52E-06 | CC | PARP6 |
| JUP | 1.84E-10 | 0.257186 | 0.292 | 0.161 | 3.58E-06 | CC | JUP |
| RHOBTB3 | 1.97E-10 | 0.306255 | 0.542 | 0.431 | 3.83E-06 | CC | RHOBTB3 |
| SMDT1 | 2.25E-10 | 0.25821 | 0.784 | 0.787 | 4.38E-06 | CC | SMDT1 |
| ACHE | 2.32E-10 | 0.301763 | 0.273 | 0.144 | 4.52E-06 | CC | ACHE |
| PALM | 3.19E-10 | 0.321141 | 0.402 | 0.287 | 6.21E-06 | CC | PALM |
| PAK2 | 3.46E-10 | 0.329036 | 0.644 | 0.636 | 6.73E-06 | CC | PAK2 |
| KLC2 | 3.72E-10 | 0.279149 | 0.428 | 0.312 | 7.24E-06 | CC | KLC2 |
| GFOD2 | 5.24E-10 | 0.289851 | 0.356 | 0.236 | 1.02E-05 | CC | GFOD2 |
| LHX5-AS1 | 5.51E-10 | 0.873356 | 0.481 | 0.394 | 1.07E-05 | CC | LHX5-AS1 |
| EFNA3 | 7.58E-10 | 0.300417 | 0.504 | 0.406 | 1.47E-05 | CC | EFNA3 |
| AASDHPP | 1.01E-09 | 0.286812 | 0.648 | 0.624 | 1.96E-05 | CC | AASDHPPT |
| RND2 | 1.03E-09 | 0.305268 | 0.443 | 0.33 | 2.00E-05 | CC | RND2 |
| PTN | 1.03E-09 | 0.34349 | 0.564 | 0.444 | 2.00E-05 | CC | PTN |
| GRAMD1A | 1.05E-09 | 0.276149 | 0.458 | 0.353 | 2.05E-05 | CC | GRAMD1A |
| BLOC1S2 | 1.06E-09 | 0.284682 | 0.674 | 0.656 | 2.06E-05 | CC | BLOC1S2 |
| LIMD2 | 1.17E-09 | 0.333014 | 0.5 | 0.402 | 2.27E-05 | CC | LIMD2 |
| BICD1 | 1.32E-09 | 0.29501 | 0.402 | 0.289 | 2.57E-05 | CC | BICD1 |
| TNRC6C | 1.75E-09 | 0.305666 | 0.36 | 0.246 | 3.40E-05 | CC | TNRC6C |
| PPP1R14E | 2.15E-09 | 0.306352 | 0.492 | 0.399 | 4.18E-05 | CC | PPP1R14B |
| ZCCHC17 | 2.17E-09 | 0.274014 | 0.659 | 0.626 | 4.22E-05 | CC | ZCCHC17 |
| HPCAL1 | 2.25E-09 | 0.311849 | 0.402 | 0.293 | 4.37E-05 | CC | HPCAL1 |
| YTHDF2 | 2.62E-09 | 0.285997 | 0.758 | 0.754 | 5.10E-05 | CC | YTHDF2 |
| MYCBP2 | 2.74E-09 | 0.323243 | 0.345 | 0.225 | 5.34E-05 | CC | MYCBP2 |
| ZMAT2 | 2.87E-09 | 0.26124 | 0.583 | 0.526 | 5.59E-05 | CC | ZMAT2 |
| GNAQ | 3.78E-09 | 0.301884 | 0.398 | 0.284 | 7.36E-05 | CC | GNAQ |
| NKAIN4 | 4.61E-09 | 0.384961 | 0.591 | 0.535 | 8.96E-05 | CC | NKAIN4 |
| COX20 | 4.78E-09 | 0.302071 | 0.576 | 0.511 | 9.31E-05 | CC | COX20 |
| KMT2A | 6.05E-09 | 0.292198 | 0.477 | 0.391 | 0.000118 | CC | KMT2A |
| AP2M1 | 7.14E-09 | 0.251035 | 0.898 | 0.91 | 0.000139 | CC | AP2M1 |
| USP22 | 7.49E-09 | 0.304961 | 0.534 | 0.482 | 0.000146 | CC | USP22 |
| ANKRD12 | 7.61E-09 | 0.287912 | 0.527 | 0.451 | 0.000148 | CC | ANKRD12 |
| CELF2 | 8.22E-09 | 0.385695 | 0.496 | 0.433 | 0.00016 | CC | CELF2 |
| SLC39A6 | 8.25E-09 | 0.271604 | 0.636 | 0.605 | 0.00016 | CC | SLC39A6 |
| SESTD1 | 8.39E-09 | 0.287961 | 0.436 | 0.34 | 0.000163 | CC | SESTD1 |

| | | | | | | | |
|---|---|---|---|---|---|---|---|
| VPS26B | 8.50E-09 | 0.294319 | 0.439 | 0.349 | 0.000165 | CC | VPS26B |
| BCL7A | 9.61E-09 | 0.275419 | 0.466 | 0.354 | 0.000187 | CC | BCL7A |
| HSDL1 | 9.63E-09 | 0.305471 | 0.447 | 0.356 | 0.000187 | CC | HSDL1 |
| SRSF8 | 1.04E-08 | 0.274996 | 0.629 | 0.595 | 0.000203 | CC | SRSF8 |
| RSBN1 | 1.05E-08 | 0.250051 | 0.549 | 0.492 | 0.000204 | CC | RSBN1 |
| PGAP1 | 1.21E-08 | 0.30259 | 0.39 | 0.272 | 0.000235 | CC | PGAP1 |
| C14orf37 | 1.25E-08 | 0.274946 | 0.386 | 0.279 | 0.000243 | CC | C14orf37 |
| PAFAH1B1 | 1.51E-08 | 0.274357 | 0.625 | 0.606 | 0.000294 | CC | PAFAH1B1 |
| PPP2R5E | 1.63E-08 | 0.27423 | 0.523 | 0.449 | 0.000316 | CC | PPP2R5E |
| PCDH9 | 1.82E-08 | 0.335371 | 0.311 | 0.185 | 0.000354 | CC | PCDH9 |
| NEUROG2 | 2.68E-08 | 0.279744 | 0.144 | 0.057 | 0.000522 | CC | NEUROG2 |
| PPP3CA | 4.17E-08 | 0.25281 | 0.318 | 0.205 | 0.000812 | CC | PPP3CA |
| TRAPPC6B | 4.34E-08 | 0.268827 | 0.345 | 0.239 | 0.000845 | CC | TRAPPC6B |
| MOAP1 | 5.25E-08 | 0.266683 | 0.5 | 0.426 | 0.001021 | CC | MOAP1 |
| KLHL241 | 5.27E-08 | 0.26372 | 0.424 | 0.33 | 0.001024 | CC | KLHL24 |
| INAFM1 | 6.01E-08 | 0.395065 | 0.413 | 0.339 | 0.001169 | CC | INAFM1 |
| EPB41 | 7.55E-08 | 0.315663 | 0.288 | 0.187 | 0.001469 | CC | EPB41 |
| FGF13 | 8.09E-08 | 0.281385 | 0.326 | 0.221 | 0.001573 | CC | FGF13 |
| TLE1 | 9.05E-08 | 0.406189 | 0.542 | 0.487 | 0.00176 | CC | TLE1 |
| UBE2L6 | 1.03E-07 | 0.394748 | 0.606 | 0.615 | 0.001994 | CC | UBE2L6 |
| NPTX2 | 1.08E-07 | 0.255559 | 0.129 | 0.051 | 0.002093 | CC | NPTX2 |
| PSMD14 | 1.13E-07 | 0.272249 | 0.72 | 0.716 | 0.002197 | CC | PSMD14 |
| TACC2 | 1.21E-07 | 0.266904 | 0.326 | 0.223 | 0.002351 | CC | TACC2 |
| BTBD10 | 1.22E-07 | 0.275716 | 0.398 | 0.311 | 0.002383 | CC | BTBD10 |
| EFNB3 | 1.29E-07 | 0.280376 | 0.39 | 0.297 | 0.002509 | CC | EFNB3 |
| PRRT2 | 1.35E-07 | 0.262142 | 0.443 | 0.363 | 0.002627 | CC | PRRT2 |
| RERE | 1.55E-07 | 0.265366 | 0.489 | 0.411 | 0.003022 | CC | RERE |
| TMEM132 | 1.97E-07 | 0.274771 | 0.455 | 0.377 | 0.003824 | CC | TMEM132A |
| ERV3-1 | 2.36E-07 | 0.251416 | 0.409 | 0.323 | 0.004594 | CC | ERV3-1 |
| ZCCHC11 | 2.42E-07 | 0.256381 | 0.576 | 0.535 | 0.0047 | CC | ZCCHC11 |
| KMT5B | 2.67E-07 | 0.259192 | 0.515 | 0.46 | 0.005189 | CC | KMT5B |
| MCTS1 | 2.70E-07 | 0.250917 | 0.625 | 0.598 | 0.005258 | CC | MCTS1 |
| IGBP1 | 3.09E-07 | 0.252581 | 0.598 | 0.568 | 0.006014 | CC | IGBP1 |
| KCNQ2 | 3.50E-07 | 0.255244 | 0.341 | 0.241 | 0.006808 | CC | KCNQ2 |
| HTATSF1 | 4.21E-07 | 0.313718 | 0.629 | 0.639 | 0.008196 | CC | HTATSF1 |
| PAK1 | 4.30E-07 | 0.278289 | 0.39 | 0.308 | 0.008373 | CC | PAK1 |
| ATAD1 | 4.99E-07 | 0.278141 | 0.617 | 0.629 | 0.009709 | CC | ATAD1 |
| 6-Mar | 5.57E-07 | 0.263454 | 0.564 | 0.548 | 0.010827 | CC | 6-Mar |
| BSDC1 | 5.94E-07 | 0.258757 | 0.508 | 0.468 | 0.011551 | CC | BSDC1 |
| DICER1 | 6.83E-07 | 0.264976 | 0.462 | 0.395 | 0.01329 | CC | DICER1 |

| | | | | | | | |
|---|---|---|---|---|---|---|---|
| TRIM33 | 7.00E-07 | 0.254719 | 0.428 | 0.344 | 0.013626 | CC | TRIM33 |
| MAPK6 | 8.24E-07 | 0.261504 | 0.595 | 0.556 | 0.01603 | CC | MAPK6 |
| SGCB | 1.02E-06 | 0.257714 | 0.356 | 0.273 | 0.019815 | CC | SGCB |
| NECAP1 | 1.31E-06 | 0.266877 | 0.409 | 0.333 | 0.025544 | CC | NECAP1 |
| TMX4 | 1.47E-06 | 0.270437 | 0.458 | 0.397 | 0.028695 | CC | TMX4 |
| CCDC28B | 1.75E-06 | 0.263753 | 0.591 | 0.566 | 0.033951 | CC | CCDC28B |
| ZFHX4 | 2.65E-06 | 0.411938 | 0.508 | 0.461 | 0.051522 | CC | ZFHX4 |
| PEG3 | 3.90E-06 | 0.361922 | 0.568 | 0.543 | 0.075832 | CC | PEG3 |
| USB1 | 3.95E-06 | 0.254678 | 0.432 | 0.383 | 0.076819 | CC | USB1 |
| CCDC88A | 5.04E-06 | 0.258718 | 0.394 | 0.326 | 0.097987 | CC | CCDC88A |
| GNAS | 6.04E-06 | 0.259103 | 0.833 | 0.832 | 0.117406 | CC | GNAS |
| ZNF462 | 1.19E-05 | 0.268324 | 0.405 | 0.338 | 0.232413 | CC | ZNF462 |
| CHD7 | 1.79E-05 | 0.252272 | 0.564 | 0.551 | 0.349158 | CC | CHD7 |
| FKBP1B | 3.27E-05 | 0.250865 | 0.398 | 0.342 | 0.636927 | CC | FKBP1B |
| SCD5 | 3.46E-05 | 0.265124 | 0.477 | 0.432 | 0.673253 | CC | SCD5 |
| NUDT11 | 5.84E-05 | 0.25131 | 0.42 | 0.375 | 1 | CC | NUDT11 |
| DCBLD2 | 7.79E-05 | 0.310423 | 0.322 | 0.253 | 1 | CC | DCBLD2 |
| JUND | 9.80E-05 | 0.251889 | 0.583 | 0.583 | 1 | CC | JUND |
| PRPSAP2 | 0.000105 | 0.258908 | 0.553 | 0.566 | 1 | CC | PRPSAP2 |
| PRPH | 0.000182 | 0.590258 | 0.159 | 0.095 | 1 | CC | PRPH |
| CDKN1C | 0.000188 | 0.278926 | 0.371 | 0.321 | 1 | CC | CDKN1C |
| SPTAN1 | 0.000272 | 0.250258 | 0.318 | 0.263 | 1 | CC | SPTAN1 |
| CNTNAP2 | 0.000404 | 0.383776 | 0.754 | 0.799 | 1 | CC | CNTNAP2 |
| 1-Jun | 0.001506 | 0.258589 | 0.712 | 0.741 | 1 | CC | JUN |
| HIST1H2B | 0.001622 | 0.271286 | 0.277 | 0.218 | 1 | CC | HIST1H2BG |
| VCAN | 0.001638 | 0.306717 | 0.322 | 0.276 | 1 | CC | VCAN |
| ARL4D | 0.00178 | 0.359239 | 0.477 | 0.479 | 1 | CC | ARL4D |
| C1orf61 | 0.005331 | 0.352226 | 0.405 | 0.362 | 1 | CC | C1orf61 |
| LMO3 | 0.007783 | 0.276761 | 0.205 | 0.159 | 1 | CC | LMO3 |
| LINC0155 | 5.24E-138 | 1.279978 | 0.733 | 0.1 | 1.02E-133 | NP1 | LINC01551 |
| FABP7 | 8.99E-109 | 1.821881 | 0.995 | 0.536 | 1.75E-104 | NP1 | FABP7 |
| SFRP1 | 5.06E-99 | 1.437351 | 0.854 | 0.285 | 9.85E-95 | NP1 | SFRP1 |
| FOXG1 | 2.60E-89 | 0.595788 | 0.519 | 0.072 | 5.05E-85 | NP1 | FOXG1 |
| C1orf611 | 2.17E-87 | 1.354853 | 0.869 | 0.318 | 4.23E-83 | NP1 | C1orf61 |
| PTPRZ1 | 4.53E-81 | 0.867545 | 0.801 | 0.254 | 8.80E-77 | NP1 | PTPRZ1 |
| TTYH1 | 6.47E-73 | 0.983069 | 0.985 | 0.802 | 1.26E-68 | NP1 | TTYH1 |
| LHX21 | 4.67E-72 | 0.654701 | 0.612 | 0.128 | 9.09E-68 | NP1 | LHX2 |
| AMBN | 3.68E-71 | 0.598229 | 0.379 | 0.044 | 7.15E-67 | NP1 | AMBN |
| ID4 | 3.49E-69 | 1.061059 | 0.956 | 0.73 | 6.79E-65 | NP1 | ID4 |
| RPS6 | 4.64E-69 | 0.524814 | 1 | 1 | 9.03E-65 | NP1 | RPS6 |

| | | | | | | | |
|---|---|---|---|---|---|---|---|
| RPS151 | 2.31E-67 | 0.445789 | 1 | 1 | 4.50E-63 | NP1 | RPS15 |
| CKB1 | 5.51E-67 | 0.749665 | 1 | 0.982 | 1.07E-62 | NP1 | CKB |
| HNRNPA1 | 4.58E-66 | 0.714952 | 1 | 0.995 | 8.91E-62 | NP1 | HNRNPA1 |
| SOX3 | 2.00E-64 | 0.766601 | 0.883 | 0.418 | 3.89E-60 | NP1 | SOX3 |
| TMSB4X1 | 2.79E-64 | 0.697019 | 1 | 1 | 5.42E-60 | NP1 | TMSB4X |
| VIM | 3.29E-63 | 0.943047 | 1 | 0.953 | 6.40E-59 | NP1 | VIM |
| PLP1 | 1.12E-62 | 0.882664 | 0.631 | 0.174 | 2.18E-58 | NP1 | PLP1 |
| NAP1L1 | 5.75E-60 | 0.557752 | 1 | 0.988 | 1.12E-55 | NP1 | NAP1L1 |
| RPSA | 1.15E-59 | 0.46747 | 1 | 0.999 | 2.23E-55 | NP1 | RPSA |
| RPL19 | 1.14E-57 | 0.370113 | 1 | 1 | 2.23E-53 | NP1 | RPL19 |
| GLI3 | 4.39E-57 | 0.60039 | 0.621 | 0.189 | 8.54E-53 | NP1 | GLI3 |
| VEPH1 | 5.82E-57 | 0.397263 | 0.456 | 0.09 | 1.13E-52 | NP1 | VEPH1 |
| NR2E1 | 7.51E-56 | 0.368815 | 0.388 | 0.065 | 1.46E-51 | NP1 | NR2E1 |
| RPS9 | 1.87E-55 | 0.395257 | 1 | 1 | 3.64E-51 | NP1 | RPS9 |
| CREB5 | 6.89E-55 | 0.645445 | 0.709 | 0.252 | 1.34E-50 | NP1 | CREB5 |
| RPL351 | 8.89E-55 | 0.365411 | 1 | 1 | 1.73E-50 | NP1 | RPL35 |
| LEF1 | 1.91E-53 | 0.498154 | 0.597 | 0.172 | 3.72E-49 | NP1 | LEF1 |
| RPL121 | 2.31E-53 | 0.429028 | 1 | 0.999 | 4.49E-49 | NP1 | RPL12 |
| RPL10A | 7.47E-53 | 0.412952 | 1 | 0.999 | 1.45E-48 | NP1 | RPL10A |
| RPLP01 | 1.01E-52 | 0.447918 | 1 | 1 | 1.96E-48 | NP1 | RPLP0 |
| SFRP2 | 1.89E-52 | 1.106286 | 0.801 | 0.371 | 3.68E-48 | NP1 | SFRP2 |
| RPS3 | 2.16E-52 | 0.404826 | 1 | 1 | 4.20E-48 | NP1 | RPS3 |
| RPS2 | 3.24E-51 | 0.360694 | 1 | 1 | 6.31E-47 | NP1 | RPS2 |
| NTRK2 | 3.36E-51 | 0.75986 | 0.485 | 0.122 | 6.54E-47 | NP1 | NTRK2 |
| LIX1 | 3.94E-49 | 0.603337 | 0.723 | 0.286 | 7.67E-45 | NP1 | LIX1 |
| RPL131 | 1.05E-47 | 0.351331 | 1 | 1 | 2.04E-43 | NP1 | RPL13 |
| FAM181A | 1.65E-47 | 0.473669 | 0.549 | 0.161 | 3.21E-43 | NP1 | FAM181A |
| RPL18A1 | 6.82E-46 | 0.351101 | 1 | 1 | 1.33E-41 | NP1 | RPL18A |
| NES | 1.73E-45 | 0.678241 | 0.893 | 0.558 | 3.37E-41 | NP1 | NES |
| RPL71 | 2.26E-45 | 0.398521 | 1 | 1 | 4.41E-41 | NP1 | RPL7 |
| VCAM1 | 1.24E-44 | 0.346338 | 0.325 | 0.057 | 2.42E-40 | NP1 | VCAM1 |
| RPL321 | 2.74E-44 | 0.333045 | 1 | 1 | 5.34E-40 | NP1 | RPL32 |
| QKI | 7.16E-44 | 0.586856 | 0.728 | 0.329 | 1.39E-39 | NP1 | QKI |
| RPL11 | 1.94E-43 | 0.312605 | 1 | 1 | 3.77E-39 | NP1 | RPL11 |
| GAS1 | 7.25E-43 | 0.546006 | 0.626 | 0.241 | 1.41E-38 | NP1 | GAS1 |
| PAX6 | 1.43E-42 | 0.677499 | 0.922 | 0.551 | 2.78E-38 | NP1 | PAX6 |
| RPL7A | 1.74E-42 | 0.381872 | 1 | 1 | 3.39E-38 | NP1 | RPL7A |
| GNB2L1 | 2.49E-42 | 0.347604 | 1 | 1 | 4.84E-38 | NP1 | GNB2L1 |
| HMGA2 | 3.72E-42 | 0.659641 | 0.723 | 0.36 | 7.24E-38 | NP1 | HMGA2 |
| SLC1A3 | 3.87E-41 | 0.447794 | 0.369 | 0.081 | 7.52E-37 | NP1 | SLC1A3 |

| | | | | | | | |
|---|---|---|---|---|---|---|---|
| RPL141 | 5.32E-41 | 0.33766 | 1 | 0.999 | 1.03E-36 | NP1 | RPL14 |
| RPS241 | 8.90E-41 | 0.45226 | 1 | 0.991 | 1.73E-36 | NP1 | RPS24 |
| PPA1 | 1.36E-40 | 0.614795 | 0.922 | 0.732 | 2.65E-36 | NP1 | PPA1 |
| ACTG11 | 4.18E-40 | 0.399708 | 1 | 1 | 8.14E-36 | NP1 | ACTG1 |
| FABP51 | 4.59E-40 | 0.717717 | 0.82 | 0.478 | 8.93E-36 | NP1 | FABP5 |
| GPM6B | 9.10E-40 | 0.658206 | 0.883 | 0.601 | 1.77E-35 | NP1 | GPM6B |
| RPS231 | 2.65E-39 | 0.325303 | 1 | 1 | 5.15E-35 | NP1 | RPS23 |
| NME1 | 4.59E-39 | 0.598512 | 0.917 | 0.729 | 8.93E-35 | NP1 | NME1 |
| ATP5G2 | 1.24E-38 | 0.319241 | 1 | 1 | 2.42E-34 | NP1 | ATP5G2 |
| RPLP1 | 1.88E-38 | 0.289416 | 1 | 1 | 3.66E-34 | NP1 | RPLP1 |
| DCT | 1.90E-38 | 0.306147 | 0.252 | 0.038 | 3.70E-34 | NP1 | DCT |
| RPL29 | 2.88E-38 | 0.331575 | 1 | 1 | 5.60E-34 | NP1 | RPL29 |
| GRIP2 | 4.46E-38 | 0.318635 | 0.32 | 0.064 | 8.68E-34 | NP1 | GRIP2 |
| RPL51 | 9.97E-38 | 0.326912 | 1 | 0.999 | 1.94E-33 | NP1 | RPL5 |
| LINC0046 | 1.66E-37 | 0.484436 | 0.617 | 0.246 | 3.22E-33 | NP1 | LINC00461 |
| RAN | 1.71E-37 | 0.417186 | 1 | 0.985 | 3.32E-33 | NP1 | RAN |
| SOX21 | 2.37E-37 | 0.353152 | 0.379 | 0.092 | 4.62E-33 | NP1 | SOX21 |
| EFNB2 | 3.45E-37 | 0.43349 | 0.524 | 0.18 | 6.71E-33 | NP1 | EFNB2 |
| RPS71 | 4.95E-37 | 0.301137 | 1 | 1 | 9.64E-33 | NP1 | RPS7 |
| RPS141 | 5.31E-37 | 0.273255 | 1 | 1 | 1.03E-32 | NP1 | RPS14 |
| TFAP2C | 6.22E-37 | 0.263891 | 0.316 | 0.062 | 1.21E-32 | NP1 | TFAP2C |
| RPS281 | 1.34E-36 | 0.295532 | 1 | 0.999 | 2.61E-32 | NP1 | RPS28 |
| RPS121 | 1.48E-36 | 0.303869 | 1 | 1 | 2.87E-32 | NP1 | RPS12 |
| HMGA1 | 6.52E-36 | 0.545455 | 0.976 | 0.906 | 1.27E-31 | NP1 | HMGA1 |
| MT2A | 7.38E-36 | 0.372224 | 0.408 | 0.109 | 1.44E-31 | NP1 | MT2A |
| TMSB15A | 9.65E-36 | 0.560942 | 0.932 | 0.63 | 1.88E-31 | NP1 | TMSB15A |
| YBX1 | 3.02E-35 | 0.355063 | 1 | 0.999 | 5.87E-31 | NP1 | YBX1 |
| RP3-395M | 4.83E-35 | 0.597733 | 0.481 | 0.158 | 9.40E-31 | NP1 | RP3-395M20.12 |
| MCM7 | 1.40E-34 | 0.592121 | 0.743 | 0.37 | 2.72E-30 | NP1 | MCM7 |
| RPL4 | 1.42E-34 | 0.320096 | 1 | 0.999 | 2.77E-30 | NP1 | RPL4 |
| RPL23A | 4.03E-34 | 0.30567 | 1 | 0.998 | 7.84E-30 | NP1 | RPL23A |
| LDHB | 6.05E-34 | 0.365733 | 1 | 0.997 | 1.18E-29 | NP1 | LDHB |
| SOX2 | 1.18E-33 | 0.507968 | 0.942 | 0.677 | 2.30E-29 | NP1 | SOX2 |
| RPS171 | 1.86E-33 | 0.296705 | 1 | 1 | 3.63E-29 | NP1 | RPS17 |
| PARP1 | 3.27E-33 | 0.458722 | 0.942 | 0.712 | 6.36E-29 | NP1 | PARP1 |
| HMGB1 | 4.27E-33 | 0.511796 | 1 | 0.98 | 8.31E-29 | NP1 | HMGB1 |
| SERBP1 | 4.91E-33 | 0.457155 | 0.966 | 0.914 | 9.54E-29 | NP1 | SERBP1 |
| MARCKSL | 4.14E-32 | 0.450077 | 1 | 0.978 | 8.06E-28 | NP1 | MARCKSL1 |
| ACTB1 | 5.59E-32 | 0.382919 | 1 | 1 | 1.09E-27 | NP1 | ACTB |
| HES5 | 5.92E-32 | 0.29948 | 0.33 | 0.079 | 1.15E-27 | NP1 | HES5 |

| | | | | | | | |
|---|---|---|---|---|---|---|---|
| RPS15A1 | 7.50E-32 | 0.2814 | 1 | 1 | 1.46E-27 | NP1 | RPS15A |
| TSPAN5 | 1.09E-31 | 0.46076 | 0.689 | 0.358 | 2.13E-27 | NP1 | TSPAN5 |
| EFNB1 | 1.13E-31 | 0.498573 | 0.762 | 0.431 | 2.20E-27 | NP1 | EFNB1 |
| RPL391 | 2.03E-31 | 0.326302 | 1 | 0.996 | 3.94E-27 | NP1 | RPL39 |
| HES4 | 3.49E-31 | 0.529726 | 0.956 | 0.749 | 6.80E-27 | NP1 | HES4 |
| RPS27A1 | 7.88E-31 | 0.3006 | 1 | 1 | 1.53E-26 | NP1 | RPS27A |
| FGFBP3 | 7.90E-31 | 0.734241 | 0.646 | 0.33 | 1.54E-26 | NP1 | FGFBP3 |
| PPIA | 1.20E-30 | 0.417057 | 0.985 | 0.979 | 2.34E-26 | NP1 | PPIA |
| APEX1 | 3.51E-30 | 0.467075 | 0.908 | 0.766 | 6.83E-26 | NP1 | APEX1 |
| RPL371 | 4.01E-30 | 0.321607 | 1 | 0.998 | 7.81E-26 | NP1 | RPL37 |
| CENPV | 1.75E-29 | 0.648342 | 0.854 | 0.635 | 3.40E-25 | NP1 | CENPV |
| C5orf30 | 2.43E-29 | 0.417999 | 0.549 | 0.237 | 4.73E-25 | NP1 | C5orf30 |
| AIF1L | 2.66E-29 | 0.327317 | 0.461 | 0.165 | 5.17E-25 | NP1 | AIF1L |
| HNRNPA2 | 2.71E-29 | 0.486599 | 0.981 | 0.917 | 5.27E-25 | NP1 | HNRNPA2B1 |
| CDK4 | 6.25E-29 | 0.446241 | 0.903 | 0.722 | 1.22E-24 | NP1 | CDK4 |
| MASP1 | 5.16E-28 | 0.31015 | 0.383 | 0.115 | 1.00E-23 | NP1 | MASP1 |
| RPL341 | 1.87E-27 | 0.266251 | 1 | 1 | 3.64E-23 | NP1 | RPL34 |
| SYNE2 | 2.79E-27 | 0.523203 | 0.718 | 0.432 | 5.44E-23 | NP1 | SYNE2 |
| RPS182 | 2.99E-27 | 0.27471 | 1 | 1 | 5.82E-23 | NP1 | RPS18 |
| CDH2 | 3.28E-27 | 0.480043 | 0.927 | 0.698 | 6.38E-23 | NP1 | CDH2 |
| LIMCH1 | 4.06E-27 | 0.342404 | 0.447 | 0.167 | 7.89E-23 | NP1 | LIMCH1 |
| SNRPD1 | 4.93E-27 | 0.433185 | 0.971 | 0.857 | 9.60E-23 | NP1 | SNRPD1 |
| NME4 | 9.42E-27 | 0.45204 | 0.893 | 0.681 | 1.83E-22 | NP1 | NME4 |
| RPS81 | 1.17E-26 | 0.260861 | 1 | 1 | 2.28E-22 | NP1 | RPS8 |
| NEUROG2 | 1.65E-26 | 0.258118 | 0.243 | 0.05 | 3.21E-22 | NP1 | NEUROG2 |
| VCAN1 | 3.91E-26 | 0.401015 | 0.558 | 0.255 | 7.61E-22 | NP1 | VCAN |
| HNRNPM | 4.97E-26 | 0.514132 | 0.893 | 0.774 | 9.66E-22 | NP1 | HNRNPM |
| RPS101 | 5.02E-26 | 0.277646 | 1 | 0.997 | 9.78E-22 | NP1 | RPS10 |
| RGMA | 1.15E-25 | 0.374238 | 0.617 | 0.314 | 2.24E-21 | NP1 | RGMA |
| GPC1 | 1.15E-25 | 0.448712 | 0.583 | 0.305 | 2.24E-21 | NP1 | GPC1 |
| RBM31 | 1.96E-25 | 0.429727 | 0.917 | 0.771 | 3.82E-21 | NP1 | RBM3 |
| TMEM97 | 6.57E-25 | 0.51314 | 0.791 | 0.519 | 1.28E-20 | NP1 | TMEM97 |
| TMEM161 | 6.80E-25 | 0.478195 | 0.641 | 0.367 | 1.32E-20 | NP1 | TMEM161B-AS1 |
| MOXD1 | 8.87E-25 | 0.294371 | 0.17 | 0.027 | 1.73E-20 | NP1 | MOXD1 |
| NPM1 | 1.02E-24 | 0.277317 | 1 | 0.999 | 1.98E-20 | NP1 | NPM1 |
| SNRPF | 1.45E-24 | 0.366157 | 0.956 | 0.884 | 2.83E-20 | NP1 | SNRPF |
| RPL231 | 2.12E-24 | 0.261636 | 1 | 0.998 | 4.12E-20 | NP1 | RPL23 |
| GNAI2 | 3.48E-24 | 0.423679 | 0.859 | 0.744 | 6.78E-20 | NP1 | GNAI2 |
| RBMX | 4.18E-24 | 0.385622 | 0.966 | 0.873 | 8.14E-20 | NP1 | RBMX |
| SET | 6.59E-24 | 0.353132 | 0.99 | 0.961 | 1.28E-19 | NP1 | SET |

| | | | | | | | |
|---|---|---|---|---|---|---|---|
| EEF1D1 | 6.64E-24 | 0.305649 | 0.995 | 0.981 | 1.29E-19 | NP1 | EEF1D |
| UHRF1 | 8.51E-24 | 0.304439 | 0.442 | 0.165 | 1.66E-19 | NP1 | UHRF1 |
| SOX9 | 3.72E-23 | 0.39855 | 0.553 | 0.28 | 7.24E-19 | NP1 | SOX9 |
| TRIM24 | 6.47E-23 | 0.399586 | 0.568 | 0.293 | 1.26E-18 | NP1 | TRIM24 |
| SMS | 7.21E-23 | 0.457017 | 0.903 | 0.789 | 1.40E-18 | NP1 | SMS |
| C1orf21 | 9.39E-23 | 0.319181 | 0.549 | 0.26 | 1.83E-18 | NP1 | C1orf21 |
| EGR1 | 1.03E-22 | 0.587754 | 0.786 | 0.523 | 2.01E-18 | NP1 | EGR1 |
| PLAGL1 | 1.21E-22 | 0.365453 | 0.51 | 0.235 | 2.35E-18 | NP1 | PLAGL1 |
| RPS3A1 | 1.40E-22 | 0.257945 | 1 | 1 | 2.73E-18 | NP1 | RPS3A |
| RHOBTB3 | 2.62E-22 | 0.381786 | 0.714 | 0.417 | 5.10E-18 | NP1 | RHOBTB3 |
| PSAT1 | 3.44E-22 | 0.40544 | 0.85 | 0.629 | 6.69E-18 | NP1 | PSAT1 |
| ZFP36L1 | 5.86E-22 | 0.432903 | 0.869 | 0.633 | 1.14E-17 | NP1 | ZFP36L1 |
| GINS2 | 5.93E-22 | 0.411661 | 0.563 | 0.28 | 1.15E-17 | NP1 | GINS2 |
| NKAIN41 | 1.56E-21 | 0.375268 | 0.786 | 0.518 | 3.04E-17 | NP1 | NKAIN4 |
| BTF3 | 1.93E-21 | 0.274942 | 0.995 | 0.965 | 3.76E-17 | NP1 | BTF3 |
| CNBP | 2.05E-21 | 0.318743 | 0.947 | 0.926 | 3.99E-17 | NP1 | CNBP |
| CTSC | 3.87E-21 | 0.290845 | 0.422 | 0.173 | 7.52E-17 | NP1 | CTSC |
| SLC25A6 | 4.69E-21 | 0.295061 | 0.995 | 0.986 | 9.11E-17 | NP1 | SLC25A6 |
| NASP | 5.16E-21 | 0.494204 | 0.777 | 0.584 | 1.00E-16 | NP1 | NASP |
| FBL | 5.37E-21 | 0.38617 | 0.879 | 0.716 | 1.05E-16 | NP1 | FBL |
| NOTCH1 | 6.34E-21 | 0.269688 | 0.301 | 0.097 | 1.23E-16 | NP1 | NOTCH1 |
| SRSF1 | 7.57E-21 | 0.39097 | 0.83 | 0.639 | 1.47E-16 | NP1 | SRSF1 |
| RPL36A1 | 7.64E-21 | 0.334996 | 0.985 | 0.945 | 1.49E-16 | NP1 | RPL36A |
| MT1X | 8.40E-21 | 0.251017 | 0.325 | 0.108 | 1.63E-16 | NP1 | MT1X |
| ITGB8 | 1.04E-20 | 0.28611 | 0.345 | 0.124 | 2.02E-16 | NP1 | ITGB8 |
| ANP32A | 1.66E-20 | 0.386909 | 0.859 | 0.707 | 3.22E-16 | NP1 | ANP32A |
| GLTSCR2 | 1.71E-20 | 0.350736 | 0.947 | 0.906 | 3.33E-16 | NP1 | GLTSCR2 |
| HMGN2 | 1.98E-20 | 0.45253 | 0.985 | 0.978 | 3.86E-16 | NP1 | HMGN2 |
| PPP1CC | 4.93E-20 | 0.365286 | 0.908 | 0.778 | 9.59E-16 | NP1 | PPP1CC |
| SRSF3 | 5.54E-20 | 0.350145 | 0.99 | 0.934 | 1.08E-15 | NP1 | SRSF3 |
| CD99 | 6.12E-20 | 0.327535 | 0.932 | 0.791 | 1.19E-15 | NP1 | CD99 |
| SORD | 7.49E-20 | 0.321697 | 0.51 | 0.26 | 1.46E-15 | NP1 | SORD |
| PABPC1 | 7.64E-20 | 0.314204 | 0.985 | 0.962 | 1.49E-15 | NP1 | PABPC1 |
| DEK | 1.06E-19 | 0.379879 | 0.859 | 0.683 | 2.06E-15 | NP1 | DEK |
| ANOS1 | 1.16E-19 | 0.267037 | 0.296 | 0.099 | 2.26E-15 | NP1 | ANOS1 |
| CDCA7L | 1.26E-19 | 0.331665 | 0.563 | 0.297 | 2.46E-15 | NP1 | CDCA7L |
| MYO10 | 2.26E-19 | 0.355467 | 0.65 | 0.4 | 4.40E-15 | NP1 | MYO10 |
| ACTR3B | 2.94E-19 | 0.330152 | 0.447 | 0.213 | 5.73E-15 | NP1 | ACTR3B |
| HNRNPU | 3.33E-19 | 0.350065 | 0.956 | 0.849 | 6.49E-15 | NP1 | HNRNPU |
| HMGN1 | 5.83E-19 | 0.315193 | 0.995 | 0.972 | 1.13E-14 | NP1 | HMGN1 |

| | | | | | | | |
|---|---|---|---|---|---|---|---|
| BOP1 | 6.20E-19 | 0.294111 | 0.451 | 0.212 | 1.21E-14 | NP1 | BOP1 |
| MPST | 6.59E-19 | 0.353337 | 0.801 | 0.616 | 1.28E-14 | NP1 | MPST |
| SNRPB | 7.50E-19 | 0.347095 | 0.937 | 0.851 | 1.46E-14 | NP1 | SNRPB |
| SYNCRIP | 7.69E-19 | 0.343648 | 0.762 | 0.564 | 1.50E-14 | NP1 | SYNCRIP |
| HNRNPK | 9.74E-19 | 0.297928 | 0.995 | 0.979 | 1.89E-14 | NP1 | HNRNPK |
| LSM7 | 1.01E-18 | 0.322564 | 0.966 | 0.938 | 1.96E-14 | NP1 | LSM7 |
| FUS | 1.03E-18 | 0.35274 | 0.971 | 0.901 | 2.00E-14 | NP1 | FUS |
| MARCKS1 | 1.06E-18 | 0.299498 | 0.966 | 0.902 | 2.06E-14 | NP1 | MARCKS |
| 11-Sep | 1.06E-18 | 0.386539 | 0.806 | 0.693 | 2.06E-14 | NP1 | 11-Sep |
| TKT | 1.31E-18 | 0.357311 | 0.942 | 0.86 | 2.54E-14 | NP1 | TKT |
| CARHSP1: | 1.43E-18 | 0.373642 | 0.874 | 0.724 | 2.79E-14 | NP1 | CARHSP1 |
| MRPL11 | 1.67E-18 | 0.351549 | 0.748 | 0.567 | 3.26E-14 | NP1 | MRPL11 |
| PHGDH | 1.92E-18 | 0.384839 | 0.835 | 0.632 | 3.73E-14 | NP1 | PHGDH |
| NKAIN3 | 2.13E-18 | 0.290613 | 0.403 | 0.168 | 4.14E-14 | NP1 | NKAIN3 |
| H2AFY21 | 2.24E-18 | 0.339146 | 0.82 | 0.623 | 4.36E-14 | NP1 | H2AFY2 |
| GCSH | 2.85E-18 | 0.338057 | 0.908 | 0.74 | 5.54E-14 | NP1 | GCSH |
| HNRNPD | 3.72E-18 | 0.348644 | 0.932 | 0.813 | 7.24E-14 | NP1 | HNRNPD |
| FEZ11 | 1.03E-17 | 0.326439 | 0.869 | 0.719 | 2.01E-13 | NP1 | FEZ1 |
| DLL1 | 1.33E-17 | 0.485895 | 0.437 | 0.206 | 2.59E-13 | NP1 | DLL1 |
| CENPH | 1.47E-17 | 0.270529 | 0.558 | 0.283 | 2.86E-13 | NP1 | CENPH |
| WNT7B | 1.57E-17 | 0.55047 | 0.451 | 0.223 | 3.05E-13 | NP1 | WNT7B |
| ACAT2 | 3.50E-17 | 0.54979 | 0.733 | 0.547 | 6.82E-13 | NP1 | ACAT2 |
| RPL22L11 | 3.84E-17 | 0.319291 | 0.757 | 0.547 | 7.47E-13 | NP1 | RPL22L1 |
| SSBP1 | 7.66E-17 | 0.315511 | 0.903 | 0.776 | 1.49E-12 | NP1 | SSBP1 |
| NCL | 8.10E-17 | 0.375395 | 0.947 | 0.892 | 1.58E-12 | NP1 | NCL |
| HNRNPDL | 9.42E-17 | 0.281576 | 0.981 | 0.984 | 1.83E-12 | NP1 | HNRNPDL |
| GPC4 | 1.15E-16 | 0.31324 | 0.621 | 0.378 | 2.23E-12 | NP1 | GPC4 |
| BTG11 | 2.73E-16 | 0.311624 | 0.947 | 0.875 | 5.31E-12 | NP1 | BTG1 |
| PUF60 | 2.82E-16 | 0.288481 | 0.932 | 0.84 | 5.49E-12 | NP1 | PUF60 |
| HMGCS11 | 2.88E-16 | 0.626122 | 0.913 | 0.806 | 5.60E-12 | NP1 | HMGCS1 |
| HNRNPA0 | 3.32E-16 | 0.277855 | 0.981 | 0.957 | 6.45E-12 | NP1 | HNRNPA0 |
| SRI | 4.61E-16 | 0.343582 | 0.903 | 0.816 | 8.97E-12 | NP1 | SRI |
| BAG1 | 5.03E-16 | 0.307708 | 0.738 | 0.533 | 9.79E-12 | NP1 | BAG1 |
| NONO | 5.17E-16 | 0.293395 | 0.942 | 0.891 | 1.01E-11 | NP1 | NONO |
| IVNS1ABP | 6.75E-16 | 0.305823 | 0.626 | 0.398 | 1.31E-11 | NP1 | IVNS1ABP |
| RAB8B | 2.20E-15 | 0.259098 | 0.403 | 0.192 | 4.27E-11 | NP1 | RAB8B |
| DOK5 | 2.26E-15 | 0.387809 | 0.51 | 0.295 | 4.40E-11 | NP1 | DOK5 |
| PTRHD1 | 2.31E-15 | 0.305241 | 0.718 | 0.523 | 4.49E-11 | NP1 | PTRHD1 |
| TLE11 | 3.25E-15 | 0.318085 | 0.68 | 0.475 | 6.32E-11 | NP1 | TLE1 |
| CLNS1A | 4.66E-15 | 0.266192 | 0.917 | 0.838 | 9.07E-11 | NP1 | CLNS1A |

| | | | | | | | |
|---|---|---|---|---|---|---|---|
| PTBP2 | 6.51E-15 | 0.274584 | 0.524 | 0.298 | 1.27E-10 | NP1 | PTBP2 |
| NR2F1 | 1.04E-14 | 0.353745 | 0.85 | 0.651 | 2.02E-10 | NP1 | NR2F1 |
| BAZ1A | 1.11E-14 | 0.280006 | 0.529 | 0.307 | 2.16E-10 | NP1 | BAZ1A |
| HNRNPA3 | 1.14E-14 | 0.311234 | 0.913 | 0.842 | 2.22E-10 | NP1 | HNRNPA3 |
| PAICS | 1.51E-14 | 0.336838 | 0.83 | 0.675 | 2.94E-10 | NP1 | PAICS |
| ODC1 | 1.85E-14 | 0.27445 | 0.981 | 0.93 | 3.61E-10 | NP1 | ODC1 |
| SNRPE | 1.97E-14 | 0.276615 | 0.981 | 0.915 | 3.84E-10 | NP1 | SNRPE |
| TMEM123 | 2.08E-14 | 0.285299 | 0.85 | 0.68 | 4.05E-10 | NP1 | TMEM123 |
| POLD2 | 9.67E-14 | 0.281325 | 0.777 | 0.631 | 1.88E-09 | NP1 | POLD2 |
| HEY1 | 1.08E-13 | 0.411279 | 0.466 | 0.257 | 2.11E-09 | NP1 | HEY1 |
| PAK11 | 3.33E-13 | 0.254655 | 0.505 | 0.299 | 6.47E-09 | NP1 | PAK1 |
| DBI | 4.84E-13 | 0.31739 | 0.859 | 0.721 | 9.41E-09 | NP1 | DBI |
| PDCD5 | 5.32E-13 | 0.293612 | 0.854 | 0.766 | 1.03E-08 | NP1 | PDCD5 |
| CELF21 | 5.44E-13 | 0.257494 | 0.631 | 0.422 | 1.06E-08 | NP1 | CELF2 |
| SSRP1 | 6.60E-13 | 0.297166 | 0.83 | 0.716 | 1.28E-08 | NP1 | SSRP1 |
| SH3BGRL | 7.08E-13 | 0.286218 | 0.893 | 0.761 | 1.38E-08 | NP1 | SH3BGRL |
| PDLIM3 | 9.15E-13 | 0.298528 | 0.354 | 0.173 | 1.78E-08 | NP1 | PDLIM3 |
| RNF130 | 1.00E-12 | 0.294081 | 0.694 | 0.518 | 1.95E-08 | NP1 | RNF130 |
| LRRN1 | 1.34E-12 | 0.289141 | 0.592 | 0.378 | 2.60E-08 | NP1 | LRRN1 |
| AKR1B1 | 1.52E-12 | 0.28302 | 0.786 | 0.634 | 2.95E-08 | NP1 | AKR1B1 |
| ILF3 | 1.73E-12 | 0.275835 | 0.85 | 0.759 | 3.37E-08 | NP1 | ILF3 |
| FDFT1 | 1.85E-12 | 0.362686 | 0.913 | 0.838 | 3.59E-08 | NP1 | FDFT1 |
| F2R | 1.92E-12 | 0.263872 | 0.466 | 0.273 | 3.73E-08 | NP1 | F2R |
| SMARCC1 | 2.22E-12 | 0.316359 | 0.68 | 0.554 | 4.32E-08 | NP1 | SMARCC1 |
| PGM1 | 2.35E-12 | 0.26568 | 0.612 | 0.416 | 4.56E-08 | NP1 | PGM1 |
| NEK6 | 3.17E-12 | 0.25173 | 0.393 | 0.215 | 6.16E-08 | NP1 | NEK6 |
| C8orf4 | 5.55E-12 | 0.291761 | 0.379 | 0.196 | 1.08E-07 | NP1 | C8orf4 |
| NAE1 | 5.66E-12 | 0.316027 | 0.699 | 0.561 | 1.10E-07 | NP1 | NAE1 |
| ATXN10 | 5.97E-12 | 0.282599 | 0.908 | 0.846 | 1.16E-07 | NP1 | ATXN10 |
| FZD2 | 7.35E-12 | 0.281757 | 0.583 | 0.42 | 1.43E-07 | NP1 | FZD2 |
| CMTM3 | 9.93E-12 | 0.252614 | 0.466 | 0.281 | 1.93E-07 | NP1 | CMTM3 |
| HMGB3 | 1.45E-11 | 0.265698 | 0.738 | 0.56 | 2.82E-07 | NP1 | HMGB3 |
| C17orf89 | 1.52E-11 | 0.256026 | 0.782 | 0.63 | 2.95E-07 | NP1 | C17orf89 |
| UBE2N | 2.95E-11 | 0.281006 | 0.845 | 0.75 | 5.73E-07 | NP1 | UBE2N |
| HNRNPA1 | 3.47E-11 | 0.2571 | 0.403 | 0.236 | 6.75E-07 | NP1 | HNRNPA1L2 |
| DHX9 | 3.52E-11 | 0.263665 | 0.612 | 0.438 | 6.86E-07 | NP1 | DHX9 |
| EMX2 | 3.64E-11 | 0.294307 | 0.767 | 0.608 | 7.08E-07 | NP1 | EMX2 |
| SRSF2 | 4.07E-11 | 0.281002 | 0.937 | 0.882 | 7.92E-07 | NP1 | SRSF2 |
| CHD71 | 4.98E-11 | 0.284994 | 0.699 | 0.538 | 9.69E-07 | NP1 | CHD7 |
| CCNB1IP1 | 5.28E-11 | 0.269171 | 0.801 | 0.672 | 1.03E-06 | NP1 | CCNB1IP1 |

| | | | | | | | |
|---|---|---|---|---|---|---|---|
| ATP1B3 | 6.42E-11 | 0.268055 | 0.922 | 0.83 | 1.25E-06 | NP1 | ATP1B3 |
| LSM6 | 8.99E-11 | 0.260998 | 0.733 | 0.58 | 1.75E-06 | NP1 | LSM6 |
| NDN | 1.10E-10 | 0.293637 | 0.859 | 0.783 | 2.14E-06 | NP1 | NDN |
| FBLN1 | 1.23E-10 | 0.329539 | 0.529 | 0.347 | 2.39E-06 | NP1 | FBLN1 |
| RBBP4 | 1.44E-10 | 0.253492 | 0.879 | 0.787 | 2.80E-06 | NP1 | RBBP4 |
| SRSF7 | 1.75E-10 | 0.264364 | 0.908 | 0.824 | 3.40E-06 | NP1 | SRSF7 |
| RSL1D1 | 1.77E-10 | 0.26366 | 0.854 | 0.752 | 3.45E-06 | NP1 | RSL1D1 |
| IGDCC31 | 1.87E-10 | 0.256756 | 0.587 | 0.407 | 3.63E-06 | NP1 | IGDCC3 |
| ILF2 | 3.95E-10 | 0.27062 | 0.888 | 0.847 | 7.68E-06 | NP1 | ILF2 |
| SFPQ | 9.42E-10 | 0.275673 | 0.791 | 0.704 | 1.83E-05 | NP1 | SFPQ |
| ANP32B | 9.81E-10 | 0.272017 | 0.723 | 0.605 | 1.91E-05 | NP1 | ANP32B |
| RAC3 | 1.52E-09 | 0.281417 | 0.573 | 0.432 | 2.96E-05 | NP1 | RAC3 |
| CCT5 | 1.65E-09 | 0.251426 | 0.82 | 0.72 | 3.21E-05 | NP1 | CCT5 |
| MSI1 | 2.14E-09 | 0.273209 | 0.568 | 0.395 | 4.16E-05 | NP1 | MSI1 |
| CD320 | 7.89E-09 | 0.254791 | 0.573 | 0.428 | 0.000153 | NP1 | CD320 |
| TCF4 | 8.04E-09 | 0.26506 | 0.553 | 0.413 | 0.000156 | NP1 | TCF4 |
| MCM3 | 2.05E-08 | 0.266498 | 0.466 | 0.314 | 0.0004 | NP1 | MCM3 |
| CHN11 | 6.26E-08 | 0.277166 | 0.476 | 0.327 | 0.001218 | NP1 | CHN1 |
| FAM107A | 4.51E-07 | 0.251046 | 0.18 | 0.082 | 0.008777 | NP1 | FAM107A |
| HNRNPH1 | 6.60E-07 | 0.310766 | 0.738 | 0.667 | 0.012834 | NP1 | HNRNPH1 |
| SLF1 | 2.71E-06 | 0.250102 | 0.374 | 0.245 | 0.052801 | NP1 | SLF1 |
| FDPS | 1.69E-05 | 0.255481 | 0.888 | 0.851 | 0.329366 | NP1 | FDPS |
| CCNA2 | 3.53E-193 | 1.143856 | 0.711 | 0.061 | 6.86E-189 | NP2 | CCNA2 |
| MKI67 | 1.84E-179 | 0.895598 | 0.624 | 0.044 | 3.59E-175 | NP2 | MKI67 |
| PBK | 7.20E-176 | 0.922936 | 0.715 | 0.073 | 1.40E-171 | NP2 | PBK |
| AURKB | 2.24E-174 | 0.994832 | 0.653 | 0.057 | 4.35E-170 | NP2 | AURKB |
| SPC25 | 2.63E-173 | 0.792144 | 0.636 | 0.051 | 5.13E-169 | NP2 | SPC25 |
| KIF2C | 2.13E-166 | 0.645958 | 0.583 | 0.041 | 4.14E-162 | NP2 | KIF2C |
| NUSAP1 | 4.59E-163 | 1.23878 | 0.789 | 0.12 | 8.94E-159 | NP2 | NUSAP1 |
| RRM2 | 2.98E-161 | 0.821039 | 0.69 | 0.073 | 5.80E-157 | NP2 | RRM2 |
| TOP2A | 3.33E-158 | 1.526104 | 0.748 | 0.107 | 6.47E-154 | NP2 | TOP2A |
| BIRC5 | 2.87E-157 | 1.334027 | 0.814 | 0.136 | 5.58E-153 | NP2 | BIRC5 |
| KIAA0101 | 2.21E-154 | 1.515721 | 0.926 | 0.215 | 4.30E-150 | NP2 | KIAA0101 |
| NUF2 | 1.61E-150 | 0.799226 | 0.645 | 0.07 | 3.14E-146 | NP2 | NUF2 |
| CENPF | 2.87E-146 | 1.247267 | 0.744 | 0.118 | 5.58E-142 | NP2 | CENPF |
| NDC80 | 7.77E-140 | 0.480182 | 0.504 | 0.036 | 1.51E-135 | NP2 | NDC80 |
| CCNB2 | 3.51E-137 | 1.189518 | 0.736 | 0.12 | 6.82E-133 | NP2 | CCNB2 |
| GTSE1 | 1.53E-134 | 0.728156 | 0.566 | 0.058 | 2.97E-130 | NP2 | GTSE1 |
| FAM64A | 7.14E-134 | 0.93434 | 0.64 | 0.086 | 1.39E-129 | NP2 | FAM64A |
| ZWINT | 3.90E-131 | 0.747044 | 0.76 | 0.134 | 7.59E-127 | NP2 | ZWINT |

| | | | | | | | |
|---|---|---|---|---|---|---|---|
| KIFC1 | 7.95E-126 | 0.610018 | 0.545 | 0.058 | 1.55E-121 | NP2 | KIFC1 |
| PLK1 | 3.35E-123 | 1.093915 | 0.562 | 0.066 | 6.51E-119 | NP2 | PLK1 |
| DEPDC1B | 4.16E-123 | 0.56388 | 0.57 | 0.068 | 8.10E-119 | NP2 | DEPDC1B |
| PRC1 | 4.41E-123 | 0.822714 | 0.657 | 0.107 | 8.59E-119 | NP2 | PRC1 |
| CDCA8 | 1.00E-122 | 0.671621 | 0.545 | 0.06 | 1.95E-118 | NP2 | CDCA8 |
| UBE2C | 2.38E-122 | 1.849412 | 0.802 | 0.199 | 4.63E-118 | NP2 | UBE2C |
| TUBA1B | 6.68E-119 | 1.391845 | 1 | 0.985 | 1.30E-114 | NP2 | TUBA1B |
| HJURP | 2.92E-116 | 0.543359 | 0.479 | 0.044 | 5.68E-112 | NP2 | HJURP |
| UBE2T | 3.86E-114 | 0.868433 | 0.822 | 0.221 | 7.51E-110 | NP2 | UBE2T |
| ASPM | 6.02E-114 | 0.488152 | 0.434 | 0.034 | 1.17E-109 | NP2 | ASPM |
| TPX2 | 2.40E-112 | 0.768904 | 0.62 | 0.098 | 4.67E-108 | NP2 | TPX2 |
| CENPA | 5.32E-112 | 0.537957 | 0.438 | 0.037 | 1.03E-107 | NP2 | CENPA |
| CDK1 | 3.07E-110 | 1.17517 | 0.769 | 0.184 | 5.97E-106 | NP2 | CDK1 |
| DLGAP5 | 1.21E-109 | 0.53063 | 0.467 | 0.045 | 2.34E-105 | NP2 | DLGAP5 |
| CENPU | 2.79E-108 | 0.47499 | 0.632 | 0.099 | 5.42E-104 | NP2 | CENPU |
| FOXM1 | 2.58E-107 | 0.358306 | 0.434 | 0.038 | 5.01E-103 | NP2 | FOXM1 |
| HMGB11 | 5.17E-107 | 1.124494 | 1 | 0.98 | 1.01E-102 | NP2 | HMGB1 |
| HMGN21 | 7.66E-107 | 1.192224 | 1 | 0.976 | 1.49E-102 | NP2 | HMGN2 |
| MELK | 9.27E-107 | 0.45242 | 0.537 | 0.067 | 1.80E-102 | NP2 | MELK |
| CDCA3 | 9.43E-107 | 0.566099 | 0.463 | 0.047 | 1.83E-102 | NP2 | CDCA3 |
| SMC4 | 6.19E-106 | 0.983638 | 0.802 | 0.236 | 1.20E-101 | NP2 | SMC4 |
| CDCA5 | 6.52E-106 | 0.50513 | 0.562 | 0.081 | 1.27E-101 | NP2 | CDCA5 |
| OIP5 | 1.23E-105 | 0.401472 | 0.479 | 0.051 | 2.39E-101 | NP2 | OIP5 |
| H2AFZ | 3.43E-105 | 1.209421 | 1 | 0.979 | 6.67E-101 | NP2 | H2AFZ |
| CDC20 | 7.62E-104 | 1.172138 | 0.665 | 0.134 | 1.48E-99 | NP2 | CDC20 |
| TK1 | 1.58E-103 | 0.420309 | 0.517 | 0.061 | 3.07E-99 | NP2 | TK1 |
| CENPH1 | 1.46E-102 | 0.851183 | 0.814 | 0.249 | 2.83E-98 | NP2 | CENPH |
| TACC3 | 1.69E-102 | 0.547172 | 0.521 | 0.068 | 3.28E-98 | NP2 | TACC3 |
| RTKN2 | 3.04E-102 | 0.421393 | 0.517 | 0.066 | 5.91E-98 | NP2 | RTKN2 |
| CENPM | 3.15E-102 | 0.551889 | 0.624 | 0.107 | 6.13E-98 | NP2 | CENPM |
| MYBL2 | 1.38E-101 | 0.450321 | 0.479 | 0.054 | 2.69E-97 | NP2 | MYBL2 |
| CDC25C | 1.97E-100 | 0.329695 | 0.364 | 0.025 | 3.83E-96 | NP2 | CDC25C |
| CENPN | 2.66E-99 | 0.473683 | 0.595 | 0.099 | 5.17E-95 | NP2 | CENPN |
| KIF20A | 4.73E-99 | 0.322738 | 0.322 | 0.017 | 9.20E-95 | NP2 | KIF20A |
| SPC24 | 5.79E-98 | 0.32597 | 0.417 | 0.039 | 1.13E-93 | NP2 | SPC24 |
| MCM71 | 3.35E-96 | 1.008856 | 0.884 | 0.347 | 6.52E-92 | NP2 | MCM7 |
| MAD2L1 | 4.20E-94 | 1.063607 | 0.847 | 0.335 | 8.18E-90 | NP2 | MAD2L1 |
| CKS1B | 4.56E-94 | 1.185043 | 0.926 | 0.507 | 8.88E-90 | NP2 | CKS1B |
| HMGB2 | 7.78E-94 | 1.407864 | 0.996 | 0.875 | 1.51E-89 | NP2 | HMGB2 |
| ATAD2 | 8.11E-94 | 0.408812 | 0.517 | 0.073 | 1.58E-89 | NP2 | ATAD2 |

| | | | | | | | |
|---|---|---|---|---|---|---|---|
| DEK1 | 1.52E-93 | 0.94045 | 0.963 | 0.669 | 2.95E-89 | NP2 | DEK |
| SPAG5 | 3.80E-93 | 0.334643 | 0.388 | 0.035 | 7.40E-89 | NP2 | SPAG5 |
| PRIM1 | 1.68E-92 | 0.463018 | 0.574 | 0.095 | 3.27E-88 | NP2 | PRIM1 |
| KIF15 | 1.00E-91 | 0.256233 | 0.355 | 0.027 | 1.94E-87 | NP2 | KIF15 |
| GMNN | 1.12E-90 | 0.642024 | 0.736 | 0.206 | 2.18E-86 | NP2 | GMNN |
| UHRF11 | 1.39E-90 | 0.584966 | 0.653 | 0.136 | 2.70E-86 | NP2 | UHRF1 |
| HNRNPA1 | 3.31E-90 | 0.715329 | 1 | 0.995 | 6.43E-86 | NP2 | HNRNPA1 |
| NCAPG | 2.33E-89 | 0.39141 | 0.417 | 0.045 | 4.54E-85 | NP2 | NCAPG |
| NASP1 | 3.77E-89 | 0.880481 | 0.963 | 0.56 | 7.33E-85 | NP2 | NASP |
| CENPE | 2.25E-88 | 0.417856 | 0.413 | 0.045 | 4.37E-84 | NP2 | CENPE |
| RAN1 | 1.55E-87 | 0.68423 | 1 | 0.984 | 3.01E-83 | NP2 | RAN |
| TROAP | 5.83E-87 | 0.436607 | 0.397 | 0.042 | 1.14E-82 | NP2 | TROAP |
| SGOL1 | 4.13E-86 | 0.456787 | 0.459 | 0.062 | 8.04E-82 | NP2 | SGOL1 |
| KIF11 | 6.06E-86 | 0.310626 | 0.384 | 0.038 | 1.18E-81 | NP2 | KIF11 |
| MND1 | 6.84E-86 | 0.295862 | 0.376 | 0.035 | 1.33E-81 | NP2 | MND1 |
| HNRNPA2 | 6.86E-85 | 0.801316 | 0.992 | 0.914 | 1.33E-80 | NP2 | HNRNPA2B1 |
| SMC2 | 7.26E-85 | 0.577639 | 0.698 | 0.18 | 1.41E-80 | NP2 | SMC2 |
| HMGA11 | 9.74E-84 | 0.856656 | 0.996 | 0.902 | 1.90E-79 | NP2 | HMGA1 |
| MCM10 | 1.65E-83 | 0.297735 | 0.393 | 0.042 | 3.21E-79 | NP2 | MCM10 |
| CENPK | 4.53E-83 | 0.519741 | 0.636 | 0.139 | 8.81E-79 | NP2 | CENPK |
| FBXO5 | 1.14E-82 | 0.499727 | 0.574 | 0.113 | 2.22E-78 | NP2 | FBXO5 |
| PPIA1 | 2.66E-82 | 0.681968 | 0.992 | 0.978 | 5.17E-78 | NP2 | PPIA |
| KIF4A | 4.34E-82 | 0.335847 | 0.38 | 0.04 | 8.45E-78 | NP2 | KIF4A |
| CENPV1 | 1.13E-81 | 0.961462 | 0.95 | 0.62 | 2.19E-77 | NP2 | CENPV |
| MIS18BP1 | 1.83E-81 | 0.448875 | 0.496 | 0.081 | 3.56E-77 | NP2 | MIS18BP1 |
| CDCA2 | 2.42E-81 | 0.313969 | 0.318 | 0.025 | 4.71E-77 | NP2 | CDCA2 |
| SKA3 | 3.62E-81 | 0.29697 | 0.38 | 0.04 | 7.05E-77 | NP2 | SKA3 |
| ORC6 | 9.32E-81 | 0.608757 | 0.785 | 0.256 | 1.81E-76 | NP2 | ORC6 |
| TUBB1 | 2.68E-80 | 0.750468 | 1 | 0.99 | 5.22E-76 | NP2 | TUBB |
| CEP55 | 4.71E-80 | 0.258493 | 0.285 | 0.018 | 9.16E-76 | NP2 | CEP55 |
| CCNB1 | 1.64E-79 | 1.254958 | 0.632 | 0.167 | 3.18E-75 | NP2 | CCNB1 |
| KIAA1524 | 7.77E-79 | 0.309112 | 0.405 | 0.05 | 1.51E-74 | NP2 | KIAA1524 |
| PSRC1 | 7.88E-79 | 0.633179 | 0.5 | 0.089 | 1.53E-74 | NP2 | PSRC1 |
| TMSB15A: | 3.57E-78 | 0.896935 | 0.979 | 0.619 | 6.94E-74 | NP2 | TMSB15A |
| FANCI | 2.20E-77 | 0.311021 | 0.417 | 0.055 | 4.27E-73 | NP2 | FANCI |
| GINS21 | 2.27E-77 | 0.678806 | 0.773 | 0.251 | 4.41E-73 | NP2 | GINS2 |
| CNPY1 | 2.54E-77 | 0.617825 | 0.347 | 0.036 | 4.94E-73 | NP2 | CNPY1 |
| TYMS | 4.85E-77 | 0.982363 | 0.86 | 0.436 | 9.43E-73 | NP2 | TYMS |
| CDKN3 | 1.13E-75 | 0.565248 | 0.541 | 0.113 | 2.20E-71 | NP2 | CDKN3 |
| PHF19 | 1.39E-75 | 0.347048 | 0.434 | 0.064 | 2.70E-71 | NP2 | PHF19 |

| | | | | | | | |
|---|---|---|---|---|---|---|---|
| FAM83D | 5.80E-75 | 0.497261 | 0.397 | 0.054 | 1.13E-70 | NP2 | FAM83D |
| LMNB1 | 7.86E-75 | 0.713119 | 0.814 | 0.323 | 1.53E-70 | NP2 | LMNB1 |
| NEK2 | 3.17E-74 | 0.445763 | 0.45 | 0.068 | 6.17E-70 | NP2 | NEK2 |
| MXD3 | 2.46E-73 | 0.459001 | 0.43 | 0.065 | 4.79E-69 | NP2 | MXD3 |
| CENPW | 2.88E-73 | 0.613704 | 0.74 | 0.233 | 5.59E-69 | NP2 | CENPW |
| BUB1 | 3.98E-73 | 0.27667 | 0.335 | 0.034 | 7.75E-69 | NP2 | BUB1 |
| ASF1B | 1.54E-72 | 0.345963 | 0.45 | 0.07 | 3.00E-68 | NP2 | ASF1B |
| LINC0122 | 4.14E-72 | 0.316005 | 0.43 | 0.064 | 8.05E-68 | NP2 | LINC01224 |
| RPS61 | 1.84E-71 | 0.478407 | 1 | 1 | 3.57E-67 | NP2 | RPS6 |
| TTK | 2.72E-71 | 0.314464 | 0.343 | 0.039 | 5.29E-67 | NP2 | TTK |
| RAD51AP | 7.02E-71 | 0.356896 | 0.421 | 0.064 | 1.37E-66 | NP2 | RAD51AP1 |
| RNASEH2 | 1.19E-70 | 0.598753 | 0.818 | 0.328 | 2.32E-66 | NP2 | RNASEH2A |
| PTTG1 | 1.29E-70 | 1.348141 | 0.876 | 0.512 | 2.51E-66 | NP2 | PTTG1 |
| CKAP2 | 2.24E-70 | 0.663187 | 0.653 | 0.199 | 4.35E-66 | NP2 | CKAP2 |
| RANBP1 | 1.63E-69 | 0.692573 | 0.992 | 0.94 | 3.16E-65 | NP2 | RANBP1 |
| H2AFX | 1.86E-69 | 0.980785 | 0.789 | 0.382 | 3.62E-65 | NP2 | H2AFX |
| KIF23 | 4.36E-68 | 0.392149 | 0.438 | 0.075 | 8.49E-64 | NP2 | KIF23 |
| SNRPB1 | 7.01E-68 | 0.663286 | 0.983 | 0.845 | 1.36E-63 | NP2 | SNRPB |
| RRM1 | 5.69E-67 | 0.679771 | 0.822 | 0.368 | 1.11E-62 | NP2 | RRM1 |
| HMGB31 | 6.19E-67 | 0.767213 | 0.913 | 0.536 | 1.20E-62 | NP2 | HMGB3 |
| HIST1H4C | 1.72E-66 | 2.070762 | 0.748 | 0.338 | 3.35E-62 | NP2 | HIST1H4C |
| CKB2 | 2.83E-66 | 0.696272 | 1 | 0.981 | 5.50E-62 | NP2 | CKB |
| PTMA1 | 7.86E-66 | 0.465519 | 1 | 1 | 1.53E-61 | NP2 | PTMA |
| RMI2 | 1.52E-64 | 0.349028 | 0.413 | 0.068 | 2.96E-60 | NP2 | RMI2 |
| CLSPN | 7.96E-64 | 0.381112 | 0.492 | 0.1 | 1.55E-59 | NP2 | CLSPN |
| PIF1 | 1.11E-63 | 0.405964 | 0.24 | 0.017 | 2.16E-59 | NP2 | PIF1 |
| NAP1L11 | 1.18E-63 | 0.495891 | 1 | 0.988 | 2.29E-59 | NP2 | NAP1L1 |
| FABP52 | 4.82E-63 | 0.659843 | 0.905 | 0.463 | 9.38E-59 | NP2 | FABP5 |
| HNRNPD1 | 6.93E-63 | 0.636481 | 0.979 | 0.805 | 1.35E-58 | NP2 | HNRNPD |
| POC1A | 7.28E-63 | 0.321053 | 0.45 | 0.085 | 1.42E-58 | NP2 | POC1A |
| PRR11 | 2.05E-62 | 0.252049 | 0.322 | 0.039 | 4.00E-58 | NP2 | PRR11 |
| RPS22 | 3.32E-62 | 0.389869 | 1 | 1 | 6.46E-58 | NP2 | RPS2 |
| NUCKS1 | 5.50E-62 | 0.66593 | 0.983 | 0.926 | 1.07E-57 | NP2 | NUCKS1 |
| DHFR | 1.66E-61 | 0.600214 | 0.665 | 0.22 | 3.22E-57 | NP2 | DHFR |
| AIF1L1 | 1.88E-61 | 0.374137 | 0.579 | 0.147 | 3.66E-57 | NP2 | AIF1L |
| ECT2 | 1.90E-61 | 0.343759 | 0.438 | 0.083 | 3.69E-57 | NP2 | ECT2 |
| HNRNPA3 | 2.05E-61 | 0.584391 | 0.975 | 0.833 | 3.98E-57 | NP2 | HNRNPA3 |
| PCNA | 4.19E-61 | 0.752521 | 0.847 | 0.444 | 8.16E-57 | NP2 | PCNA |
| DTL | 9.77E-61 | 0.300745 | 0.372 | 0.058 | 1.90E-56 | NP2 | DTL |
| CDC45 | 1.72E-60 | 0.269032 | 0.36 | 0.052 | 3.35E-56 | NP2 | CDC45 |

| | | | | | | | |
|---|---|---|---|---|---|---|---|
| RPSA1 | 5.11E-60 | 0.42666 | 1 | 0.999 | 9.94E-56 | NP2 | RPSA |
| VRK1 | 5.14E-60 | 0.550436 | 0.736 | 0.279 | 1.00E-55 | NP2 | VRK1 |
| KIF20B | 1.02E-59 | 0.525054 | 0.599 | 0.17 | 1.98E-55 | NP2 | KIF20B |
| FANCD2 | 1.59E-59 | 0.2691 | 0.355 | 0.052 | 3.10E-55 | NP2 | FANCD2 |
| STMN11 | 3.92E-59 | 0.687649 | 1 | 0.977 | 7.62E-55 | NP2 | STMN1 |
| SNRPD11 | 3.98E-59 | 0.60621 | 0.975 | 0.855 | 7.75E-55 | NP2 | SNRPD1 |
| CDCA7 | 1.36E-58 | 0.318214 | 0.355 | 0.054 | 2.65E-54 | NP2 | CDCA7 |
| KPNA2 | 2.25E-58 | 1.440837 | 0.921 | 0.666 | 4.37E-54 | NP2 | KPNA2 |
| ANP32B1 | 2.29E-58 | 0.617457 | 0.893 | 0.583 | 4.46E-54 | NP2 | ANP32B |
| NES1 | 1.05E-57 | 0.73454 | 0.917 | 0.549 | 2.05E-53 | NP2 | NES |
| PARPBP | 2.14E-57 | 0.284711 | 0.368 | 0.059 | 4.17E-53 | NP2 | PARPBP |
| FANCA | 4.22E-57 | 0.283598 | 0.405 | 0.074 | 8.22E-53 | NP2 | FANCA |
| C19orf48 | 6.43E-57 | 0.535249 | 0.818 | 0.376 | 1.25E-52 | NP2 | C19orf48 |
| H2AFV | 6.89E-57 | 0.577044 | 0.95 | 0.764 | 1.34E-52 | NP2 | H2AFV |
| SRSF31 | 9.90E-57 | 0.546047 | 0.988 | 0.933 | 1.93E-52 | NP2 | SRSF3 |
| MCM2 | 1.21E-56 | 0.423249 | 0.558 | 0.149 | 2.35E-52 | NP2 | MCM2 |
| HELLS | 1.48E-56 | 0.501679 | 0.603 | 0.184 | 2.88E-52 | NP2 | HELLS |
| NKAIN31 | 6.58E-56 | 0.368428 | 0.566 | 0.145 | 1.28E-51 | NP2 | NKAIN3 |
| SGOL2 | 7.48E-56 | 0.393522 | 0.376 | 0.066 | 1.46E-51 | NP2 | SGOL2 |
| GINS4 | 1.25E-55 | 0.264414 | 0.384 | 0.068 | 2.42E-51 | NP2 | GINS4 |
| HNRNPDL | 6.58E-55 | 0.462838 | 1 | 0.982 | 1.28E-50 | NP2 | HNRNPDL |
| TRIP13 | 7.15E-55 | 0.298341 | 0.426 | 0.087 | 1.39E-50 | NP2 | TRIP13 |
| VIM1 | 1.03E-54 | 0.633219 | 0.996 | 0.953 | 2.00E-50 | NP2 | VIM |
| CHAF1A | 1.79E-54 | 0.337749 | 0.475 | 0.11 | 3.47E-50 | NP2 | CHAF1A |
| CDC25B | 2.42E-54 | 0.443172 | 0.479 | 0.116 | 4.71E-50 | NP2 | CDC25B |
| CDT1 | 7.48E-54 | 0.397107 | 0.525 | 0.139 | 1.45E-49 | NP2 | CDT1 |
| NMU | 9.41E-54 | 0.492818 | 0.698 | 0.252 | 1.83E-49 | NP2 | NMU |
| LRR1 | 1.20E-53 | 0.296324 | 0.442 | 0.096 | 2.34E-49 | NP2 | LRR1 |
| ACTB2 | 1.69E-53 | 0.463681 | 1 | 1 | 3.28E-49 | NP2 | ACTB |
| USP1 | 1.83E-53 | 0.518771 | 0.736 | 0.311 | 3.56E-49 | NP2 | USP1 |
| TMPO | 7.09E-53 | 0.571163 | 0.764 | 0.365 | 1.38E-48 | NP2 | TMPO |
| RPS91 | 1.75E-52 | 0.326678 | 1 | 1 | 3.41E-48 | NP2 | RPS9 |
| SUMO21 | 3.00E-52 | 0.412261 | 1 | 0.994 | 5.83E-48 | NP2 | SUMO2 |
| CDCA4 | 1.77E-51 | 0.455766 | 0.616 | 0.211 | 3.45E-47 | NP2 | CDCA4 |
| SRSF21 | 3.58E-51 | 0.563398 | 0.979 | 0.876 | 6.96E-47 | NP2 | SRSF2 |
| PSIP1 | 7.91E-51 | 0.520513 | 0.95 | 0.779 | 1.54E-46 | NP2 | PSIP1 |
| RPS31 | 8.72E-51 | 0.34667 | 1 | 1 | 1.70E-46 | NP2 | RPS3 |
| GPSM2 | 1.21E-50 | 0.451654 | 0.479 | 0.126 | 2.35E-46 | NP2 | GPSM2 |
| RPS152 | 2.84E-50 | 0.324707 | 1 | 1 | 5.52E-46 | NP2 | RPS15 |
| NME11 | 1.55E-49 | 0.567421 | 0.959 | 0.721 | 3.02E-45 | NP2 | NME1 |

| | | | | | | | |
|---|---|---|---|---|---|---|---|
| TIMELESS | 3.80E-49 | 0.338986 | 0.541 | 0.155 | 7.40E-45 | NP2 | TIMELESS |
| SERBP11 | 6.76E-49 | 0.467518 | 0.992 | 0.91 | 1.31E-44 | NP2 | SERBP1 |
| HMGN11 | 6.34E-48 | 0.46586 | 0.992 | 0.972 | 1.23E-43 | NP2 | HMGN1 |
| PHGDH1 | 7.89E-48 | 0.657632 | 0.917 | 0.619 | 1.53E-43 | NP2 | PHGDH |
| RPA3 | 9.38E-48 | 0.566824 | 0.888 | 0.581 | 1.82E-43 | NP2 | RPA3 |
| TUBB6 | 2.06E-47 | 0.511775 | 0.525 | 0.166 | 4.00E-43 | NP2 | TUBB6 |
| HES41 | 3.09E-47 | 0.675914 | 0.963 | 0.745 | 6.02E-43 | NP2 | HES4 |
| PARP11 | 4.21E-47 | 0.529642 | 0.942 | 0.709 | 8.20E-43 | NP2 | PARP1 |
| HN11 | 6.41E-47 | 0.528646 | 0.983 | 0.863 | 1.25E-42 | NP2 | HN1 |
| DNAJC9 | 2.34E-46 | 0.509567 | 0.752 | 0.352 | 4.55E-42 | NP2 | DNAJC9 |
| HES61 | 2.43E-46 | 0.816196 | 0.806 | 0.399 | 4.73E-42 | NP2 | HES6 |
| HNRNPM1 | 2.76E-46 | 0.529393 | 0.946 | 0.766 | 5.36E-42 | NP2 | HNRNPM |
| CDC6 | 2.94E-46 | 0.304048 | 0.351 | 0.069 | 5.72E-42 | NP2 | CDC6 |
| CDCA7L1 | 4.47E-46 | 0.434045 | 0.694 | 0.277 | 8.69E-42 | NP2 | CDCA7L |
| FUS1 | 4.90E-46 | 0.485438 | 0.979 | 0.899 | 9.53E-42 | NP2 | FUS |
| AKR1B11 | 6.42E-46 | 0.488961 | 0.884 | 0.62 | 1.25E-41 | NP2 | AKR1B1 |
| KIF22 | 8.31E-46 | 0.628786 | 0.736 | 0.38 | 1.62E-41 | NP2 | KIF22 |
| RPL23A1 | 1.78E-45 | 0.339632 | 1 | 0.998 | 3.47E-41 | NP2 | RPL23A |
| MCM5 | 2.32E-45 | 0.361326 | 0.471 | 0.123 | 4.51E-41 | NP2 | MCM5 |
| FANCG | 5.53E-45 | 0.254619 | 0.438 | 0.11 | 1.08E-40 | NP2 | FANCG |
| TTYH11 | 1.85E-44 | 0.583268 | 0.983 | 0.799 | 3.60E-40 | NP2 | TTYH1 |
| RPL122 | 3.66E-44 | 0.35167 | 1 | 0.999 | 7.13E-40 | NP2 | RPL12 |
| TUBB2B1 | 6.17E-44 | 0.352279 | 1 | 0.981 | 1.20E-39 | NP2 | TUBB2B |
| YBX11 | 7.13E-44 | 0.356309 | 1 | 0.999 | 1.39E-39 | NP2 | YBX1 |
| EXOSC8 | 7.42E-44 | 0.512344 | 0.777 | 0.423 | 1.44E-39 | NP2 | EXOSC8 |
| EZH2 | 7.51E-44 | 0.445843 | 0.657 | 0.275 | 1.46E-39 | NP2 | EZH2 |
| SAE1 | 8.19E-44 | 0.514346 | 0.888 | 0.611 | 1.59E-39 | NP2 | SAE1 |
| RACGAP1 | 1.06E-43 | 0.42965 | 0.512 | 0.163 | 2.06E-39 | NP2 | RACGAP1 |
| TMEM971 | 3.17E-43 | 0.52918 | 0.864 | 0.506 | 6.18E-39 | NP2 | TMEM97 |
| PTN1 | 4.81E-43 | 0.857928 | 0.789 | 0.42 | 9.36E-39 | NP2 | PTN |
| FGFBP31 | 6.76E-43 | 0.694837 | 0.682 | 0.32 | 1.31E-38 | NP2 | FGFBP3 |
| RBMX1 | 1.35E-42 | 0.517606 | 0.967 | 0.872 | 2.63E-38 | NP2 | RBMX |
| RCC1 | 5.15E-42 | 0.317169 | 0.459 | 0.132 | 1.00E-37 | NP2 | RCC1 |
| HNRNPA0 | 5.68E-42 | 0.431251 | 0.988 | 0.955 | 1.11E-37 | NP2 | HNRNPA0 |
| SSRP11 | 1.73E-41 | 0.494162 | 0.921 | 0.704 | 3.37E-37 | NP2 | SSRP1 |
| MMS22L | 2.88E-41 | 0.259551 | 0.38 | 0.088 | 5.61E-37 | NP2 | MMS22L |
| COMMD4 | 1.35E-40 | 0.432196 | 0.731 | 0.374 | 2.63E-36 | NP2 | COMMD4 |
| CRNDE | 1.48E-40 | 0.366312 | 0.479 | 0.148 | 2.89E-36 | NP2 | CRNDE |
| MCM31 | 1.68E-40 | 0.497112 | 0.657 | 0.289 | 3.26E-36 | NP2 | MCM3 |
| RFX4 | 2.41E-40 | 0.37794 | 0.496 | 0.15 | 4.68E-36 | NP2 | RFX4 |

| | | | | | | | |
|---|---|---|---|---|---|---|---|
| SNRPE1 | 3.66E-40 | 0.408674 | 0.983 | 0.914 | 7.11E-36 | NP2 | SNRPE |
| RPLP02 | 4.64E-40 | 0.329115 | 1 | 1 | 9.03E-36 | NP2 | RPLP0 |
| POLD1 | 5.01E-40 | 0.283004 | 0.446 | 0.128 | 9.75E-36 | NP2 | POLD1 |
| MCM4 | 5.28E-40 | 0.321883 | 0.446 | 0.129 | 1.03E-35 | NP2 | MCM4 |
| QKI1 | 8.35E-40 | 0.402542 | 0.711 | 0.324 | 1.63E-35 | NP2 | QKI |
| CTSC1 | 1.79E-39 | 0.290089 | 0.508 | 0.159 | 3.48E-35 | NP2 | CTSC |
| MGST1 | 1.90E-39 | 0.656789 | 0.636 | 0.297 | 3.69E-35 | NP2 | MGST1 |
| AURKA | 2.19E-39 | 0.51511 | 0.355 | 0.087 | 4.26E-35 | NP2 | AURKA |
| CSE1L | 3.23E-39 | 0.424332 | 0.707 | 0.345 | 6.28E-35 | NP2 | CSE1L |
| ILF21 | 3.55E-39 | 0.439178 | 0.959 | 0.838 | 6.91E-35 | NP2 | ILF2 |
| GNAI21 | 3.90E-39 | 0.511193 | 0.938 | 0.733 | 7.58E-35 | NP2 | GNAI2 |
| SFPQ1 | 4.74E-39 | 0.522517 | 0.909 | 0.689 | 9.23E-35 | NP2 | SFPQ |
| LBR | 6.07E-39 | 0.535155 | 0.736 | 0.394 | 1.18E-34 | NP2 | LBR |
| CDK41 | 9.96E-39 | 0.511412 | 0.913 | 0.718 | 1.94E-34 | NP2 | CDK4 |
| UBE2I | 1.01E-38 | 0.402212 | 0.979 | 0.85 | 1.97E-34 | NP2 | UBE2I |
| H2AFY22 | 1.20E-38 | 0.463185 | 0.884 | 0.612 | 2.33E-34 | NP2 | H2AFY2 |
| NUDT1 | 1.45E-38 | 0.395207 | 0.669 | 0.303 | 2.83E-34 | NP2 | NUDT1 |
| MT2A1 | 1.49E-38 | 0.428134 | 0.397 | 0.106 | 2.91E-34 | NP2 | MT2A |
| PRKDC | 1.55E-38 | 0.442327 | 0.831 | 0.485 | 3.02E-34 | NP2 | PRKDC |
| SNRPF1 | 4.54E-38 | 0.418674 | 0.975 | 0.88 | 8.83E-34 | NP2 | SNRPF |
| RNASEH2I | 4.88E-38 | 0.251127 | 0.455 | 0.131 | 9.49E-34 | NP2 | RNASEH2B |
| SRSF71 | 5.11E-38 | 0.469652 | 0.959 | 0.817 | 9.94E-34 | NP2 | SRSF7 |
| CCDC34 | 2.44E-37 | 0.449578 | 0.748 | 0.375 | 4.75E-33 | NP2 | CCDC34 |
| MARCKSL | 4.20E-37 | 0.411659 | 1 | 0.978 | 8.17E-33 | NP2 | MARCKSL1 |
| IDH2 | 8.29E-37 | 0.465887 | 0.913 | 0.676 | 1.61E-32 | NP2 | IDH2 |
| RFC3 | 1.49E-36 | 0.349797 | 0.45 | 0.144 | 2.90E-32 | NP2 | RFC3 |
| DBF4 | 1.51E-36 | 0.375665 | 0.554 | 0.213 | 2.94E-32 | NP2 | DBF4 |
| KCNQ21 | 1.69E-36 | 0.314006 | 0.579 | 0.215 | 3.28E-32 | NP2 | KCNQ2 |
| CKS22 | 2.37E-36 | 0.724602 | 0.926 | 0.768 | 4.60E-32 | NP2 | CKS2 |
| LDHB1 | 2.53E-36 | 0.348408 | 1 | 0.997 | 4.92E-32 | NP2 | LDHB |
| H3F3A1 | 2.57E-36 | 0.310193 | 1 | 0.998 | 4.99E-32 | NP2 | H3F3A |
| 10-Sep | 6.36E-36 | 0.290932 | 0.545 | 0.203 | 1.24E-31 | NP2 | 10-Sep |
| SET1 | 7.96E-36 | 0.433328 | 0.996 | 0.96 | 1.55E-31 | NP2 | SET |
| CMTM7 | 1.16E-35 | 0.257843 | 0.397 | 0.105 | 2.25E-31 | NP2 | CMTM7 |
| H2AFY | 1.29E-35 | 0.434491 | 0.975 | 0.866 | 2.51E-31 | NP2 | H2AFY |
| HNRNPU1 | 1.46E-35 | 0.405561 | 0.975 | 0.845 | 2.84E-31 | NP2 | HNRNPU |
| RHOBTB3 | 2.36E-35 | 0.395378 | 0.769 | 0.406 | 4.58E-31 | NP2 | RHOBTB3 |
| POU3F21 | 2.88E-35 | 0.316692 | 0.537 | 0.185 | 5.60E-31 | NP2 | POU3F2 |
| VCAN2 | 2.95E-35 | 0.291646 | 0.62 | 0.243 | 5.73E-31 | NP2 | VCAN |
| RBBP41 | 4.75E-35 | 0.429189 | 0.946 | 0.777 | 9.23E-31 | NP2 | RBBP4 |

| | | | | | | | |
|---|---|---|---|---|---|---|---|
| HNRNPR | 6.75E-35 | 0.404157 | 0.959 | 0.879 | 1.31E-30 | NP2 | HNRNPR |
| LSM61 | 8.31E-35 | 0.42399 | 0.835 | 0.565 | 1.62E-30 | NP2 | LSM6 |
| FBL1 | 9.77E-35 | 0.416028 | 0.921 | 0.709 | 1.90E-30 | NP2 | FBL |
| NPM11 | 1.13E-34 | 0.341407 | 1 | 0.999 | 2.20E-30 | NP2 | NPM1 |
| XRCC6 | 1.36E-34 | 0.391326 | 0.983 | 0.883 | 2.64E-30 | NP2 | XRCC6 |
| CBX5 | 1.96E-34 | 0.461816 | 0.86 | 0.62 | 3.81E-30 | NP2 | CBX5 |
| EEF1D2 | 1.98E-34 | 0.318223 | 0.996 | 0.981 | 3.85E-30 | NP2 | EEF1D |
| FBLN11 | 2.07E-34 | 0.491908 | 0.661 | 0.329 | 4.03E-30 | NP2 | FBLN1 |
| BCHE | 2.08E-34 | 0.261202 | 0.351 | 0.087 | 4.05E-30 | NP2 | BCHE |
| DUT | 2.23E-34 | 0.600838 | 0.851 | 0.648 | 4.35E-30 | NP2 | DUT |
| CCT51 | 3.23E-34 | 0.421168 | 0.901 | 0.709 | 6.29E-30 | NP2 | CCT5 |
| PSAT11 | 4.48E-34 | 0.543735 | 0.876 | 0.622 | 8.71E-30 | NP2 | PSAT1 |
| GYPC | 6.10E-34 | 0.418914 | 0.558 | 0.228 | 1.19E-29 | NP2 | GYPC |
| PMAIP1 | 7.20E-34 | 0.383192 | 0.504 | 0.18 | 1.40E-29 | NP2 | PMAIP1 |
| HNRNPK1 | 1.03E-33 | 0.338143 | 1 | 0.978 | 1.99E-29 | NP2 | HNRNPK |
| AMBN1 | 1.26E-33 | 0.330137 | 0.264 | 0.051 | 2.45E-29 | NP2 | AMBN |
| BARD1 | 1.71E-33 | 0.295626 | 0.521 | 0.192 | 3.34E-29 | NP2 | BARD1 |
| CDK2 | 3.13E-33 | 0.268114 | 0.488 | 0.17 | 6.08E-29 | NP2 | CDK2 |
| SRSF11 | 3.91E-33 | 0.423054 | 0.88 | 0.63 | 7.60E-29 | NP2 | SRSF1 |
| FABP71 | 7.39E-33 | 0.979983 | 0.826 | 0.548 | 1.44E-28 | NP2 | FABP7 |
| MDK | 1.65E-32 | 0.49747 | 0.996 | 0.948 | 3.22E-28 | NP2 | MDK |
| CHEK1 | 1.71E-32 | 0.306534 | 0.517 | 0.192 | 3.33E-28 | NP2 | CHEK1 |
| BUB3 | 2.50E-32 | 0.548022 | 0.872 | 0.682 | 4.86E-28 | NP2 | BUB3 |
| HN1L | 3.30E-32 | 0.369688 | 0.674 | 0.34 | 6.42E-28 | NP2 | HN1L |
| KNSTRN | 4.32E-32 | 0.425151 | 0.446 | 0.155 | 8.41E-28 | NP2 | KNSTRN |
| FGF17 | 4.65E-32 | 0.663385 | 0.157 | 0.018 | 9.05E-28 | NP2 | FGF17 |
| ACAT21 | 6.10E-32 | 0.555278 | 0.81 | 0.535 | 1.19E-27 | NP2 | ACAT2 |
| MARCKS2 | 6.70E-32 | 0.444241 | 0.983 | 0.899 | 1.30E-27 | NP2 | MARCKS |
| CNTFR1 | 6.87E-32 | 0.380223 | 0.777 | 0.418 | 1.34E-27 | NP2 | CNTFR |
| SKA2 | 7.10E-32 | 0.425541 | 0.727 | 0.416 | 1.38E-27 | NP2 | SKA2 |
| SYNCRIP1 | 7.47E-32 | 0.40602 | 0.839 | 0.552 | 1.45E-27 | NP2 | SYNCRIP |
| HAUS1 | 7.94E-32 | 0.376264 | 0.764 | 0.436 | 1.54E-27 | NP2 | HAUS1 |
| ANP32E | 1.12E-31 | 0.484142 | 0.831 | 0.612 | 2.18E-27 | NP2 | ANP32E |
| NSMCE4A | 1.20E-31 | 0.366049 | 0.727 | 0.389 | 2.34E-27 | NP2 | NSMCE4A |
| UBE2S | 3.09E-31 | 0.762407 | 0.868 | 0.726 | 6.01E-27 | NP2 | UBE2S |
| CBX1 | 5.16E-31 | 0.41648 | 0.959 | 0.816 | 1.00E-26 | NP2 | CBX1 |
| LSM4 | 5.51E-31 | 0.373572 | 0.983 | 0.928 | 1.07E-26 | NP2 | LSM4 |
| TPM4 | 6.27E-31 | 0.403792 | 0.872 | 0.587 | 1.22E-26 | NP2 | TPM4 |
| FAM181A1 | 6.52E-31 | 0.254973 | 0.479 | 0.163 | 1.27E-26 | NP2 | FAM181A |
| LSM5 | 6.66E-31 | 0.410004 | 0.959 | 0.832 | 1.30E-26 | NP2 | LSM5 |

| | | | | | | | |
|---|---|---|---|---|---|---|---|
| ERH | 6.84E-31 | 0.34155 | 0.996 | 0.974 | 1.33E-26 | NP2 | ERH |
| SFRP21 | 1.38E-30 | 0.294349 | 0.731 | 0.372 | 2.69E-26 | NP2 | SFRP2 |
| BOC | 1.45E-30 | 0.264402 | 0.434 | 0.147 | 2.83E-26 | NP2 | BOC |
| SMC3 | 1.57E-30 | 0.395392 | 0.847 | 0.61 | 3.05E-26 | NP2 | SMC3 |
| PTPRZ11 | 2.75E-30 | 0.38685 | 0.607 | 0.267 | 5.35E-26 | NP2 | PTPRZ1 |
| KHDRBS1 | 3.58E-30 | 0.338851 | 0.971 | 0.889 | 6.97E-26 | NP2 | KHDRBS1 |
| PTX3 | 4.64E-30 | 0.298696 | 0.368 | 0.109 | 9.02E-26 | NP2 | PTX3 |
| SRSF10 | 5.23E-30 | 0.400096 | 0.921 | 0.771 | 1.02E-25 | NP2 | SRSF10 |
| TRA2B | 6.61E-30 | 0.374258 | 0.967 | 0.832 | 1.29E-25 | NP2 | TRA2B |
| BAZ1A1 | 7.22E-30 | 0.301231 | 0.64 | 0.291 | 1.40E-25 | NP2 | BAZ1A |
| RPA1 | 8.87E-30 | 0.353045 | 0.723 | 0.408 | 1.73E-25 | NP2 | RPA1 |
| CARHSP1: | 1.55E-29 | 0.427132 | 0.913 | 0.717 | 3.02E-25 | NP2 | CARHSP1 |
| PPP1CC1 | 1.96E-29 | 0.410195 | 0.926 | 0.774 | 3.82E-25 | NP2 | PPP1CC |
| POLD21 | 2.93E-29 | 0.37329 | 0.88 | 0.617 | 5.70E-25 | NP2 | POLD2 |
| ZFP36L11 | 3.06E-29 | 0.446314 | 0.897 | 0.625 | 5.96E-25 | NP2 | ZFP36L1 |
| ANP32A1 | 7.10E-29 | 0.376594 | 0.893 | 0.7 | 1.38E-24 | NP2 | ANP32A |
| SRPK1 | 7.81E-29 | 0.290363 | 0.554 | 0.235 | 1.52E-24 | NP2 | SRPK1 |
| SOX31 | 1.12E-28 | 0.474829 | 0.727 | 0.428 | 2.18E-24 | NP2 | SOX3 |
| TMEM106 | 1.13E-28 | 0.360484 | 0.802 | 0.496 | 2.19E-24 | NP2 | TMEM106C |
| MIS18A | 1.53E-28 | 0.300216 | 0.628 | 0.308 | 2.98E-24 | NP2 | MIS18A |
| SCRG1 | 1.97E-28 | 0.301858 | 0.19 | 0.032 | 3.83E-24 | NP2 | SCRG1 |
| HNRNPAB | 2.74E-28 | 0.411882 | 0.793 | 0.542 | 5.33E-24 | NP2 | HNRNPAB |
| PA2G4 | 4.17E-28 | 0.384657 | 0.926 | 0.793 | 8.12E-24 | NP2 | PA2G4 |
| SYNE21 | 4.45E-28 | 0.368229 | 0.756 | 0.423 | 8.65E-24 | NP2 | SYNE2 |
| TP53 | 4.63E-28 | 0.278415 | 0.533 | 0.22 | 9.00E-24 | NP2 | TP53 |
| DBI1 | 5.30E-28 | 0.444708 | 0.893 | 0.715 | 1.03E-23 | NP2 | DBI |
| RFC2 | 6.72E-28 | 0.318335 | 0.558 | 0.248 | 1.31E-23 | NP2 | RFC2 |
| CCNB1IP1 | 8.42E-28 | 0.370234 | 0.893 | 0.659 | 1.64E-23 | NP2 | CCNB1IP1 |
| HDAC21 | 8.88E-28 | 0.323393 | 0.983 | 0.903 | 1.73E-23 | NP2 | HDAC2 |
| SFRP11 | 1.23E-27 | 0.79907 | 0.579 | 0.307 | 2.40E-23 | NP2 | SFRP1 |
| DTYMK | 1.94E-27 | 0.429575 | 0.764 | 0.527 | 3.77E-23 | NP2 | DTYMK |
| SUPT16H | 2.47E-27 | 0.418444 | 0.764 | 0.498 | 4.80E-23 | NP2 | SUPT16H |
| ARL6IP6 | 3.53E-27 | 0.397965 | 0.773 | 0.488 | 6.87E-23 | NP2 | ARL6IP6 |
| DDX39A | 5.82E-27 | 0.43233 | 0.727 | 0.467 | 1.13E-22 | NP2 | DDX39A |
| PLAGL11 | 1.06E-26 | 0.274034 | 0.533 | 0.227 | 2.07E-22 | NP2 | PLAGL1 |
| LIMD21 | 1.09E-26 | 0.290206 | 0.731 | 0.377 | 2.11E-22 | NP2 | LIMD2 |
| DKC1 | 1.16E-26 | 0.320841 | 0.752 | 0.451 | 2.26E-22 | NP2 | DKC1 |
| CTPS1 | 1.33E-26 | 0.312193 | 0.558 | 0.249 | 2.59E-22 | NP2 | CTPS1 |
| HMGA21 | 1.45E-26 | 0.407675 | 0.678 | 0.359 | 2.82E-22 | NP2 | HMGA2 |
| RBBP7 | 1.60E-26 | 0.381831 | 0.917 | 0.775 | 3.12E-22 | NP2 | RBBP7 |

| | | | | | | | |
|---|---|---|---|---|---|---|---|
| NCL1 | 1.62E-26 | 0.410144 | 0.955 | 0.891 | 3.16E-22 | NP2 | NCL |
| TUBB4B | 1.67E-26 | 0.575553 | 0.938 | 0.838 | 3.24E-22 | NP2 | TUBB4B |
| SMARCC1 | 3.55E-26 | 0.349671 | 0.818 | 0.536 | 6.91E-22 | NP2 | SMARCC1 |
| CBX3 | 3.82E-26 | 0.368702 | 0.917 | 0.767 | 7.42E-22 | NP2 | CBX3 |
| PFN2 | 4.14E-26 | 0.368056 | 0.996 | 0.925 | 8.05E-22 | NP2 | PFN2 |
| SAP30 | 6.56E-26 | 0.29234 | 0.558 | 0.259 | 1.28E-21 | NP2 | SAP30 |
| H1FX | 6.67E-26 | 0.548596 | 0.913 | 0.77 | 1.30E-21 | NP2 | H1FX |
| SNRPA | 9.60E-26 | 0.370589 | 0.806 | 0.557 | 1.87E-21 | NP2 | SNRPA |
| SOX22 | 9.87E-26 | 0.42393 | 0.897 | 0.678 | 1.92E-21 | NP2 | SOX2 |
| TMEM237 | 1.16E-25 | 0.305223 | 0.624 | 0.322 | 2.26E-21 | NP2 | TMEM237 |
| FEN1 | 1.80E-25 | 0.353647 | 0.64 | 0.342 | 3.51E-21 | NP2 | FEN1 |
| SEPHS1 | 2.24E-25 | 0.253815 | 0.537 | 0.238 | 4.35E-21 | NP2 | SEPHS1 |
| NUP107 | 2.92E-25 | 0.263923 | 0.537 | 0.236 | 5.68E-21 | NP2 | NUP107 |
| PMF1 | 7.32E-25 | 0.330325 | 0.793 | 0.518 | 1.42E-20 | NP2 | PMF1 |
| RAD21 | 9.24E-25 | 0.440954 | 0.868 | 0.731 | 1.80E-20 | NP2 | RAD21 |
| CXCR4 | 1.28E-24 | 0.618806 | 0.62 | 0.356 | 2.49E-20 | NP2 | CXCR4 |
| ARGLU1 | 1.31E-24 | 0.347781 | 0.93 | 0.728 | 2.54E-20 | NP2 | ARGLU1 |
| HNRNPH3 | 1.95E-24 | 0.321949 | 0.959 | 0.835 | 3.78E-20 | NP2 | HNRNPH3 |
| WEE1 | 2.07E-24 | 0.273203 | 0.512 | 0.225 | 4.02E-20 | NP2 | WEE1 |
| CDK5RAP: | 3.02E-24 | 0.254564 | 0.479 | 0.202 | 5.88E-20 | NP2 | CDK5RAP2 |
| HIRIP3 | 4.23E-24 | 0.269344 | 0.562 | 0.267 | 8.23E-20 | NP2 | HIRIP3 |
| CNBP1 | 7.72E-24 | 0.305949 | 0.992 | 0.921 | 1.50E-19 | NP2 | CNBP |
| DNMT1 | 1.03E-23 | 0.350606 | 0.649 | 0.386 | 2.00E-19 | NP2 | DNMT1 |
| IGF2BP1 | 1.04E-23 | 0.297657 | 0.583 | 0.289 | 2.03E-19 | NP2 | IGF2BP1 |
| PPIF | 1.17E-23 | 0.274363 | 0.616 | 0.316 | 2.27E-19 | NP2 | PPIF |
| ACTL6A | 1.41E-23 | 0.353727 | 0.777 | 0.532 | 2.74E-19 | NP2 | ACTL6A |
| RFC4 | 1.47E-23 | 0.254559 | 0.508 | 0.228 | 2.86E-19 | NP2 | RFC4 |
| GPM6B1 | 1.98E-23 | 0.37059 | 0.826 | 0.602 | 3.85E-19 | NP2 | GPM6B |
| AHCY | 4.00E-23 | 0.329721 | 0.831 | 0.616 | 7.79E-19 | NP2 | AHCY |
| RPA2 | 4.74E-23 | 0.342529 | 0.793 | 0.558 | 9.21E-19 | NP2 | RPA2 |
| STMN31 | 6.51E-23 | 0.260704 | 0.653 | 0.325 | 1.27E-18 | NP2 | STMN3 |
| CPSF3 | 9.20E-23 | 0.263056 | 0.574 | 0.287 | 1.79E-18 | NP2 | CPSF3 |
| LINC0046: | 1.17E-22 | 0.304149 | 0.533 | 0.249 | 2.28E-18 | NP2 | LINC00461 |
| TIMM10 | 1.36E-22 | 0.314861 | 0.752 | 0.486 | 2.65E-18 | NP2 | TIMM10 |
| PRDX2 | 1.37E-22 | 0.264234 | 1 | 0.985 | 2.67E-18 | NP2 | PRDX2 |
| PFN1 | 2.99E-22 | 0.279804 | 0.996 | 0.969 | 5.82E-18 | NP2 | PFN1 |
| LEF11 | 3.00E-22 | 0.270529 | 0.446 | 0.182 | 5.84E-18 | NP2 | LEF1 |
| CHD72 | 3.21E-22 | 0.302934 | 0.798 | 0.524 | 6.24E-18 | NP2 | CHD7 |
| ASRGL1 | 4.88E-22 | 0.310922 | 0.669 | 0.392 | 9.49E-18 | NP2 | ASRGL1 |
| CAMTA1 | 6.44E-22 | 0.297927 | 0.926 | 0.763 | 1.25E-17 | NP2 | CAMTA1 |

| | | | | | | | |
|---|---|---|---|---|---|---|---|
| ILF31 | 7.73E-22 | 0.317585 | 0.88 | 0.754 | 1.50E-17 | NP2 | ILF3 |
| NONO1 | 7.86E-22 | 0.280505 | 0.967 | 0.887 | 1.53E-17 | NP2 | NONO |
| FKBP3 | 7.90E-22 | 0.323464 | 0.926 | 0.713 | 1.54E-17 | NP2 | FKBP3 |
| SEPT111 | 1.15E-21 | 0.347786 | 0.884 | 0.682 | 2.23E-17 | NP2 | 11-Sep |
| RHNO1 | 1.62E-21 | 0.314875 | 0.653 | 0.385 | 3.15E-17 | NP2 | RHNO1 |
| GDI2 | 2.79E-21 | 0.286314 | 0.979 | 0.893 | 5.43E-17 | NP2 | GDI2 |
| RP3-395M | 3.60E-21 | 0.25245 | 0.413 | 0.16 | 7.01E-17 | NP2 | RP3-395M20.12 |
| GNG11 | 4.07E-21 | 0.326095 | 0.475 | 0.218 | 7.92E-17 | NP2 | GNG11 |
| MRPL111 | 5.32E-21 | 0.315461 | 0.793 | 0.558 | 1.03E-16 | NP2 | MRPL11 |
| PLEKHO1 | 5.58E-21 | 0.269314 | 0.661 | 0.366 | 1.09E-16 | NP2 | PLEKHO1 |
| DCTPP1 | 6.91E-21 | 0.323998 | 0.727 | 0.469 | 1.34E-16 | NP2 | DCTPP1 |
| EIF1AX | 8.94E-21 | 0.329578 | 0.901 | 0.785 | 1.74E-16 | NP2 | EIF1AX |
| RDX | 9.20E-21 | 0.328587 | 0.88 | 0.666 | 1.79E-16 | NP2 | RDX |
| GCSH1 | 9.53E-21 | 0.340116 | 0.88 | 0.74 | 1.85E-16 | NP2 | GCSH |
| LIX11 | 1.07E-20 | 0.286181 | 0.579 | 0.295 | 2.08E-16 | NP2 | LIX1 |
| HAT1 | 1.29E-20 | 0.356016 | 0.694 | 0.445 | 2.51E-16 | NP2 | HAT1 |
| HSPD1 | 1.90E-20 | 0.33718 | 1 | 0.941 | 3.69E-16 | NP2 | HSPD1 |
| PSMA2 | 2.82E-20 | 0.280852 | 0.975 | 0.922 | 5.48E-16 | NP2 | PSMA2 |
| LIG1 | 3.30E-20 | 0.292601 | 0.483 | 0.236 | 6.43E-16 | NP2 | LIG1 |
| DERA | 3.71E-20 | 0.251561 | 0.471 | 0.222 | 7.22E-16 | NP2 | DERA |
| DCXR | 4.52E-20 | 0.337037 | 0.909 | 0.769 | 8.79E-16 | NP2 | DCXR |
| ELAVL1 | 5.27E-20 | 0.305961 | 0.905 | 0.765 | 1.03E-15 | NP2 | ELAVL1 |
| APEX11 | 8.26E-20 | 0.329411 | 0.901 | 0.764 | 1.61E-15 | NP2 | APEX1 |
| PDCD51 | 8.80E-20 | 0.284584 | 0.901 | 0.759 | 1.71E-15 | NP2 | PDCD5 |
| RBM32 | 9.13E-20 | 0.319923 | 0.901 | 0.77 | 1.78E-15 | NP2 | RBM3 |
| POLR2J | 9.23E-20 | 0.289476 | 0.93 | 0.835 | 1.80E-15 | NP2 | POLR2J |
| PAICS1 | 1.33E-19 | 0.332427 | 0.843 | 0.67 | 2.59E-15 | NP2 | PAICS |
| PEA15 | 1.41E-19 | 0.288666 | 0.636 | 0.36 | 2.75E-15 | NP2 | PEA15 |
| LINC0155 | 1.92E-19 | 0.396469 | 0.355 | 0.133 | 3.74E-15 | NP2 | LINC01551 |
| CKAP5 | 2.31E-19 | 0.270084 | 0.455 | 0.209 | 4.50E-15 | NP2 | CKAP5 |
| PSMA4 | 2.58E-19 | 0.296834 | 0.979 | 0.932 | 5.03E-15 | NP2 | PSMA4 |
| NME41 | 2.96E-19 | 0.326208 | 0.868 | 0.68 | 5.75E-15 | NP2 | NME4 |
| TEAD2 | 3.19E-19 | 0.280506 | 0.591 | 0.337 | 6.20E-15 | NP2 | TEAD2 |
| NT5DC2 | 3.33E-19 | 0.300633 | 0.574 | 0.323 | 6.49E-15 | NP2 | NT5DC2 |
| RNF1301 | 6.63E-19 | 0.28877 | 0.748 | 0.508 | 1.29E-14 | NP2 | RNF130 |
| SMARCB1 | 9.03E-19 | 0.250156 | 0.971 | 0.86 | 1.76E-14 | NP2 | SMARCB1 |
| C5orf301 | 9.50E-19 | 0.254527 | 0.496 | 0.238 | 1.85E-14 | NP2 | C5orf30 |
| ZNF7381 | 9.56E-19 | 0.288661 | 0.769 | 0.527 | 1.86E-14 | NP2 | ZNF738 |
| CCDC167 | 1.12E-18 | 0.298865 | 0.868 | 0.694 | 2.17E-14 | NP2 | CCDC167 |
| MRPL51 | 1.12E-18 | 0.272083 | 0.926 | 0.856 | 2.18E-14 | NP2 | MRPL51 |

| | | | | | | | |
|---|---|---|---|---|---|---|---|
| C20orf27 | 1.30E-18 | 0.290664 | 0.727 | 0.475 | 2.53E-14 | NP2 | C20orf27 |
| HMGXB4 | 1.52E-18 | 0.290113 | 0.674 | 0.435 | 2.96E-14 | NP2 | HMGXB4 |
| SRSF9 | 1.84E-18 | 0.252551 | 0.963 | 0.911 | 3.58E-14 | NP2 | SRSF9 |
| SNRNP40 | 1.90E-18 | 0.297404 | 0.785 | 0.563 | 3.69E-14 | NP2 | SNRNP40 |
| RNF26 | 2.09E-18 | 0.318668 | 0.599 | 0.365 | 4.06E-14 | NP2 | RNF26 |
| FDFT11 | 2.14E-18 | 0.418361 | 0.917 | 0.836 | 4.17E-14 | NP2 | FDFT1 |
| STRA13 | 2.57E-18 | 0.322801 | 0.769 | 0.553 | 4.99E-14 | NP2 | STRA13 |
| SEC11A | 2.64E-18 | 0.271585 | 0.988 | 0.917 | 5.14E-14 | NP2 | SEC11A |
| C4orf27 | 3.20E-18 | 0.281899 | 0.752 | 0.513 | 6.23E-14 | NP2 | C4orf27 |
| LHX22 | 3.36E-18 | 0.400565 | 0.36 | 0.148 | 6.53E-14 | NP2 | LHX2 |
| MPHOSPH | 4.79E-18 | 0.262192 | 0.64 | 0.368 | 9.31E-14 | NP2 | MPHOSPH6 |
| PPA11 | 4.96E-18 | 0.292913 | 0.897 | 0.732 | 9.64E-14 | NP2 | PPA1 |
| CALD1 | 5.06E-18 | 0.25425 | 0.814 | 0.573 | 9.84E-14 | NP2 | CALD1 |
| ODC11 | 5.62E-18 | 0.273815 | 0.971 | 0.931 | 1.09E-13 | NP2 | ODC1 |
| DRAXIN1 | 6.42E-18 | 0.325798 | 0.603 | 0.34 | 1.25E-13 | NP2 | DRAXIN |
| PTBP1 | 6.72E-18 | 0.2823 | 0.798 | 0.588 | 1.31E-13 | NP2 | PTBP1 |
| RHEB | 9.34E-18 | 0.267213 | 0.979 | 0.909 | 1.82E-13 | NP2 | RHEB |
| CTNNAL1 | 1.03E-17 | 0.273748 | 0.599 | 0.342 | 2.00E-13 | NP2 | CTNNAL1 |
| CA14 | 1.42E-17 | 0.297035 | 0.521 | 0.282 | 2.77E-13 | NP2 | CA14 |
| SNRNP25 | 1.60E-17 | 0.281821 | 0.789 | 0.572 | 3.12E-13 | NP2 | SNRNP25 |
| SREK1 | 2.68E-17 | 0.266538 | 0.698 | 0.465 | 5.21E-13 | NP2 | SREK1 |
| HP1BP3 | 4.00E-17 | 0.333244 | 0.798 | 0.592 | 7.78E-13 | NP2 | HP1BP3 |
| UBE2N1 | 4.13E-17 | 0.311553 | 0.872 | 0.745 | 8.04E-13 | NP2 | UBE2N |
| PXMP2 | 4.88E-17 | 0.254544 | 0.566 | 0.313 | 9.49E-13 | NP2 | PXMP2 |
| GAS11 | 6.24E-17 | 0.255669 | 0.492 | 0.25 | 1.21E-12 | NP2 | GAS1 |
| LSM2 | 7.64E-17 | 0.258896 | 0.884 | 0.75 | 1.49E-12 | NP2 | LSM2 |
| KPNB1 | 8.38E-17 | 0.293853 | 0.876 | 0.767 | 1.63E-12 | NP2 | KPNB1 |
| DHX91 | 1.15E-16 | 0.267458 | 0.669 | 0.428 | 2.24E-12 | NP2 | DHX9 |
| TALDO1 | 1.71E-16 | 0.276181 | 0.86 | 0.702 | 3.32E-12 | NP2 | TALDO1 |
| ENY2 | 2.05E-16 | 0.250781 | 0.942 | 0.848 | 3.99E-12 | NP2 | ENY2 |
| EWSR1 | 2.72E-16 | 0.267025 | 0.855 | 0.706 | 5.30E-12 | NP2 | EWSR1 |
| TPGS21 | 2.92E-16 | 0.262381 | 0.868 | 0.705 | 5.68E-12 | NP2 | TPGS2 |
| RNPS1 | 4.94E-16 | 0.260841 | 0.942 | 0.859 | 9.60E-12 | NP2 | RNPS1 |
| PBX11 | 7.15E-16 | 0.261476 | 0.719 | 0.493 | 1.39E-11 | NP2 | PBX1 |
| MAGOH | 7.16E-16 | 0.265695 | 0.868 | 0.66 | 1.39E-11 | NP2 | MAGOH |
| SLC16A1 | 7.37E-16 | 0.261968 | 0.591 | 0.361 | 1.43E-11 | NP2 | SLC16A1 |
| G3BP1 | 7.98E-16 | 0.293116 | 0.752 | 0.547 | 1.55E-11 | NP2 | G3BP1 |
| IL13RA2 | 9.66E-16 | 0.297993 | 0.256 | 0.089 | 1.88E-11 | NP2 | IL13RA2 |
| CLYBL | 1.57E-15 | 0.285373 | 0.343 | 0.154 | 3.05E-11 | NP2 | CLYBL |
| MAPK1IP1 | 1.62E-15 | 0.259613 | 0.843 | 0.715 | 3.16E-11 | NP2 | MAPK1IP1L |

| | | | | | | | |
|---|---|---|---|---|---|---|---|
| PABPC11 | 2.02E-15 | 0.257034 | 0.988 | 0.961 | 3.93E-11 | NP2 | PABPC1 |
| MAD2L2 | 2.03E-15 | 0.269455 | 0.851 | 0.713 | 3.96E-11 | NP2 | MAD2L2 |
| BLOC1S1 | 3.83E-15 | 0.271558 | 0.839 | 0.647 | 7.46E-11 | NP2 | BLOC1S1 |
| XRN2 | 4.31E-15 | 0.256854 | 0.847 | 0.662 | 8.38E-11 | NP2 | XRN2 |
| ID41 | 4.44E-15 | 0.426762 | 0.88 | 0.735 | 8.64E-11 | NP2 | ID4 |
| NAE11 | 6.09E-15 | 0.262129 | 0.744 | 0.553 | 1.18E-10 | NP2 | NAE1 |
| SIVA11 | 8.53E-15 | 0.25291 | 0.901 | 0.762 | 1.66E-10 | NP2 | SIVA1 |
| PPP2CA | 1.35E-14 | 0.265245 | 0.884 | 0.759 | 2.62E-10 | NP2 | PPP2CA |
| HNRNPH1 | 3.75E-14 | 0.323937 | 0.835 | 0.655 | 7.30E-10 | NP2 | HNRNPH1 |
| UQCC2 | 4.64E-14 | 0.256252 | 0.905 | 0.727 | 9.03E-10 | NP2 | UQCC2 |
| FDPS1 | 1.13E-13 | 0.278499 | 0.93 | 0.845 | 2.20E-09 | NP2 | FDPS |
| EIF4EBP1 | 3.08E-13 | 0.336835 | 0.723 | 0.535 | 5.99E-09 | NP2 | EIF4EBP1 |
| HEY11 | 3.97E-13 | 0.260717 | 0.463 | 0.253 | 7.73E-09 | NP2 | HEY1 |
| EFNB11 | 1.29E-12 | 0.262041 | 0.653 | 0.437 | 2.51E-08 | NP2 | EFNB1 |
| GGCT | 1.86E-12 | 0.253772 | 0.723 | 0.524 | 3.62E-08 | NP2 | GGCT |
| EZR | 1.86E-12 | 0.25637 | 0.636 | 0.45 | 3.62E-08 | NP2 | EZR |
| SHMT21 | 5.85E-12 | 0.261938 | 0.694 | 0.47 | 1.14E-07 | NP2 | SHMT2 |
| METRN | 2.42E-11 | 0.32306 | 0.835 | 0.707 | 4.71E-07 | NP2 | METRN |
| HMGCS12 | 9.67E-11 | 0.355032 | 0.88 | 0.808 | 1.88E-06 | NP2 | HMGCS1 |
| PON2 | 1.06E-10 | 0.252872 | 0.715 | 0.537 | 2.07E-06 | NP2 | PON2 |
| DLL11 | 5.59E-10 | 0.261431 | 0.38 | 0.208 | 1.09E-05 | NP2 | DLL1 |
| HSP90AA1 | 1.40E-08 | 0.33908 | 1 | 0.994 | 0.000273 | NP2 | HSP90AA1 |
| NUDCD2 | 8.05E-08 | 0.250343 | 0.661 | 0.563 | 0.001565 | NP2 | NUDCD2 |
| ARL4A | 2.12E-07 | 0.256764 | 0.715 | 0.595 | 0.004134 | NP2 | ARL4A |
| CCND2 | 2.28E-07 | 0.316767 | 0.591 | 0.465 | 0.004434 | NP2 | CCND2 |
| C1orf612 | 6.13E-07 | 0.507551 | 0.45 | 0.357 | 0.011929 | NP2 | C1orf61 |
| ID2 | 1.42E-05 | 0.33437 | 0.802 | 0.725 | 0.277063 | NP2 | ID2 |
| ID3 | 1.60E-05 | 0.305246 | 0.876 | 0.775 | 0.311201 | NP2 | ID3 |
| EGR11 | 1.93E-05 | 0.334943 | 0.628 | 0.536 | 0.37592 | NP2 | EGR1 |
| ARL6IP1 | 4.88E-05 | 0.600071 | 0.938 | 0.929 | 0.94993 | NP2 | ARL6IP1 |
| DLK1 | 0.000468 | 0.672196 | 0.756 | 0.713 | 1 | NP2 | DLK1 |
| TUBA1C | 0.001137 | 0.27162 | 0.438 | 0.352 | 1 | NP2 | TUBA1C |
| MDK1 | 3.44E-99 | 0.977414 | 1 | 0.947 | 6.68E-95 | CE | MDK |
| TPBG | 1.02E-76 | 1.228418 | 0.97 | 0.665 | 1.99E-72 | CE | TPBG |
| ZIC1 | 3.26E-74 | 0.861437 | 0.784 | 0.364 | 6.34E-70 | CE | ZIC1 |
| RP4-665J2 | 4.73E-67 | 0.762105 | 0.583 | 0.16 | 9.21E-63 | CE | RP4-665J23.1 |
| BARHL21 | 1.19E-61 | 0.552992 | 0.371 | 0.06 | 2.31E-57 | CE | BARHL2 |
| ZIC4 | 1.52E-56 | 0.657627 | 0.595 | 0.212 | 2.96E-52 | CE | ZIC4 |
| CBLN1 | 1.34E-55 | 0.754651 | 0.318 | 0.05 | 2.61E-51 | CE | CBLN1 |
| PCDH91 | 1.72E-51 | 0.611172 | 0.519 | 0.158 | 3.35E-47 | CE | PCDH9 |

| | | | | | | | |
|---|---|---|---|---|---|---|---|
| WNT2B | 7.22E-43 | 0.773696 | 0.326 | 0.072 | 1.40E-38 | CE | WNT2B |
| VIM2 | 3.61E-42 | 0.737098 | 0.996 | 0.952 | 7.03E-38 | CE | VIM |
| CIRBP1 | 2.42E-41 | 0.378484 | 0.996 | 0.987 | 4.70E-37 | CE | CIRBP |
| MYL61 | 3.19E-41 | 0.477703 | 1 | 0.988 | 6.21E-37 | CE | MYL6 |
| ZFP36L12 | 4.54E-41 | 0.572067 | 0.89 | 0.623 | 8.84E-37 | CE | ZFP36L1 |
| PAX61 | 3.93E-40 | 0.780062 | 0.799 | 0.556 | 7.64E-36 | CE | PAX6 |
| WFIKKN1 | 1.80E-39 | 0.484121 | 0.576 | 0.242 | 3.50E-35 | CE | WFIKKN1 |
| SRP14 | 3.91E-39 | 0.319561 | 1 | 0.999 | 7.61E-35 | CE | SRP14 |
| RP11-849I | 9.54E-37 | 0.321373 | 0.25 | 0.046 | 1.86E-32 | CE | RP11-849I19.1 |
| CA10 | 1.29E-33 | 0.25668 | 0.178 | 0.024 | 2.52E-29 | CE | CA10 |
| PAX3 | 1.89E-33 | 0.319492 | 0.223 | 0.04 | 3.67E-29 | CE | PAX3 |
| SPARCL1 | 7.21E-33 | 0.266365 | 0.129 | 0.011 | 1.40E-28 | CE | SPARCL1 |
| CNTNAP2 | 6.75E-32 | 0.518885 | 0.928 | 0.777 | 1.31E-27 | CE | CNTNAP2 |
| CDH21 | 2.52E-31 | 0.534358 | 0.879 | 0.698 | 4.90E-27 | CE | CDH2 |
| RAB34 | 9.83E-31 | 0.41889 | 0.606 | 0.328 | 1.91E-26 | CE | RAB34 |
| FSTL1 | 3.94E-30 | 0.374213 | 0.636 | 0.332 | 7.66E-26 | CE | FSTL1 |
| BMP4 | 5.02E-28 | 0.331558 | 0.239 | 0.056 | 9.77E-24 | CE | BMP4 |
| TGIF1 | 5.32E-28 | 0.50666 | 0.746 | 0.521 | 1.04E-23 | CE | TGIF1 |
| CDH6 | 1.59E-27 | 0.425334 | 0.352 | 0.123 | 3.09E-23 | CE | CDH6 |
| SPRY1 | 3.04E-27 | 0.457327 | 0.564 | 0.292 | 5.92E-23 | CE | SPRY1 |
| WLS | 4.62E-27 | 0.433037 | 0.864 | 0.594 | 8.98E-23 | CE | WLS |
| CLU | 6.46E-27 | 0.510809 | 0.955 | 0.924 | 1.26E-22 | CE | CLU |
| AC009501 | 7.93E-27 | 0.355759 | 0.379 | 0.14 | 1.54E-22 | CE | AC009501.4 |
| PPDPF | 1.12E-25 | 0.264001 | 0.996 | 0.994 | 2.18E-21 | CE | PPDPF |
| TNNT1 | 2.35E-25 | 0.494043 | 0.606 | 0.347 | 4.57E-21 | CE | TNNT1 |
| ANXA2 | 3.66E-25 | 0.51952 | 0.758 | 0.538 | 7.11E-21 | CE | ANXA2 |
| SH3BP51 | 1.90E-24 | 0.366037 | 0.481 | 0.223 | 3.69E-20 | CE | SH3BP5 |
| CD9 | 3.91E-24 | 0.382712 | 0.443 | 0.2 | 7.61E-20 | CE | CD9 |
| H3F3A2 | 4.07E-24 | 0.254756 | 1 | 0.998 | 7.91E-20 | CE | H3F3A |
| MARCKSL | 4.55E-24 | 0.307546 | 0.996 | 0.978 | 8.86E-20 | CE | MARCKSL1 |
| SH3BGRL | 2.59E-23 | 0.381299 | 0.894 | 0.757 | 5.04E-19 | CE | SH3BGRL |
| ID31 | 4.27E-23 | 0.731049 | 0.883 | 0.773 | 8.30E-19 | CE | ID3 |
| PLTP | 1.82E-21 | 0.373821 | 0.909 | 0.741 | 3.54E-17 | CE | PLTP |
| RCN2 | 1.27E-20 | 0.251228 | 0.996 | 0.979 | 2.48E-16 | CE | RCN2 |
| SYT41 | 1.72E-20 | 0.430576 | 0.277 | 0.09 | 3.34E-16 | CE | SYT4 |
| ATP2B1 | 1.83E-20 | 0.41174 | 0.735 | 0.557 | 3.55E-16 | CE | ATP2B1 |
| MARCKS3 | 1.87E-20 | 0.321752 | 0.981 | 0.899 | 3.64E-16 | CE | MARCKS |
| KCNJ2 | 2.73E-20 | 0.252264 | 0.197 | 0.05 | 5.32E-16 | CE | KCNJ2 |
| RP13-436I | 8.84E-20 | 0.317921 | 0.428 | 0.209 | 1.72E-15 | CE | RP13-436F16.1 |
| SDK2 | 2.03E-19 | 0.348869 | 0.477 | 0.262 | 3.95E-15 | CE | SDK2 |

| | | | | | | | |
|---|---|---|---|---|---|---|---|
| SPARC | 2.18E-19 | 0.413672 | 0.92 | 0.802 | 4.25E-15 | CE | SPARC |
| TSPAN6 | 2.75E-19 | 0.384319 | 0.799 | 0.656 | 5.35E-15 | CE | TSPAN6 |
| MT-CO3 | 8.59E-19 | 0.336659 | 0.996 | 0.981 | 1.67E-14 | CE | MT-CO3 |
| APP1 | 8.68E-19 | 0.344257 | 0.939 | 0.871 | 1.69E-14 | CE | APP |
| NFIA | 1.44E-18 | 0.365083 | 0.348 | 0.158 | 2.80E-14 | CE | NFIA |
| METRN1 | 1.93E-18 | 0.340681 | 0.841 | 0.705 | 3.76E-14 | CE | METRN |
| ZFHX41 | 2.34E-18 | 0.322591 | 0.655 | 0.443 | 4.55E-14 | CE | ZFHX4 |
| TENM3 | 2.39E-18 | 0.278003 | 0.314 | 0.13 | 4.64E-14 | CE | TENM3 |
| SEC11A1 | 4.31E-18 | 0.265526 | 0.936 | 0.923 | 8.39E-14 | CE | SEC11A |
| SPINT2 | 4.48E-18 | 0.400058 | 0.856 | 0.742 | 8.71E-14 | CE | SPINT2 |
| COL9A1 | 7.14E-18 | 0.342585 | 0.28 | 0.109 | 1.39E-13 | CE | COL9A1 |
| MAGED2 | 1.07E-17 | 0.277266 | 0.981 | 0.951 | 2.07E-13 | CE | MAGED2 |
| COL9A2 | 1.30E-17 | 0.257678 | 0.292 | 0.114 | 2.53E-13 | CE | COL9A2 |
| DECR1 | 1.82E-17 | 0.338593 | 0.731 | 0.543 | 3.55E-13 | CE | DECR1 |
| IFITM3 | 1.83E-17 | 0.386062 | 0.826 | 0.668 | 3.56E-13 | CE | IFITM3 |
| OTX1 | 2.11E-17 | 0.263842 | 0.379 | 0.182 | 4.11E-13 | CE | OTX1 |
| NTPCR | 2.41E-17 | 0.321981 | 0.723 | 0.565 | 4.68E-13 | CE | NTPCR |
| CLYBL1 | 3.41E-17 | 0.310622 | 0.337 | 0.152 | 6.63E-13 | CE | CLYBL |
| IGFBP5 | 3.99E-17 | 0.584079 | 0.697 | 0.442 | 7.75E-13 | CE | IGFBP5 |
| LIX12 | 5.51E-17 | 0.370749 | 0.515 | 0.3 | 1.07E-12 | CE | LIX1 |
| ID1 | 6.54E-17 | 0.631999 | 0.788 | 0.689 | 1.27E-12 | CE | ID1 |
| CLIC1 | 8.02E-17 | 0.305863 | 0.905 | 0.847 | 1.56E-12 | CE | CLIC1 |
| FLRT3 | 8.64E-17 | 0.25412 | 0.337 | 0.151 | 1.68E-12 | CE | FLRT3 |
| CPE1 | 1.48E-16 | 0.396078 | 0.803 | 0.703 | 2.88E-12 | CE | CPE |
| TSC22D11 | 1.62E-16 | 0.349861 | 0.924 | 0.895 | 3.15E-12 | CE | TSC22D1 |
| MSI2 | 3.65E-16 | 0.335079 | 0.739 | 0.588 | 7.10E-12 | CE | MSI2 |
| H1F0 | 5.29E-16 | 0.36431 | 0.549 | 0.343 | 1.03E-11 | CE | H1F0 |
| CTNNB1 | 6.27E-16 | 0.403892 | 0.716 | 0.544 | 1.22E-11 | CE | CTNNB1 |
| SCD51 | 6.69E-16 | 0.323109 | 0.606 | 0.416 | 1.30E-11 | CE | SCD5 |
| MFAP2 | 6.96E-16 | 0.326775 | 0.61 | 0.432 | 1.35E-11 | CE | MFAP2 |
| PGLS | 9.01E-16 | 0.313386 | 0.848 | 0.785 | 1.75E-11 | CE | PGLS |
| PDIA6 | 1.10E-15 | 0.283881 | 0.962 | 0.909 | 2.13E-11 | CE | PDIA6 |
| TMEM38B | 2.17E-15 | 0.330949 | 0.746 | 0.605 | 4.23E-11 | CE | TMEM38B |
| CITED2 | 2.95E-15 | 0.405756 | 0.811 | 0.681 | 5.75E-11 | CE | CITED2 |
| SERPING1 | 3.50E-15 | 0.268648 | 0.5 | 0.315 | 6.81E-11 | CE | SERPING1 |
| MT-CYB | 4.56E-15 | 0.327039 | 0.996 | 0.962 | 8.88E-11 | CE | MT-CYB |
| GLUL | 4.77E-15 | 0.360652 | 0.932 | 0.854 | 9.29E-11 | CE | GLUL |
| ZIC2 | 4.78E-15 | 0.334837 | 0.458 | 0.281 | 9.30E-11 | CE | ZIC2 |
| ANXA6 | 6.60E-15 | 0.291532 | 0.648 | 0.436 | 1.28E-10 | CE | ANXA6 |
| SOX23 | 6.63E-15 | 0.454174 | 0.803 | 0.687 | 1.29E-10 | CE | SOX2 |

| | | | | | | | |
|---|---|---|---|---|---|---|---|
| TM7SF2 | 8.27E-15 | 0.356395 | 0.773 | 0.642 | 1.61E-10 | CE | TM7SF2 |
| TLE4 | 1.40E-14 | 0.351475 | 0.519 | 0.343 | 2.72E-10 | CE | TLE4 |
| SFRP22 | 1.40E-14 | 0.549895 | 0.557 | 0.39 | 2.72E-10 | CE | SFRP2 |
| COL6A2 | 1.57E-14 | 0.288798 | 0.432 | 0.253 | 3.04E-10 | CE | COL6A2 |
| SDC2 | 1.97E-14 | 0.386953 | 0.644 | 0.473 | 3.84E-10 | CE | SDC2 |
| SYTL1 | 3.06E-14 | 0.266341 | 0.379 | 0.198 | 5.96E-10 | CE | SYTL1 |
| COL6A1 | 8.13E-14 | 0.277071 | 0.701 | 0.563 | 1.58E-09 | CE | COL6A1 |
| GGH | 1.16E-13 | 0.273516 | 0.788 | 0.685 | 2.25E-09 | CE | GGH |
| PGD | 1.22E-13 | 0.266771 | 0.837 | 0.71 | 2.38E-09 | CE | PGD |
| C7orf50 | 3.33E-13 | 0.252666 | 0.879 | 0.807 | 6.47E-09 | CE | C7orf50 |
| PDIA3 | 3.86E-13 | 0.300195 | 0.89 | 0.87 | 7.51E-09 | CE | PDIA3 |
| PFN21 | 7.80E-13 | 0.282337 | 0.977 | 0.927 | 1.52E-08 | CE | PFN2 |
| MED28 | 1.09E-12 | 0.25082 | 0.83 | 0.748 | 2.12E-08 | CE | MED28 |
| GOLM1 | 1.25E-12 | 0.308797 | 0.689 | 0.592 | 2.43E-08 | CE | GOLM1 |
| DLK11 | 1.25E-12 | 1.102142 | 0.784 | 0.709 | 2.43E-08 | CE | DLK1 |
| TCF7L2 | 1.64E-12 | 0.452213 | 0.534 | 0.359 | 3.19E-08 | CE | TCF7L2 |
| NRN11 | 1.70E-12 | 0.573871 | 0.47 | 0.295 | 3.30E-08 | CE | NRN1 |
| NAT14 | 1.83E-12 | 0.271556 | 0.648 | 0.51 | 3.56E-08 | CE | NAT14 |
| EGFL6 | 2.38E-12 | 0.353748 | 0.163 | 0.053 | 4.64E-08 | CE | EGFL6 |
| RASL11B | 2.42E-12 | 0.314152 | 0.227 | 0.096 | 4.71E-08 | CE | RASL11B |
| HACD3 | 2.48E-12 | 0.255171 | 0.924 | 0.858 | 4.83E-08 | CE | HACD3 |
| MT-ATP6 | 4.64E-12 | 0.254893 | 0.977 | 0.902 | 9.03E-08 | CE | MT-ATP6 |
| PERP | 7.81E-12 | 0.287725 | 0.549 | 0.388 | 1.52E-07 | CE | PERP |
| ESD | 8.27E-12 | 0.287716 | 0.837 | 0.794 | 1.61E-07 | CE | ESD |
| NLRP1 | 8.86E-12 | 0.362629 | 0.837 | 0.753 | 1.72E-07 | CE | NLRP1 |
| TUBA1C1 | 1.12E-11 | 0.336221 | 0.508 | 0.342 | 2.17E-07 | CE | TUBA1C |
| CD59 | 2.10E-11 | 0.258449 | 0.674 | 0.566 | 4.08E-07 | CE | CD59 |
| HYAL2 | 2.97E-11 | 0.263842 | 0.617 | 0.49 | 5.78E-07 | CE | HYAL2 |
| LGMN | 3.03E-11 | 0.269835 | 0.723 | 0.629 | 5.90E-07 | CE | LGMN |
| S100A11 | 3.18E-11 | 0.368995 | 0.356 | 0.201 | 6.19E-07 | CE | S100A11 |
| S100A6 | 3.51E-11 | 0.318981 | 0.436 | 0.277 | 6.83E-07 | CE | S100A6 |
| CXCR41 | 3.63E-11 | 0.409292 | 0.53 | 0.364 | 7.05E-07 | CE | CXCR4 |
| CDC421 | 3.81E-11 | 0.255074 | 0.883 | 0.818 | 7.42E-07 | CE | CDC42 |
| EMP3 | 6.47E-11 | 0.305781 | 0.58 | 0.432 | 1.26E-06 | CE | EMP3 |
| CYBA | 6.98E-11 | 0.255261 | 0.72 | 0.578 | 1.36E-06 | CE | CYBA |
| PTN2 | 1.06E-10 | 0.361373 | 0.61 | 0.439 | 2.07E-06 | CE | PTN |
| GAS6 | 1.39E-10 | 0.293461 | 0.417 | 0.272 | 2.71E-06 | CE | GAS6 |
| KCNQ1OT | 1.42E-10 | 0.257629 | 0.841 | 0.708 | 2.76E-06 | CE | KCNQ1OT1 |
| SLC22A17 | 2.60E-10 | 0.273675 | 0.811 | 0.704 | 5.06E-06 | CE | SLC22A17 |
| CPVL | 3.97E-10 | 0.259835 | 0.326 | 0.187 | 7.72E-06 | CE | CPVL |

| | | | | | | | |
|---|---|---|---|---|---|---|---|
| NES2 | 4.47E-10 | 0.312245 | 0.712 | 0.571 | 8.69E-06 | CE | NES |
| HLA-A | 5.26E-10 | 0.26274 | 0.67 | 0.555 | 1.02E-05 | CE | HLA-A |
| CCNG2 | 5.69E-10 | 0.328415 | 0.496 | 0.37 | 1.11E-05 | CE | CCNG2 |
| OBSL1 | 7.92E-10 | 0.260787 | 0.53 | 0.398 | 1.54E-05 | CE | OBSL1 |
| CA141 | 8.98E-10 | 0.327666 | 0.432 | 0.29 | 1.75E-05 | CE | CA14 |
| NELL2 | 1.03E-09 | 0.322904 | 0.523 | 0.387 | 2.01E-05 | CE | NELL2 |
| PRCP | 1.34E-09 | 0.291876 | 0.595 | 0.481 | 2.61E-05 | CE | PRCP |
| APLP2 | 1.47E-09 | 0.283548 | 0.86 | 0.834 | 2.85E-05 | CE | APLP2 |
| MGST11 | 2.29E-09 | 0.460826 | 0.451 | 0.317 | 4.46E-05 | CE | MGST1 |
| NME3 | 3.15E-09 | 0.256058 | 0.689 | 0.606 | 6.13E-05 | CE | NME3 |
| CTSL | 5.28E-09 | 0.254458 | 0.64 | 0.556 | 0.000103 | CE | CTSL |
| SNCA | 7.39E-09 | 0.327495 | 0.462 | 0.328 | 0.000144 | CE | SNCA |
| UNC119 | 9.05E-09 | 0.384669 | 0.754 | 0.695 | 0.000176 | CE | UNC119 |
| TPM1 | 1.15E-08 | 0.293092 | 0.674 | 0.565 | 0.000223 | CE | TPM1 |
| SAT12 | 1.38E-08 | 0.425627 | 0.917 | 0.902 | 0.000269 | CE | SAT1 |
| ID21 | 1.62E-08 | 0.37533 | 0.758 | 0.73 | 0.000315 | CE | ID2 |
| LYPD1 | 1.65E-08 | 0.439511 | 0.538 | 0.404 | 0.000322 | CE | LYPD1 |
| ITM2C | 2.85E-08 | 0.271396 | 0.795 | 0.782 | 0.000555 | CE | ITM2C |
| BTG12 | 3.28E-08 | 0.292337 | 0.924 | 0.876 | 0.000638 | CE | BTG1 |
| GNB3 | 4.08E-08 | 0.38463 | 0.314 | 0.197 | 0.000793 | CE | GNB3 |
| RHOC | 4.38E-08 | 0.278698 | 0.754 | 0.717 | 0.000852 | CE | RHOC |
| LGALS11 | 6.67E-08 | 0.309811 | 0.36 | 0.231 | 0.001298 | CE | LGALS1 |
| CRB2 | 6.86E-08 | 0.28996 | 0.489 | 0.352 | 0.001334 | CE | CRB2 |
| MAP1LC3A | 8.90E-08 | 0.281323 | 0.549 | 0.445 | 0.001732 | CE | MAP1LC3A |
| RSPO1 | 1.51E-07 | 0.356499 | 0.523 | 0.404 | 0.002943 | CE | RSPO1 |
| PON21 | 5.86E-07 | 0.255403 | 0.644 | 0.544 | 0.01141 | CE | PON2 |
| GJA1 | 6.08E-07 | 0.320508 | 0.473 | 0.362 | 0.011832 | CE | GJA1 |
| ASCL1 | 8.26E-07 | 0.477138 | 0.239 | 0.137 | 0.016078 | CE | ASCL1 |
| BST21 | 9.45E-06 | 0.313394 | 0.383 | 0.292 | 0.183923 | CE | BST2 |
| PLEKHA5 | 5.39E-05 | 0.258446 | 0.496 | 0.426 | 1 | CE | PLEKHA5 |
| HES42 | 0.000367 | 0.28748 | 0.769 | 0.767 | 1 | CE | HES4 |
| TRH | 0.005407 | 0.433144 | 0.674 | 0.675 | 1 | CE | TRH |
| MCIDAS | 3.19E-220 | 0.45137 | 0.615 | 0.004 | 6.21E-216 | MB1 | MCIDAS |
| STMND1 | 5.88E-209 | 0.769079 | 0.821 | 0.014 | 1.14E-204 | MB1 | STMND1 |
| CDC20B | 4.02E-192 | 1.984533 | 0.974 | 0.027 | 7.83E-188 | MB1 | CDC20B |
| LINC0151 | 2.33E-136 | 0.78111 | 0.795 | 0.026 | 4.53E-132 | MB1 | LINC01513 |
| TP53AIP1 | 1.40E-132 | 0.2594 | 0.41 | 0.004 | 2.72E-128 | MB1 | TP53AIP1 |
| FOXN4 | 2.35E-128 | 0.30723 | 0.513 | 0.009 | 4.57E-124 | MB1 | FOXN4 |
| RIBC2 | 6.81E-113 | 0.791377 | 0.718 | 0.027 | 1.33E-108 | MB1 | RIBC2 |
| C7orf57 | 3.49E-104 | 0.790467 | 0.872 | 0.048 | 6.80E-100 | MB1 | C7orf57 |

| | | | | | | | | |
|---|---|---|---|---|---|---|---|---|
| NEK10 | 3.01E-97 | 0.546194 | 0.795 | 0.04 | 5.86E-93 | MB1 | | NEK10 |
| LRRC26 | 4.57E-97 | 1.220754 | 0.923 | 0.062 | 8.89E-93 | MB1 | | LRRC26 |
| DNAH12 | 7.53E-97 | 0.542331 | 0.667 | 0.027 | 1.47E-92 | MB1 | | DNAH12 |
| 10-Mar | 2.88E-95 | 0.669984 | 0.821 | 0.046 | 5.60E-91 | MB1 | | 10-Mar |
| C11orf97 | 1.01E-93 | 0.892408 | 0.846 | 0.05 | 1.97E-89 | MB1 | | C11orf97 |
| CCDC39 | 3.45E-89 | 0.592701 | 0.692 | 0.033 | 6.71E-85 | MB1 | | CCDC39 |
| ZNF295-A | 5.80E-88 | 0.689642 | 0.487 | 0.015 | 1.13E-83 | MB1 | | ZNF295-AS1 |
| CFAP161 | 6.60E-85 | 0.410939 | 0.59 | 0.024 | 1.28E-80 | MB1 | | CFAP161 |
| TEKT1 | 5.30E-83 | 1.125017 | 0.949 | 0.081 | 1.03E-78 | MB1 | | TEKT1 |
| NEK21 | 1.66E-82 | 1.211167 | 1 | 0.093 | 3.24E-78 | MB1 | | NEK2 |
| FAM221B | 5.09E-81 | 0.429619 | 0.513 | 0.019 | 9.90E-77 | MB1 | | FAM221B |
| C20orf85 | 1.59E-80 | 1.24753 | 0.846 | 0.062 | 3.09E-76 | MB1 | | C20orf85 |
| LRRC46 | 2.18E-80 | 1.194875 | 0.897 | 0.074 | 4.25E-76 | MB1 | | LRRC46 |
| ROPN1L | 2.80E-79 | 1.753832 | 0.974 | 0.096 | 5.45E-75 | MB1 | | ROPN1L |
| CCDC65 | 5.20E-78 | 0.598175 | 0.744 | 0.047 | 1.01E-73 | MB1 | | CCDC65 |
| STOML3 | 1.40E-77 | 0.94571 | 0.795 | 0.053 | 2.72E-73 | MB1 | | STOML3 |
| DNAH9 | 8.00E-77 | 0.920913 | 0.923 | 0.081 | 1.56E-72 | MB1 | | DNAH9 |
| CFAP52 | 1.26E-76 | 0.904572 | 0.949 | 0.085 | 2.46E-72 | MB1 | | CFAP52 |
| KIF19 | 2.69E-76 | 0.916347 | 0.846 | 0.067 | 5.23E-72 | MB1 | | KIF19 |
| LRRC71 | 1.15E-75 | 0.346919 | 0.59 | 0.028 | 2.24E-71 | MB1 | | LRRC71 |
| C11orf88 | 5.55E-75 | 1.754036 | 0.949 | 0.095 | 1.08E-70 | MB1 | | C11orf88 |
| CFAP47 | 1.23E-74 | 0.261713 | 0.513 | 0.02 | 2.39E-70 | MB1 | | CFAP47 |
| TMEM31 | 1.89E-72 | 1.154415 | 0.821 | 0.065 | 3.69E-68 | MB1 | | TMEM31 |
| CCNO | 3.18E-71 | 3.123437 | 1 | 0.12 | 6.18E-67 | MB1 | | CCNO |
| TTC29 | 3.79E-70 | 0.467735 | 0.641 | 0.037 | 7.36E-66 | MB1 | | TTC29 |
| C11orf16 | 6.30E-70 | 0.267204 | 0.333 | 0.008 | 1.23E-65 | MB1 | | C11orf16 |
| UBXN10 | 6.96E-70 | 0.526371 | 0.769 | 0.058 | 1.35E-65 | MB1 | | UBXN10 |
| AKAP14 | 2.02E-69 | 0.76845 | 0.769 | 0.059 | 3.93E-65 | MB1 | | AKAP14 |
| PLK4 | 1.23E-66 | 0.626895 | 0.692 | 0.049 | 2.40E-62 | MB1 | | PLK4 |
| CCDC146 | 1.10E-65 | 0.53568 | 0.744 | 0.057 | 2.15E-61 | MB1 | | CCDC146 |
| SPAG17 | 2.25E-65 | 0.68081 | 0.821 | 0.072 | 4.38E-61 | MB1 | | SPAG17 |
| ADGB | 2.69E-65 | 0.402736 | 0.538 | 0.027 | 5.23E-61 | MB1 | | ADGB |
| RP11-295I | 1.68E-63 | 0.816542 | 0.846 | 0.08 | 3.26E-59 | MB1 | | RP11-295M3.4 |
| DNAAF1 | 1.06E-61 | 0.722816 | 0.821 | 0.076 | 2.06E-57 | MB1 | | DNAAF1 |
| LINC0048 | 1.90E-61 | 0.615199 | 0.538 | 0.031 | 3.70E-57 | MB1 | | LINC00488 |
| LRP2BP | 3.25E-59 | 0.787583 | 0.744 | 0.067 | 6.33E-55 | MB1 | | LRP2BP |
| CCDC170 | 3.40E-58 | 0.83589 | 0.795 | 0.077 | 6.62E-54 | MB1 | | CCDC170 |
| FAM166B | 6.51E-58 | 0.937368 | 0.821 | 0.086 | 1.27E-53 | MB1 | | FAM166B |
| WDR63 | 1.73E-57 | 0.425129 | 0.692 | 0.055 | 3.37E-53 | MB1 | | WDR63 |
| CFAP45 | 2.61E-57 | 1.12626 | 0.949 | 0.128 | 5.07E-53 | MB1 | | CFAP45 |

| | | | | | | | |
|---|---|---|---|---|---|---|---|
| PTPRC | 1.33E-56 | 0.324959 | 0.436 | 0.02 | 2.58E-52 | MB1 | PTPRC |
| C4orf22 | 5.20E-56 | 0.405911 | 0.487 | 0.027 | 1.01E-51 | MB1 | C4orf22 |
| TEKT4 | 1.55E-55 | 0.290325 | 0.41 | 0.018 | 3.01E-51 | MB1 | TEKT4 |
| SPATA18 | 2.30E-54 | 0.622961 | 0.667 | 0.056 | 4.48E-50 | MB1 | SPATA18 |
| USP13 | 3.57E-54 | 1.272062 | 0.949 | 0.145 | 6.95E-50 | MB1 | USP13 |
| CERKL | 3.81E-54 | 0.302153 | 0.41 | 0.019 | 7.40E-50 | MB1 | CERKL |
| KATNAL2 | 6.38E-54 | 0.4598 | 0.744 | 0.069 | 1.24E-49 | MB1 | KATNAL2 |
| E2F7 | 1.78E-53 | 0.525376 | 0.59 | 0.043 | 3.46E-49 | MB1 | E2F7 |
| CCNA1 | 3.41E-53 | 0.973875 | 0.821 | 0.094 | 6.63E-49 | MB1 | CCNA1 |
| ZNF474 | 1.01E-52 | 0.336083 | 0.513 | 0.032 | 1.96E-48 | MB1 | ZNF474 |
| C22orf42 | 4.00E-51 | 0.422823 | 0.41 | 0.02 | 7.79E-47 | MB1 | C22orf42 |
| C9orf135 | 1.16E-50 | 0.944587 | 0.692 | 0.069 | 2.25E-46 | MB1 | C9orf135 |
| DAW1 | 1.24E-50 | 0.324109 | 0.487 | 0.03 | 2.42E-46 | MB1 | DAW1 |
| TCTEX1D1 | 1.65E-50 | 0.945356 | 0.923 | 0.13 | 3.20E-46 | MB1 | TCTEX1D1 |
| FAM216B | 2.17E-50 | 0.76072 | 0.564 | 0.042 | 4.22E-46 | MB1 | FAM216B |
| MUC12 | 2.67E-50 | 0.800903 | 0.718 | 0.073 | 5.20E-46 | MB1 | MUC12 |
| DNAAF3 | 3.10E-50 | 1.078766 | 0.949 | 0.156 | 6.02E-46 | MB1 | DNAAF3 |
| ZMYND10 | 4.27E-50 | 1.928903 | 1 | 0.195 | 8.31E-46 | MB1 | ZMYND10 |
| EFCAB1 | 7.81E-50 | 1.158467 | 0.974 | 0.165 | 1.52E-45 | MB1 | EFCAB1 |
| WDR38 | 2.25E-49 | 0.637606 | 0.667 | 0.062 | 4.38E-45 | MB1 | WDR38 |
| TEX26 | 2.80E-49 | 0.671107 | 0.564 | 0.045 | 5.44E-45 | MB1 | TEX26 |
| CFAP100 | 3.29E-49 | 0.29023 | 0.462 | 0.027 | 6.41E-45 | MB1 | CFAP100 |
| BBOF1 | 6.86E-49 | 0.799897 | 0.974 | 0.156 | 1.33E-44 | MB1 | BBOF1 |
| STIL | 8.31E-49 | 0.388169 | 0.538 | 0.039 | 1.62E-44 | MB1 | STIL |
| FAM92B | 9.01E-49 | 0.875867 | 0.897 | 0.124 | 1.75E-44 | MB1 | FAM92B |
| SPEF1 | 1.21E-48 | 0.974022 | 0.923 | 0.145 | 2.35E-44 | MB1 | SPEF1 |
| AK7 | 1.65E-48 | 0.407113 | 0.692 | 0.066 | 3.21E-44 | MB1 | AK7 |
| PPP1R32 | 8.72E-48 | 0.757447 | 0.846 | 0.111 | 1.70E-43 | MB1 | PPP1R32 |
| CEP152 | 1.64E-47 | 0.35211 | 0.564 | 0.044 | 3.19E-43 | MB1 | CEP152 |
| DEPDC1 | 2.10E-47 | 0.501531 | 0.615 | 0.055 | 4.09E-43 | MB1 | DEPDC1 |
| SPAG6 | 2.90E-47 | 1.217836 | 0.974 | 0.174 | 5.64E-43 | MB1 | SPAG6 |
| CFAP65 | 2.99E-47 | 0.476809 | 0.744 | 0.081 | 5.81E-43 | MB1 | CFAP65 |
| MELK1 | 7.25E-47 | 0.704635 | 0.821 | 0.103 | 1.41E-42 | MB1 | MELK |
| FAM183A | 8.84E-47 | 1.701825 | 1 | 0.198 | 1.72E-42 | MB1 | FAM183A |
| CFAP74 | 2.29E-46 | 0.402736 | 0.615 | 0.055 | 4.46E-42 | MB1 | CFAP74 |
| CCDC78 | 3.30E-46 | 0.539364 | 0.59 | 0.05 | 6.42E-42 | MB1 | CCDC78 |
| ENKUR | 6.21E-46 | 0.531808 | 0.615 | 0.056 | 1.21E-41 | MB1 | ENKUR |
| SGOL21 | 6.64E-46 | 0.530403 | 0.769 | 0.087 | 1.29E-41 | MB1 | SGOL2 |
| ANKRD66 | 1.15E-45 | 0.570596 | 0.513 | 0.039 | 2.24E-41 | MB1 | ANKRD66 |
| DNAH11 | 1.39E-45 | 0.398376 | 0.59 | 0.05 | 2.70E-41 | MB1 | DNAH11 |

| | | | | | | | |
|---|---|---|---|---|---|---|---|
| RP11-834( | 4.80E-45 | 0.398062 | 0.462 | 0.03 | 9.33E-41 | MB1 | RP11-834C11.4 |
| MAP3K19 | 6.03E-45 | 0.492087 | 0.564 | 0.048 | 1.17E-40 | MB1 | MAP3K19 |
| TSNAXIP1 | 8.50E-45 | 0.437836 | 0.641 | 0.063 | 1.65E-40 | MB1 | TSNAXIP1 |
| C6orf118 | 7.60E-44 | 0.906502 | 0.897 | 0.147 | 1.48E-39 | MB1 | C6orf118 |
| RAMP1 | 1.79E-43 | 0.400428 | 0.538 | 0.043 | 3.48E-39 | MB1 | RAMP1 |
| RSPH4A | 1.92E-43 | 0.729585 | 0.692 | 0.08 | 3.73E-39 | MB1 | RSPH4A |
| RP11-542( | 4.88E-43 | 0.256687 | 0.154 | 0.002 | 9.49E-39 | MB1 | RP11-542G1.1 |
| CDC201 | 5.69E-43 | 1.217197 | 0.974 | 0.175 | 1.11E-38 | MB1 | CDC20 |
| CDK11 | 1.63E-42 | 1.982993 | 1 | 0.232 | 3.17E-38 | MB1 | CDK1 |
| RP11-356I | 1.79E-42 | 1.188845 | 0.718 | 0.085 | 3.49E-38 | MB1 | RP11-356K23.1 |
| C2orf50 | 1.85E-42 | 0.347677 | 0.59 | 0.054 | 3.60E-38 | MB1 | C2orf50 |
| CCDC96 | 4.83E-42 | 0.485364 | 0.692 | 0.081 | 9.39E-38 | MB1 | CCDC96 |
| LRRC6 | 1.77E-41 | 0.370127 | 0.641 | 0.066 | 3.44E-37 | MB1 | LRRC6 |
| HIST1H2B | 2.50E-41 | 0.61585 | 0.667 | 0.074 | 4.87E-37 | MB1 | HIST1H2BJ |
| SPAG1 | 2.52E-41 | 0.600514 | 0.744 | 0.098 | 4.89E-37 | MB1 | SPAG1 |
| CAPSL | 2.76E-41 | 1.168913 | 0.872 | 0.149 | 5.37E-37 | MB1 | CAPSL |
| ERICH2 | 6.73E-41 | 0.26362 | 0.436 | 0.03 | 1.31E-36 | MB1 | ERICH2 |
| DYX1C1 | 1.37E-40 | 1.011961 | 1 | 0.246 | 2.66E-36 | MB1 | DYX1C1 |
| DNAI1 | 1.87E-40 | 0.358468 | 0.487 | 0.039 | 3.64E-36 | MB1 | DNAI1 |
| TSGA10 | 2.52E-40 | 0.650134 | 0.744 | 0.1 | 4.90E-36 | MB1 | TSGA10 |
| CFAP221 | 3.60E-40 | 0.489438 | 0.744 | 0.097 | 7.01E-36 | MB1 | CFAP221 |
| WDR49 | 4.07E-40 | 0.367553 | 0.513 | 0.043 | 7.92E-36 | MB1 | WDR49 |
| LINC0064: | 5.13E-40 | 0.345173 | 0.538 | 0.048 | 9.98E-36 | MB1 | LINC00643 |
| CFAP77 | 8.26E-40 | 0.968222 | 0.949 | 0.187 | 1.61E-35 | MB1 | CFAP77 |
| DYNLRB2 | 2.82E-39 | 0.872785 | 0.821 | 0.132 | 5.49E-35 | MB1 | DYNLRB2 |
| C11orf70 | 3.02E-39 | 0.552327 | 0.462 | 0.037 | 5.88E-35 | MB1 | C11orf70 |
| SPATA17 | 3.13E-39 | 0.628102 | 0.795 | 0.111 | 6.09E-35 | MB1 | SPATA17 |
| CCDC180 | 6.04E-39 | 0.437221 | 0.641 | 0.072 | 1.18E-34 | MB1 | CCDC180 |
| RIBC1 | 7.62E-39 | 0.367377 | 0.615 | 0.066 | 1.48E-34 | MB1 | RIBC1 |
| CFAP206 | 9.77E-39 | 0.3465 | 0.436 | 0.033 | 1.90E-34 | MB1 | CFAP206 |
| DLGAP51 | 1.42E-38 | 0.583971 | 0.667 | 0.079 | 2.77E-34 | MB1 | DLGAP5 |
| SLC32A11 | 1.46E-38 | 0.314403 | 0.487 | 0.039 | 2.84E-34 | MB1 | SLC32A1 |
| RSPH1 | 1.63E-38 | 1.392213 | 0.949 | 0.214 | 3.17E-34 | MB1 | RSPH1 |
| GSTA3 | 1.83E-38 | 0.500502 | 0.436 | 0.033 | 3.57E-34 | MB1 | GSTA3 |
| CFAP57 | 2.06E-38 | 0.31312 | 0.41 | 0.028 | 4.02E-34 | MB1 | CFAP57 |
| AK9 | 3.44E-38 | 0.567296 | 0.769 | 0.109 | 6.69E-34 | MB1 | AK9 |
| CFAP73 | 8.00E-38 | 0.490848 | 0.462 | 0.038 | 1.56E-33 | MB1 | CFAP73 |
| MAL2 | 1.38E-37 | 0.647995 | 0.692 | 0.09 | 2.69E-33 | MB1 | MAL2 |
| RUNDC3B | 1.61E-37 | 0.359481 | 0.564 | 0.055 | 3.14E-33 | MB1 | RUNDC3B |
| NUP62CL | 2.80E-37 | 0.713064 | 0.846 | 0.147 | 5.44E-33 | MB1 | NUP62CL |

| | | | | | | | |
|---|---|---|---|---|---|---|---|
| NKX6-1 | 2.90E-37 | 0.292411 | 0.308 | 0.016 | 5.65E-33 | MB1 | NKX6-1 |
| CCDC171 | 4.44E-37 | 0.516092 | 0.667 | 0.083 | 8.63E-33 | MB1 | CCDC171 |
| C5orf49 | 1.10E-36 | 1.926369 | 1 | 0.319 | 2.14E-32 | MB1 | C5orf49 |
| RSPH9 | 1.14E-36 | 1.207366 | 0.974 | 0.246 | 2.23E-32 | MB1 | RSPH9 |
| PRR29 | 1.38E-36 | 0.453909 | 0.564 | 0.059 | 2.69E-32 | MB1 | PRR29 |
| TTLL10 | 1.62E-36 | 0.380507 | 0.564 | 0.059 | 3.15E-32 | MB1 | TTLL10 |
| ANLN | 1.86E-36 | 0.362556 | 0.538 | 0.052 | 3.62E-32 | MB1 | ANLN |
| C1orf194 | 1.88E-36 | 1.088977 | 0.872 | 0.17 | 3.66E-32 | MB1 | C1orf194 |
| DRC1 | 2.48E-36 | 0.685678 | 0.872 | 0.15 | 4.83E-32 | MB1 | DRC1 |
| C4orf47 | 2.53E-36 | 0.565679 | 0.744 | 0.11 | 4.93E-32 | MB1 | C4orf47 |
| SGOL11 | 2.98E-36 | 0.472878 | 0.718 | 0.093 | 5.80E-32 | MB1 | SGOL1 |
| RP11-263I | 3.02E-36 | 1.159849 | 0.897 | 0.203 | 5.87E-32 | MB1 | RP11-263K19.4 |
| CCNB11 | 3.71E-36 | 0.804005 | 0.974 | 0.203 | 7.22E-32 | MB1 | CCNB1 |
| CFAP46 | 7.55E-36 | 0.652236 | 0.795 | 0.132 | 1.47E-31 | MB1 | CFAP46 |
| ARMC3 | 8.27E-36 | 0.725194 | 0.769 | 0.122 | 1.61E-31 | MB1 | ARMC3 |
| FANK1 | 9.98E-36 | 0.914195 | 0.846 | 0.165 | 1.94E-31 | MB1 | FANK1 |
| CASC1 | 1.58E-35 | 0.382896 | 0.59 | 0.066 | 3.07E-31 | MB1 | CASC1 |
| CCDC173 | 1.65E-35 | 0.866371 | 0.872 | 0.167 | 3.21E-31 | MB1 | CCDC173 |
| CCDC113 | 1.73E-35 | 0.533561 | 0.795 | 0.124 | 3.37E-31 | MB1 | CCDC113 |
| C9orf24 | 3.05E-35 | 1.919505 | 1 | 0.305 | 5.94E-31 | MB1 | C9orf24 |
| CFAP126 | 5.66E-35 | 1.836642 | 1 | 0.335 | 1.10E-30 | MB1 | CFAP126 |
| C5AR1 | 6.60E-35 | 0.339375 | 0.513 | 0.05 | 1.28E-30 | MB1 | C5AR1 |
| CCDC74A | 8.08E-35 | 1.254747 | 0.974 | 0.303 | 1.57E-30 | MB1 | CCDC74A |
| KDELC2 | 8.63E-35 | 0.433857 | 0.615 | 0.075 | 1.68E-30 | MB1 | KDELC2 |
| TSPAN2 | 1.23E-34 | 0.357705 | 0.564 | 0.06 | 2.39E-30 | MB1 | TSPAN2 |
| LKAAEAR1 | 1.65E-34 | 0.419698 | 0.538 | 0.058 | 3.21E-30 | MB1 | LKAAEAR1 |
| SPEF2 | 1.86E-34 | 0.615097 | 0.769 | 0.126 | 3.62E-30 | MB1 | SPEF2 |
| LRRC10B | 3.24E-34 | 0.358002 | 0.615 | 0.074 | 6.31E-30 | MB1 | LRRC10B |
| SPA17 | 4.43E-34 | 0.80844 | 0.923 | 0.204 | 8.63E-30 | MB1 | SPA17 |
| STON2 | 9.21E-34 | 0.426603 | 0.59 | 0.071 | 1.79E-29 | MB1 | STON2 |
| PACRG | 1.07E-33 | 0.739069 | 0.872 | 0.174 | 2.08E-29 | MB1 | PACRG |
| FOXJ1 | 1.80E-33 | 2.137904 | 1 | 0.373 | 3.51E-29 | MB1 | FOXJ1 |
| C22orf15 | 4.13E-33 | 0.628803 | 0.744 | 0.115 | 8.04E-29 | MB1 | C22orf15 |
| FAM188B | 5.55E-33 | 0.270414 | 0.462 | 0.042 | 1.08E-28 | MB1 | FAM188B |
| C9orf116 | 5.97E-33 | 1.395086 | 1 | 0.343 | 1.16E-28 | MB1 | C9orf116 |
| NPHP1 | 7.65E-33 | 0.699378 | 0.872 | 0.185 | 1.49E-28 | MB1 | NPHP1 |
| CDS1 | 1.20E-32 | 0.619153 | 0.692 | 0.107 | 2.33E-28 | MB1 | CDS1 |
| DDAH1 | 1.28E-32 | 0.748542 | 0.872 | 0.183 | 2.49E-28 | MB1 | DDAH1 |
| RIIAD1 | 1.35E-32 | 0.577714 | 0.667 | 0.097 | 2.64E-28 | MB1 | RIIAD1 |
| ABLIM1 | 1.59E-32 | 0.448664 | 0.744 | 0.11 | 3.08E-28 | MB1 | ABLIM1 |

| | | | | | | | |
|---|---|---|---|---|---|---|---|
| KIF9 | 2.06E-32 | 1.616219 | 1 | 0.374 | 4.01E-28 | MB1 | KIF9 |
| CENPF1 | 2.16E-32 | 0.759938 | 0.897 | 0.17 | 4.20E-28 | MB1 | CENPF |
| IQCG | 2.44E-32 | 0.65124 | 0.872 | 0.176 | 4.74E-28 | MB1 | IQCG |
| TMEM231 | 5.02E-32 | 0.904191 | 0.923 | 0.253 | 9.77E-28 | MB1 | TMEM231 |
| OSCP1 | 5.09E-32 | 0.989525 | 0.923 | 0.236 | 9.91E-28 | MB1 | OSCP1 |
| NME5 | 5.79E-32 | 0.884811 | 0.846 | 0.18 | 1.13E-27 | MB1 | NME5 |
| SKA31 | 7.74E-32 | 0.37471 | 0.564 | 0.067 | 1.51E-27 | MB1 | SKA3 |
| C10orf107 | 9.29E-32 | 0.351232 | 0.513 | 0.056 | 1.81E-27 | MB1 | C10orf107 |
| TMEM232 | 2.44E-31 | 0.331946 | 0.436 | 0.041 | 4.74E-27 | MB1 | TMEM232 |
| CCDC157 | 2.91E-31 | 0.635038 | 0.795 | 0.153 | 5.65E-27 | MB1 | CCDC157 |
| PPP1R42 | 3.23E-31 | 0.417111 | 0.59 | 0.077 | 6.28E-27 | MB1 | PPP1R42 |
| LINC0109 | 5.56E-31 | 0.40195 | 0.282 | 0.016 | 1.08E-26 | MB1 | LINC01091 |
| FAM81B | 1.24E-30 | 0.621851 | 0.821 | 0.154 | 2.41E-26 | MB1 | FAM81B |
| B9D2 | 1.25E-30 | 0.666123 | 0.872 | 0.196 | 2.44E-26 | MB1 | B9D2 |
| ODF3B | 1.38E-30 | 0.97323 | 1 | 0.275 | 2.68E-26 | MB1 | ODF3B |
| WDR66 | 1.76E-30 | 0.360821 | 0.513 | 0.058 | 3.42E-26 | MB1 | WDR66 |
| BUB11 | 2.08E-30 | 0.363557 | 0.513 | 0.058 | 4.05E-26 | MB1 | BUB1 |
| GRAMD3 | 2.81E-30 | 0.667638 | 0.641 | 0.096 | 5.47E-26 | MB1 | GRAMD3 |
| LRWD1 | 4.96E-30 | 0.979707 | 0.872 | 0.216 | 9.65E-26 | MB1 | LRWD1 |
| AC007405 | 5.65E-30 | 0.635687 | 0.769 | 0.146 | 1.10E-25 | MB1 | AC007405.6 |
| RPGR | 5.93E-30 | 0.618445 | 0.872 | 0.184 | 1.15E-25 | MB1 | RPGR |
| ULK4 | 8.20E-30 | 0.38261 | 0.538 | 0.065 | 1.59E-25 | MB1 | ULK4 |
| TPPP3 | 1.23E-29 | 2.898846 | 1 | 0.486 | 2.39E-25 | MB1 | TPPP3 |
| NUF21 | 3.41E-29 | 0.438262 | 0.744 | 0.119 | 6.64E-25 | MB1 | NUF2 |
| RBKS | 4.27E-29 | 0.496308 | 0.692 | 0.11 | 8.31E-25 | MB1 | RBKS |
| UBE2C1 | 8.58E-29 | 0.893445 | 0.974 | 0.249 | 1.67E-24 | MB1 | UBE2C |
| SYT51 | 1.44E-28 | 0.621545 | 0.769 | 0.14 | 2.80E-24 | MB1 | SYT5 |
| CENPK1 | 1.78E-28 | 0.685264 | 0.846 | 0.18 | 3.46E-24 | MB1 | CENPK |
| PIH1D2 | 1.81E-28 | 0.297289 | 0.487 | 0.054 | 3.52E-24 | MB1 | PIH1D2 |
| NRAV | 2.39E-28 | 0.444654 | 0.641 | 0.096 | 4.65E-24 | MB1 | NRAV |
| CCDC341 | 2.57E-28 | 1.162627 | 1 | 0.403 | 4.99E-24 | MB1 | CCDC34 |
| CEP120 | 3.19E-28 | 0.307531 | 0.462 | 0.05 | 6.21E-24 | MB1 | CEP120 |
| TOP2A1 | 3.92E-28 | 0.70743 | 0.846 | 0.162 | 7.62E-24 | MB1 | TOP2A |
| SPAG8 | 4.74E-28 | 0.608886 | 0.744 | 0.138 | 9.22E-24 | MB1 | SPAG8 |
| TEKT2 | 5.83E-28 | 1.057686 | 0.872 | 0.225 | 1.13E-23 | MB1 | TEKT2 |
| IFT22 | 1.77E-27 | 1.445138 | 1 | 0.637 | 3.43E-23 | MB1 | IFT22 |
| CCDC138 | 2.07E-27 | 0.480889 | 0.718 | 0.127 | 4.04E-23 | MB1 | CCDC138 |
| IQCD | 5.23E-27 | 0.41026 | 0.692 | 0.119 | 1.02E-22 | MB1 | IQCD |
| MORN2 | 8.94E-27 | 1.145576 | 1 | 0.542 | 1.74E-22 | MB1 | MORN2 |
| ZDHHC1 | 9.04E-27 | 0.413478 | 0.59 | 0.087 | 1.76E-22 | MB1 | ZDHHC1 |

| | | | | | | | |
|---|---|---|---|---|---|---|---|
| AC079922 | 9.56E-27 | 0.30532 | 0.462 | 0.053 | 1.86E-22 | MB1 | AC079922.3 |
| MORN5 | 1.10E-26 | 0.699826 | 0.795 | 0.178 | 2.13E-22 | MB1 | MORN5 |
| PIFO | 1.44E-26 | 1.412762 | 1 | 0.468 | 2.81E-22 | MB1 | PIFO |
| C1orf158 | 2.00E-26 | 0.382618 | 0.462 | 0.055 | 3.89E-22 | MB1 | C1orf158 |
| TSPAN19 | 2.87E-26 | 0.476327 | 0.59 | 0.091 | 5.59E-22 | MB1 | TSPAN19 |
| CEP126 | 2.92E-26 | 0.487984 | 0.769 | 0.152 | 5.67E-22 | MB1 | CEP126 |
| LDLRAD4 | 2.97E-26 | 0.294641 | 0.385 | 0.038 | 5.77E-22 | MB1 | LDLRAD4 |
| DNALI1 | 3.51E-26 | 1.176368 | 1 | 0.492 | 6.84E-22 | MB1 | DNALI1 |
| PSENEN | 4.92E-26 | 1.68484 | 1 | 0.748 | 9.57E-22 | MB1 | PSENEN |
| RP11-368I | 5.83E-26 | 0.260318 | 0.385 | 0.038 | 1.13E-21 | MB1 | RP11-368I7.4 |
| RUVBL2 | 8.07E-26 | 1.412316 | 1 | 0.68 | 1.57E-21 | MB1 | RUVBL2 |
| HSPBP1 | 1.01E-25 | 1.172455 | 0.974 | 0.521 | 1.96E-21 | MB1 | HSPBP1 |
| EVL | 1.31E-25 | 1.51965 | 0.949 | 0.412 | 2.55E-21 | MB1 | EVL |
| CFAP43 | 1.45E-25 | 0.323545 | 0.436 | 0.049 | 2.82E-21 | MB1 | CFAP43 |
| DYNLL1 | 1.47E-25 | 1.44216 | 1 | 0.975 | 2.86E-21 | MB1 | DYNLL1 |
| ARL6 | 2.03E-25 | 0.411706 | 0.59 | 0.091 | 3.96E-21 | MB1 | ARL6 |
| TTC26 | 2.22E-25 | 0.500422 | 0.641 | 0.109 | 4.31E-21 | MB1 | TTC26 |
| PTGES3 | 2.55E-25 | 1.467345 | 1 | 0.971 | 4.96E-21 | MB1 | PTGES3 |
| CCDC60 | 2.92E-25 | 0.386802 | 0.436 | 0.051 | 5.67E-21 | MB1 | CCDC60 |
| CENPW1 | 2.92E-25 | 0.769139 | 0.949 | 0.275 | 5.68E-21 | MB1 | CENPW |
| VRK3 | 2.95E-25 | 0.801815 | 0.974 | 0.377 | 5.74E-21 | MB1 | VRK3 |
| SPATA33 | 3.94E-25 | 0.971312 | 0.949 | 0.339 | 7.66E-21 | MB1 | SPATA33 |
| RAB36 | 4.33E-25 | 0.458947 | 0.615 | 0.105 | 8.42E-21 | MB1 | RAB36 |
| BBS12 | 4.41E-25 | 0.37243 | 0.564 | 0.085 | 8.59E-21 | MB1 | BBS12 |
| C14orf79 | 4.87E-25 | 0.532816 | 0.692 | 0.139 | 9.47E-21 | MB1 | C14orf79 |
| PLK11 | 6.12E-25 | 0.587556 | 0.641 | 0.108 | 1.19E-20 | MB1 | PLK1 |
| AK8 | 6.33E-25 | 0.34621 | 0.436 | 0.052 | 1.23E-20 | MB1 | AK8 |
| CEP135 | 6.34E-25 | 0.341643 | 0.59 | 0.091 | 1.23E-20 | MB1 | CEP135 |
| ENO4 | 6.44E-25 | 0.399865 | 0.59 | 0.092 | 1.25E-20 | MB1 | ENO4 |
| WDR93 | 7.32E-25 | 0.28649 | 0.462 | 0.057 | 1.42E-20 | MB1 | WDR93 |
| CBY1 | 7.32E-25 | 1.198555 | 0.974 | 0.519 | 1.42E-20 | MB1 | CBY1 |
| WDR78 | 8.07E-25 | 0.402541 | 0.538 | 0.079 | 1.57E-20 | MB1 | WDR78 |
| CCDC30 | 9.92E-25 | 0.45895 | 0.769 | 0.164 | 1.93E-20 | MB1 | CCDC30 |
| RP11-620. | 1.18E-24 | 1.398783 | 1 | 0.586 | 2.30E-20 | MB1 | RP11-620J15.3 |
| IQCC | 1.36E-24 | 0.302269 | 0.513 | 0.071 | 2.65E-20 | MB1 | IQCC |
| PIM2 | 1.71E-24 | 0.724963 | 0.897 | 0.254 | 3.33E-20 | MB1 | PIM2 |
| SYT10 | 1.75E-24 | 0.432003 | 0.487 | 0.065 | 3.40E-20 | MB1 | SYT10 |
| FBXW9 | 1.76E-24 | 0.819994 | 0.949 | 0.336 | 3.42E-20 | MB1 | FBXW9 |
| P4HTM | 3.60E-24 | 1.158929 | 0.974 | 0.519 | 7.01E-20 | MB1 | P4HTM |
| SYDE2 | 1.04E-23 | 0.391967 | 0.59 | 0.098 | 2.03E-19 | MB1 | SYDE2 |

| | | | | | | | |
|---|---|---|---|---|---|---|---|
| CDK2AP21 | 1.06E-23 | 1.890098 | 0.949 | 0.515 | 2.06E-19 | MB1 | CDK2AP2 |
| EFHC1 | 1.20E-23 | 1.007983 | 0.974 | 0.435 | 2.33E-19 | MB1 | EFHC1 |
| FUZ | 1.45E-23 | 0.889909 | 0.897 | 0.308 | 2.82E-19 | MB1 | FUZ |
| AGPAT2 | 1.55E-23 | 0.811865 | 0.974 | 0.395 | 3.01E-19 | MB1 | AGPAT2 |
| ODF2L | 2.16E-23 | 0.979138 | 0.923 | 0.36 | 4.21E-19 | MB1 | ODF2L |
| CKAP2L | 2.51E-23 | 0.308942 | 0.41 | 0.048 | 4.88E-19 | MB1 | CKAP2L |
| C9orf9 | 2.52E-23 | 1.010953 | 0.872 | 0.276 | 4.90E-19 | MB1 | C9orf9 |
| C17orf97 | 2.77E-23 | 0.476739 | 0.718 | 0.147 | 5.38E-19 | MB1 | C17orf97 |
| TMEM14B | 3.18E-23 | 1.037823 | 1 | 0.829 | 6.19E-19 | MB1 | TMEM14B |
| BAIAP3 | 3.25E-23 | 0.448479 | 0.744 | 0.16 | 6.32E-19 | MB1 | BAIAP3 |
| C21orf58 | 3.58E-23 | 0.397514 | 0.718 | 0.147 | 6.97E-19 | MB1 | C21orf58 |
| RP11-128I | 3.91E-23 | 0.412653 | 0.462 | 0.063 | 7.61E-19 | MB1 | RP11-128M1.1 |
| PPIF1 | 5.03E-23 | 0.884327 | 0.949 | 0.337 | 9.78E-19 | MB1 | PPIF |
| POC1A1 | 5.34E-23 | 0.383269 | 0.641 | 0.114 | 1.04E-18 | MB1 | POC1A |
| DUSP18 | 6.88E-23 | 0.425689 | 0.744 | 0.161 | 1.34E-18 | MB1 | DUSP18 |
| WRAP53 | 8.94E-23 | 0.585135 | 0.846 | 0.241 | 1.74E-18 | MB1 | WRAP53 |
| LRRIQ1 | 9.32E-23 | 0.591762 | 0.718 | 0.169 | 1.81E-18 | MB1 | LRRIQ1 |
| TTC25 | 1.17E-22 | 0.321418 | 0.59 | 0.1 | 2.27E-18 | MB1 | TTC25 |
| FAM47E | 1.47E-22 | 0.359728 | 0.538 | 0.087 | 2.86E-18 | MB1 | FAM47E |
| TUBB4B1 | 1.69E-22 | 1.706094 | 0.974 | 0.846 | 3.28E-18 | MB1 | TUBB4B |
| CEP164 | 2.05E-22 | 0.564171 | 0.769 | 0.192 | 3.99E-18 | MB1 | CEP164 |
| GALNS | 2.60E-22 | 0.433312 | 0.692 | 0.146 | 5.06E-18 | MB1 | GALNS |
| LRRC73 | 3.01E-22 | 0.655603 | 0.897 | 0.262 | 5.85E-18 | MB1 | LRRC73 |
| CNIH2 | 4.00E-22 | 0.432642 | 0.667 | 0.131 | 7.78E-18 | MB1 | CNIH2 |
| CCDC181 | 4.41E-22 | 0.742516 | 0.923 | 0.326 | 8.57E-18 | MB1 | CCDC181 |
| NUDC | 4.62E-22 | 1.153811 | 1 | 0.892 | 8.99E-18 | MB1 | NUDC |
| CCNB21 | 8.00E-22 | 0.559382 | 0.769 | 0.173 | 1.56E-17 | MB1 | CCNB2 |
| TCTEX1D2 | 8.21E-22 | 1.086208 | 1 | 0.68 | 1.60E-17 | MB1 | TCTEX1D2 |
| HYDIN | 8.97E-22 | 0.343651 | 0.59 | 0.1 | 1.75E-17 | MB1 | HYDIN |
| SLF11 | 1.35E-21 | 0.691753 | 0.872 | 0.245 | 2.62E-17 | MB1 | SLF1 |
| CKS1B1 | 1.42E-21 | 1.14731 | 1 | 0.542 | 2.75E-17 | MB1 | CKS1B |
| IQCK | 2.03E-21 | 0.797775 | 0.949 | 0.353 | 3.94E-17 | MB1 | IQCK |
| WDR92 | 2.20E-21 | 0.349221 | 0.667 | 0.133 | 4.29E-17 | MB1 | WDR92 |
| LRRC23 | 2.76E-21 | 0.868634 | 0.923 | 0.34 | 5.38E-17 | MB1 | LRRC23 |
| STOX1 | 2.96E-21 | 0.359359 | 0.564 | 0.1 | 5.75E-17 | MB1 | STOX1 |
| IFT57 | 3.32E-21 | 0.858962 | 0.974 | 0.545 | 6.46E-17 | MB1 | IFT57 |
| SMC41 | 3.34E-21 | 0.625617 | 0.923 | 0.284 | 6.50E-17 | MB1 | SMC4 |
| UBXN11 | 3.72E-21 | 0.81556 | 0.897 | 0.353 | 7.24E-17 | MB1 | UBXN11 |
| C21orf62 | 4.09E-21 | 0.516037 | 0.718 | 0.155 | 7.95E-17 | MB1 | C21orf62 |
| ASF1B1 | 4.66E-21 | 0.332074 | 0.59 | 0.101 | 9.06E-17 | MB1 | ASF1B |

| | | | | | | | |
|---|---|---|---|---|---|---|---|
| GK | 5.27E-21 | 0.427428 | 0.641 | 0.134 | 1.03E-16 | MB1 | GK |
| DRC3 | 5.40E-21 | 0.350144 | 0.538 | 0.089 | 1.05E-16 | MB1 | DRC3 |
| TMEM67 | 5.74E-21 | 0.61037 | 0.846 | 0.243 | 1.12E-16 | MB1 | TMEM67 |
| CETN2 | 5.81E-21 | 1.063732 | 1 | 0.84 | 1.13E-16 | MB1 | CETN2 |
| DPCD | 7.28E-21 | 0.994382 | 1 | 0.644 | 1.42E-16 | MB1 | DPCD |
| PPIL6 | 9.62E-21 | 0.754171 | 0.897 | 0.273 | 1.87E-16 | MB1 | PPIL6 |
| MNS1 | 1.22E-20 | 0.272357 | 0.282 | 0.026 | 2.38E-16 | MB1 | MNS1 |
| MLF1 | 1.28E-20 | 1.000657 | 1 | 0.759 | 2.49E-16 | MB1 | MLF1 |
| AKR7A2 | 1.83E-20 | 0.99517 | 1 | 0.765 | 3.57E-16 | MB1 | AKR7A2 |
| PLCE1 | 1.86E-20 | 0.263706 | 0.487 | 0.075 | 3.62E-16 | MB1 | PLCE1 |
| BIRC51 | 2.94E-20 | 0.570354 | 0.821 | 0.196 | 5.72E-16 | MB1 | BIRC5 |
| CNTRL | 3.05E-20 | 0.350329 | 0.615 | 0.119 | 5.93E-16 | MB1 | CNTRL |
| TCTEX1D4 | 3.43E-20 | 0.381002 | 0.538 | 0.092 | 6.67E-16 | MB1 | TCTEX1D4 |
| DYNC2H1 | 3.73E-20 | 0.440313 | 0.692 | 0.157 | 7.26E-16 | MB1 | DYNC2H1 |
| C14orf142 | 4.88E-20 | 0.85097 | 0.846 | 0.303 | 9.50E-16 | MB1 | C14orf142 |
| SMKR1 | 5.47E-20 | 0.25261 | 0.462 | 0.068 | 1.06E-15 | MB1 | SMKR1 |
| BCO2 | 8.24E-20 | 0.299584 | 0.538 | 0.095 | 1.60E-15 | MB1 | BCO2 |
| RCAN3 | 8.46E-20 | 0.406723 | 0.59 | 0.114 | 1.65E-15 | MB1 | RCAN3 |
| HIST1H2B | 8.62E-20 | 0.555673 | 0.41 | 0.058 | 1.68E-15 | MB1 | HIST1H2BC |
| MDH1B | 8.99E-20 | 0.329342 | 0.564 | 0.104 | 1.75E-15 | MB1 | MDH1B |
| RFX2 | 9.13E-20 | 0.504499 | 0.718 | 0.186 | 1.78E-15 | MB1 | RFX2 |
| ANKRD42 | 9.64E-20 | 0.393122 | 0.667 | 0.14 | 1.88E-15 | MB1 | ANKRD42 |
| TMEM107 | 1.02E-19 | 0.743902 | 0.949 | 0.389 | 1.99E-15 | MB1 | TMEM107 |
| MAD2L11 | 1.59E-19 | 0.804142 | 0.949 | 0.378 | 3.09E-15 | MB1 | MAD2L1 |
| FAM174A | 1.62E-19 | 0.642347 | 0.872 | 0.31 | 3.16E-15 | MB1 | FAM174A |
| LZTFL1 | 1.74E-19 | 0.496935 | 0.846 | 0.26 | 3.38E-15 | MB1 | LZTFL1 |
| CCDC153 | 1.83E-19 | 0.546386 | 0.718 | 0.176 | 3.55E-15 | MB1 | CCDC153 |
| CDADC1 | 2.05E-19 | 0.464982 | 0.769 | 0.206 | 3.99E-15 | MB1 | CDADC1 |
| AURKA1 | 2.62E-19 | 0.46951 | 0.564 | 0.107 | 5.09E-15 | MB1 | AURKA |
| C2orf73 | 3.24E-19 | 0.295575 | 0.333 | 0.039 | 6.31E-15 | MB1 | C2orf73 |
| RPE65 | 3.40E-19 | 0.437838 | 0.538 | 0.095 | 6.62E-15 | MB1 | RPE65 |
| BTG3 | 3.54E-19 | 1.278344 | 1 | 0.769 | 6.89E-15 | MB1 | BTG3 |
| IFT43 | 3.68E-19 | 0.836654 | 0.974 | 0.57 | 7.17E-15 | MB1 | IFT43 |
| PBK1 | 4.05E-19 | 0.364006 | 0.667 | 0.131 | 7.89E-15 | MB1 | PBK |
| PITPNM1 | 4.19E-19 | 0.589375 | 0.744 | 0.204 | 8.16E-15 | MB1 | PITPNM1 |
| AHI1 | 4.27E-19 | 0.784509 | 0.923 | 0.382 | 8.30E-15 | MB1 | AHI1 |
| TRAF3IP2 | 4.54E-19 | 0.352199 | 0.462 | 0.074 | 8.84E-15 | MB1 | TRAF3IP2 |
| PTTG11 | 4.94E-19 | 0.980355 | 0.974 | 0.542 | 9.60E-15 | MB1 | PTTG1 |
| TMPO-AS | 5.17E-19 | 0.267716 | 0.41 | 0.059 | 1.01E-14 | MB1 | TMPO-AS1 |
| MDFIC | 5.33E-19 | 0.288098 | 0.487 | 0.079 | 1.04E-14 | MB1 | MDFIC |

| | | | | | | | |
|---|---|---|---|---|---|---|---|
| MINOS1 | 5.42E-19 | 0.770016 | 1 | 0.779 | 1.05E-14 | MB1 | MINOS1 |
| CEP19 | 6.43E-19 | 0.528055 | 0.692 | 0.174 | 1.25E-14 | MB1 | CEP19 |
| FAM229B | 6.83E-19 | 0.93827 | 1 | 0.728 | 1.33E-14 | MB1 | FAM229B |
| TRAF3IP1 | 7.56E-19 | 0.454303 | 0.692 | 0.165 | 1.47E-14 | MB1 | TRAF3IP1 |
| HIPK1 | 7.77E-19 | 0.430908 | 0.821 | 0.222 | 1.51E-14 | MB1 | HIPK1 |
| SPAG16 | 8.47E-19 | 0.830146 | 1 | 0.592 | 1.65E-14 | MB1 | SPAG16 |
| ANXA1 | 9.12E-19 | 0.813674 | 0.385 | 0.054 | 1.77E-14 | MB1 | ANXA1 |
| LRP11 | 9.89E-19 | 0.479801 | 0.744 | 0.19 | 1.92E-14 | MB1 | LRP11 |
| RP11-242I | 1.43E-18 | 0.292387 | 0.667 | 0.145 | 2.78E-14 | MB1 | RP11-242D8.1 |
| C21orf59 | 1.44E-18 | 0.99041 | 0.923 | 0.552 | 2.79E-14 | MB1 | C21orf59 |
| EFCAB11 | 1.62E-18 | 0.325004 | 0.692 | 0.157 | 3.15E-14 | MB1 | EFCAB11 |
| WDR54 | 2.03E-18 | 0.797336 | 1 | 0.763 | 3.94E-14 | MB1 | WDR54 |
| WDR60 | 2.28E-18 | 0.66913 | 0.949 | 0.42 | 4.43E-14 | MB1 | WDR60 |
| CFDP1 | 2.36E-18 | 1.091261 | 1 | 0.769 | 4.59E-14 | MB1 | CFDP1 |
| CCDC189 | 4.61E-18 | 0.464944 | 0.667 | 0.164 | 8.96E-14 | MB1 | CCDC189 |
| MOK | 4.87E-18 | 0.508995 | 0.744 | 0.22 | 9.47E-14 | MB1 | MOK |
| TAGLN2 | 5.34E-18 | 0.933341 | 1 | 0.617 | 1.04E-13 | MB1 | TAGLN2 |
| NKD1 | 6.32E-18 | 0.287537 | 0.487 | 0.084 | 1.23E-13 | MB1 | NKD1 |
| IFT122 | 7.25E-18 | 0.348819 | 0.641 | 0.147 | 1.41E-13 | MB1 | IFT122 |
| PSIP11 | 7.50E-18 | 0.982428 | 0.974 | 0.794 | 1.46E-13 | MB1 | PSIP1 |
| KIF231 | 7.83E-18 | 0.36191 | 0.538 | 0.105 | 1.52E-13 | MB1 | KIF23 |
| KIF221 | 9.67E-18 | 0.82955 | 0.897 | 0.409 | 1.88E-13 | MB1 | KIF22 |
| SYNE1 | 1.16E-17 | 0.327968 | 0.513 | 0.094 | 2.25E-13 | MB1 | SYNE1 |
| ICT1 | 1.27E-17 | 0.668919 | 0.872 | 0.383 | 2.47E-13 | MB1 | ICT1 |
| TMEM17 | 1.29E-17 | 0.41197 | 0.667 | 0.164 | 2.51E-13 | MB1 | TMEM17 |
| REV3L | 1.59E-17 | 0.569017 | 0.897 | 0.32 | 3.08E-13 | MB1 | REV3L |
| C15orf65 | 1.81E-17 | 0.393243 | 0.615 | 0.135 | 3.53E-13 | MB1 | C15orf65 |
| SLC4A8 | 1.90E-17 | 0.394432 | 0.718 | 0.179 | 3.69E-13 | MB1 | SLC4A8 |
| GLB1L | 2.48E-17 | 0.33313 | 0.59 | 0.129 | 4.83E-13 | MB1 | GLB1L |
| AK1 | 2.73E-17 | 1.114251 | 0.949 | 0.619 | 5.31E-13 | MB1 | AK1 |
| CFAP36 | 2.81E-17 | 0.656753 | 0.974 | 0.604 | 5.46E-13 | MB1 | CFAP36 |
| SLC12A2 | 3.19E-17 | 0.339278 | 0.667 | 0.158 | 6.20E-13 | MB1 | SLC12A2 |
| COPRS | 3.65E-17 | 0.983971 | 0.949 | 0.544 | 7.09E-13 | MB1 | COPRS |
| DYNLT11 | 3.65E-17 | 0.771324 | 1 | 0.879 | 7.10E-13 | MB1 | DYNLT1 |
| RGCC | 3.91E-17 | 0.4184 | 0.436 | 0.075 | 7.61E-13 | MB1 | RGCC |
| TUBG1 | 4.91E-17 | 0.755857 | 0.923 | 0.514 | 9.56E-13 | MB1 | TUBG1 |
| PCM1 | 6.54E-17 | 0.767717 | 0.949 | 0.578 | 1.27E-12 | MB1 | PCM1 |
| CSPP1 | 7.42E-17 | 0.36752 | 0.667 | 0.164 | 1.44E-12 | MB1 | CSPP1 |
| MYCBP | 7.50E-17 | 0.387938 | 0.59 | 0.134 | 1.46E-12 | MB1 | MYCBP |
| TPX21 | 9.00E-17 | 0.328595 | 0.641 | 0.144 | 1.75E-12 | MB1 | TPX2 |

| | | | | | | | |
|---|---|---|---|---|---|---|---|
| TPST2 | 1.06E-16 | 0.355806 | 0.744 | 0.197 | 2.06E-12 | MB1 | TPST2 |
| IK | 1.56E-16 | 0.728605 | 0.974 | 0.657 | 3.03E-12 | MB1 | IK |
| PROM1 | 1.88E-16 | 0.329975 | 0.436 | 0.076 | 3.65E-12 | MB1 | PROM1 |
| MDM1 | 2.16E-16 | 0.492719 | 0.769 | 0.231 | 4.20E-12 | MB1 | MDM1 |
| DALRD3 | 2.17E-16 | 0.715009 | 0.974 | 0.585 | 4.23E-12 | MB1 | DALRD3 |
| MAPK15 | 2.64E-16 | 0.370024 | 0.538 | 0.117 | 5.14E-12 | MB1 | MAPK15 |
| NDUFAF3 | 2.68E-16 | 0.734135 | 0.974 | 0.826 | 5.21E-12 | MB1 | NDUFAF3 |
| CCDC74B | 2.72E-16 | 0.594077 | 0.692 | 0.214 | 5.29E-12 | MB1 | CCDC74B |
| B9D1 | 2.75E-16 | 0.752446 | 0.897 | 0.41 | 5.34E-12 | MB1 | B9D1 |
| UTRN | 2.96E-16 | 0.438327 | 0.667 | 0.174 | 5.76E-12 | MB1 | UTRN |
| PHTF1 | 3.02E-16 | 0.573364 | 0.846 | 0.314 | 5.87E-12 | MB1 | PHTF1 |
| IFT52 | 3.03E-16 | 0.650292 | 0.974 | 0.574 | 5.89E-12 | MB1 | IFT52 |
| YWHAH1 | 3.42E-16 | 0.746964 | 1 | 0.755 | 6.65E-12 | MB1 | YWHAH |
| CALM11 | 4.10E-16 | 0.807908 | 1 | 0.965 | 7.99E-12 | MB1 | CALM1 |
| OAT1 | 4.39E-16 | 0.769823 | 0.974 | 0.532 | 8.54E-12 | MB1 | OAT |
| SNRPG | 4.45E-16 | 0.621492 | 1 | 0.894 | 8.66E-12 | MB1 | SNRPG |
| ZNRF1 | 4.99E-16 | 0.370959 | 0.513 | 0.105 | 9.71E-12 | MB1 | ZNRF1 |
| MLLT1 | 5.09E-16 | 0.793453 | 0.897 | 0.477 | 9.90E-12 | MB1 | MLLT1 |
| RSPH3 | 5.23E-16 | 0.355609 | 0.641 | 0.163 | 1.02E-11 | MB1 | RSPH3 |
| STPG1 | 5.44E-16 | 0.318549 | 0.564 | 0.127 | 1.06E-11 | MB1 | STPG1 |
| ECT21 | 5.45E-16 | 0.301553 | 0.538 | 0.113 | 1.06E-11 | MB1 | ECT2 |
| IGFBP7 | 5.53E-16 | 0.310309 | 0.513 | 0.097 | 1.08E-11 | MB1 | IGFBP7 |
| IPO13 | 8.37E-16 | 0.348963 | 0.667 | 0.174 | 1.63E-11 | MB1 | IPO13 |
| ZBBX | 8.47E-16 | 0.267999 | 0.487 | 0.095 | 1.65E-11 | MB1 | ZBBX |
| HSPH12 | 9.37E-16 | 0.962177 | 0.974 | 0.549 | 1.82E-11 | MB1 | HSPH1 |
| AC007325 | 9.66E-16 | 0.643128 | 0.897 | 0.363 | 1.88E-11 | MB1 | AC007325.4 |
| KIF20B1 | 1.05E-15 | 0.293119 | 0.769 | 0.205 | 2.04E-11 | MB1 | KIF20B |
| CEP83 | 1.18E-15 | 0.445382 | 0.641 | 0.17 | 2.29E-11 | MB1 | CEP83 |
| GLT8D1 | 1.46E-15 | 0.64214 | 0.923 | 0.48 | 2.84E-11 | MB1 | GLT8D1 |
| ZNHIT2 | 1.56E-15 | 0.503987 | 0.795 | 0.278 | 3.04E-11 | MB1 | ZNHIT2 |
| AGTRAP | 1.65E-15 | 0.76396 | 0.821 | 0.408 | 3.21E-11 | MB1 | AGTRAP |
| PAQR4 | 1.67E-15 | 0.505471 | 0.667 | 0.192 | 3.24E-11 | MB1 | PAQR4 |
| CTD-2015 | 1.71E-15 | 0.424488 | 0.641 | 0.164 | 3.33E-11 | MB1 | CTD-2015H6.3 |
| CC2D2A | 1.82E-15 | 0.354505 | 0.769 | 0.225 | 3.54E-11 | MB1 | CC2D2A |
| ARHGAP1ﾐ | 2.10E-15 | 0.420354 | 0.385 | 0.066 | 4.08E-11 | MB1 | ARHGAP18 |
| RANBP11 | 2.17E-15 | 0.638261 | 1 | 0.944 | 4.21E-11 | MB1 | RANBP1 |
| C20orf96 | 2.61E-15 | 0.721391 | 0.872 | 0.395 | 5.08E-11 | MB1 | C20orf96 |
| GMPPB | 2.78E-15 | 0.445447 | 0.718 | 0.228 | 5.40E-11 | MB1 | GMPPB |
| MIPEP | 2.83E-15 | 0.299911 | 0.615 | 0.153 | 5.50E-11 | MB1 | MIPEP |
| TM7SF3 | 3.02E-15 | 0.655965 | 0.949 | 0.474 | 5.88E-11 | MB1 | TM7SF3 |

| | | | | | | | |
|---|---|---|---|---|---|---|---|
| RFK | 3.82E-15 | 0.669071 | 0.974 | 0.498 | 7.43E-11 | MB1 | RFK |
| RUVBL1 | 4.01E-15 | 0.739101 | 0.949 | 0.517 | 7.80E-11 | MB1 | RUVBL1 |
| HSPA8 | 4.95E-15 | 0.830111 | 1 | 0.981 | 9.62E-11 | MB1 | HSPA8 |
| CSRP2 | 5.42E-15 | 0.81533 | 0.974 | 0.656 | 1.05E-10 | MB1 | CSRP2 |
| FRMD8 | 5.72E-15 | 0.250443 | 0.385 | 0.066 | 1.11E-10 | MB1 | FRMD8 |
| PSMC3 | 6.06E-15 | 0.824398 | 0.949 | 0.742 | 1.18E-10 | MB1 | PSMC3 |
| PKIG | 7.47E-15 | 0.648436 | 0.949 | 0.516 | 1.45E-10 | MB1 | PKIG |
| ANXA21 | 8.00E-15 | 0.859518 | 0.923 | 0.557 | 1.56E-10 | MB1 | ANXA2 |
| ZNF487 | 8.70E-15 | 0.342357 | 0.564 | 0.138 | 1.69E-10 | MB1 | ZNF487 |
| VDAC3 | 8.83E-15 | 0.685836 | 1 | 0.746 | 1.72E-10 | MB1 | VDAC3 |
| S100A10 | 1.36E-14 | 0.356011 | 0.487 | 0.102 | 2.64E-10 | MB1 | S100A10 |
| CEP41 | 1.71E-14 | 0.591424 | 0.795 | 0.3 | 3.33E-10 | MB1 | CEP41 |
| TCTN2 | 1.71E-14 | 0.266243 | 0.436 | 0.081 | 3.34E-10 | MB1 | TCTN2 |
| GCLM | 1.72E-14 | 0.55066 | 0.769 | 0.291 | 3.34E-10 | MB1 | GCLM |
| SAXO2 | 1.73E-14 | 0.288224 | 0.462 | 0.095 | 3.36E-10 | MB1 | SAXO2 |
| ODF21 | 1.77E-14 | 0.623299 | 0.872 | 0.445 | 3.44E-10 | MB1 | ODF2 |
| CDKN2D1 | 2.48E-14 | 0.589891 | 0.821 | 0.302 | 4.83E-10 | MB1 | CDKN2D |
| FOLR1 | 2.58E-14 | 0.491184 | 0.667 | 0.181 | 5.03E-10 | MB1 | FOLR1 |
| HSPE1 | 3.02E-14 | 0.69591 | 1 | 0.909 | 5.88E-10 | MB1 | HSPE1 |
| SLC25A14 | 3.04E-14 | 0.457181 | 0.667 | 0.205 | 5.91E-10 | MB1 | SLC25A14 |
| TRIP131 | 3.77E-14 | 0.3989 | 0.513 | 0.115 | 7.34E-10 | MB1 | TRIP13 |
| PSME2 | 4.45E-14 | 0.640484 | 0.974 | 0.591 | 8.66E-10 | MB1 | PSME2 |
| DTNA | 4.70E-14 | 0.538761 | 0.795 | 0.315 | 9.14E-10 | MB1 | DTNA |
| CALM2 | 5.06E-14 | 0.758083 | 1 | 0.997 | 9.84E-10 | MB1 | CALM2 |
| TCTN1 | 7.59E-14 | 0.632124 | 0.846 | 0.377 | 1.48E-09 | MB1 | TCTN1 |
| DNAL1 | 8.05E-14 | 0.45598 | 0.769 | 0.269 | 1.57E-09 | MB1 | DNAL1 |
| CLHC1 | 8.64E-14 | 0.32071 | 0.615 | 0.165 | 1.68E-09 | MB1 | CLHC1 |
| NME7 | 8.90E-14 | 0.571312 | 1 | 0.456 | 1.73E-09 | MB1 | NME7 |
| PSD | 1.04E-13 | 0.324142 | 0.462 | 0.098 | 2.02E-09 | MB1 | PSD |
| CDT11 | 1.15E-13 | 0.354409 | 0.641 | 0.171 | 2.23E-09 | MB1 | CDT1 |
| RP11-60L: | 1.36E-13 | 0.292184 | 0.333 | 0.054 | 2.64E-09 | MB1 | RP11-60L3.6 |
| FABP6 | 1.53E-13 | 0.274379 | 0.308 | 0.046 | 2.98E-09 | MB1 | FABP6 |
| EMC4 | 1.86E-13 | 0.64418 | 0.974 | 0.781 | 3.63E-09 | MB1 | EMC4 |
| MAATS1 | 1.92E-13 | 0.293578 | 0.513 | 0.117 | 3.73E-09 | MB1 | MAATS1 |
| FAM227A | 2.00E-13 | 0.334247 | 0.59 | 0.157 | 3.89E-09 | MB1 | FAM227A |
| EFCAB2 | 2.29E-13 | 0.411479 | 0.564 | 0.153 | 4.45E-09 | MB1 | EFCAB2 |
| UBE2S1 | 2.76E-13 | 0.991722 | 0.974 | 0.736 | 5.37E-09 | MB1 | UBE2S |
| UBN2 | 3.13E-13 | 0.260306 | 0.641 | 0.17 | 6.08E-09 | MB1 | UBN2 |
| ANKRD54 | 3.29E-13 | 0.630972 | 0.795 | 0.371 | 6.39E-09 | MB1 | ANKRD54 |
| EIF2S21 | 3.41E-13 | 0.618778 | 1 | 0.83 | 6.63E-09 | MB1 | EIF2S2 |

| | | | | | | | |
|---|---|---|---|---|---|---|---|
| MEAF61 | 3.86E-13 | 0.634498 | 1 | 0.856 | 7.51E-09 | MB1 | MEAF6 |
| YBX3 | 4.92E-13 | 0.551469 | 0.923 | 0.464 | 9.58E-09 | MB1 | YBX3 |
| DPY30 | 6.35E-13 | 0.716864 | 0.974 | 0.719 | 1.24E-08 | MB1 | DPY30 |
| ACTR6 | 7.05E-13 | 0.591295 | 0.821 | 0.361 | 1.37E-08 | MB1 | ACTR6 |
| HSBP1 | 7.15E-13 | 0.561277 | 1 | 0.981 | 1.39E-08 | MB1 | HSBP1 |
| SNAPC11 | 8.27E-13 | 0.524222 | 0.872 | 0.401 | 1.61E-08 | MB1 | SNAPC1 |
| C11orf74 | 8.64E-13 | 0.516893 | 0.949 | 0.501 | 1.68E-08 | MB1 | C11orf74 |
| COG7 | 8.99E-13 | 0.390949 | 0.615 | 0.19 | 1.75E-08 | MB1 | COG7 |
| RNASEH2/ | 9.16E-13 | 0.652441 | 0.821 | 0.371 | 1.78E-08 | MB1 | RNASEH2A |
| MIS18A1 | 9.36E-13 | 0.494754 | 0.821 | 0.333 | 1.82E-08 | MB1 | MIS18A |
| PNMA11 | 1.02E-12 | 0.785347 | 0.923 | 0.629 | 1.98E-08 | MB1 | PNMA1 |
| WDR34 | 1.03E-12 | 0.626708 | 0.923 | 0.479 | 2.00E-08 | MB1 | WDR34 |
| KCTD17 | 1.15E-12 | 0.449124 | 0.692 | 0.245 | 2.24E-08 | MB1 | KCTD17 |
| WRB | 1.18E-12 | 0.654582 | 1 | 0.845 | 2.30E-08 | MB1 | WRB |
| CLMN | 1.29E-12 | 0.309904 | 0.462 | 0.104 | 2.50E-08 | MB1 | CLMN |
| IFT27 | 1.34E-12 | 0.608893 | 0.949 | 0.641 | 2.60E-08 | MB1 | IFT27 |
| CHIC2 | 1.60E-12 | 0.316594 | 0.718 | 0.213 | 3.12E-08 | MB1 | CHIC2 |
| SLC41A1 | 1.80E-12 | 0.279861 | 0.513 | 0.127 | 3.50E-08 | MB1 | SLC41A1 |
| DZIP3 | 1.95E-12 | 0.586327 | 0.846 | 0.408 | 3.79E-08 | MB1 | DZIP3 |
| SLC13A4 | 2.16E-12 | 0.330389 | 0.641 | 0.179 | 4.21E-08 | MB1 | SLC13A4 |
| STYXL1 | 2.53E-12 | 0.318827 | 0.615 | 0.175 | 4.91E-08 | MB1 | STYXL1 |
| C5orf15 | 2.57E-12 | 0.599275 | 0.949 | 0.6 | 4.99E-08 | MB1 | C5orf15 |
| BUD312 | 2.79E-12 | 0.629988 | 0.974 | 0.85 | 5.43E-08 | MB1 | BUD31 |
| HNRNPF | 2.95E-12 | 0.614866 | 0.974 | 0.764 | 5.73E-08 | MB1 | HNRNPF |
| APITD1 | 3.14E-12 | 0.256469 | 0.538 | 0.137 | 6.11E-08 | MB1 | APITD1 |
| LRTOMT | 3.51E-12 | 0.413547 | 0.641 | 0.199 | 6.83E-08 | MB1 | LRTOMT |
| COMMD8 | 4.08E-12 | 0.40062 | 0.846 | 0.354 | 7.93E-08 | MB1 | COMMD8 |
| IFT46 | 4.14E-12 | 0.418112 | 0.744 | 0.271 | 8.05E-08 | MB1 | IFT46 |
| S100A111 | 4.94E-12 | 0.369899 | 0.667 | 0.211 | 9.62E-08 | MB1 | S100A11 |
| SRI1 | 5.09E-12 | 0.559128 | 1 | 0.821 | 9.90E-08 | MB1 | SRI |
| FAM181B | 5.71E-12 | 0.304651 | 0.59 | 0.165 | 1.11E-07 | MB1 | FAM181B |
| RASSF8-A | 6.32E-12 | 0.280729 | 0.641 | 0.186 | 1.23E-07 | MB1 | RASSF8-AS1 |
| CIB12 | 6.39E-12 | 0.600722 | 1 | 0.745 | 1.24E-07 | MB1 | CIB1 |
| CABIN1 | 6.82E-12 | 0.345857 | 0.692 | 0.222 | 1.33E-07 | MB1 | CABIN1 |
| RPL39L | 7.90E-12 | 0.411783 | 0.718 | 0.258 | 1.54E-07 | MB1 | RPL39L |
| NUDT15 | 8.03E-12 | 0.491593 | 0.769 | 0.302 | 1.56E-07 | MB1 | NUDT15 |
| CCDC191 | 8.97E-12 | 0.365161 | 0.538 | 0.151 | 1.74E-07 | MB1 | CCDC191 |
| TTLL5 | 9.05E-12 | 0.258702 | 0.436 | 0.101 | 1.76E-07 | MB1 | TTLL5 |
| RABL2B | 9.07E-12 | 0.530919 | 0.795 | 0.351 | 1.76E-07 | MB1 | RABL2B |
| SSB | 9.31E-12 | 0.59508 | 1 | 0.846 | 1.81E-07 | MB1 | SSB |

| | | | | | | | |
|---|---|---|---|---|---|---|---|
| CKS23 | 1.18E-11 | 0.867998 | 0.974 | 0.781 | 2.30E-07 | MB1 | CKS2 |
| TACC31 | 1.19E-11 | 0.328094 | 0.462 | 0.109 | 2.31E-07 | MB1 | TACC3 |
| IGFBPL11 | 1.37E-11 | 0.250135 | 0.462 | 0.109 | 2.66E-07 | MB1 | IGFBPL1 |
| ABAT | 1.44E-11 | 0.359679 | 0.769 | 0.288 | 2.80E-07 | MB1 | ABAT |
| KATNB1 | 1.46E-11 | 0.472747 | 0.821 | 0.327 | 2.84E-07 | MB1 | KATNB1 |
| TMX41 | 1.85E-11 | 0.475135 | 0.846 | 0.397 | 3.59E-07 | MB1 | TMX4 |
| HACD31 | 1.85E-11 | 0.739764 | 0.974 | 0.864 | 3.61E-07 | MB1 | HACD3 |
| CFAP20 | 2.13E-11 | 0.570587 | 0.949 | 0.694 | 4.15E-07 | MB1 | CFAP20 |
| SUPT20H | 2.18E-11 | 0.512794 | 0.872 | 0.46 | 4.24E-07 | MB1 | SUPT20H |
| STRBP | 2.20E-11 | 0.494814 | 0.795 | 0.341 | 4.28E-07 | MB1 | STRBP |
| ADD3 | 2.33E-11 | 0.46469 | 0.846 | 0.361 | 4.52E-07 | MB1 | ADD3 |
| INAFM11 | 2.40E-11 | 0.36587 | 0.872 | 0.338 | 4.67E-07 | MB1 | INAFM1 |
| TUBA1B1 | 2.50E-11 | 0.540265 | 1 | 0.986 | 4.87E-07 | MB1 | TUBA1B |
| CLUAP1 | 2.61E-11 | 0.46264 | 0.872 | 0.386 | 5.09E-07 | MB1 | CLUAP1 |
| GSKIP | 2.66E-11 | 0.532615 | 0.872 | 0.425 | 5.18E-07 | MB1 | GSKIP |
| NAA20 | 2.71E-11 | 0.546241 | 0.949 | 0.602 | 5.28E-07 | MB1 | NAA20 |
| DHFR1 | 2.75E-11 | 0.375838 | 0.744 | 0.258 | 5.34E-07 | MB1 | DHFR |
| CEP78 | 2.99E-11 | 0.412681 | 0.615 | 0.2 | 5.81E-07 | MB1 | CEP78 |
| CEP95 | 3.05E-11 | 0.33061 | 0.718 | 0.241 | 5.93E-07 | MB1 | CEP95 |
| RNF6 | 3.56E-11 | 0.333045 | 0.667 | 0.224 | 6.93E-07 | MB1 | RNF6 |
| CTXN1 | 3.73E-11 | 0.498588 | 0.872 | 0.424 | 7.26E-07 | MB1 | CTXN1 |
| PPID | 3.85E-11 | 0.51673 | 0.949 | 0.531 | 7.49E-07 | MB1 | PPID |
| PLTP1 | 3.98E-11 | 0.654169 | 0.974 | 0.756 | 7.75E-07 | MB1 | PLTP |
| MRPL57 | 4.60E-11 | 0.511294 | 1 | 0.839 | 8.95E-07 | MB1 | MRPL57 |
| DRAM2 | 4.71E-11 | 0.455248 | 0.949 | 0.606 | 9.17E-07 | MB1 | DRAM2 |
| OTUD4 | 4.75E-11 | 0.358827 | 0.487 | 0.132 | 9.25E-07 | MB1 | OTUD4 |
| IQCE | 4.78E-11 | 0.344494 | 0.667 | 0.232 | 9.31E-07 | MB1 | IQCE |
| ZNF599 | 7.10E-11 | 0.298146 | 0.538 | 0.156 | 1.38E-06 | MB1 | ZNF599 |
| ANXA11 | 7.18E-11 | 0.39706 | 0.667 | 0.231 | 1.40E-06 | MB1 | ANXA11 |
| KLHDC9 | 8.24E-11 | 0.426182 | 0.744 | 0.286 | 1.60E-06 | MB1 | KLHDC9 |
| DNAJA1 | 8.92E-11 | 0.530142 | 0.974 | 0.889 | 1.73E-06 | MB1 | DNAJA1 |
| PRADC1 | 8.98E-11 | 0.51703 | 0.769 | 0.336 | 1.75E-06 | MB1 | PRADC1 |
| CEP350 | 9.67E-11 | 0.350698 | 0.692 | 0.251 | 1.88E-06 | MB1 | CEP350 |
| DBF41 | 1.01E-10 | 0.398269 | 0.667 | 0.241 | 1.96E-06 | MB1 | DBF4 |
| DNAL4 | 1.06E-10 | 0.461218 | 0.795 | 0.365 | 2.07E-06 | MB1 | DNAL4 |
| UQCRFS1 | 1.19E-10 | 0.440092 | 0.974 | 0.899 | 2.31E-06 | MB1 | UQCRFS1 |
| IFT81 | 1.26E-10 | 0.417503 | 0.795 | 0.339 | 2.44E-06 | MB1 | IFT81 |
| COQ4 | 1.26E-10 | 0.539788 | 0.949 | 0.61 | 2.45E-06 | MB1 | COQ4 |
| C12orf76 | 1.32E-10 | 0.480957 | 0.949 | 0.61 | 2.56E-06 | MB1 | C12orf76 |
| ACTN1 | 1.36E-10 | 0.458499 | 0.795 | 0.351 | 2.65E-06 | MB1 | ACTN1 |

| | | | | | | | |
|---|---|---|---|---|---|---|---|
| TNFAIP8L | 1.38E-10 | 0.307325 | 0.41 | 0.098 | 2.69E-06 | MB1 | TNFAIP8L1 |
| FBRS | 1.39E-10 | 0.274748 | 0.538 | 0.158 | 2.71E-06 | MB1 | FBRS |
| THOC7 | 1.42E-10 | 0.454546 | 0.897 | 0.468 | 2.76E-06 | MB1 | THOC7 |
| ZSCAN1 | 1.46E-10 | 0.514472 | 0.744 | 0.333 | 2.83E-06 | MB1 | ZSCAN1 |
| CISD2 | 1.67E-10 | 0.572255 | 0.974 | 0.687 | 3.25E-06 | MB1 | CISD2 |
| EZR1 | 1.94E-10 | 0.563961 | 0.821 | 0.463 | 3.78E-06 | MB1 | EZR |
| LYRM5 | 1.98E-10 | 0.343043 | 0.744 | 0.28 | 3.86E-06 | MB1 | LYRM5 |
| HSP90AA1 | 2.11E-10 | 0.950899 | 1 | 0.994 | 4.11E-06 | MB1 | HSP90AA1 |
| CHCHD2 | 2.37E-10 | 0.53913 | 1 | 0.987 | 4.62E-06 | MB1 | CHCHD2 |
| ACTN4 | 2.48E-10 | 0.4342 | 0.769 | 0.351 | 4.83E-06 | MB1 | ACTN4 |
| LCA5 | 2.53E-10 | 0.268901 | 0.692 | 0.233 | 4.93E-06 | MB1 | LCA5 |
| ROGDI | 2.60E-10 | 0.461171 | 0.795 | 0.39 | 5.06E-06 | MB1 | ROGDI |
| TACC1 | 2.77E-10 | 0.289568 | 0.564 | 0.174 | 5.38E-06 | MB1 | TACC1 |
| STK33 | 2.80E-10 | 0.369413 | 0.692 | 0.274 | 5.45E-06 | MB1 | STK33 |
| TEX9 | 3.10E-10 | 0.355741 | 0.59 | 0.204 | 6.03E-06 | MB1 | TEX9 |
| MRPL43 | 3.37E-10 | 0.800089 | 0.923 | 0.68 | 6.55E-06 | MB1 | MRPL43 |
| BBIP1 | 3.69E-10 | 0.457076 | 0.821 | 0.375 | 7.17E-06 | MB1 | BBIP1 |
| G2E3 | 3.73E-10 | 0.358454 | 0.615 | 0.21 | 7.26E-06 | MB1 | G2E3 |
| MCAT | 3.96E-10 | 0.268813 | 0.744 | 0.274 | 7.71E-06 | MB1 | MCAT |
| CCDC24 | 3.98E-10 | 0.256956 | 0.513 | 0.146 | 7.74E-06 | MB1 | CCDC24 |
| DHRS4 | 4.53E-10 | 0.256291 | 0.41 | 0.101 | 8.82E-06 | MB1 | DHRS4 |
| GALNT18 | 4.73E-10 | 0.295216 | 0.59 | 0.188 | 9.21E-06 | MB1 | GALNT18 |
| ZNF584 | 4.79E-10 | 0.320209 | 0.59 | 0.2 | 9.32E-06 | MB1 | ZNF584 |
| FAM161A | 5.11E-10 | 0.393289 | 0.744 | 0.302 | 9.93E-06 | MB1 | FAM161A |
| FAM206A | 5.14E-10 | 0.284425 | 0.513 | 0.146 | 1.00E-05 | MB1 | FAM206A |
| TCEB2 | 5.15E-10 | 0.451947 | 1 | 0.994 | 1.00E-05 | MB1 | TCEB2 |
| TTR | 6.11E-10 | 0.812439 | 1 | 1 | 1.19E-05 | MB1 | TTR |
| CCDC14 | 6.36E-10 | 0.557515 | 0.769 | 0.337 | 1.24E-05 | MB1 | CCDC14 |
| CHAF1A1 | 6.73E-10 | 0.284224 | 0.513 | 0.142 | 1.31E-05 | MB1 | CHAF1A |
| SMUG1 | 7.11E-10 | 0.387389 | 0.744 | 0.305 | 1.38E-05 | MB1 | SMUG1 |
| TMEM138 | 8.47E-10 | 0.382142 | 0.744 | 0.301 | 1.65E-05 | MB1 | TMEM138 |
| GPR162 | 1.00E-09 | 0.392096 | 0.641 | 0.232 | 1.95E-05 | MB1 | GPR162 |
| ACADVL | 1.22E-09 | 0.639839 | 0.923 | 0.69 | 2.38E-05 | MB1 | ACADVL |
| PSMD10 | 1.32E-09 | 0.46789 | 0.949 | 0.598 | 2.57E-05 | MB1 | PSMD10 |
| PTRH1 | 1.35E-09 | 0.351631 | 0.692 | 0.275 | 2.63E-05 | MB1 | PTRH1 |
| CLU1 | 1.42E-09 | 0.574358 | 1 | 0.926 | 2.76E-05 | MB1 | CLU |
| MEA1 | 1.43E-09 | 0.453196 | 0.923 | 0.658 | 2.78E-05 | MB1 | MEA1 |
| C5orf45 | 1.58E-09 | 0.413211 | 0.769 | 0.33 | 3.07E-05 | MB1 | C5orf45 |
| TRPM3 | 1.58E-09 | 0.428396 | 0.846 | 0.372 | 3.08E-05 | MB1 | TRPM3 |
| FAM216A | 1.66E-09 | 0.326184 | 0.59 | 0.202 | 3.24E-05 | MB1 | FAM216A |

| | | | | | | | |
|---|---|---|---|---|---|---|---|
| LINC0049: | 1.70E-09 | 0.541217 | 0.974 | 0.821 | 3.32E-05 | MB1 | LINC00493 |
| TULP3 | 1.74E-09 | 0.378957 | 0.744 | 0.324 | 3.38E-05 | MB1 | TULP3 |
| SPCS11 | 1.80E-09 | 0.387976 | 0.974 | 0.946 | 3.49E-05 | MB1 | SPCS1 |
| PPP1R7 | 1.91E-09 | 0.400531 | 0.897 | 0.5 | 3.71E-05 | MB1 | PPP1R7 |
| TEX30 | 1.96E-09 | 0.317123 | 0.641 | 0.218 | 3.82E-05 | MB1 | TEX30 |
| DYNC2LI1 | 2.06E-09 | 0.492262 | 0.846 | 0.484 | 4.01E-05 | MB1 | DYNC2LI1 |
| UQCC21 | 2.09E-09 | 0.445921 | 0.974 | 0.741 | 4.07E-05 | MB1 | UQCC2 |
| ANAPC7 | 2.44E-09 | 0.267956 | 0.667 | 0.245 | 4.74E-05 | MB1 | ANAPC7 |
| SRSF91 | 2.62E-09 | 0.447888 | 1 | 0.915 | 5.09E-05 | MB1 | SRSF9 |
| H2AFJ | 3.32E-09 | 0.413194 | 0.821 | 0.402 | 6.45E-05 | MB1 | H2AFJ |
| MAZ | 3.33E-09 | 0.400716 | 0.872 | 0.449 | 6.47E-05 | MB1 | MAZ |
| TUSC3 | 3.34E-09 | 0.43699 | 1 | 0.726 | 6.50E-05 | MB1 | TUSC3 |
| LMO2 | 3.41E-09 | 0.275326 | 0.513 | 0.157 | 6.64E-05 | MB1 | LMO2 |
| CDCA81 | 3.63E-09 | 0.256579 | 0.41 | 0.105 | 7.07E-05 | MB1 | CDCA8 |
| NOP56 | 3.67E-09 | 0.514079 | 0.897 | 0.632 | 7.14E-05 | MB1 | NOP56 |
| TECR | 3.86E-09 | 0.48007 | 1 | 0.846 | 7.51E-05 | MB1 | TECR |
| LINC0009/ | 4.15E-09 | 0.383809 | 0.769 | 0.331 | 8.06E-05 | MB1 | LINC00094 |
| PRPS2 | 4.72E-09 | 0.264593 | 0.462 | 0.136 | 9.18E-05 | MB1 | PRPS2 |
| NQO1 | 4.81E-09 | 0.564776 | 0.846 | 0.446 | 9.36E-05 | MB1 | NQO1 |
| SPPL2B | 5.19E-09 | 0.316816 | 0.692 | 0.285 | 0.000101 | MB1 | SPPL2B |
| RWDD4 | 5.57E-09 | 0.475928 | 0.923 | 0.575 | 0.000108 | MB1 | RWDD4 |
| TMEM106 | 6.13E-09 | 0.477278 | 0.872 | 0.522 | 0.000119 | MB1 | TMEM106C |
| DERL3 | 6.14E-09 | 0.352849 | 0.462 | 0.144 | 0.000119 | MB1 | DERL3 |
| ARMC9 | 6.18E-09 | 0.414479 | 0.59 | 0.23 | 0.00012 | MB1 | ARMC9 |
| EID2B | 6.26E-09 | 0.289915 | 0.564 | 0.184 | 0.000122 | MB1 | EID2B |
| MBOAT7 | 6.41E-09 | 0.383766 | 0.897 | 0.459 | 0.000125 | MB1 | MBOAT7 |
| PPP1R2 | 6.51E-09 | 0.78882 | 0.846 | 0.546 | 0.000127 | MB1 | PPP1R2 |
| KLHDC8B | 6.60E-09 | 0.561645 | 0.949 | 0.718 | 0.000128 | MB1 | KLHDC8B |
| MRPS33 | 6.71E-09 | 0.514869 | 0.974 | 0.586 | 0.000131 | MB1 | MRPS33 |
| PBXIP1 | 8.09E-09 | 0.493062 | 0.641 | 0.284 | 0.000157 | MB1 | PBXIP1 |
| MAPRE31 | 9.07E-09 | 0.325364 | 0.667 | 0.278 | 0.000176 | MB1 | MAPRE3 |
| PLIN3 | 9.14E-09 | 0.400283 | 0.923 | 0.517 | 0.000178 | MB1 | PLIN3 |
| SLC35A21 | 9.54E-09 | 0.402584 | 0.923 | 0.471 | 0.000186 | MB1 | SLC35A2 |
| MAP9 | 1.04E-08 | 0.369115 | 0.897 | 0.521 | 0.000203 | MB1 | MAP9 |
| HSPD11 | 1.05E-08 | 0.551821 | 0.974 | 0.947 | 0.000205 | MB1 | HSPD1 |
| EIF2B4 | 1.10E-08 | 0.318578 | 0.692 | 0.295 | 0.000214 | MB1 | EIF2B4 |
| C12orf75 | 1.11E-08 | 0.519389 | 0.872 | 0.487 | 0.000217 | MB1 | C12orf75 |
| GTF2E2 | 1.12E-08 | 0.401893 | 0.769 | 0.359 | 0.000218 | MB1 | GTF2E2 |
| TTC5 | 1.16E-08 | 0.383042 | 0.769 | 0.362 | 0.000225 | MB1 | TTC5 |
| MRPS6 | 1.31E-08 | 0.462202 | 0.974 | 0.811 | 0.000254 | MB1 | MRPS6 |

| | | | | | | | |
|---|---|---|---|---|---|---|---|
| ANKRD37: | 1.71E-08 | 0.334609 | 0.615 | 0.229 | 0.000333 | MB1 | ANKRD37 |
| GALK2 | 1.77E-08 | 0.347748 | 0.615 | 0.241 | 0.000345 | MB1 | GALK2 |
| POLR2I | 1.84E-08 | 0.515496 | 1 | 0.84 | 0.000358 | MB1 | POLR2I |
| PRKAR2A | 2.02E-08 | 0.496738 | 0.795 | 0.464 | 0.000394 | MB1 | PRKAR2A |
| SURF2 | 2.17E-08 | 0.309435 | 0.821 | 0.392 | 0.000423 | MB1 | SURF2 |
| SLC25A4 | 2.23E-08 | 0.525319 | 0.923 | 0.637 | 0.000433 | MB1 | SLC25A4 |
| DNPH1 | 2.32E-08 | 0.420986 | 0.949 | 0.647 | 0.000452 | MB1 | DNPH1 |
| RHBDD2 | 2.45E-08 | 0.454108 | 1 | 0.866 | 0.000476 | MB1 | RHBDD2 |
| MRFAP1 | 2.47E-08 | 0.409568 | 0.949 | 0.902 | 0.000481 | MB1 | MRFAP1 |
| SRGAP2B | 2.69E-08 | 0.432668 | 0.538 | 0.201 | 0.000523 | MB1 | SRGAP2B |
| IDH21 | 2.94E-08 | 0.468567 | 0.897 | 0.697 | 0.000571 | MB1 | IDH2 |
| AHSA1 | 3.06E-08 | 0.586117 | 0.872 | 0.607 | 0.000596 | MB1 | AHSA1 |
| APOO | 3.29E-08 | 0.454306 | 0.872 | 0.577 | 0.00064 | MB1 | APOO |
| ELMOD2 | 3.68E-08 | 0.313221 | 0.538 | 0.189 | 0.000715 | MB1 | ELMOD2 |
| YWHAE | 3.83E-08 | 0.355102 | 1 | 0.99 | 0.000744 | MB1 | YWHAE |
| LACTB2 | 4.20E-08 | 0.390215 | 0.667 | 0.283 | 0.000818 | MB1 | LACTB2 |
| UQCR11 | 4.41E-08 | 0.409833 | 1 | 0.957 | 0.000857 | MB1 | UQCR11 |
| HIST1H1C | 4.53E-08 | 0.591313 | 0.795 | 0.418 | 0.000881 | MB1 | HIST1H1C |
| CCDC109E | 4.65E-08 | 0.399045 | 0.641 | 0.294 | 0.000904 | MB1 | CCDC109B |
| LRRC49 | 4.83E-08 | 0.273353 | 0.667 | 0.266 | 0.000939 | MB1 | LRRC49 |
| PARD6A | 5.02E-08 | 0.283168 | 0.538 | 0.187 | 0.000977 | MB1 | PARD6A |
| CTBS | 5.31E-08 | 0.267221 | 0.667 | 0.282 | 0.001033 | MB1 | CTBS |
| AKAP9 | 5.99E-08 | 0.603139 | 0.923 | 0.668 | 0.001165 | MB1 | AKAP9 |
| RMDN3 | 6.47E-08 | 0.28163 | 0.821 | 0.374 | 0.001258 | MB1 | RMDN3 |
| RACGAP1: | 7.13E-08 | 0.346303 | 0.538 | 0.193 | 0.001387 | MB1 | RACGAP1 |
| ISCA2 | 9.40E-08 | 0.378616 | 0.923 | 0.584 | 0.001829 | MB1 | ISCA2 |
| NDUFA8 | 1.04E-07 | 0.416097 | 0.949 | 0.663 | 0.002015 | MB1 | NDUFA8 |
| NEK1 | 1.19E-07 | 0.271353 | 0.538 | 0.191 | 0.002306 | MB1 | NEK1 |
| WBP11 | 1.21E-07 | 0.440835 | 0.821 | 0.57 | 0.002358 | MB1 | WBP11 |
| CNTLN | 1.25E-07 | 0.293786 | 0.564 | 0.224 | 0.002436 | MB1 | CNTLN |
| KPNA21 | 1.27E-07 | 0.375887 | 0.949 | 0.688 | 0.002462 | MB1 | KPNA2 |
| SLC22A17 | 1.38E-07 | 0.435622 | 0.949 | 0.712 | 0.002675 | MB1 | SLC22A17 |
| RABL2A | 1.43E-07 | 0.360131 | 0.615 | 0.268 | 0.002777 | MB1 | RABL2A |
| CYSTM1 | 1.47E-07 | 0.424543 | 0.949 | 0.761 | 0.002868 | MB1 | CYSTM1 |
| TFDP1 | 1.50E-07 | 0.388952 | 0.692 | 0.315 | 0.002927 | MB1 | TFDP1 |
| CADM11 | 1.61E-07 | 0.399484 | 0.872 | 0.563 | 0.003137 | MB1 | CADM1 |
| PARK7 | 1.72E-07 | 0.376155 | 1 | 0.951 | 0.003345 | MB1 | PARK7 |
| ATP5C1 | 1.81E-07 | 0.331445 | 0.974 | 0.899 | 0.003529 | MB1 | ATP5C1 |
| COX171 | 1.83E-07 | 0.409457 | 1 | 0.758 | 0.00356 | MB1 | COX17 |
| MAPK101 | 1.83E-07 | 0.286582 | 0.795 | 0.365 | 0.003568 | MB1 | MAPK10 |

| | | | | | | | |
|---|---|---|---|---|---|---|---|
| SUGP2 | 2.04E-07 | 0.326255 | 0.821 | 0.393 | 0.003966 | MB1 | SUGP2 |
| TTC8 | 2.08E-07 | 0.279151 | 0.59 | 0.232 | 0.004049 | MB1 | TTC8 |
| ACO2 | 2.25E-07 | 0.420373 | 0.974 | 0.682 | 0.004372 | MB1 | ACO2 |
| MAP61 | 2.43E-07 | 0.299213 | 0.795 | 0.392 | 0.004734 | MB1 | MAP6 |
| PCNXL4 | 2.67E-07 | 0.302661 | 0.769 | 0.395 | 0.005188 | MB1 | PCNXL4 |
| PSMB5 | 2.67E-07 | 0.380923 | 1 | 0.926 | 0.005197 | MB1 | PSMB5 |
| PPP1R16A | 2.81E-07 | 0.314115 | 0.641 | 0.3 | 0.005469 | MB1 | PPP1R16A |
| UCP2 | 2.91E-07 | 0.477095 | 0.692 | 0.333 | 0.005667 | MB1 | UCP2 |
| RARRES2 | 2.92E-07 | 0.404934 | 0.692 | 0.318 | 0.005687 | MB1 | RARRES2 |
| SS18 | 2.98E-07 | 0.292129 | 0.769 | 0.403 | 0.0058 | MB1 | SS18 |
| CCP110 | 3.28E-07 | 0.337301 | 0.538 | 0.212 | 0.006382 | MB1 | CCP110 |
| SPINT21 | 3.36E-07 | 0.428505 | 0.949 | 0.752 | 0.006546 | MB1 | SPINT2 |
| WDR73 | 3.44E-07 | 0.298751 | 0.795 | 0.409 | 0.006683 | MB1 | WDR73 |
| ZNHIT1 | 3.55E-07 | 0.328589 | 0.974 | 0.876 | 0.006901 | MB1 | ZNHIT1 |
| GOT1 | 3.60E-07 | 0.307125 | 0.795 | 0.444 | 0.006995 | MB1 | GOT1 |
| MED25 | 3.81E-07 | 0.369101 | 0.795 | 0.411 | 0.007409 | MB1 | MED25 |
| PCSK1N1 | 3.90E-07 | 0.473379 | 0.974 | 0.817 | 0.007585 | MB1 | PCSK1N |
| ENDOG | 3.91E-07 | 0.340714 | 0.59 | 0.256 | 0.007597 | MB1 | ENDOG |
| POLR2L | 4.07E-07 | 0.400913 | 1 | 0.918 | 0.007911 | MB1 | POLR2L |
| PPP4R3A | 4.40E-07 | 0.541086 | 0.667 | 0.347 | 0.00855 | MB1 | PPP4R3A |
| TEX264 | 4.72E-07 | 0.311893 | 0.795 | 0.393 | 0.009187 | MB1 | TEX264 |
| IGFBP6 | 4.83E-07 | 0.314404 | 0.41 | 0.137 | 0.009391 | MB1 | IGFBP6 |
| COA1 | 4.88E-07 | 0.336974 | 0.974 | 0.779 | 0.009499 | MB1 | COA1 |
| TMBIM4 | 5.12E-07 | 0.378282 | 0.923 | 0.536 | 0.009951 | MB1 | TMBIM4 |
| H2AFV1 | 5.30E-07 | 0.373335 | 0.923 | 0.781 | 0.010312 | MB1 | H2AFV |
| EMC3 | 5.34E-07 | 0.372331 | 0.821 | 0.526 | 0.010386 | MB1 | EMC3 |
| DUSP14 | 5.48E-07 | 0.342024 | 0.667 | 0.312 | 0.010666 | MB1 | DUSP14 |
| TMEM261 | 5.68E-07 | 0.370235 | 0.949 | 0.708 | 0.011047 | MB1 | TMEM261 |
| CCT2 | 5.77E-07 | 0.380826 | 0.974 | 0.907 | 0.011227 | MB1 | CCT2 |
| SSRP12 | 5.80E-07 | 0.410618 | 0.923 | 0.723 | 0.01129 | MB1 | SSRP1 |
| C9orf72 | 5.83E-07 | 0.36632 | 0.718 | 0.347 | 0.011334 | MB1 | C9orf72 |
| ACYP1 | 6.20E-07 | 0.356077 | 0.769 | 0.415 | 0.012052 | MB1 | ACYP1 |
| MORF4L2: | 6.36E-07 | 0.340214 | 1 | 0.996 | 0.012364 | MB1 | MORF4L2 |
| GFOD21 | 6.58E-07 | 0.264064 | 0.59 | 0.244 | 0.012804 | MB1 | GFOD2 |
| MPC2 | 6.70E-07 | 0.46667 | 0.897 | 0.701 | 0.013039 | MB1 | MPC2 |
| CIR11 | 6.95E-07 | 0.358289 | 0.795 | 0.43 | 0.013521 | MB1 | CIR1 |
| CLDN5 | 7.22E-07 | 0.411981 | 0.641 | 0.279 | 0.014055 | MB1 | CLDN5 |
| C21orf2 | 7.27E-07 | 0.309943 | 0.744 | 0.396 | 0.014142 | MB1 | C21orf2 |
| MAP1A1 | 7.47E-07 | 0.418019 | 0.769 | 0.504 | 0.014527 | MB1 | MAP1A |
| DIXDC1 | 7.73E-07 | 0.272439 | 0.667 | 0.298 | 0.015031 | MB1 | DIXDC1 |

| | | | | | | | |
|---|---|---|---|---|---|---|---|
| CPNE2 | 8.06E-07 | 0.254269 | 0.615 | 0.265 | 0.015676 | MB1 | CPNE2 |
| HTRA1 | 8.11E-07 | 0.282516 | 0.615 | 0.267 | 0.015782 | MB1 | HTRA1 |
| PITPNA | 8.27E-07 | 0.302341 | 0.615 | 0.291 | 0.016088 | MB1 | PITPNA |
| RESP18 | 8.49E-07 | 0.931081 | 0.41 | 0.152 | 0.016514 | MB1 | RESP18 |
| BORCS7 | 8.82E-07 | 0.343809 | 0.846 | 0.531 | 0.017158 | MB1 | BORCS7 |
| ISYNA1 | 8.95E-07 | 0.435535 | 0.974 | 0.879 | 0.017417 | MB1 | ISYNA1 |
| PSMB1 | 9.10E-07 | 0.27112 | 1 | 0.965 | 0.017701 | MB1 | PSMB1 |
| MED31 | 9.68E-07 | 0.395383 | 0.769 | 0.399 | 0.018839 | MB1 | MED31 |
| PTN3 | 9.72E-07 | 0.4618 | 0.821 | 0.452 | 0.018907 | MB1 | PTN |
| YPEL5 | 1.02E-06 | 0.416012 | 0.949 | 0.689 | 0.019746 | MB1 | YPEL5 |
| HIST1H2A | 1.10E-06 | 0.456677 | 0.744 | 0.35 | 0.021333 | MB1 | HIST1H2AC |
| GPX7 | 1.12E-06 | 0.267729 | 0.564 | 0.229 | 0.021798 | MB1 | GPX7 |
| ADIPOR11 | 1.16E-06 | 0.384288 | 0.897 | 0.62 | 0.02255 | MB1 | ADIPOR1 |
| NDUFC2 | 1.25E-06 | 0.316918 | 1 | 0.947 | 0.02434 | MB1 | NDUFC2 |
| ATP11C | 1.31E-06 | 0.355303 | 0.692 | 0.337 | 0.025409 | MB1 | ATP11C |
| AGO2 | 1.45E-06 | 0.285508 | 0.615 | 0.265 | 0.028124 | MB1 | AGO2 |
| EBNA1BP2 | 1.54E-06 | 0.3063 | 0.897 | 0.579 | 0.029941 | MB1 | EBNA1BP2 |
| CALM31 | 1.56E-06 | 0.320887 | 0.974 | 0.943 | 0.03036 | MB1 | CALM3 |
| TXNDC12 | 1.60E-06 | 0.34053 | 0.974 | 0.727 | 0.031101 | MB1 | TXNDC12 |
| GCHFR | 1.65E-06 | 0.380586 | 0.923 | 0.572 | 0.032048 | MB1 | GCHFR |
| SNX3 | 1.66E-06 | 0.32971 | 1 | 0.911 | 0.032339 | MB1 | SNX3 |
| SMIM19 | 1.67E-06 | 0.356844 | 0.821 | 0.52 | 0.032557 | MB1 | SMIM19 |
| G6PC3 | 1.68E-06 | 0.394625 | 0.974 | 0.691 | 0.03262 | MB1 | G6PC3 |
| ARL13B | 1.71E-06 | 0.282251 | 0.615 | 0.282 | 0.033235 | MB1 | ARL13B |
| CARHSP1 | 1.77E-06 | 0.348885 | 0.974 | 0.733 | 0.034457 | MB1 | CARHSP1 |
| PAIP2 | 1.81E-06 | 0.363568 | 0.974 | 0.872 | 0.035165 | MB1 | PAIP2 |
| HSPB111 | 1.81E-06 | 0.607924 | 0.897 | 0.562 | 0.035268 | MB1 | HSPB11 |
| H2AFY1 | 1.83E-06 | 0.382508 | 0.974 | 0.876 | 0.03563 | MB1 | H2AFY |
| GTF2A2 | 1.87E-06 | 0.333539 | 1 | 0.777 | 0.036398 | MB1 | GTF2A2 |
| ZSCAN18 | 1.92E-06 | 0.337762 | 0.923 | 0.673 | 0.037365 | MB1 | ZSCAN18 |
| COX7A2 | 2.07E-06 | 0.340848 | 1 | 0.997 | 0.040242 | MB1 | COX7A2 |
| PPME1 | 2.13E-06 | 0.338631 | 0.769 | 0.399 | 0.041342 | MB1 | PPME1 |
| PEX2 | 2.15E-06 | 0.376896 | 0.897 | 0.643 | 0.041754 | MB1 | PEX2 |
| SNX17 | 2.32E-06 | 0.313128 | 0.897 | 0.556 | 0.045038 | MB1 | SNX17 |
| COL18A1 | 2.46E-06 | 0.252649 | 0.769 | 0.386 | 0.047926 | MB1 | COL18A1 |
| ATXN7L3B | 2.48E-06 | 0.294151 | 0.769 | 0.457 | 0.048172 | MB1 | ATXN7L3B |
| DYNLRB1 | 2.58E-06 | 0.328314 | 1 | 0.887 | 0.050149 | MB1 | DYNLRB1 |
| CCT7 | 2.62E-06 | 0.35352 | 1 | 0.867 | 0.050906 | MB1 | CCT7 |
| COX5B | 2.63E-06 | 0.272122 | 1 | 0.984 | 0.051178 | MB1 | COX5B |
| BOLA3 | 2.70E-06 | 0.300256 | 0.846 | 0.481 | 0.052553 | MB1 | BOLA3 |

| | | | | | | | |
|---|---|---|---|---|---|---|---|
| SHFM1 | 2.71E-06 | 0.338129 | 0.974 | 0.874 | 0.052665 | MB1 | SHFM1 |
| C11orf49 | 2.84E-06 | 0.334373 | 0.872 | 0.593 | 0.055329 | MB1 | C11orf49 |
| VRK11 | 2.93E-06 | 0.318196 | 0.641 | 0.32 | 0.05699 | MB1 | VRK1 |
| BCL2L12 | 3.07E-06 | 0.277928 | 0.641 | 0.303 | 0.059701 | MB1 | BCL2L12 |
| IDH3A | 3.07E-06 | 0.317816 | 0.641 | 0.309 | 0.059782 | MB1 | IDH3A |
| NELFE1 | 3.10E-06 | 0.351839 | 0.949 | 0.73 | 0.060303 | MB1 | NELFE |
| ANXA61 | 3.24E-06 | 0.281873 | 0.846 | 0.453 | 0.063112 | MB1 | ANXA6 |
| CHMP51 | 3.25E-06 | 0.331507 | 1 | 0.892 | 0.063162 | MB1 | CHMP5 |
| ERCC1 | 3.41E-06 | 0.357072 | 0.923 | 0.624 | 0.066266 | MB1 | ERCC1 |
| UBE2M | 3.43E-06 | 0.295828 | 0.692 | 0.362 | 0.06672 | MB1 | UBE2M |
| MEST | 3.52E-06 | 0.442226 | 1 | 0.935 | 0.068483 | MB1 | MEST |
| PIN1 | 3.69E-06 | 0.32259 | 0.974 | 0.789 | 0.071791 | MB1 | PIN1 |
| MTFP1 | 3.69E-06 | 0.275447 | 0.564 | 0.25 | 0.071805 | MB1 | MTFP1 |
| TMX21 | 3.81E-06 | 0.283263 | 0.872 | 0.548 | 0.074199 | MB1 | TMX2 |
| MRPL22 | 3.93E-06 | 0.436423 | 0.846 | 0.615 | 0.076419 | MB1 | MRPL22 |
| COPS8 | 4.04E-06 | 0.384779 | 1 | 0.807 | 0.078519 | MB1 | COPS8 |
| CHCHD6 | 4.04E-06 | 0.314084 | 0.923 | 0.644 | 0.07859 | MB1 | CHCHD6 |
| LINC0063: | 4.24E-06 | 0.302648 | 0.974 | 0.584 | 0.082405 | MB1 | LINC00632 |
| RFX3 | 4.28E-06 | 0.367851 | 0.615 | 0.309 | 0.083356 | MB1 | RFX3 |
| HIST1H2A | 4.33E-06 | 0.253405 | 0.41 | 0.149 | 0.084299 | MB1 | HIST1H2AE |
| PLD3 | 4.37E-06 | 0.373809 | 1 | 0.923 | 0.085018 | MB1 | PLD3 |
| C7orf55 | 4.38E-06 | 0.337891 | 0.795 | 0.553 | 0.085133 | MB1 | C7orf55 |
| WARS | 4.41E-06 | 0.259333 | 0.615 | 0.273 | 0.085778 | MB1 | WARS |
| PDLIM4 | 4.58E-06 | 0.311472 | 0.718 | 0.395 | 0.089046 | MB1 | PDLIM4 |
| TPR | 4.93E-06 | 0.302117 | 0.923 | 0.625 | 0.095824 | MB1 | TPR |
| SSNA1 | 5.26E-06 | 0.386448 | 0.872 | 0.673 | 0.102284 | MB1 | SSNA1 |
| KTN1 | 5.46E-06 | 0.354315 | 0.872 | 0.611 | 0.106133 | MB1 | KTN1 |
| VAMP7 | 5.49E-06 | 0.263973 | 0.615 | 0.266 | 0.106736 | MB1 | VAMP7 |
| C4orf271 | 5.69E-06 | 0.348142 | 0.846 | 0.532 | 0.110721 | MB1 | C4orf27 |
| CCDC82 | 5.85E-06 | 0.30822 | 0.692 | 0.349 | 0.113811 | MB1 | CCDC82 |
| SPRTN | 5.97E-06 | 0.320065 | 0.487 | 0.211 | 0.116214 | MB1 | SPRTN |
| CD47 | 6.04E-06 | 0.273346 | 0.718 | 0.395 | 0.117509 | MB1 | CD47 |
| MKKS | 6.52E-06 | 0.297584 | 0.872 | 0.543 | 0.126863 | MB1 | MKKS |
| HAGH | 7.14E-06 | 0.28211 | 0.872 | 0.568 | 0.138836 | MB1 | HAGH |
| GCC2 | 7.18E-06 | 0.25908 | 0.821 | 0.439 | 0.139723 | MB1 | GCC2 |
| CTSL1 | 7.42E-06 | 0.34398 | 0.872 | 0.56 | 0.144268 | MB1 | CTSL |
| NGFRAP1 | 7.49E-06 | 0.251694 | 1 | 0.99 | 0.145793 | MB1 | NGFRAP1 |
| OAZ21 | 7.64E-06 | 0.31831 | 1 | 0.807 | 0.148718 | MB1 | OAZ2 |
| GPAA1 | 7.73E-06 | 0.322147 | 0.872 | 0.653 | 0.150297 | MB1 | GPAA1 |
| COQ9 | 8.32E-06 | 0.290542 | 0.769 | 0.444 | 0.161952 | MB1 | COQ9 |

| | | | | | | | |
|---|---|---|---|---|---|---|---|
| IFI27L1 | 8.98E-06 | 0.301675 | 0.949 | 0.654 | 0.174715 | MB1 | IFI27L1 |
| ECEL1 | 1.02E-05 | 0.400092 | 0.872 | 0.567 | 0.198547 | MB1 | ECEL1 |
| CCDC91 | 1.04E-05 | 0.264058 | 0.615 | 0.275 | 0.202832 | MB1 | CCDC91 |
| RBM14 | 1.14E-05 | 0.341703 | 0.718 | 0.392 | 0.221932 | MB1 | RBM14 |
| RAB7A | 1.15E-05 | 0.276391 | 0.974 | 0.866 | 0.224107 | MB1 | RAB7A |
| H3F3B2 | 1.19E-05 | 0.488375 | 1 | 1 | 0.23095 | MB1 | H3F3B |
| DNAJB61 | 1.19E-05 | 0.297329 | 1 | 0.885 | 0.232407 | MB1 | DNAJB6 |
| CHCHD5 | 1.22E-05 | 0.300606 | 0.949 | 0.715 | 0.237064 | MB1 | CHCHD5 |
| SLC43A2 | 1.32E-05 | 0.302049 | 0.641 | 0.337 | 0.256302 | MB1 | SLC43A2 |
| MT-ND2 | 1.32E-05 | 0.461552 | 1 | 0.972 | 0.256978 | MB1 | MT-ND2 |
| SOD1 | 1.38E-05 | 0.258587 | 1 | 0.958 | 0.268718 | MB1 | SOD1 |
| RB1CC1 | 1.39E-05 | 0.316269 | 0.718 | 0.37 | 0.27049 | MB1 | RB1CC1 |
| GNPAT | 1.43E-05 | 0.298591 | 0.718 | 0.376 | 0.278276 | MB1 | GNPAT |
| MT-ND4 | 1.51E-05 | 0.352388 | 1 | 0.967 | 0.294609 | MB1 | MT-ND4 |
| HTR2C | 1.55E-05 | 0.353612 | 0.564 | 0.257 | 0.300888 | MB1 | HTR2C |
| AIG1 | 1.89E-05 | 0.264342 | 0.769 | 0.442 | 0.368146 | MB1 | AIG1 |
| SAP18 | 2.02E-05 | 0.252161 | 1 | 0.955 | 0.39236 | MB1 | SAP18 |
| UBB | 2.46E-05 | 0.316042 | 1 | 0.983 | 0.47872 | MB1 | UBB |
| CDC37L1 | 2.73E-05 | 0.264022 | 0.718 | 0.384 | 0.531833 | MB1 | CDC37L1 |
| ERGIC2 | 2.94E-05 | 0.276456 | 0.846 | 0.505 | 0.571388 | MB1 | ERGIC2 |
| PTOV1 | 3.09E-05 | 0.297295 | 0.949 | 0.86 | 0.601106 | MB1 | PTOV1 |
| NDUFB5 | 3.35E-05 | 0.350078 | 0.949 | 0.827 | 0.651294 | MB1 | NDUFB5 |
| C19orf70 | 3.39E-05 | 0.30499 | 0.949 | 0.718 | 0.660451 | MB1 | C19orf70 |
| FSTL11 | 3.40E-05 | 0.271201 | 0.667 | 0.361 | 0.662271 | MB1 | FSTL1 |
| CHMP2B1 | 3.55E-05 | 0.323037 | 0.846 | 0.593 | 0.690447 | MB1 | CHMP2B |
| CCT52 | 3.58E-05 | 0.326399 | 0.923 | 0.725 | 0.695727 | MB1 | CCT5 |
| MT-CYB1 | 3.81E-05 | 0.414904 | 1 | 0.965 | 0.740678 | MB1 | MT-CYB |
| ERI3 | 3.92E-05 | 0.274318 | 0.923 | 0.659 | 0.763057 | MB1 | ERI3 |
| RALBP1 | 4.04E-05 | 0.262628 | 0.795 | 0.464 | 0.785046 | MB1 | RALBP1 |
| TXN | 4.23E-05 | 0.351636 | 1 | 0.965 | 0.822478 | MB1 | TXN |
| AURKAIP1 | 4.31E-05 | 0.280785 | 1 | 0.898 | 0.837504 | MB1 | AURKAIP1 |
| GPN3 | 4.46E-05 | 0.252602 | 0.744 | 0.395 | 0.867925 | MB1 | GPN3 |
| ADI1 | 4.84E-05 | 0.300849 | 0.692 | 0.367 | 0.941626 | MB1 | ADI1 |
| GLRX3 | 4.93E-05 | 0.313486 | 0.846 | 0.621 | 0.958803 | MB1 | GLRX3 |
| WIPI2 | 5.00E-05 | 0.292115 | 0.897 | 0.641 | 0.972152 | MB1 | WIPI2 |
| PRR13 | 5.20E-05 | 0.345298 | 0.846 | 0.648 | 1 | MB1 | PRR13 |
| MRPS18C | 5.69E-05 | 0.291644 | 0.923 | 0.619 | 1 | MB1 | MRPS18C |
| LYRM1 | 5.79E-05 | 0.274469 | 0.692 | 0.344 | 1 | MB1 | LYRM1 |
| PPP1R15A | 5.87E-05 | 0.417715 | 0.897 | 0.683 | 1 | MB1 | PPP1R15A |
| MT-CO2 | 5.95E-05 | 0.413112 | 1 | 0.988 | 1 | MB1 | MT-CO2 |

| | | | | | | | |
|---|---|---|---|---|---|---|---|
| GNB5 | 6.18E-05 | 0.275286 | 0.615 | 0.346 | 1 | MB1 | GNB5 |
| C4orf3 | 6.51E-05 | 0.319476 | 0.974 | 0.871 | 1 | MB1 | C4orf3 |
| HS2ST1 | 7.11E-05 | 0.264311 | 0.564 | 0.271 | 1 | MB1 | HS2ST1 |
| NDUFA9 | 7.13E-05 | 0.298753 | 0.795 | 0.587 | 1 | MB1 | NDUFA9 |
| MED21 | 7.71E-05 | 0.28313 | 0.795 | 0.446 | 1 | MB1 | MED21 |
| MRPL20 | 9.43E-05 | 0.253314 | 1 | 0.769 | 1 | MB1 | MRPL20 |
| CNIH1 | 9.54E-05 | 0.251996 | 0.949 | 0.781 | 1 | MB1 | CNIH1 |
| COX6C | 9.60E-05 | 0.303871 | 0.974 | 0.983 | 1 | MB1 | COX6C |
| ARL3 | 0.000107 | 0.312245 | 0.949 | 0.764 | 1 | MB1 | ARL3 |
| CHKA | 0.000109 | 0.297531 | 0.692 | 0.383 | 1 | MB1 | CHKA |
| FAM118A | 0.000119 | 0.342303 | 0.744 | 0.507 | 1 | MB1 | FAM118A |
| DDX39A1 | 0.000132 | 0.314056 | 0.769 | 0.489 | 1 | MB1 | DDX39A |
| NUCB21 | 0.000137 | 0.327347 | 0.872 | 0.706 | 1 | MB1 | NUCB2 |
| JKAMP | 0.000137 | 0.294066 | 0.821 | 0.562 | 1 | MB1 | JKAMP |
| APOE | 0.000147 | 0.34505 | 0.974 | 0.853 | 1 | MB1 | APOE |
| WDR13 | 0.000149 | 0.302595 | 0.821 | 0.552 | 1 | MB1 | WDR13 |
| ATP6V1D2 | 0.000153 | 0.254028 | 0.949 | 0.743 | 1 | MB1 | ATP6V1D |
| NDUFV2 | 0.000154 | 0.256431 | 0.923 | 0.785 | 1 | MB1 | NDUFV2 |
| CHURC1 | 0.000158 | 0.376614 | 0.692 | 0.456 | 1 | MB1 | CHURC1 |
| AZI2 | 0.000166 | 0.262866 | 0.692 | 0.446 | 1 | MB1 | AZI2 |
| TCEA2 | 0.000169 | 0.280441 | 0.872 | 0.637 | 1 | MB1 | TCEA2 |
| SERPINF1 | 0.00018 | 0.264549 | 0.897 | 0.608 | 1 | MB1 | SERPINF1 |
| WBSCR22 | 0.000182 | 0.251573 | 0.897 | 0.692 | 1 | MB1 | WBSCR22 |
| UBXN4 | 0.000185 | 0.268296 | 0.949 | 0.827 | 1 | MB1 | UBXN4 |
| CCT4 | 0.000191 | 0.319221 | 0.949 | 0.919 | 1 | MB1 | CCT4 |
| PRKAR1A | 0.000201 | 0.353977 | 0.949 | 0.813 | 1 | MB1 | PRKAR1A |
| GNG5 | 0.000209 | 0.26878 | 1 | 0.909 | 1 | MB1 | GNG5 |
| ALDOA2 | 0.00021 | 0.37855 | 1 | 0.989 | 1 | MB1 | ALDOA |
| PTMA2 | 0.00022 | 0.408916 | 1 | 1 | 1 | MB1 | PTMA |
| PINK1 | 0.000235 | 0.25109 | 0.641 | 0.37 | 1 | MB1 | PINK1 |
| TAX1BP1 | 0.000238 | 0.254289 | 0.949 | 0.78 | 1 | MB1 | TAX1BP1 |
| PSMA41 | 0.000248 | 0.286986 | 1 | 0.935 | 1 | MB1 | PSMA4 |
| SLC39A3 | 0.00027 | 0.278642 | 0.795 | 0.502 | 1 | MB1 | SLC39A3 |
| PTBP11 | 0.000271 | 0.280499 | 0.846 | 0.606 | 1 | MB1 | PTBP1 |
| TSTD1 | 0.000271 | 0.313059 | 0.667 | 0.405 | 1 | MB1 | TSTD1 |
| CALY | 0.000292 | 0.391514 | 0.564 | 0.306 | 1 | MB1 | CALY |
| DECR11 | 0.000299 | 0.317735 | 0.846 | 0.56 | 1 | MB1 | DECR1 |
| SNRPE2 | 0.000311 | 0.290135 | 1 | 0.92 | 1 | MB1 | SNRPE |
| ARL2 | 0.000322 | 0.275629 | 0.897 | 0.765 | 1 | MB1 | ARL2 |
| USP8 | 0.000358 | 0.250487 | 0.744 | 0.425 | 1 | MB1 | USP8 |

| | | | | | | | |
|---|---|---|---|---|---|---|---|
| SLC36A4 | 0.000377 | 0.259214 | 0.692 | 0.404 | 1 | MB1 | SLC36A4 |
| CHMP4B | 0.000442 | 0.276378 | 0.923 | 0.634 | 1 | MB1 | CHMP4B |
| SSBP4 | 0.000447 | 0.251195 | 0.769 | 0.554 | 1 | MB1 | SSBP4 |
| CACYBP | 0.000562 | 0.283762 | 0.949 | 0.775 | 1 | MB1 | CACYBP |
| CHORDC1 | 0.000611 | 0.28051 | 0.718 | 0.417 | 1 | MB1 | CHORDC1 |
| STIP1 | 0.000636 | 0.389318 | 0.795 | 0.642 | 1 | MB1 | STIP1 |
| METRN2 | 0.000741 | 0.260902 | 0.923 | 0.717 | 1 | MB1 | METRN |
| CETN3 | 0.00076 | 0.297355 | 0.667 | 0.424 | 1 | MB1 | CETN3 |
| SPARC1 | 0.000981 | 0.265837 | 0.949 | 0.813 | 1 | MB1 | SPARC |
| TCP12 | 0.001084 | 0.278549 | 0.974 | 0.856 | 1 | MB1 | TCP1 |
| NDUFB1 | 0.00115 | 0.25167 | 0.923 | 0.76 | 1 | MB1 | NDUFB1 |
| UFC1 | 0.001162 | 0.276692 | 0.897 | 0.701 | 1 | MB1 | UFC1 |
| SIVA12 | 0.001218 | 0.295432 | 0.923 | 0.774 | 1 | MB1 | SIVA1 |
| MT-ND3 | 0.001411 | 0.314489 | 0.974 | 0.799 | 1 | MB1 | MT-ND3 |
| COX7B | 0.001449 | 0.28266 | 0.897 | 0.788 | 1 | MB1 | COX7B |
| DPH3 | 0.001803 | 0.259954 | 0.769 | 0.555 | 1 | MB1 | DPH3 |
| MT-CO1 | 0.001885 | 0.290195 | 1 | 0.993 | 1 | MB1 | MT-CO1 |
| C16orf13 | 0.001956 | 0.26239 | 0.949 | 0.784 | 1 | MB1 | C16orf13 |
| MT-ND1 | 0.002161 | 0.316797 | 1 | 0.927 | 1 | MB1 | MT-ND1 |
| LYPLA2 | 0.002249 | 0.252658 | 0.718 | 0.507 | 1 | MB1 | LYPLA2 |
| MT-CO31 | 0.002456 | 0.342773 | 1 | 0.983 | 1 | MB1 | MT-CO3 |
| ORC4 | 0.003038 | 0.263476 | 0.718 | 0.511 | 1 | MB1 | ORC4 |
| HSP90B11 | 0.00478 | 0.289545 | 1 | 0.909 | 1 | MB1 | HSP90B1 |
| GGH1 | 0.009156 | 0.259589 | 0.872 | 0.693 | 1 | MB1 | GGH |
| C11orf881 | 1.18E-180 | 0.976448 | 0.801 | 0.061 | 2.29E-176 | MB2 | C11orf88 |
| TEKT11 | 1.63E-179 | 0.67959 | 0.755 | 0.05 | 3.17E-175 | MB2 | TEKT1 |
| ROPN1L1 | 1.29E-163 | 0.869568 | 0.775 | 0.065 | 2.51E-159 | MB2 | ROPN1L |
| FAM166B1 | 7.56E-161 | 0.588493 | 0.728 | 0.055 | 1.47E-156 | MB2 | FAM166B |
| STOML31 | 1.16E-146 | 0.79602 | 0.563 | 0.031 | 2.26E-142 | MB2 | STOML3 |
| ANKRD661 | 1.54E-143 | 0.407169 | 0.47 | 0.017 | 3.00E-139 | MB2 | ANKRD66 |
| C11orf971 | 3.23E-142 | 0.540956 | 0.55 | 0.03 | 6.29E-138 | MB2 | C11orf97 |
| DYNLRB21 | 4.96E-138 | 0.732319 | 0.808 | 0.098 | 9.65E-134 | MB2 | DYNLRB2 |
| FAM216B1 | 4.27E-137 | 0.42024 | 0.483 | 0.021 | 8.30E-133 | MB2 | FAM216B |
| CFAP521 | 3.37E-136 | 0.545121 | 0.682 | 0.06 | 6.56E-132 | MB2 | CFAP52 |
| FAM183A1 | 4.06E-136 | 1.414784 | 0.921 | 0.163 | 7.89E-132 | MB2 | FAM183A |
| C1orf1941 | 1.12E-135 | 0.952164 | 0.887 | 0.134 | 2.19E-131 | MB2 | C1orf194 |
| SPAG61 | 1.28E-135 | 0.8596 | 0.901 | 0.139 | 2.48E-131 | MB2 | SPAG6 |
| CCDC1701 | 8.85E-134 | 0.537283 | 0.636 | 0.052 | 1.72E-129 | MB2 | CCDC170 |
| RSPH11 | 7.01E-130 | 1.21431 | 0.927 | 0.178 | 1.36E-125 | MB2 | RSPH1 |
| ZMYND10 | 2.25E-128 | 1.085644 | 0.914 | 0.16 | 4.37E-124 | MB2 | ZMYND10 |

| | | | | | | | |
|---|---|---|---|---|---|---|---|
| DNAAF11 | 1.00E-122 | 0.597321 | 0.609 | 0.053 | 1.95E-118 | MB2 | DNAAF1 |
| C20orf851 | 2.89E-121 | 0.697837 | 0.556 | 0.042 | 5.62E-117 | MB2 | C20orf85 |
| CAPSL1 | 1.23E-120 | 0.798383 | 0.808 | 0.116 | 2.38E-116 | MB2 | CAPSL |
| SPAG171 | 1.57E-119 | 0.468084 | 0.589 | 0.049 | 3.05E-115 | MB2 | SPAG17 |
| WDR381 | 2.67E-118 | 0.482002 | 0.536 | 0.04 | 5.19E-114 | MB2 | WDR38 |
| CFAP771 | 1.15E-116 | 0.826835 | 0.868 | 0.154 | 2.24E-112 | MB2 | CFAP77 |
| C9orf1351 | 6.22E-116 | 0.504504 | 0.563 | 0.046 | 1.21E-111 | MB2 | C9orf135 |
| DRC31 | 1.73E-115 | 0.504271 | 0.623 | 0.06 | 3.37E-111 | MB2 | DRC3 |
| TCTEX1D4 | 4.82E-115 | 0.503892 | 0.629 | 0.063 | 9.38E-111 | MB2 | TCTEX1D4 |
| TCTEX1D1 | 2.15E-114 | 0.673457 | 0.755 | 0.101 | 4.19E-110 | MB2 | TCTEX1D1 |
| SPEF11 | 2.22E-113 | 0.645087 | 0.781 | 0.115 | 4.32E-109 | MB2 | SPEF1 |
| SPAG81 | 5.08E-113 | 0.602171 | 0.762 | 0.106 | 9.89E-109 | MB2 | SPAG8 |
| FOLR11 | 4.24E-112 | 1.079087 | 0.828 | 0.145 | 8.25E-108 | MB2 | FOLR1 |
| CFAP451 | 2.04E-107 | 0.670024 | 0.728 | 0.102 | 3.96E-103 | MB2 | CFAP45 |
| DNAH91 | 4.23E-104 | 0.491128 | 0.589 | 0.061 | 8.22E-100 | MB2 | DNAH9 |
| RP11-356I | 9.22E-103 | 1.202945 | 0.576 | 0.062 | 1.79E-98 | MB2 | RP11-356K23.1 |
| RP11-295I | 1.66E-102 | 0.569685 | 0.576 | 0.06 | 3.24E-98 | MB2 | RP11-295M3.4 |
| ARMC31 | 2.79E-102 | 0.532054 | 0.695 | 0.094 | 5.42E-98 | MB2 | ARMC3 |
| CFAP1261 | 6.99E-102 | 1.369582 | 0.974 | 0.303 | 1.36E-97 | MB2 | CFAP126 |
| DRC11 | 7.37E-102 | 0.677803 | 0.762 | 0.12 | 1.43E-97 | MB2 | DRC1 |
| SLC13A41 | 5.38E-99 | 0.952791 | 0.788 | 0.145 | 1.05E-94 | MB2 | SLC13A4 |
| ENO41 | 2.15E-96 | 0.41718 | 0.589 | 0.066 | 4.19E-92 | MB2 | ENO4 |
| C9orf241 | 8.16E-96 | 1.645537 | 0.921 | 0.275 | 1.59E-91 | MB2 | C9orf24 |
| RPE651 | 2.25E-95 | 0.743539 | 0.583 | 0.07 | 4.38E-91 | MB2 | RPE65 |
| PRR291 | 2.26E-94 | 0.310479 | 0.477 | 0.04 | 4.40E-90 | MB2 | PRR29 |
| CCDC781 | 1.02E-93 | 0.565948 | 0.437 | 0.033 | 1.98E-89 | MB2 | CCDC78 |
| C5orf491 | 5.31E-93 | 1.17349 | 0.94 | 0.289 | 1.03E-88 | MB2 | C5orf49 |
| MAP3K19 | 6.86E-93 | 0.279031 | 0.43 | 0.031 | 1.34E-88 | MB2 | MAP3K19 |
| PIFO1 | 2.99E-91 | 1.427714 | 0.987 | 0.442 | 5.82E-87 | MB2 | PIFO |
| GSTA31 | 3.39E-89 | 0.386998 | 0.344 | 0.018 | 6.59E-85 | MB2 | GSTA3 |
| C9orf1161 | 8.40E-88 | 1.022488 | 0.934 | 0.314 | 1.63E-83 | MB2 | C9orf116 |
| C6orf1181 | 1.01E-87 | 0.534303 | 0.715 | 0.121 | 1.97E-83 | MB2 | C6orf118 |
| ODF3B1 | 2.96E-85 | 0.902624 | 0.887 | 0.246 | 5.76E-81 | MB2 | ODF3B |
| LRRC461 | 4.74E-84 | 0.419355 | 0.517 | 0.058 | 9.23E-80 | MB2 | LRRC46 |
| FAM92B1 | 8.01E-84 | 0.617205 | 0.642 | 0.103 | 1.56E-79 | MB2 | FAM92B |
| DNAH7 | 9.25E-84 | 0.309457 | 0.49 | 0.05 | 1.80E-79 | MB2 | DNAH7 |
| KIF91 | 2.82E-83 | 1.070552 | 0.934 | 0.347 | 5.49E-79 | MB2 | KIF9 |
| TPPP31 | 7.07E-83 | 1.612216 | 0.987 | 0.461 | 1.37E-78 | MB2 | TPPP3 |
| TTC291 | 8.92E-83 | 0.252187 | 0.371 | 0.025 | 1.73E-78 | MB2 | TTC29 |
| EFCAB12 | 3.31E-81 | 0.617382 | 0.735 | 0.14 | 6.44E-77 | MB2 | EFCAB1 |

| | | | | | | | |
|---|---|---|---|---|---|---|---|
| AKAP141 | 4.52E-81 | 0.35111 | 0.457 | 0.044 | 8.80E-77 | MB2 | AKAP14 |
| PACRG1 | 4.63E-80 | 0.590705 | 0.735 | 0.148 | 9.01E-76 | MB2 | PACRG |
| FANK11 | 8.91E-80 | 0.59388 | 0.715 | 0.139 | 1.73E-75 | MB2 | FANK1 |
| CFAP53 | 4.61E-78 | 0.405635 | 0.576 | 0.081 | 8.96E-74 | MB2 | CFAP53 |
| MORN51 | 3.06E-76 | 0.577631 | 0.735 | 0.15 | 5.95E-72 | MB2 | MORN5 |
| FAM81B1 | 1.49E-75 | 0.567522 | 0.689 | 0.129 | 2.90E-71 | MB2 | FAM81B |
| PPP1R421 | 3.80E-73 | 0.295609 | 0.483 | 0.058 | 7.40E-69 | MB2 | PPP1R42 |
| LRRC231 | 1.52E-72 | 0.772823 | 0.921 | 0.31 | 2.95E-68 | MB2 | LRRC23 |
| SPATA181 | 2.15E-72 | 0.319333 | 0.417 | 0.041 | 4.18E-68 | MB2 | SPATA18 |
| SLC4A5 | 3.00E-71 | 0.498041 | 0.656 | 0.122 | 5.83E-67 | MB2 | SLC4A5 |
| DNAH111 | 2.26E-70 | 0.267872 | 0.391 | 0.036 | 4.39E-66 | MB2 | DNAH11 |
| CCDC1461 | 7.71E-70 | 0.304713 | 0.417 | 0.044 | 1.50E-65 | MB2 | CCDC146 |
| CETN21 | 3.18E-67 | 0.937552 | 1 | 0.832 | 6.18E-63 | MB2 | CETN2 |
| TRPM31 | 4.34E-66 | 1.40219 | 0.901 | 0.344 | 8.44E-62 | MB2 | TRPM3 |
| DNALI11 | 5.34E-66 | 0.885134 | 0.94 | 0.47 | 1.04E-61 | MB2 | DNALI1 |
| UBXN111 | 5.66E-65 | 0.746124 | 0.901 | 0.325 | 1.10E-60 | MB2 | UBXN11 |
| C7orf571 | 8.90E-65 | 0.251293 | 0.384 | 0.039 | 1.73E-60 | MB2 | C7orf57 |
| OMG | 1.73E-64 | 0.38753 | 0.291 | 0.021 | 3.37E-60 | MB2 | OMG |
| FABP61 | 3.64E-64 | 0.416023 | 0.338 | 0.03 | 7.09E-60 | MB2 | FABP6 |
| SMIM5 | 3.16E-63 | 0.326565 | 0.437 | 0.055 | 6.14E-59 | MB2 | SMIM5 |
| SLC17A8 | 1.40E-62 | 0.396133 | 0.43 | 0.055 | 2.71E-58 | MB2 | SLC17A8 |
| CFAP461 | 1.32E-61 | 0.389702 | 0.596 | 0.112 | 2.57E-57 | MB2 | CFAP46 |
| EFHC11 | 3.99E-61 | 0.797431 | 0.94 | 0.41 | 7.75E-57 | MB2 | EFHC1 |
| CFAP70 | 1.35E-60 | 0.289237 | 0.49 | 0.074 | 2.63E-56 | MB2 | CFAP70 |
| MLF11 | 1.73E-60 | 0.854493 | 0.993 | 0.747 | 3.36E-56 | MB2 | MLF1 |
| CD302 | 7.53E-60 | 0.60568 | 0.636 | 0.136 | 1.47E-55 | MB2 | CD302 |
| C22orf151 | 2.21E-59 | 0.412345 | 0.543 | 0.097 | 4.29E-55 | MB2 | C22orf15 |
| CCDC1731 | 7.17E-58 | 0.504296 | 0.642 | 0.147 | 1.40E-53 | MB2 | CCDC173 |
| C1orf168 | 1.67E-57 | 0.40224 | 0.543 | 0.097 | 3.24E-53 | MB2 | C1orf168 |
| TCTN11 | 1.01E-56 | 0.661169 | 0.907 | 0.349 | 1.97E-52 | MB2 | TCTN1 |
| IQCG1 | 2.37E-56 | 0.447844 | 0.662 | 0.155 | 4.62E-52 | MB2 | IQCG |
| CEP1261 | 5.28E-56 | 0.417959 | 0.603 | 0.132 | 1.03E-51 | MB2 | CEP126 |
| MAPK151 | 7.96E-56 | 0.347283 | 0.53 | 0.096 | 1.55E-51 | MB2 | MAPK15 |
| GRAMD31 | 2.01E-55 | 0.438196 | 0.477 | 0.08 | 3.91E-51 | MB2 | GRAMD3 |
| HYDIN1 | 2.15E-55 | 0.333841 | 0.49 | 0.082 | 4.18E-51 | MB2 | HYDIN |
| IFT221 | 3.51E-55 | 0.695781 | 0.98 | 0.619 | 6.83E-51 | MB2 | IFT22 |
| NRAV1 | 5.00E-55 | 0.289714 | 0.483 | 0.079 | 9.73E-51 | MB2 | NRAV |
| SPAG11 | 1.02E-53 | 0.290479 | 0.483 | 0.082 | 1.98E-49 | MB2 | SPAG1 |
| SPATA171 | 7.42E-53 | 0.303614 | 0.517 | 0.095 | 1.44E-48 | MB2 | SPATA17 |
| CCDC1531 | 9.02E-53 | 0.441024 | 0.642 | 0.154 | 1.76E-48 | MB2 | CCDC153 |

| | | | | | | | |
|---|---|---|---|---|---|---|---|
| MDH1B1 | 1.25E-51 | 0.309235 | 0.483 | 0.086 | 2.44E-47 | MB2 | MDH1B |
| CFAP2211 | 2.56E-51 | 0.29285 | 0.477 | 0.082 | 4.98E-47 | MB2 | CFAP221 |
| DNAAF31 | 4.99E-51 | 0.495755 | 0.596 | 0.139 | 9.72E-47 | MB2 | DNAAF3 |
| RABL2A1 | 1.10E-50 | 0.481405 | 0.768 | 0.24 | 2.15E-46 | MB2 | RABL2A |
| TEKT21 | 2.79E-50 | 0.562627 | 0.702 | 0.203 | 5.42E-46 | MB2 | TEKT2 |
| BBOF11 | 4.47E-50 | 0.357883 | 0.609 | 0.139 | 8.70E-46 | MB2 | BBOF1 |
| WDR781 | 7.16E-50 | 0.252347 | 0.417 | 0.064 | 1.39E-45 | MB2 | WDR78 |
| CASC11 | 9.71E-50 | 0.267377 | 0.384 | 0.054 | 1.89E-45 | MB2 | CASC1 |
| HSBP11 | 2.27E-49 | 0.610917 | 1 | 0.98 | 4.42E-45 | MB2 | HSBP1 |
| PSENEN1 | 3.15E-49 | 0.708308 | 0.987 | 0.736 | 6.12E-45 | MB2 | PSENEN |
| SULF1 | 4.58E-49 | 0.473403 | 0.722 | 0.211 | 8.91E-45 | MB2 | SULF1 |
| FOXJ11 | 4.97E-49 | 0.67817 | 0.868 | 0.351 | 9.68E-45 | MB2 | FOXJ1 |
| AC007325 | 4.12E-48 | 0.671022 | 0.841 | 0.34 | 8.02E-44 | MB2 | AC007325.4 |
| RPGR1 | 4.17E-48 | 0.351268 | 0.649 | 0.164 | 8.11E-44 | MB2 | RPGR |
| PLTP2 | 7.72E-48 | 0.79349 | 0.98 | 0.745 | 1.50E-43 | MB2 | PLTP |
| SPA171 | 2.40E-47 | 0.422201 | 0.669 | 0.185 | 4.68E-43 | MB2 | SPA17 |
| CFAP651 | 3.14E-47 | 0.29784 | 0.417 | 0.069 | 6.11E-43 | MB2 | CFAP65 |
| HTR2C1 | 3.46E-47 | 1.160565 | 0.702 | 0.232 | 6.72E-43 | MB2 | HTR2C |
| B9D11 | 7.32E-47 | 0.579535 | 0.874 | 0.387 | 1.42E-42 | MB2 | B9D1 |
| ATP11C1 | 4.30E-46 | 0.66089 | 0.808 | 0.31 | 8.36E-42 | MB2 | ATP11C |
| FBXW91 | 2.13E-45 | 0.487715 | 0.801 | 0.314 | 4.15E-41 | MB2 | FBXW9 |
| C9orf91 | 2.51E-45 | 0.464154 | 0.768 | 0.253 | 4.88E-41 | MB2 | C9orf9 |
| ZBBX1 | 2.65E-45 | 0.261858 | 0.437 | 0.078 | 5.16E-41 | MB2 | ZBBX |
| ZNF4871 | 3.77E-45 | 0.271998 | 0.543 | 0.118 | 7.34E-41 | MB2 | ZNF487 |
| ANXA62 | 7.43E-45 | 0.586597 | 0.907 | 0.429 | 1.44E-40 | MB2 | ANXA6 |
| OSCP11 | 8.87E-45 | 0.455492 | 0.695 | 0.216 | 1.73E-40 | MB2 | OSCP1 |
| EFHC2 | 1.97E-44 | 0.285304 | 0.464 | 0.092 | 3.84E-40 | MB2 | EFHC2 |
| SPEF21 | 5.75E-44 | 0.315873 | 0.51 | 0.111 | 1.12E-39 | MB2 | SPEF2 |
| MORN21 | 8.15E-44 | 0.628646 | 0.921 | 0.524 | 1.58E-39 | MB2 | MORN2 |
| UQCR111 | 8.61E-44 | 0.525964 | 1 | 0.955 | 1.68E-39 | MB2 | UQCR11 |
| PPIL61 | 9.06E-44 | 0.468652 | 0.742 | 0.252 | 1.76E-39 | MB2 | PPIL6 |
| ZBED5-AS | 1.41E-43 | 0.327199 | 0.55 | 0.128 | 2.74E-39 | MB2 | ZBED5-AS1 |
| C14orf142 | 2.64E-43 | 0.458491 | 0.775 | 0.28 | 5.14E-39 | MB2 | C14orf142 |
| SOD3 | 3.82E-43 | 0.340522 | 0.424 | 0.079 | 7.43E-39 | MB2 | SOD3 |
| CCDC1571 | 9.40E-43 | 0.353936 | 0.563 | 0.136 | 1.83E-38 | MB2 | CCDC157 |
| ANXA12 | 1.28E-41 | 0.488035 | 0.311 | 0.042 | 2.49E-37 | MB2 | ANXA1 |
| GMPR | 1.09E-40 | 0.443987 | 0.649 | 0.195 | 2.12E-36 | MB2 | GMPR |
| TUBB4B2 | 2.07E-40 | 0.744143 | 0.993 | 0.838 | 4.03E-36 | MB2 | TUBB4B |
| ANXA22 | 2.18E-40 | 0.814656 | 0.921 | 0.538 | 4.25E-36 | MB2 | ANXA2 |
| MITF | 3.56E-40 | 0.413271 | 0.656 | 0.19 | 6.92E-36 | MB2 | MITF |

| | | | | | | | |
|---|---|---|---|---|---|---|---|
| DYX1C11 | 3.95E-40 | 0.408332 | 0.709 | 0.227 | 7.69E-36 | MB2 | DYX1C1 |
| MLLT12 | 4.44E-40 | 0.545111 | 0.914 | 0.455 | 8.64E-36 | MB2 | MLLT1 |
| CA2 | 5.09E-40 | 1.083807 | 0.907 | 0.517 | 9.89E-36 | MB2 | CA2 |
| RDH10 | 5.25E-40 | 0.299385 | 0.457 | 0.097 | 1.02E-35 | MB2 | RDH10 |
| C17orf971 | 6.92E-40 | 0.34771 | 0.536 | 0.13 | 1.35E-35 | MB2 | C17orf97 |
| SULT1A1 | 1.16E-39 | 0.27979 | 0.55 | 0.133 | 2.26E-35 | MB2 | SULT1A1 |
| IGFBP71 | 1.93E-39 | 0.874304 | 0.404 | 0.083 | 3.75E-35 | MB2 | IGFBP7 |
| PPP1R321 | 3.48E-39 | 0.286413 | 0.464 | 0.1 | 6.76E-35 | MB2 | PPP1R32 |
| SLC7A8 | 3.61E-39 | 0.343742 | 0.609 | 0.173 | 7.02E-35 | MB2 | SLC7A8 |
| TTR1 | 4.60E-39 | 1.0686 | 1 | 1 | 8.95E-35 | MB2 | TTR |
| ZFYVE16 | 1.04E-38 | 0.583104 | 0.821 | 0.379 | 2.02E-34 | MB2 | ZFYVE16 |
| FAM229B1 | 1.51E-38 | 0.670284 | 0.987 | 0.715 | 2.94E-34 | MB2 | FAM229B |
| CCDC1131 | 1.78E-38 | 0.280968 | 0.483 | 0.111 | 3.46E-34 | MB2 | CCDC113 |
| FAM227A1 | 2.02E-38 | 0.256272 | 0.55 | 0.137 | 3.92E-34 | MB2 | FAM227A |
| TAGLN21 | 6.27E-38 | 0.587748 | 0.947 | 0.601 | 1.22E-33 | MB2 | TAGLN2 |
| DPCD1 | 6.43E-38 | 0.623623 | 0.947 | 0.63 | 1.25E-33 | MB2 | DPCD |
| MRPS61 | 7.70E-38 | 0.683986 | 0.974 | 0.803 | 1.50E-33 | MB2 | MRPS6 |
| CCDC40 | 1.32E-37 | 0.261469 | 0.464 | 0.107 | 2.57E-33 | MB2 | CCDC40 |
| P4HTM1 | 2.26E-37 | 0.540105 | 0.927 | 0.499 | 4.40E-33 | MB2 | P4HTM |
| NUP62CL1 | 2.36E-37 | 0.328151 | 0.53 | 0.134 | 4.59E-33 | MB2 | NUP62CL |
| RBM47 | 3.97E-37 | 0.513517 | 0.762 | 0.302 | 7.72E-33 | MB2 | RBM47 |
| CEP831 | 3.97E-37 | 0.311864 | 0.556 | 0.151 | 7.73E-33 | MB2 | CEP83 |
| C21orf621 | 4.85E-37 | 0.306918 | 0.536 | 0.139 | 9.43E-33 | MB2 | C21orf62 |
| IFT431 | 5.09E-37 | 0.561381 | 0.921 | 0.553 | 9.90E-33 | MB2 | IFT43 |
| RSPH91 | 7.44E-37 | 0.453921 | 0.675 | 0.229 | 1.45E-32 | MB2 | RSPH9 |
| PALMD | 1.60E-36 | 0.294997 | 0.57 | 0.152 | 3.10E-32 | MB2 | PALMD |
| DYNC2H1 | 1.77E-36 | 0.26721 | 0.543 | 0.139 | 3.45E-32 | MB2 | DYNC2H1 |
| IGFBP61 | 1.81E-36 | 0.498471 | 0.477 | 0.119 | 3.51E-32 | MB2 | IGFBP6 |
| CCNO1 | 2.64E-36 | 0.416292 | 0.477 | 0.111 | 5.14E-32 | MB2 | CCNO |
| AKR7A21 | 2.84E-36 | 0.579539 | 0.967 | 0.756 | 5.53E-32 | MB2 | AKR7A2 |
| AKAP91 | 4.90E-36 | 0.56872 | 0.98 | 0.651 | 9.53E-32 | MB2 | AKAP9 |
| SLC16A10 | 9.18E-36 | 0.29693 | 0.43 | 0.095 | 1.79E-31 | MB2 | SLC16A10 |
| CLDN51 | 1.11E-35 | 0.806744 | 0.695 | 0.257 | 2.16E-31 | MB2 | CLDN5 |
| SYDE21 | 1.68E-35 | 0.272079 | 0.411 | 0.086 | 3.26E-31 | MB2 | SYDE2 |
| HIST1H2A | 3.07E-35 | 0.467271 | 0.51 | 0.129 | 5.96E-31 | MB2 | HIST1H2AE |
| CCDC241 | 4.59E-35 | 0.25189 | 0.503 | 0.128 | 8.93E-31 | MB2 | CCDC24 |
| SPAG161 | 8.89E-35 | 0.539716 | 0.921 | 0.576 | 1.73E-30 | MB2 | SPAG16 |
| LRTOMT1 | 1.07E-34 | 0.366392 | 0.583 | 0.181 | 2.09E-30 | MB2 | LRTOMT |
| TCEB21 | 1.78E-34 | 0.395067 | 1 | 0.994 | 3.45E-30 | MB2 | TCEB2 |
| KCNJ13 | 2.26E-34 | 0.424622 | 0.517 | 0.138 | 4.39E-30 | MB2 | KCNJ13 |

| | | | | | | | |
|---|---|---|---|---|---|---|---|
| ANXA111 | 2.44E-34 | 0.332943 | 0.642 | 0.211 | 4.74E-30 | MB2 | ANXA11 |
| TM7SF21 | 4.47E-34 | 0.574363 | 0.947 | 0.637 | 8.69E-30 | MB2 | TM7SF2 |
| RGCC1 | 6.27E-34 | 0.285838 | 0.344 | 0.063 | 1.22E-29 | MB2 | RGCC |
| PTRH11 | 7.46E-34 | 0.389912 | 0.689 | 0.253 | 1.45E-29 | MB2 | PTRH1 |
| DUSP181 | 1.35E-33 | 0.261238 | 0.536 | 0.146 | 2.62E-29 | MB2 | DUSP18 |
| PITPNM11 | 1.75E-33 | 0.362795 | 0.596 | 0.187 | 3.40E-29 | MB2 | PITPNM1 |
| SCAMP1-/ | 2.35E-33 | 0.276853 | 0.536 | 0.151 | 4.57E-29 | MB2 | SCAMP1-AS1 |
| SERINC5 | 3.22E-33 | 0.377219 | 0.576 | 0.171 | 6.27E-29 | MB2 | SERINC5 |
| GCHFR1 | 3.66E-33 | 0.587421 | 0.921 | 0.554 | 7.12E-29 | MB2 | GCHFR |
| TEX91 | 4.47E-33 | 0.318651 | 0.589 | 0.184 | 8.70E-29 | MB2 | TEX9 |
| COPRS1 | 5.41E-33 | 0.472646 | 0.901 | 0.526 | 1.05E-28 | MB2 | COPRS |
| CCDC74B1 | 1.36E-32 | 0.38897 | 0.603 | 0.196 | 2.64E-28 | MB2 | CCDC74B |
| SHTN1 | 1.79E-32 | 0.38858 | 0.715 | 0.268 | 3.47E-28 | MB2 | SHTN1 |
| HIST1H2B | 3.61E-32 | 0.257706 | 0.291 | 0.048 | 7.02E-28 | MB2 | HIST1H2BC |
| EPHX1 | 4.99E-32 | 0.332637 | 0.669 | 0.242 | 9.70E-28 | MB2 | EPHX1 |
| TMCO3 | 1.03E-31 | 0.586197 | 0.788 | 0.407 | 2.01E-27 | MB2 | TMCO3 |
| LINC0098 | 2.23E-31 | 0.466359 | 0.781 | 0.325 | 4.34E-27 | MB2 | LINC00982 |
| LRRIQ11 | 2.33E-31 | 0.297477 | 0.53 | 0.154 | 4.53E-27 | MB2 | LRRIQ1 |
| COL4A5 | 2.39E-31 | 0.435746 | 0.795 | 0.361 | 4.66E-27 | MB2 | COL4A5 |
| ADD31 | 2.82E-31 | 0.411277 | 0.801 | 0.34 | 5.49E-27 | MB2 | ADD3 |
| MDM11 | 8.00E-31 | 0.315969 | 0.629 | 0.213 | 1.56E-26 | MB2 | MDM1 |
| RP11-89K | 8.29E-31 | 0.320134 | 0.616 | 0.198 | 1.61E-26 | MB2 | RP11-89K21.1 |
| SLC5A3 | 1.23E-30 | 0.443314 | 0.742 | 0.334 | 2.39E-26 | MB2 | SLC5A3 |
| RBMS1 | 2.82E-30 | 0.357948 | 0.695 | 0.252 | 5.49E-26 | MB2 | RBMS1 |
| RAB11FIP | 4.71E-30 | 0.27971 | 0.517 | 0.156 | 9.17E-26 | MB2 | RAB11FIP4 |
| PHACTR2 | 5.42E-30 | 0.361649 | 0.556 | 0.181 | 1.05E-25 | MB2 | PHACTR2 |
| TCTEX1D2 | 5.80E-30 | 0.501896 | 0.954 | 0.666 | 1.13E-25 | MB2 | TCTEX1D2 |
| DMKN | 8.27E-30 | 0.349019 | 0.589 | 0.193 | 1.61E-25 | MB2 | DMKN |
| LRRC731 | 1.41E-29 | 0.394021 | 0.642 | 0.247 | 2.74E-25 | MB2 | LRRC73 |
| POLR2L1 | 2.94E-29 | 0.479882 | 0.993 | 0.915 | 5.73E-25 | MB2 | POLR2L |
| SYT17 | 3.10E-29 | 0.29529 | 0.55 | 0.171 | 6.04E-25 | MB2 | SYT17 |
| MOK1 | 3.31E-29 | 0.311754 | 0.603 | 0.203 | 6.44E-25 | MB2 | MOK |
| MYL5 | 3.67E-29 | 0.253381 | 0.536 | 0.165 | 7.14E-25 | MB2 | MYL5 |
| NQO11 | 4.65E-29 | 0.531645 | 0.854 | 0.425 | 9.05E-25 | MB2 | NQO1 |
| SPATA331 | 5.54E-29 | 0.356253 | 0.768 | 0.32 | 1.08E-24 | MB2 | SPATA33 |
| FUZ1 | 5.55E-29 | 0.379223 | 0.702 | 0.291 | 1.08E-24 | MB2 | FUZ |
| IGFBP51 | 1.09E-28 | 0.512214 | 0.854 | 0.444 | 2.12E-24 | MB2 | IGFBP5 |
| MMP2 | 1.14E-28 | 0.407617 | 0.55 | 0.181 | 2.21E-24 | MB2 | MMP2 |
| TMEM231 | 1.27E-28 | 0.392488 | 0.629 | 0.239 | 2.46E-24 | MB2 | TMEM231 |
| NME51 | 3.47E-28 | 0.294544 | 0.536 | 0.167 | 6.76E-24 | MB2 | NME5 |

| | | | | | | | |
|---|---|---|---|---|---|---|---|
| EFCAB21 | 5.77E-28 | 0.257921 | 0.477 | 0.138 | 1.12E-23 | MB2 | EFCAB2 |
| GALNT181 | 8.48E-28 | 0.276188 | 0.543 | 0.17 | 1.65E-23 | MB2 | GALNT18 |
| RP11-620. | 9.96E-28 | 0.453507 | 0.914 | 0.571 | 1.94E-23 | MB2 | RP11-620J15.3 |
| IQCK1 | 1.04E-27 | 0.408076 | 0.755 | 0.335 | 2.03E-23 | MB2 | IQCK |
| UQCR10 | 1.05E-27 | 0.374518 | 1 | 0.987 | 2.04E-23 | MB2 | UQCR10 |
| CBY11 | 1.22E-27 | 0.44378 | 0.861 | 0.503 | 2.38E-23 | MB2 | CBY1 |
| CFDP11 | 1.24E-27 | 0.61674 | 0.94 | 0.761 | 2.41E-23 | MB2 | CFDP1 |
| SYTL11 | 1.98E-27 | 0.316269 | 0.57 | 0.194 | 3.86E-23 | MB2 | SYTL1 |
| GSTK1 | 2.10E-27 | 0.424288 | 0.768 | 0.38 | 4.08E-23 | MB2 | GSTK1 |
| DPY301 | 2.71E-27 | 0.449111 | 0.927 | 0.709 | 5.27E-23 | MB2 | DPY30 |
| MT-CO21 | 4.13E-27 | 0.63464 | 0.987 | 0.988 | 8.03E-23 | MB2 | MT-CO2 |
| WFIKKN1: | 7.86E-27 | 0.312424 | 0.669 | 0.253 | 1.53E-22 | MB2 | WFIKKN1 |
| SOD11 | 7.99E-27 | 0.379203 | 1 | 0.956 | 1.56E-22 | MB2 | SOD1 |
| TSTD11 | 8.80E-27 | 0.480765 | 0.768 | 0.385 | 1.71E-22 | MB2 | TSTD1 |
| IFT461 | 1.32E-26 | 0.351199 | 0.629 | 0.254 | 2.56E-22 | MB2 | IFT46 |
| LGI1 | 2.62E-26 | 0.276673 | 0.563 | 0.186 | 5.09E-22 | MB2 | LGI1 |
| KRT8 | 2.67E-26 | 0.470633 | 0.854 | 0.469 | 5.20E-22 | MB2 | KRT8 |
| GCLM1 | 2.89E-26 | 0.348512 | 0.662 | 0.273 | 5.62E-22 | MB2 | GCLM |
| TMEM107 | 4.08E-26 | 0.364484 | 0.768 | 0.372 | 7.94E-22 | MB2 | TMEM107 |
| ATP1B11 | 5.49E-26 | 0.466845 | 0.901 | 0.605 | 1.07E-21 | MB2 | ATP1B1 |
| VGLL4 | 6.87E-26 | 0.445037 | 0.921 | 0.594 | 1.34E-21 | MB2 | VGLL4 |
| DNPH11 | 7.49E-26 | 0.440966 | 0.927 | 0.634 | 1.46E-21 | MB2 | DNPH1 |
| WDR341 | 7.63E-26 | 0.412432 | 0.841 | 0.462 | 1.48E-21 | MB2 | WDR34 |
| RABL2B1 | 8.30E-26 | 0.403285 | 0.722 | 0.333 | 1.62E-21 | MB2 | RABL2B |
| EVL1 | 1.20E-25 | 0.438714 | 0.781 | 0.396 | 2.34E-21 | MB2 | EVL |
| AGTRAP1 | 1.39E-25 | 0.407827 | 0.781 | 0.39 | 2.70E-21 | MB2 | AGTRAP |
| LYPLA21 | 1.46E-25 | 0.484409 | 0.828 | 0.489 | 2.84E-21 | MB2 | LYPLA2 |
| NFIA1 | 1.75E-25 | 0.26371 | 0.503 | 0.157 | 3.41E-21 | MB2 | NFIA |
| TBC1D9 | 2.15E-25 | 0.324191 | 0.576 | 0.215 | 4.19E-21 | MB2 | TBC1D9 |
| CLU2 | 3.21E-25 | 0.524589 | 1 | 0.923 | 6.24E-21 | MB2 | CLU |
| MT-ND41 | 4.46E-25 | 0.545281 | 0.98 | 0.967 | 8.69E-21 | MB2 | MT-ND4 |
| CYSTM11 | 4.49E-25 | 0.458314 | 0.94 | 0.752 | 8.73E-21 | MB2 | CYSTM1 |
| ZNF436-A | 5.06E-25 | 0.399229 | 0.762 | 0.36 | 9.85E-21 | MB2 | ZNF436-AS1 |
| NUDT7 | 6.46E-25 | 0.281584 | 0.497 | 0.167 | 1.26E-20 | MB2 | NUDT7 |
| TMEM261 | 6.86E-25 | 0.436568 | 0.94 | 0.696 | 1.33E-20 | MB2 | TMEM261 |
| CC2D2A1 | 7.13E-25 | 0.293832 | 0.576 | 0.211 | 1.39E-20 | MB2 | CC2D2A |
| CCDC74A: | 8.36E-25 | 0.32321 | 0.675 | 0.289 | 1.63E-20 | MB2 | CCDC74A |
| SIL1 | 9.75E-25 | 0.31586 | 0.702 | 0.296 | 1.90E-20 | MB2 | SIL1 |
| FSTL12 | 1.90E-24 | 0.331218 | 0.762 | 0.339 | 3.69E-20 | MB2 | FSTL1 |
| CST3 | 2.41E-24 | 0.406413 | 0.993 | 0.899 | 4.68E-20 | MB2 | CST3 |

| | | | | | | | |
|---|---|---|---|---|---|---|---|
| KRT18 | 2.54E-24 | 0.426815 | 0.748 | 0.35 | 4.94E-20 | MB2 | KRT18 |
| OSBPL1A | 3.13E-24 | 0.367988 | 0.755 | 0.359 | 6.08E-20 | MB2 | OSBPL1A |
| CNKSR3 | 5.28E-24 | 0.280821 | 0.457 | 0.145 | 1.03E-19 | MB2 | CNKSR3 |
| C21orf591 | 6.03E-24 | 0.378571 | 0.874 | 0.537 | 1.17E-19 | MB2 | C21orf59 |
| CCDC1891 | 9.52E-24 | 0.293089 | 0.47 | 0.152 | 1.85E-19 | MB2 | CCDC189 |
| CALM21 | 1.05E-23 | 0.399087 | 1 | 0.997 | 2.03E-19 | MB2 | CALM2 |
| DALRD31 | 1.43E-23 | 0.403644 | 0.901 | 0.571 | 2.78E-19 | MB2 | DALRD3 |
| PPIC | 1.55E-23 | 0.309259 | 0.616 | 0.24 | 3.02E-19 | MB2 | PPIC |
| C11orf741 | 2.00E-23 | 0.419024 | 0.841 | 0.486 | 3.88E-19 | MB2 | C11orf74 |
| ZNHIT21 | 2.13E-23 | 0.295384 | 0.642 | 0.262 | 4.14E-19 | MB2 | ZNHIT2 |
| PKIG1 | 2.81E-23 | 0.380515 | 0.841 | 0.502 | 5.47E-19 | MB2 | PKIG |
| DGCR6 | 3.19E-23 | 0.273346 | 0.57 | 0.205 | 6.20E-19 | MB2 | DGCR6 |
| HSPBP11 | 5.90E-23 | 0.361462 | 0.881 | 0.505 | 1.15E-18 | MB2 | HSPBP1 |
| ODF2L1 | 6.21E-23 | 0.353976 | 0.715 | 0.345 | 1.21E-18 | MB2 | ODF2L |
| SESN3 | 7.40E-23 | 0.358474 | 0.815 | 0.405 | 1.44E-18 | MB2 | SESN3 |
| GSTP1 | 7.84E-23 | 0.350128 | 1 | 0.994 | 1.53E-18 | MB2 | GSTP1 |
| H2AFJ1 | 8.46E-23 | 0.350439 | 0.781 | 0.383 | 1.65E-18 | MB2 | H2AFJ |
| TUSC31 | 8.70E-23 | 0.462406 | 0.94 | 0.716 | 1.69E-18 | MB2 | TUSC3 |
| COX6C1 | 8.91E-23 | 0.372875 | 1 | 0.982 | 1.73E-18 | MB2 | COX6C |
| MT-CO11 | 1.07E-22 | 0.450535 | 1 | 0.993 | 2.08E-18 | MB2 | MT-CO1 |
| TMEM671 | 1.20E-22 | 0.263738 | 0.589 | 0.23 | 2.33E-18 | MB2 | TMEM67 |
| ITM2B | 1.46E-22 | 0.401872 | 0.987 | 0.946 | 2.84E-18 | MB2 | ITM2B |
| MT-ND11 | 1.49E-22 | 0.563165 | 0.96 | 0.926 | 2.90E-18 | MB2 | MT-ND1 |
| ZDBF2 | 2.47E-22 | 0.332994 | 0.53 | 0.207 | 4.80E-18 | MB2 | ZDBF2 |
| MT-ND21 | 2.75E-22 | 0.576729 | 0.987 | 0.972 | 5.34E-18 | MB2 | MT-ND2 |
| GJA11 | 3.09E-22 | 0.389398 | 0.748 | 0.349 | 6.00E-18 | MB2 | GJA1 |
| LRRN2 | 4.03E-22 | 0.278389 | 0.616 | 0.239 | 7.84E-18 | MB2 | LRRN2 |
| CLDN3 | 4.35E-22 | 0.368994 | 0.709 | 0.325 | 8.46E-18 | MB2 | CLDN3 |
| NDUFB7 | 4.53E-22 | 0.371397 | 0.98 | 0.895 | 8.82E-18 | MB2 | NDUFB7 |
| FGFR2 | 6.66E-22 | 0.272766 | 0.609 | 0.248 | 1.30E-17 | MB2 | FGFR2 |
| S100A112 | 1.00E-21 | 0.423767 | 0.523 | 0.198 | 1.95E-17 | MB2 | S100A11 |
| BMP7 | 1.10E-21 | 0.420295 | 0.775 | 0.421 | 2.14E-17 | MB2 | BMP7 |
| CHCHD10 | 1.31E-21 | 0.486918 | 0.907 | 0.713 | 2.55E-17 | MB2 | CHCHD10 |
| BSG | 1.46E-21 | 0.315664 | 1 | 0.981 | 2.83E-17 | MB2 | BSG |
| DYNLL11 | 1.97E-21 | 0.454096 | 1 | 0.974 | 3.84E-17 | MB2 | DYNLL1 |
| EPHX2 | 2.25E-21 | 0.319138 | 0.742 | 0.376 | 4.38E-17 | MB2 | EPHX2 |
| CTD-2015 | 3.68E-21 | 0.252149 | 0.45 | 0.153 | 7.16E-17 | MB2 | CTD-2015H6.3 |
| SLIT2 | 4.24E-21 | 0.298125 | 0.642 | 0.263 | 8.24E-17 | MB2 | SLIT2 |
| NDUFB51 | 6.46E-21 | 0.356313 | 0.974 | 0.82 | 1.26E-16 | MB2 | NDUFB5 |
| MT-ND31 | 7.66E-21 | 0.491548 | 0.914 | 0.794 | 1.49E-16 | MB2 | MT-ND3 |

| | | | | | | | |
|---|---|---|---|---|---|---|---|
| COX5B1 | 8.33E-21 | 0.292387 | 1 | 0.983 | 1.62E-16 | MB2 | COX5B |
| SLC22A17 | 1.10E-20 | 0.42467 | 0.921 | 0.702 | 2.13E-16 | MB2 | SLC22A17 |
| CTBS1 | 1.12E-20 | 0.255711 | 0.642 | 0.264 | 2.18E-16 | MB2 | CTBS |
| ANKRD54[illegible] | 1.34E-20 | 0.295658 | 0.709 | 0.356 | 2.60E-16 | MB2 | ANKRD54 |
| RFX31 | 3.15E-20 | 0.297681 | 0.656 | 0.291 | 6.14E-16 | MB2 | RFX3 |
| WDR601 | 3.20E-20 | 0.396114 | 0.762 | 0.406 | 6.23E-16 | MB2 | WDR60 |
| LINC0046[illegible] | 3.73E-20 | 0.298133 | 0.649 | 0.293 | 7.26E-16 | MB2 | LINC00467 |
| DYNLT12 | 5.83E-20 | 0.390267 | 0.98 | 0.874 | 1.13E-15 | MB2 | DYNLT1 |
| PDLIM41 | 5.94E-20 | 0.331845 | 0.735 | 0.377 | 1.16E-15 | MB2 | PDLIM4 |
| SEPW1 | 6.40E-20 | 0.446157 | 0.947 | 0.831 | 1.24E-15 | MB2 | SEPW1 |
| RARRES21 | 6.42E-20 | 0.340413 | 0.662 | 0.301 | 1.25E-15 | MB2 | RARRES2 |
| SPINT22 | 7.95E-20 | 0.394346 | 0.94 | 0.742 | 1.55E-15 | MB2 | SPINT2 |
| CCDC1811 | 8.91E-20 | 0.261828 | 0.689 | 0.312 | 1.73E-15 | MB2 | CCDC181 |
| TPBG1 | 9.69E-20 | 0.344516 | 0.987 | 0.68 | 1.89E-15 | MB2 | TPBG |
| C19orf701 | 9.79E-20 | 0.346459 | 0.921 | 0.708 | 1.90E-15 | MB2 | C19orf70 |
| ATPIF1 | 1.17E-19 | 0.32291 | 1 | 0.956 | 2.28E-15 | MB2 | ATPIF1 |
| FTH1 | 1.47E-19 | 0.311685 | 1 | 1 | 2.86E-15 | MB2 | FTH1 |
| PGAP11 | 1.73E-19 | 0.271037 | 0.609 | 0.263 | 3.36E-15 | MB2 | PGAP1 |
| C5orf151 | 2.18E-19 | 0.336865 | 0.894 | 0.586 | 4.24E-15 | MB2 | C5orf15 |
| FXYD1 | 2.34E-19 | 0.389905 | 0.563 | 0.247 | 4.54E-15 | MB2 | FXYD1 |
| PRNP | 2.49E-19 | 0.496828 | 0.808 | 0.53 | 4.84E-15 | MB2 | PRNP |
| GALNT1 | 2.76E-19 | 0.250617 | 0.589 | 0.245 | 5.37E-15 | MB2 | GALNT1 |
| ORAI3 | 3.10E-19 | 0.300605 | 0.675 | 0.325 | 6.03E-15 | MB2 | ORAI3 |
| ZSCAN11 | 3.21E-19 | 0.286428 | 0.662 | 0.317 | 6.25E-15 | MB2 | ZSCAN1 |
| G6PC31 | 3.38E-19 | 0.361807 | 0.947 | 0.678 | 6.57E-15 | MB2 | G6PC3 |
| PRPS1 | 3.68E-19 | 0.489241 | 0.874 | 0.637 | 7.16E-15 | MB2 | PRPS1 |
| NDUFB11 | 4.64E-19 | 0.382341 | 0.927 | 0.752 | 9.03E-15 | MB2 | NDUFB1 |
| PTGR1 | 4.97E-19 | 0.302905 | 0.709 | 0.347 | 9.67E-15 | MB2 | PTGR1 |
| IFI27L11 | 5.00E-19 | 0.368104 | 0.901 | 0.642 | 9.72E-15 | MB2 | IFI27L1 |
| FAM118A[illegible] | 6.32E-19 | 0.362755 | 0.808 | 0.491 | 1.23E-14 | MB2 | FAM118A |
| VAT1L | 6.63E-19 | 0.352355 | 0.589 | 0.273 | 1.29E-14 | MB2 | VAT1L |
| WRB1 | 9.34E-19 | 0.336357 | 0.967 | 0.839 | 1.82E-14 | MB2 | WRB |
| RHBDD21 | 1.18E-18 | 0.363691 | 0.98 | 0.861 | 2.30E-14 | MB2 | RHBDD2 |
| ATP5G3 | 1.22E-18 | 0.296023 | 1 | 0.986 | 2.37E-14 | MB2 | ATP5G3 |
| PLD31 | 1.49E-18 | 0.436764 | 0.993 | 0.919 | 2.90E-14 | MB2 | PLD3 |
| HMGN3 | 2.28E-18 | 0.346547 | 0.987 | 0.933 | 4.44E-14 | MB2 | HMGN3 |
| SDC21 | 2.38E-18 | 0.274726 | 0.841 | 0.469 | 4.62E-14 | MB2 | SDC2 |
| COQ41 | 2.51E-18 | 0.341146 | 0.894 | 0.596 | 4.88E-14 | MB2 | COQ4 |
| SERPINF1 | 2.54E-18 | 0.537716 | 0.874 | 0.595 | 4.95E-14 | MB2 | SERPINF1 |
| WNK1 | 3.12E-18 | 0.354085 | 0.702 | 0.385 | 6.06E-14 | MB2 | WNK1 |

| | | | | | | | |
|---|---|---|---|---|---|---|---|
| KLHDC8B | 3.13E-18 | 0.389852 | 0.934 | 0.707 | 6.10E-14 | MB2 | KLHDC8B |
| CTXN11 | 4.12E-18 | 0.35033 | 0.742 | 0.41 | 8.02E-14 | MB2 | CTXN1 |
| PCBD1 | 4.52E-18 | 0.352594 | 0.894 | 0.608 | 8.80E-14 | MB2 | PCBD1 |
| PARD6B | 4.80E-18 | 0.254112 | 0.536 | 0.223 | 9.34E-14 | MB2 | PARD6B |
| OTX2 | 5.26E-18 | 0.292402 | 0.808 | 0.426 | 1.02E-13 | MB2 | OTX2 |
| BOLA31 | 6.33E-18 | 0.35342 | 0.775 | 0.467 | 1.23E-13 | MB2 | BOLA3 |
| CYBA1 | 6.51E-18 | 0.377241 | 0.861 | 0.576 | 1.27E-13 | MB2 | CYBA |
| TMEM179 | 8.06E-18 | 0.312315 | 0.795 | 0.456 | 1.57E-13 | MB2 | TMEM179B |
| MPC21 | 8.48E-18 | 0.346296 | 0.914 | 0.69 | 1.65E-13 | MB2 | MPC2 |
| CALY1 | 8.81E-18 | 0.327972 | 0.629 | 0.289 | 1.71E-13 | MB2 | CALY |
| POLR2I1 | 1.29E-17 | 0.359834 | 0.967 | 0.834 | 2.50E-13 | MB2 | POLR2I |
| MT-CO32 | 2.01E-17 | 0.457479 | 0.987 | 0.983 | 3.91E-13 | MB2 | MT-CO3 |
| MMP24-A | 2.05E-17 | 0.343767 | 0.768 | 0.44 | 3.98E-13 | MB2 | MMP24-AS1 |
| ABHD11 | 2.90E-17 | 0.262931 | 0.722 | 0.366 | 5.65E-13 | MB2 | ABHD11 |
| C21orf21 | 3.09E-17 | 0.25107 | 0.748 | 0.378 | 6.01E-13 | MB2 | C21orf2 |
| MSI21 | 3.16E-17 | 0.343602 | 0.887 | 0.586 | 6.14E-13 | MB2 | MSI2 |
| IFI27L2 | 4.04E-17 | 0.408344 | 0.934 | 0.792 | 7.85E-13 | MB2 | IFI27L2 |
| ACADVL1 | 4.17E-17 | 0.372088 | 0.934 | 0.678 | 8.12E-13 | MB2 | ACADVL |
| MRPS331 | 5.01E-17 | 0.335329 | 0.861 | 0.574 | 9.75E-13 | MB2 | MRPS33 |
| ADAMTS6 | 7.37E-17 | 0.258869 | 0.497 | 0.203 | 1.43E-12 | MB2 | ADAMTS6 |
| EEF1B2 | 9.47E-17 | 0.372587 | 0.993 | 0.949 | 1.84E-12 | MB2 | EEF1B2 |
| APRT | 9.60E-17 | 0.326455 | 0.914 | 0.65 | 1.87E-12 | MB2 | APRT |
| GAS61 | 9.69E-17 | 0.253905 | 0.589 | 0.267 | 1.88E-12 | MB2 | GAS6 |
| HIGD2A | 1.12E-16 | 0.338865 | 0.96 | 0.832 | 2.17E-12 | MB2 | HIGD2A |
| MRPL571 | 1.79E-16 | 0.323605 | 0.98 | 0.832 | 3.48E-12 | MB2 | MRPL57 |
| TMEM88 | 2.00E-16 | 0.313972 | 0.483 | 0.201 | 3.90E-12 | MB2 | TMEM88 |
| NME31 | 2.11E-16 | 0.357011 | 0.868 | 0.598 | 4.11E-12 | MB2 | NME3 |
| PKM | 2.52E-16 | 0.318666 | 1 | 0.988 | 4.90E-12 | MB2 | PKM |
| WDR541 | 2.59E-16 | 0.38329 | 0.934 | 0.755 | 5.03E-12 | MB2 | WDR54 |
| RBP11 | 2.69E-16 | 0.331854 | 0.901 | 0.651 | 5.23E-12 | MB2 | RBP1 |
| C20orf961 | 3.70E-16 | 0.255319 | 0.715 | 0.381 | 7.20E-12 | MB2 | C20orf96 |
| CCNA11 | 4.72E-16 | 0.255664 | 0.305 | 0.092 | 9.17E-12 | MB2 | CCNA1 |
| C4orf31 | 4.84E-16 | 0.38386 | 0.967 | 0.866 | 9.42E-12 | MB2 | C4orf3 |
| CDO1 | 5.62E-16 | 0.321399 | 0.887 | 0.608 | 1.09E-11 | MB2 | CDO1 |
| MT-CYB2 | 9.42E-16 | 0.420084 | 0.987 | 0.964 | 1.83E-11 | MB2 | MT-CYB |
| COX8A | 1.29E-15 | 0.266062 | 1 | 0.972 | 2.50E-11 | MB2 | COX8A |
| NDUFC21 | 1.34E-15 | 0.25522 | 0.987 | 0.945 | 2.60E-11 | MB2 | NDUFC2 |
| HIST1H1C | 1.43E-15 | 0.967531 | 0.682 | 0.407 | 2.79E-11 | MB2 | HIST1H1C |
| LMNA | 1.67E-15 | 0.309481 | 0.675 | 0.361 | 3.25E-11 | MB2 | LMNA |
| NDUFA3 | 1.97E-15 | 0.336649 | 0.98 | 0.798 | 3.82E-11 | MB2 | NDUFA3 |

| | | | | | | | |
|---|---|---|---|---|---|---|---|
| SERPINB6 | 2.20E-15 | 0.286157 | 0.715 | 0.403 | 4.29E-11 | MB2 | SERPINB6 |
| MTX1 | 2.35E-15 | 0.266834 | 0.795 | 0.465 | 4.58E-11 | MB2 | MTX1 |
| MRPS25 | 2.95E-15 | 0.251552 | 0.656 | 0.357 | 5.73E-11 | MB2 | MRPS25 |
| NDUFA81 | 3.04E-15 | 0.304273 | 0.874 | 0.654 | 5.91E-11 | MB2 | NDUFA8 |
| DECR12 | 3.89E-15 | 0.303116 | 0.861 | 0.544 | 7.56E-11 | MB2 | DECR1 |
| CEP411 | 3.98E-15 | 0.273698 | 0.589 | 0.289 | 7.74E-11 | MB2 | CEP41 |
| NDUFC1 | 6.13E-15 | 0.326589 | 0.954 | 0.787 | 1.19E-10 | MB2 | NDUFC1 |
| STUB1 | 6.19E-15 | 0.292036 | 0.901 | 0.669 | 1.20E-10 | MB2 | STUB1 |
| C5orf451 | 6.90E-15 | 0.254189 | 0.649 | 0.316 | 1.34E-10 | MB2 | C5orf45 |
| SSBP41 | 7.83E-15 | 0.314836 | 0.821 | 0.539 | 1.52E-10 | MB2 | SSBP4 |
| NDUFB2 | 8.84E-15 | 0.301407 | 0.98 | 0.918 | 1.72E-10 | MB2 | NDUFB2 |
| ARL4C | 1.01E-14 | 0.314245 | 0.675 | 0.376 | 1.97E-10 | MB2 | ARL4C |
| NUDC1 | 1.18E-14 | 0.328211 | 0.974 | 0.888 | 2.29E-10 | MB2 | NUDC |
| CADM12 | 1.48E-14 | 0.312172 | 0.861 | 0.548 | 2.88E-10 | MB2 | CADM1 |
| RUVBL11 | 1.62E-14 | 0.293509 | 0.821 | 0.504 | 3.16E-10 | MB2 | RUVBL1 |
| ECI1 | 1.77E-14 | 0.257617 | 0.861 | 0.554 | 3.45E-10 | MB2 | ECI1 |
| ZFP36L2 | 2.03E-14 | 0.360545 | 0.815 | 0.544 | 3.95E-10 | MB2 | ZFP36L2 |
| HACD32 | 2.17E-14 | 0.293968 | 0.967 | 0.859 | 4.22E-10 | MB2 | HACD3 |
| C12orf751 | 2.26E-14 | 0.329176 | 0.768 | 0.475 | 4.39E-10 | MB2 | C12orf75 |
| DRAM21 | 2.93E-14 | 0.301779 | 0.901 | 0.592 | 5.70E-10 | MB2 | DRAM2 |
| CIB13 | 3.09E-14 | 0.288837 | 0.98 | 0.734 | 6.01E-10 | MB2 | CIB1 |
| EGFL61 | 3.63E-14 | 0.335213 | 0.212 | 0.056 | 7.07E-10 | MB2 | EGFL6 |
| APOA1BP | 3.76E-14 | 0.289541 | 0.781 | 0.464 | 7.32E-10 | MB2 | APOA1BP |
| WLS1 | 4.06E-14 | 0.309462 | 0.947 | 0.602 | 7.89E-10 | MB2 | WLS |
| MAP1LC3/ | 4.67E-14 | 0.273245 | 0.755 | 0.437 | 9.09E-10 | MB2 | MAP1LC3A |
| C1QBP | 4.71E-14 | 0.271961 | 0.967 | 0.814 | 9.16E-10 | MB2 | C1QBP |
| ECEL11 | 5.83E-14 | 0.346631 | 0.854 | 0.553 | 1.13E-09 | MB2 | ECEL1 |
| ANAPC16 | 6.78E-14 | 0.277324 | 0.967 | 0.832 | 1.32E-09 | MB2 | ANAPC16 |
| APLP21 | 7.05E-14 | 0.310704 | 0.993 | 0.826 | 1.37E-09 | MB2 | APLP2 |
| NDUFV21 | 7.16E-14 | 0.293323 | 0.947 | 0.776 | 1.39E-09 | MB2 | NDUFV2 |
| TNNT11 | 7.40E-14 | 0.253097 | 0.675 | 0.356 | 1.44E-09 | MB2 | TNNT1 |
| TMBIM41 | 7.51E-14 | 0.293893 | 0.795 | 0.525 | 1.46E-09 | MB2 | TMBIM4 |
| HRSP12 | 9.00E-14 | 0.273406 | 0.762 | 0.454 | 1.75E-09 | MB2 | HRSP12 |
| PCP4 | 9.28E-14 | 0.458193 | 0.57 | 0.298 | 1.81E-09 | MB2 | PCP4 |
| COX7B1 | 1.03E-13 | 0.312934 | 0.94 | 0.78 | 2.00E-09 | MB2 | COX7B |
| GPC3 | 1.69E-13 | 0.288272 | 0.775 | 0.463 | 3.29E-09 | MB2 | GPC3 |
| UQCRB | 1.81E-13 | 0.279706 | 0.907 | 0.727 | 3.52E-09 | MB2 | UQCRB |
| IER3 | 1.85E-13 | 0.277196 | 0.49 | 0.221 | 3.61E-09 | MB2 | IER3 |
| CD991 | 1.99E-13 | 0.34384 | 0.967 | 0.793 | 3.87E-09 | MB2 | CD99 |
| SNX171 | 3.00E-13 | 0.259601 | 0.834 | 0.542 | 5.84E-09 | MB2 | SNX17 |

| | | | | | | | |
|---|---|---|---|---|---|---|---|
| GEM | 3.09E-13 | 0.253925 | 0.45 | 0.196 | 6.01E-09 | MB2 | GEM |
| ATP5I | 3.19E-13 | 0.260684 | 0.987 | 0.955 | 6.21E-09 | MB2 | ATP5I |
| RP13-436I | 3.89E-13 | 0.257799 | 0.47 | 0.217 | 7.57E-09 | MB2 | RP13-436F16.1 |
| CCDC51 | 4.04E-13 | 0.250093 | 0.649 | 0.352 | 7.86E-09 | MB2 | CCDC51 |
| FEZ2 | 5.13E-13 | 0.271008 | 0.808 | 0.515 | 9.99E-09 | MB2 | FEZ2 |
| TMEM134 | 5.19E-13 | 0.286356 | 0.821 | 0.557 | 1.01E-08 | MB2 | TMEM134 |
| MRPL28 | 5.21E-13 | 0.258477 | 0.967 | 0.759 | 1.01E-08 | MB2 | MRPL28 |
| LAMTOR4 | 5.40E-13 | 0.280619 | 0.967 | 0.855 | 1.05E-08 | MB2 | LAMTOR4 |
| MINOS11 | 6.49E-13 | 0.311098 | 0.94 | 0.772 | 1.26E-08 | MB2 | MINOS1 |
| ENSA | 6.72E-13 | 0.290143 | 0.868 | 0.656 | 1.31E-08 | MB2 | ENSA |
| TSC22D3 | 8.71E-13 | 0.284162 | 0.901 | 0.696 | 1.70E-08 | MB2 | TSC22D3 |
| METRN3 | 8.81E-13 | 0.329641 | 0.901 | 0.708 | 1.71E-08 | MB2 | METRN |
| YBX31 | 9.73E-13 | 0.330955 | 0.715 | 0.455 | 1.89E-08 | MB2 | YBX3 |
| TMEM230 | 1.85E-12 | 0.271693 | 0.94 | 0.761 | 3.60E-08 | MB2 | TMEM230 |
| RAMP2 | 2.00E-12 | 0.265195 | 0.464 | 0.222 | 3.89E-08 | MB2 | RAMP2 |
| MZT2A | 2.86E-12 | 0.264816 | 0.967 | 0.799 | 5.57E-08 | MB2 | MZT2A |
| AP006222. | 3.14E-12 | 0.252054 | 0.682 | 0.406 | 6.11E-08 | MB2 | AP006222.2 |
| TIMM8B | 3.27E-12 | 0.259557 | 0.94 | 0.772 | 6.37E-08 | MB2 | TIMM8B |
| LRPAP1 | 3.34E-12 | 0.275536 | 0.94 | 0.746 | 6.50E-08 | MB2 | LRPAP1 |
| AP2M11 | 3.44E-12 | 0.257326 | 0.98 | 0.903 | 6.70E-08 | MB2 | AP2M1 |
| IFITM31 | 3.55E-12 | 0.27566 | 0.861 | 0.674 | 6.90E-08 | MB2 | IFITM3 |
| EMC2 | 7.25E-12 | 0.263824 | 0.715 | 0.411 | 1.41E-07 | MB2 | EMC2 |
| SERF21 | 8.20E-12 | 0.307978 | 1 | 0.998 | 1.60E-07 | MB2 | SERF2 |
| BCAS4 | 1.15E-11 | 0.251767 | 0.722 | 0.44 | 2.24E-07 | MB2 | BCAS4 |
| DGCR6L | 1.37E-11 | 0.272203 | 0.881 | 0.649 | 2.67E-07 | MB2 | DGCR6L |
| ZSCAN181 | 1.87E-11 | 0.261771 | 0.907 | 0.662 | 3.65E-07 | MB2 | ZSCAN18 |
| TMEM14B | 2.01E-11 | 0.267064 | 0.974 | 0.822 | 3.90E-07 | MB2 | TMEM14B |
| KTN11 | 2.19E-11 | 0.266758 | 0.828 | 0.6 | 4.26E-07 | MB2 | KTN1 |
| IFT571 | 2.85E-11 | 0.257769 | 0.788 | 0.536 | 5.55E-07 | MB2 | IFT57 |
| CHID1 | 4.90E-11 | 0.261142 | 0.861 | 0.704 | 9.54E-07 | MB2 | CHID1 |
| RUVBL21 | 1.83E-10 | 0.26638 | 0.901 | 0.67 | 3.57E-06 | MB2 | RUVBL2 |
| CXCL14 | 2.31E-10 | 0.257945 | 0.576 | 0.317 | 4.50E-06 | MB2 | CXCL14 |
| AK11 | 2.80E-10 | 0.333649 | 0.815 | 0.611 | 5.45E-06 | MB2 | AK1 |
| MT-ATP61 | 3.58E-10 | 0.302823 | 0.96 | 0.907 | 6.97E-06 | MB2 | MT-ATP6 |
| MEST1 | 6.13E-10 | 0.254012 | 0.993 | 0.932 | 1.19E-05 | MB2 | MEST |
| HES1 | 6.39E-10 | 0.360146 | 0.934 | 0.734 | 1.24E-05 | MB2 | HES1 |
| COL6A11 | 9.07E-10 | 0.267157 | 0.834 | 0.561 | 1.76E-05 | MB2 | COL6A1 |
| IFT271 | 2.40E-09 | 0.256912 | 0.834 | 0.633 | 4.67E-05 | MB2 | IFT27 |
| TAGLN | 3.59E-09 | 0.618523 | 0.377 | 0.182 | 6.98E-05 | MB2 | TAGLN |
| TPM11 | 5.69E-09 | 0.415795 | 0.762 | 0.565 | 0.000111 | MB2 | TPM1 |

| | | | | | | | |
|---|---|---|---|---|---|---|---|
| MT-ND5 | 6.94E-09 | 0.29372 | 0.927 | 0.805 | 0.000135 | MB2 | MT-ND5 |
| ARL4A1 | 1.22E-08 | 0.281422 | 0.821 | 0.593 | 0.000237 | MB2 | ARL4A |
| HSPB112 | 2.85E-07 | 0.282733 | 0.795 | 0.552 | 0.005537 | MB2 | HSPB11 |
| P4HA12 | 0.000116 | 0.304936 | 0.775 | 0.624 | 1 | MB2 | P4HA1 |
| TK11 | 1.50E-110 | 0.793603 | 0.859 | 0.082 | 2.93E-106 | MB3 | TK1 |
| ASF1B2 | 3.78E-99 | 0.726572 | 0.821 | 0.085 | 7.36E-95 | MB3 | ASF1B |
| PKMYT1 | 2.93E-77 | 0.381912 | 0.577 | 0.048 | 5.70E-73 | MB3 | PKMYT1 |
| CDC451 | 2.64E-75 | 0.456335 | 0.654 | 0.064 | 5.14E-71 | MB3 | CDC45 |
| ZWINT1 | 1.59E-65 | 0.874928 | 0.91 | 0.174 | 3.09E-61 | MB3 | ZWINT |
| MCM101 | 1.22E-61 | 0.356277 | 0.577 | 0.061 | 2.37E-57 | MB3 | MCM10 |
| CENPU1 | 1.42E-61 | 0.573864 | 0.808 | 0.131 | 2.77E-57 | MB3 | CENPU |
| KIAA0101 | 1.35E-60 | 1.547897 | 0.987 | 0.265 | 2.63E-56 | MB3 | KIAA0101 |
| MYBL21 | 3.22E-60 | 0.477287 | 0.641 | 0.079 | 6.27E-56 | MB3 | MYBL2 |
| ATAD21 | 2.60E-59 | 0.483823 | 0.705 | 0.099 | 5.07E-55 | MB3 | ATAD2 |
| CDC61 | 1.03E-58 | 0.494964 | 0.628 | 0.08 | 2.00E-54 | MB3 | CDC6 |
| DSCC1 | 1.38E-57 | 0.279644 | 0.487 | 0.046 | 2.69E-53 | MB3 | DSCC1 |
| E2F1 | 9.14E-57 | 0.388194 | 0.551 | 0.063 | 1.78E-52 | MB3 | E2F1 |
| FAM64A1 | 1.27E-56 | 0.55097 | 0.769 | 0.121 | 2.48E-52 | MB3 | FAM64A |
| RRM21 | 1.38E-56 | 0.981671 | 0.718 | 0.116 | 2.68E-52 | MB3 | RRM2 |
| SPC251 | 1.96E-56 | 0.501172 | 0.679 | 0.092 | 3.81E-52 | MB3 | SPC25 |
| RMI21 | 5.53E-56 | 0.462143 | 0.641 | 0.085 | 1.07E-51 | MB3 | RMI2 |
| MELK2 | 3.84E-55 | 0.43075 | 0.679 | 0.096 | 7.48E-51 | MB3 | MELK |
| CENPK2 | 4.22E-53 | 0.531243 | 0.859 | 0.168 | 8.20E-49 | MB3 | CENPK |
| CLSPN1 | 1.81E-52 | 0.50461 | 0.731 | 0.12 | 3.51E-48 | MB3 | CLSPN |
| RAD51AP | 6.07E-52 | 0.3606 | 0.615 | 0.083 | 1.18E-47 | MB3 | RAD51AP1 |
| UHRF12 | 5.45E-51 | 0.527293 | 0.859 | 0.166 | 1.06E-46 | MB3 | UHRF1 |
| POC1A2 | 6.25E-51 | 0.373448 | 0.679 | 0.104 | 1.22E-46 | MB3 | POC1A |
| PSMC3IP | 1.25E-50 | 0.451895 | 0.718 | 0.125 | 2.44E-46 | MB3 | PSMC3IP |
| CENPM1 | 2.63E-50 | 0.542595 | 0.756 | 0.139 | 5.13E-46 | MB3 | CENPM |
| TCF19 | 8.00E-50 | 0.263046 | 0.474 | 0.05 | 1.56E-45 | MB3 | TCF19 |
| GTSE11 | 8.82E-50 | 0.45791 | 0.641 | 0.092 | 1.72E-45 | MB3 | GTSE1 |
| CHAF1A2 | 2.89E-48 | 0.420148 | 0.731 | 0.128 | 5.62E-44 | MB3 | CHAF1A |
| RTKN21 | 1.20E-47 | 0.370445 | 0.628 | 0.094 | 2.33E-43 | MB3 | RTKN2 |
| CCNE1 | 4.39E-47 | 0.278342 | 0.449 | 0.048 | 8.54E-43 | MB3 | CCNE1 |
| PBK2 | 6.11E-47 | 0.53126 | 0.705 | 0.12 | 1.19E-42 | MB3 | PBK |
| NUSAP11 | 8.50E-47 | 0.632964 | 0.821 | 0.168 | 1.65E-42 | MB3 | NUSAP1 |
| DHFR2 | 1.35E-46 | 0.744428 | 0.897 | 0.244 | 2.62E-42 | MB3 | DHFR |
| CDK12 | 3.32E-46 | 0.674381 | 0.91 | 0.221 | 6.45E-42 | MB3 | CDK1 |
| DTL1 | 7.44E-46 | 0.306803 | 0.551 | 0.074 | 1.45E-41 | MB3 | DTL |
| GINS22 | 1.24E-45 | 0.792638 | 0.949 | 0.283 | 2.41E-41 | MB3 | GINS2 |

| | | | | | | | |
|---|---|---|---|---|---|---|---|
| FANCA1 | 2.03E-45 | 0.304581 | 0.603 | 0.091 | 3.94E-41 | MB3 | FANCA |
| HELLS1 | 1.13E-44 | 0.512428 | 0.885 | 0.205 | 2.20E-40 | MB3 | HELLS |
| CDCA51 | 2.06E-44 | 0.374625 | 0.667 | 0.112 | 4.00E-40 | MB3 | CDCA5 |
| FANCI1 | 9.52E-44 | 0.335968 | 0.538 | 0.077 | 1.85E-39 | MB3 | FANCI |
| KIFC11 | 1.27E-43 | 0.332375 | 0.603 | 0.091 | 2.47E-39 | MB3 | KIFC1 |
| CCNA21 | 1.09E-42 | 0.474922 | 0.654 | 0.11 | 2.12E-38 | MB3 | CCNA2 |
| TYMS1 | 1.94E-42 | 1.187594 | 0.974 | 0.463 | 3.77E-38 | MB3 | TYMS |
| CENPQ | 1.97E-41 | 0.296395 | 0.705 | 0.128 | 3.83E-37 | MB3 | CENPQ |
| UBE2T1 | 3.53E-41 | 0.742822 | 0.897 | 0.262 | 6.87E-37 | MB3 | UBE2T |
| CDCA82 | 9.14E-40 | 0.289894 | 0.59 | 0.094 | 1.78E-35 | MB3 | CDCA8 |
| MCM41 | 6.53E-39 | 0.368087 | 0.705 | 0.143 | 1.27E-34 | MB3 | MCM4 |
| RFC31 | 1.98E-38 | 0.404117 | 0.731 | 0.156 | 3.86E-34 | MB3 | RFC3 |
| KIF232 | 2.14E-38 | 0.322617 | 0.577 | 0.096 | 4.17E-34 | MB3 | KIF23 |
| GMNN1 | 3.39E-38 | 0.574691 | 0.872 | 0.24 | 6.59E-34 | MB3 | GMNN |
| MCM51 | 4.87E-38 | 0.41741 | 0.692 | 0.14 | 9.48E-34 | MB3 | MCM5 |
| UBE2C2 | 5.74E-38 | 0.669154 | 0.859 | 0.24 | 1.12E-33 | MB3 | UBE2C |
| BIRC52 | 5.83E-38 | 0.711794 | 0.795 | 0.186 | 1.13E-33 | MB3 | BIRC5 |
| CENPW2 | 5.74E-37 | 0.606835 | 0.885 | 0.265 | 1.12E-32 | MB3 | CENPW |
| DNAJC91 | 8.17E-37 | 0.676792 | 0.962 | 0.374 | 1.59E-32 | MB3 | DNAJC9 |
| MCM21 | 3.99E-36 | 0.433208 | 0.756 | 0.172 | 7.77E-32 | MB3 | MCM2 |
| MAD2L12 | 1.73E-35 | 0.668377 | 1 | 0.367 | 3.36E-31 | MB3 | MAD2L1 |
| CKS1B2 | 2.01E-35 | 0.893021 | 1 | 0.535 | 3.91E-31 | MB3 | CKS1B |
| CHAF1B | 3.01E-35 | 0.257023 | 0.526 | 0.087 | 5.85E-31 | MB3 | CHAF1B |
| PCNA1 | 4.30E-35 | 0.915605 | 0.962 | 0.469 | 8.36E-31 | MB3 | PCNA |
| CENPN1 | 5.55E-35 | 0.32786 | 0.654 | 0.133 | 1.08E-30 | MB3 | CENPN |
| PRC11 | 2.91E-34 | 0.379619 | 0.679 | 0.146 | 5.65E-30 | MB3 | PRC1 |
| H2AFZ1 | 4.64E-34 | 0.949007 | 1 | 0.981 | 9.03E-30 | MB3 | H2AFZ |
| CDT12 | 5.10E-34 | 0.405856 | 0.705 | 0.161 | 9.93E-30 | MB3 | CDT1 |
| MCM32 | 5.55E-34 | 0.615329 | 0.91 | 0.307 | 1.08E-29 | MB3 | MCM3 |
| RRM11 | 5.70E-34 | 0.720444 | 0.962 | 0.396 | 1.11E-29 | MB3 | RRM1 |
| MKI671 | 7.75E-34 | 0.431119 | 0.526 | 0.089 | 1.51E-29 | MB3 | MKI67 |
| TACC32 | 6.50E-33 | 0.294145 | 0.551 | 0.1 | 1.26E-28 | MB3 | TACC3 |
| RFC21 | 9.63E-33 | 0.513861 | 0.846 | 0.261 | 1.87E-28 | MB3 | RFC2 |
| FBXO51 | 3.96E-32 | 0.42559 | 0.641 | 0.144 | 7.71E-28 | MB3 | FBXO5 |
| LIG11 | 1.04E-31 | 0.500532 | 0.808 | 0.243 | 2.02E-27 | MB3 | LIG1 |
| MCM72 | 1.26E-31 | 0.796839 | 0.923 | 0.385 | 2.44E-27 | MB3 | MCM7 |
| S100B | 8.52E-31 | 0.444045 | 0.731 | 0.192 | 1.66E-26 | MB3 | S100B |
| CENPH2 | 3.16E-30 | 0.58031 | 0.859 | 0.288 | 6.15E-26 | MB3 | CENPH |
| SMC42 | 3.45E-30 | 0.58644 | 0.846 | 0.275 | 6.70E-26 | MB3 | SMC4 |
| PRIM11 | 5.59E-30 | 0.306785 | 0.615 | 0.128 | 1.09E-25 | MB3 | PRIM1 |

| | | | | | | | |
|---|---|---|---|---|---|---|---|
| SMC21 | 5.86E-30 | 0.471062 | 0.756 | 0.216 | 1.14E-25 | MB3 | SMC2 |
| ORC61 | 3.47E-29 | 0.501811 | 0.859 | 0.292 | 6.74E-25 | MB3 | ORC6 |
| RNASEH2/ | 3.49E-29 | 0.698894 | 0.872 | 0.362 | 6.79E-25 | MB3 | RNASEH2A |
| POLA2 | 3.52E-29 | 0.265872 | 0.5 | 0.096 | 6.85E-25 | MB3 | POLA2 |
| MXD31 | 3.63E-29 | 0.286885 | 0.487 | 0.09 | 7.07E-25 | MB3 | MXD3 |
| DSN1 | 5.35E-29 | 0.301549 | 0.628 | 0.145 | 1.04E-24 | MB3 | DSN1 |
| SNRPB3 | 8.03E-29 | 0.629708 | 1 | 0.854 | 1.56E-24 | MB3 | SNRPB |
| TOP2A2 | 1.94E-28 | 0.582641 | 0.641 | 0.157 | 3.77E-24 | MB3 | TOP2A |
| MND11 | 2.44E-28 | 0.274263 | 0.385 | 0.06 | 4.74E-24 | MB3 | MND1 |
| HIST1H4C | 2.70E-28 | 1.439506 | 0.872 | 0.363 | 5.25E-24 | MB3 | HIST1H4C |
| CENPF2 | 5.49E-28 | 0.386613 | 0.667 | 0.166 | 1.07E-23 | MB3 | CENPF |
| DEK2 | 5.63E-28 | 0.692508 | 0.987 | 0.689 | 1.10E-23 | MB3 | DEK |
| RP11-227I | 5.85E-28 | 0.529471 | 0.641 | 0.163 | 1.14E-23 | MB3 | RP11-227H15.5 |
| TUBA1B2 | 8.09E-28 | 0.789961 | 1 | 0.986 | 1.57E-23 | MB3 | TUBA1B |
| IMPA2 | 1.24E-27 | 0.250632 | 0.603 | 0.134 | 2.41E-23 | MB3 | IMPA2 |
| BARD11 | 3.84E-27 | 0.376864 | 0.731 | 0.208 | 7.48E-23 | MB3 | BARD1 |
| HJURP1 | 5.20E-27 | 0.266614 | 0.436 | 0.077 | 1.01E-22 | MB3 | HJURP |
| FANCG1 | 6.21E-27 | 0.255805 | 0.577 | 0.129 | 1.21E-22 | MB3 | FANCG |
| HMGN22 | 1.16E-26 | 0.74927 | 1 | 0.978 | 2.25E-22 | MB3 | HMGN2 |
| PXMP21 | 2.48E-26 | 0.537422 | 0.846 | 0.322 | 4.83E-22 | MB3 | PXMP2 |
| C16orf59 | 2.75E-26 | 0.285328 | 0.513 | 0.11 | 5.35E-22 | MB3 | C16orf59 |
| WDR342 | 4.60E-26 | 0.623405 | 0.923 | 0.472 | 8.94E-22 | MB3 | WDR34 |
| RFC5 | 7.56E-26 | 0.358538 | 0.744 | 0.227 | 1.47E-21 | MB3 | RFC5 |
| NUF22 | 1.06E-25 | 0.261383 | 0.538 | 0.116 | 2.07E-21 | MB3 | NUF2 |
| BRI3BP | 1.37E-25 | 0.258029 | 0.487 | 0.103 | 2.66E-21 | MB3 | BRI3BP |
| POLD11 | 3.19E-25 | 0.266926 | 0.603 | 0.146 | 6.21E-21 | MB3 | POLD1 |
| TMEM106 | 3.63E-25 | 0.656471 | 0.923 | 0.514 | 7.07E-21 | MB3 | TMEM106C |
| CDCA31 | 4.08E-25 | 0.309194 | 0.423 | 0.079 | 7.94E-21 | MB3 | CDCA3 |
| HMGB12 | 7.56E-25 | 0.603767 | 1 | 0.981 | 1.47E-20 | MB3 | HMGB1 |
| TUBB2 | 2.97E-24 | 0.637909 | 1 | 0.991 | 5.78E-20 | MB3 | TUBB |
| GGCT1 | 3.21E-24 | 0.538138 | 0.987 | 0.529 | 6.25E-20 | MB3 | GGCT |
| PLS3 | 5.14E-24 | 0.919279 | 0.987 | 0.719 | 1.00E-19 | MB3 | PLS3 |
| AURKB1 | 9.22E-24 | 0.373176 | 0.487 | 0.105 | 1.79E-19 | MB3 | AURKB |
| DUT1 | 1.79E-23 | 0.711663 | 0.987 | 0.658 | 3.48E-19 | MB3 | DUT |
| NUDT12 | 2.80E-23 | 0.411098 | 0.846 | 0.323 | 5.45E-19 | MB3 | NUDT1 |
| MCM6 | 2.96E-23 | 0.269476 | 0.641 | 0.17 | 5.76E-19 | MB3 | MCM6 |
| COL4A6 | 3.22E-23 | 0.580202 | 0.936 | 0.459 | 6.26E-19 | MB3 | COL4A6 |
| C19orf481 | 3.25E-23 | 0.515402 | 0.872 | 0.406 | 6.33E-19 | MB3 | C19orf48 |
| PPP2R2B | 5.44E-23 | 0.598947 | 0.846 | 0.343 | 1.06E-18 | MB3 | PPP2R2B |
| RFC41 | 5.77E-23 | 0.382107 | 0.731 | 0.24 | 1.12E-18 | MB3 | RFC4 |

| | | | | | | | |
|---|---|---|---|---|---|---|---|
| USP12 | 5.90E-23 | 0.473957 | 0.846 | 0.338 | 1.15E-18 | MB3 | USP1 |
| TUBG11 | 6.73E-23 | 0.532638 | 0.962 | 0.506 | 1.31E-18 | MB3 | TUBG1 |
| MIS18BP1 | 6.99E-23 | 0.256787 | 0.5 | 0.111 | 1.36E-18 | MB3 | MIS18BP1 |
| TIMELESS | 2.21E-22 | 0.309368 | 0.628 | 0.18 | 4.29E-18 | MB3 | TIMELESS |
| KIF20B2 | 3.65E-22 | 0.273294 | 0.692 | 0.198 | 7.09E-18 | MB3 | KIF20B |
| NETO2 | 5.01E-22 | 0.318664 | 0.654 | 0.194 | 9.74E-18 | MB3 | NETO2 |
| CDK21 | 5.22E-22 | 0.288585 | 0.654 | 0.187 | 1.02E-17 | MB3 | CDK2 |
| LRR11 | 7.53E-22 | 0.262994 | 0.5 | 0.119 | 1.46E-17 | MB3 | LRR1 |
| LSM41 | 5.11E-21 | 0.444478 | 1 | 0.931 | 9.95E-17 | MB3 | LSM4 |
| CLN6 | 5.24E-21 | 0.266186 | 0.59 | 0.165 | 1.02E-16 | MB3 | CLN6 |
| NASP2 | 5.55E-21 | 0.543426 | 0.974 | 0.588 | 1.08E-16 | MB3 | NASP |
| CCDC342 | 8.30E-21 | 0.428536 | 0.885 | 0.397 | 1.61E-16 | MB3 | CCDC34 |
| ASRGL11 | 8.58E-21 | 0.469399 | 0.885 | 0.404 | 1.67E-16 | MB3 | ASRGL1 |
| PFN11 | 9.69E-21 | 0.440239 | 1 | 0.97 | 1.89E-16 | MB3 | PFN1 |
| SNRNP251 | 1.46E-20 | 0.460657 | 0.962 | 0.582 | 2.84E-16 | MB3 | SNRNP25 |
| ARL6IP61 | 1.46E-20 | 0.514608 | 0.923 | 0.504 | 2.84E-16 | MB3 | ARL6IP6 |
| VRK12 | 1.54E-20 | 0.420991 | 0.795 | 0.31 | 3.00E-16 | MB3 | VRK1 |
| TMEM972 | 1.97E-20 | 0.538218 | 0.949 | 0.529 | 3.83E-16 | MB3 | TMEM97 |
| CCDC109B | 3.33E-20 | 0.408693 | 0.756 | 0.284 | 6.49E-16 | MB3 | CCDC109B |
| HIRIP31 | 6.42E-20 | 0.332222 | 0.756 | 0.281 | 1.25E-15 | MB3 | HIRIP3 |
| FEN11 | 7.09E-20 | 0.477686 | 0.833 | 0.357 | 1.38E-15 | MB3 | FEN1 |
| PKM1 | 2.33E-19 | 0.579257 | 1 | 0.988 | 4.53E-15 | MB3 | PKM |
| CDKN31 | 3.27E-19 | 0.349918 | 0.513 | 0.145 | 6.36E-15 | MB3 | CDKN3 |
| H2AFX1 | 3.63E-19 | 0.548979 | 0.846 | 0.409 | 7.07E-15 | MB3 | H2AFX |
| RPA11 | 5.08E-19 | 0.412907 | 0.885 | 0.425 | 9.88E-15 | MB3 | RPA1 |
| RANBP12 | 7.04E-19 | 0.494352 | 1 | 0.943 | 1.37E-14 | MB3 | RANBP1 |
| CEP781 | 7.47E-19 | 0.272327 | 0.628 | 0.193 | 1.45E-14 | MB3 | CEP78 |
| TMEM98 | 7.82E-19 | 0.4926 | 0.987 | 0.669 | 1.52E-14 | MB3 | TMEM98 |
| SIGMAR1 | 1.03E-18 | 0.365897 | 0.846 | 0.369 | 2.00E-14 | MB3 | SIGMAR1 |
| EZH21 | 1.59E-18 | 0.329258 | 0.795 | 0.298 | 3.09E-14 | MB3 | EZH2 |
| TFDP11 | 1.66E-18 | 0.30997 | 0.795 | 0.305 | 3.22E-14 | MB3 | TFDP1 |
| LOXL1 | 1.82E-18 | 0.497972 | 0.897 | 0.437 | 3.54E-14 | MB3 | LOXL1 |
| RPA31 | 2.63E-18 | 0.452811 | 0.974 | 0.6 | 5.11E-14 | MB3 | RPA3 |
| DTYMK1 | 2.69E-18 | 0.522587 | 0.897 | 0.539 | 5.24E-14 | MB3 | DTYMK |
| WLS2 | 3.61E-18 | 0.623183 | 0.949 | 0.613 | 7.02E-14 | MB3 | WLS |
| CTNNAL11 | 4.62E-18 | 0.362729 | 0.821 | 0.353 | 8.98E-14 | MB3 | CTNNAL1 |
| HES11 | 8.04E-18 | 0.597618 | 1 | 0.738 | 1.56E-13 | MB3 | HES1 |
| MFGE8 | 8.65E-18 | 0.443079 | 0.833 | 0.406 | 1.68E-13 | MB3 | MFGE8 |
| DNMT11 | 1.28E-17 | 0.375712 | 0.872 | 0.397 | 2.49E-13 | MB3 | DNMT1 |
| TPX22 | 1.39E-17 | 0.348538 | 0.487 | 0.141 | 2.71E-13 | MB3 | TPX2 |

| | | | | | | | |
|---|---|---|---|---|---|---|---|
| ARL21 | 1.41E-17 | 0.501631 | 0.987 | 0.76 | 2.74E-13 | MB3 | ARL2 |
| PALMD1 | 1.70E-17 | 0.258003 | 0.551 | 0.166 | 3.30E-13 | MB3 | PALMD |
| HMGA22 | 1.87E-17 | 0.391568 | 0.846 | 0.376 | 3.63E-13 | MB3 | HMGA2 |
| EFHD1 | 2.13E-17 | 0.253888 | 0.59 | 0.185 | 4.14E-13 | MB3 | EFHD1 |
| NGFRAP1: | 2.18E-17 | 0.373187 | 1 | 0.99 | 4.24E-13 | MB3 | NGFRAP1 |
| GGH2 | 2.19E-17 | 0.415908 | 1 | 0.686 | 4.27E-13 | MB3 | GGH |
| STRA131 | 2.98E-17 | 0.541868 | 0.897 | 0.565 | 5.80E-13 | MB3 | STRA13 |
| DBI2 | 4.73E-17 | 0.485662 | 0.962 | 0.725 | 9.21E-13 | MB3 | DBI |
| POLR3K | 5.75E-17 | 0.370634 | 0.769 | 0.329 | 1.12E-12 | MB3 | POLR3K |
| GSTP11 | 8.25E-17 | 0.364619 | 1 | 0.994 | 1.60E-12 | MB3 | GSTP1 |
| NMU1 | 8.33E-17 | 0.582593 | 0.718 | 0.284 | 1.62E-12 | MB3 | NMU |
| RPL39L1 | 1.14E-16 | 0.313572 | 0.679 | 0.251 | 2.22E-12 | MB3 | RPL39L |
| TMEM237 | 1.15E-16 | 0.367746 | 0.795 | 0.338 | 2.23E-12 | MB3 | TMEM237 |
| TCEAL2 | 1.37E-16 | 0.373084 | 0.744 | 0.305 | 2.67E-12 | MB3 | TCEAL2 |
| H2AFY3 | 1.92E-16 | 0.423799 | 1 | 0.873 | 3.73E-12 | MB3 | H2AFY |
| SLC37A4 | 1.96E-16 | 0.309874 | 0.756 | 0.299 | 3.81E-12 | MB3 | SLC37A4 |
| COMMD4: | 2.96E-16 | 0.432299 | 0.808 | 0.397 | 5.76E-12 | MB3 | COMMD4 |
| SMCO4 | 3.56E-16 | 0.444774 | 0.782 | 0.367 | 6.92E-12 | MB3 | SMCO4 |
| RPA21 | 3.57E-16 | 0.444553 | 0.962 | 0.569 | 6.94E-12 | MB3 | RPA2 |
| FN3KRP | 3.99E-16 | 0.326748 | 0.744 | 0.308 | 7.76E-12 | MB3 | FN3KRP |
| DCK | 5.44E-16 | 0.384086 | 0.897 | 0.457 | 1.06E-11 | MB3 | DCK |
| CTGF | 6.95E-16 | 0.527974 | 0.859 | 0.416 | 1.35E-11 | MB3 | CTGF |
| SRSF22 | 7.53E-16 | 0.426192 | 1 | 0.883 | 1.47E-11 | MB3 | SRSF2 |
| AP2S1 | 7.56E-16 | 0.409822 | 1 | 0.892 | 1.47E-11 | MB3 | AP2S1 |
| IDH22 | 1.13E-15 | 0.477164 | 0.987 | 0.69 | 2.19E-11 | MB3 | IDH2 |
| SLC5A6 | 1.19E-15 | 0.387044 | 0.705 | 0.299 | 2.31E-11 | MB3 | SLC5A6 |
| NOTCH2 | 1.20E-15 | 0.250827 | 0.654 | 0.236 | 2.34E-11 | MB3 | NOTCH2 |
| ABHD2 | 1.21E-15 | 0.27836 | 0.615 | 0.213 | 2.35E-11 | MB3 | ABHD2 |
| CDCA41 | 1.22E-15 | 0.329569 | 0.641 | 0.239 | 2.38E-11 | MB3 | CDCA4 |
| IFITM32 | 1.45E-15 | 0.485373 | 0.987 | 0.675 | 2.81E-11 | MB3 | IFITM3 |
| CRABP2 | 1.46E-15 | 0.550052 | 1 | 0.933 | 2.84E-11 | MB3 | CRABP2 |
| PRTG | 1.72E-15 | 0.288212 | 0.731 | 0.285 | 3.36E-11 | MB3 | PRTG |
| RSPO11 | 1.96E-15 | 0.412216 | 0.872 | 0.402 | 3.81E-11 | MB3 | RSPO1 |
| COL4A51 | 2.17E-15 | 0.315594 | 0.846 | 0.374 | 4.23E-11 | MB3 | COL4A5 |
| HAT11 | 2.24E-15 | 0.371331 | 0.872 | 0.457 | 4.35E-11 | MB3 | HAT1 |
| CTPS11 | 2.54E-15 | 0.270865 | 0.705 | 0.266 | 4.93E-11 | MB3 | CTPS1 |
| SLC20A1 | 2.55E-15 | 0.2988 | 0.769 | 0.335 | 4.95E-11 | MB3 | SLC20A1 |
| DNPH12 | 2.66E-15 | 0.40229 | 0.962 | 0.642 | 5.17E-11 | MB3 | DNPH1 |
| SLIT21 | 3.39E-15 | 0.271146 | 0.705 | 0.273 | 6.60E-11 | MB3 | SLIT2 |
| MINOS12 | 3.49E-15 | 0.430352 | 0.962 | 0.777 | 6.78E-11 | MB3 | MINOS1 |

| | | | | | | | |
|---|---|---|---|---|---|---|---|
| GPC31 | 3.62E-15 | 0.542609 | 0.885 | 0.469 | 7.05E-11 | MB3 | GPC3 |
| BAMBI | 4.95E-15 | 0.349019 | 0.705 | 0.287 | 9.64E-11 | MB3 | BAMBI |
| NRM | 5.18E-15 | 0.261938 | 0.756 | 0.306 | 1.01E-10 | MB3 | NRM |
| ATP11C2 | 5.52E-15 | 0.314624 | 0.808 | 0.326 | 1.07E-10 | MB3 | ATP11C |
| ECEL12 | 6.96E-15 | 0.502258 | 0.974 | 0.558 | 1.35E-10 | MB3 | ECEL1 |
| GULP1 | 7.04E-15 | 0.403254 | 0.949 | 0.539 | 1.37E-10 | MB3 | GULP1 |
| EEF1B21 | 8.62E-15 | 0.344245 | 1 | 0.95 | 1.68E-10 | MB3 | EEF1B2 |
| PHGDH2 | 1.01E-14 | 0.478547 | 0.962 | 0.639 | 1.96E-10 | MB3 | PHGDH |
| PTTG12 | 1.50E-14 | 0.356284 | 0.885 | 0.538 | 2.93E-10 | MB3 | PTTG1 |
| KLHDC8B | 1.58E-14 | 0.467268 | 0.987 | 0.712 | 3.07E-10 | MB3 | KLHDC8B |
| EBP | 1.62E-14 | 0.415878 | 0.91 | 0.552 | 3.14E-10 | MB3 | EBP |
| SPATS2L | 1.76E-14 | 0.316519 | 0.756 | 0.334 | 3.42E-10 | MB3 | SPATS2L |
| TUBB4B3 | 1.89E-14 | 0.430887 | 0.987 | 0.843 | 3.67E-10 | MB3 | TUBB4B |
| CYR61 | 2.03E-14 | 0.608451 | 0.923 | 0.583 | 3.95E-10 | MB3 | CYR61 |
| HMGA12 | 2.37E-14 | 0.43268 | 0.987 | 0.909 | 4.61E-10 | MB3 | HMGA1 |
| RSPO2 | 2.60E-14 | 0.451711 | 0.936 | 0.489 | 5.06E-10 | MB3 | RSPO2 |
| DCTPP11 | 3.37E-14 | 0.326181 | 0.923 | 0.481 | 6.56E-10 | MB3 | DCTPP1 |
| PCP41 | 3.74E-14 | 0.590894 | 0.667 | 0.303 | 7.28E-10 | MB3 | PCP4 |
| HINT1 | 4.10E-14 | 0.268119 | 1 | 0.998 | 7.97E-10 | MB3 | HINT1 |
| RHOH | 4.29E-14 | 0.258892 | 0.564 | 0.202 | 8.35E-10 | MB3 | RHOH |
| TMEM147 | 4.53E-14 | 0.368691 | 1 | 0.866 | 8.81E-10 | MB3 | TMEM147 |
| ACOT71 | 5.55E-14 | 0.327333 | 0.872 | 0.438 | 1.08E-09 | MB3 | ACOT7 |
| NUCKS11 | 5.83E-14 | 0.363993 | 1 | 0.93 | 1.13E-09 | MB3 | NUCKS1 |
| PGRMC1 | 6.31E-14 | 0.543136 | 1 | 0.876 | 1.23E-09 | MB3 | PGRMC1 |
| CSE1L1 | 6.45E-14 | 0.304208 | 0.821 | 0.367 | 1.25E-09 | MB3 | CSE1L |
| SAE11 | 7.70E-14 | 0.398857 | 0.962 | 0.629 | 1.50E-09 | MB3 | SAE1 |
| GLO1 | 1.02E-13 | 0.399047 | 0.987 | 0.891 | 1.99E-09 | MB3 | GLO1 |
| MT-CYB3 | 1.34E-13 | 0.496848 | 1 | 0.965 | 2.60E-09 | MB3 | MT-CYB |
| TIMM101 | 1.72E-13 | 0.313966 | 0.897 | 0.501 | 3.35E-09 | MB3 | TIMM10 |
| POLR2L2 | 2.09E-13 | 0.397071 | 1 | 0.917 | 4.06E-09 | MB3 | POLR2L |
| PGAM1 | 2.10E-13 | 0.361103 | 1 | 0.927 | 4.09E-09 | MB3 | PGAM1 |
| MEA11 | 2.28E-13 | 0.362224 | 0.962 | 0.652 | 4.43E-09 | MB3 | MEA1 |
| RAN2 | 3.72E-13 | 0.294598 | 1 | 0.985 | 7.24E-09 | MB3 | RAN |
| ZDHHC12 | 3.91E-13 | 0.314609 | 0.654 | 0.296 | 7.62E-09 | MB3 | ZDHHC12 |
| NENF | 3.92E-13 | 0.318544 | 0.859 | 0.46 | 7.62E-09 | MB3 | NENF |
| SNRPD12 | 4.00E-13 | 0.342958 | 1 | 0.863 | 7.79E-09 | MB3 | SNRPD1 |
| PRPS11 | 4.03E-13 | 0.420499 | 0.962 | 0.642 | 7.85E-09 | MB3 | PRPS1 |
| TMEM160 | 4.06E-13 | 0.404363 | 0.962 | 0.734 | 7.91E-09 | MB3 | TMEM160 |
| LIM2 | 4.88E-13 | 0.254005 | 0.462 | 0.149 | 9.50E-09 | MB3 | LIM2 |
| PSMC31 | 5.02E-13 | 0.397186 | 0.962 | 0.738 | 9.77E-09 | MB3 | PSMC3 |

| | | | | | | | |
|---|---|---|---|---|---|---|---|
| IFI27L12 | 5.19E-13 | 0.37503 | 0.936 | 0.649 | 1.01E-08 | MB3 | IFI27L1 |
| FAAP20 | 5.95E-13 | 0.360023 | 0.962 | 0.621 | 1.16E-08 | MB3 | FAAP20 |
| BSG1 | 7.00E-13 | 0.307977 | 1 | 0.981 | 1.36E-08 | MB3 | BSG |
| SNRPE3 | 7.73E-13 | 0.364178 | 1 | 0.918 | 1.50E-08 | MB3 | SNRPE |
| LMNB11 | 8.93E-13 | 0.255145 | 0.808 | 0.359 | 1.74E-08 | MB3 | LMNB1 |
| PARP12 | 9.81E-13 | 0.423463 | 0.987 | 0.724 | 1.91E-08 | MB3 | PARP1 |
| UQCC22 | 1.03E-12 | 0.358946 | 0.974 | 0.737 | 2.01E-08 | MB3 | UQCC2 |
| HSPB113 | 1.20E-12 | 0.357413 | 0.897 | 0.556 | 2.34E-08 | MB3 | HSPB11 |
| SDC22 | 1.35E-12 | 0.315264 | 0.91 | 0.478 | 2.62E-08 | MB3 | SDC2 |
| NSMCE4A | 1.35E-12 | 0.308166 | 0.795 | 0.411 | 2.63E-08 | MB3 | NSMCE4A |
| NME42 | 1.46E-12 | 0.36669 | 0.949 | 0.691 | 2.84E-08 | MB3 | NME4 |
| STRADB | 1.71E-12 | 0.313618 | 0.833 | 0.44 | 3.32E-08 | MB3 | STRADB |
| YEATS4 | 1.71E-12 | 0.299085 | 0.897 | 0.485 | 3.33E-08 | MB3 | YEATS4 |
| GMPR1 | 1.76E-12 | 0.295476 | 0.551 | 0.213 | 3.43E-08 | MB3 | GMPR |
| PIN11 | 1.77E-12 | 0.396345 | 0.974 | 0.786 | 3.44E-08 | MB3 | PIN1 |
| TMPO1 | 1.78E-12 | 0.274097 | 0.821 | 0.392 | 3.47E-08 | MB3 | TMPO |
| ANAPC11 | 1.95E-12 | 0.350668 | 1 | 0.925 | 3.79E-08 | MB3 | ANAPC11 |
| BMP71 | 1.98E-12 | 0.323467 | 0.821 | 0.431 | 3.86E-08 | MB3 | BMP7 |
| BLOC1S11 | 2.15E-12 | 0.402277 | 0.987 | 0.655 | 4.18E-08 | MB3 | BLOC1S1 |
| KIF222 | 2.69E-12 | 0.287561 | 0.833 | 0.403 | 5.23E-08 | MB3 | KIF22 |
| MRPS62 | 2.82E-12 | 0.388565 | 1 | 0.807 | 5.48E-08 | MB3 | MRPS6 |
| AGTRAP2 | 3.16E-12 | 0.347869 | 0.782 | 0.402 | 6.16E-08 | MB3 | AGTRAP |
| ITM2B1 | 3.56E-12 | 0.451167 | 1 | 0.947 | 6.92E-08 | MB3 | ITM2B |
| TSPAN4 | 3.66E-12 | 0.393815 | 0.885 | 0.561 | 7.11E-08 | MB3 | TSPAN4 |
| CDK42 | 3.98E-12 | 0.378495 | 0.987 | 0.73 | 7.74E-08 | MB3 | CDK4 |
| SAC3D1 | 4.12E-12 | 0.336253 | 0.833 | 0.436 | 8.02E-08 | MB3 | SAC3D1 |
| LRRN11 | 4.18E-12 | 0.284906 | 0.821 | 0.382 | 8.13E-08 | MB3 | LRRN1 |
| CDO11 | 4.26E-12 | 0.463387 | 0.949 | 0.615 | 8.29E-08 | MB3 | CDO1 |
| RBBP42 | 4.44E-12 | 0.324983 | 0.987 | 0.788 | 8.64E-08 | MB3 | RBBP4 |
| CKAP21 | 4.59E-12 | 0.278477 | 0.59 | 0.234 | 8.93E-08 | MB3 | CKAP2 |
| RALY | 6.43E-12 | 0.343091 | 0.923 | 0.736 | 1.25E-07 | MB3 | RALY |
| TMEM881 | 7.87E-12 | 0.357404 | 0.526 | 0.208 | 1.53E-07 | MB3 | TMEM88 |
| CMC2 | 7.89E-12 | 0.294145 | 0.885 | 0.47 | 1.54E-07 | MB3 | CMC2 |
| LY6H | 7.95E-12 | 0.344851 | 0.769 | 0.392 | 1.55E-07 | MB3 | LY6H |
| SPATA332 | 8.01E-12 | 0.286532 | 0.705 | 0.337 | 1.56E-07 | MB3 | SPATA33 |
| CARHSP1 | 1.40E-11 | 0.354904 | 0.949 | 0.73 | 2.73E-07 | MB3 | CARHSP1 |
| HACD33 | 1.45E-11 | 0.346938 | 1 | 0.861 | 2.82E-07 | MB3 | HACD3 |
| ECHDC1 | 1.51E-11 | 0.294162 | 0.808 | 0.458 | 2.94E-07 | MB3 | ECHDC1 |
| PA2G41 | 1.69E-11 | 0.328722 | 0.974 | 0.801 | 3.28E-07 | MB3 | PA2G4 |
| ACAT22 | 1.72E-11 | 0.39421 | 0.897 | 0.552 | 3.34E-07 | MB3 | ACAT2 |

| | | | | | | | |
|---|---|---|---|---|---|---|---|
| TMEM107 | 1.83E-11 | 0.285313 | 0.769 | 0.385 | 3.56E-07 | MB3 | TMEM107 |
| MFAP21 | 2.33E-11 | 0.307505 | 0.833 | 0.438 | 4.53E-07 | MB3 | MFAP2 |
| ACYP11 | 2.93E-11 | 0.266988 | 0.782 | 0.408 | 5.70E-07 | MB3 | ACYP1 |
| CBX51 | 2.94E-11 | 0.376778 | 0.923 | 0.635 | 5.72E-07 | MB3 | CBX5 |
| RSPO3 | 3.24E-11 | 0.339947 | 0.487 | 0.181 | 6.31E-07 | MB3 | RSPO3 |
| PDLIM7 | 3.97E-11 | 0.357393 | 0.974 | 0.761 | 7.72E-07 | MB3 | PDLIM7 |
| ABHD12 | 4.22E-11 | 0.279041 | 0.705 | 0.337 | 8.21E-07 | MB3 | ABHD12 |
| PPP1CC2 | 4.23E-11 | 0.32026 | 1 | 0.782 | 8.23E-07 | MB3 | PPP1CC |
| PRDX21 | 4.83E-11 | 0.255179 | 1 | 0.986 | 9.40E-07 | MB3 | PRDX2 |
| GPAA11 | 6.34E-11 | 0.31709 | 0.949 | 0.647 | 1.23E-06 | MB3 | GPAA1 |
| NDUFA11 | 6.86E-11 | 0.336828 | 1 | 0.879 | 1.34E-06 | MB3 | NDUFA11 |
| HNRNPD2 | 7.12E-11 | 0.324587 | 1 | 0.817 | 1.38E-06 | MB3 | HNRNPD |
| SPARC2 | 8.54E-11 | 0.3966 | 1 | 0.809 | 1.66E-06 | MB3 | SPARC |
| ANP32B2 | 8.93E-11 | 0.318501 | 0.936 | 0.604 | 1.74E-06 | MB3 | ANP32B |
| CYB5B | 9.15E-11 | 0.341193 | 0.936 | 0.641 | 1.78E-06 | MB3 | CYB5B |
| UNG | 1.14E-10 | 0.336926 | 0.654 | 0.325 | 2.21E-06 | MB3 | UNG |
| HMGB21 | 1.15E-10 | 0.359149 | 0.987 | 0.884 | 2.25E-06 | MB3 | HMGB2 |
| SLBP | 1.32E-10 | 0.332785 | 0.872 | 0.532 | 2.56E-06 | MB3 | SLBP |
| PIM1 | 1.47E-10 | 0.252767 | 0.679 | 0.308 | 2.87E-06 | MB3 | PIM1 |
| PFN22 | 1.56E-10 | 0.312474 | 1 | 0.93 | 3.03E-06 | MB3 | PFN2 |
| MT-CO33 | 1.57E-10 | 0.360942 | 1 | 0.982 | 3.06E-06 | MB3 | MT-CO3 |
| COPRS2 | 1.66E-10 | 0.27729 | 0.897 | 0.539 | 3.23E-06 | MB3 | COPRS |
| COTL1 | 1.70E-10 | 0.309322 | 0.949 | 0.828 | 3.30E-06 | MB3 | COTL1 |
| CA21 | 1.99E-10 | 0.309109 | 0.872 | 0.531 | 3.88E-06 | MB3 | CA2 |
| TCTEX1D2 | 2.31E-10 | 0.315364 | 0.974 | 0.675 | 4.50E-06 | MB3 | TCTEX1D2 |
| TSPAN3 | 2.91E-10 | 0.281368 | 1 | 0.911 | 5.66E-06 | MB3 | TSPAN3 |
| SOST | 2.96E-10 | 0.266839 | 0.282 | 0.078 | 5.76E-06 | MB3 | SOST |
| APOE1 | 2.97E-10 | 0.540543 | 1 | 0.85 | 5.77E-06 | MB3 | APOE |
| TPM12 | 3.59E-10 | 0.377162 | 0.936 | 0.565 | 6.98E-06 | MB3 | TPM1 |
| VAT1L1 | 3.63E-10 | 0.272791 | 0.628 | 0.282 | 7.07E-06 | MB3 | VAT1L |
| POLD22 | 5.22E-10 | 0.30462 | 0.949 | 0.633 | 1.01E-05 | MB3 | POLD2 |
| SNRPG1 | 5.58E-10 | 0.267332 | 1 | 0.892 | 1.08E-05 | MB3 | SNRPG |
| PRDX3 | 6.13E-10 | 0.272851 | 0.987 | 0.807 | 1.19E-05 | MB3 | PRDX3 |
| ACAA2 | 6.52E-10 | 0.286524 | 0.821 | 0.464 | 1.27E-05 | MB3 | ACAA2 |
| RBP12 | 8.15E-10 | 0.344402 | 0.923 | 0.658 | 1.58E-05 | MB3 | RBP1 |
| RBBP71 | 8.67E-10 | 0.338463 | 0.974 | 0.783 | 1.69E-05 | MB3 | RBBP7 |
| RNASEH2( | 9.20E-10 | 0.259642 | 0.833 | 0.476 | 1.79E-05 | MB3 | RNASEH2C |
| APOA1BP1 | 9.99E-10 | 0.27022 | 0.821 | 0.472 | 1.94E-05 | MB3 | APOA1BP |
| PPP1CA | 1.91E-09 | 0.269901 | 0.987 | 0.789 | 3.71E-05 | MB3 | PPP1CA |
| CSRP21 | 1.95E-09 | 0.349809 | 0.923 | 0.652 | 3.80E-05 | MB3 | CSRP2 |

| | | | | | | | |
|---|---|---|---|---|---|---|---|
| FAM89A | 2.08E-09 | 0.549427 | 0.564 | 0.262 | 4.04E-05 | MB3 | FAM89A |
| NDUFB11 | 2.21E-09 | 0.260604 | 1 | 0.949 | 4.29E-05 | MB3 | NDUFB11 |
| SIVA13 | 2.35E-09 | 0.295381 | 0.987 | 0.769 | 4.57E-05 | MB3 | SIVA1 |
| KIF21A | 2.44E-09 | 0.277977 | 0.91 | 0.559 | 4.75E-05 | MB3 | KIF21A |
| RPN2 | 2.49E-09 | 0.271733 | 1 | 0.855 | 4.84E-05 | MB3 | RPN2 |
| TTR2 | 2.53E-09 | 0.643907 | 1 | 1 | 4.93E-05 | MB3 | TTR |
| FUS2 | 3.24E-09 | 0.277631 | 1 | 0.904 | 6.30E-05 | MB3 | FUS |
| NQO12 | 3.37E-09 | 0.293572 | 0.808 | 0.44 | 6.55E-05 | MB3 | NQO1 |
| METTL9 | 3.86E-09 | 0.283699 | 0.987 | 0.847 | 7.51E-05 | MB3 | METTL9 |
| MT-CO22 | 4.26E-09 | 0.322914 | 1 | 0.988 | 8.29E-05 | MB3 | MT-CO2 |
| EMX21 | 4.88E-09 | 0.288146 | 0.962 | 0.61 | 9.50E-05 | MB3 | EMX2 |
| UBE2S2 | 5.53E-09 | 0.367465 | 0.936 | 0.734 | 0.000108 | MB3 | UBE2S |
| ATP1B31 | 9.10E-09 | 0.297572 | 0.974 | 0.833 | 0.000177 | MB3 | ATP1B3 |
| PPIA2 | 9.17E-09 | 0.260316 | 1 | 0.979 | 0.000178 | MB3 | PPIA |
| PTMA3 | 1.12E-08 | 0.253287 | 1 | 1 | 0.000217 | MB3 | PTMA |
| TRPM32 | 1.29E-08 | 0.375073 | 0.692 | 0.369 | 0.00025 | MB3 | TRPM3 |
| KPNB11 | 1.63E-08 | 0.286501 | 0.987 | 0.771 | 0.000317 | MB3 | KPNB1 |
| PIGX | 1.64E-08 | 0.267961 | 0.795 | 0.477 | 0.000319 | MB3 | PIGX |
| CYBA2 | 1.71E-08 | 0.280904 | 0.897 | 0.584 | 0.000333 | MB3 | CYBA |
| SSRP13 | 1.90E-08 | 0.268448 | 0.949 | 0.719 | 0.000369 | MB3 | SSRP1 |
| CISD1 | 2.13E-08 | 0.266633 | 0.949 | 0.64 | 0.000415 | MB3 | CISD1 |
| GAMT | 2.23E-08 | 0.262557 | 0.846 | 0.522 | 0.000434 | MB3 | GAMT |
| PIH1D1 | 2.31E-08 | 0.284033 | 0.91 | 0.6 | 0.000449 | MB3 | PIH1D1 |
| NAE12 | 2.43E-08 | 0.256082 | 0.872 | 0.562 | 0.000472 | MB3 | NAE1 |
| GTF3A | 2.50E-08 | 0.25327 | 0.846 | 0.496 | 0.000487 | MB3 | GTF3A |
| NPC22 | 2.85E-08 | 0.307099 | 1 | 0.901 | 0.000555 | MB3 | NPC2 |
| CAMTA11 | 3.21E-08 | 0.26673 | 0.974 | 0.773 | 0.000624 | MB3 | CAMTA1 |
| SLC25A5 | 4.04E-08 | 0.262725 | 1 | 0.946 | 0.000785 | MB3 | SLC25A5 |
| C12orf752 | 4.36E-08 | 0.281778 | 0.782 | 0.484 | 0.000848 | MB3 | C12orf75 |
| H2AFV2 | 4.92E-08 | 0.272073 | 0.962 | 0.777 | 0.000957 | MB3 | H2AFV |
| HNRNPAB | 5.91E-08 | 0.256005 | 0.846 | 0.558 | 0.00115 | MB3 | HNRNPAB |
| MT-ATP62 | 6.42E-08 | 0.286009 | 0.974 | 0.908 | 0.001249 | MB3 | MT-ATP6 |
| PSIP12 | 6.79E-08 | 0.253521 | 1 | 0.79 | 0.001321 | MB3 | PSIP1 |
| RHBDD22 | 6.97E-08 | 0.260451 | 1 | 0.864 | 0.001355 | MB3 | RHBDD2 |
| EIF1AX1 | 7.25E-08 | 0.285645 | 0.962 | 0.791 | 0.00141 | MB3 | EIF1AX |
| MT-ND22 | 7.48E-08 | 0.252773 | 1 | 0.972 | 0.001455 | MB3 | MT-ND2 |
| BAD | 9.64E-08 | 0.273906 | 0.936 | 0.717 | 0.001875 | MB3 | BAD |
| CLDN52 | 1.05E-07 | 0.329473 | 0.564 | 0.275 | 0.002033 | MB3 | CLDN5 |
| LY6E | 1.13E-07 | 0.258589 | 0.769 | 0.462 | 0.002203 | MB3 | LY6E |
| FADS11 | 1.37E-07 | 0.354258 | 0.949 | 0.794 | 0.002656 | MB3 | FADS1 |

| | | | | | | | |
|---|---|---|---|---|---|---|---|
| PCSK1N2 | 1.44E-07 | 0.294619 | 1 | 0.814 | 0.002796 | MB3 | PCSK1N |
| H1FX1 | 1.57E-07 | 0.326712 | 0.974 | 0.779 | 0.003054 | MB3 | H1FX |
| ITGAE | 1.63E-07 | 0.259844 | 0.833 | 0.498 | 0.003179 | MB3 | ITGAE |
| FAM213A | 1.64E-07 | 0.281937 | 0.974 | 0.787 | 0.003185 | MB3 | FAM213A |
| PAICS2 | 1.77E-07 | 0.294922 | 0.885 | 0.682 | 0.003448 | MB3 | PAICS |
| MT-ND32 | 1.79E-07 | 0.297628 | 0.974 | 0.796 | 0.003483 | MB3 | MT-ND3 |
| PTOV11 | 1.79E-07 | 0.270542 | 0.974 | 0.858 | 0.003492 | MB3 | PTOV1 |
| EGR12 | 1.81E-07 | 0.262373 | 0.872 | 0.535 | 0.003527 | MB3 | EGR1 |
| STUB11 | 1.96E-07 | 0.257249 | 0.923 | 0.676 | 0.00382 | MB3 | STUB1 |
| C7orf501 | 2.04E-07 | 0.266991 | 0.962 | 0.81 | 0.003962 | MB3 | C7orf50 |
| MZT2A1 | 2.04E-07 | 0.284847 | 0.962 | 0.804 | 0.003972 | MB3 | MZT2A |
| PRKDC1 | 2.06E-07 | 0.25865 | 0.821 | 0.51 | 0.004006 | MB3 | PRKDC |
| LYPD11 | 2.25E-07 | 0.308329 | 0.718 | 0.409 | 0.004385 | MB3 | LYPD1 |
| SLC22A17 | 2.31E-07 | 0.268216 | 0.962 | 0.708 | 0.004484 | MB3 | SLC22A17 |
| DDT | 3.02E-07 | 0.261308 | 0.962 | 0.757 | 0.005876 | MB3 | DDT |
| FBL2 | 3.42E-07 | 0.261267 | 0.936 | 0.723 | 0.006661 | MB3 | FBL |
| ZFP36L21 | 3.84E-07 | 0.294085 | 0.833 | 0.552 | 0.00747 | MB3 | ZFP36L2 |
| TXNDC17 | 6.07E-07 | 0.288526 | 0.962 | 0.764 | 0.011808 | MB3 | TXNDC17 |
| FDPS2 | 6.78E-07 | 0.325383 | 1 | 0.849 | 0.013188 | MB3 | FDPS |
| MT-ND42 | 1.86E-06 | 0.287913 | 1 | 0.966 | 0.036162 | MB3 | MT-ND4 |
| MASP11 | 4.49E-06 | 0.270341 | 0.308 | 0.132 | 0.087274 | MB3 | MASP1 |
| CNTNAP2: | 4.83E-06 | 0.290142 | 0.974 | 0.787 | 0.093905 | MB3 | CNTNAP2 |
| TSPAN131 | 6.27E-06 | 0.306932 | 0.897 | 0.674 | 0.122024 | MB3 | TSPAN13 |
| HES43 | 8.45E-06 | 0.369599 | 0.974 | 0.76 | 0.164342 | MB3 | HES4 |
| HTR2C2 | 1.24E-05 | 0.365658 | 0.462 | 0.255 | 0.241216 | MB3 | HTR2C |
| SERPINF1 | 0.000139 | 0.251454 | 0.872 | 0.604 | 1 | MB3 | SERPINF1 |
| TTYH12 | 0.000152 | 0.380406 | 0.949 | 0.814 | 1 | MB3 | TTYH1 |
| LDHA1 | 0.001232 | 0.291443 | 0.808 | 0.595 | 1 | MB3 | LDHA |
| PLS31 | 1.05E-182 | 1.520152 | 0.994 | 0.659 | 2.05E-178 | MB4 | PLS3 |
| CRABP21 | 1.33E-172 | 1.049189 | 1 | 0.918 | 2.60E-168 | MB4 | CRABP2 |
| ECEL13 | 5.36E-150 | 0.972983 | 0.963 | 0.471 | 1.04E-145 | MB4 | ECEL1 |
| TTR3 | 3.70E-149 | 1.738435 | 1 | 1 | 7.19E-145 | MB4 | TTR |
| PKM2 | 1.65E-148 | 0.945353 | 1 | 0.985 | 3.21E-144 | MB4 | PKM |
| RSPO21 | 2.35E-141 | 0.985349 | 0.928 | 0.394 | 4.57E-137 | MB4 | RSPO2 |
| ITM2B2 | 7.34E-130 | 0.773744 | 0.996 | 0.937 | 1.43E-125 | MB4 | ITM2B |
| KLHDC8B | 3.09E-115 | 0.697196 | 0.969 | 0.657 | 6.01E-111 | MB4 | KLHDC8B |
| LINC0098: | 6.15E-107 | 0.636279 | 0.742 | 0.254 | 1.20E-102 | MB4 | LINC00982 |
| IGFBP52 | 4.06E-106 | 0.705765 | 0.866 | 0.368 | 7.91E-102 | MB4 | IGFBP5 |
| CA22 | 2.68E-104 | 0.919934 | 0.886 | 0.453 | 5.21E-100 | MB4 | CA2 |
| CLDN53 | 2.27E-103 | 1.069273 | 0.649 | 0.19 | 4.41E-99 | MB4 | CLDN5 |

| | | | | | | | |
|---|---|---|---|---|---|---|---|
| GULP11 | 2.92E-102 | 0.616027 | 0.901 | 0.462 | 5.69E-98 | MB4 | GULP1 |
| RP11-227I | 1.68E-97 | 0.557717 | 0.494 | 0.097 | 3.27E-93 | MB4 | RP11-227H15.5 |
| PPP2R2B1 | 1.30E-96 | 0.629423 | 0.731 | 0.263 | 2.53E-92 | MB4 | PPP2R2B |
| WLS3 | 2.39E-93 | 0.728638 | 0.921 | 0.547 | 4.65E-89 | MB4 | WLS |
| CXCL141 | 1.03E-92 | 0.874943 | 0.688 | 0.242 | 2.00E-88 | MB4 | CXCL14 |
| NDNF | 2.99E-91 | 0.446896 | 0.587 | 0.159 | 5.82E-87 | MB4 | NDNF |
| RSPO12 | 1.23E-89 | 0.64648 | 0.798 | 0.319 | 2.40E-85 | MB4 | RSPO1 |
| TPBG2 | 1.66E-87 | 0.629968 | 0.973 | 0.628 | 3.23E-83 | MB4 | TPBG |
| PALMD2 | 5.00E-85 | 0.338315 | 0.477 | 0.102 | 9.74E-81 | MB4 | PALMD |
| ATP11C3 | 1.09E-83 | 0.482938 | 0.709 | 0.247 | 2.12E-79 | MB4 | ATP11C |
| MITF1 | 4.22E-82 | 0.450133 | 0.533 | 0.139 | 8.21E-78 | MB4 | MITF |
| APOE2 | 1.39E-81 | 0.786187 | 0.988 | 0.821 | 2.70E-77 | MB4 | APOE |
| RP11-89K | 1.39E-81 | 0.491441 | 0.531 | 0.145 | 2.71E-77 | MB4 | RP11-89K21.1 |
| GPC32 | 1.51E-80 | 0.658574 | 0.816 | 0.397 | 2.94E-76 | MB4 | GPC3 |
| PCP42 | 3.36E-79 | 0.719656 | 0.653 | 0.227 | 6.53E-75 | MB4 | PCP4 |
| HTR2C3 | 1.36E-78 | 0.786171 | 0.585 | 0.178 | 2.65E-74 | MB4 | HTR2C |
| GSTP12 | 2.73E-77 | 0.407982 | 1 | 0.993 | 5.31E-73 | MB4 | GSTP1 |
| SPARC3 | 1.42E-76 | 0.562296 | 0.975 | 0.774 | 2.76E-72 | MB4 | SPARC |
| SIX3 | 3.41E-74 | 0.473367 | 0.715 | 0.292 | 6.64E-70 | MB4 | SIX3 |
| BMP72 | 5.81E-70 | 0.432462 | 0.781 | 0.356 | 1.13E-65 | MB4 | BMP7 |
| NR4A2 | 8.08E-68 | 0.347889 | 0.417 | 0.096 | 1.57E-63 | MB4 | NR4A2 |
| CDO12 | 5.12E-67 | 0.605653 | 0.857 | 0.566 | 9.96E-63 | MB4 | CDO1 |
| LMX1A | 6.94E-67 | 0.335447 | 0.519 | 0.156 | 1.35E-62 | MB4 | LMX1A |
| HES12 | 1.79E-66 | 0.724053 | 0.95 | 0.694 | 3.48E-62 | MB4 | HES1 |
| PDGFRB | 3.43E-66 | 0.295991 | 0.397 | 0.088 | 6.67E-62 | MB4 | PDGFRB |
| LYPD12 | 6.32E-66 | 0.59273 | 0.727 | 0.339 | 1.23E-61 | MB4 | LYPD1 |
| RBM471 | 2.79E-64 | 0.397979 | 0.643 | 0.25 | 5.42E-60 | MB4 | RBM47 |
| SERPINF1 | 6.40E-64 | 0.741676 | 0.853 | 0.551 | 1.24E-59 | MB4 | SERPINF1 |
| FGFR21 | 1.99E-63 | 0.375094 | 0.566 | 0.195 | 3.87E-59 | MB4 | FGFR2 |
| TRPM33 | 4.13E-63 | 0.793058 | 0.682 | 0.301 | 8.04E-59 | MB4 | TRPM3 |
| GDF5 | 2.64E-61 | 0.266293 | 0.347 | 0.071 | 5.13E-57 | MB4 | GDF5 |
| EMX22 | 4.60E-61 | 0.460242 | 0.928 | 0.543 | 8.96E-57 | MB4 | EMX2 |
| LIM21 | 7.80E-61 | 0.33544 | 0.399 | 0.097 | 1.52E-56 | MB4 | LIM2 |
| CYR611 | 4.04E-60 | 0.660683 | 0.835 | 0.531 | 7.86E-56 | MB4 | CYR61 |
| DCX1 | 1.08E-58 | 0.278935 | 0.909 | 0.49 | 2.09E-54 | MB4 | DCX |
| PIFO2 | 1.77E-58 | 0.304616 | 0.802 | 0.392 | 3.44E-54 | MB4 | PIFO |
| FAM89A1 | 2.53E-58 | 0.834887 | 0.539 | 0.203 | 4.93E-54 | MB4 | FAM89A |
| ZFYVE161 | 1.32E-57 | 0.398967 | 0.702 | 0.331 | 2.57E-53 | MB4 | ZFYVE16 |
| LINC0099ᵋ | 1.54E-57 | 0.443488 | 0.946 | 0.695 | 2.99E-53 | MB4 | LINC00998 |
| KCNJ131 | 3.26E-57 | 0.403258 | 0.397 | 0.102 | 6.33E-53 | MB4 | KCNJ13 |

| | | | | | | | |
|---|---|---|---|---|---|---|---|
| CFAP1262 | 4.67E-57 | 0.267279 | 0.667 | 0.263 | 9.08E-53 | MB4 | CFAP126 |
| OTX21 | 1.16E-55 | 0.421633 | 0.736 | 0.376 | 2.26E-51 | MB4 | OTX2 |
| SMCO41 | 1.36E-55 | 0.413183 | 0.657 | 0.309 | 2.66E-51 | MB4 | SMCO4 |
| LOXL11 | 2.18E-55 | 0.462317 | 0.727 | 0.381 | 4.25E-51 | MB4 | LOXL1 |
| PRPS12 | 6.95E-55 | 0.477182 | 0.874 | 0.595 | 1.35E-50 | MB4 | PRPS1 |
| CLU3 | 1.04E-54 | 0.37835 | 0.994 | 0.91 | 2.03E-50 | MB4 | CLU |
| COL4A61 | 1.24E-54 | 0.413254 | 0.777 | 0.397 | 2.40E-50 | MB4 | COL4A6 |
| SLC13A42 | 1.62E-54 | 0.26856 | 0.436 | 0.121 | 3.14E-50 | MB4 | SLC13A4 |
| IFITM33 | 3.78E-54 | 0.477346 | 0.926 | 0.623 | 7.36E-50 | MB4 | IFITM3 |
| PDLIM71 | 8.21E-54 | 0.439403 | 0.924 | 0.728 | 1.60E-49 | MB4 | PDLIM7 |
| LGI11 | 8.36E-54 | 0.339145 | 0.463 | 0.145 | 1.63E-49 | MB4 | LGI1 |
| ID11 | 1.32E-53 | 0.541584 | 0.919 | 0.643 | 2.57E-49 | MB4 | ID1 |
| HOPX | 9.03E-53 | 0.384985 | 0.271 | 0.048 | 1.76E-48 | MB4 | HOPX |
| HSBP12 | 6.54E-52 | 0.310892 | 1 | 0.977 | 1.27E-47 | MB4 | HSBP1 |
| MLLT13 | 1.19E-51 | 0.409507 | 0.758 | 0.413 | 2.31E-47 | MB4 | MLLT1 |
| RPRM | 1.91E-51 | 0.507227 | 0.607 | 0.271 | 3.72E-47 | MB4 | RPRM |
| LINC0150: | 5.20E-51 | 0.311653 | 0.444 | 0.146 | 1.01E-46 | MB4 | LINC01503 |
| PGM5P4-/ | 7.87E-51 | 0.268495 | 0.287 | 0.057 | 1.53E-46 | MB4 | PGM5P4-AS1 |
| CST31 | 1.62E-50 | 0.372731 | 0.981 | 0.886 | 3.15E-46 | MB4 | CST3 |
| NQO13 | 1.64E-50 | 0.478436 | 0.717 | 0.384 | 3.19E-46 | MB4 | NQO1 |
| ID32 | 3.22E-50 | 0.520151 | 0.961 | 0.74 | 6.25E-46 | MB4 | ID3 |
| SHTN11 | 8.87E-50 | 0.305168 | 0.572 | 0.226 | 1.72E-45 | MB4 | SHTN1 |
| MEST2 | 2.68E-49 | 0.530058 | 0.99 | 0.922 | 5.22E-45 | MB4 | MEST |
| SOST1 | 4.25E-49 | 0.603124 | 0.248 | 0.042 | 8.27E-45 | MB4 | SOST |
| LY6H1 | 7.20E-49 | 0.372827 | 0.682 | 0.332 | 1.40E-44 | MB4 | LY6H |
| VAT1L2 | 1.54E-48 | 0.356717 | 0.558 | 0.225 | 3.00E-44 | MB4 | VAT1L |
| PCSK1N3 | 5.13E-48 | 0.395068 | 0.957 | 0.785 | 9.97E-44 | MB4 | PCSK1N |
| MRPS63 | 3.22E-46 | 0.388578 | 0.944 | 0.78 | 6.27E-42 | MB4 | MRPS6 |
| DMRT3 | 4.94E-46 | 0.306247 | 0.572 | 0.247 | 9.61E-42 | MB4 | DMRT3 |
| SLC22A17 | 6.42E-46 | 0.410163 | 0.899 | 0.668 | 1.25E-41 | MB4 | SLC22A17 |
| CNTNAP2: | 2.67E-45 | 0.459928 | 0.95 | 0.753 | 5.20E-41 | MB4 | CNTNAP2 |
| LRRN21 | 3.89E-45 | 0.267804 | 0.51 | 0.199 | 7.57E-41 | MB4 | LRRN2 |
| CD3021 | 1.44E-44 | 0.303494 | 0.378 | 0.114 | 2.80E-40 | MB4 | CD302 |
| PLD32 | 2.89E-44 | 0.354912 | 0.986 | 0.908 | 5.62E-40 | MB4 | PLD3 |
| PRNP1 | 2.90E-44 | 0.382886 | 0.787 | 0.485 | 5.64E-40 | MB4 | PRNP |
| BDH2 | 3.51E-44 | 0.382973 | 0.831 | 0.544 | 6.82E-40 | MB4 | BDH2 |
| FXYD11 | 9.10E-44 | 0.331721 | 0.504 | 0.205 | 1.77E-39 | MB4 | FXYD1 |
| PSAP | 2.32E-43 | 0.295356 | 0.994 | 0.965 | 4.51E-39 | MB4 | PSAP |
| RHOH1 | 3.55E-43 | 0.260999 | 0.442 | 0.155 | 6.90E-39 | MB4 | RHOH |
| TAGLN22 | 3.01E-42 | 0.342718 | 0.864 | 0.561 | 5.85E-38 | MB4 | TAGLN2 |

| | | | | | | | |
|---|---|---|---|---|---|---|---|
| COTL11 | 3.99E-42 | 0.35015 | 0.946 | 0.802 | 7.77E-38 | MB4 | COTL1 |
| CTGF1 | 4.35E-42 | 0.63205 | 0.665 | 0.369 | 8.46E-38 | MB4 | CTGF |
| TMEM882 | 9.06E-42 | 0.390466 | 0.436 | 0.162 | 1.76E-37 | MB4 | TMEM88 |
| IFI27L21 | 2.85E-41 | 0.358346 | 0.924 | 0.769 | 5.55E-37 | MB4 | IFI27L2 |
| LHFP | 6.04E-41 | 0.316229 | 0.676 | 0.363 | 1.17E-36 | MB4 | LHFP |
| IFI27L13 | 2.22E-40 | 0.394983 | 0.837 | 0.613 | 4.32E-36 | MB4 | IFI27L1 |
| BAMBI1 | 5.92E-40 | 0.317257 | 0.537 | 0.24 | 1.15E-35 | MB4 | BAMBI |
| PLTP3 | 7.96E-40 | 0.320058 | 0.921 | 0.718 | 1.55E-35 | MB4 | PLTP |
| RHBDD23 | 1.73E-39 | 0.3082 | 0.957 | 0.845 | 3.37E-35 | MB4 | RHBDD2 |
| ZNF436-A | 2.52E-39 | 0.302837 | 0.634 | 0.321 | 4.89E-35 | MB4 | ZNF436-AS1 |
| TSPAN41 | 2.98E-39 | 0.379741 | 0.764 | 0.522 | 5.80E-35 | MB4 | TSPAN4 |
| MT-CO34 | 1.81E-38 | 0.37213 | 1 | 0.979 | 3.52E-34 | MB4 | MT-CO3 |
| SLIT22 | 5.34E-38 | 0.31282 | 0.514 | 0.229 | 1.04E-33 | MB4 | SLIT2 |
| ANXA63 | 1.02E-37 | 0.349882 | 0.692 | 0.399 | 1.98E-33 | MB4 | ANXA6 |
| LSAMP | 1.11E-37 | 0.340692 | 0.725 | 0.418 | 2.16E-33 | MB4 | LSAMP |
| IFI6 | 1.44E-37 | 0.30844 | 0.62 | 0.339 | 2.81E-33 | MB4 | IFI6 |
| BCAS41 | 1.65E-37 | 0.296039 | 0.69 | 0.398 | 3.21E-33 | MB4 | BCAS4 |
| CRYL1 | 2.78E-37 | 0.332941 | 0.655 | 0.365 | 5.41E-33 | MB4 | CRYL1 |
| ZFP36L22 | 4.14E-37 | 0.391773 | 0.783 | 0.504 | 8.06E-33 | MB4 | ZFP36L2 |
| CALB1 | 5.45E-37 | 0.620847 | 0.444 | 0.187 | 1.06E-32 | MB4 | CALB1 |
| GJA12 | 8.93E-37 | 0.279689 | 0.622 | 0.31 | 1.74E-32 | MB4 | GJA1 |
| PGRMC11 | 1.94E-36 | 0.548847 | 0.959 | 0.86 | 3.77E-32 | MB4 | PGRMC1 |
| TM7SF22 | 2.12E-36 | 0.340447 | 0.866 | 0.602 | 4.13E-32 | MB4 | TM7SF2 |
| GMPR2 | 4.97E-36 | 0.261292 | 0.436 | 0.169 | 9.67E-32 | MB4 | GMPR |
| DNPH13 | 5.35E-36 | 0.314567 | 0.878 | 0.594 | 1.04E-31 | MB4 | DNPH1 |
| ABHD21 | 6.41E-36 | 0.259955 | 0.432 | 0.172 | 1.25E-31 | MB4 | ABHD2 |
| VGLL41 | 7.77E-36 | 0.33529 | 0.814 | 0.564 | 1.51E-31 | MB4 | VGLL4 |
| APLP22 | 1.44E-35 | 0.333097 | 0.942 | 0.809 | 2.80E-31 | MB4 | APLP2 |
| COL4A52 | 4.19E-35 | 0.293061 | 0.632 | 0.326 | 8.15E-31 | MB4 | COL4A5 |
| STRA6 | 5.59E-35 | 0.256415 | 0.403 | 0.157 | 1.09E-30 | MB4 | STRA6 |
| NREP1 | 8.23E-35 | 0.303848 | 0.963 | 0.798 | 1.60E-30 | MB4 | NREP |
| CCND1 | 1.25E-34 | 0.333905 | 0.835 | 0.609 | 2.43E-30 | MB4 | CCND1 |
| C1QTNF4 | 1.44E-34 | 0.270948 | 0.541 | 0.255 | 2.80E-30 | MB4 | C1QTNF4 |
| CNN3 | 1.78E-34 | 0.368341 | 0.986 | 0.919 | 3.47E-30 | MB4 | CNN3 |
| PDLIM5 | 1.38E-33 | 0.283305 | 0.678 | 0.401 | 2.68E-29 | MB4 | PDLIM5 |
| ATOX1 | 1.59E-32 | 0.287061 | 0.946 | 0.854 | 3.09E-28 | MB4 | ATOX1 |
| SESN31 | 3.86E-32 | 0.285512 | 0.659 | 0.372 | 7.52E-28 | MB4 | SESN3 |
| DNER1 | 4.14E-32 | 0.403892 | 0.872 | 0.665 | 8.05E-28 | MB4 | DNER |
| COL1A2 | 4.95E-32 | 0.369581 | 0.39 | 0.154 | 9.63E-28 | MB4 | COL1A2 |
| MFAP22 | 5.89E-32 | 0.267639 | 0.694 | 0.389 | 1.15E-27 | MB4 | MFAP2 |

| | | | | | | | |
|---|---|---|---|---|---|---|---|
| HRSP121 | 7.86E-32 | 0.30476 | 0.686 | 0.418 | 1.53E-27 | MB4 | HRSP12 |
| TMEM981 | 7.93E-32 | 0.337498 | 0.87 | 0.63 | 1.54E-27 | MB4 | TMEM98 |
| MT-ND33 | 1.25E-31 | 0.365151 | 0.928 | 0.769 | 2.44E-27 | MB4 | MT-ND3 |
| SPINT23 | 2.04E-31 | 0.313264 | 0.934 | 0.708 | 3.97E-27 | MB4 | SPINT2 |
| HK1 | 3.10E-31 | 0.296244 | 0.764 | 0.51 | 6.02E-27 | MB4 | HK1 |
| PCBD11 | 3.93E-31 | 0.302473 | 0.795 | 0.582 | 7.64E-27 | MB4 | PCBD1 |
| MT-CYB4 | 6.22E-31 | 0.383499 | 0.994 | 0.959 | 1.21E-26 | MB4 | MT-CYB |
| MRPL281 | 1.20E-30 | 0.272974 | 0.917 | 0.735 | 2.34E-26 | MB4 | MRPL28 |
| CYSTM12 | 1.39E-30 | 0.289135 | 0.924 | 0.723 | 2.71E-26 | MB4 | CYSTM1 |
| NARS2 | 3.81E-30 | 0.269707 | 0.649 | 0.378 | 7.42E-26 | MB4 | NARS2 |
| CAPZA2 | 4.45E-30 | 0.287014 | 0.952 | 0.886 | 8.65E-26 | MB4 | CAPZA2 |
| ZNF5031 | 3.14E-29 | 0.30204 | 0.855 | 0.569 | 6.10E-25 | MB4 | ZNF503 |
| NUCB1 | 6.91E-29 | 0.269591 | 0.866 | 0.69 | 1.34E-24 | MB4 | NUCB1 |
| TSPAN31 | 1.34E-28 | 0.27003 | 0.961 | 0.902 | 2.61E-24 | MB4 | TSPAN3 |
| EMX2OS | 4.10E-28 | 0.257328 | 0.599 | 0.335 | 7.97E-24 | MB4 | EMX2OS |
| AC007325 | 5.31E-28 | 0.252607 | 0.585 | 0.317 | 1.03E-23 | MB4 | AC007325.4 |
| ENO31 | 1.14E-27 | 0.272724 | 0.643 | 0.361 | 2.21E-23 | MB4 | ENO3 |
| C1orf122 | 1.46E-27 | 0.254653 | 0.946 | 0.826 | 2.85E-23 | MB4 | C1orf122 |
| POLR2L3 | 7.33E-27 | 0.277045 | 0.967 | 0.908 | 1.43E-22 | MB4 | POLR2L |
| KRT81 | 9.17E-27 | 0.316994 | 0.686 | 0.443 | 1.78E-22 | MB4 | KRT8 |
| MT-ND23 | 1.81E-26 | 0.324931 | 0.994 | 0.967 | 3.52E-22 | MB4 | MT-ND2 |
| SLC5A61 | 2.59E-26 | 0.320518 | 0.488 | 0.267 | 5.04E-22 | MB4 | SLC5A6 |
| RBP13 | 5.15E-25 | 0.298245 | 0.851 | 0.619 | 1.00E-20 | MB4 | RBP1 |
| PEG10 | 4.41E-23 | 0.413817 | 0.87 | 0.704 | 8.57E-19 | MB4 | PEG10 |
| XRCC5 | 1.30E-22 | 0.310004 | 0.969 | 0.883 | 2.53E-18 | MB4 | XRCC5 |
| MT-ATP63 | 4.48E-22 | 0.331782 | 0.965 | 0.896 | 8.72E-18 | MB4 | MT-ATP6 |
| MT-ND6 | 6.39E-22 | 0.291512 | 0.61 | 0.393 | 1.24E-17 | MB4 | MT-ND6 |
| TXNDC171 | 7.49E-22 | 0.252505 | 0.888 | 0.74 | 1.46E-17 | MB4 | TXNDC17 |
| TMCO31 | 1.80E-21 | 0.264486 | 0.624 | 0.382 | 3.50E-17 | MB4 | TMCO3 |
| MT-CO12 | 4.50E-21 | 0.262912 | 0.994 | 0.993 | 8.76E-17 | MB4 | MT-CO1 |
| HIST1H2A | 2.37E-19 | 0.334346 | 0.521 | 0.313 | 4.61E-15 | MB4 | HIST1H2AC |
| KIF21A1 | 1.36E-18 | 0.251336 | 0.719 | 0.533 | 2.64E-14 | MB4 | KIF21A |
| MT-CO23 | 5.51E-18 | 0.273314 | 1 | 0.985 | 1.07E-13 | MB4 | MT-CO2 |
| IGFBP2 | 1.45E-17 | 0.273699 | 0.884 | 0.718 | 2.82E-13 | MB4 | IGFBP2 |
| LDHA2 | 1.98E-08 | 0.357893 | 0.698 | 0.577 | 0.000385 | MB4 | LDHA |
